# Integrated View of Baseline Protein Expression in Human Tissues using public Data Independent Acquisition datasets

**DOI:** 10.1101/2024.09.16.613191

**Authors:** Ananth Prakash, Andrew Collins, Liora Vilmovsky, Silvie Fexova, Andrew R. Jones, Juan Antonio Vizcaino

## Abstract

The PRIDE database is the largest public data repository of mass spectrometry-based proteomics data and currently stores more than 40,000 datasets covering a wide range of organisms, experimental techniques and biological conditions. During the past few years, PRIDE has seen a significant increase in the amount of submitted Data-Independent Acquisition (DIA) proteomics datasets. This provides an excellent opportunity for large scale data reanalysis and reuse.

We have reanalysed 15 public label-free DIA datasets across various healthy human tissues, to provide a state-of-the-art view of the human proteome in baseline conditions (without any perturbations). We computed baseline protein abundances and compared them across various tissues, samples and datasets. Our second aim was to compare protein abundances obtained here from the results of previous analyses using human baseline Data-Dependent Acquisition (DDA) datasets. We observed a good correlation across some tissues, especially in liver and colon but weak correlations were found in others, such as lung and pancreas. The reanalysed results including protein abundance values and curated metadata are made available to view and download from the resource Expression Atlas.

**For TOC Only:** 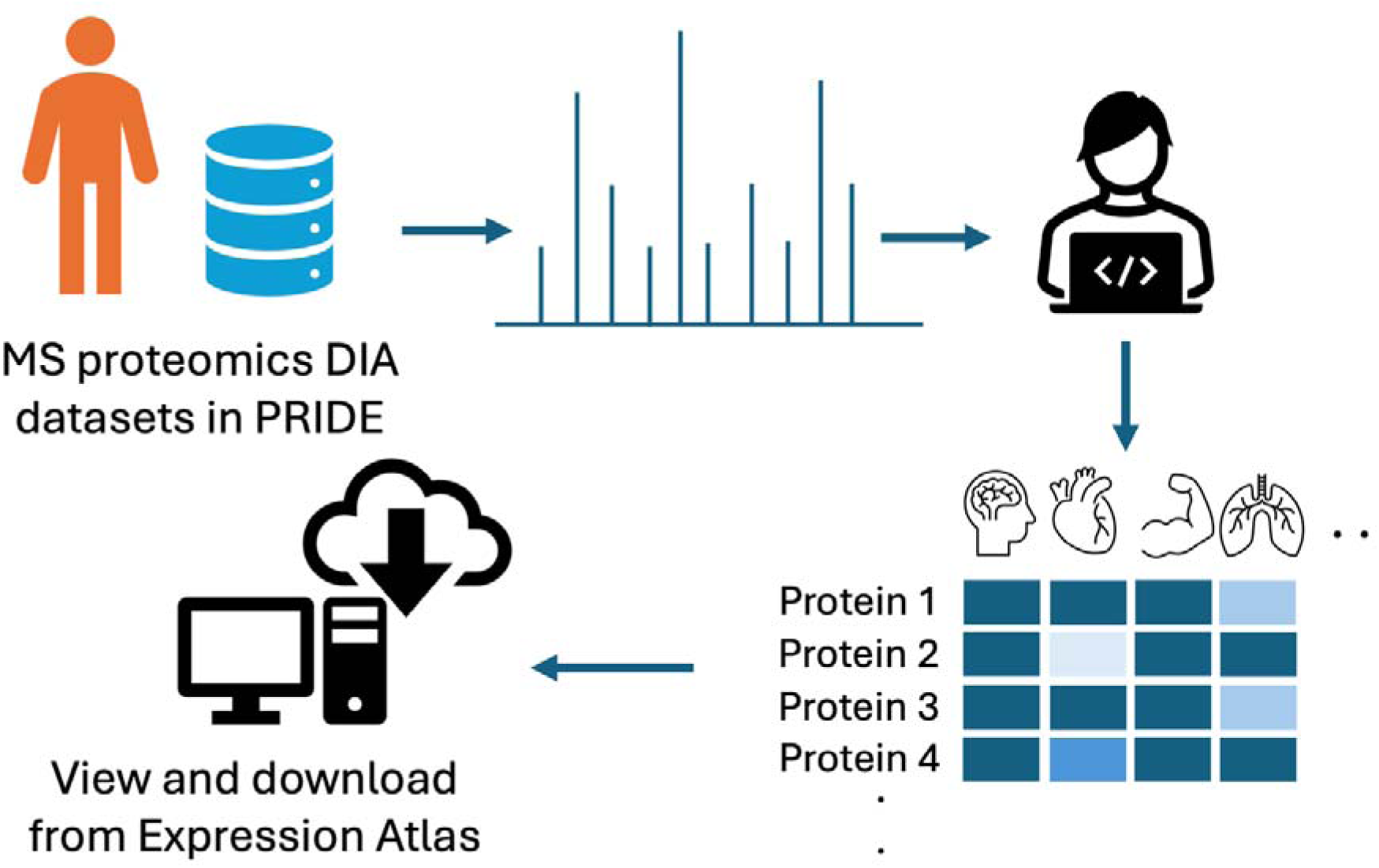

## Introduction

The most popular quantitative approach for Mass Spectrometry (MS)-based proteomics has historically been data-dependent acquisition (DDA) bottom-up proteomics. However, Data Independent Acquisition (DIA) approaches^1^ have matured enormously, thanks to multiple technical developments in e.g. sample preparation, improvements in instrumentation and computational data analysis approaches^2^. In contrast to DDA, where only the most intense peptide ions are measured, in DIA, fragmentation products are generated from every peptide ion that is sampled in the MS^1^ scans. As a result, a DIA analysis approach can potentially provide a more comprehensive quantitative view on the proteome when compared to DDA approaches.

There are two main approaches of DIA proteomics data analysis: spectrum-centric and peptide-centric. Spectrum-centric methods follow the DDA general approach, by generating “pseudo-MS/MS spectra” where fragment ions are associated with the precursor ion from which they are most likely derived. The pseudo spectra can be analysed like DDA data, for instance through typical sequence database searching tools. Peptide-centric methods are currently much more widely used and rely on deciding in advance which peptidoforms may be present in the sample, by using a spectral library^2^. There are three main approaches for generating the spectral libraries that can be used for DIA analysis. First, it is possible to perform a deeply fractionated DDA analysis using the same instrumentation as will be used by the DIA experiments, to generate an *experimentally matched* spectral library. Second, there are some publicly available libraries, created from a given type of samples, for instance a “pan-species” spectral library, assembled from multiple MS runs^3^. The third approach consists of the use of *in silico* generated spectral libraries, generated using artificial intelligence (AI)-trained models, having learnt retention times and peptide intensities from past DDA datasets^2^.

In parallel to other developments, for the last decade or so, the proteomics community has learnt to follow and implement open data practices. The PRIDE database^4^, one of the founding members of the ProteomeXchange consortium^5^, is the most popular proteomics data repository worldwide. The availability of extensive public proteomics datasets in PRIDE and other ProteomeXchange resources has triggered multiple applications, such as meta-analysis studies including some studies applying different AI approaches^6,7^. Additionally, by systematically reanalysing public datasets, original findings can be updated, confirmed and/or strengthened^8^. Moreover, novel insights beyond the scope of the original studies can be obtained through alternative reanalysis strategies, for instance in the case of proteogenomics approaches^9^.

In this context, we are facilitating comparison of transcriptomics and proteomics data in the resource Expression Atlas^10^, so that protein expression abundance information is made accessible in the long-term to life scientists, including those not experts in proteomics. We have already performed combined analyses of baseline (i.e. without any perturbation) protein expression studies of public DDA experiments coming from human^11^, and from mouse and rat^12^ and pig tissues^13^, as model organisms. Additionally, we have also performed combined analyses of DDA experiments generated from cell lines and tumour tissue^14^, and more recently, a study focused on colorectal cancer related datasets as an approach for biomarker discovery^15^. In all cases, protein abundance results have been made available through Expression Atlas. There, protein abundance data can be visualised together with gene expression information, although rarely from the same samples.

However, in parallel with the trends in the field, an increasing fraction of data deposited in PRIDE and in other ProteomeXchange resources comes from DIA approaches. In this context, a few years back, we carried out a pilot study including ten DIA human datasets including mostly cell lines (all of them, SWATH-MS experiments from SCIEX instruments)^16^, using an analysis approach based on the use of the “pan-human” spectral library^17^.

Here, we report the reanalysis and integration of 15 public label-free DIA human baseline tissue datasets including 178 healthy/normal control samples. The objective of this study is two-fold: on one hand, we aim to facilitate access to protein abundance data from baseline human tissues coming from state-of-the-art DIA approaches. On the other, we want to study their similarity with analogous protein abundance data generated using DDA approaches. Unlike our previous study involving DIA datasets^16^, we used *in silico* generated spectral libraries for the analysis. The protein abundance results, as in our previous studies, have been incorporated into Expression Atlas. Additionally, we made a comparison of the expression values coming from DIA datasets and the protein abundance results from previous DDA studies including human baseline tissues^18,19^, for instance using ProteomicsDB data and also our previous combined analysis of DDA public human datasets^11^.

## Methods

### Datasets

Public MS-based human DIA proteomics datasets were queried from the PRIDE database in January 2023. Out of the 288 datasets that were available, several datasets were selected for downstream reanalysis. The selection criteria included: (i) non-enriched samples, only coming from tissues (not including cell lines or fluid samples such as blood/plasma, cerebrospinal fluid, etc.); (ii) datasets from SCIEX (TripleTOF series) and Thermo Fisher Scientific (QExactive series and Orbitrap) instruments; (iii) availability of experimental metadata to link samples to their respective data files; and (iv) in the case of datasets from SCIEX instruments, only datasets that had both .wiff and .scan files deposited were considered (this is a requirement that was added for all SCIEX data submissions to PRIDE in 2021, but there are some previously submitted datasets that do not comply with this requirement). Where enough experimental metadata was not made available in the initial data submissions and/or in the corresponding publications we contacted their respective authors to obtain this information. At the end, we selected 15 datasets for reanalysis (Table 1).

**Table 1.** Summary of protein identifications among datasets. * Healthy/normal control samples only. **†** Post-processed results from control samples, wherein each peptide has at least 2 peptides mapping to it. **‡** Post-processed results from control samples, where a gene has at least two unique peptide mappings. Protein expression data in Expression Atlas can be accessed using the link https://wwwdev.ebi.ac.uk/gxa/experiments/E-PROT-XXX/Results, where XXX should be replaced with an E-PROT identifier.

| Expression Atlas accession number | PRIDE proteomics dataset identifier | Tissue | Mass spectrometer | Fractionation | No. of MS runs* | Number of samples* | Number of PSMs* | Number of unique peptides † | Number of unique genes (canonical proteins) ‡ |
| --- | --- | --- | --- | --- | --- | --- | --- | --- | --- |
| E-PROT-137 | PXD031419 <sup>29</sup> | Brain | TripleTOF 5600 | No | 75 | 19 | 1,424,053 | 49,709 | 5,069 |
| E-PROT-138 | PXD012254 <sup>30</sup> | Colon | Q Exactive | No | 19 | 19 | 871,632 | 53,273 | 5,567 |
| E-PROT-139 | PXD001506 <sup>31</sup> | Duodenum | TripleTOF 5600 | No | 6 | 6 | 104,435 | 14,424 | 1,955 |
| E-PROT-140 | PXD001764 <sup>32</sup> | Esophagus | TripleTOF 5600 | No | 3 | 3 | 58,131 | 15,545 | 2,546 |
| E-PROT-141 | PXD019594 <sup>33</sup> | Heart | TripleTOF 5600 | No | 10 | 10 | 93,177 | 10,456 | 1,131 |
| E-PROT-142 | PXD025705 <sup>34</sup> | Liver | Q Exactive Plus | No | 33 | 11 | 3,099,160 | 85,833 | 6,403 |
| E-PROT-143 | PXD004684 <sup>35</sup> | Lung | Q Exactive | No | 8 | 4 | 155,588 | 25,179 | 2,905 |
| E-PROT-144 | PXD032076 <sup>36</sup> | Pancreas | Orbitrap Fusion | No | 25 | 25 | 417,648 | 27,139 | 3,507 |
| E-PROT-145 | PXD025431 <sup>37</sup> | Skin | Q Exactive HF | No | 10 | 10 | 437,823 | 48,343 | 4,134 |
| E-PROT-146 | PXD002732 <sup>38</sup> | Thyroid | TripleTOF 5600 | No | 8 | 8 | 160,326 | 21,965 | 2,999 |
| E-PROT-147 | PXD022872 <sup>29</sup> | Brain | TripleTOF 5600 | No | 71 | 11 | 1,432,467 | 50,605 | 5,119 |
| E-PROT-148 | PXD033060 <sup>39</sup> | Brain | TripleTOF 5600 | Yes | 17 | 17 | 621,054 | 44,306 | 4,987 |
| E-PROT-149 | PXD018830 <sup>40</sup> | Breast epithelium | Q Exactive | No | 4 | 4 | 113,729 | 27,855 | 4,159 |
| E-PROT-150 | PXD039665 <sup>41</sup> | Skin | Orbitrap Exploris 480 | No | 13 | 13 | 124,942 | 13,463 | 1,759 |
| E-PROT-151 | PXD034908 <sup>42</sup> | Skeletal muscle | TripleTOF 6600 | No | 54 | 18 | 1,174,230 | 18,406 | 1,413 |
| **Total** | **15 datasets** | **12 tissues** |  |  | **356 MS runs** | **178 Samples** |  |  |  |

Metadata was annotated using Annotare^20^ and saved as Investigation Description Format (IDF) and Sample and Data Relationship Format (SDRF) files. IDF comprises description about dataset experiment protocol, contact information of investigators, and publication details, while SDRF contains information of samples including case/control, donor age, gender, treatment conditions, experimental factors, etc. Both IDF and SDRF files were integrated into Expression Atlas.

### Proteomics raw data processing

To process spectral data on a Linux (Rocky Linux 8) platform, the vendor raw files were first converted to mzML format using conversion tools. For Thermo Fisher Scientific instruments (.raw), Thermorawfileparser^21^ 1.4.2 was used with default settings. For SCIEX instruments (.wiff and.scan) ProteoWizard’s msConvert^22^ conversion tool was used with the “peakPicking vendor” option enabled.

As a target database, the UniProt human ‘one protein sequence per gene’ database (UP000005640, May 2023) with 20,838 sequences was used to which cRAP contaminant sequences^23^ (245 sequences) were added. The target database was used in generating an *in silico* spectral library using DIA-NN^24^ (version 1.8.1) with default parameters. We also used an entrapment database^25,26^ wherein *Arabidopsis thaliana* UniProt ‘one protein sequence per gene’ database (UP000006548, December 2022) with 41,621 sequences, which was added to the target human protein search database. Similarly, an *in silico* spectral library of the entrapment database was generated with default parameters using DIA-NN.

To improve run time performance, we designed our analysis by separating a conventional DIA-NN^24^ run into two phases so that it could be reanalysed efficiently on a Linux high performance computing (HPC) platform, using the Slurm job scheduler^27^. Prior to performing the identification for any given dataset, a calibration run was first performed to identify the search tolerances. By default, DIA-NN will auto-detect the tolerances using the first sample in the batch. This leads to issues when running DIA-NN in parallel as each sample will be run with different tolerance settings. To solve this, we ran DIA-NN using default auto-detect settings on a single sample against the target database. The output of this calibration run provided the MS1, MS2, and scan window values that would be used for the full run.

Using the tolerance values identified above, we then employed a multi-node approach to process all raw files in parallel across the HPC cluster. This parallel process allowed us to reduce the run time of DIA-NN’s first phase to the runtime of a single file - a speed-up factor directly proportional to the number of files in each dataset. In this first phase, each MS run was searched against UniProt human with an *Arabidopsis thaliana* entrapment database and the *in silico* spectral library mentioned above. During this phase, cross-run normalisation and MaxLFQ-based protein quantification were disabled (--no-norm, --no-maxlfq), match between runs was not performed, and main report not generated (--no-main-report). Q-value was set to 0.01, peptide cleavage sites assigned to trypsin and the rest of the parameters were set to default. At the completion of the first phase, protein quantification (.quant files) from individual runs were saved in a temporary directory. During the second phase, the existing. quant files generated from the first phase (--use-quant) were collated and a cross-run analysis performed on a single node, wherein cross-run normalisation, match-between runs, MaxLFQ-based protein quantification were performed and main report files, including protein groups, gene groups, unique genes and precursors matrices, were generated.

In a conventional DIA-NN run, each input file is sequentially analysed against the spectral library for protein identification and quantification. By leveraging multi-node processing in the first phase wherein each file was simultaneously analysed and then collating the.quant files, this significantly reduced the total run time by approximately 50% (Figure S1 in Supporting File 1).

### Postprocessing

The report.tsv output file from DIA-NN was used as the basis for post-processing. Contaminants were removed along with mappings to more than one protein or gene identifier. In each MS run we removed entries with fewer than two unique peptide sequences per protein and the abundances of proteins were aggregated using their median values within each MS run in a dataset. DIA-NN outputs abundances as label-free quantification (LFQ) values. We converted LFQ to intensity based absolute quantification (iBAQ) values by normalising the LFQ abundances with the theoretical number of tryptic peptides of each canonical protein. As explained in previous publications^11–13^, we represent protein abundances as abundances of their respective parent gene names, which we will use as equivalent to ‘canonical proteins’ as described in UniProt (https://www.uniprot.org/help/canonical_and_isoforms). For ease of comparison of protein abundances across tissues and datasets with potentially large batch effects between different experiments, iBAQ protein abundances were converted into ranked bins^11^. Briefly, iBAQ abundances were numerically sorted and grouped into five bins of equal size. Proteins in bin 1 were of lowest abundance and those in bin 5 were highly abundant. The heatmap of samples was generated using binned abundance values. Post-processing was performed using R scripts.

### Integration into Expression Atlas

Post-processed results comprising protein abundances, expressed as their canonical gene identifiers (proteins were mapped to gene identifiers by DIA-NN), were integrated into Expression Atlas^28^ along with metadata files (IDF and SDRF) and a quality assessment summary. Expression Atlas dataset identifiers (E-PROT) for each dataset are shown in Table 1.

### Comparison of baseline protein abundances generated using DDA data

Ensembl gene identifiers of the respective canonical proteins were used for comparing proteins identified between this study and previous DDA studies performed in human baseline tissues. Ensembl gene identifiers and normalised iBAQ protein abundance values in supplementary table 2 available from^11^ was gathered for tissues that are common between the two studies for comparison purposes. The iBAQ protein abundances of DIA datasets were then compared with the fraction of total normalised iBAQ protein abundances from DDA datasets.

Additionally, normalised protein intensities from ProteomicsDB^18^ were queried for tissues that were in common in our study (8 tissues). Values were obtained using the ProteomicsDB Application Programming Interface (downloaded in April 2022). For different tissue samples we aggregated the normalised intensities using the median of their respective tissues. The intensities were log2 normalised and compared.

Furthermore, protein abundances calculated across various baseline human tissues using the TMT-labelling method were obtained from^19^ (Supplementary file ‘NIHMS1624446- supplement-2’, sheet: ‘C protein normalized abundance’). Protein abundances of the respective tissues measured across different TMT channels and MS runs were aggregated using the median and log2 transformed. Different tissue samples from esophagus, heart, brain and colon were aggregated into their respective tissues. Binned protein abundances of the samples were used for calculating Pearson correlation values within each tissue. For pairwise comparison of samples between different datasets only the abundances (binned values) of proteins commonly identified between two samples were considered. Brain and skin were the only tissues in this study which are available in multiple datasets. The correlation of protein abundances between brain and skin samples from multiple datasets was calculated and median correlation values are presented. Correlations were calculated using the R programming language.

### Comparison of missing values between DDA and DIA datasets

To compare the proportion of missing protein abundance values between DDA and DIA techniques we used the protein abundances from human DDA samples calculated in our previous study^11^. Protein abundances from DDA samples were represented as their canonical gene identifiers as described for DIA samples above. To compare the completeness of protein detection among samples of their respective tissues within a dataset, first for a given tissue we calculated the total number of missing abundances (NA) among its samples across all canonical proteins and then normalised it by the total number of observations, as shown in the following equation below:

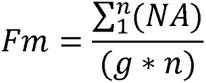

where ‘Fm’ is the fraction of missingness, ‘n’ is total number of samples of a particular tissue within a dataset, ‘g’ is the total number of genes identified in those respective tissue samples within the dataset, ‘NA’ is the missing abundance of a canonical protein. Where a dataset had more than one tissue, the ‘Fm’ was calculated individually for each group of tissue samples and then a median of ‘Fm’ over all tissues was calculated for the dataset.

### False Discovery Rate analysis using entrapment database

All datasets were analysed individually. To estimate the protein False Discovery Rate (FDR) across datasets we used an *Arabidopsis thaliana* entrapment search database as explained above. From the protein group matrix report output file from DIA-NN, we treated the protein groups that had gene identifiers belonging exclusively to *Arabidopsis thaliana* as decoys and the rest of the protein groups were regarded as targets. The distribution of number of datasets in which all decoys and targets were found in common were computed and the FDR for a target protein identified across a number of datasets was calculated using the equation:

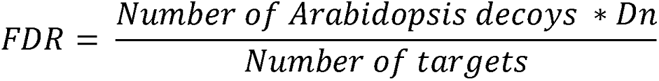

where ‘Dn’ = 0.5, is the database normalising factor i.e., size of UniProt human one protein per gene set database (20,838 sequences) normalised per the size of the UniProt *Arabidopsis thaliana* one protein per gene set database (41,621 sequences).

## Results

### DIA human proteomics datasets reanalysis

We selected 15 DIA proteomics datasets, which represented the available range in the public domain for human tissues in healthy/baseline conditions. The spectra from these datasets were acquired either using SCIEX or Thermo Fisher Scientific instruments, as explained in ‘Methods’. In total there were 1,361 MS runs from all samples in aggregated datasets, which included 356 MS runs from 178 healthy or normal control samples. Each dataset was analysed individually and post-processed. The complete list of datasets along with the number of peptides and proteins identified and quantified are shown in Table 1. Protein abundances computed only from healthy or normal tissue samples are discussed in this manuscript. However, abundances from all samples along with a quality assessment summary and sample metadata are made available in Expression Atlas (https://www.ebi.ac.uk/gxa/) to view and download. The overall data reanalysis protocol is summarised in Figure 1.

**Figure 1.**
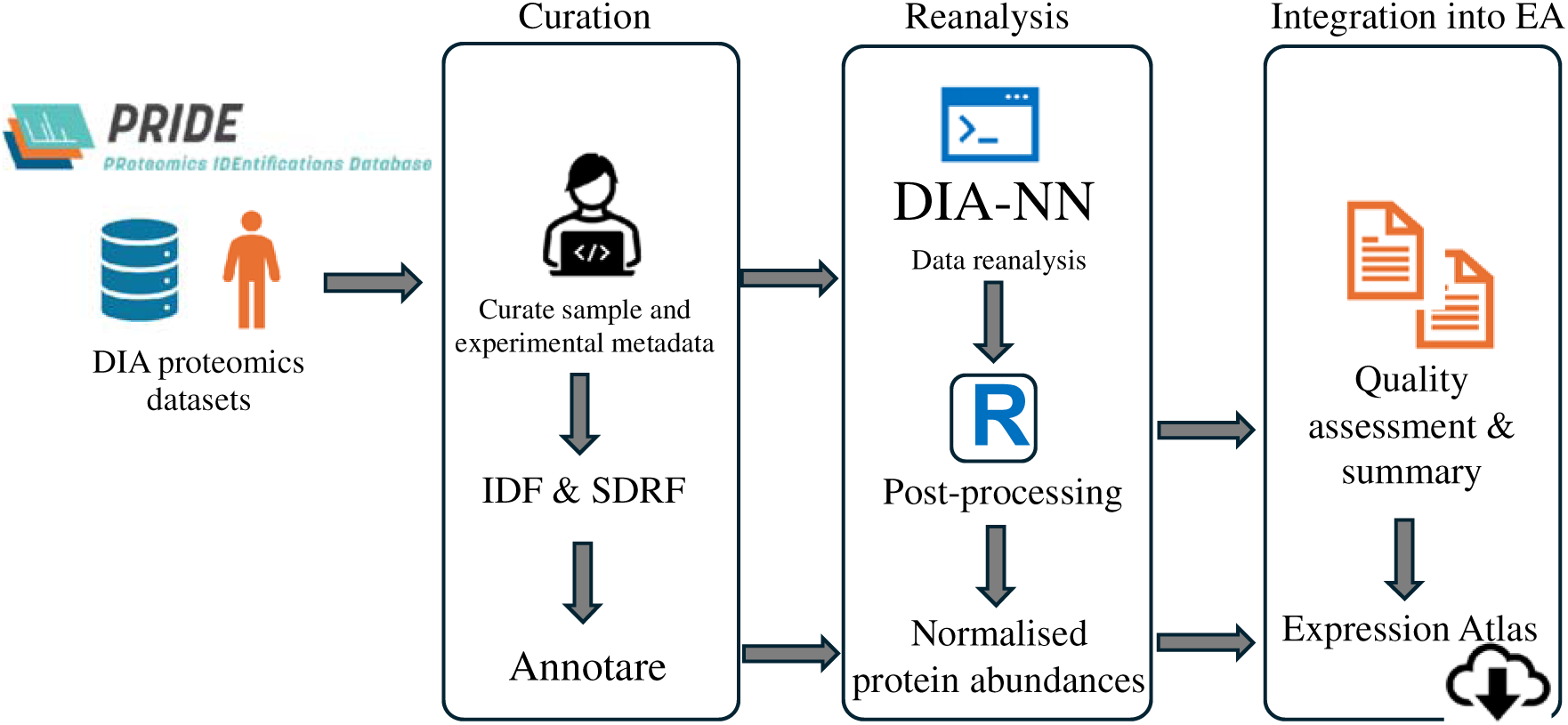
Overview of DIA datasets reanalysis pipeline. EA = Expression Atlas

### Protein coverage across samples

From each sample analysed in a dataset, we kept those proteins which were identified by at least two peptides. Those mapping to more than one gene identifier were also filtered out for downstream analysis. We identified a total of 9,299 canonical proteins, of which 521 proteins were observed in all 12 tissues and 2,679 were only found in one tissue (Table S1 in Supporting File 2). We identified the largest number of proteins in liver (6,401 proteins, 68.8%) followed by brain (6,018, 64.7%, aggregated over three datasets) and colon (5,564, 59.8%). The fewest proteins were identified in heart (1,131, 12.2%), skeletal muscle (1,412, 15.2%) and duodenum (1,955, 21.0%) (Figure 2A). As seen in Table 1, each tissue is represented by one dataset, except brain which is represented by 3 datasets: PXD022872 (5,119, 55.0%), PXD031419 (5,069, 54.5%) and PXD033060 (4,987, 53.6%), and skin, which is represented by 2 datasets: PXD025431 (4,132, 44.4%) and PXD039665 (1,748, 18.8%). We observed significant variations in the protein abundances across each tissue. Figure 2B shows the aggregated (median) protein abundances of each tissue over several samples. Among datasets, PXD025705 (6,401, 68.8%) from liver, had the largest number of proteins identified, followed by PXD012254 (5,564, 59.8%), from colon. The protein abundances in each dataset vary reflecting those observed for every tissue.

**Figure 2.**
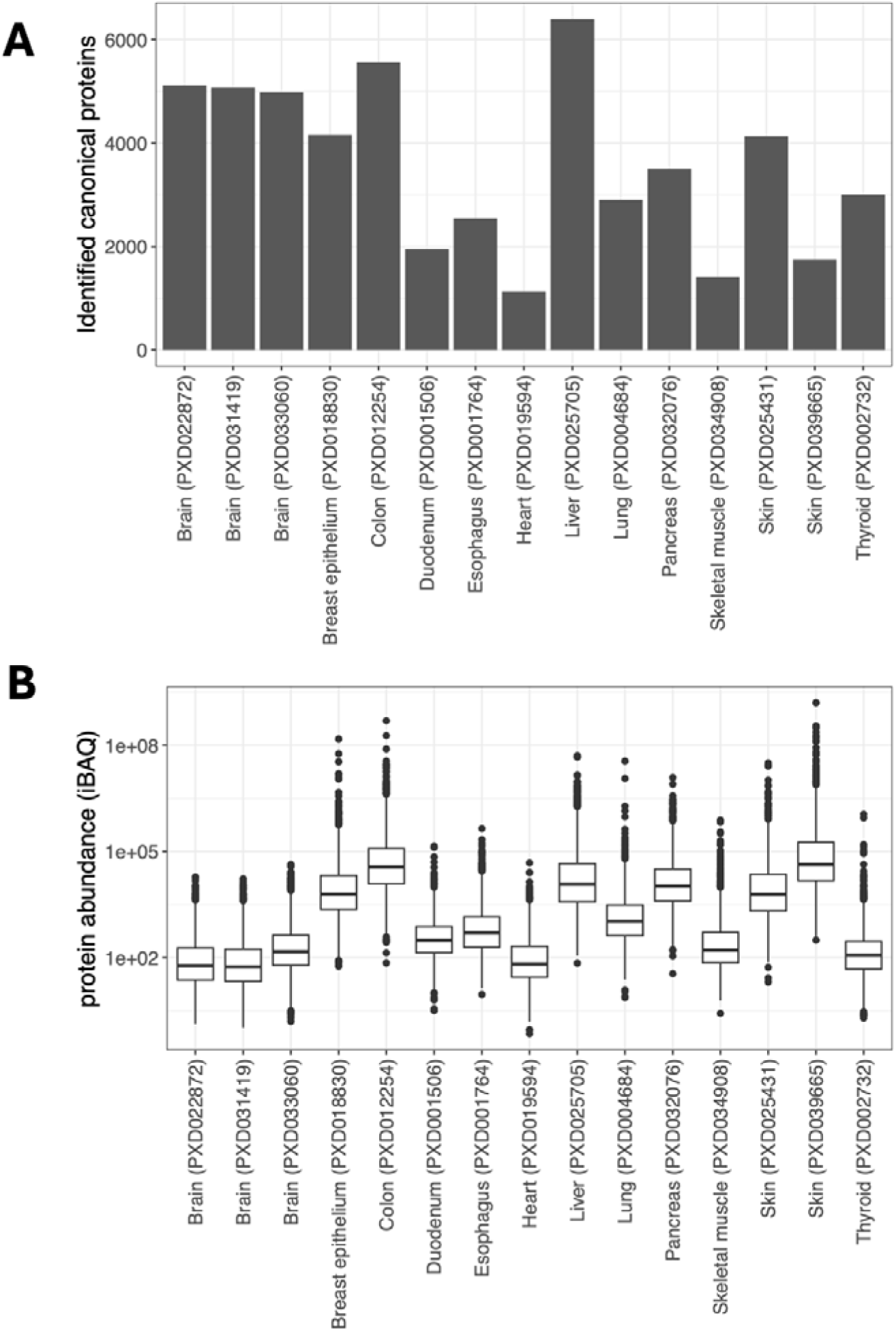
Distribution of protein identification and abundances across tissues and datasets (A) Number of canonical proteins identified across different tissues and datasets. (B) iBAQ protein abundances of canonical proteins across different tissues and datasets.

### Protein abundance comparison across tissues

To compare protein abundances across different tissues we first transformed the LFQ protein abundance values obtained from DIA-NN to iBAQ values as explained in the ‘Methods’ section. To make easier the comparison across datasets and tissues we grouped iBAQ values equally into five categorical bins. Proteins within bin 1 are of lowest abundances and those in bin 5 are of highest abundances. These binned abundances are available in Table S2 in Supporting File 2.

We carried out pairwise comparison across all samples (n=178) using the binned protein abundance values (Figure 3). We observed moderate correlation of protein abundances in brain (median R^2^=0.40) and low correlation of protein abundances in skin samples (median R^2^=0.20). Due to the large number of brain samples included in the aggregated dataset (n=47), there were more data points for computing Pearson correlation values when compared to pairwise sample comparisons made between different tissues. In comparison, the correlation of protein abundances in DDA brain samples from our previous study^11^ was slightly higher (median R^2^=0.61). It is important to note that the number of DDA brain samples was then much larger (n=339) than those in analysed here by DIA approaches (n=47).

**Figure 3.**
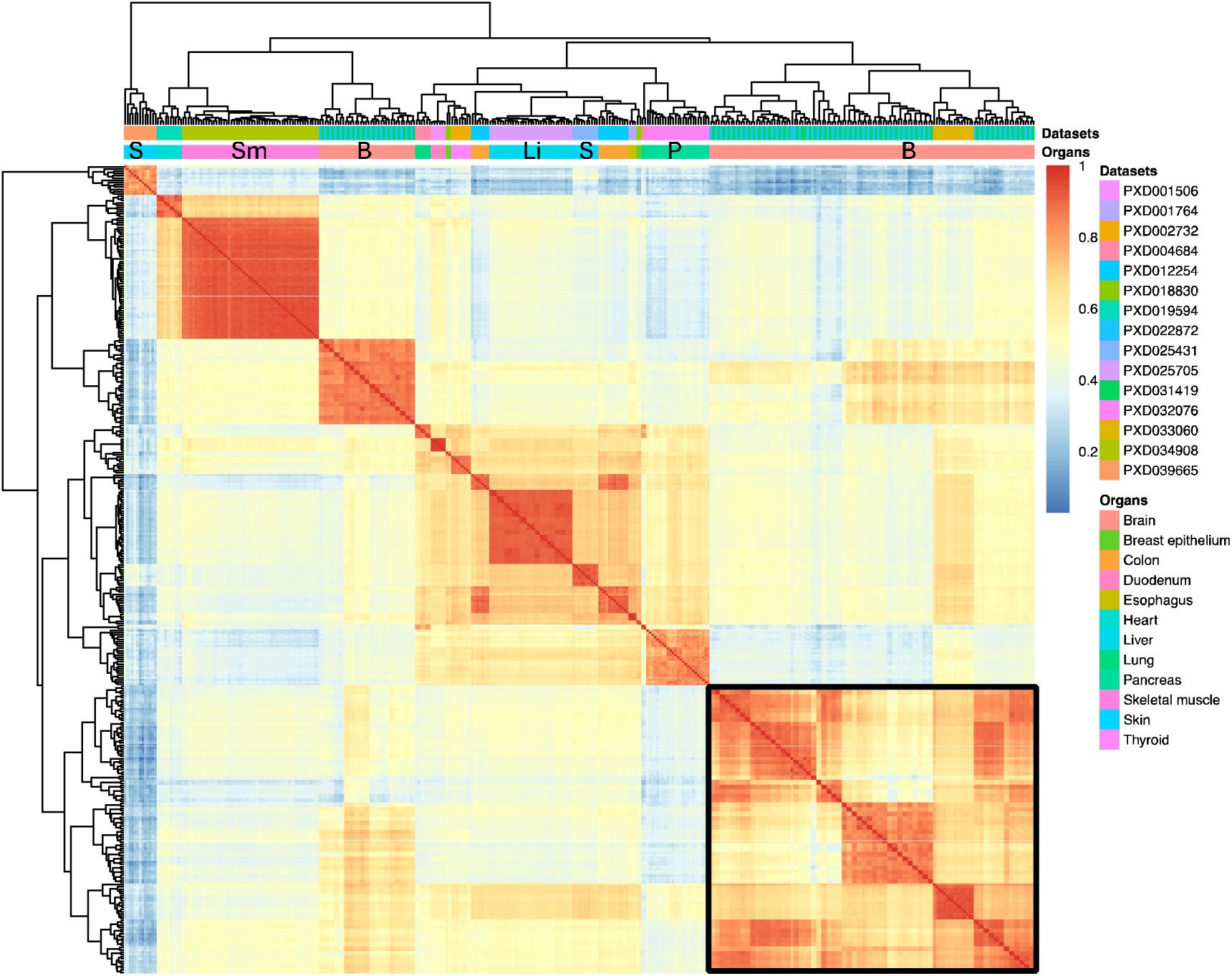
Heatmap of binned protein abundances across all samples between various tissues and datasets. Brain samples clustered together are highlighted using a black border. S: Skin, Sm: Skeletal muscle, B: Brain, Li: Liver, P: Pancreas.

### *Arabidopsis thaliana* entrapment analysis

We designed this work to reanalyse each dataset individually, as in previous studies. The protein FDR was set to 1% with the Benjamini-Hochberg correction for multiple hypothesis testing within each dataset. Although this is a stringent filter, if protein observations from multiple datasets are collated at the end, this could result in a protein FDR much greater than 1%. To check this, we used an entrapment target sequence approach, wherein *Arabidopsis thaliana* protein sequences were added to the target human sequence database. The results from the reanalyses of datasets were searched against the entrapment database for protein groups comprised exclusively of canonical *Arabidopsis thaliana* proteins (false targets). We found a total of 1,367 decoy protein groups across all 15 datasets including 1,018 decoy protein groups which were observed in only one dataset (Figure 4). The largest fraction of decoy hits was found in dataset PXD039665 (4.3%) and the lowest in dataset PXD012254 (1.1%). From the number of decoys detected in common across all datasets, we estimated a resulting combined protein FDR of less than 1% when proteins were observed in at least 7 different datasets and an FDR of less than 5% when proteins were observed in at least 3 datasets. In all cases, *Arabidopsis thaliana* proteins with an evidence of at least 2 peptides were considered. When using the aggregated results from all datasets, column C (‘Present_in_number_of_datasets’), in Tables S1 and S2 (in Supporting File 2) should be used to filter canonical proteins observed in a different number of datasets for obtaining the appropriate values of the ‘combined protein FDR’. We concluded that the protein FDR values were comparable to our previous human DDA study and appropriate for the objectives of this work. We recommend that DIA analyses use an entrapment approach such as this, to ensure that there is robust control of the FDR, following any post-processing or merging steps that may be done.

**Figure 4.**
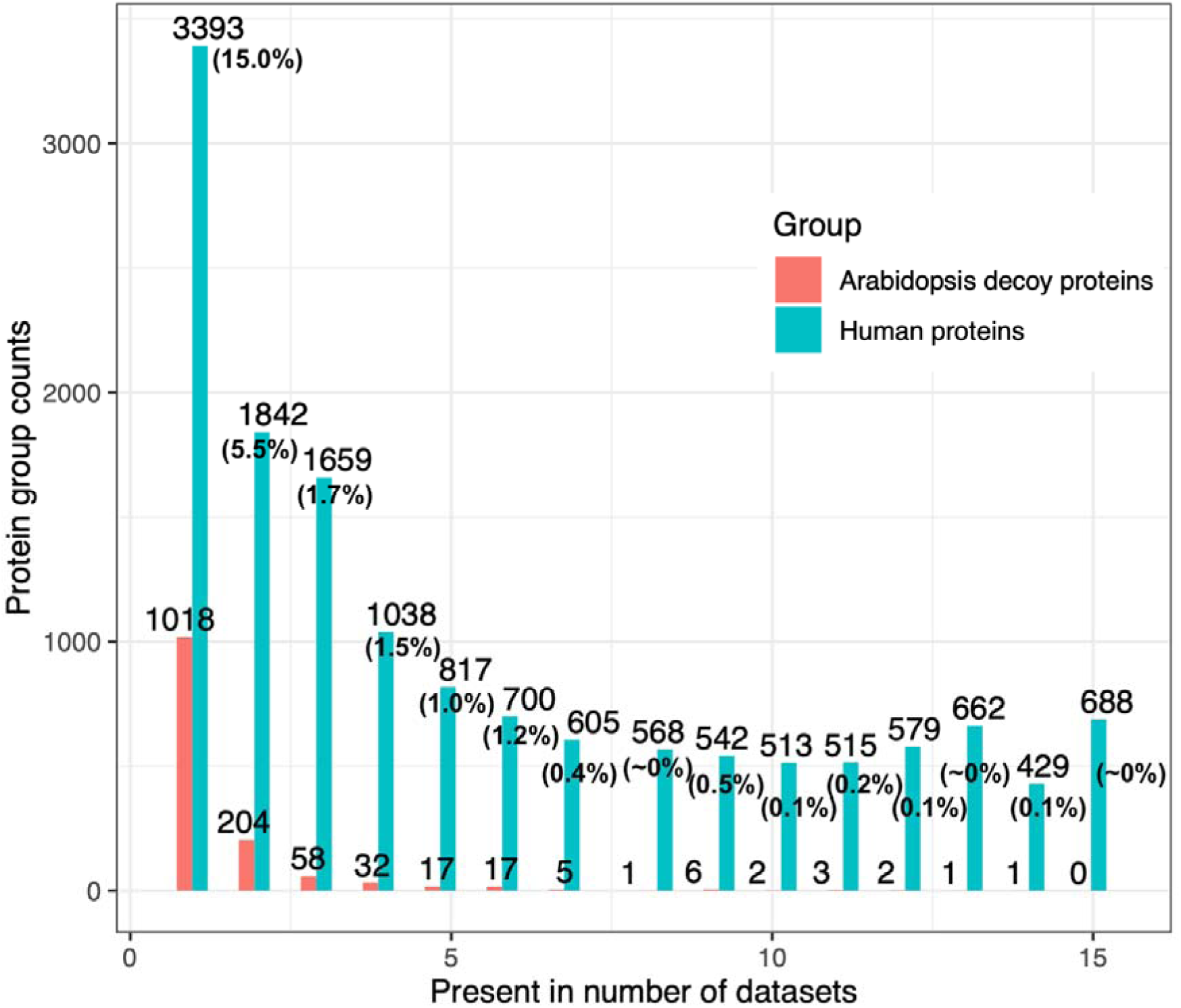
(A) Distribution of decoy and target protein groups present across all datasets identified using the *Arabidopsis thaliana* entrapment search database. The protein FDR values of the target proteins present in common across different numbers of datasets are shown in parenthesis The calculation of the FDR is described in the ‘Methods’ section.

### Protein abundance comparison between the DIA datasets and previous DDA studies

We had previously reanalysed 24 public proteomics dataset representing 31 healthy human tissues^11^ coming from DDA studies. We first compared the number of canonical proteins identified between datasets analysed using DDA and DIA (see ‘Methods’). We had identified a total of 13,071 canonical proteins from DDA studies and 9,299 canonical proteins from the reanalysis of the DIA datasets in this study, all in baseline conditions. When comparing DDA and DIA results, we found 8,449 (64.6%) common canonical proteins obtained from the two groups of datasets, with 4,621 (35.3%) canonical proteins identified only in DDA datasets and 853 (9.1%) canonical proteins identified only in DIA datasets (Table S3 in Supporting File 2, Figure S2 in Supporting File 1). It is important to note here that in comparison to the DIA datasets, there were many more MS runs and samples analysed in the DDA datasets, and additionally many of the DDA datasets were fractionated increasing the depth and coverage of the study. By analysing the missingness of protein abundance values among samples and comparing it with samples in DDA datasets from our earlier study^11^ we observed similar trends in the fraction of missing values among samples analysed by DIA and DDA techniques, implying that DIA does not confer substantial advantages in terms of avoiding missing values, at least in the datasets we analysed (Figure S3 in Supporting File 1).

We then compared protein abundances between DIA datasets and DDA datasets from^11^. Protein abundances from DDA datasets were computed as iBAQ values and, as mentioned in ‘Methods’, the abundances from DIA datasets were transformed from LFQ to iBAQ values. We observed strong correlation of protein abundances in various tissues including colon (R^2^=0.64), liver (R^2^=0.63) and brain (R^2^=0.55), and weak correlation in lung (R^2^=0.24), pancreas (R^2^=0.33), duodenum (R^2^=0.34) and thyroid (R^2^=0.35) (Figure 5). We also compared the binned protein abundances of the commonly identified proteins between tissues in DIA and DDA datasets^11^(Supporting File 3). We observed that highly abundant proteins (of bins 4 and 5) showed a higher similarity in protein abundances between DIA and DDA datasets, when compared to lower abundant proteins (of bins 1 to 3) (Figure S4 in Supporting File 1). The protein abundance profiles in Supporting File 3 can be useful in identifying proteins with similar or dissimilar abundances between different tissues in DIA and DDA datasets. For example, in Supporting File 3 it can be observed that mitochondrial protein ATP synthase group of subunits (gene name: ATP5F1A, UniProt accession: P25705) was detected in both DIA and DDA datasets to be highly expressed across all tissues consistently. However, the gamma-aminobutyric acid receptor subunits (gene name: GABRA1, UniProt accession: P14867) was exclusively identified in brain samples from both DIA and DDA datasets.

**Figure 5.**
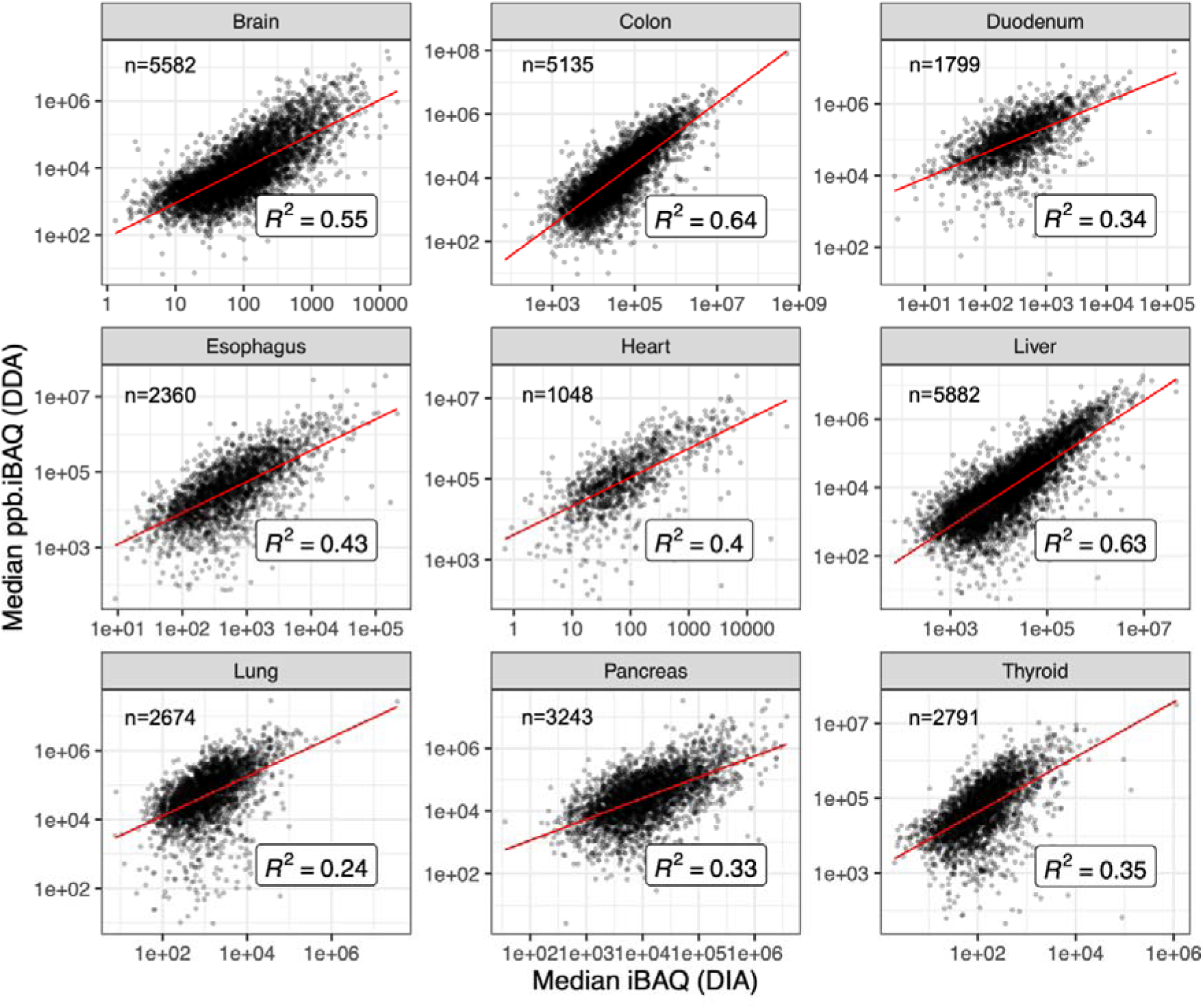
Correlation of protein abundances between human baseline DIA and the DDA datasets from a previous study^11^. *n* shows the number of data points (common canonical proteins) considered in each panel.

We then compared our results with protein abundance data from DDA studies available in ProteomicsDB^18^. We observed a good correlation in protein abundance across various tissues (Figure S5 in Supporting File 1). The strongest correlations were observed in liver (R^2^=0.71) and colon (R^2^=0.68) and the lowest in thyroid (R^2^=0.39). We also compared abundances with protein expression data across various human tissues at baseline conditions from a large-scale study using TMT^19^. In this case, we did not observe a strong correlation across various tissues (Figure S6 in Supporting File 1).

## Discussion

Here we present a detailed reanalysis of 15 DIA proteomics datasets from 12 human tissues in baseline conditions. We selected datasets from PRIDE, manually curated them connecting the samples with the raw files, and generated protein abundances for all of them. The objective was two-fold. First, we wanted to facilitate access to protein abundance data from state-of-the-art DIA proteomics approaches. Second, we wanted to explore their comparability with analogous protein abundance data generated using DDA approaches.

Different benchmarking studies have been published, pointing out advantages and disadvantages of using different approaches (and software tools) for DIA analysis (e.g. ^43–45^). High-quality spectral libraries can be generated *in silico* in tools such as DIA-NN including every peptide sequence. However, using a public or a ‘matched experimental’ library, containing peptidoforms likely to be present in the sample (e.g. for particular biological conditions, tissues or fluids) would likely provide a better statistical power, at the expense of losing a few low abundant peptidoforms absent from a DDA library. An *in silico* spectral library can be potentially much bigger, and thus may give lower sensitivity overall. We here assessed the feasibility of using them to obtain reliable results. It is important to highlight that the use of *in silico* spectral libraries makes public data reanalysis efforts feasible (like in the case of DDA datasets), without having to rely on the availability of e.g. a ‘pan-species’ public library. The other alternative option is that submitters make their in-house spectral libraries publicly available as well, but at present, this does not happen very often. Indeed, although spectral libraries can be submitted to ProteomeXchange resources (and PRIDE in particular) as part of DIA datasets, it is not mandatory to do it and then, it is not a common practice yet. However, there are cases where they are made available, and this is a key factor for enabling the reproducibility the results of the original studies.

The approach followed in this study is different from our previous reanalysis effort of DIA datasets, where the ‘pan-human’ library was used^17^, but also a different analysis software (the pipeline was based on OpenSWATH)^46^. No common datasets were included in both studies. In the current study, by using an entrapment database, we checked that the resulting protein FDR per dataset was at an adequate level.

We also compared the protein abundance values in DIA datasets with the results generated from previous DDA studies. One caveat is that at present, the number of public DIA human datasets from baseline human tissues is much smaller when compared with DDA. The results of this analysis were heterogeneous, with good level of overall correlation for some of the tissues (especially for liver and colon), but much lower for others (e.g. lung and pancreas). Also, we studied the proteins that were detected by either DDA, DIA or both approaches, and compared their level of expression across the common tissues in either DDA or DIA (see Supporting File 3). Finally, we observed a similar percentage of protein abundance missing values among samples analysed by DIA and DDA techniques. It needs to be taken into account that there are some limitations in this comparison. On one hand, samples are not matched between the different studies. On the other hand, for our previous DDA metanalysis study, we used a different protein sequence database, including several proteins per gene.

Lastly, there was only one DIA dataset where fractionation was used. The objective of this study was not to identify tissue specific proteins. Given that most tissues reanalysed here are only represented by one dataset (apart from brain and skin), we think that confidently identifying protein tissue specificity is not feasible (data not shown). However, if the main objective of a study is to find tissue-specific proteins, it would be possible to combine downstream the results found in this study with the findings in our analogous previous DDA study^11^.

Whereas the reuse of public DIA proteomics datasets is relatively common for benchmarking efforts in the context of development of software tools and analysis approaches (e.g. ^47–49^) data reanalysis of DIA datasets is still very limited, unlike in the case of DDA datasets. However, the trend is changing, as more data make it into the public domain. Some recent efforts such as the development of the open quantMS pipeline^50^ can facilitate the reanalysis of large datasets.

In conclusion, we here present a metaanalysis study of public DIA human datasets generated from tissues in baseline conditions, from the PRIDE database. Analogous studies of protein abundance in model organisms (such as those possible for DDA data) are not feasible yet because the amount of DIA baseline data from them is still small. The resulting protein abundance data has been made available via Expression Atlas.

## Code availability

https://github.com/Ananth-Prakash/PRIDE-DIA-DataReuse

## Data availability

Supporting File 1: https://drive.google.com/file/d/1MVYesCbW7gHYv7IgFEVGBpIZUx28mREG/view?usp=sharing

Supporting File 2: https://docs.google.com/spreadsheets/d/1_PtA0Ly9FG27RadlVngi_6cw_clKCYV/edit?usp=sharing&ouid=107719713788096579220&rtpof=true&sd=true

Supporting File 3: https://drive.google.com/file/d/1B2cbA0WsvI-AZRAs5y-Zxd6mAz8SZJXD/view?usp=sharing

## Supplementary data

Supplementary Figure S1: Comparison of the DIA-NN runtimes of an example dataset when run in sequential and parallel modes.

Supplementary Figure S2: Venn diagrams showing the number of canonical proteins identified by DIA in this study and by DDA in our previous study^11^ in various tissue samples.

Supplementary Figure S3: Comparison of missing values in samples between DDA^11^ and DIA datasets.

Supplementary Figure S4: Comparison of binned protein abundances in tissues from DIA and DDA datasets.

Supplementary Figure S5: Comparison of protein expression across various human tissues in baseline conditions analysed from DIA datasets (this study) and from ProteomicsDB.

Supplementary Figure S6: Comparison of protein expression across various human tissues at baseline condition analysed from DIA datasets (this study) and from Jiang *et al.*^19^ using the TMT-labelling method.

Supplementary Table 1: Canonical protein abundances (iBAQ) across various tissues.

Supplementary Table 2: Binned canonical protein abundances across various tissues.

Supplementary Table 3: Canonical proteins identified by DIA only, DDA only and by both techniques.

## Acknowledgements

First, we would like to thank all data submitters who made their datasets available via PRIDE and ProteomeXchange. This work has been funded by the BBSRC/NSF grant ‘DIA-Exchange’ [BB/X001911/1 and BB/X002020/1], BBSRC ‘GRAPPA’ [BB/T019670/1 and

BB/T019557/1], Wellcome [grant number 221401/Z/20/Z] and EMBL core funding.

## Abbreviations

EA: Expression Atlas
DDA: Data Dependent Acquisition
DIA: Data Independent Acquisition
FDR: False Discovery Rate
iBAQ: intensity-Based Absolute Quantification
IDF: Investigation Description Format
LFQ: Label-Free Quantification
ppb: parts per billion
SDRF: Sample and Data Relationship Format

