## Supporting File 1 for "Integrated View of Baseline Protein Expression in Human Tissues using public Data Independent Acquisition datasets"

**TABLE OF CONTENT**

**S-2** Table of Content

**S-3** **Figure S1:** Comparison of DIA-NN runtimes of an example dataset when run in sequential and parallel modes.

**S-4** **Figure S2:** Venn diagrams showing the number of canonical proteins identified by DIA in this study and by DDA in our previous study in various tissue samples.

**S-5** **Figure S3:** Comparison of missing values in samples between DDA and DIA datasets.

**S-6** **Figure S4:** Comparison of binned protein abundances in tissues from DIA and DDA datasets.

**S-7** **Figure S5:** Comparison of protein expression across various human tissues in baseline conditions analysed from DIA datasets (this study) and from ProteomicsDB.

**S-8** **Figure S6:** Comparison of protein expression across various human tissues at baseline condition analysed from DIA datasets (this study) and from Jiang *et al.* using the TMT-labelling method.

**S-9** Supplementary tables

**Supplementary Table 1:** Canonical protein abundances (iBAQ) across various tissues.

**Supplementary Table 2:** Binned canonical protein abundances across various tissues.

**Supplementary Table 3:** Canonical proteins identified by DIA only, DDA only and by both techniques.

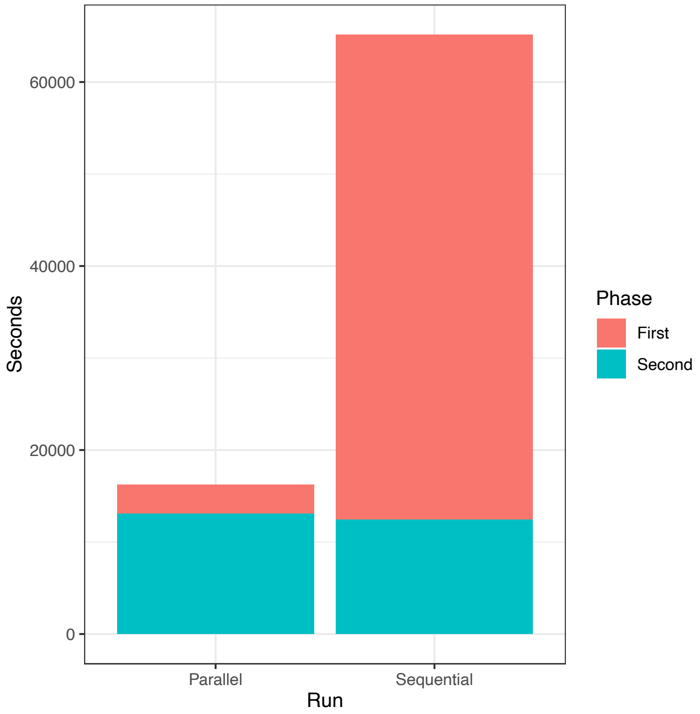


**Figure S1.** Comparison of DIA-NN runtimes of an example dataset PXD032076 (with 173 MS runs) when run in sequential and parallel modes. For sequential run 1x 80 core system was used for the complete run through. For parallel run 8 cores were used for the first phase, then 80 cores for the second phase. The first phase value in parallel = Maximum(all runs)

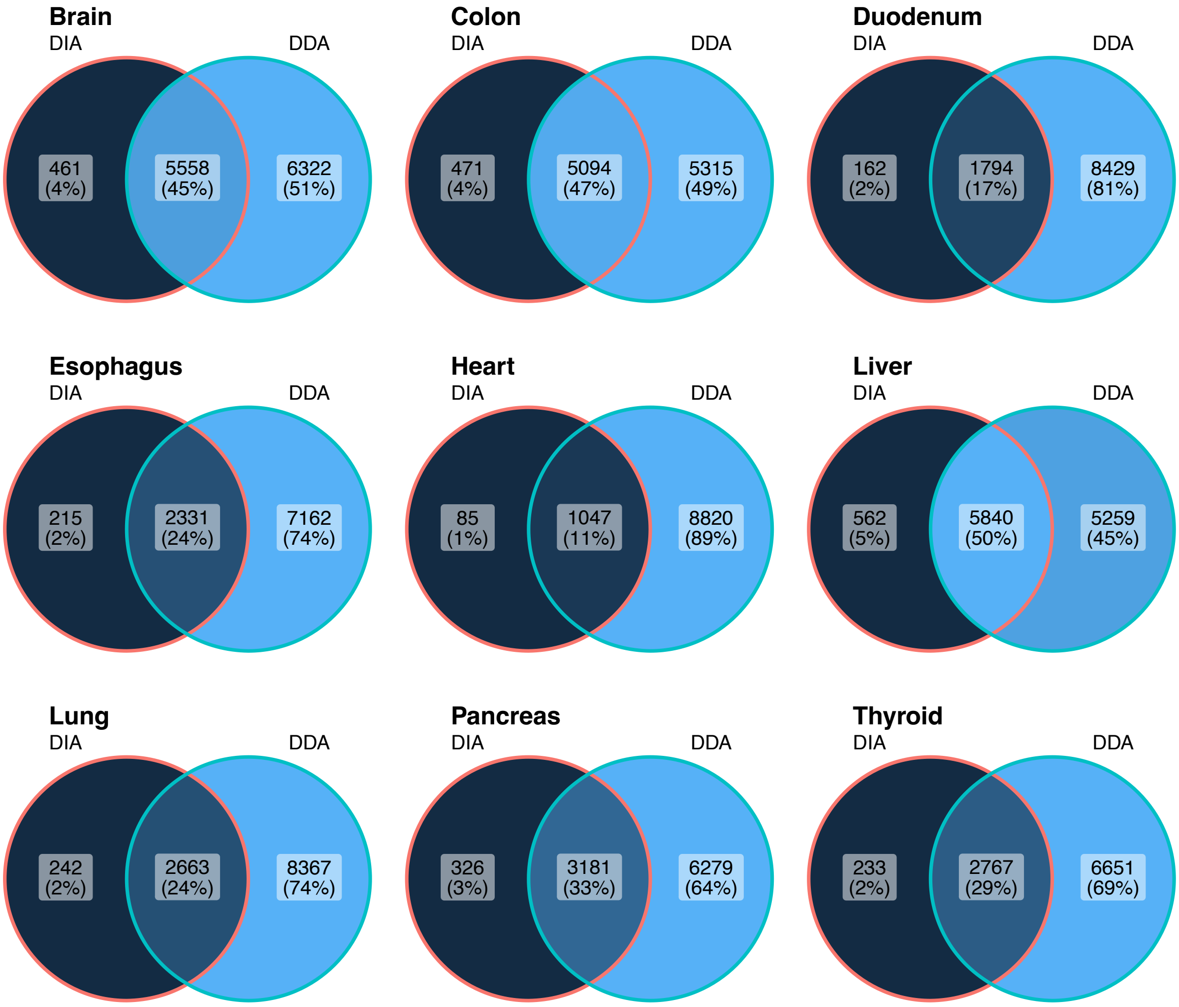


**Figure S2.** Venn diagrams showing the number of canonical proteins identified across various tissues by DIA in this study and by DDA in our previous study[1]. UniProt protein identifiers were mapped to their respective Ensembl gene symbols, representing canonical proteins, which were used for comparison.

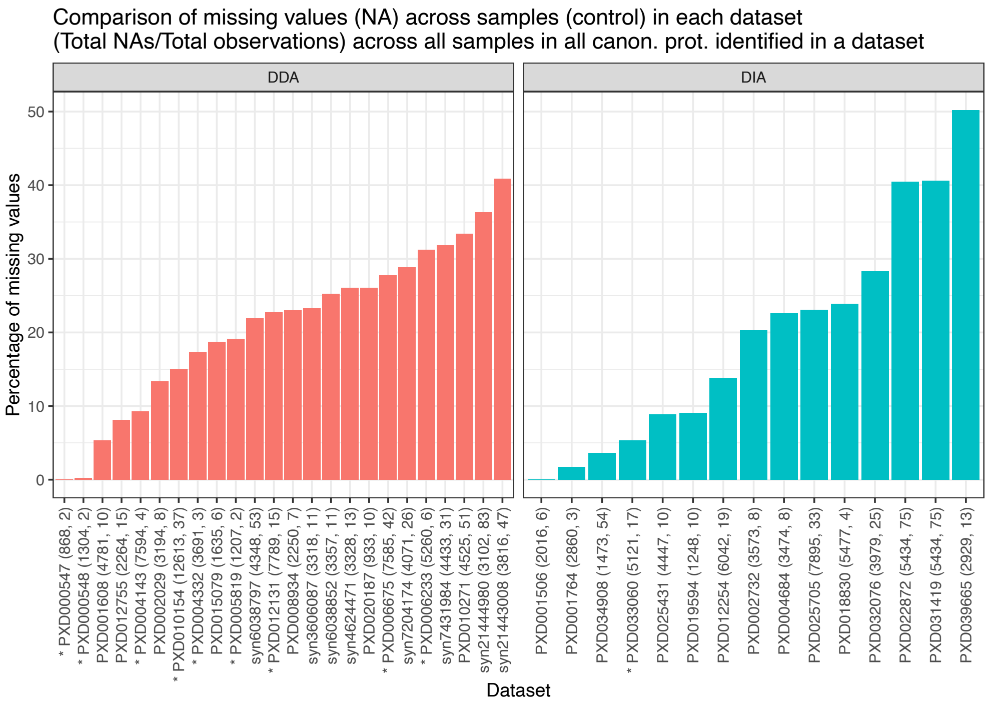


A

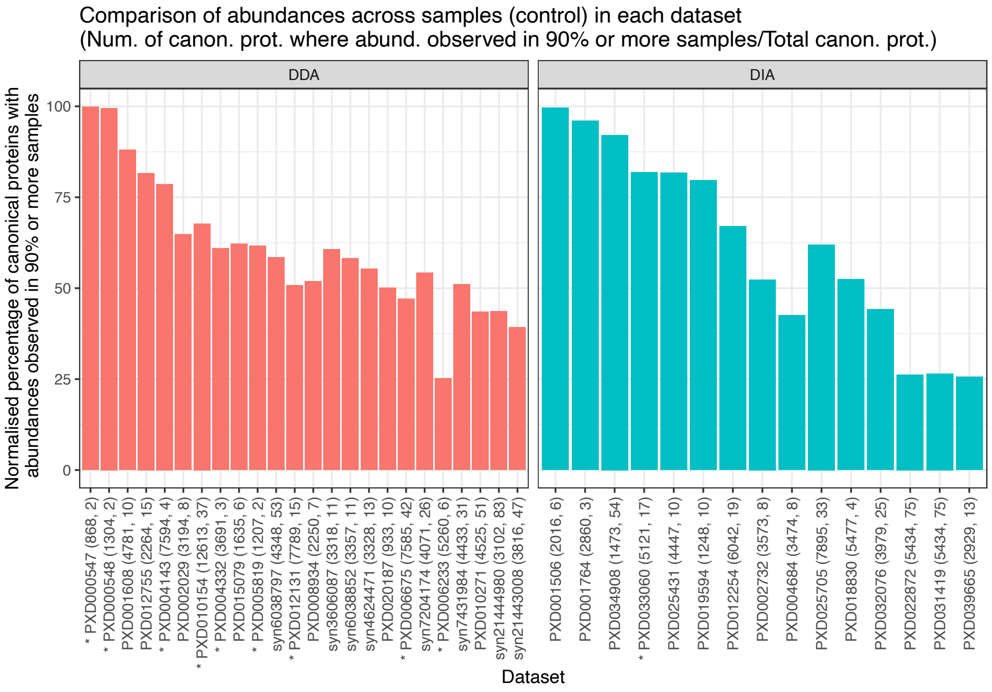


B

**Figure S3. (A)** Percentage of missing protein abundances in samples analysed from DDA[1] and DIA datasets. **(B)** Percentage of total canonical proteins in a dataset in which at least 90 percent samples had an abundance. * denotes datasets in which samples are fractionated. Numbers with parenthesis denotes number of genes identified and number of samples in that dataset respectively.

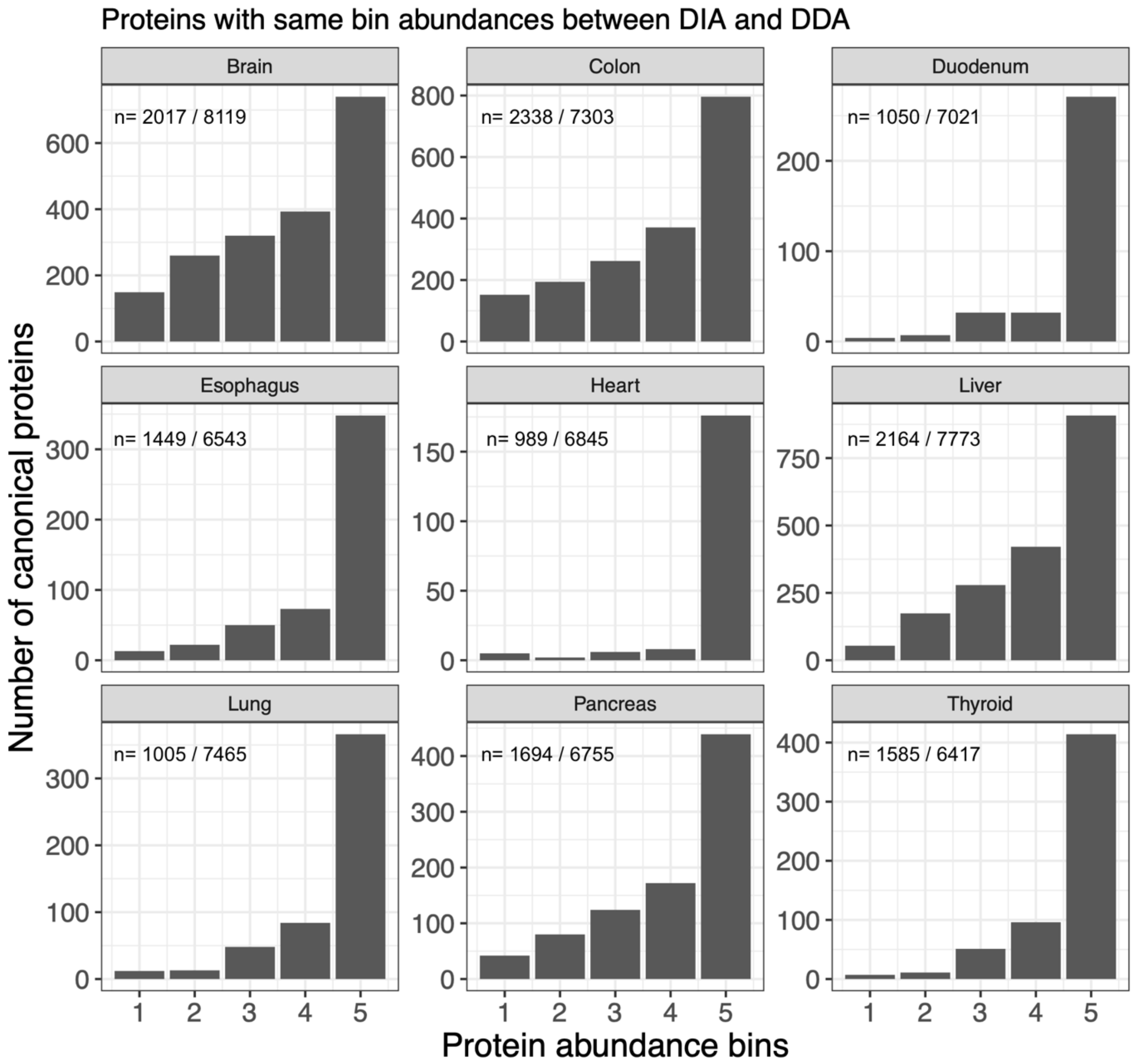


**Figure S4.** Comparison of binned protein abundances in tissues from DIA and DDA datasets. The number of canonical proteins that have the same binned protein abundances from DIA datasets and from our previous study with DDA[1] datasets. The number 'n' in each panel shows the (pairs with same binned values between DIA and DDA) / (total number of pairs with a binned value in either DIA or DDA).

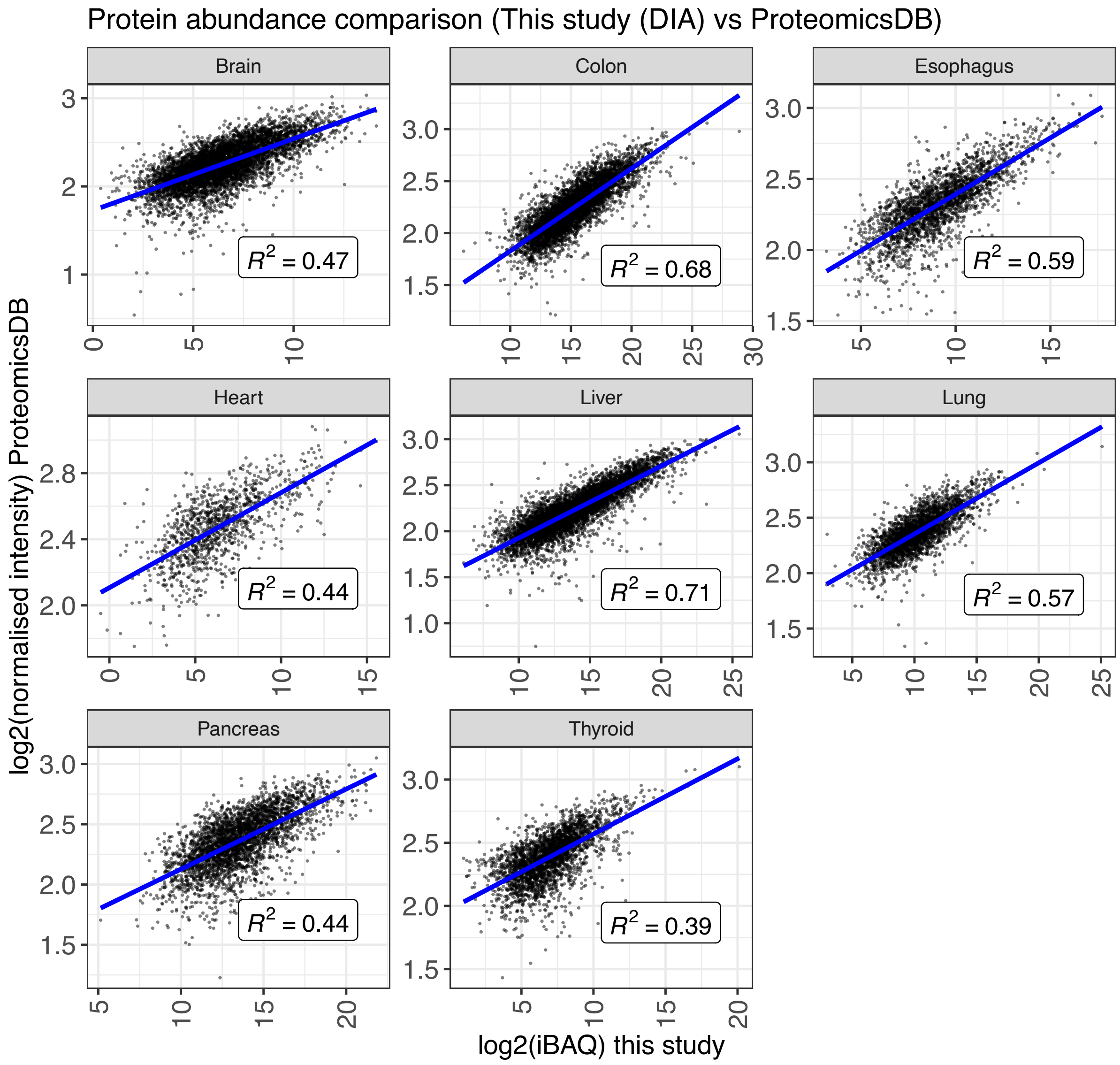


**Figure S5.** Comparison of protein expression across various human organs at baseline condition analysed from DIA in this study and from ProteomicsDB.

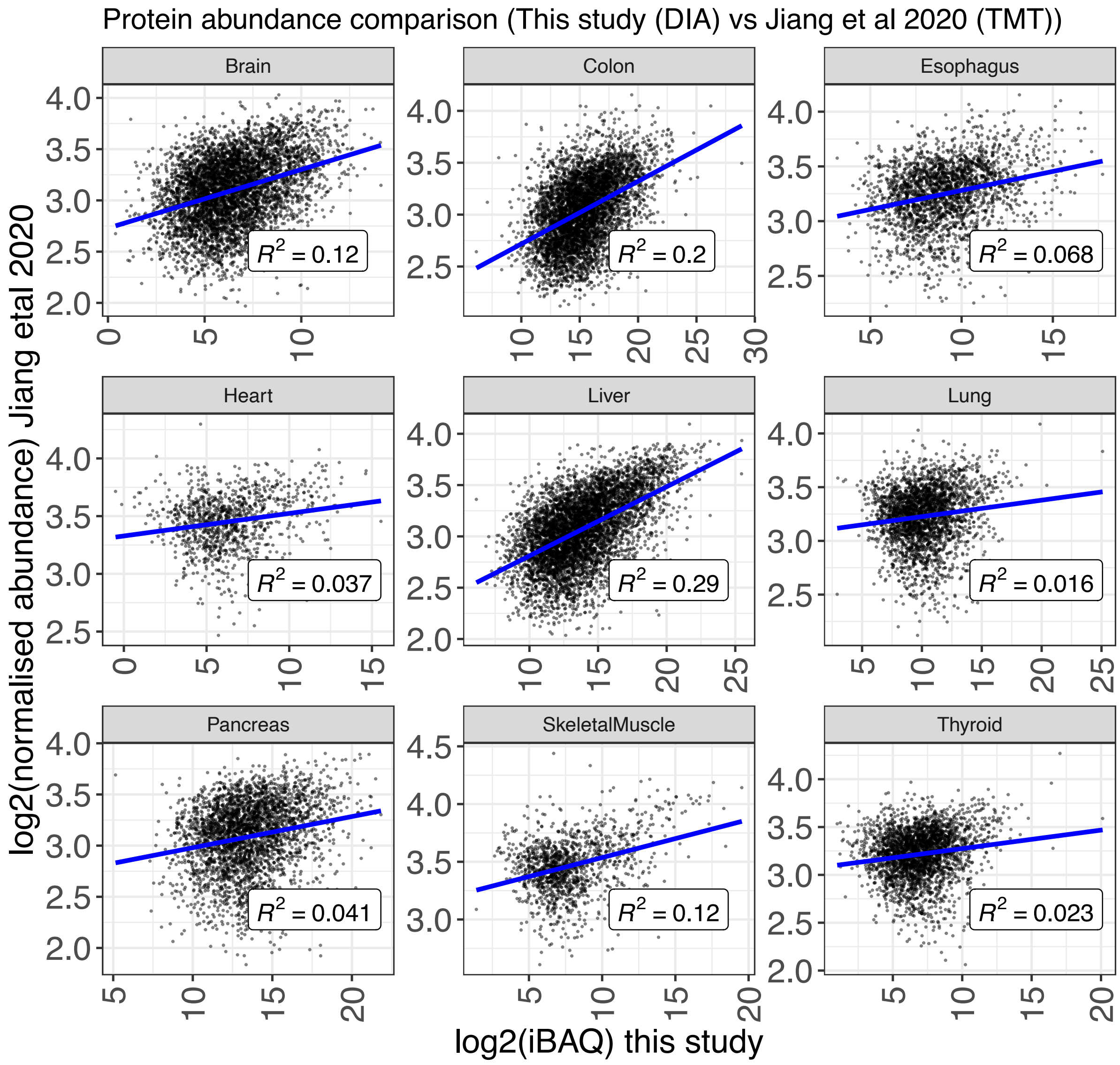


**Figure S6.** Comparison of protein expression across various human organs at baseline condition analysed from DIA in this study and from Jiang et. al.[2] using TMT-labelling method.

Supplementary tables

**Supplementary Table 1:** Canonical protein abundances (iBAQ) across various tissues.

**Supplementary Table 2:** Binned canonical protein abundances across various tissues.

**Supplementary Table 3:** Canonical proteins identified by DIA only, DDA only and by both techniques.
