## Supporting File 3 for "Integrated View of Baseline Protein Expression in Human Tissues using public Data Independent Acquisition datasets"

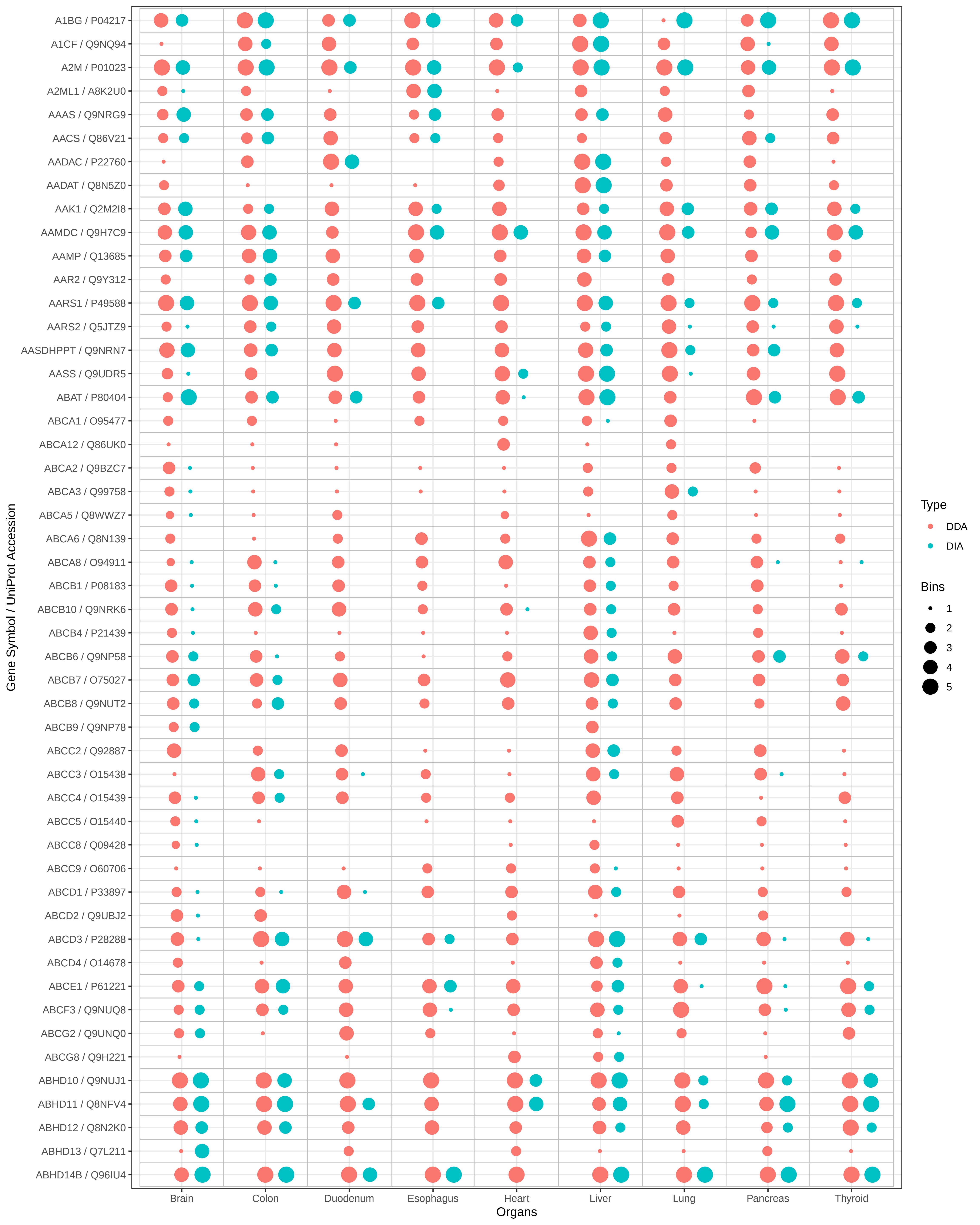
A1BG / P04217
A1CF / Q9NQ94
A2M / P01023
A2ML1 / A8K2U0
AAAS / Q9NRG9
AACS / Q86V21
AADAC / P22760
AADAT / Q8N5Z0
AAK1 / Q2M2I8
AAMDC / Q9H7C9
AAMP / Q13685
AAR2 / Q9Y312
AARS1 / P49588
AARS2 / Q5JTZ9
AASDHPPT / Q9NRN7
AASS / Q9UDR5
ABAT / P80404
ABCA1 / O95477
ABCA12 / Q86UK0
ABCA2 / Q9BZC7
ABCA3 / Q99758
ABCA5 / Q8WWZ7
ABCA6 / Q8N139
ABCA8 / O94911
ABCB1 / P08183
ABCB10 / Q9NRK6
ABCB4 / P21439
ABCB6 / Q9NP58
ABCB7 / O75027
ABCB8 / Q9NUT2
ABCB9 / Q9NP78
ABCC2 / Q92887
ABCC3 / O15438
ABCC4 / O15439
ABCC5 / O15440
ABCC8 / Q09428
ABCC9 / O60706
ABCD1 / P33897
ABCD2 / Q9UBJ2
ABCD3 / P28288
ABCD4 / O14678
ABCE1 / P61221
ABCF3 / Q9NUQ8
ABCG2 / Q9UNQ0
ABCG8 / Q9H221
ABHD10 / Q9NUJ1
ABHD11 / Q8NFV4
ABHD12 / Q8N2K0
ABHD13 / Q7L211
ABHD14B / Q96IU4
Gene Symbol / UniProt Accession
Brain
Colon
Duodenum
Esophagus
Heart
Liver
Lung
Pancreas
Thyroid
Organs
Type
DDA
DIA
Bins
1
2
3
4
5

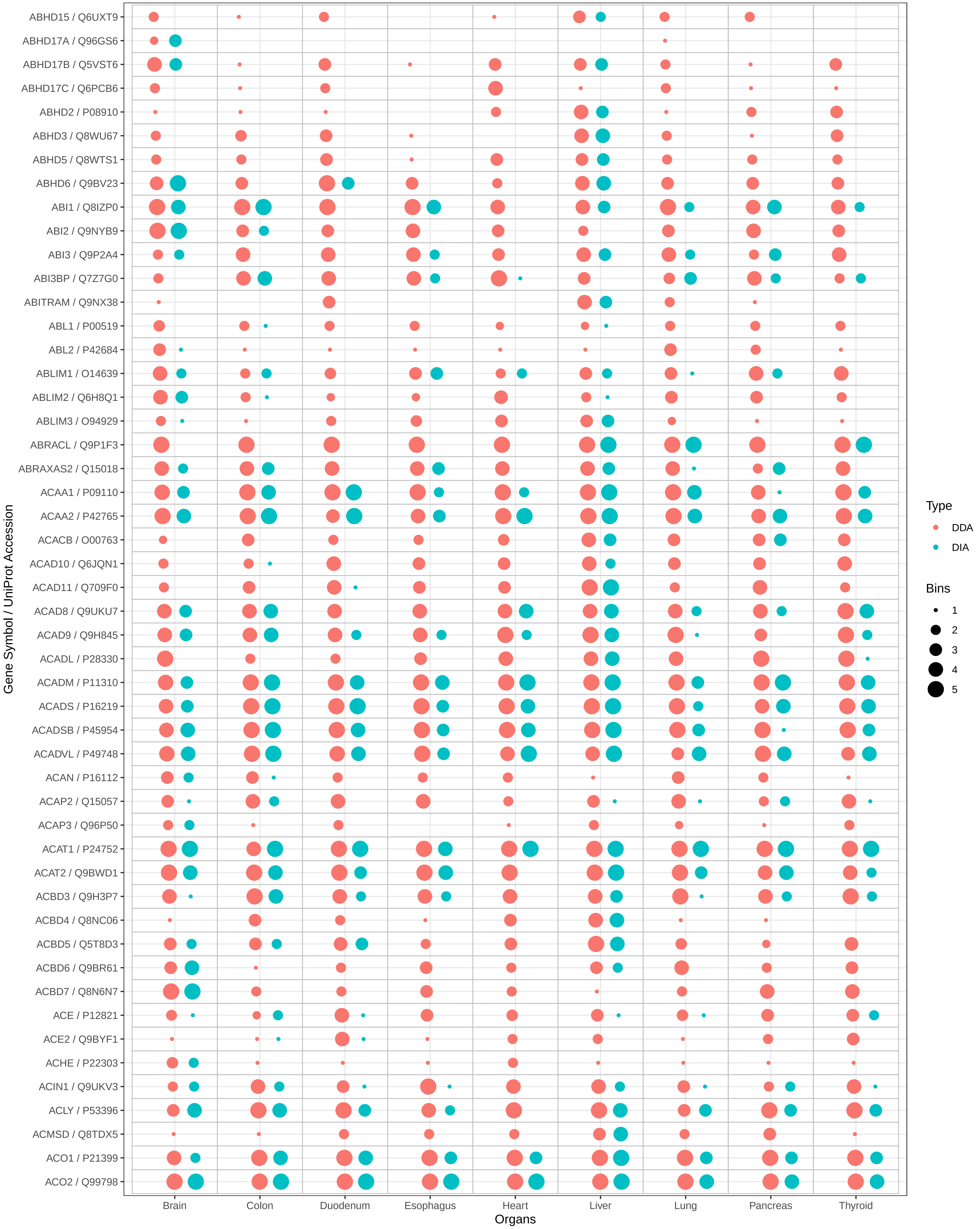
ABHD15 / Q6UXT9
ABHD17A / Q96GS6
ABHD17B / Q5VST6
ABHD17C / Q6PCB6
ABHD2 / P08910
ABHD3 / Q8WU67
ABHD5 / Q8WTS1
ABHD6 / Q9BV23
ABI1 / Q8IZP0
ABI2 / Q9NYB9
ABI3 / Q9P2A4
ABI3BP / Q7Z7G0
ABITRAM / Q9NX38
ABL1 / P00519
ABL2 / P42684
ABLIM1 / O14639
ABLIM2 / Q6H8Q1
ABLIM3 / O94929
ABRACL / Q9P1F3
ABRAXAS2 / Q15018
ACAA1 / P09110
ACAA2 / P42765
ACACB / O00763
ACAD10 / Q6JQN1
ACAD11 / Q709F0
ACAD8 / Q9UKU7
ACAD9 / Q9H845
ACADL / P28330
ACADM / P11310
ACADS / P16219
ACADSB / P45954
ACADVL / P49748
ACAN / P16112
ACAP2 / Q15057
ACAP3 / Q96P50
ACAT1 / P24752
ACAT2 / Q9BWD1
ACBD3 / Q9H3P7
ACBD4 / Q8NC06
ACBD5 / Q5T8D3
ACBD6 / Q9BR61
ACBD7 / Q8N6N7
ACE / P12821
ACE2 / Q9BYF1
ACHE / P22303
ACIN1 / Q9UKV3
ACLY / P53396
ACMSD / Q8TDX5
ACO1 / P21399
ACO2 / Q99798
Gene Symbol / UniProt Accession
Brain
Colon
Duodenum
Esophagus
Heart
Liver
Lung
Pancreas
Thyroid
Organs
Type
DDA
DIA
Bins
1
2
3
4
5

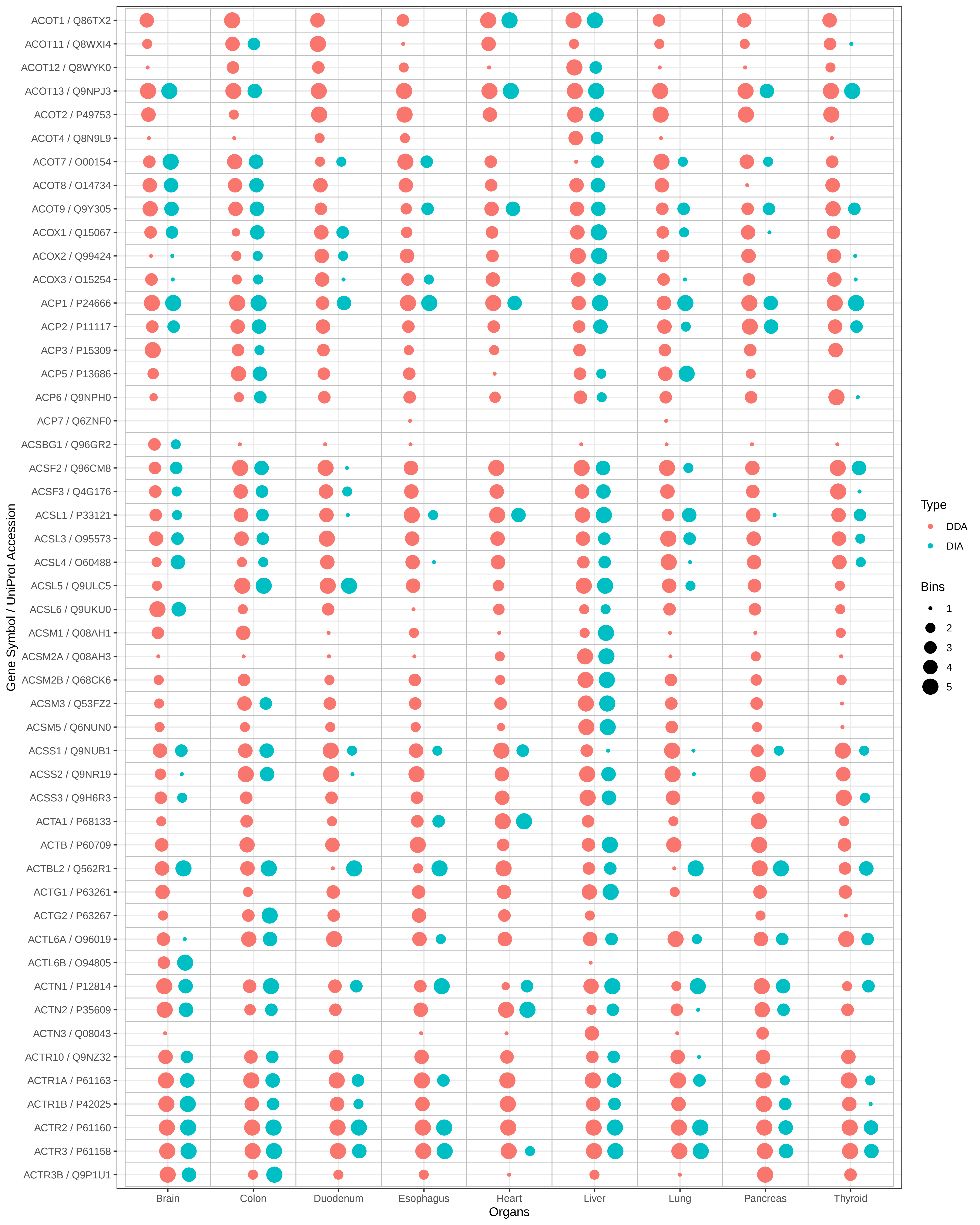

ACOT1 / Q86TX2
ACOT11 / Q8WXI4
ACOT12 / Q8WYK0
ACOT13 / Q9NPJ3
ACOT2 / P49753
ACOT4 / Q8N9L9
ACOT7 / O00154
ACOT8 / O14734
ACOT9 / Q9Y305
ACOX1 / Q15067
ACOX2 / Q99424
ACOX3 / O15254
ACP1 / P24666
ACP2 / P11117
ACP3 / P15309
ACP5 / P13686
ACP6 / Q9NPH0
ACP7 / Q6ZNF0
ACSBG1 / Q96GR2
ACSF2 / Q96CM8
ACSF3 / Q4G176
ACSL1 / P33121
ACSL3 / O95573
ACSL4 / O60488
ACSL5 / Q9ULC5
ACSL6 / Q9UKU0
ACSM1 / Q08AH1
ACSM2A / Q08AH3
ACSM2B / Q68CK6
ACSM3 / Q53FZ2
ACSM5 / Q6NUN0
ACSS1 / Q9NUB1
ACSS2 / Q9NR19
ACSS3 / Q9H6R3
ACTA1 / P68133
ACTB / P60709
ACTBL2 / Q562R1
ACTG1 / P63261
ACTG2 / P63267
ACTL6A / O96019
ACTL6B / O94805
ACTN1 / P12814
ACTN2 / P35609
ACTN3 / Q08043
ACTR10 / Q9NZ32
ACTR1A / P61163
ACTR1B / P42025
ACTR2 / P61160
ACTR3 / P61158
ACTR3B / Q9P1U1
Gene Symbol / UniProt Accession
Brain
Colon
Duodenum
Esophagus
Heart
Liver
Lung
Pancreas
Thyroid
Organs
Type
DDA
DIA
Bins
1
2
3
4
5

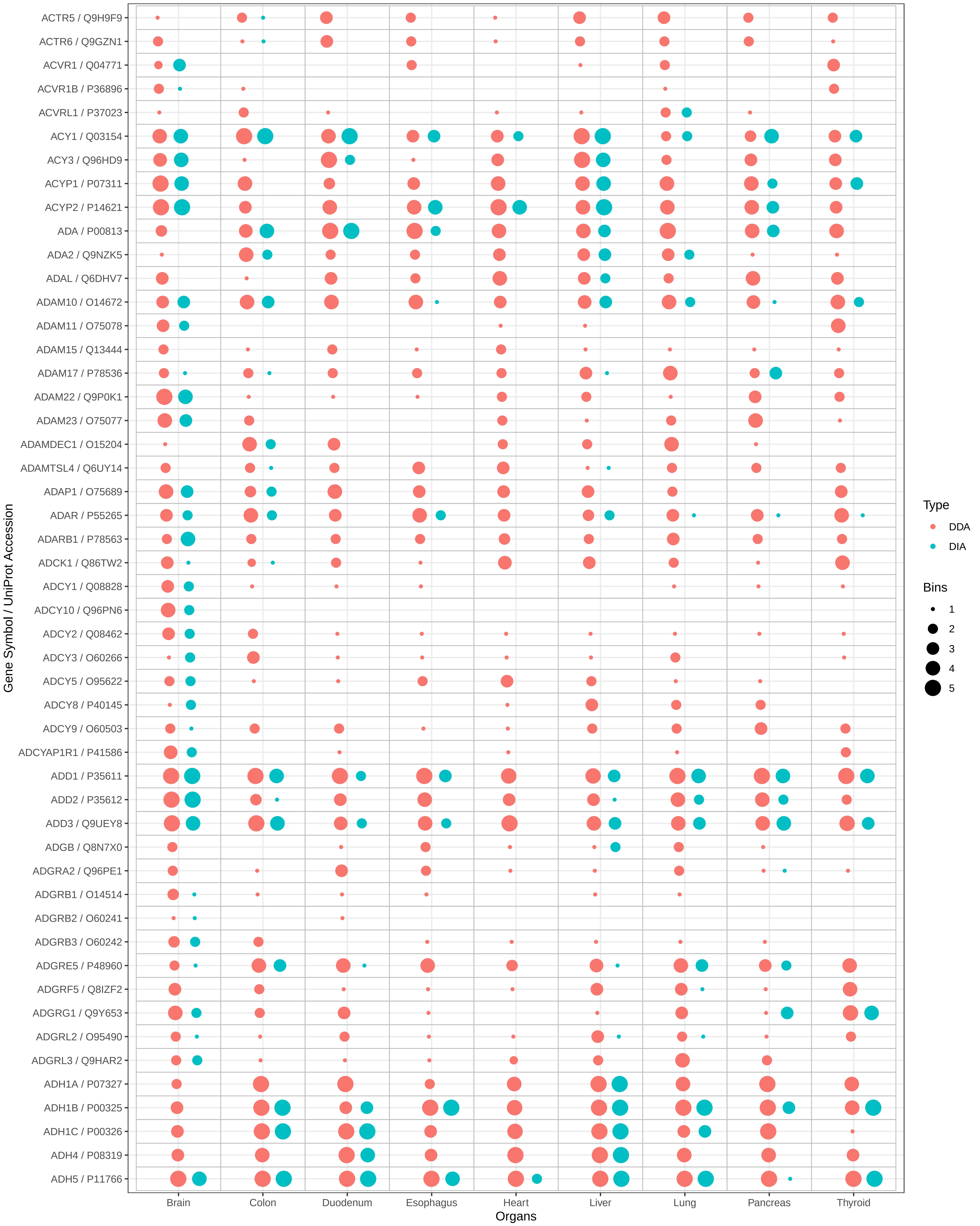

ACTR5 / Q9H9F9
ACTR6 / Q9GZN1
ACVR1 / Q04771
ACVR1B / P36896
ACVRL1 / P37023
ACY1 / Q03154
ACY3 / Q96HD9
ACYP1 / P07311
ACYP2 / P14621
ADA / P00813
ADA2 / Q9NZK5
ADAL / Q6DHV7
ADAM10 / O14672
ADAM11 / O75078
ADAM15 / Q13444
ADAM17 / P78536
ADAM22 / Q9P0K1
ADAM23 / O75077
ADAMDEC1 / O15204
ADAMTSL4 / Q6UY14
ADAP1 / O75689
ADAR / P55265
ADARB1 / P78563
ADCK1 / Q86TW2
ADCY1 / Q08828
ADCY10 / Q96PN6
ADCY2 / Q08462
ADCY3 / O60266
ADCY5 / O95622
ADCY8 / P40145
ADCY9 / O60503
ADCYAP1R1 / P41586
ADD1 / P35611
ADD2 / P35612
ADD3 / Q9UEY8
ADGB / Q8N7X0
ADGRA2 / Q96PE1
ADGRB1 / O14514
ADGRB2 / O60241
ADGRB3 / O60242
ADGRE5 / P48960
ADGRF5 / Q8IZF2
ADGRG1 / Q9Y653
ADGRL2 / O95490
ADGRL3 / Q9HAR2
ADH1A / P07327
ADH1B / P00325
ADH1C / P00326
ADH4 / P08319
ADH5 / P11766
Gene Symbol / UniProt Accession
Brain
Colon
Duodenum
Esophagus
Heart
Liver
Lung
Pancreas
Thyroid
Organs
Type
DDA
DIA
Bins
1
2
3
4
5

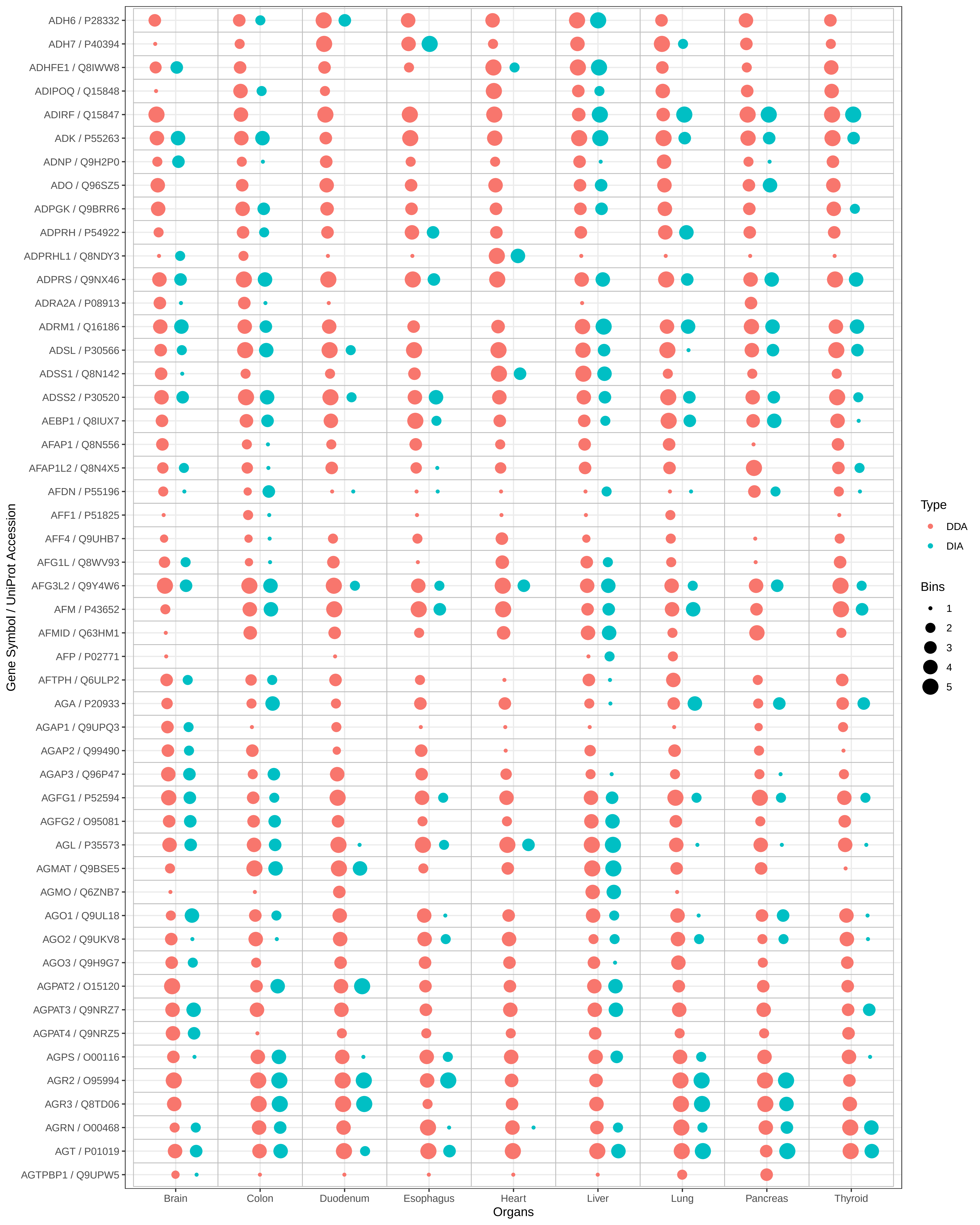

ADH6 / P28332
ADH7 / P40394
ADHFE1 / Q8IWW8
ADIPOQ / Q15848
ADIRF / Q15847
ADK / P55263
ADNP / Q9H2P0
ADO / Q96SZ5
ADPGK / Q9BRR6
ADPRH / P54922
ADPRHL1 / Q8NDY3
ADPRS / Q9NX46
ADRA2A / P08913
ADRM1 / Q16186
ADSL / P30566
ADSS1 / Q8N142
ADSS2 / P30520
AEBP1 / Q8IUX7
AFAP1 / Q8N556
AFAP1L2 / Q8N4X5
AFDN / P55196
AFF1 / P51825
AFF4 / Q9UHB7
AFG1L / Q8WV93
AFG3L2 / Q9Y4W6
AFM / P43652
AFMID / Q63HM1
AFP / P02771
AFTPH / Q6ULP2
AGA / P20933
AGAP1 / Q9UPQ3
AGAP2 / Q99490
AGAP3 / Q96P47
AGFG1 / P52594
AGFG2 / O95081
AGL / P35573
AGMAT / Q9BSE5
AGMO / Q6ZNB7
AGO1 / Q9UL18
AGO2 / Q9UKV8
AGO3 / Q9H9G7
AGPAT2 / O15120
AGPAT3 / Q9NRZ7
AGPAT4 / Q9NRZ5
AGPS / O00116
AGR2 / O95994
AGR3 / Q8TD06
AGRN / O00468
AGT / P01019
AGTPBP1 / Q9UPW5
Gene Symbol / UniProt Accession
Brain
Colon
Duodenum
Esophagus
Heart
Liver
Lung
Pancreas
Thyroid
Organs
Type
DDA
DIA
Bins
1
2
3
4
5

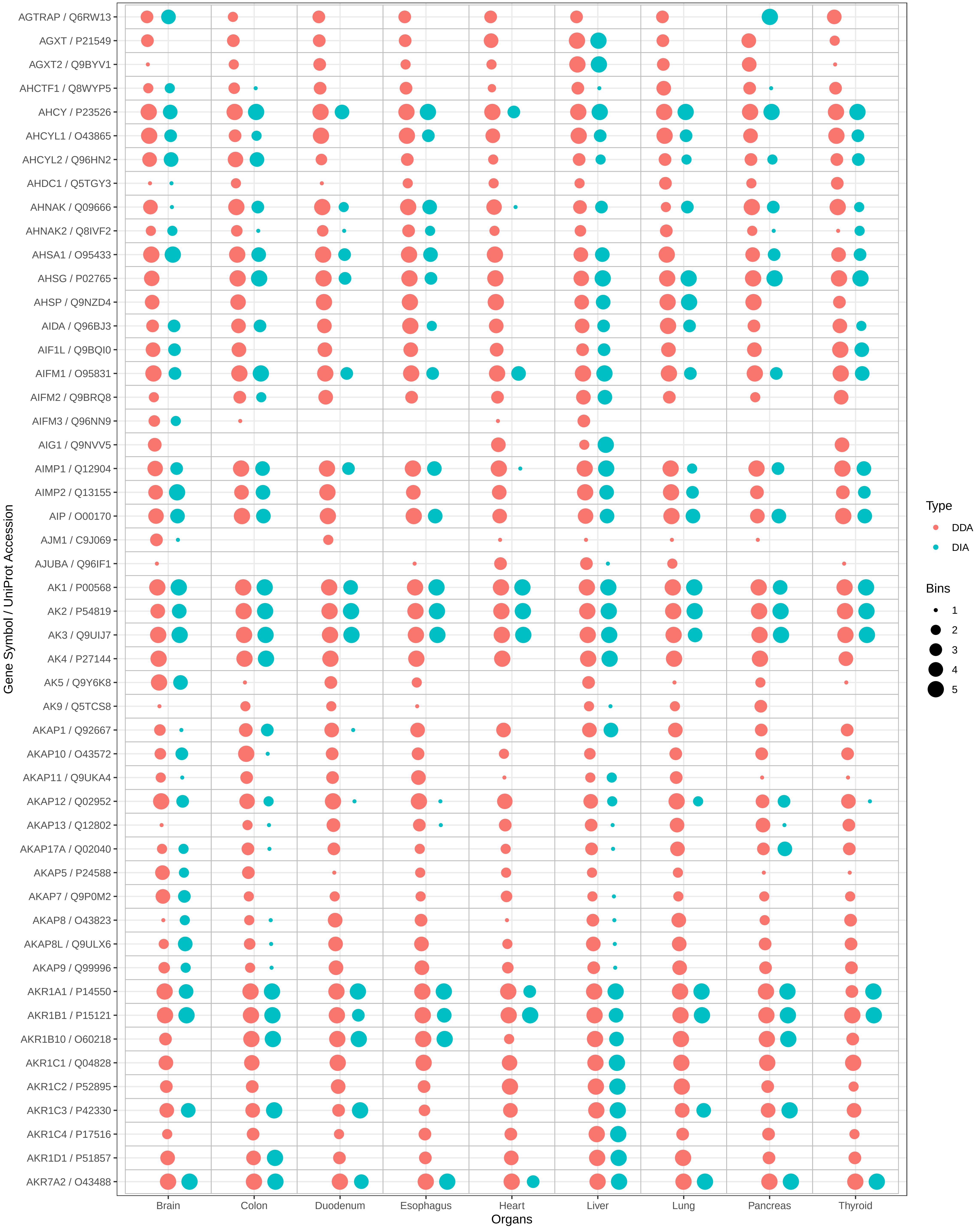
AGTRAP / Q6RW13
AGXT / P21549
AGXT2 / Q9BYV1
AHCTF1 / Q8WYP5
AHCY / P23526
AHCYL1 / O43865
AHCYL2 / Q96HN2
AHDC1 / Q5TGY3
AHNAK / Q09666
AHNAK2 / Q8IVF2
AHSA1 / O95433
AHSG / P02765
AHSP / Q9NZD4
AIDA / Q96BJ3
AIF1L / Q9BQI0
AIFM1 / O95831
AIFM2 / Q9BRQ8
AIFM3 / Q96NN9
AIG1 / Q9NVV5
AIMP1 / Q12904
AIMP2 / Q13155
AIP / O00170
AJM1 / C9J069
AJUBA / Q96IF1
AK1 / P00568
AK2 / P54819
AK3 / Q9UIJ7
AK4 / P27144
AK5 / Q9Y6K8
AK9 / Q5TCS8
AKAP1 / Q92667
AKAP10 / O43572
AKAP11 / Q9UKA4
AKAP12 / Q02952
AKAP13 / Q12802
AKAP17A / Q02040
AKAP5 / P24588
AKAP7 / Q9P0M2
AKAP8 / O43823
AKAP8L / Q9ULX6
AKAP9 / Q99996
AKR1A1 / P14550
AKR1B1 / P15121
AKR1B10 / O60218
AKR1C1 / Q04828
AKR1C2 / P52895
AKR1C3 / P42330
AKR1C4 / P17516
AKR1D1 / P51857
AKR7A2 / O43488
Gene Symbol / UniProt Accession
Brain
Colon
Duodenum
Esophagus
Heart
Liver
Lung
Pancreas
Thyroid
Organs
Type
DDA
DIA
Bins
1
2
3
4
5

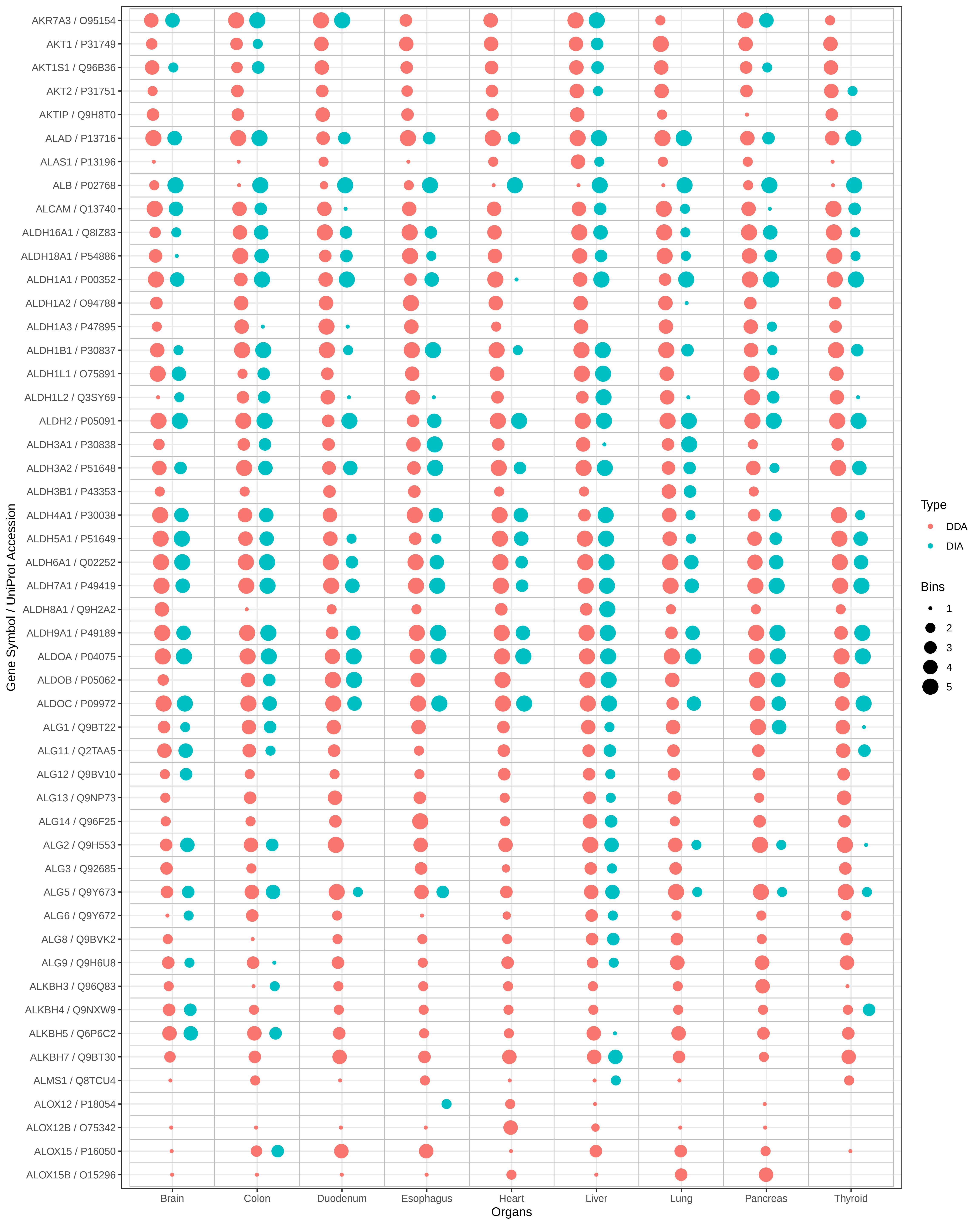

AKR7A3 / O95154
AKT1 / P31749
AKT1S1 / Q96B36
AKT2 / P31751
AKTIP / Q9H8T0
ALAD / P13716
ALAS1 / P13196
ALB / P02768
ALCAM / Q13740
ALDH16A1 / Q8IZ83
ALDH18A1 / P54886
ALDH1A1 / P00352
ALDH1A2 / O94788
ALDH1A3 / P47895
ALDH1B1 / P30837
ALDH1L1 / O75891
ALDH1L2 / Q3SY69
ALDH2 / P05091
ALDH3A1 / P30838
ALDH3A2 / P51648
ALDH3B1 / P43353
ALDH4A1 / P30038
ALDH5A1 / P51649
ALDH6A1 / Q02252
ALDH7A1 / P49419
ALDH8A1 / Q9H2A2
ALDH9A1 / P49189
ALDOA / P04075
ALDOB / P05062
ALDOC / P09972
ALG1 / Q9BT22
ALG11 / Q2TAA5
ALG12 / Q9BV10
ALG13 / Q9NP73
ALG14 / Q96F25
ALG2 / Q9H553
ALG3 / Q92685
ALG5 / Q9Y673
ALG6 / Q9Y672
ALG8 / Q9BVK2
ALG9 / Q9H6U8
ALKBH3 / Q96Q83
ALKBH4 / Q9NXW9
ALKBH5 / Q6P6C2
ALKBH7 / Q9BT30
ALMS1 / Q8TCU4
ALOX12 / P18054
ALOX12B / O75342
ALOX15 / P16050
ALOX15B / O15296
Gene Symbol / UniProt Accession
Brain
Colon
Duodenum
Esophagus
Heart
Liver
Lung
Pancreas
Thyroid
Organs
Type
DDA
DIA
Bins
1
2
3
4
5

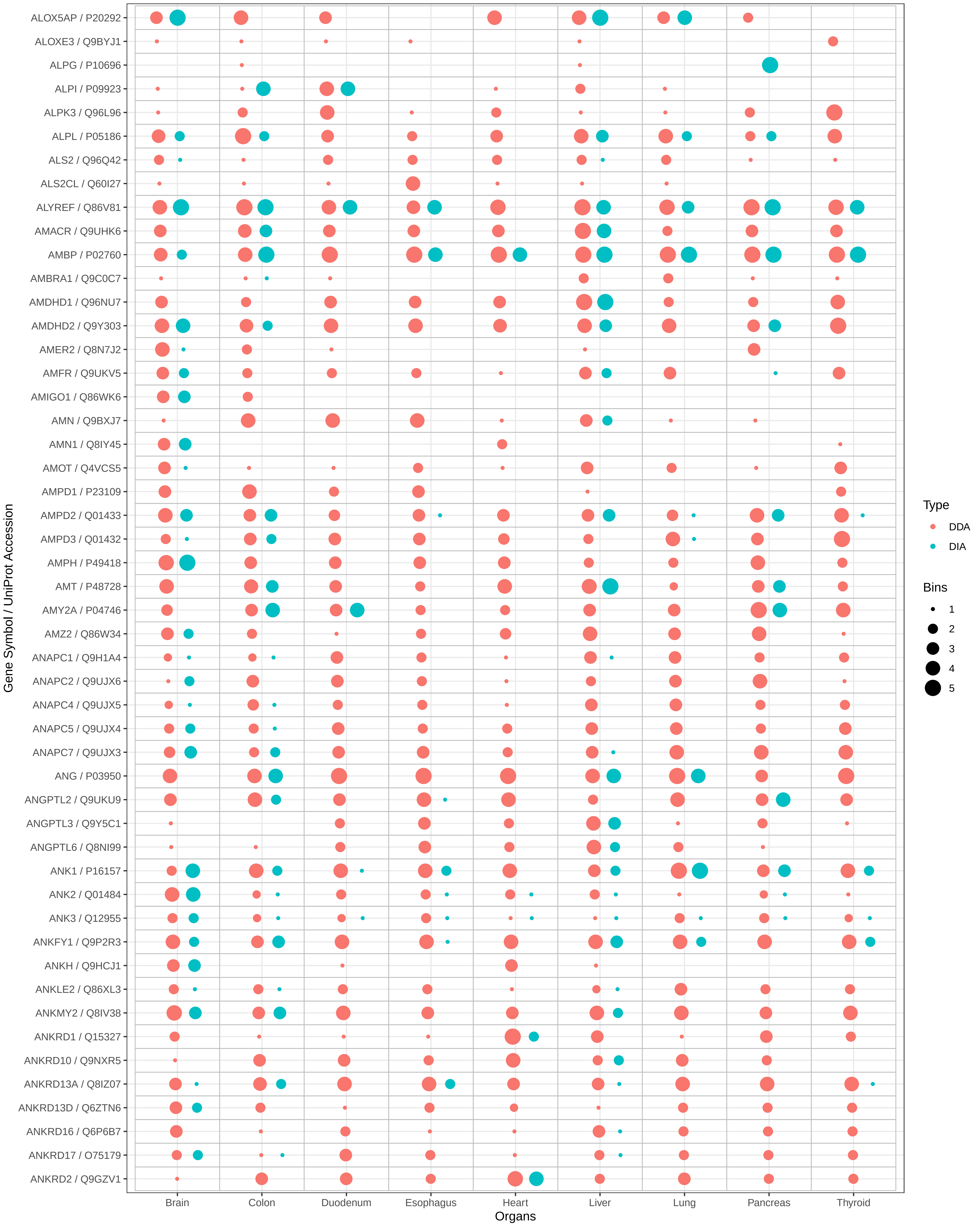
ALOX5AP / P20292
ALOXE3 / Q9BYJ1
ALPG / P10696
ALPI / P09923
ALPK3 / Q96L96
ALPL / P05186
ALS2 / Q96Q42
ALS2CL / Q60I27
ALYREF / Q86V81
AMACR / Q9UHK6
AMBP / P02760
AMBRA1 / Q9C0C7
AMDHD1 / Q96NU7
AMDHD2 / Q9Y303
AMER2 / Q8N7J2
AMFR / Q9UKV5
AMIGO1 / Q86WK6
AMN / Q9BXJ7
AMN1 / Q8IY45
AMOT / Q4VCS5
AMPD1 / P23109
AMPD2 / Q01433
AMPD3 / Q01432
AMPH / P49418
AMT / P48728
AMY2A / P04746
AMZ2 / Q86W34
ANAPC1 / Q9H1A4
ANAPC2 / Q9UJX6
ANAPC4 / Q9UJX5
ANAPC5 / Q9UJX4
ANAPC7 / Q9UJX3
ANG / P03950
ANGPTL2 / Q9UKU9
ANGPTL3 / Q9Y5C1
ANGPTL6 / Q8NI99
ANK1 / P16157
ANK2 / Q01484
ANK3 / Q12955
ANKFY1 / Q9P2R3
ANKH / Q9HCJ1
ANKLE2 / Q86XL3
ANKMY2 / Q8IV38
ANKRD1 / Q15327
ANKRD10 / Q9NXR5
ANKRD13A / Q8IZ07
ANKRD13D / Q6ZTN6
ANKRD16 / Q6P6B7
ANKRD17 / O75179
ANKRD2 / Q9GZV1
Gene Symbol / UniProt Accession
Brain
Colon
Duodenum
Esophagus
Heart
Liver
Lung
Pancreas
Thyroid
Organs
Type
DDA
DIA
Bins
1
2
3
4
5

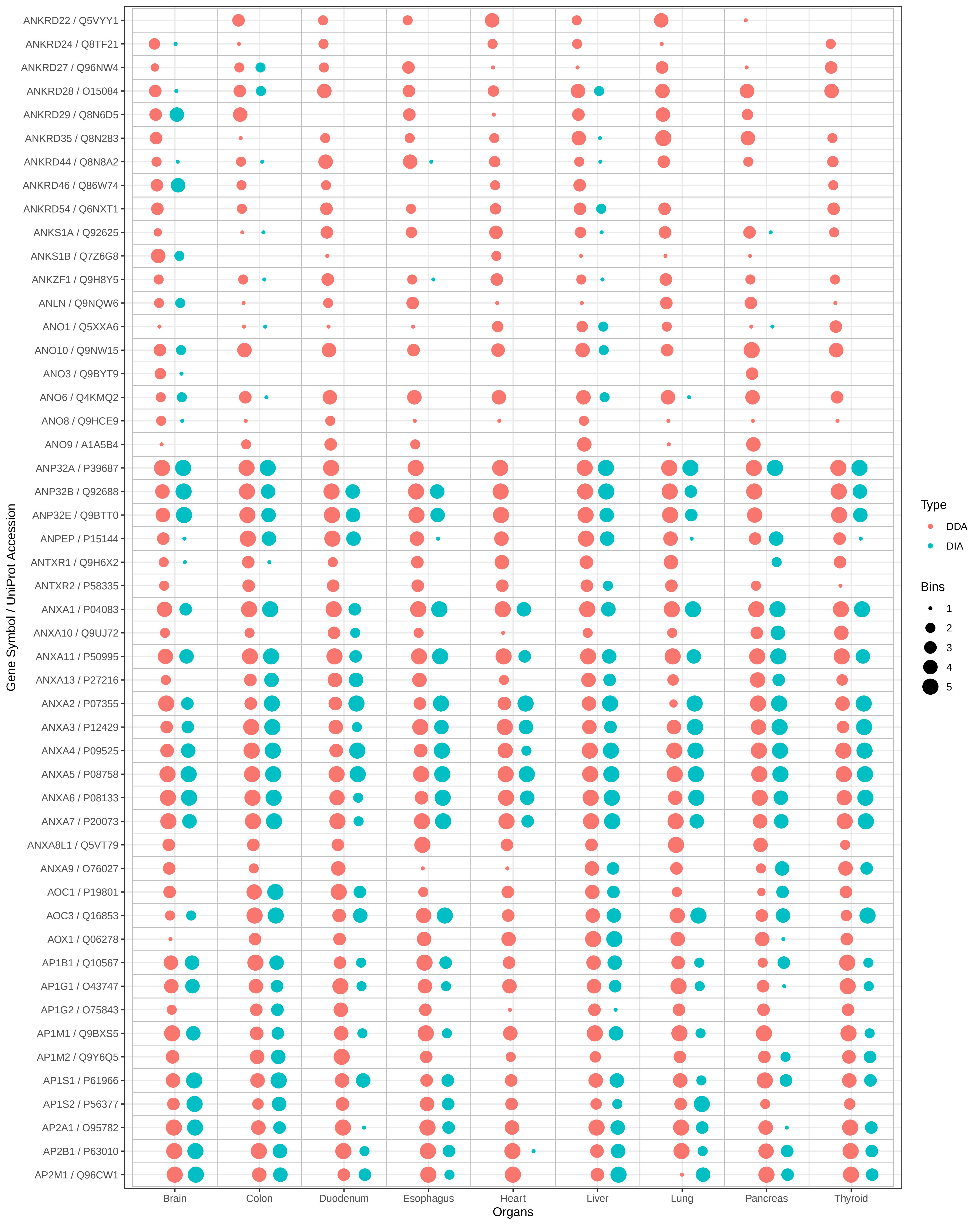

ANKRD22 / Q5VYY1
ANKRD24 / Q8TF21
ANKRD27 / Q96NW4
ANKRD28 / O15084
ANKRD29 / Q8N6D5
ANKRD35 / Q8N283
ANKRD44 / Q8N8A2
ANKRD46 / Q86W74
ANKRD54 / Q6NXT1
ANKS1A / Q92625
ANKS1B / Q7Z6G8
ANKZF1 / Q9H8Y5
ANLN / Q9NQW6
ANO1 / Q5XXA6
ANO10 / Q9NW15
ANO3 / Q9BYT9
ANO6 / Q4KMQ2
ANO8 / Q9HCE9
ANO9 / A1A5B4
ANP32A / P39687
ANP32B / Q92688
ANP32E / Q9BTT0
ANPEP / P15144
ANTXR1 / Q9H6X2
ANTXR2 / P58335
ANXA1 / P04083
ANXA10 / Q9UJ72
ANXA11 / P50995
ANXA13 / P27216
ANXA2 / P07355
ANXA3 / P12429
ANXA4 / P09525
ANXA5 / P08758
ANXA6 / P08133
ANXA7 / P20073
ANXA8L1 / Q5VT79
ANXA9 / O76027
AOC1 / P19801
AOC3 / Q16853
AOX1 / Q06278
AP1B1 / Q10567
AP1G1 / O43747
AP1G2 / O75843
AP1M1 / Q9BXS5
AP1M2 / Q9Y6Q5
AP1S1 / P61966
AP1S2 / P56377
AP2A1 / O95782
AP2B1 / P63010
AP2M1 / Q96CW1
Gene Symbol / UniProt Accession
Brain
Colon
Duodenum
Esophagus
Heart
Liver
Lung
Pancreas
Thyroid
Organs
Type
DDA
DIA
Bins
1
2
3
4
5

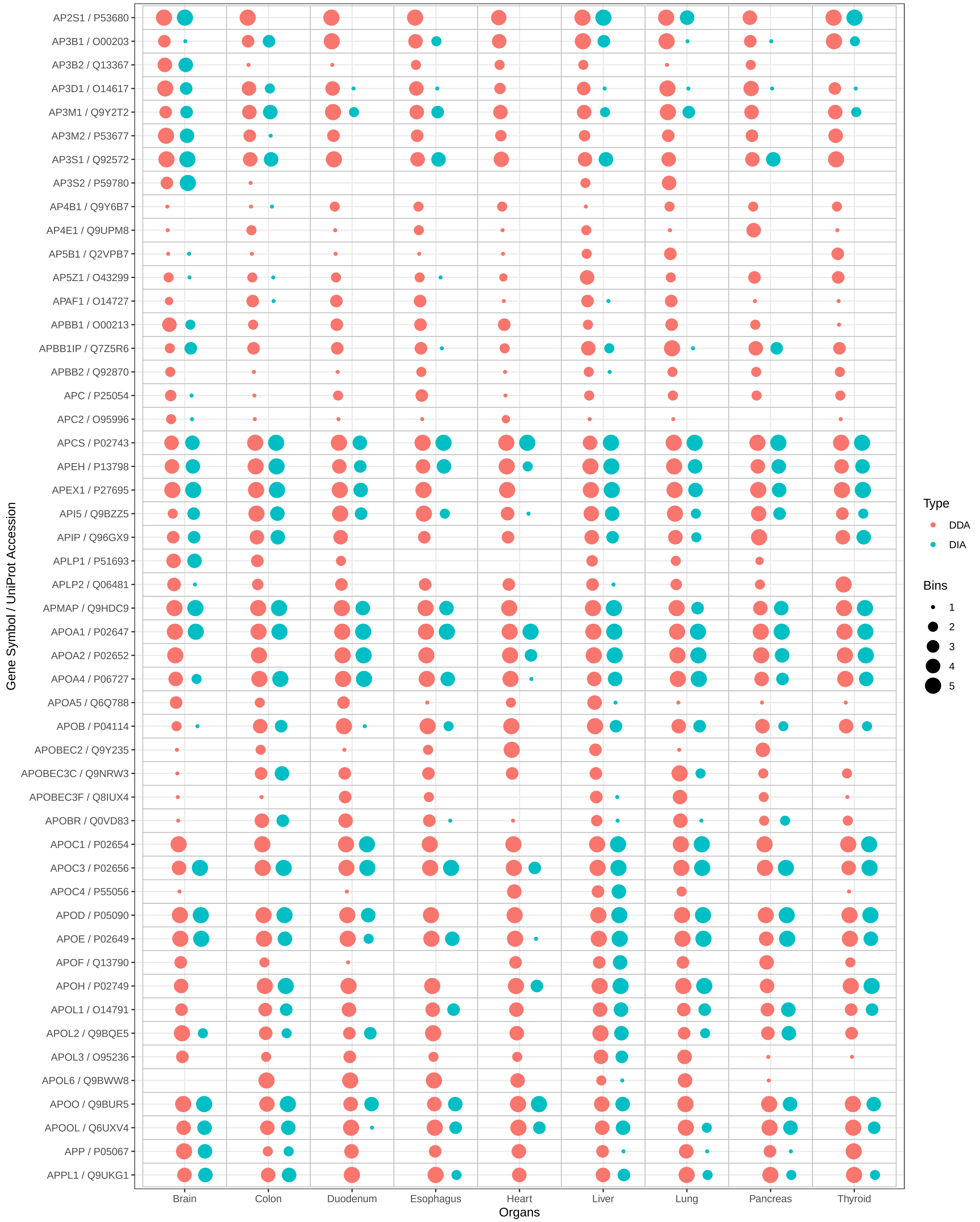

AP2S1 / P53680
AP3B1 / O00203
AP3B2 / Q13367
AP3D1 / O14617
AP3M1 / Q9Y2T2
AP3M2 / P53677
AP3S1 / Q92572
AP3S2 / P59780
AP4B1 / Q9Y6B7
AP4E1 / Q9UPM8
AP5B1 / Q2VPB7
AP5Z1 / O43299
APAF1 / O14727
APBB1 / O00213
APBB1IP / Q7Z5R6
APBB2 / Q92870
APC / P25054
APC2 / O95996
APCS / P02743
APEH / P13798
APEX1 / P27695
API5 / Q9BZZ5
APIP / Q96GX9
APLP1 / P51693
APLP2 / Q06481
APMAP / Q9HDC9
APOA1 / P02647
APOA2 / P02652
APOA4 / P06727
APOA5 / Q6Q788
APOB / P04114
APOBEC2 / Q9Y235
APOBEC3C / Q9NRW3
APOBEC3F / Q8IUX4
APOBR / Q0VD83
APOC1 / P02654
APOC3 / P02656
APOC4 / P55056
APOD / P05090
APOE / P02649
APOF / Q13790
APOH / P02749
APOL1 / O14791
APOL2 / Q9BQE5
APOL3 / O95236
APOL6 / Q9BWW8
APOO / Q9BUR5
APOOL / Q6UXV4
APP / P05067
APPL1 / Q9UKG1
Gene Symbol / UniProt Accession
Brain
Colon
Duodenum
Esophagus
Heart
Liver
Lung
Pancreas
Thyroid
Organs
Type
DDA
DIA
Bins
1
2
3
4
5

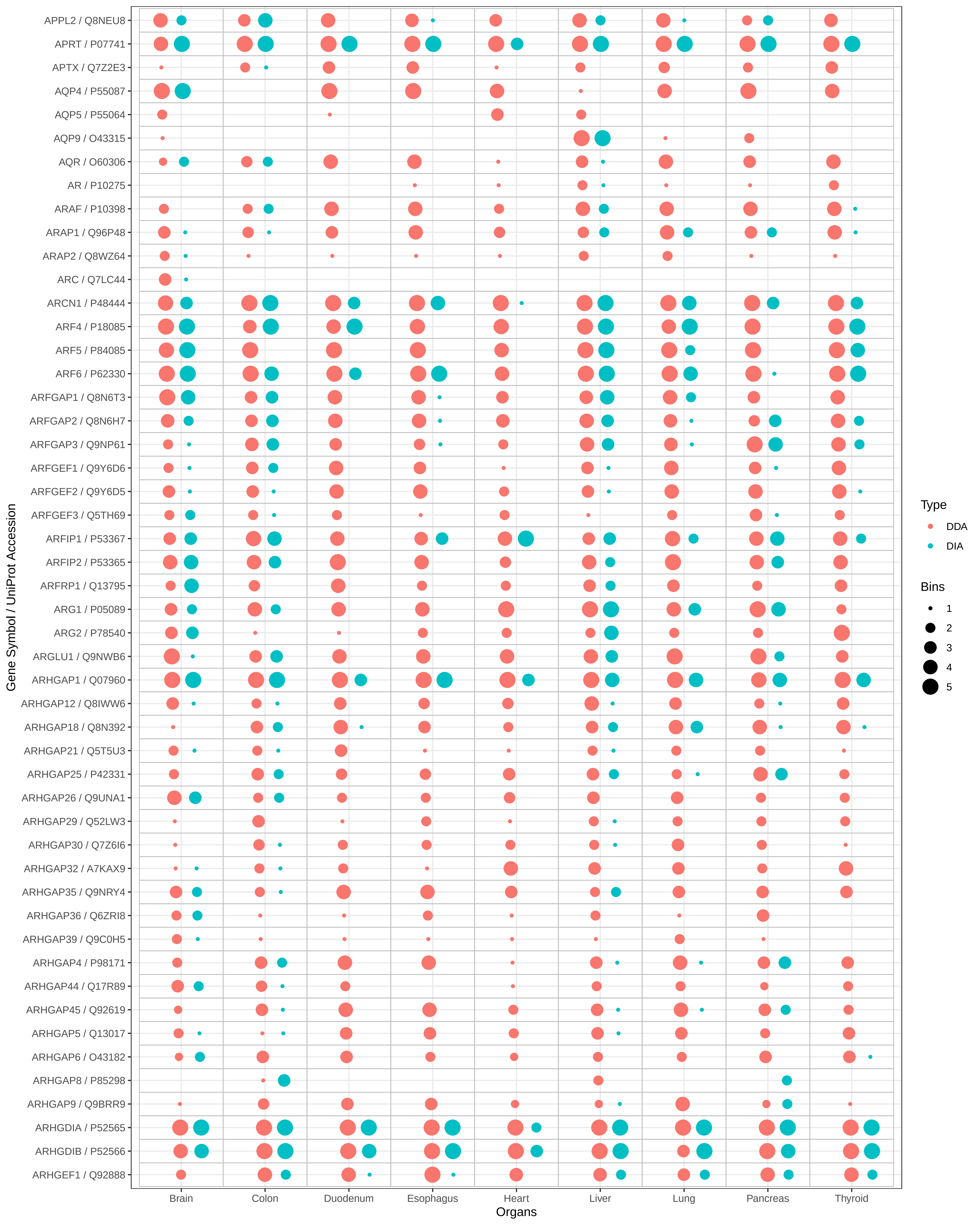

APPL2 / Q8NEU8
APRT / P07741
APTX / Q7Z2E3
AQP4 / P55087
AQP5 / P55064
AQP9 / O43315
AQR / O60306
AR / P10275
ARAF / P10398
ARAP1 / Q96P48
ARAP2 / Q8WZ64
ARC / Q7LC44
ARCN1 / P48444
ARF4 / P18085
ARF5 / P84085
ARF6 / P62330
ARFGAP1 / Q8N6T3
ARFGAP2 / Q8N6H7
ARFGAP3 / Q9NP61
ARFGEF1 / Q9Y6D6
ARFGEF2 / Q9Y6D5
ARFGEF3 / Q5TH69
ARFIP1 / P53367
ARFIP2 / P53365
ARFRP1 / Q13795
ARG1 / P05089
ARG2 / P78540
ARGLU1 / Q9NWB6
ARHGAP1 / Q07960
ARHGAP12 / Q8IWW6
ARHGAP18 / Q8N392
ARHGAP21 / Q5T5U3
ARHGAP25 / P42331
ARHGAP26 / Q9UNA1
ARHGAP29 / Q52LW3
ARHGAP30 / Q7Z6I6
ARHGAP32 / A7KAX9
ARHGAP35 / Q9NRY4
ARHGAP36 / Q6ZRI8
ARHGAP39 / Q9C0H5
ARHGAP4 / P98171
ARHGAP44 / Q17R89
ARHGAP45 / Q92619
ARHGAP5 / Q13017
ARHGAP6 / O43182
ARHGAP8 / P85298
ARHGAP9 / Q9BRR9
ARHGDIA / P52565
ARHGDIB / P52566
ARHGEF1 / Q92888
Gene Symbol / UniProt Accession
Brain
Colon
Duodenum
Esophagus
Heart
Liver
Lung
Pancreas
Thyroid
Organs
Type
DDA
DIA
Bins
1
2
3
4
5

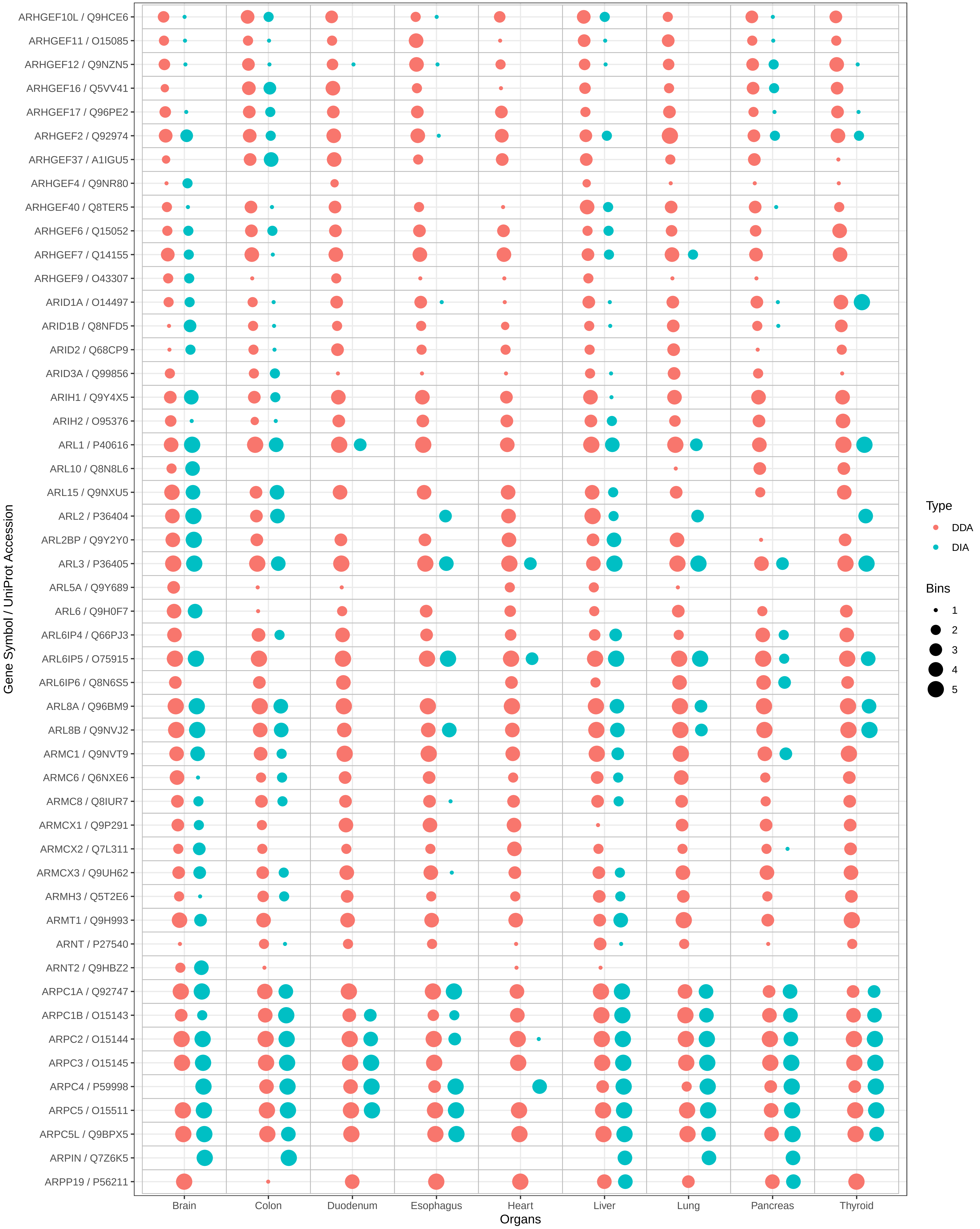

ARHGEF10L / Q9HCE6
ARHGEF11 / O15085
ARHGEF12 / Q9NZN5
ARHGEF16 / Q5VV41
ARHGEF17 / Q96PE2
ARHGEF2 / Q92974
ARHGEF37 / A1IGU5
ARHGEF4 / Q9NR80
ARHGEF40 / Q8TER5
ARHGEF6 / Q15052
ARHGEF7 / Q14155
ARHGEF9 / O43307
ARID1A / O14497
ARID1B / Q8NFD5
ARID2 / Q68CP9
ARID3A / Q99856
ARIH1 / Q9Y4X5
ARIH2 / O95376
ARL1 / P40616
ARL10 / Q8N8L6
ARL15 / Q9NXU5
ARL2 / P36404
ARL2BP / Q9Y2Y0
ARL3 / P36405
ARL5A / Q9Y689
ARL6 / Q9H0F7
ARL6IP4 / Q66PJ3
ARL6IP5 / O75915
ARL6IP6 / Q8N6S5
ARL8A / Q96BM9
ARL8B / Q9NVJ2
ARMC1 / Q9NVT9
ARMC6 / Q6NXE6
ARMC8 / Q8IUR7
ARMCX1 / Q9P291
ARMCX2 / Q7L311
ARMCX3 / Q9UH62
ARMH3 / Q5T2E6
ARMT1 / Q9H993
ARNT / P27540
ARNT2 / Q9HBZ2
ARPC1A / Q92747
ARPC1B / O15143
ARPC2 / O15144
ARPC3 / O15145
ARPC4 / P59998
ARPC5 / O15511
ARPC5L / Q9BPX5
ARPIN / Q7Z6K5
ARPP19 / P56211
Gene Symbol / UniProt Accession
Brain
Colon
Duodenum
Esophagus
Heart
Liver
Lung
Pancreas
Thyroid
Organs
Type
DDA
DIA
Bins
1
2
3
4
5

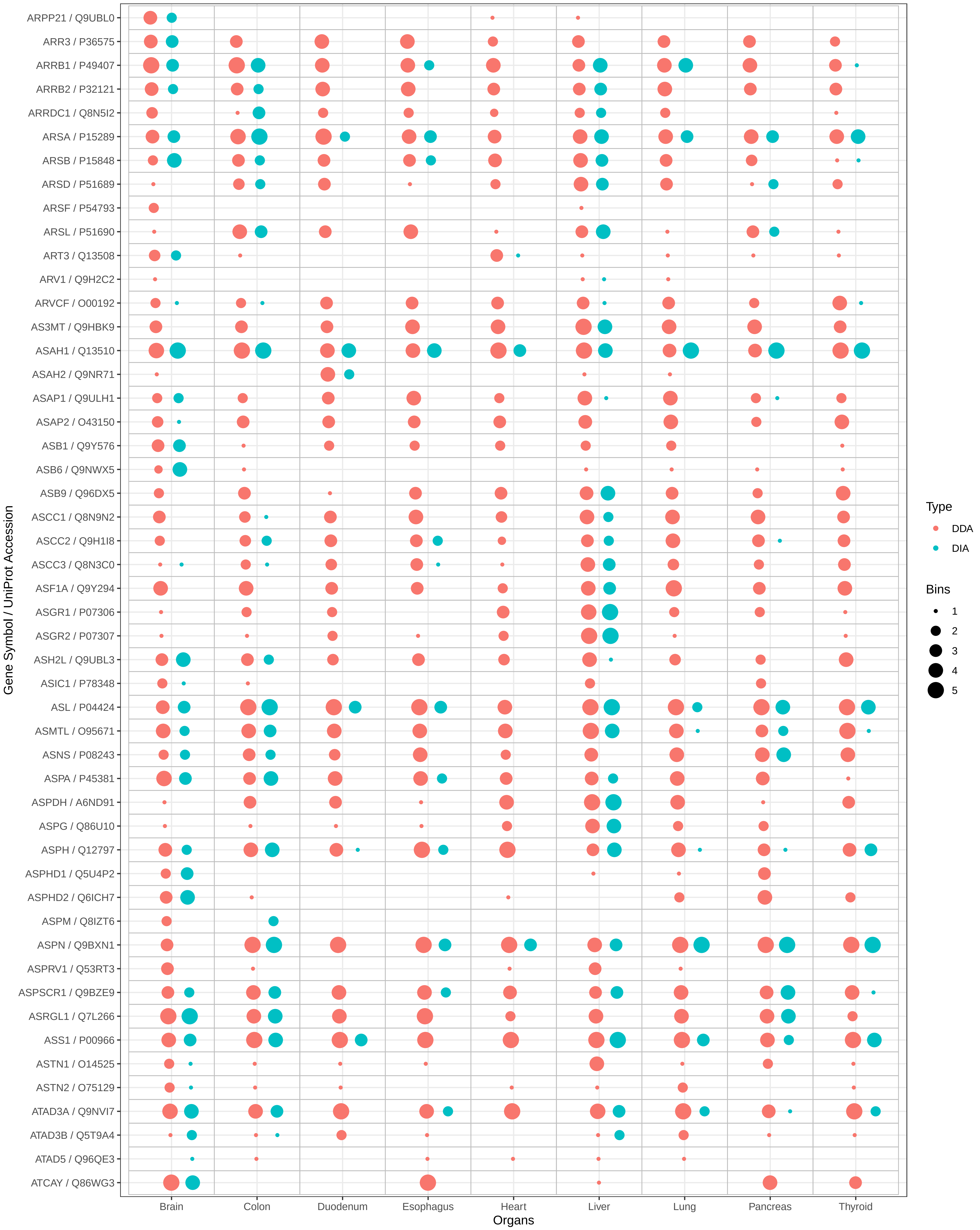

ARPP21 / Q9UBL0
ARR3 / P36575
ARRB1 / P49407
ARRB2 / P32121
ARRDC1 / Q8N5I2
ARSA / P15289
ARSB / P15848
ARSD / P51689
ARSF / P54793
ARSL / P51690
ART3 / Q13508
ARV1 / Q9H2C2
ARVCF / O00192
AS3MT / Q9HBK9
ASAH1 / Q13510
ASAH2 / Q9NR71
ASAP1 / Q9ULH1
ASAP2 / O43150
ASB1 / Q9Y576
ASB6 / Q9NWX5
ASB9 / Q96DX5
ASCC1 / Q8N9N2
ASCC2 / Q9H1I8
ASCC3 / Q8N3C0
ASF1A / Q9Y294
ASGR1 / P07306
ASGR2 / P07307
ASH2L / Q9UBL3
ASIC1 / P78348
ASL / P04424
ASMTL / O95671
ASNS / P08243
ASPA / P45381
ASPDH / A6ND91
ASPG / Q86U10
ASPH / Q12797
ASPHD1 / Q5U4P2
ASPHD2 / Q6ICH7
ASPM / Q8IZT6
ASPN / Q9BXN1
ASPRV1 / Q53RT3
ASPSCR1 / Q9BZE9
ASRGL1 / Q7L266
ASS1 / P00966
ASTN1 / O14525
ASTN2 / O75129
ATAD3A / Q9NVI7
ATAD3B / Q5T9A4
ATAD5 / Q96QE3
ATCAY / Q86WG3
Gene Symbol / UniProt Accession
Brain
Colon
Duodenum
Esophagus
Heart
Liver
Lung
Pancreas
Thyroid
Organs
Type
DDA
DIA
Bins
1
2
3
4
5

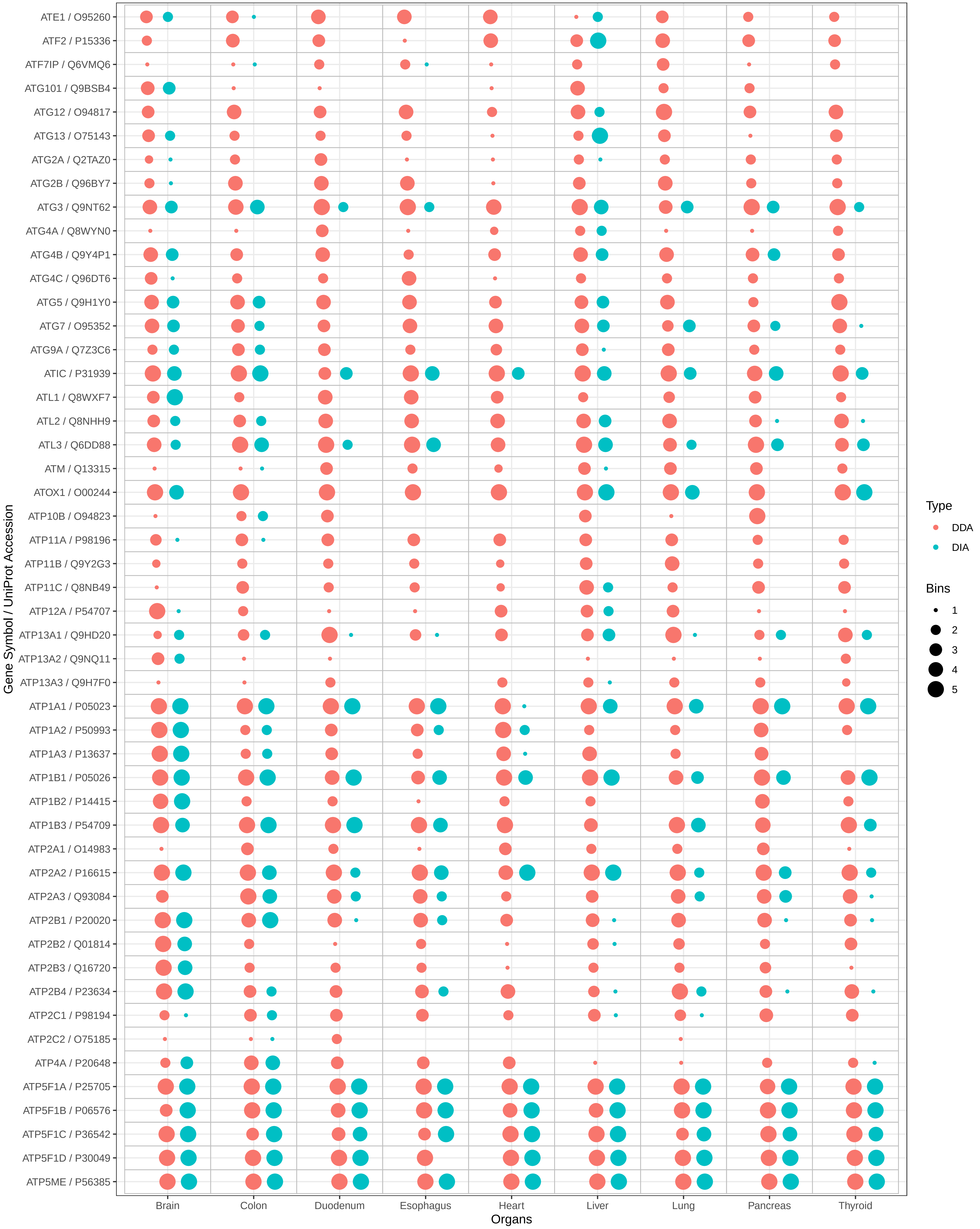

ATE1 / O95260
ATF2 / P15336
ATF7IP / Q6VMQ6
ATG101 / Q9BSB4
ATG12 / O94817
ATG13 / O75143
ATG2A / Q2TAZ0
ATG2B / Q96BY7
ATG3 / Q9NT62
ATG4A / Q8WYN0
ATG4B / Q9Y4P1
ATG4C / Q96DT6
ATG5 / Q9H1Y0
ATG7 / O95352
ATG9A / Q7Z3C6
ATIC / P31939
ATL1 / Q8WXF7
ATL2 / Q8NHH9
ATL3 / Q6DD88
ATM / Q13315
ATOX1 / O00244
ATP10B / O94823
ATP11A / P98196
ATP11B / Q9Y2G3
ATP11C / Q8NB49
ATP12A / P54707
ATP13A1 / Q9HD20
ATP13A2 / Q9NQ11
ATP13A3 / Q9H7F0
ATP1A1 / P05023
ATP1A2 / P50993
ATP1A3 / P13637
ATP1B1 / P05026
ATP1B2 / P14415
ATP1B3 / P54709
ATP2A1 / O14983
ATP2A2 / P16615
ATP2A3 / Q93084
ATP2B1 / P20020
ATP2B2 / Q01814
ATP2B3 / Q16720
ATP2B4 / P23634
ATP2C1 / P98194
ATP2C2 / O75185
ATP4A / P20648
ATP5F1A / P25705
ATP5F1B / P06576
ATP5F1C / P36542
ATP5F1D / P30049
ATP5ME / P56385
Gene Symbol / UniProt Accession
Brain
Colon
Duodenum
Esophagus
Heart
Liver
Lung
Pancreas
Thyroid
Organs
Type
DDA
DIA
Bins
1
2
3
4
5

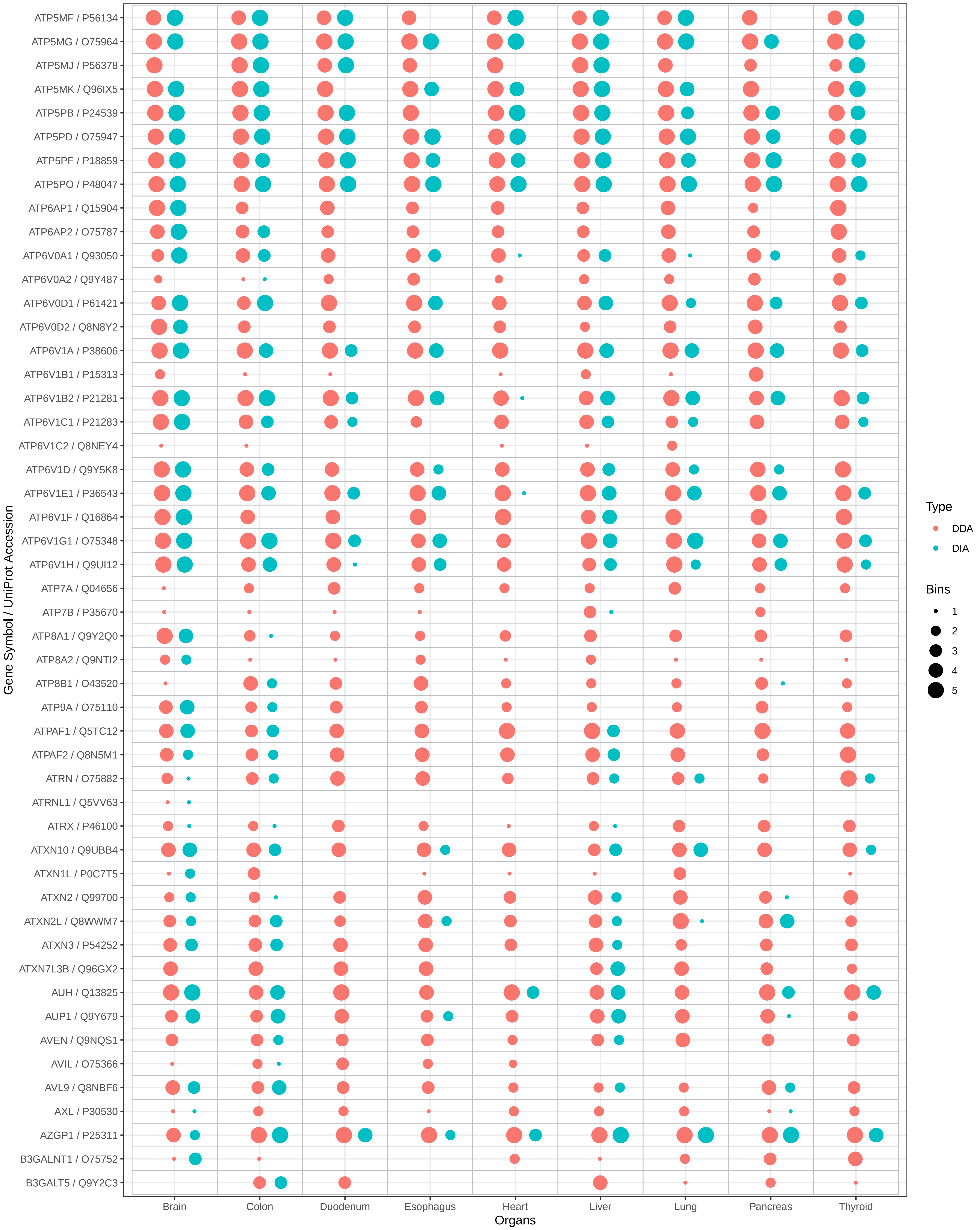
ATP5MF / P56134
ATP5MG / O75964
ATP5MJ / P56378
ATP5MK / Q96IX5
ATP5PB / P24539
ATP5PD / O75947
ATP5PF / P18859
ATP5PO / P48047
ATP6AP1 / Q15904
ATP6AP2 / O75787
ATP6V0A1 / Q93050
ATP6V0A2 / Q9Y487
ATP6V0D1 / P61421
ATP6V0D2 / Q8N8Y2
ATP6V1A / P38606
ATP6V1B1 / P15313
ATP6V1B2 / P21281
ATP6V1C1 / P21283
ATP6V1C2 / Q8NEY4
ATP6V1D / Q9Y5K8
ATP6V1E1 / P36543
ATP6V1F / Q16864
ATP6V1G1 / O75348
ATP6V1H / Q9UI12
ATP7A / Q04656
ATP7B / P35670
ATP8A1 / Q9Y2Q0
ATP8A2 / Q9NTI2
ATP8B1 / O43520
ATP9A / O75110
ATPAF1 / Q5TC12
ATPAF2 / Q8N5M1
ATRN / O75882
ATRNL1 / Q5VV63
ATRX / P46100
ATXN10 / Q9UBB4
ATXN1L / P0C7T5
ATXN2 / Q99700
ATXN2L / Q8WWM7
ATXN3 / P54252
ATXN7L3B / Q96GX2
AUH / Q13825
AUP1 / Q9Y679
AVEN / Q9NQS1
AVIL / O75366
AVL9 / Q8NBF6
AXL / P30530
AZGP1 / P25311
B3GALNT1 / O75752
B3GALT5 / Q9Y2C3
Gene Symbol / UniProt Accession
Brain
Colon
Duodenum
Esophagus
Heart
Liver
Lung
Pancreas
Thyroid
Organs
Type
DDA
DIA
Bins
1
2
3
4
5

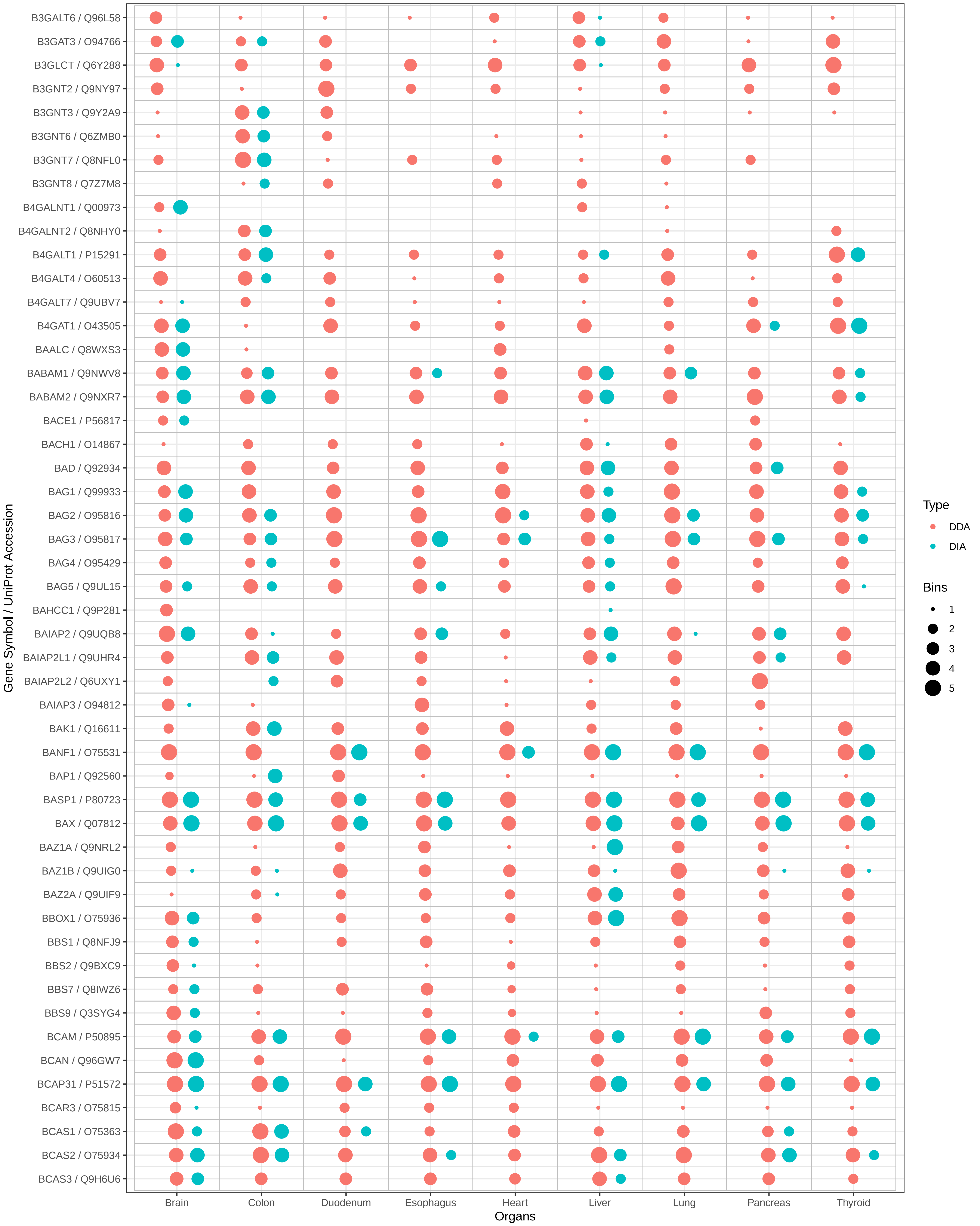
B3GALT6 / Q96L58
B3GAT3 / O94766
B3GLCT / Q6Y288
B3GNT2 / Q9NY97
B3GNT3 / Q9Y2A9
B3GNT6 / Q6ZMB0
B3GNT7 / Q8NFL0
B3GNT8 / Q7Z7M8
B4GALNT1 / Q00973
B4GALNT2 / Q8NHY0
B4GALT1 / P15291
B4GALT4 / O60513
B4GALT7 / Q9UBV7
B4GAT1 / O43505
BAALC / Q8WXS3
BABAM1 / Q9NWV8
BABAM2 / Q9NXR7
BACE1 / P56817
BACH1 / O14867
BAD / Q92934
BAG1 / Q99933
BAG2 / O95816
BAG3 / O95817
BAG4 / O95429
BAG5 / Q9UL15
BAHCC1 / Q9P281
BAIAP2 / Q9UQB8
BAIAP2L1 / Q9UHR4
BAIAP2L2 / Q6UXY1
BAIAP3 / O94812
BAK1 / Q16611
BANF1 / O75531
BAP1 / Q92560
BASP1 / P80723
BAX / Q07812
BAZ1A / Q9NRL2
BAZ1B / Q9UIG0
BAZ2A / Q9UIF9
BBOX1 / O75936
BBS1 / Q8NFJ9
BBS2 / Q9BXC9
BBS7 / Q8IWZ6
BBS9 / Q3SYG4
BCAM / P50895
BCAN / Q96GW7
BCAP31 / P51572
BCAR3 / O75815
BCAS1 / O75363
BCAS2 / O75934
BCAS3 / Q9H6U6
Gene Symbol / UniProt Accession
Brain
Colon
Duodenum
Esophagus
Heart
Liver
Lung
Pancreas
Thyroid
Organs
Type
DDA
DIA
Bins
1
2
3
4
5

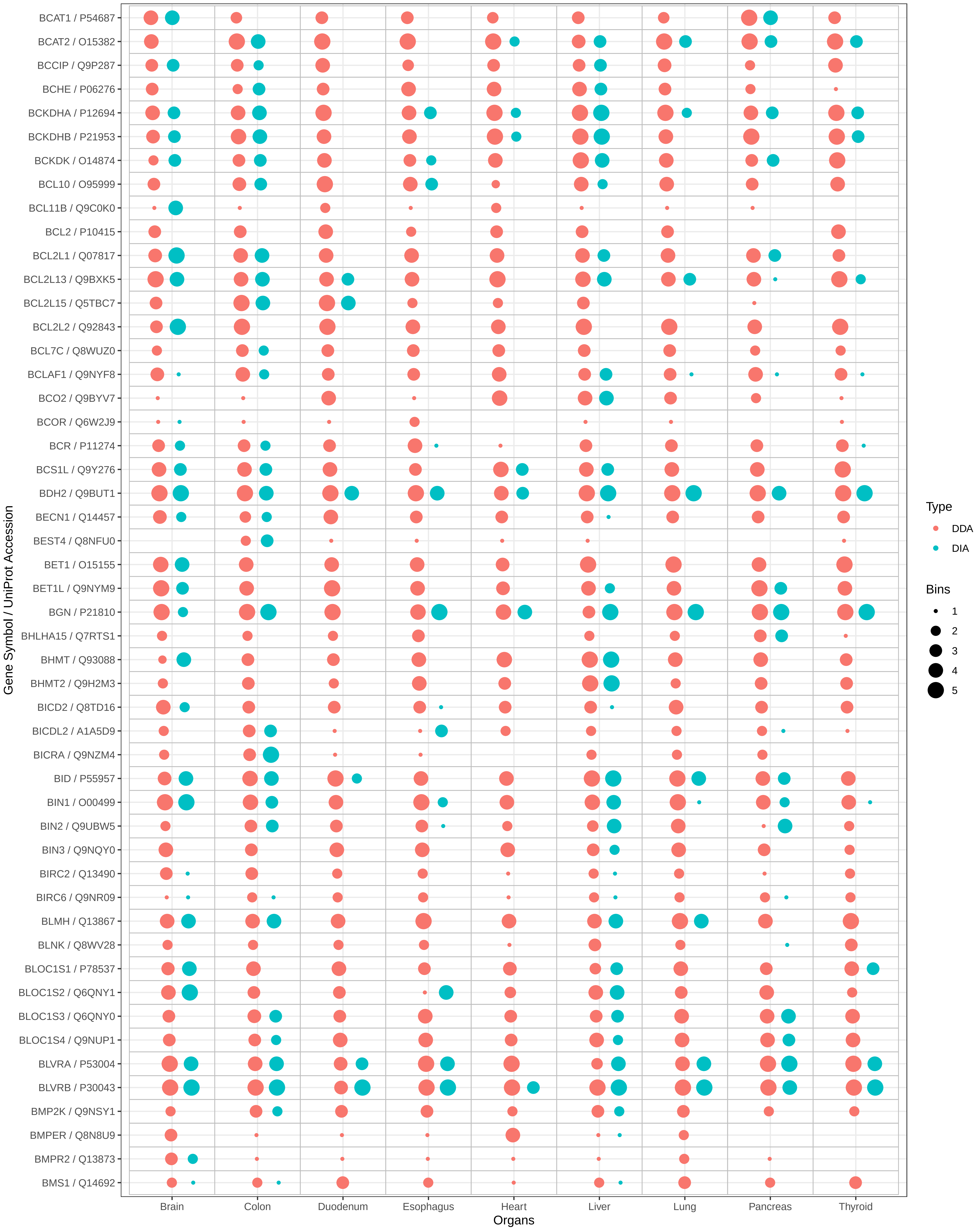
BCAT1 / P54687
BCAT2 / O15382
BCCIP / Q9P287
BCHE / P06276
BCKDHA / P12694
BCKDHB / P21953
BCKDK / O14874
BCL10 / O95999
BCL11B / Q9C0K0
BCL2 / P10415
BCL2L1 / Q07817
BCL2L13 / Q9BXK5
BCL2L15 / Q5TBC7
BCL2L2 / Q92843
BCL7C / Q8WUZ0
BCLAF1 / Q9NYF8
BCO2 / Q9BYV7
BCOR / Q6W2J9
BCR / P11274
BCS1L / Q9Y276
BDH2 / Q9BUT1
BECN1 / Q14457
BEST4 / Q8NFU0
BET1 / O15155
BET1L / Q9NYM9
BGN / P21810
BHLHA15 / Q7RTS1
BHMT / Q93088
BHMT2 / Q9H2M3
BICD2 / Q8TD16
BICDL2 / A1A5D9
BICRA / Q9NZM4
BID / P55957
BIN1 / O00499
BIN2 / Q9UBW5
BIN3 / Q9NQY0
BIRC2 / Q13490
BIRC6 / Q9NR09
BLMH / Q13867
BLNK / Q8WV28
BLOC1S1 / P78537
BLOC1S2 / Q6QNY1
BLOC1S3 / Q6QNY0
BLOC1S4 / Q9NUP1
BLVRA / P53004
BLVRB / P30043
BMP2K / Q9NSY1
BMPER / Q8N8U9
BMPR2 / Q13873
BMS1 / Q14692
Gene Symbol / UniProt Accession
Brain
Colon
Duodenum
Esophagus
Heart
Liver
Lung
Pancreas
Thyroid
Organs
Type
DDA
DIA
Bins
1
2
3
4
5

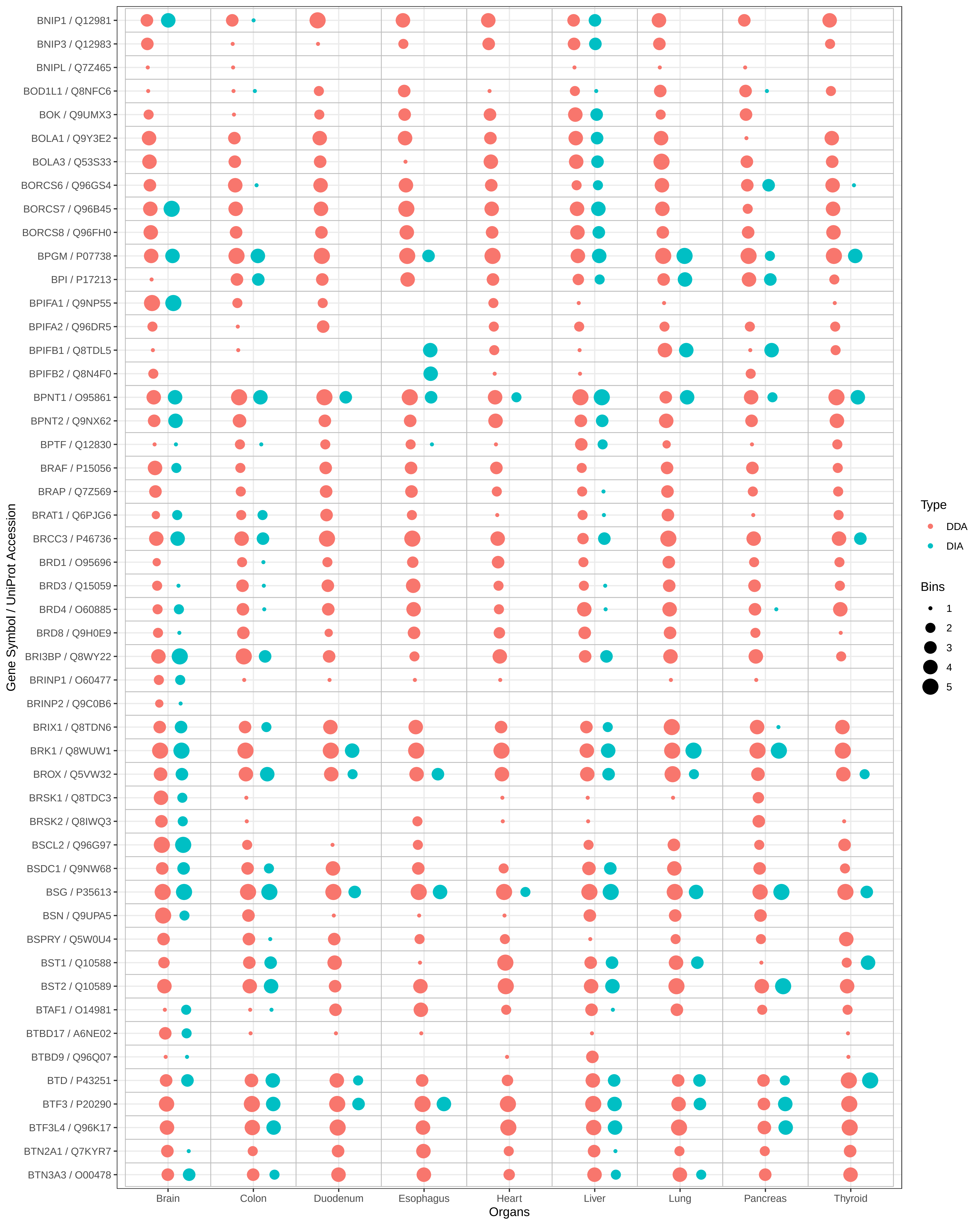

BNIP1 / Q12981
BNIP3 / Q12983
BNIPL / Q7Z465
BOD1L1 / Q8NFC6
BOK / Q9UMX3
BOLA1 / Q9Y3E2
BOLA3 / Q53S33
BORCS6 / Q96GS4
BORCS7 / Q96B45
BORCS8 / Q96FH0
BPGM / P07738
BPI / P17213
BPIFA1 / Q9NP55
BPIFA2 / Q96DR5
BPIFB1 / Q8TDL5
BPIFB2 / Q8N4F0
BPNT1 / O95861
BPNT2 / Q9NX62
BPTF / Q12830
BRAF / P15056
BRAP / Q7Z569
BRAT1 / Q6PJG6
BRCC3 / P46736
BRD1 / O95696
BRD3 / Q15059
BRD4 / O60885
BRD8 / Q9H0E9
BRI3BP / Q8WY22
BRINP1 / O60477
BRINP2 / Q9C0B6
BRIX1 / Q8TDN6
BRK1 / Q8WUW1
BROX / Q5VW32
BRSK1 / Q8TDC3
BRSK2 / Q8IWQ3
BSCL2 / Q96G97
BSDC1 / Q9NW68
BSG / P35613
BSN / Q9UPA5
BSPRY / Q5W0U4
BST1 / Q10588
BST2 / Q10589
BTAF1 / O14981
BTBD17 / A6NE02
BTBD9 / Q96Q07
BTD / P43251
BTF3 / P20290
BTF3L4 / Q96K17
BTN2A1 / Q7KYR7
BTN3A3 / O00478
Gene Symbol / UniProt Accession
Brain
Colon
Duodenum
Esophagus
Heart
Liver
Lung
Pancreas
Thyroid
Organs
Type
DDA
DIA
Bins
1
2
3
4
5

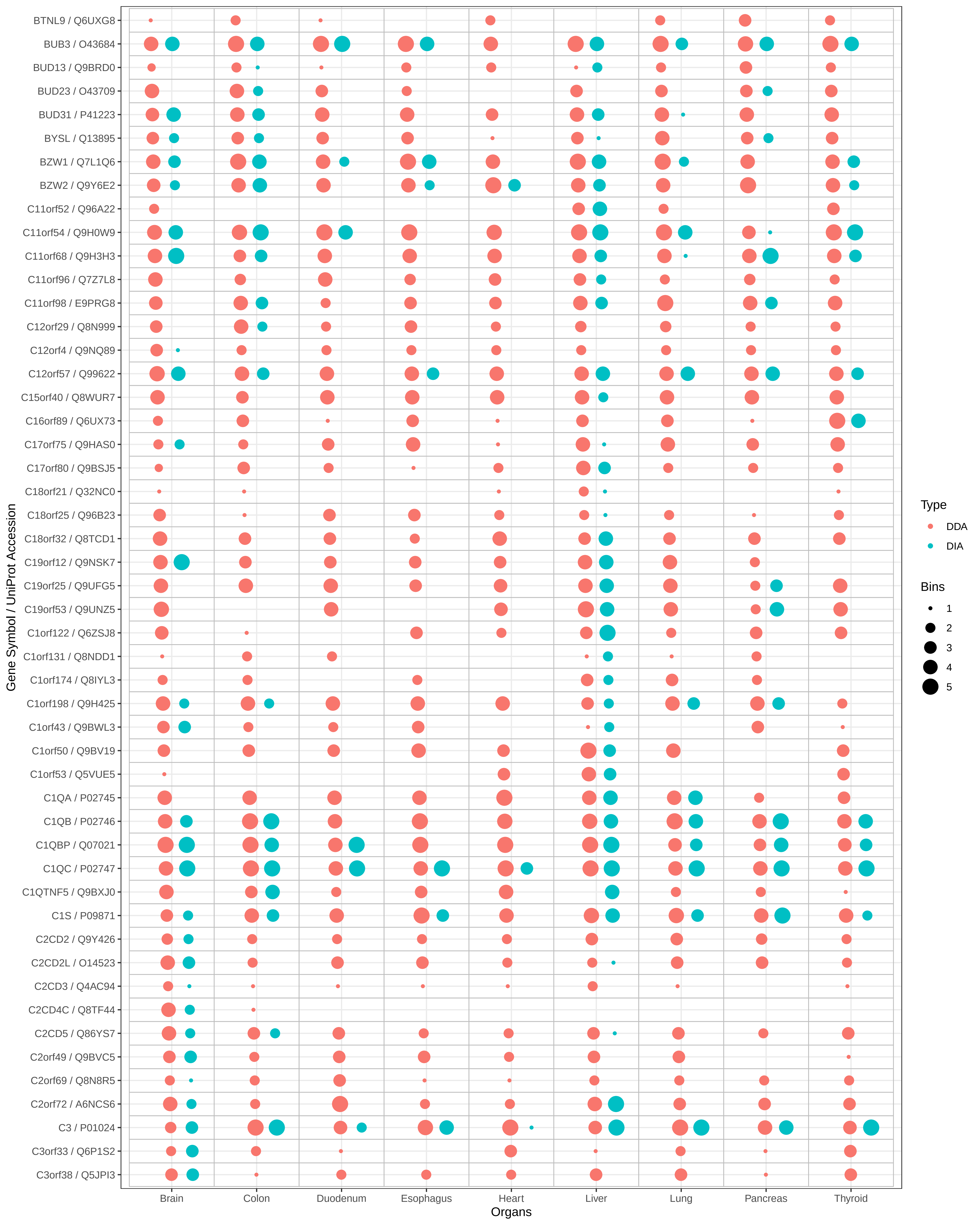

BTNL9 / Q6UXG8
BUB3 / O43684
BUD13 / Q9BRD0
BUD23 / O43709
BUD31 / P41223
BYSL / Q13895
BZW1 / Q7L1Q6
BZW2 / Q9Y6E2
C11orf52 / Q96A22
C11orf54 / Q9H0W9
C11orf68 / Q9H3H3
C11orf96 / Q7Z7L8
C11orf98 / E9PRG8
C12orf29 / Q8N999
C12orf4 / Q9NQ89
C12orf57 / Q99622
C15orf40 / Q8WUR7
C16orf89 / Q6UX73
C17orf75 / Q9HAS0
C17orf80 / Q9BSJ5
C18orf21 / Q32NC0
C18orf25 / Q96B23
C18orf32 / Q8TCD1
C19orf12 / Q9NSK7
C19orf25 / Q9UFG5
C19orf53 / Q9UNZ5
C1orf122 / Q6ZSJ8
C1orf131 / Q8NDD1
C1orf174 / Q8IYL3
C1orf198 / Q9H425
C1orf43 / Q9BWL3
C1orf50 / Q9BV19
C1orf53 / Q5VUE5
C1QA / P02745
C1QB / P02746
C1QBP / Q07021
C1QC / P02747
C1QTNF5 / Q9BXJ0
C1S / P09871
C2CD2 / Q9Y426
C2CD2L / O14523
C2CD3 / Q4AC94
C2CD4C / Q8TF44
C2CD5 / Q86YS7
C2orf49 / Q9BVC5
C2orf69 / Q8N8R5
C2orf72 / A6NCS6
C3 / P01024
C3orf33 / Q6P1S2
C3orf38 / Q5JPI3
Gene Symbol / UniProt Accession
Brain
Colon
Duodenum
Esophagus
Heart
Liver
Lung
Pancreas
Thyroid
Organs
Type
DDA
DIA
Bins
1
2
3
4
5

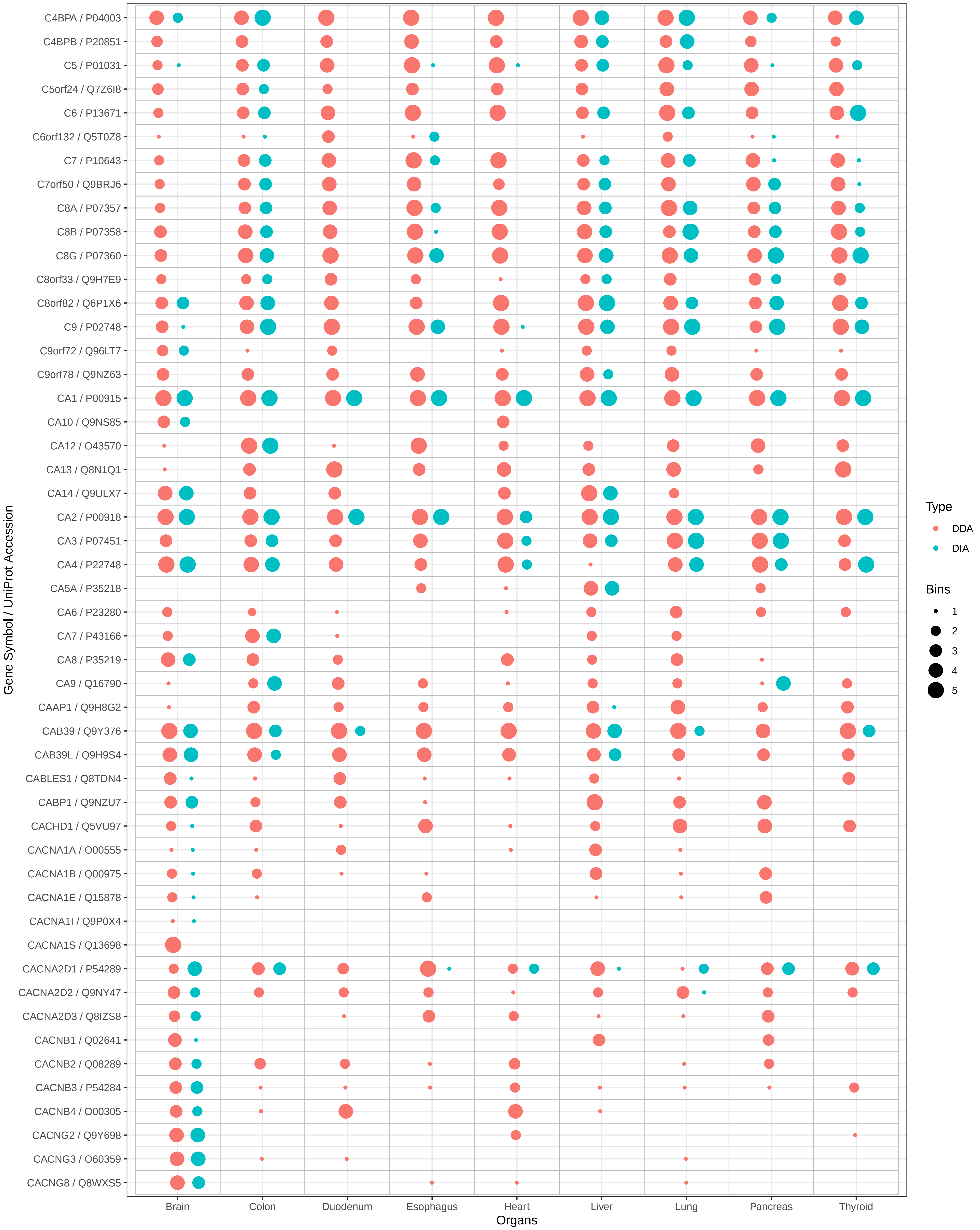

C4BPA / P04003
C4BPB / P20851
C5 / P01031
C5orf24 / Q7Z6I8
C6 / P13671
C6orf132 / Q5T0Z8
C7 / P10643
C7orf50 / Q9BRJ6
C8A / P07357
C8B / P07358
C8G / P07360
C8orf33 / Q9H7E9
C8orf82 / Q6P1X6
C9 / P02748
C9orf72 / Q96LT7
C9orf78 / Q9NZ63
CA1 / P00915
CA10 / Q9NS85
CA12 / O43570
CA13 / Q8N1Q1
CA14 / Q9ULX7
CA2 / P00918
CA3 / P07451
CA4 / P22748
CA5A / P35218
CA6 / P23280
CA7 / P43166
CA8 / P35219
CA9 / Q16790
CAAP1 / Q9H8G2
CAB39 / Q9Y376
CAB39L / Q9H9S4
CABLES1 / Q8TDN4
CABP1 / Q9NZU7
CACHD1 / Q5VU97
CACNA1A / O00555
CACNA1B / Q00975
CACNA1E / Q15878
CACNA1I / Q9P0X4
CACNA1S / Q13698
CACNA2D1 / P54289
CACNA2D2 / Q9NY47
CACNA2D3 / Q8IZS8
CACNB1 / Q02641
CACNB2 / Q08289
CACNB3 / P54284
CACNB4 / O00305
CACNG2 / Q9Y698
CACNG3 / O60359
CACNG8 / Q8WXS5
Gene Symbol / UniProt Accession
Brain
Colon
Duodenum
Esophagus
Heart
Liver
Lung
Pancreas
Thyroid
Organs
Type
DDA
DIA
Bins
1
2
3
4
5

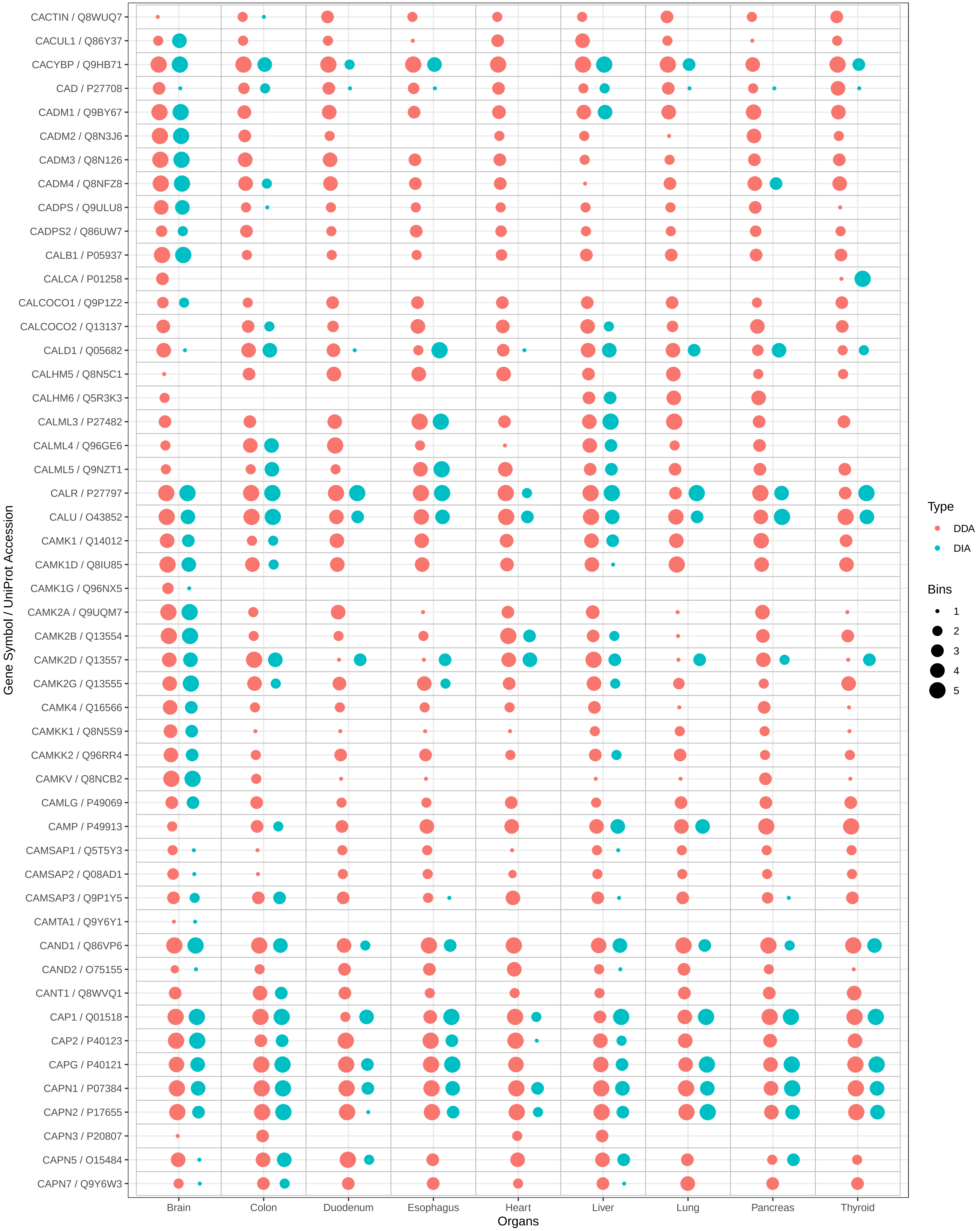

CACTIN / Q8WUQ7
CACUL1 / Q86Y37
CACYBP / Q9HB71
CAD / P27708
CADM1 / Q9BY67
CADM2 / Q8N3J6
CADM3 / Q8N126
CADM4 / Q8NFZ8
CADPS / Q9ULU8
CADPS2 / Q86UW7
CALB1 / P05937
CALCA / P01258
CALCOCO1 / Q9P1Z2
CALCOCO2 / Q13137
CALD1 / Q05682
CALHM5 / Q8N5C1
CALHM6 / Q5R3K3
CALML3 / P27482
CALML4 / Q96GE6
CALML5 / Q9NZT1
CALR / P27797
CALU / O43852
CAMK1 / Q14012
CAMK1D / Q8IU85
CAMK1G / Q96NX5
CAMK2A / Q9UQM7
CAMK2B / Q13554
CAMK2D / Q13557
CAMK2G / Q13555
CAMK4 / Q16566
CAMKK1 / Q8N5S9
CAMKK2 / Q96RR4
CAMKV / Q8NCB2
CAMLG / P49069
CAMP / P49913
CAMSAP1 / Q5T5Y3
CAMSAP2 / Q08AD1
CAMSAP3 / Q9P1Y5
CAMTA1 / Q9Y6Y1
CAND1 / Q86VP6
CAND2 / O75155
CANT1 / Q8WVQ1
CAP1 / Q01518
CAP2 / P40123
CAPG / P40121
CAPN1 / P07384
CAPN2 / P17655
CAPN3 / P20807
CAPN5 / O15484
CAPN7 / Q9Y6W3
Gene Symbol / UniProt Accession
Brain
Colon
Duodenum
Esophagus
Heart
Liver
Lung
Pancreas
Thyroid
Organs
Type
DDA
DIA
Bins
1
2
3
4
5

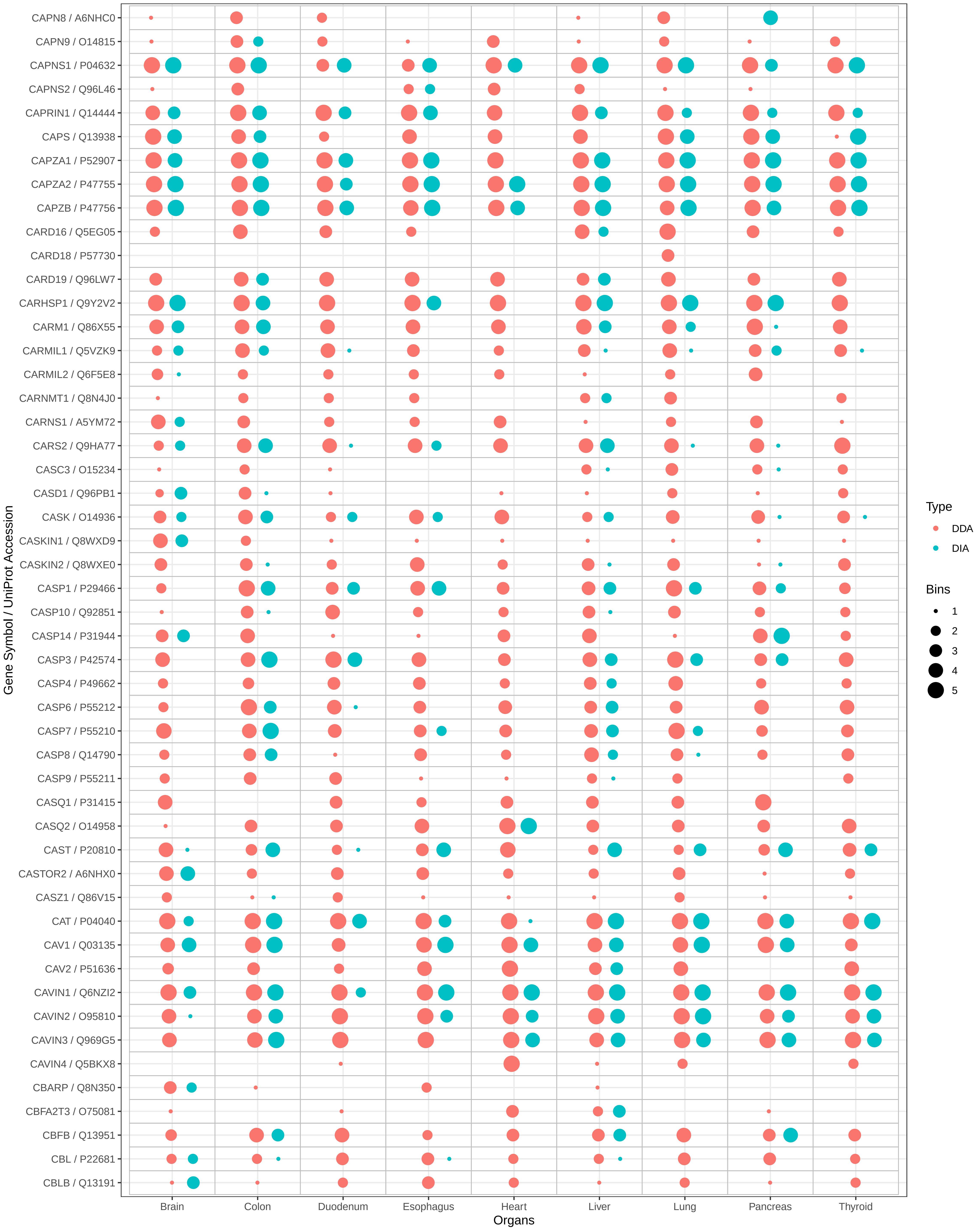
CAPN8 / A6NHC0
CAPN9 / O14815
CAPNS1 / P04632
CAPNS2 / Q96L46
CAPRIN1 / Q14444
CAPS / Q13938
CAPZA1 / P52907
CAPZA2 / P47755
CAPZB / P47756
CARD16 / Q5EG05
CARD18 / P57730
CARD19 / Q96LW7
CARHSP1 / Q9Y2V2
CARM1 / Q86X55
CARMIL1 / Q5VZK9
CARMIL2 / Q6F5E8
CARNMT1 / Q8N4J0
CARNS1 / A5YM72
CARS2 / Q9HA77
CASC3 / O15234
CASD1 / Q96PB1
CASK / O14936
CASKIN1 / Q8WXD9
CASKIN2 / Q8WXE0
CASP1 / P29466
CASP10 / Q92851
CASP14 / P31944
CASP3 / P42574
CASP4 / P49662
CASP6 / P55212
CASP7 / P55210
CASP8 / Q14790
CASP9 / P55211
CASQ1 / P31415
CASQ2 / O14958
CAST / P20810
CASTOR2 / A6NHX0
CASZ1 / Q86V15
CAT / P04040
CAV1 / Q03135
CAV2 / P51636
CAVIN1 / Q6NZI2
CAVIN2 / O95810
CAVIN3 / Q969G5
CAVIN4 / Q5BKX8
CBARP / Q8N350
CBFA2T3 / O75081
CBFB / Q13951
CBL / P22681
CBLB / Q13191
Gene Symbol / UniProt Accession
Brain
Colon
Duodenum
Esophagus
Heart
Liver
Lung
Pancreas
Thyroid
Organs
Type
DDA
DIA
Bins
1
2
3
4
5

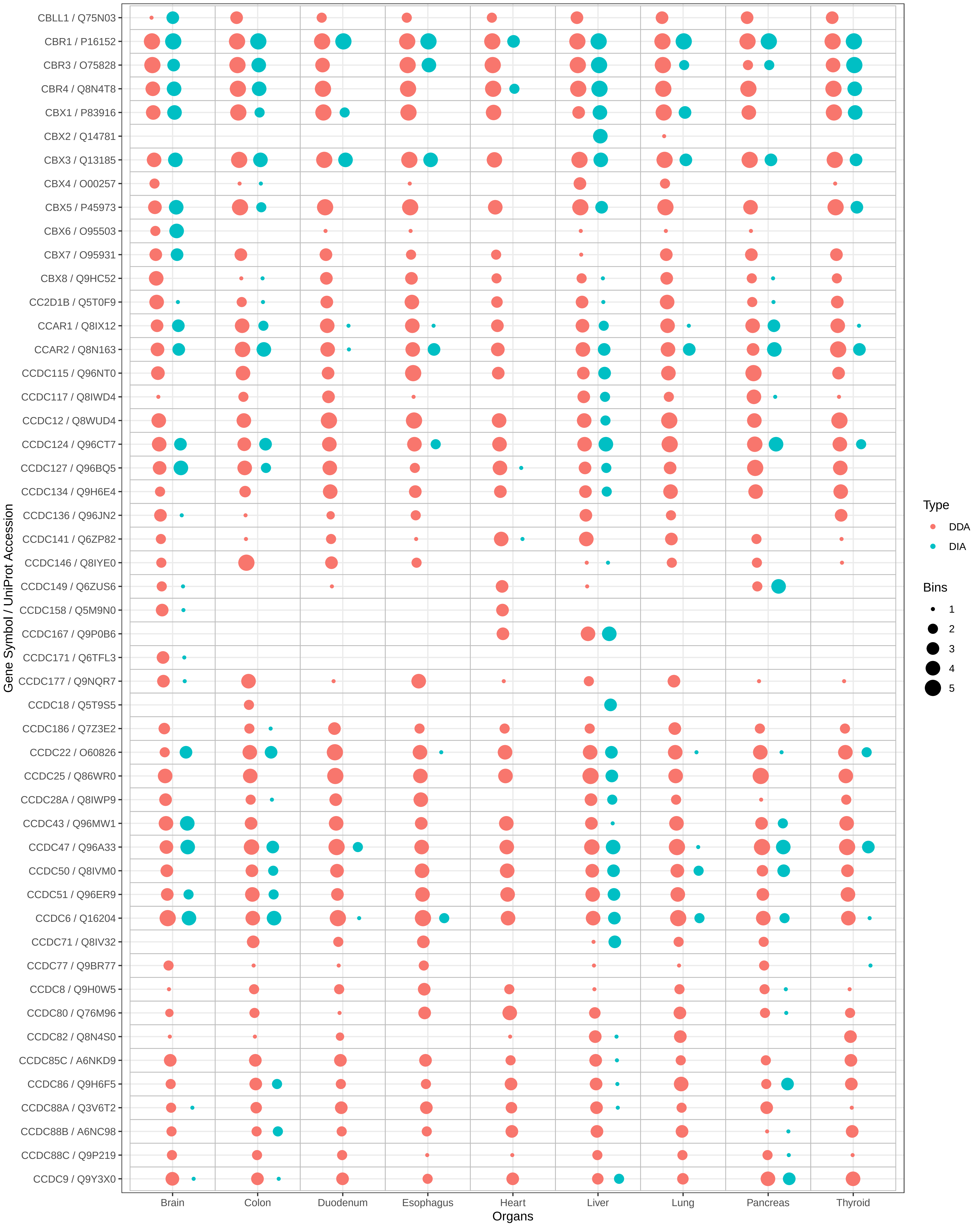

CBLL1 / Q75N03
CBR1 / P16152
CBR3 / O75828
CBR4 / Q8N4T8
CBX1 / P83916
CBX2 / Q14781
CBX3 / Q13185
CBX4 / O00257
CBX5 / P45973
CBX6 / O95503
CBX7 / O95931
CBX8 / Q9HC52
CC2D1B / Q5T0F9
CCAR1 / Q8IX12
CCAR2 / Q8N163
CCDC115 / Q96NT0
CCDC117 / Q8IWD4
CCDC12 / Q8WUD4
CCDC124 / Q96CT7
CCDC127 / Q96BQ5
CCDC134 / Q9H6E4
CCDC136 / Q96JN2
CCDC141 / Q6ZP82
CCDC146 / Q8IYE0
CCDC149 / Q6ZUS6
CCDC158 / Q5M9N0
CCDC167 / Q9P0B6
CCDC171 / Q6TFL3
CCDC177 / Q9NQR7
CCDC18 / Q5T9S5
CCDC186 / Q7Z3E2
CCDC22 / O60826
CCDC25 / Q86WR0
CCDC28A / Q8IWP9
CCDC43 / Q96MW1
CCDC47 / Q96A33
CCDC50 / Q8IVM0
CCDC51 / Q96ER9
CCDC6 / Q16204
CCDC71 / Q8IV32
CCDC77 / Q9BR77
CCDC8 / Q9H0W5
CCDC80 / Q76M96
CCDC82 / Q8N4S0
CCDC85C / A6NKD9
CCDC86 / Q9H6F5
CCDC88A / Q3V6T2
CCDC88B / A6NC98
CCDC88C / Q9P219
CCDC9 / Q9Y3X0
Gene Symbol / UniProt Accession
Brain
Colon
Duodenum
Esophagus
Heart
Liver
Lung
Pancreas
Thyroid
Organs
Type
DDA
DIA
Bins
1
2
3
4
5

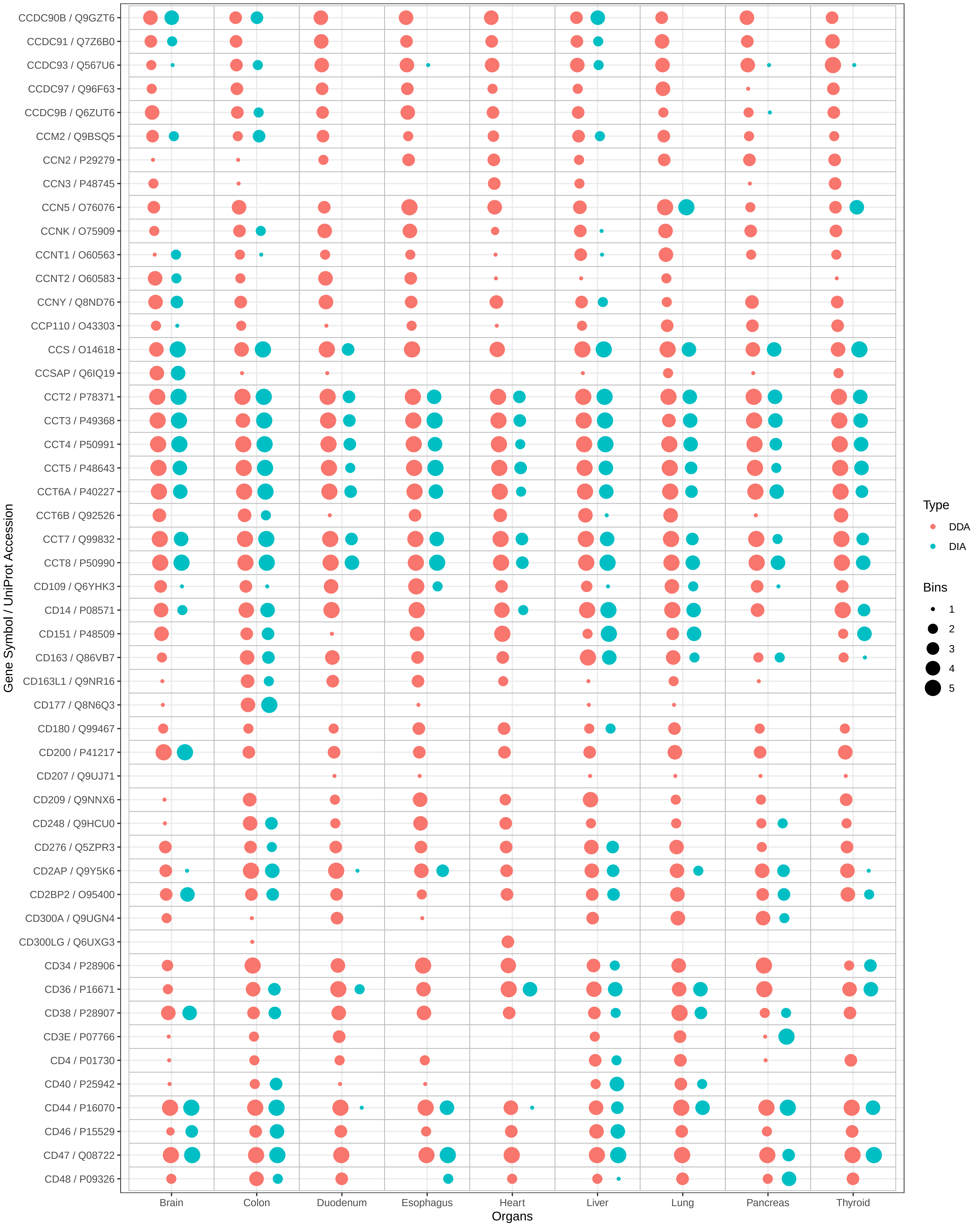
CCDC90B / Q9GZT6
CCDC91 / Q7Z6B0
CCDC93 / Q567U6
CCDC97 / Q96F63
CCDC9B / Q6ZUT6
CCM2 / Q9BSQ5
CCN2 / P29279
CCN3 / P48745
CCN5 / O76076
CCNK / O75909
CCNT1 / O60563
CCNT2 / O60583
CCNY / Q8ND76
CCP110 / O43303
CCS / O14618
CCSAP / Q6IQ19
CCT2 / P78371
CCT3 / P49368
CCT4 / P50991
CCT5 / P48643
CCT6A / P40227
CCT6B / Q92526
CCT7 / Q99832
CCT8 / P50990
CD109 / Q6YHK3
CD14 / P08571
CD151 / P48509
CD163 / Q86VB7
CD163L1 / Q9NR16
CD177 / Q8N6Q3
CD180 / Q99467
CD200 / P41217
CD207 / Q9UJ71
CD209 / Q9NNX6
CD248 / Q9HCU0
CD276 / Q5ZPR3
CD2AP / Q9Y5K6
CD2BP2 / O95400
CD300A / Q9UGN4
CD300LG / Q6UXG3
CD34 / P28906
CD36 / P16671
CD38 / P28907
CD3E / P07766
CD4 / P01730
CD40 / P25942
CD44 / P16070
CD46 / P15529
CD47 / Q08722
CD48 / P09326
Gene Symbol / UniProt Accession
Brain
Colon
Duodenum
Esophagus
Heart
Liver
Lung
Pancreas
Thyroid
Organs
Type
DDA
DIA
Bins
1
2
3
4
5

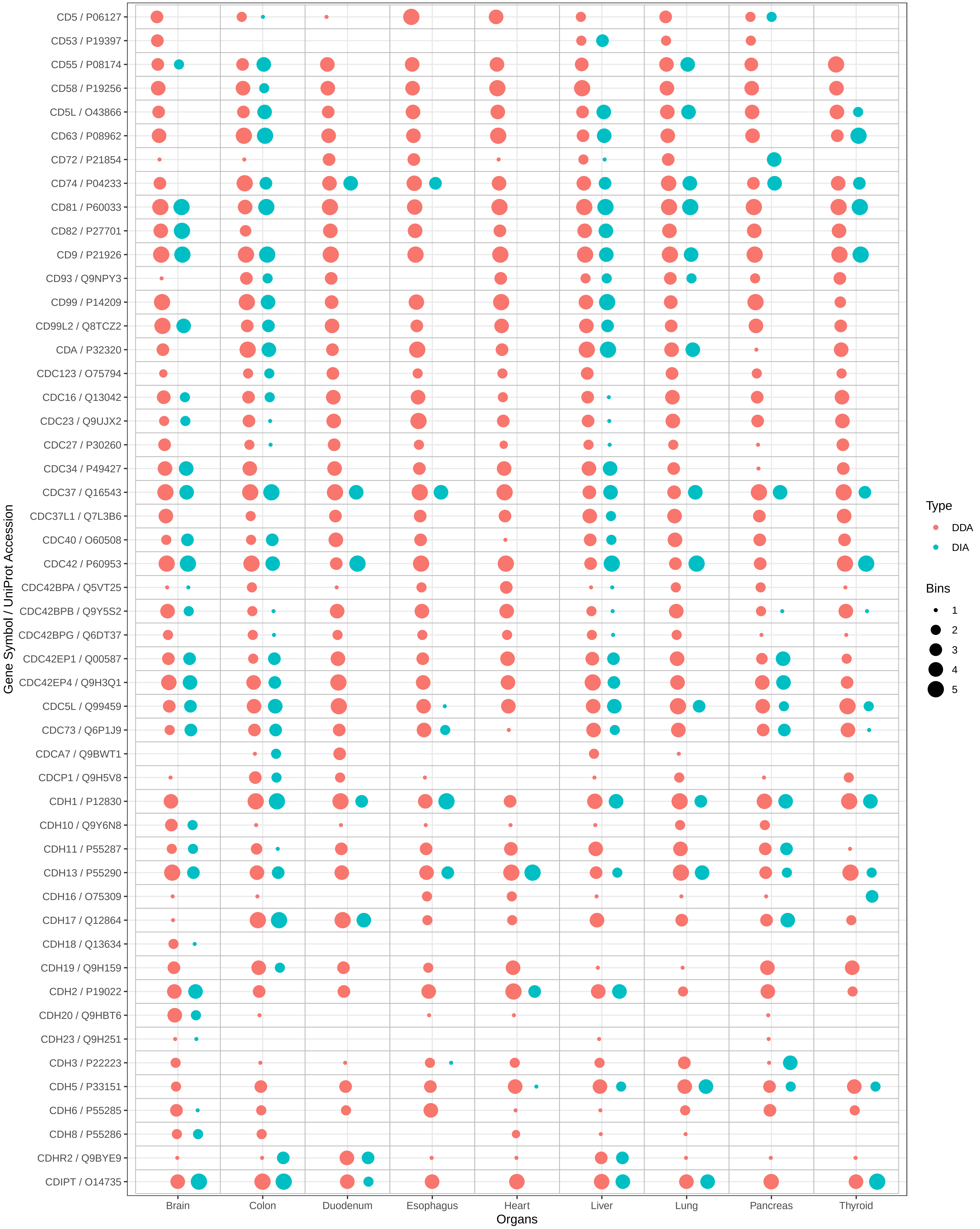
CD5 / P06127
CD53 / P19397
CD55 / P08174
CD58 / P19256
CD5L / O43866
CD63 / P08962
CD72 / P21854
CD74 / P04233
CD81 / P60033
CD82 / P27701
CD9 / P21926
CD93 / Q9NPY3
CD99 / P14209
CD99L2 / Q8TCZ2
CDA / P32320
CDC123 / O75794
CDC16 / Q13042
CDC23 / Q9UJX2
CDC27 / P30260
CDC34 / P49427
CDC37 / Q16543
CDC37L1 / Q7L3B6
CDC40 / O60508
CDC42 / P60953
CDC42BPA / Q5VT25
CDC42BPB / Q9Y5S2
CDC42BPG / Q6DT37
CDC42EP1 / Q00587
CDC42EP4 / Q9H3Q1
CDC5L / Q99459
CDC73 / Q6P1J9
CDCA7 / Q9BWT1
CDCP1 / Q9H5V8
CDH1 / P12830
CDH10 / Q9Y6N8
CDH11 / P55287
CDH13 / P55290
CDH16 / O75309
CDH17 / Q12864
CDH18 / Q13634
CDH19 / Q9H159
CDH2 / P19022
CDH20 / Q9HBT6
CDH23 / Q9H251
CDH3 / P22223
CDH5 / P33151
CDH6 / P55285
CDH8 / P55286
CDHR2 / Q9BYE9
CDIPT / O14735
Gene Symbol / UniProt Accession
Brain
Colon
Duodenum
Esophagus
Heart
Liver
Lung
Pancreas
Thyroid
Organs
Type
DDA
DIA
Bins
1
2
3
4
5

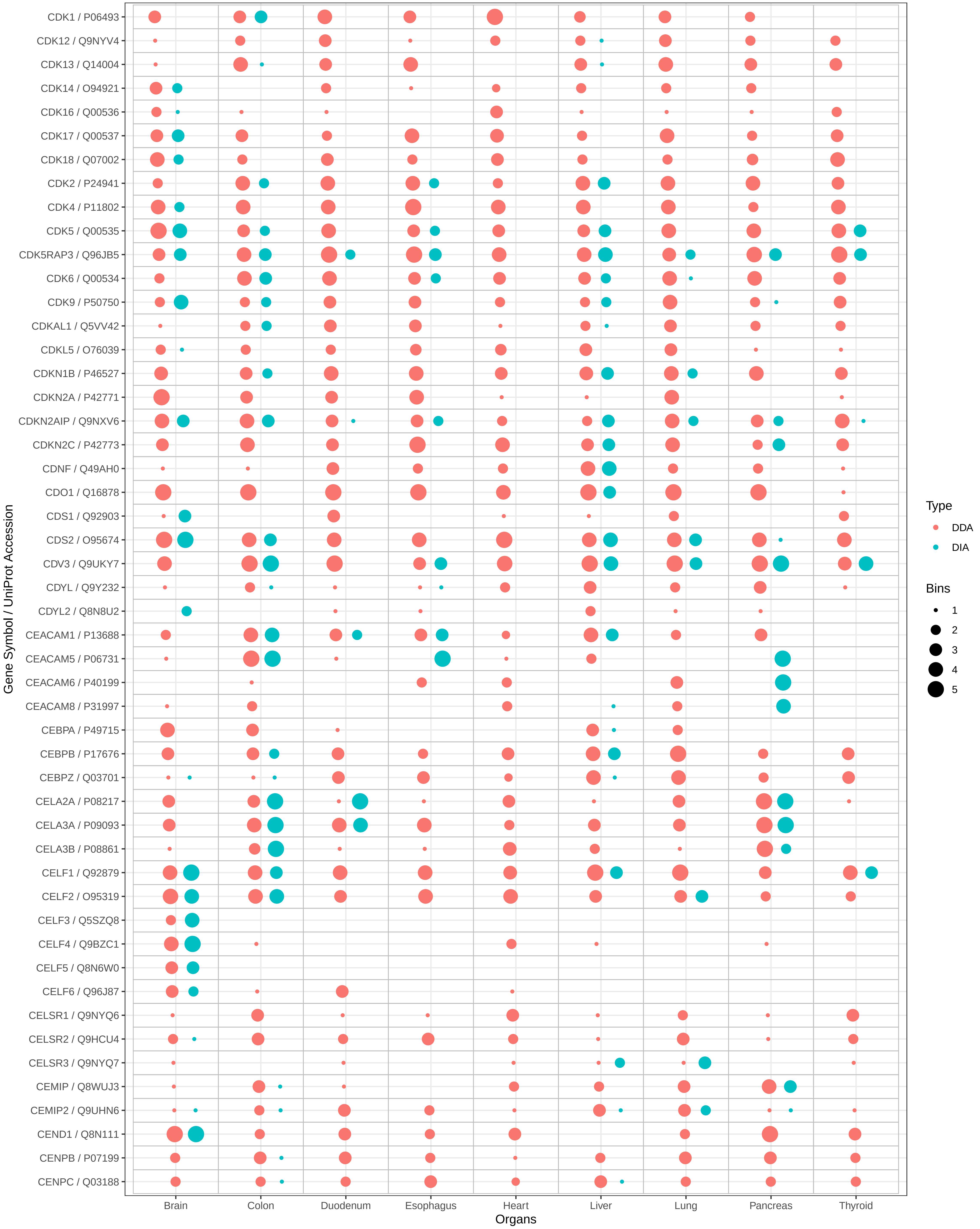

CDK1 / P06493
CDK12 / Q9NYV4
CDK13 / Q14004
CDK14 / O94921
CDK16 / Q00536
CDK17 / Q00537
CDK18 / Q07002
CDK2 / P24941
CDK4 / P11802
CDK5 / Q00535
CDK5RAP3 / Q96JB5
CDK6 / Q00534
CDK9 / P50750
CDKAL1 / Q5VV42
CDKL5 / O76039
CDKN1B / P46527
CDKN2A / P42771
CDKN2AIP / Q9NXV6
CDKN2C / P42773
CDNF / Q49AH0
CDO1 / Q16878
CDS1 / Q92903
CDS2 / O95674
CDV3 / Q9UKY7
CDYL / Q9Y232
CDYL2 / Q8N8U2
CEACAM1 / P13688
CEACAM5 / P06731
CEACAM6 / P40199
CEACAM8 / P31997
CEBPA / P49715
CEBPB / P17676
CEBPZ / Q03701
CELA2A / P08217
CELA3A / P09093
CELA3B / P08861
CELF1 / Q92879
CELF2 / O95319
CELF3 / Q5SZQ8
CELF4 / Q9BZC1
CELF5 / Q8N6W0
CELF6 / Q96J87
CELSR1 / Q9NYQ6
CELSR2 / Q9HCU4
CELSR3 / Q9NYQ7
CEMIP / Q8WUJ3
CEMIP2 / Q9UHN6
CEND1 / Q8N111
CENPB / P07199
CENPC / Q03188
Gene Symbol / UniProt Accession
Brain
Colon
Duodenum
Esophagus
Heart
Liver
Lung
Pancreas
Thyroid
Organs
Type
DDA
DIA
Bins
1
2
3
4
5

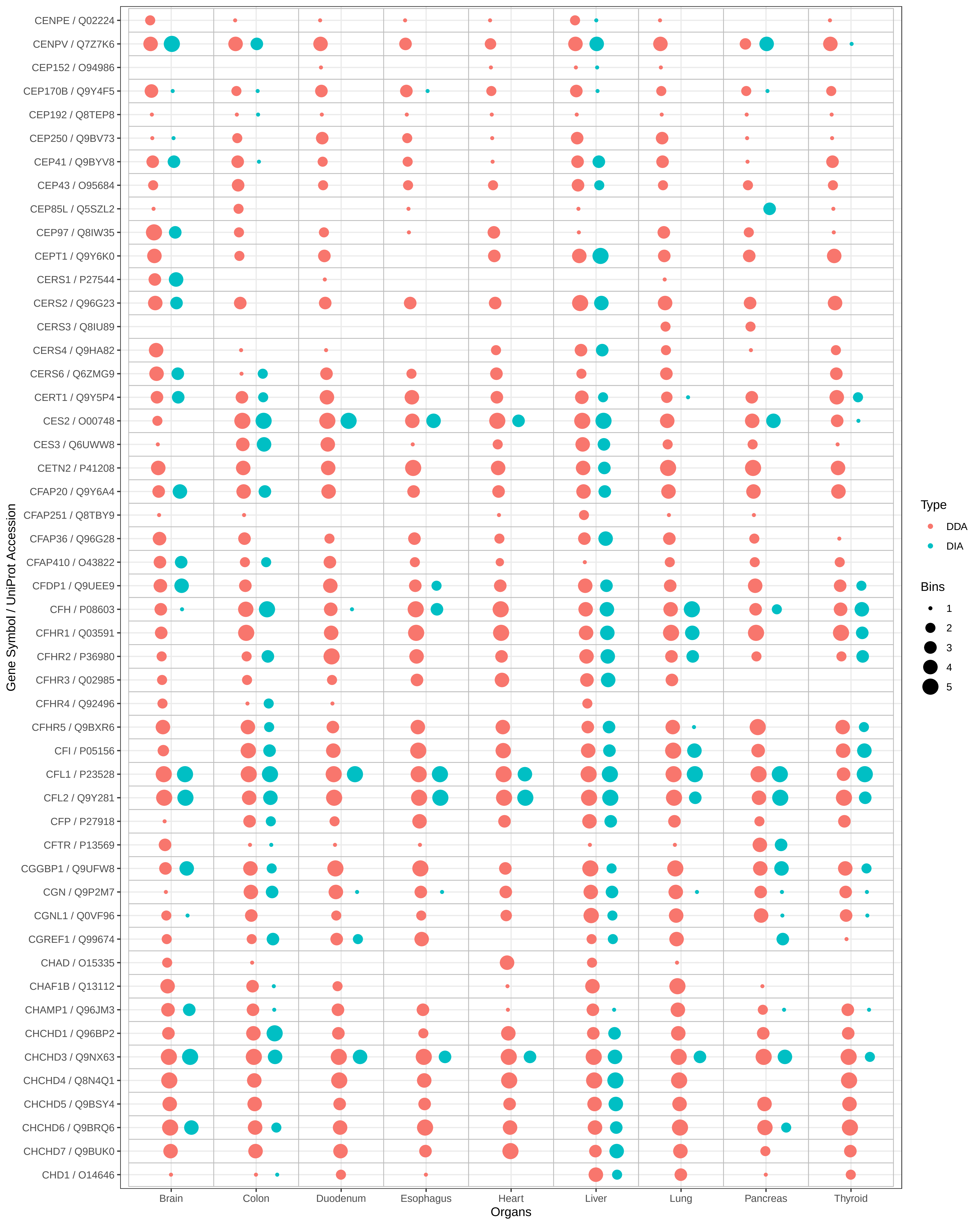
CENPE / Q02224
CENPV / Q7Z7K6
CEP152 / O94986
CEP170B / Q9Y4F5
CEP192 / Q8TEP8
CEP250 / Q9BV73
CEP41 / Q9BYV8
CEP43 / O95684
CEP85L / Q5SZL2
CEP97 / Q8IW35
CEPT1 / Q9Y6K0
CERS1 / P27544
CERS2 / Q96G23
CERS3 / Q8IU89
CERS4 / Q9HA82
CERS6 / Q6ZMG9
CERT1 / Q9Y5P4
CES2 / O00748
CES3 / Q6UWW8
CETN2 / P41208
CFAP20 / Q9Y6A4
CFAP251 / Q8TBY9
CFAP36 / Q96G28
CFAP410 / O43822
CFDP1 / Q9UEE9
CFH / P08603
CFHR1 / Q03591
CFHR2 / P36980
CFHR3 / Q02985
CFHR4 / Q92496
CFHR5 / Q9BXR6
CFI / P05156
CFL1 / P23528
CFL2 / Q9Y281
CFP / P27918
CFTR / P13569
CGGBP1 / Q9UFW8
CGN / Q9P2M7
CGNL1 / Q0VF96
CGREF1 / Q99674
CHAD / O15335
CHAF1B / Q13112
CHAMP1 / Q96JM3
CHCHD1 / Q96BP2
CHCHD3 / Q9NX63
CHCHD4 / Q8N4Q1
CHCHD5 / Q9BSY4
CHCHD6 / Q9BRQ6
CHCHD7 / Q9BUK0
CHD1 / O14646
Gene Symbol / UniProt Accession
Brain
Colon
Duodenum
Esophagus
Heart
Liver
Lung
Pancreas
Thyroid
Organs
Type
DDA
DIA
Bins
1
2
3
4
5

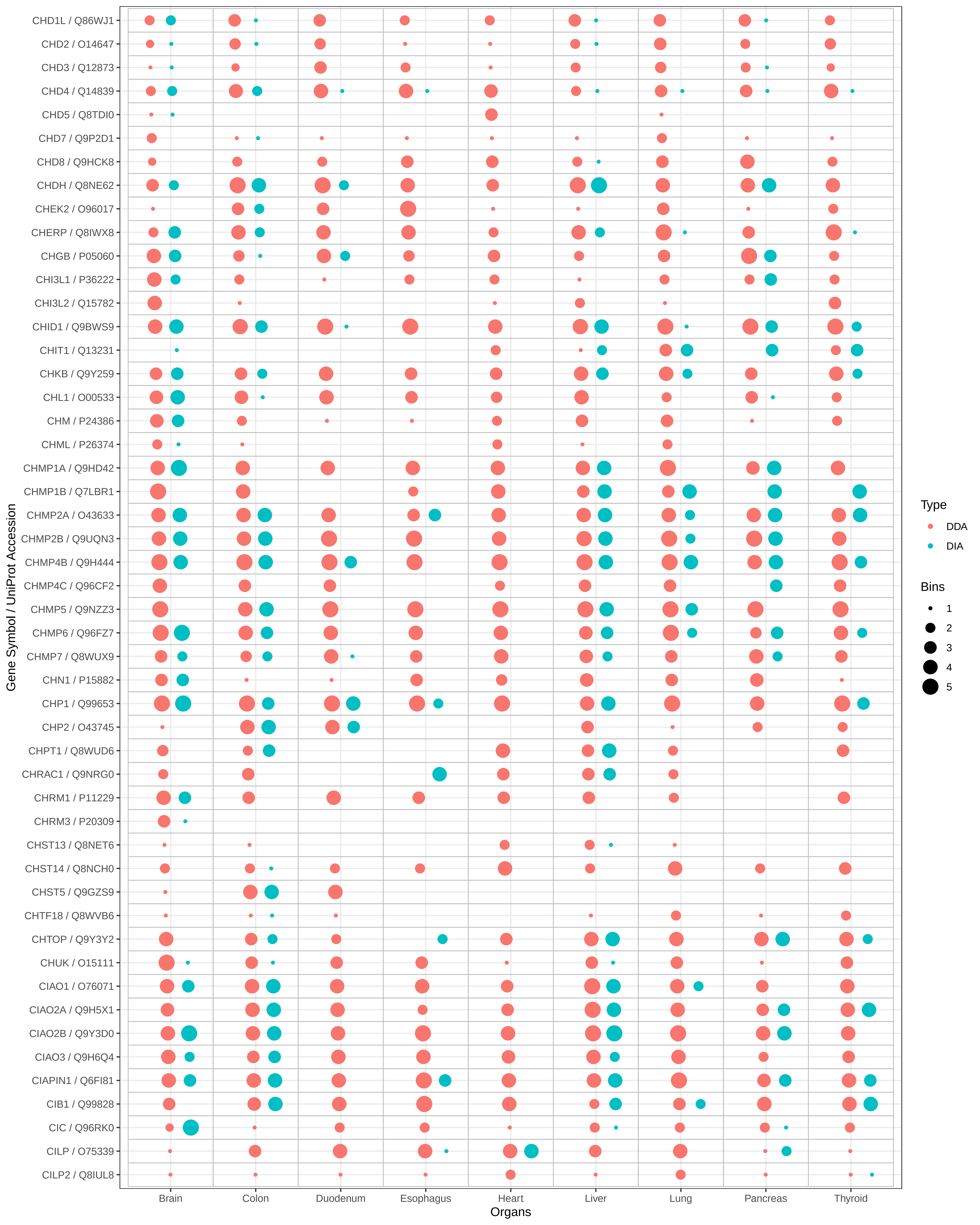

CHD1L / Q86WJ1
CHD2 / O14647
CHD3 / Q12873
CHD4 / Q14839
CHD5 / Q8TDI0
CHD7 / Q9P2D1
CHD8 / Q9HCK8
CHDH / Q8NE62
CHEK2 / O96017
CHERP / Q8IWX8
CHGB / P05060
CHI3L1 / P36222
CHI3L2 / Q15782
CHID1 / Q9BWS9
CHIT1 / Q13231
CHKB / Q9Y259
CHL1 / O00533
CHM / P24386
CHML / P26374
CHMP1A / Q9HD42
CHMP1B / Q7LBR1
CHMP2A / O43633
CHMP2B / Q9UQN3
CHMP4B / Q9H444
CHMP4C / Q96CF2
CHMP5 / Q9NZZ3
CHMP6 / Q96FZ7
CHMP7 / Q8WUX9
CHN1 / P15882
CHP1 / Q99653
CHP2 / O43745
CHPT1 / Q8WUD6
CHRAC1 / Q9NRG0
CHRM1 / P11229
CHRM3 / P20309
CHST13 / Q8NET6
CHST14 / Q8NCH0
CHST5 / Q9GZS9
CHTF18 / Q8WVB6
CHTOP / Q9Y3Y2
CHUK / O15111
CIAO1 / O76071
CIAO2A / Q9H5X1
CIAO2B / Q9Y3D0
CIAO3 / Q9H6Q4
CIAPIN1 / Q6FI81
CIB1 / Q99828
CIC / Q96RK0
CILP / O75339
CILP2 / Q8IUL8
Gene Symbol / UniProt Accession
Brain
Colon
Duodenum
Esophagus
Heart
Liver
Lung
Pancreas
Thyroid
Organs
Type
DDA
DIA
Bins
1
2
3
4
5

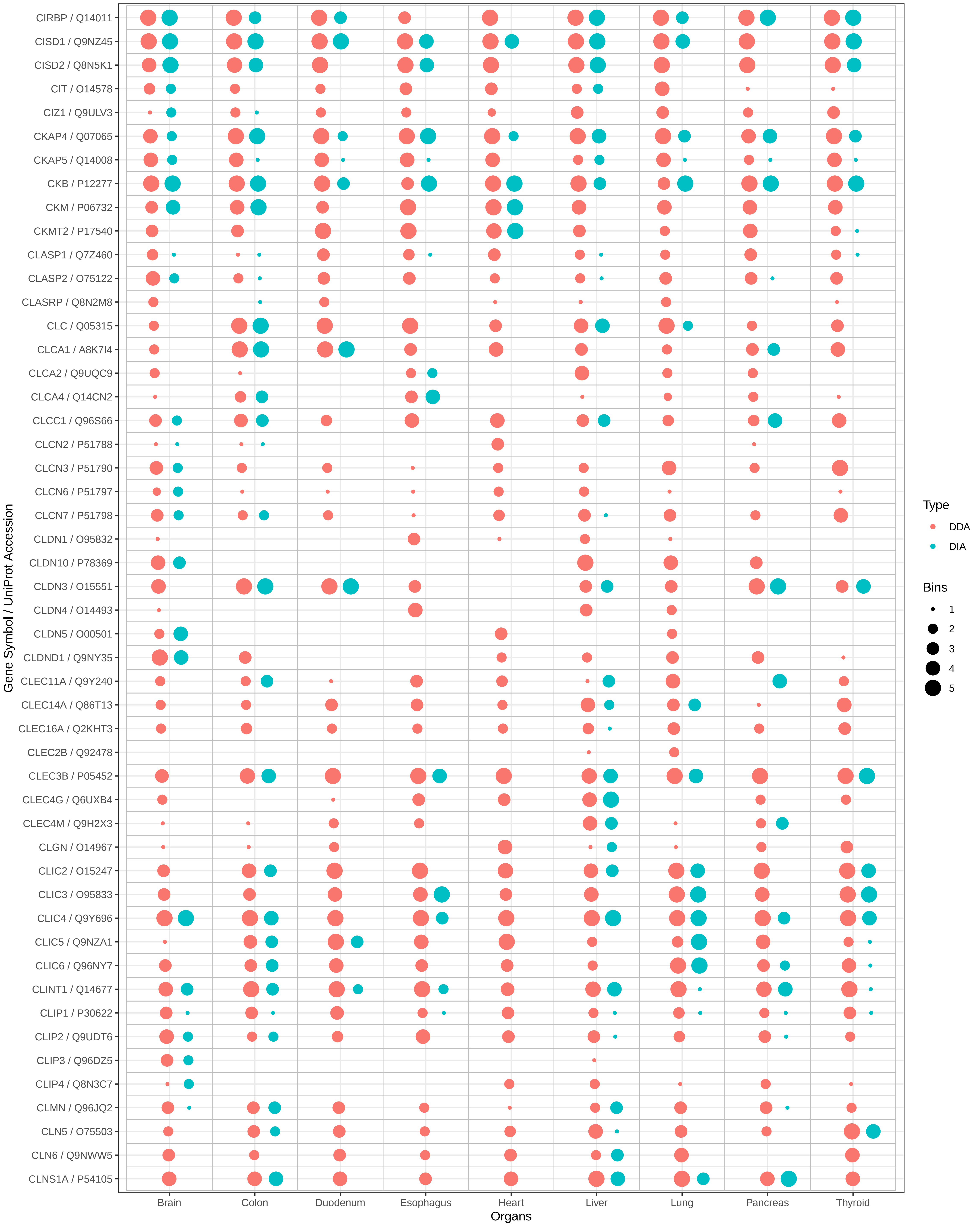
CIRBP / Q14011
CISD1 / Q9NZ45
CISD2 / Q8N5K1
CIT / O14578
CIZ1 / Q9ULV3
CKAP4 / Q07065
CKAP5 / Q14008
CKB / P12277
CKM / P06732
CKMT2 / P17540
CLASP1 / Q7Z460
CLASP2 / O75122
CLASRP / Q8N2M8
CLC / Q05315
CLCA1 / A8K7I4
CLCA2 / Q9UQC9
CLCA4 / Q14CN2
CLCC1 / Q96S66
CLCN2 / P51788
CLCN3 / P51790
CLCN6 / P51797
CLCN7 / P51798
CLDN1 / O95832
CLDN10 / P78369
CLDN3 / O15551
CLDN4 / O14493
CLDN5 / O00501
CLDND1 / Q9NY35
CLEC11A / Q9Y240
CLEC14A / Q86T13
CLEC16A / Q2KHT3
CLEC2B / Q92478
CLEC3B / P05452
CLEC4G / Q6UXB4
CLEC4M / Q9H2X3
CLGN / O14967
CLIC2 / O15247
CLIC3 / O95833
CLIC4 / Q9Y696
CLIC5 / Q9NZA1
CLIC6 / Q96NY7
CLINT1 / Q14677
CLIP1 / P30622
CLIP2 / Q9UDT6
CLIP3 / Q96DZ5
CLIP4 / Q8N3C7
CLMN / Q96JQ2
CLN5 / O75503
CLN6 / Q9NWW5
CLNS1A / P54105
Gene Symbol / UniProt Accession
Brain
Colon
Duodenum
Esophagus
Heart
Liver
Lung
Pancreas
Thyroid
Organs
Type
DDA
DIA
Bins
1
2
3
4
5

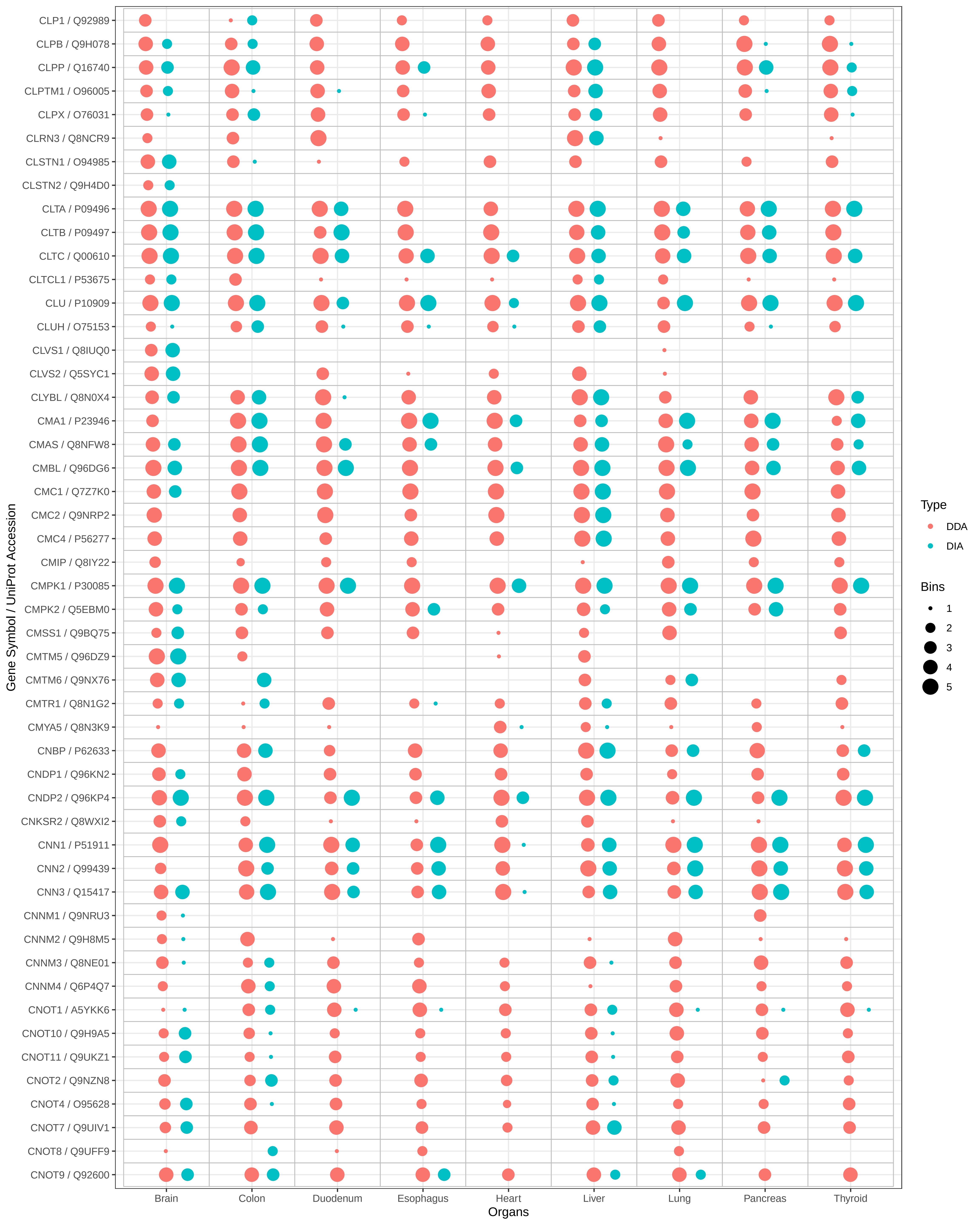

CLP1 / Q92989
CLPB / Q9H078
CLPP / Q16740
CLPTM1 / O96005
CLPX / O76031
CLRN3 / Q8NCR9
CLSTN1 / O94985
CLSTN2 / Q9H4D0
CLTA / P09496
CLTB / P09497
CLTC / Q00610
CLTCL1 / P53675
CLU / P10909
CLUH / O75153
CLVS1 / Q8IUQ0
CLVS2 / Q5SYC1
CLYBL / Q8N0X4
CMA1 / P23946
CMAS / Q8NFW8
CMBL / Q96DG6
CMC1 / Q7Z7K0
CMC2 / Q9NRP2
CMC4 / P56277
CMIP / Q8IY22
CMPK1 / P30085
CMPK2 / Q5EBM0
CMSS1 / Q9BQ75
CMTM5 / Q96DZ9
CMTM6 / Q9NX76
CMTR1 / Q8N1G2
CMYA5 / Q8N3K9
CNBP / P62633
CNDP1 / Q96KN2
CNDP2 / Q96KP4
CNKSR2 / Q8WXI2
CNN1 / P51911
CNN2 / P99439
CNN3 / Q15417
CNNM1 / Q9NRU3
CNNM2 / Q9H8M5
CNNM3 / Q8NE01
CNNM4 / Q6P4Q7
CNOT1 / A5YKK6
CNOT10 / Q9H9A5
CNOT11 / Q9UKZ1
CNOT2 / Q9NZN8
CNOT4 / O95628
CNOT7 / Q9UIV1
CNOT8 / Q9UFF9
CNOT9 / Q92600
Gene Symbol / UniProt Accession
Brain
Colon
Duodenum
Esophagus
Heart
Liver
Lung
Pancreas
Thyroid
Organs
Type
DDA
DIA
Bins
1
2
3
4
5

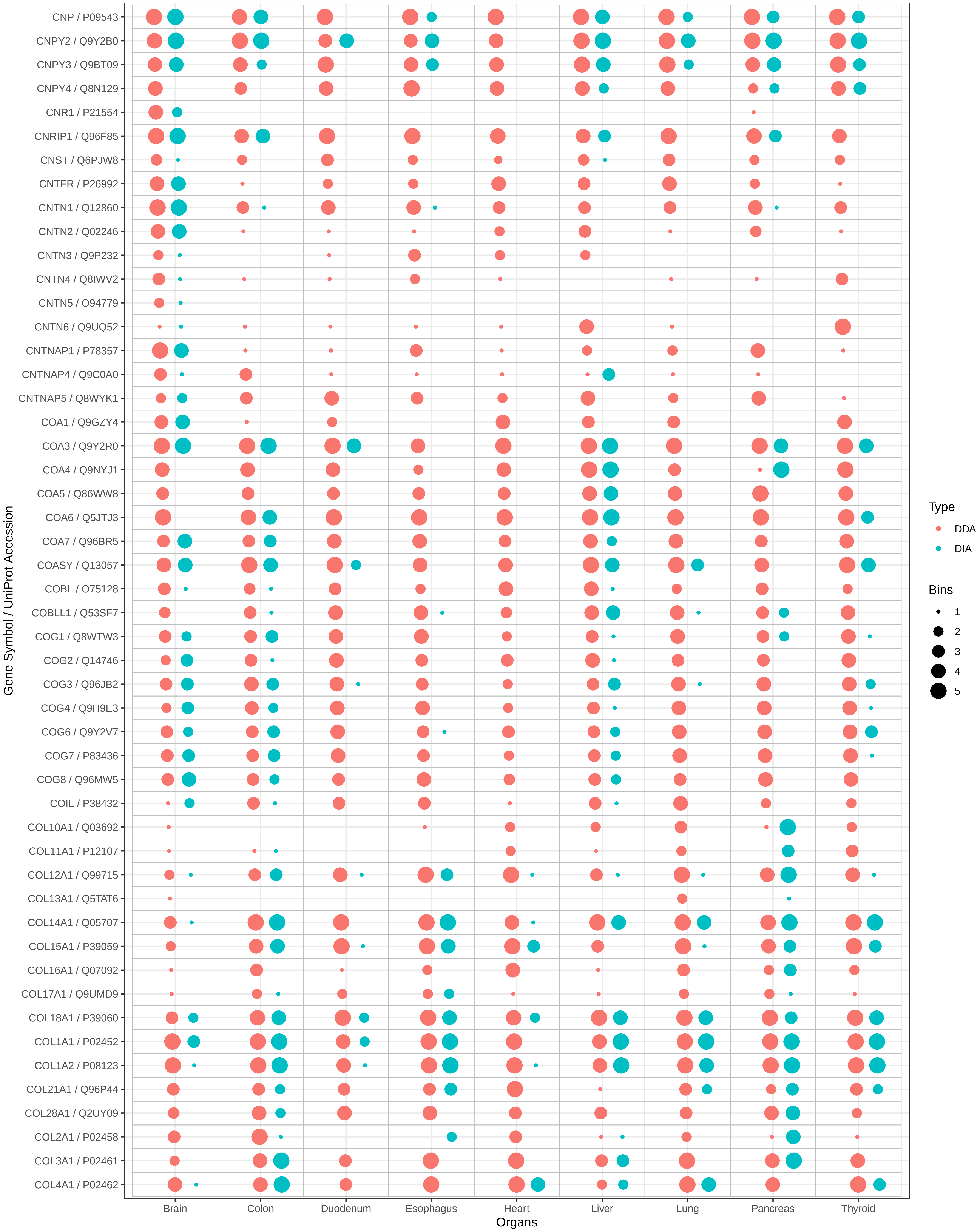

CNP / P09543
CNPY2 / Q9Y2B0
CNPY3 / Q9BT09
CNPY4 / Q8N129
CNR1 / P21554
CNRIP1 / Q96F85
CNST / Q6PJW8
CNTFR / P26992
CNTN1 / Q12860
CNTN2 / Q02246
CNTN3 / Q9P232
CNTN4 / Q8IWV2
CNTN5 / O94779
CNTN6 / Q9UQ52
CNTNAP1 / P78357
CNTNAP4 / Q9C0A0
CNTNAP5 / Q8WYK1
COA1 / Q9GZY4
COA3 / Q9Y2R0
COA4 / Q9NYJ1
COA5 / Q86WW8
COA6 / Q5JTJ3
COA7 / Q96BR5
COASY / Q13057
COBL / O75128
COBLL1 / Q53SF7
COG1 / Q8WTW3
COG2 / Q14746
COG3 / Q96JB2
COG4 / Q9H9E3
COG6 / Q9Y2V7
COG7 / P83436
COG8 / Q96MW5
COIL / P38432
COL10A1 / Q03692
COL11A1 / P12107
COL12A1 / Q99715
COL13A1 / Q5TAT6
COL14A1 / Q05707
COL15A1 / P39059
COL16A1 / Q07092
COL17A1 / Q9UMD9
COL18A1 / P39060
COL1A1 / P02452
COL1A2 / P08123
COL21A1 / Q96P44
COL28A1 / Q2UY09
COL2A1 / P02458
COL3A1 / P02461
COL4A1 / P02462
Gene Symbol / UniProt Accession
Brain
Colon
Duodenum
Esophagus
Heart
Liver
Lung
Pancreas
Thyroid
Organs
Type
DDA
DIA
Bins
1
2
3
4
5

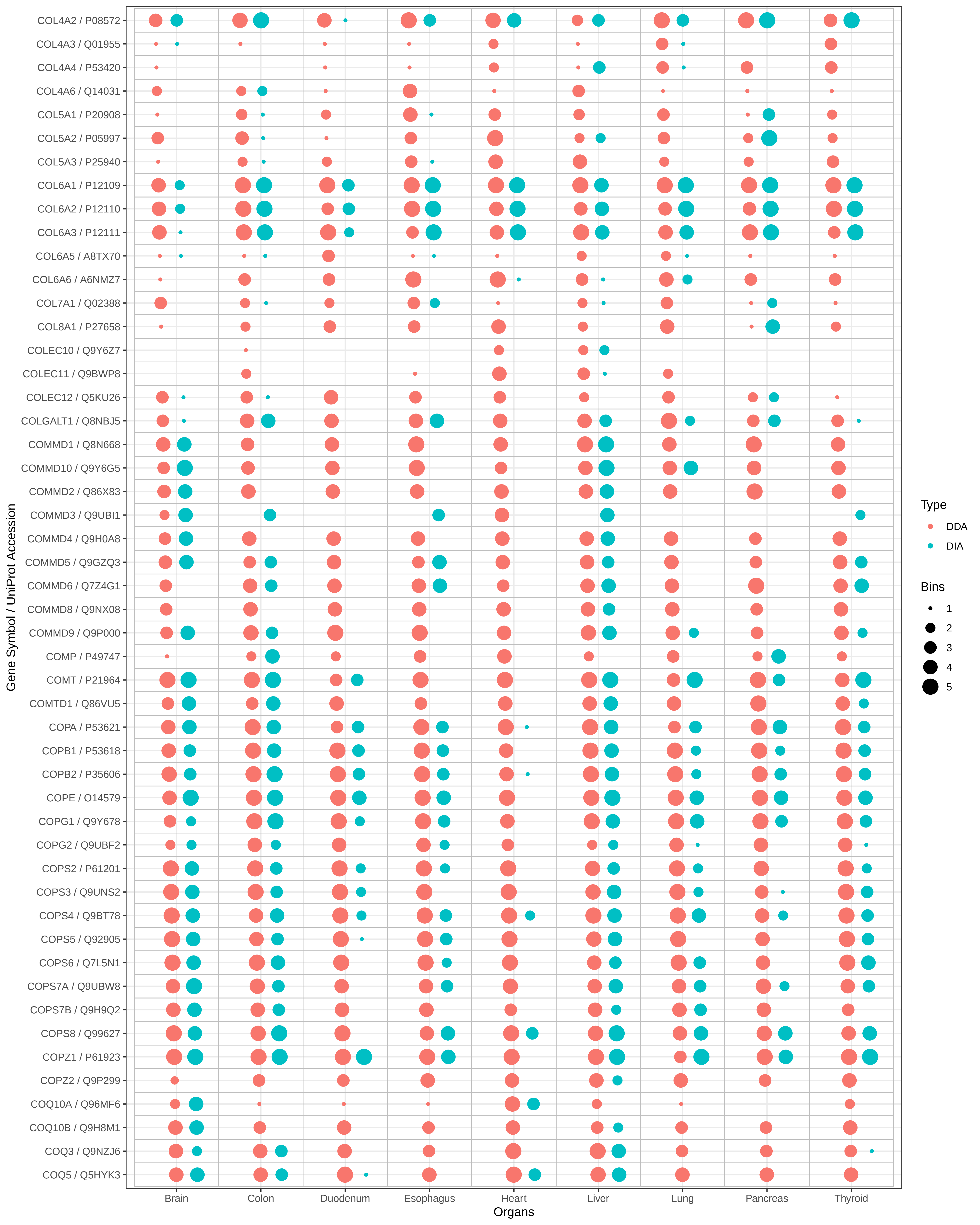

COL4A2 / P08572
COL4A3 / Q01955
COL4A4 / P53420
COL4A6 / Q14031
COL5A1 / P20908
COL5A2 / P05997
COL5A3 / P25940
COL6A1 / P12109
COL6A2 / P12110
COL6A3 / P12111
COL6A5 / A8TX70
COL6A6 / A6NMZ7
COL7A1 / Q02388
COL8A1 / P27658
COLEC10 / Q9Y6Z7
COLEC11 / Q9BWP8
COLEC12 / Q5KU26
COLGALT1 / Q8NBJ5
COMMD1 / Q8N668
COMMD10 / Q9Y6G5
COMMD2 / Q86X83
COMMD3 / Q9UBI1
COMMD4 / Q9H0A8
COMMD5 / Q9GZQ3
COMMD6 / Q7Z4G1
COMMD8 / Q9NX08
COMMD9 / Q9P000
COMP / P49747
COMT / P21964
COMTD1 / Q86VU5
COPA / P53621
COPB1 / P53618
COPB2 / P35606
COPE / O14579
COPG1 / Q9Y678
COPG2 / Q9UBF2
COPS2 / P61201
COPS3 / Q9UNS2
COPS4 / Q9BT78
COPS5 / Q92905
COPS6 / Q7L5N1
COPS7A / Q9UBW8
COPS7B / Q9H9Q2
COPS8 / Q99627
COPZ1 / P61923
COPZ2 / Q9P299
COQ10A / Q96MF6
COQ10B / Q9H8M1
COQ3 / Q9NZJ6
COQ5 / Q5HYK3
Gene Symbol / UniProt Accession
Brain
Colon
Duodenum
Esophagus
Heart
Liver
Lung
Pancreas
Thyroid
Organs
Type
DDA
DIA
Bins
1
2
3
4
5

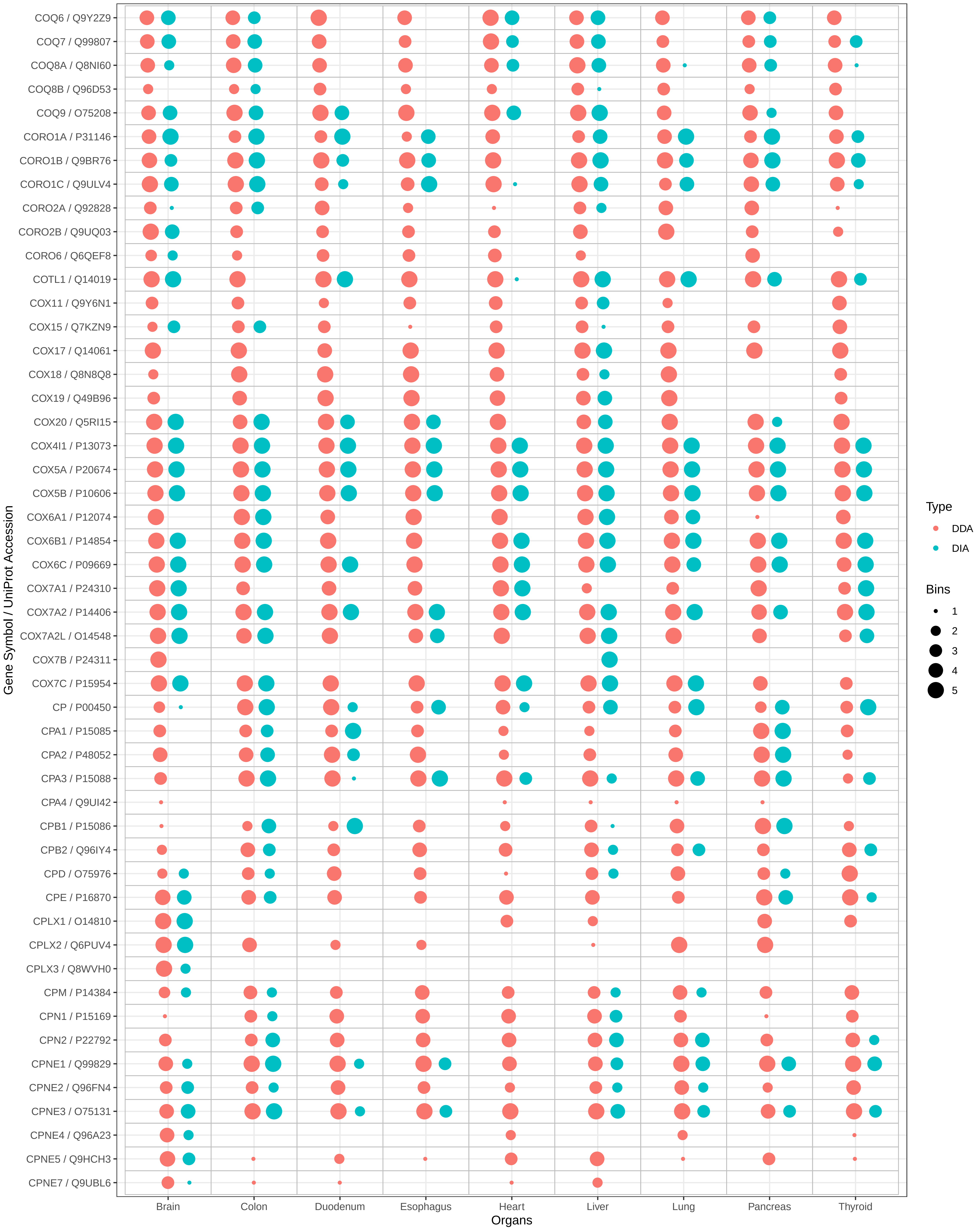
COQ6 / Q9Y2Z9
COQ7 / Q99807
COQ8A / Q8NI60
COQ8B / Q96D53
COQ9 / O75208
CORO1A / P31146
CORO1B / Q9BR76
CORO1C / Q9ULV4
CORO2A / Q92828
CORO2B / Q9UQ03
CORO6 / Q6QEF8
COTL1 / Q14019
COX11 / Q9Y6N1
COX15 / Q7KZN9
COX17 / Q14061
COX18 / Q8N8Q8
COX19 / Q49B96
COX20 / Q5RI15
COX4I1 / P13073
COX5A / P20674
COX5B / P10606
COX6A1 / P12074
COX6B1 / P14854
COX6C / P09669
COX7A1 / P24310
COX7A2 / P14406
COX7A2L / O14548
COX7B / P24311
COX7C / P15954
CP / P00450
CPA1 / P15085
CPA2 / P48052
CPA3 / P15088
CPA4 / Q9UI42
CPB1 / P15086
CPB2 / Q96IY4
CPD / O75976
CPE / P16870
CPLX1 / O14810
CPLX2 / Q6PUV4
CPLX3 / Q8WVH0
CPM / P14384
CPN1 / P15169
CPN2 / P22792
CPNE1 / Q99829
CPNE2 / Q96FN4
CPNE3 / O75131
CPNE4 / Q96A23
CPNE5 / Q9HCH3
CPNE7 / Q9UBL6
Gene Symbol / UniProt Accession
Brain
Colon
Duodenum
Esophagus
Heart
Liver
Lung
Pancreas
Thyroid
Organs
Type
DDA
DIA
Bins
1
2
3
4
5

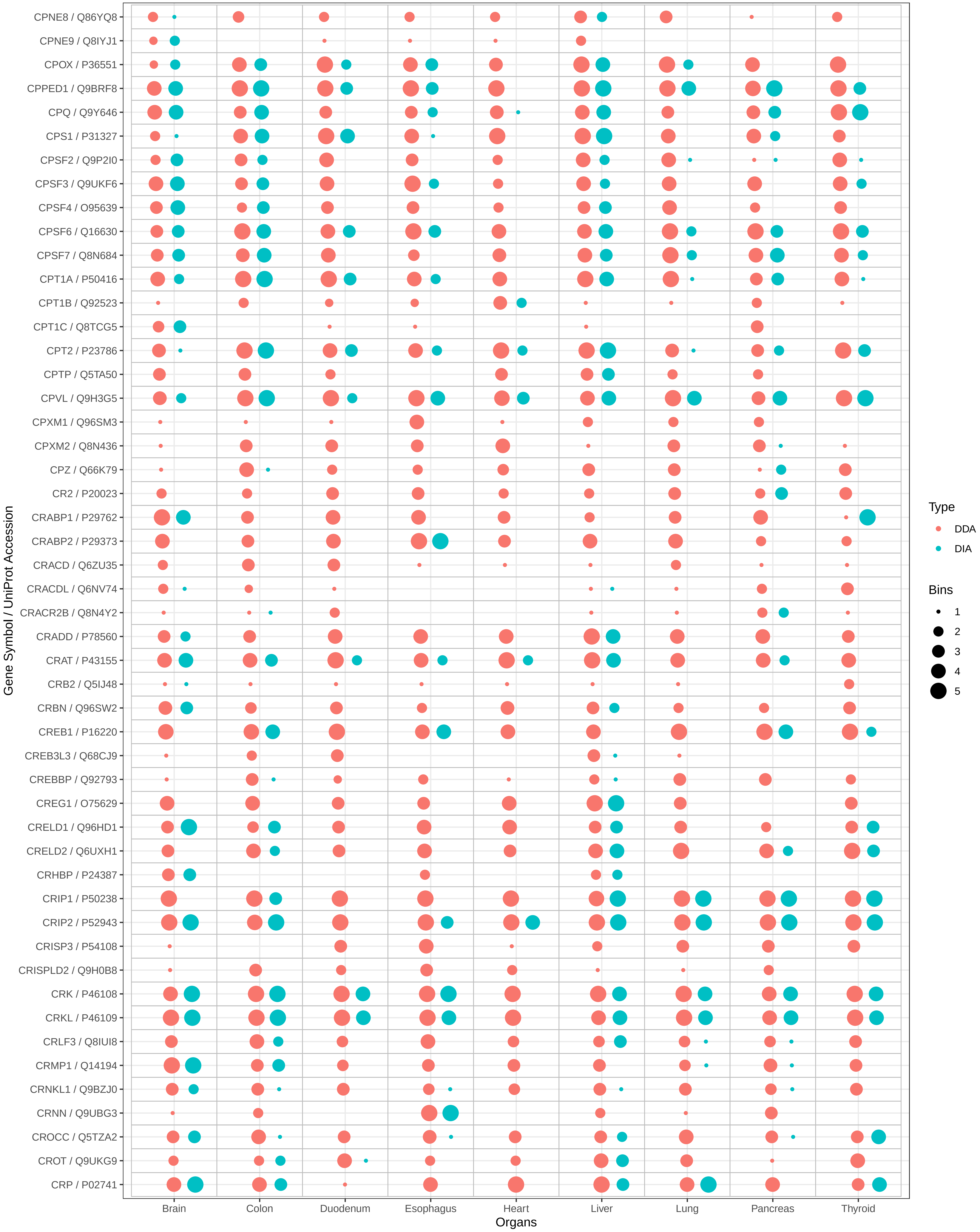

CPNE8 / Q86YQ8
CPNE9 / Q8IYJ1
CPOX / P36551
CPPED1 / Q9BRF8
CPQ / Q9Y646
CPS1 / P31327
CPSF2 / Q9P2I0
CPSF3 / Q9UKF6
CPSF4 / O95639
CPSF6 / Q16630
CPSF7 / Q8N684
CPT1A / P50416
CPT1B / Q92523
CPT1C / Q8TCG5
CPT2 / P23786
CPTP / Q5TA50
CPVL / Q9H3G5
CPXM1 / Q96SM3
CPXM2 / Q8N436
CPZ / Q66K79
CR2 / P20023
CRABP1 / P29762
CRABP2 / P29373
CRACD / Q6ZU35
CRACDL / Q6NV74
CRACR2B / Q8N4Y2
CRADD / P78560
CRAT / P43155
CRB2 / Q5IJ48
CRBN / Q96SW2
CREB1 / P16220
CREB3L3 / Q68CJ9
CREBBP / Q92793
CREG1 / O75629
CRELD1 / Q96HD1
CRELD2 / Q6UXH1
CRHBP / P24387
CRIP1 / P50238
CRIP2 / P52943
CRISP3 / P54108
CRISPLD2 / Q9H0B8
CRK / P46108
CRKL / P46109
CRLF3 / Q8IUI8
CRMP1 / Q14194
CRNKL1 / Q9BZJ0
CRNN / Q9UBG3
CROCC / Q5TZA2
CROT / Q9UKG9
CRP / P02741
Gene Symbol / UniProt Accession
Brain
Colon
Duodenum
Esophagus
Heart
Liver
Lung
Pancreas
Thyroid
Organs
Type
DDA
DIA
Bins
1
2
3
4
5

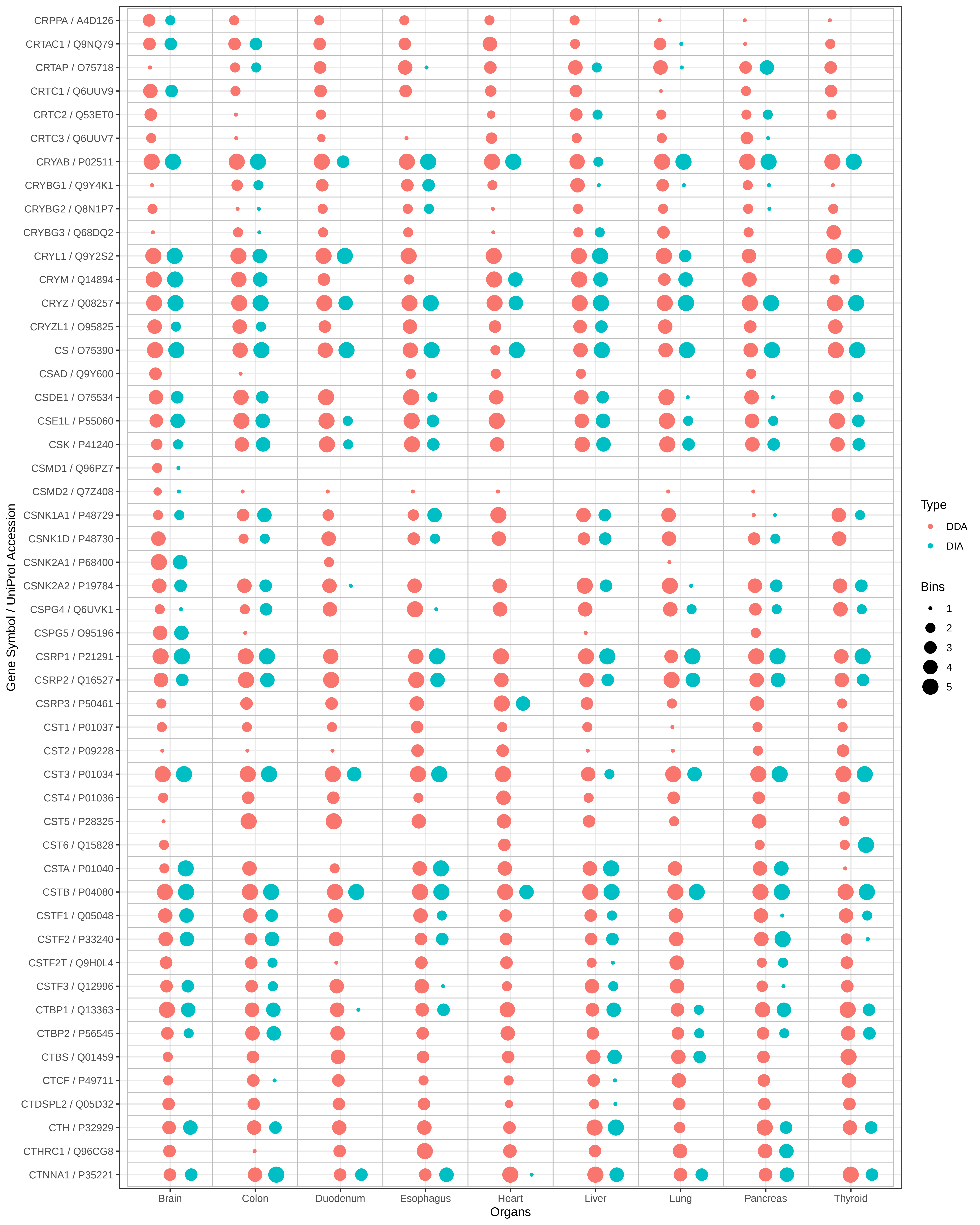
Gene Symbol / UniProt Accession
CRPPA / A4D126
CRTAC1 / Q9NQ79
CRTAP / O75718
CRTC1 / Q6UUV9
CRTC2 / Q53ET0
CRTC3 / Q6UUV7
CRYAB / P02511
CRYBG1 / Q9Y4K1
CRYBG2 / Q8N1P7
CRYBG3 / Q68DQ2
CRYL1 / Q9Y2S2
CRYM / Q14894
CRYZ / Q08257
CRYZL1 / O95825
CS / O75390
CSAD / Q9Y600
CSDE1 / O75534
CSE1L / P55060
CSK / P41240
CSMD1 / Q96PZ7
CSMD2 / Q7Z408
CSNK1A1 / P48729
CSNK1D / P48730
CSNK2A1 / P68400
CSNK2A2 / P19784
CSPG4 / Q6UVK1
CSPG5 / O95196
CSRP1 / P21291
CSRP2 / Q16527
CSRP3 / P50461
CST1 / P01037
CST2 / P09228
CST3 / P01034
CST4 / P01036
CST5 / P28325
CST6 / Q15828
CSTA / P01040
CSTB / P04080
CSTF1 / Q05048
CSTF2 / P33240
CSTF2T / Q9H0L4
CSTF3 / Q12996
CTBP1 / Q13363
CTBP2 / P56545
CTBS / Q01459
CTCF / P49711
CTDSPL2 / Q05D32
CTH / P32929
CTHRC1 / Q96CG8
CTNNA1 / P35221
Brain
Colon
Duodenum
Esophagus
Heart
Liver
Lung
Pancreas
Thyroid
Organs
Type
DDA
DIA
Bins
1
2
3
4
5

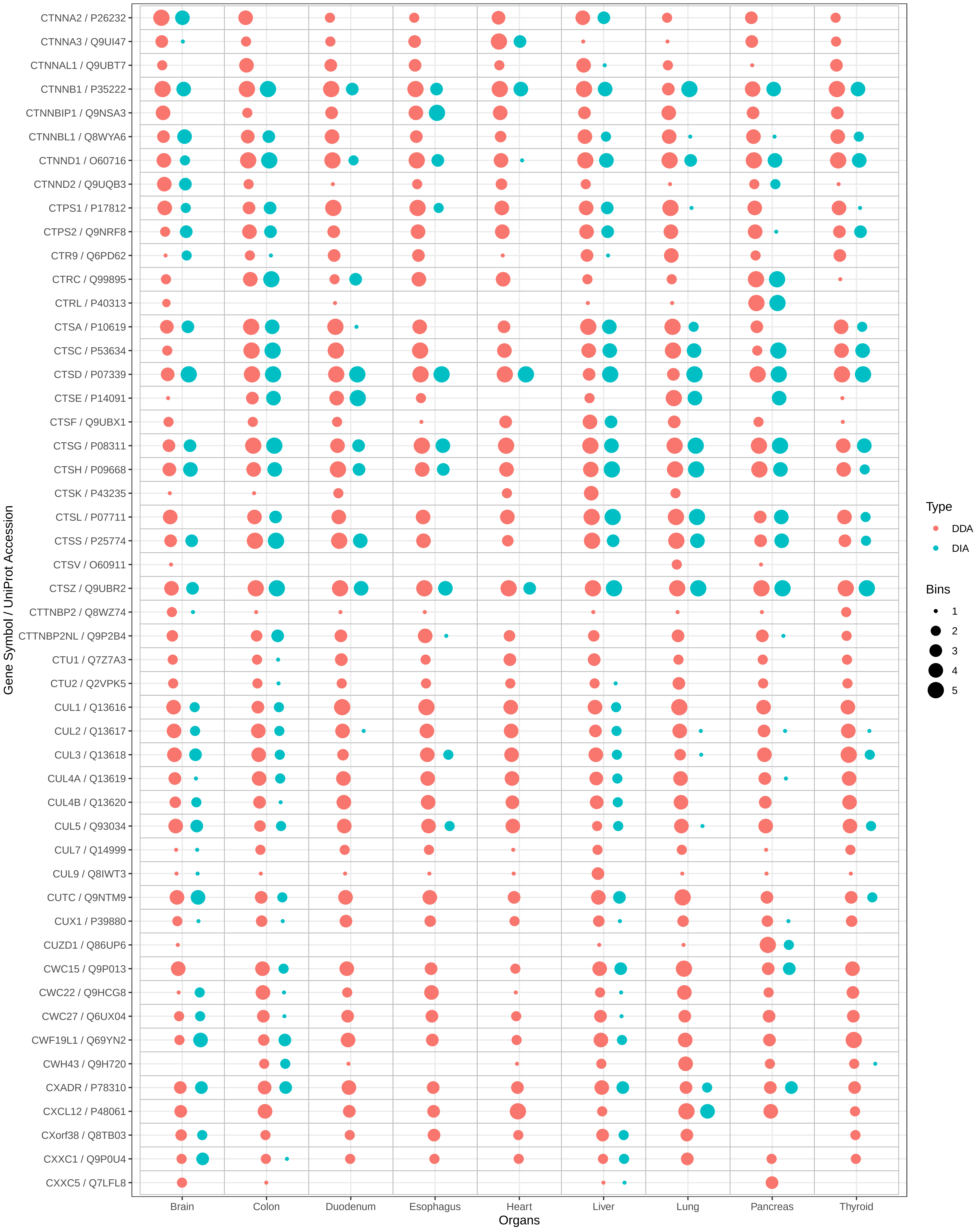
CTNNA2 / P26232
CTNNA3 / Q9UI47
CTNNAL1 / Q9UBT7
CTNNB1 / P35222
CTNNBIP1 / Q9NSA3
CTNNBL1 / Q8WYA6
CTNND1 / O60716
CTNND2 / Q9UQB3
CTPS1 / P17812
CTPS2 / Q9NRF8
CTR9 / Q6PD62
CTRC / Q99895
CTRL / P40313
CTSA / P10619
CTSC / P53634
CTSD / P07339
CTSE / P14091
CTSF / Q9UBX1
CTSG / P08311
CTSH / P09668
CTSK / P43235
CTSL / P07711
CTSS / P25774
CTSV / O60911
CTSZ / Q9UBR2
CTTNBP2 / Q8WZ74
CTTNBP2NL / Q9P2B4
CTU1 / Q7Z7A3
CTU2 / Q2VPK5
CUL1 / Q13616
CUL2 / Q13617
CUL3 / Q13618
CUL4A / Q13619
CUL4B / Q13620
CUL5 / Q93034
CUL7 / Q14999
CUL9 / Q8IWT3
CUTC / Q9NTM9
CUX1 / P39880
CUZD1 / Q86UP6
CWC15 / Q9P013
CWC22 / Q9HCG8
CWC27 / Q6UX04
CWF19L1 / Q69YN2
CWH43 / Q9H720
CXADR / P78310
CXCL12 / P48061
CXorf38 / Q8TB03
CXXC1 / Q9P0U4
CXXC5 / Q7LFL8
Gene Symbol / UniProt Accession
Brain
Colon
Duodenum
Esophagus
Heart
Liver
Lung
Pancreas
Thyroid
Organs
Type
DDA
DIA
Bins
1
2
3
4
5

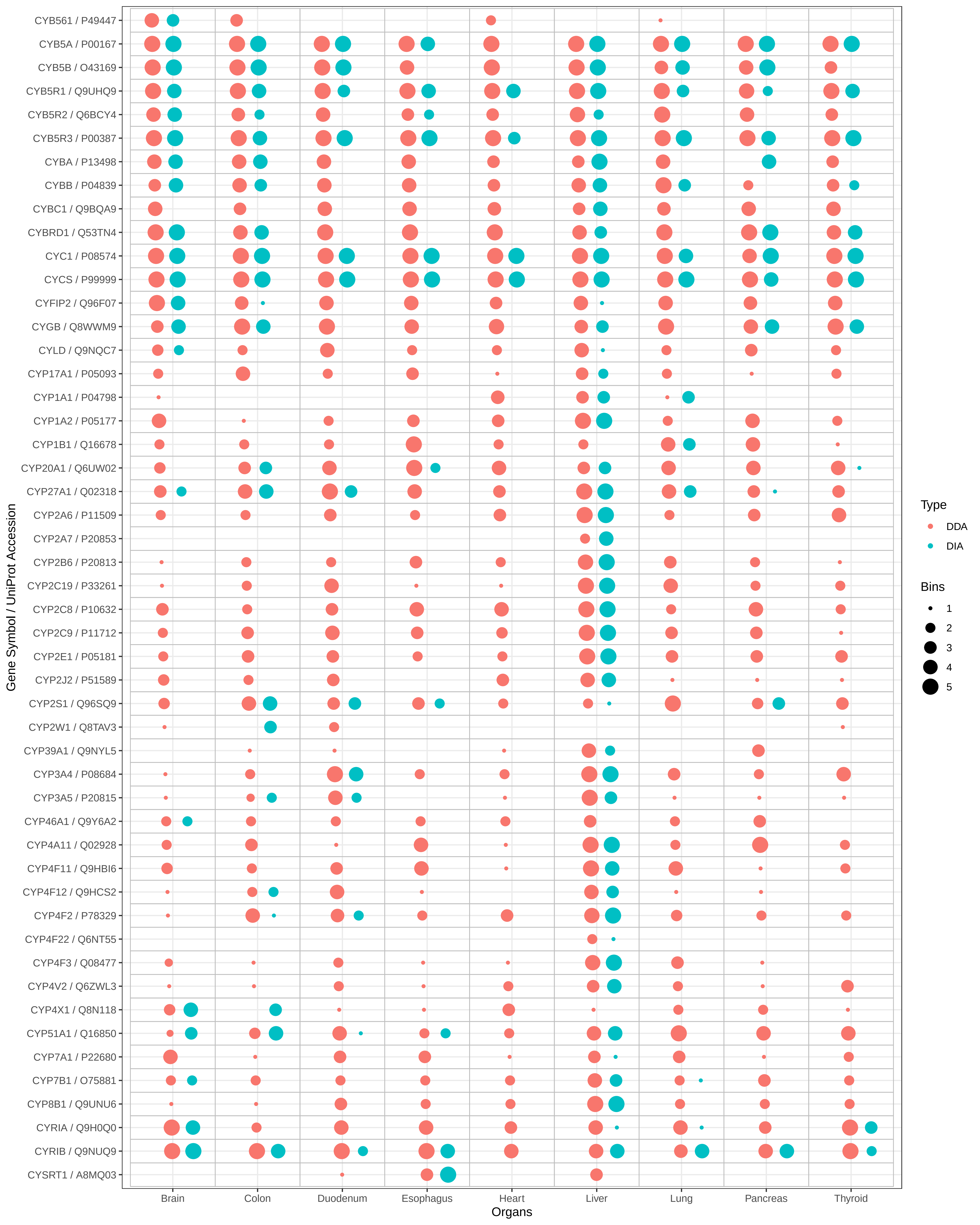
CYB561 / P49447
CYB5A / P00167
CYB5B / O43169
CYB5R1 / Q9UHQ9
CYB5R2 / Q6BCY4
CYB5R3 / P00387
CYBA / P13498
CYBB / P04839
CYBC1 / Q9BQA9
CYBRD1 / Q53TN4
CYC1 / P08574
CYCS / P99999
CYFIP2 / Q96F07
CYGB / Q8WWM9
CYLD / Q9NQC7
CYP17A1 / P05093
CYP1A1 / P04798
CYP1A2 / P05177
CYP1B1 / Q16678
CYP20A1 / Q6UW02
CYP27A1 / Q02318
CYP2A6 / P11509
CYP2A7 / P20853
CYP2B6 / P20813
CYP2C19 / P33261
CYP2C8 / P10632
CYP2C9 / P11712
CYP2E1 / P05181
CYP2J2 / P51589
CYP2S1 / Q96SQ9
CYP2W1 / Q8TAV3
CYP39A1 / Q9NYL5
CYP3A4 / P08684
CYP3A5 / P20815
CYP46A1 / Q9Y6A2
CYP4A11 / Q02928
CYP4F11 / Q9HBI6
CYP4F12 / Q9HCS2
CYP4F2 / P78329
CYP4F22 / Q6NT55
CYP4F3 / Q08477
CYP4V2 / Q6ZWL3
CYP4X1 / Q8N118
CYP51A1 / Q16850
CYP7A1 / P22680
CYP7B1 / O75881
CYP8B1 / Q9UNU6
CYRIA / Q9H0Q0
CYRIB / Q9NUQ9
CYSRT1 / A8MQ03
Gene Symbol / UniProt Accession
Brain
Colon
Duodenum
Esophagus
Heart
Liver
Lung
Pancreas
Thyroid
Organs
Type
DDA
DIA
Bins
1
2
3
4
5

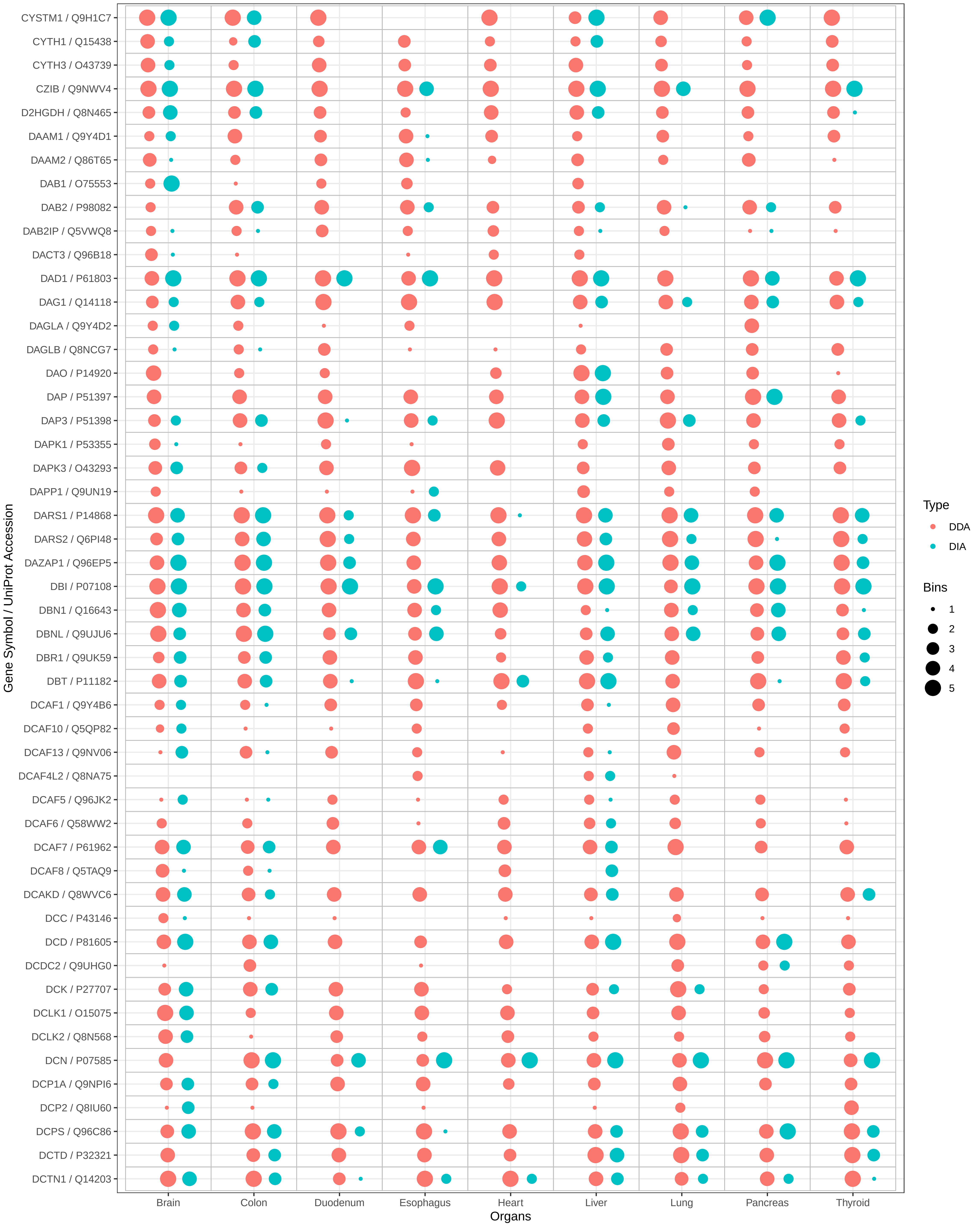

CYSTM1 / Q9H1C7
CYTH1 / Q15438
CYTH3 / O43739
CZIB / Q9NWV4
D2HGDH / Q8N465
DAAM1 / Q9Y4D1
DAAM2 / Q86T65
DAB1 / O75553
DAB2 / P98082
DAB2IP / Q5VWQ8
DACT3 / Q96B18
DAD1 / P61803
DAG1 / Q14118
DAGLA / Q9Y4D2
DAGLB / Q8NCG7
DAO / P14920
DAP / P51397
DAP3 / P51398
DAPK1 / P53355
DAPK3 / O43293
DAPP1 / Q9UN19
DARS1 / P14868
DARS2 / Q6PI48
DAZAP1 / Q96EP5
DBI / P07108
DBN1 / Q16643
DBNL / Q9UJU6
DBR1 / Q9UK59
DBT / P11182
DCAF1 / Q9Y4B6
DCAF10 / Q5QP82
DCAF13 / Q9NV06
DCAF4L2 / Q8NA75
DCAF5 / Q96JK2
DCAF6 / Q58WW2
DCAF7 / P61962
DCAF8 / Q5TAQ9
DCAKD / Q8WVC6
DCC / P43146
DCD / P81605
DCDC2 / Q9UHG0
DCK / P27707
DCLK1 / O15075
DCLK2 / Q8N568
DCN / P07585
DCP1A / Q9NPI6
DCP2 / Q8IU60
DCPS / Q96C86
DCTD / P32321
DCTN1 / Q14203
Gene Symbol / UniProt Accession
Brain
Colon
Duodenum
Esophagus
Heart
Liver
Lung
Pancreas
Thyroid
Organs
Type
DDA
DIA
Bins
1
2
3
4
5

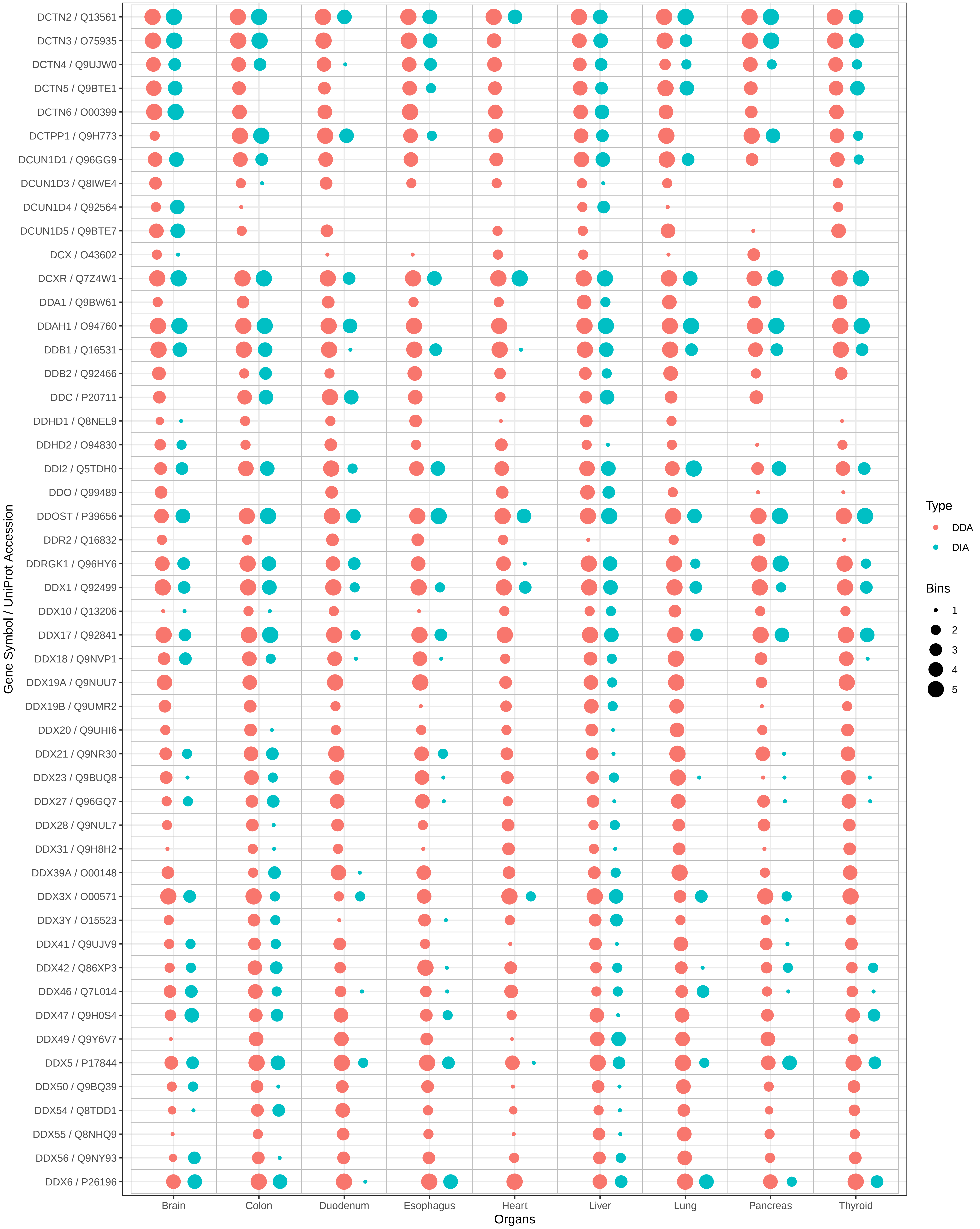
DCTN2 / Q13561
DCTN3 / O75935
DCTN4 / Q9UJW0
DCTN5 / Q9BTE1
DCTN6 / O00399
DCTPP1 / Q9H773
DCUN1D1 / Q96GG9
DCUN1D3 / Q8IWE4
DCUN1D4 / Q92564
DCUN1D5 / Q9BTE7
DCX / O43602
DCXR / Q7Z4W1
DDA1 / Q9BW61
DDAH1 / O94760
DDB1 / Q16531
DDB2 / Q92466
DDC / P20711
DDHD1 / Q8NEL9
DDHD2 / O94830
DDI2 / Q5TDH0
DDO / Q99489
DDOST / P39656
DDR2 / Q16832
DDRGK1 / Q96HY6
DDX1 / Q92499
DDX10 / Q13206
DDX17 / Q92841
DDX18 / Q9NVP1
DDX19A / Q9NUU7
DDX19B / Q9UMR2
DDX20 / Q9UHI6
DDX21 / Q9NR30
DDX23 / Q9BUQ8
DDX27 / Q96GQ7
DDX28 / Q9NUL7
DDX31 / Q9H8H2
DDX39A / O00148
DDX3X / O00571
DDX3Y / O15523
DDX41 / Q9UJV9
DDX42 / Q86XP3
DDX46 / Q7L014
DDX47 / Q9H0S4
DDX49 / Q9Y6V7
DDX5 / P17844
DDX50 / Q9BQ39
DDX54 / Q8TDD1
DDX55 / Q8NHQ9
DDX56 / Q9NY93
DDX6 / P26196
Gene Symbol / UniProt Accession
Brain
Colon
Duodenum
Esophagus
Heart
Liver
Lung
Pancreas
Thyroid
Organs
Type
DDA
DIA
Bins
1
2
3
4
5

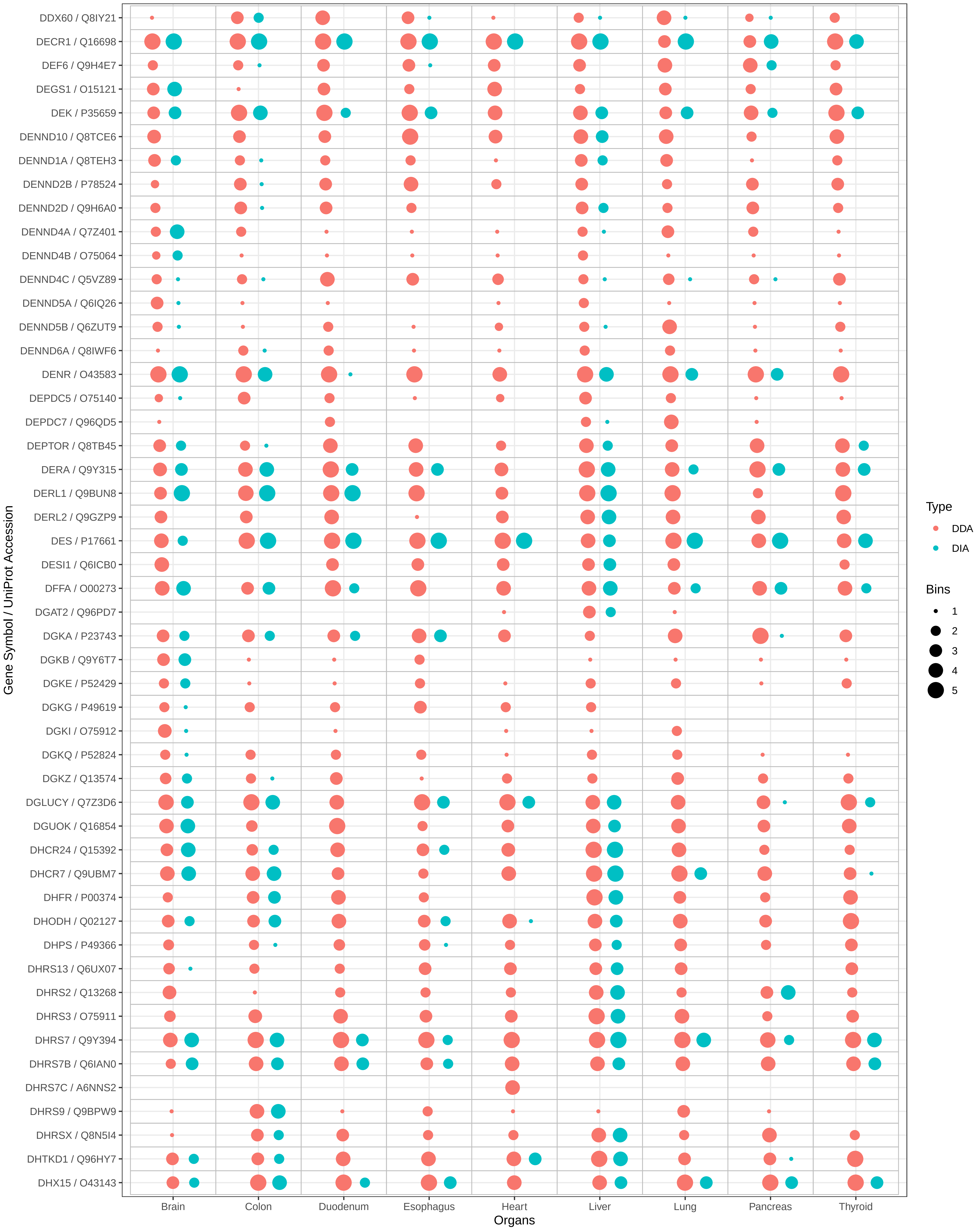

DDX60 / Q8IY21
DECR1 / Q16698
DEF6 / Q9H4E7
DEGS1 / O15121
DEK / P35659
DENND10 / Q8TCE6
DENND1A / Q8TEH3
DENND2B / P78524
DENND2D / Q9H6A0
DENND4A / Q7Z401
DENND4B / O75064
DENND4C / Q5VZ89
DENND5A / Q6IQ26
DENND5B / Q6ZUT9
DENND6A / Q8IWF6
DENR / O43583
DEPDC5 / O75140
DEPDC7 / Q96QD5
DEPTOR / Q8TB45
DERA / Q9Y315
DERL1 / Q9BUN8
DERL2 / Q9GZP9
DES / P17661
DESI1 / Q6ICB0
DFFA / O00273
DGAT2 / Q96PD7
DGKA / P23743
DGKB / Q9Y6T7
DGKE / P52429
DGKG / P49619
DGKI / O75912
DGKQ / P52824
DGKZ / Q13574
DGLUCY / Q7Z3D6
DGUOK / Q16854
DHCR24 / Q15392
DHCR7 / Q9UBM7
DHFR / P00374
DHODH / Q02127
DHPS / P49366
DHRS13 / Q6UX07
DHRS2 / Q13268
DHRS3 / O75911
DHRS7 / Q9Y394
DHRS7B / Q6IAN0
DHRS7C / A6NNS2
DHRS9 / Q9BPW9
DHRSX / Q8N5I4
DHTKD1 / Q96HY7
DHX15 / O43143
Gene Symbol / UniProt Accession
Brain
Colon
Duodenum
Esophagus
Heart
Liver
Lung
Pancreas
Thyroid
Organs
Type
DDA
DIA
Bins
1
2
3
4
5

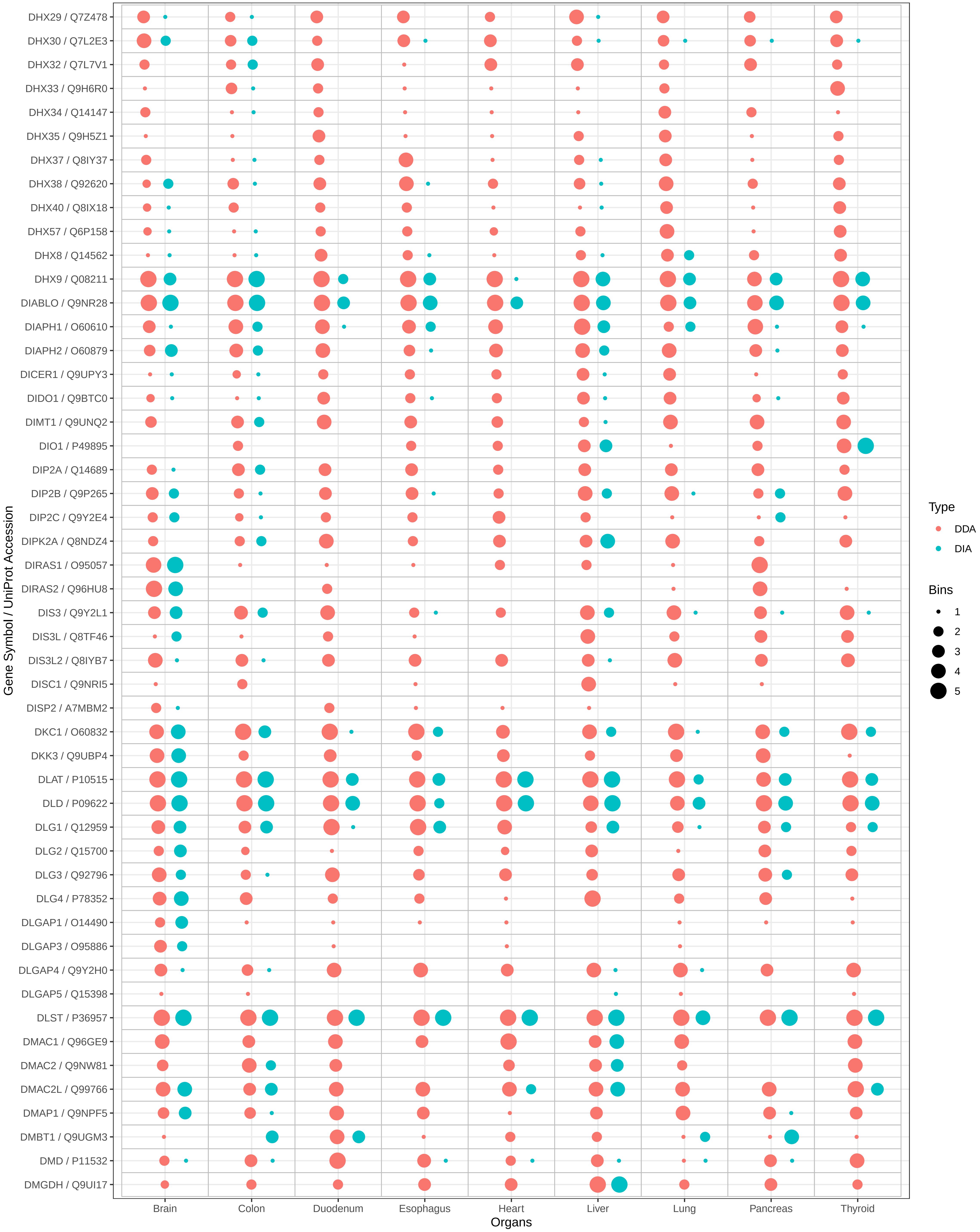

DHX29 / Q7Z478
DHX30 / Q7L2E3
DHX32 / Q7L7V1
DHX33 / Q9H6R0
DHX34 / Q14147
DHX35 / Q9H5Z1
DHX37 / Q8IY37
DHX38 / Q92620
DHX40 / Q8IX18
DHX57 / Q6P158
DHX8 / Q14562
DHX9 / Q08211
DIABLO / Q9NR28
DIAPH1 / O60610
DIAPH2 / O60879
DICER1 / Q9UPY3
DIDO1 / Q9BTC0
DIMT1 / Q9UNQ2
DIO1 / P49895
DIP2A / Q14689
DIP2B / Q9P265
DIP2C / Q9Y2E4
DIPK2A / Q8NDZ4
DIRAS1 / O95057
DIRAS2 / Q96HU8
DIS3 / Q9Y2L1
DIS3L / Q8TF46
DIS3L2 / Q8IYB7
DISC1 / Q9NRI5
DISP2 / A7MBM2
DKC1 / O60832
DKK3 / Q9UBP4
DLAT / P10515
DLD / P09622
DLG1 / Q12959
DLG2 / Q15700
DLG3 / Q92796
DLG4 / P78352
DLGAP1 / O14490
DLGAP3 / O95886
DLGAP4 / Q9Y2H0
DLGAP5 / Q15398
DLST / P36957
DMAC1 / Q96GE9
DMAC2 / Q9NW81
DMAC2L / Q99766
DMAP1 / Q9NPF5
DMBT1 / Q9UGM3
DMD / P11532
DMGDH / Q9UI17
Gene Symbol / UniProt Accession
Brain
Colon
Duodenum
Esophagus
Heart
Liver
Lung
Pancreas
Thyroid
Organs
Type
DDA
DIA
Bins
1
2
3
4
5

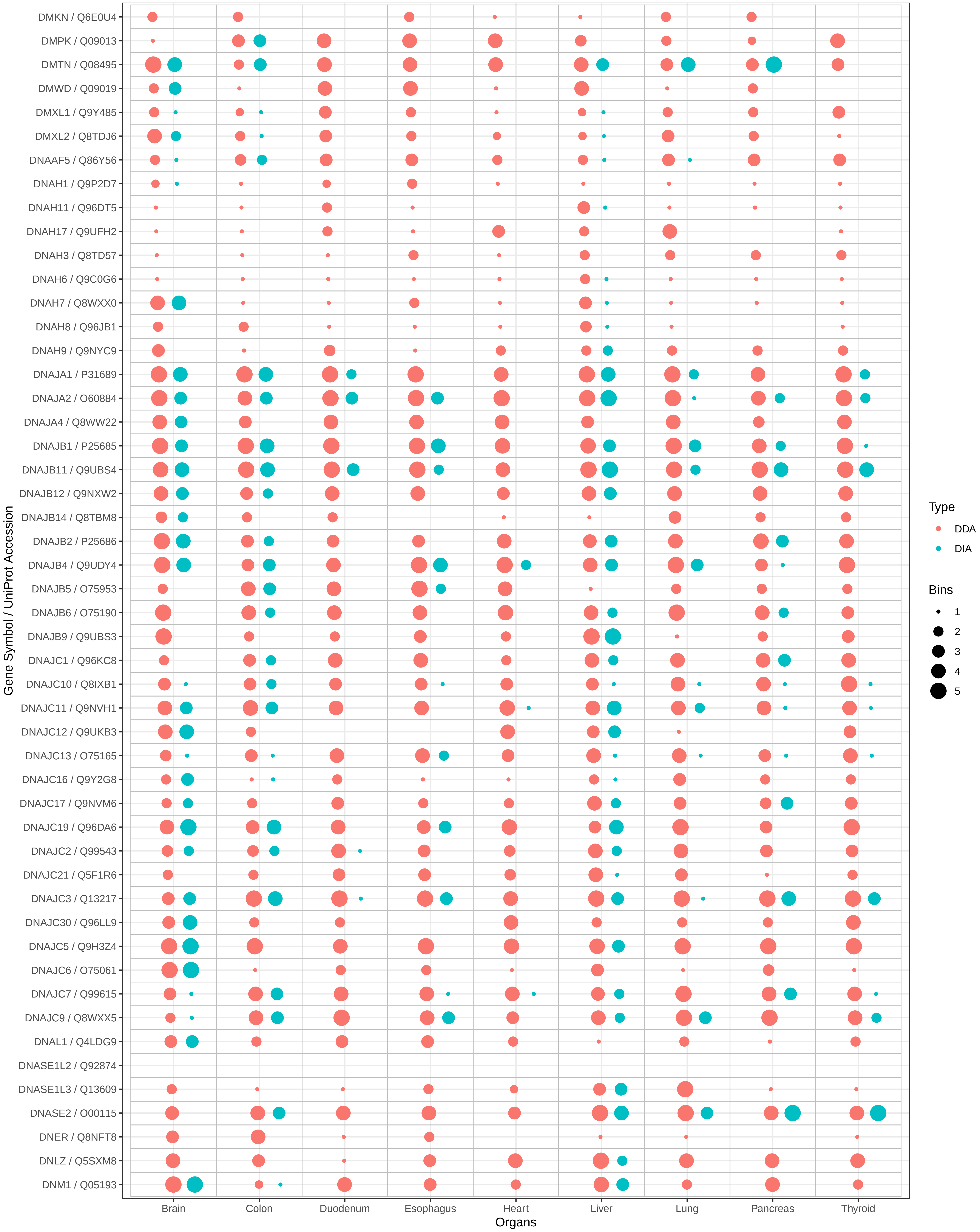
DMKN / Q6E0U4
DMPK / Q09013
DMTN / Q08495
DMWD / Q09019
DMXL1 / Q9Y485
DMXL2 / Q8TDJ6
DNAAF5 / Q86Y56
DNAH1 / Q9P2D7
DNAH11 / Q96DT5
DNAH17 / Q9UFH2
DNAH3 / Q8TD57
DNAH6 / Q9C0G6
DNAH7 / Q8WXX0
DNAH8 / Q96JB1
DNAH9 / Q9NYC9
DNAJA1 / P31689
DNAJA2 / O60884
DNAJA4 / Q8WW22
DNAJB1 / P25685
DNAJB11 / Q9UBS4
DNAJB12 / Q9NXW2
DNAJB14 / Q8TBM8
DNAJB2 / P25686
DNAJB4 / Q9UDY4
DNAJB5 / O75953
DNAJB6 / O75190
DNAJB9 / Q9UBS3
DNAJC1 / Q96KC8
DNAJC10 / Q8IXB1
DNAJC11 / Q9NVH1
DNAJC12 / Q9UKB3
DNAJC13 / O75165
DNAJC16 / Q9Y2G8
DNAJC17 / Q9NVM6
DNAJC19 / Q96DA6
DNAJC2 / Q99543
DNAJC21 / Q5F1R6
DNAJC3 / Q13217
DNAJC30 / Q96LL9
DNAJC5 / Q9H3Z4
DNAJC6 / O75061
DNAJC7 / Q99615
DNAJC9 / Q8WXX5
DNAL1 / Q4LDG9
DNASE1L2 / Q92874
DNASE1L3 / Q13609
DNASE2 / O00115
DNER / Q8NFT8
DNLZ / Q5SXM8
DNM1 / Q05193
Gene Symbol / UniProt Accession
Brain
Colon
Duodenum
Esophagus
Heart
Liver
Lung
Pancreas
Thyroid
Organs
Type
DDA
DIA
Bins
1
2
3
4
5

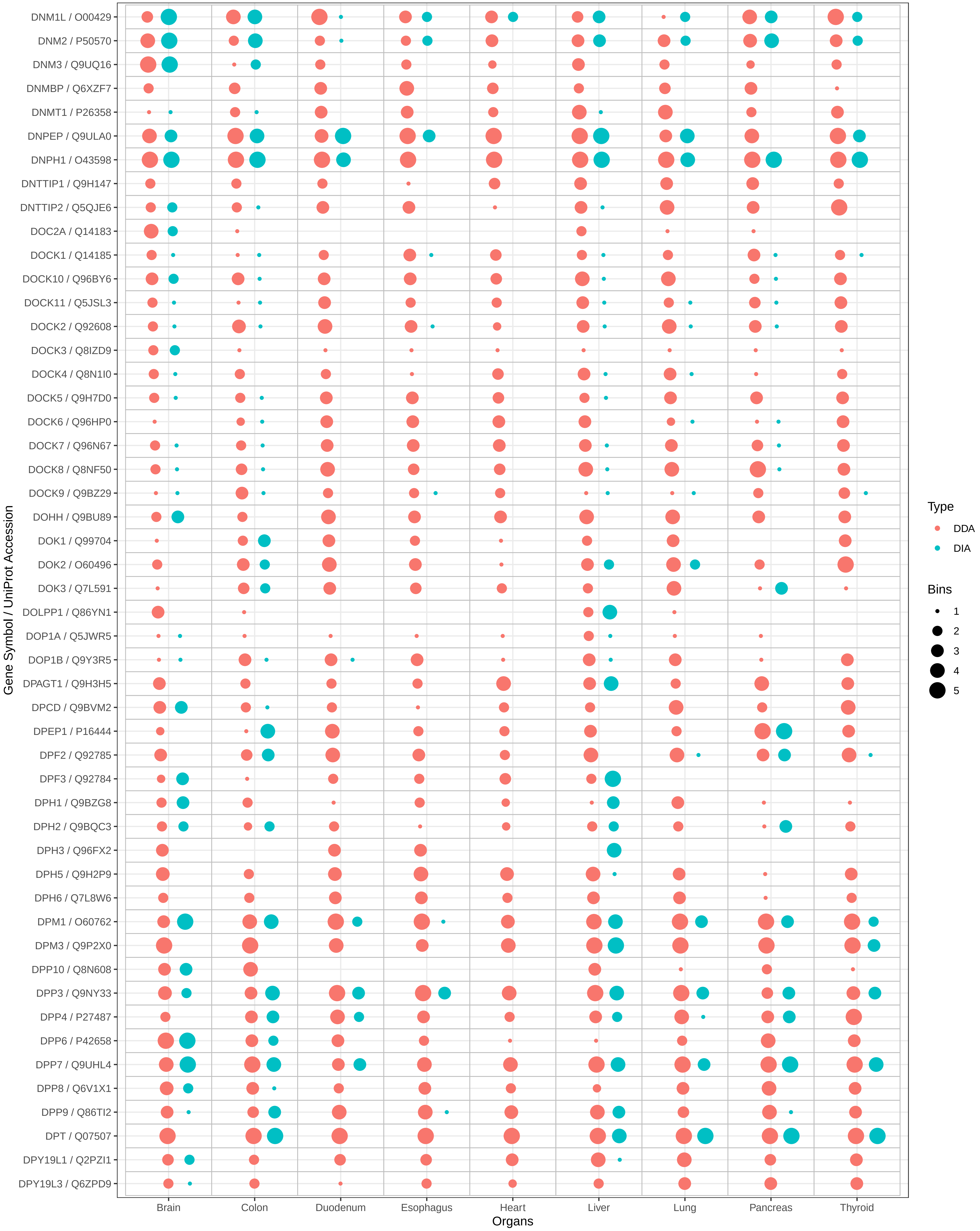

DNM1L / O00429
DNM2 / P50570
DNM3 / Q9UQ16
DNMBP / Q6XZF7
DNMT1 / P26358
DNPEP / Q9ULA0
DNPH1 / O43598
DNTTIP1 / Q9H147
DNTTIP2 / Q5QJE6
DOC2A / Q14183
DOCK1 / Q14185
DOCK10 / Q96BY6
DOCK11 / Q5JSL3
DOCK2 / Q92608
DOCK3 / Q8IZD9
DOCK4 / Q8N1I0
DOCK5 / Q9H7D0
DOCK6 / Q96HP0
DOCK7 / Q96N67
DOCK8 / Q8NF50
DOCK9 / Q9BZ29
DOHH / Q9BU89
DOK1 / Q99704
DOK2 / O60496
DOK3 / Q7L591
DOLPP1 / Q86YN1
DOP1A / Q5JWR5
DOP1B / Q9Y3R5
DPAGT1 / Q9H3H5
DPCD / Q9BVM2
DPEP1 / P16444
DPF2 / Q92785
DPF3 / Q92784
DPH1 / Q9BZG8
DPH2 / Q9BQC3
DPH3 / Q96FX2
DPH5 / Q9H2P9
DPH6 / Q7L8W6
DPM1 / O60762
DPM3 / Q9P2X0
DPP10 / Q8N608
DPP3 / Q9NY33
DPP4 / P27487
DPP6 / P42658
DPP7 / Q9UHL4
DPP8 / Q6V1X1
DPP9 / Q86TI2
DPT / Q07507
DPY19L1 / Q2PZI1
DPY19L3 / Q6ZPD9
Gene Symbol / UniProt Accession
Brain
Colon
Duodenum
Esophagus
Heart
Liver
Lung
Pancreas
Thyroid
Organs
Type
DDA
DIA
Bins
1
2
3
4
5

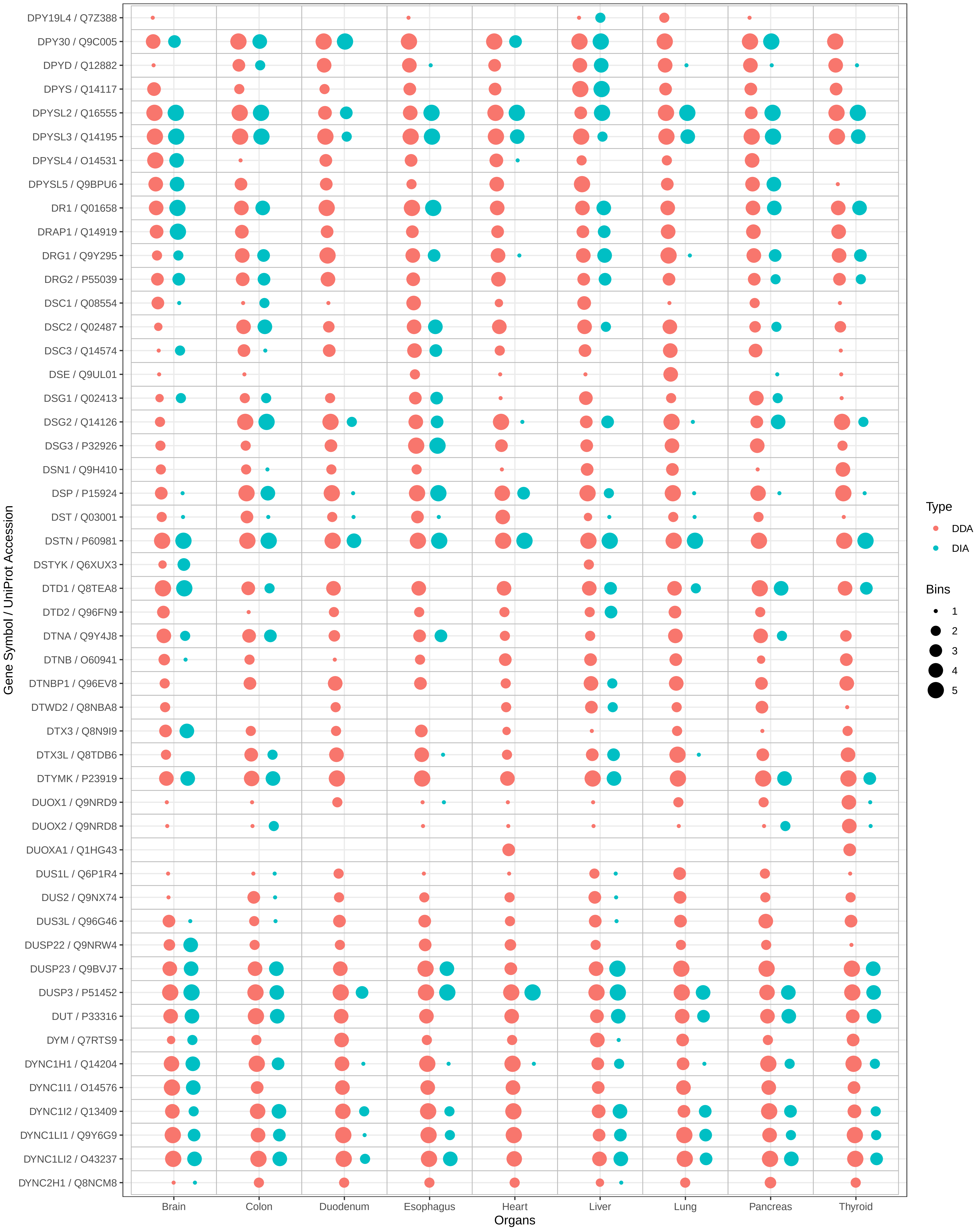

DPY19L4 / Q7Z388
DPY30 / Q9C005
DPYD / Q12882
DPYS / Q14117
DPYSL2 / Q16555
DPYSL3 / Q14195
DPYSL4 / O14531
DPYSL5 / Q9BPU6
DR1 / Q01658
DRAP1 / Q14919
DRG1 / Q9Y295
DRG2 / P55039
DSC1 / Q08554
DSC2 / Q02487
DSC3 / Q14574
DSE / Q9UL01
DSG1 / Q02413
DSG2 / Q14126
DSG3 / P32926
DSN1 / Q9H410
DSP / P15924
DST / Q03001
DSTN / P60981
DSTYK / Q6XUX3
DTD1 / Q8TEA8
DTD2 / Q96FN9
DTNA / Q9Y4J8
DTNB / O60941
DTNBP1 / Q96EV8
DTWD2 / Q8NBA8
DTX3 / Q8N9I9
DTX3L / Q8TDB6
DTYMK / P23919
DUOX1 / Q9NRD9
DUOX2 / Q9NRD8
DUOXA1 / Q1HG43
DUS1L / Q6P1R4
DUS2 / Q9NX74
DUS3L / Q96G46
DUSP22 / Q9NRW4
DUSP23 / Q9BVJ7
DUSP3 / P51452
DUT / P33316
DYM / Q7RTS9
DYNC1H1 / Q14204
DYNC1I1 / O14576
DYNC1I2 / Q13409
DYNC1LI1 / Q9Y6G9
DYNC1LI2 / O43237
DYNC2H1 / Q8NCM8
Gene Symbol / UniProt Accession
Brain
Colon
Duodenum
Esophagus
Heart
Liver
Lung
Pancreas
Thyroid
Organs
Type
DDA
DIA
Bins
1
2
3
4
5

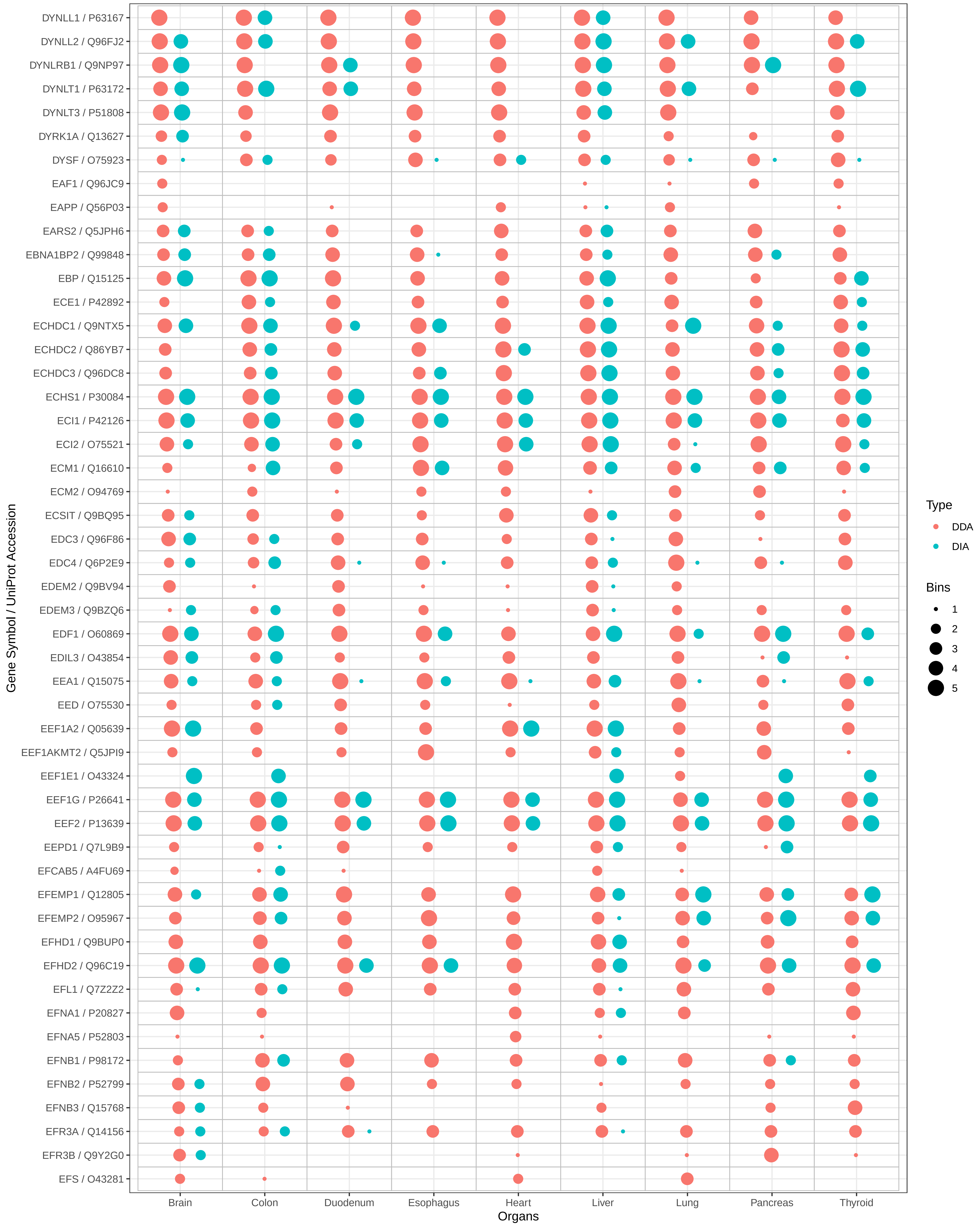

DYNLL1 / P63167
DYNLL2 / Q96FJ2
DYNLRB1 / Q9NP97
DYNLT1 / P63172
DYNLT3 / P51808
DYRK1A / Q13627
DYSF / O75923
EAF1 / Q96JC9
EAPP / Q56P03
EARS2 / Q5JPH6
EBNA1BP2 / Q99848
EBP / Q15125
ECE1 / P42892
ECHDC1 / Q9NTX5
ECHDC2 / Q86YB7
ECHDC3 / Q96DC8
ECHS1 / P30084
ECI1 / P42126
ECI2 / O75521
ECM1 / Q16610
ECM2 / O94769
ECSIT / Q9BQ95
EDC3 / Q96F86
EDC4 / Q6P2E9
EDEM2 / Q9BV94
EDEM3 / Q9BZQ6
EDF1 / O60869
EDIL3 / O43854
EEA1 / Q15075
EED / O75530
EEF1A2 / Q05639
EEF1AKMT2 / Q5JPI9
EEF1E1 / O43324
EEF1G / P26641
EEF2 / P13639
EEPD1 / Q7L9B9
EFCAB5 / A4FU69
EFEMP1 / Q12805
EFEMP2 / O95967
EFHD1 / Q9BUP0
EFHD2 / Q96C19
EFL1 / Q7Z2Z2
EFNA1 / P20827
EFNA5 / P52803
EFNB1 / P98172
EFNB2 / P52799
EFNB3 / Q15768
EFR3A / Q14156
EFR3B / Q9Y2G0
EFS / O43281
Gene Symbol / UniProt Accession
Brain
Colon
Duodenum
Esophagus
Heart
Liver
Lung
Pancreas
Thyroid
Organs
Type
DDA
DIA
Bins
1
2
3
4
5

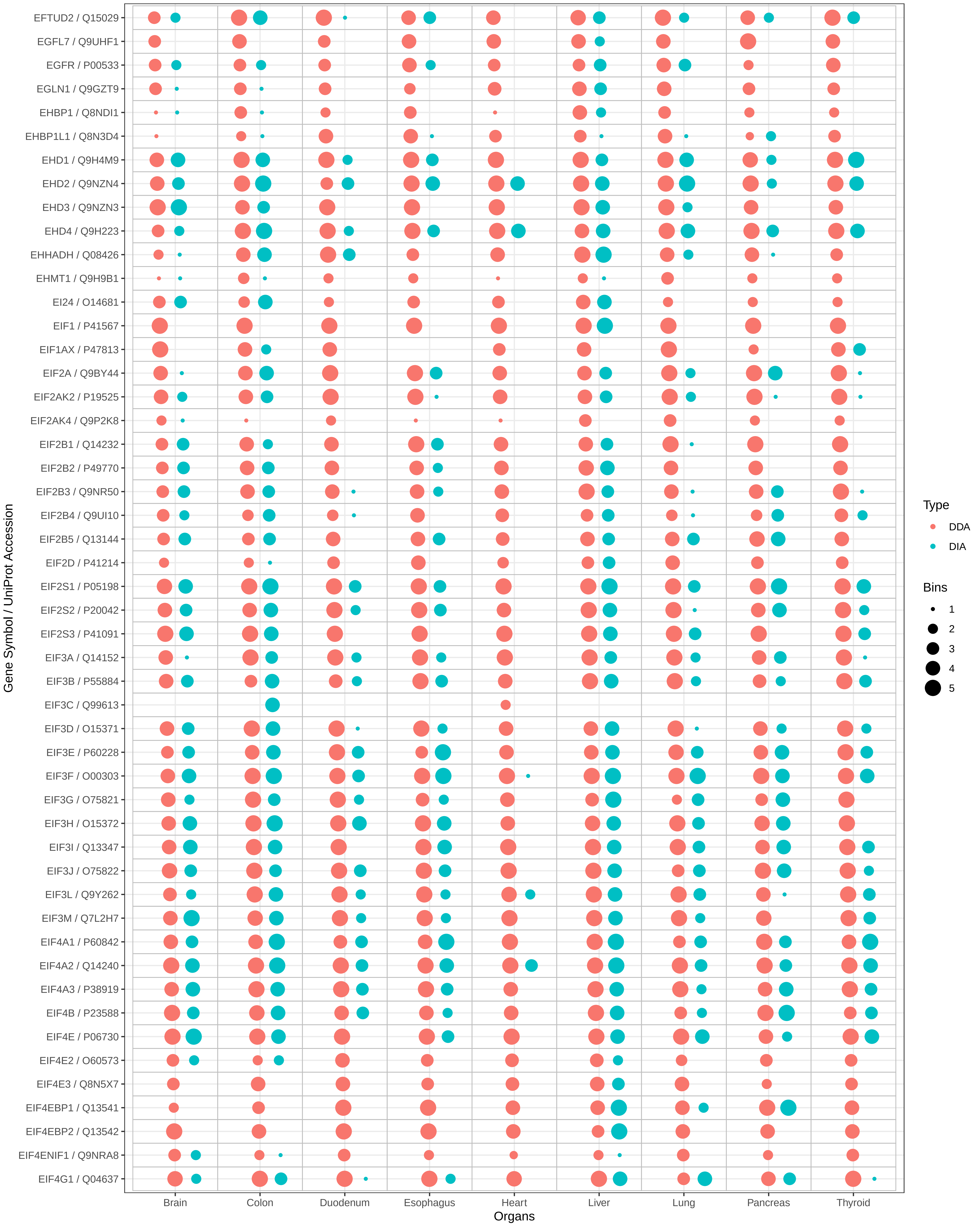

EFTUD2 / Q15029
EGFL7 / Q9UHF1
EGFR / P00533
EGLN1 / Q9GZT9
EHBP1 / Q8NDI1
EHBP1L1 / Q8N3D4
EHD1 / Q9H4M9
EHD2 / Q9NZN4
EHD3 / Q9NZN3
EHD4 / Q9H223
EHHADH / Q08426
EHMT1 / Q9H9B1
EI24 / O14681
EIF1 / P41567
EIF1AX / P47813
EIF2A / Q9BY44
EIF2AK2 / P19525
EIF2AK4 / Q9P2K8
EIF2B1 / Q14232
EIF2B2 / P49770
EIF2B3 / Q9NR50
EIF2B4 / Q9UI10
EIF2B5 / Q13144
EIF2D / P41214
EIF2S1 / P05198
EIF2S2 / P20042
EIF2S3 / P41091
EIF3A / Q14152
EIF3B / P55884
EIF3C / Q99613
EIF3D / O15371
EIF3E / P60228
EIF3F / O00303
EIF3G / O75821
EIF3H / O15372
EIF3I / Q13347
EIF3J / O75822
EIF3L / Q9Y262
EIF3M / Q7L2H7
EIF4A1 / P60842
EIF4A2 / Q14240
EIF4A3 / P38919
EIF4B / P23588
EIF4E / P06730
EIF4E2 / O60573
EIF4E3 / Q8N5X7
EIF4EBP1 / Q13541
EIF4EBP2 / Q13542
EIF4ENIF1 / Q9NRA8
EIF4G1 / Q04637
Gene Symbol / UniProt Accession
Brain
Colon
Duodenum
Esophagus
Heart
Liver
Lung
Pancreas
Thyroid
Organs
Type
DDA
DIA
Bins
1
2
3
4
5

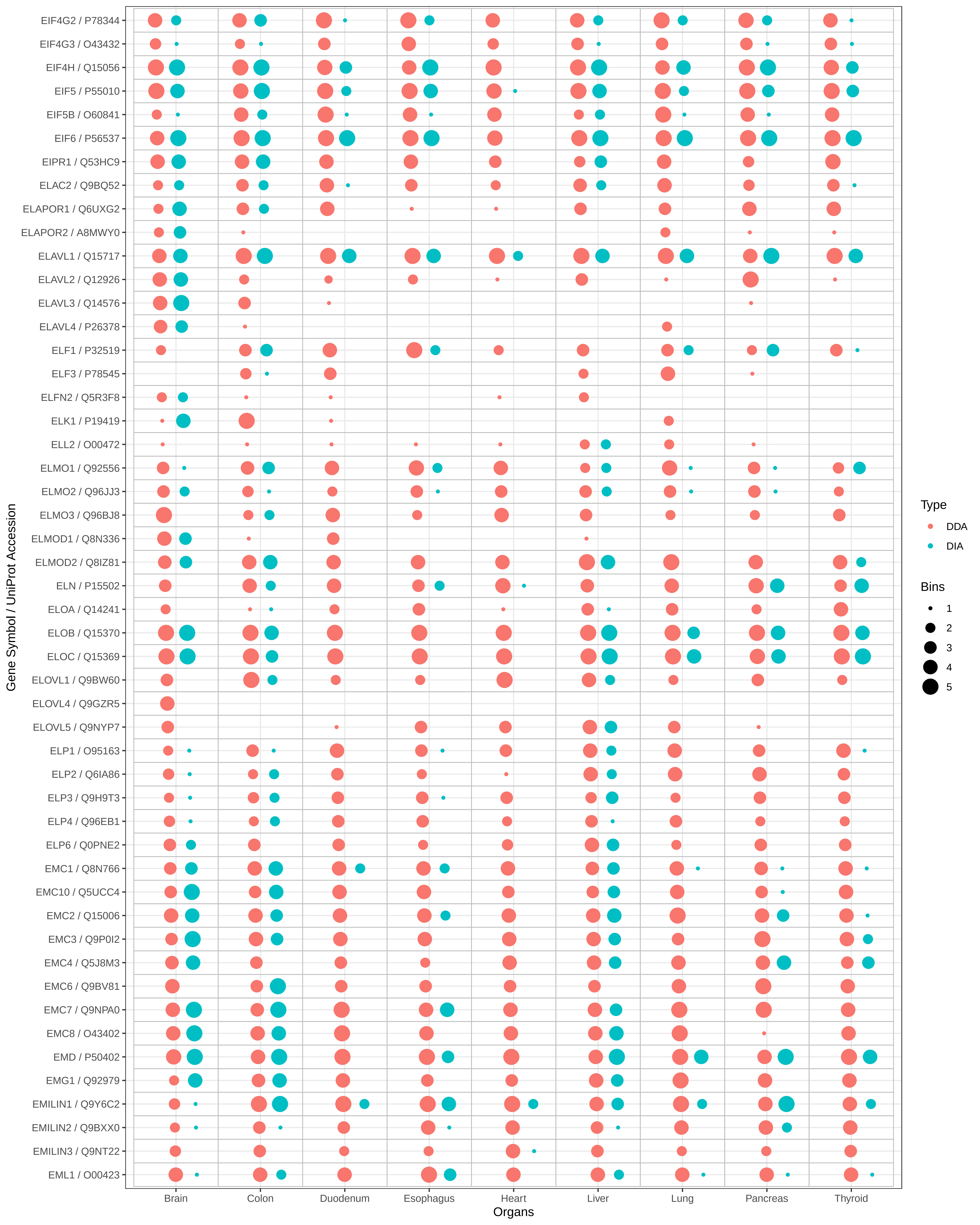

EIF4G2 / P78344
EIF4G3 / O43432
EIF4H / Q15056
EIF5 / P55010
EIF5B / O60841
EIF6 / P56537
EIPR1 / Q53HC9
ELAC2 / Q9BQ52
ELAPOR1 / Q6UXG2
ELAPOR2 / A8MWY0
ELAVL1 / Q15717
ELAVL2 / Q12926
ELAVL3 / Q14576
ELAVL4 / P26378
ELF1 / P32519
ELF3 / P78545
ELFN2 / Q5R3F8
ELK1 / P19419
ELL2 / O00472
ELMO1 / Q92556
ELMO2 / Q96JJ3
ELMO3 / Q96BJ8
ELMOD1 / Q8N336
ELMOD2 / Q8IZ81
ELN / P15502
ELOA / Q14241
ELOB / Q15370
ELOC / Q15369
ELOVL1 / Q9BW60
ELOVL4 / Q9GZR5
ELOVL5 / Q9NYP7
ELP1 / O95163
ELP2 / Q6IA86
ELP3 / Q9H9T3
ELP4 / Q96EB1
ELP6 / Q0PNE2
EMC1 / Q8N766
EMC10 / Q5UCC4
EMC2 / Q15006
EMC3 / Q9P0I2
EMC4 / Q5J8M3
EMC6 / Q9BV81
EMC7 / Q9NPA0
EMC8 / O43402
EMD / P50402
EMG1 / Q92979
EMILIN1 / Q9Y6C2
EMILIN2 / Q9BXX0
EMILIN3 / Q9NT22
EML1 / O00423
Gene Symbol / UniProt Accession
Brain
Colon
Duodenum
Esophagus
Heart
Liver
Lung
Pancreas
Thyroid
Organs
Type
DDA
DIA
Bins
1
2
3
4
5

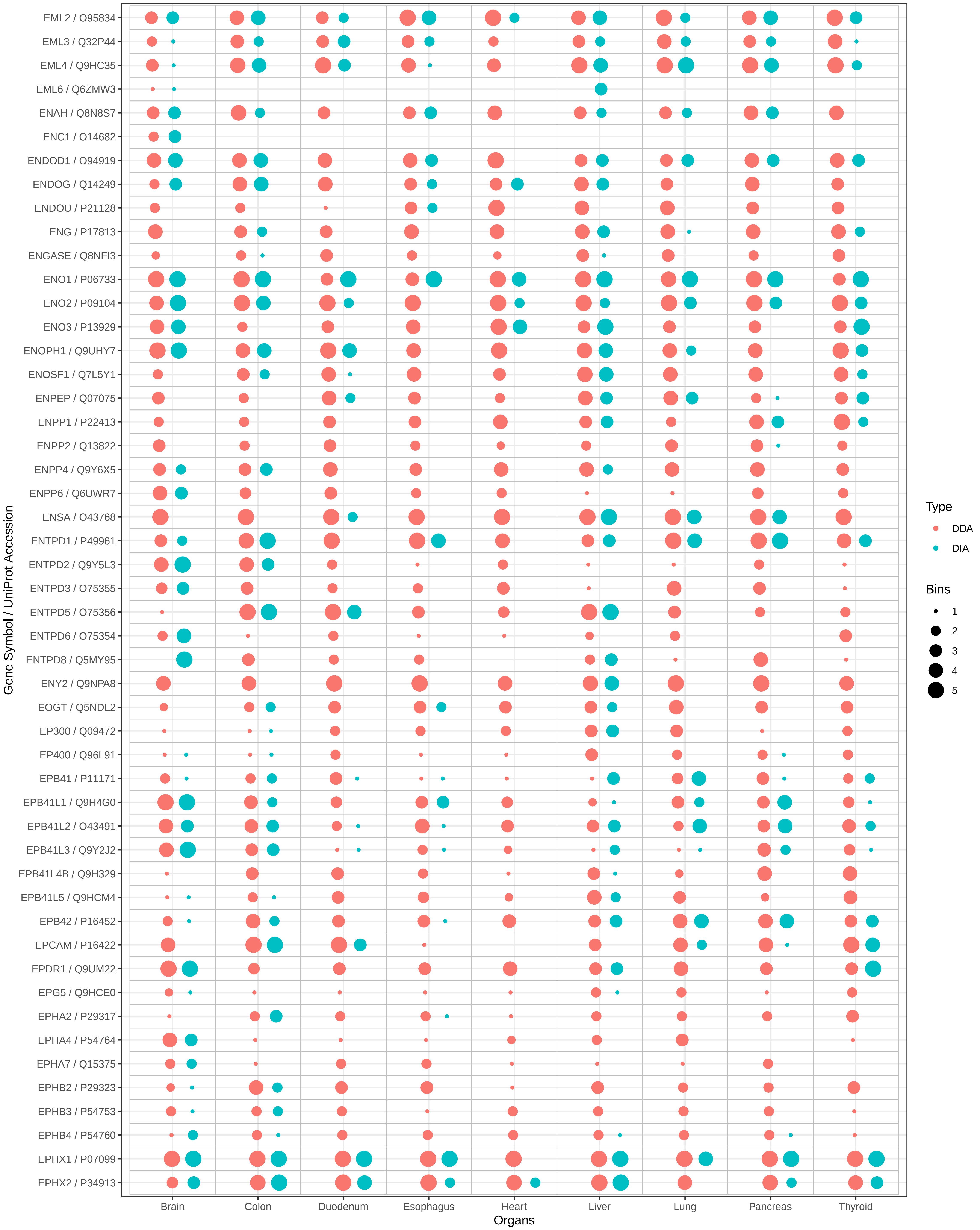

EML2 / O95834
EML3 / Q32P44
EML4 / Q9HC35
EML6 / Q6ZMW3
ENAH / Q8N8S7
ENC1 / O14682
ENDOD1 / O94919
ENDOG / Q14249
ENDOU / P21128
ENG / P17813
ENGASE / Q8NFI3
ENO1 / P06733
ENO2 / P09104
ENO3 / P13929
ENOPH1 / Q9UHY7
ENOSF1 / Q7L5Y1
ENPEP / Q07075
ENPP1 / P22413
ENPP2 / Q13822
ENPP4 / Q9Y6X5
ENPP6 / Q6UWR7
ENSA / O43768
ENTPD1 / P49961
ENTPD2 / Q9Y5L3
ENTPD3 / O75355
ENTPD5 / O75356
ENTPD6 / O75354
ENTPD8 / Q5MY95
ENY2 / Q9NPA8
EOGT / Q5NDL2
EP300 / Q09472
EP400 / Q96L91
EPB41 / P11171
EPB41L1 / Q9H4G0
EPB41L2 / O43491
EPB41L3 / Q9Y2J2
EPB41L4B / Q9H329
EPB41L5 / Q9HCM4
EPB42 / P16452
EPCAM / P16422
EPDR1 / Q9UM22
EPG5 / Q9HCE0
EPHA2 / P29317
EPHA4 / P54764
EPHA7 / Q15375
EPHB2 / P29323
EPHB3 / P54753
EPHB4 / P54760
EPHX1 / P07099
EPHX2 / P34913
Gene Symbol / UniProt Accession
Brain
Colon
Duodenum
Esophagus
Heart
Liver
Lung
Pancreas
Thyroid
Organs
Type
DDA
DIA
Bins
1
2
3
4
5

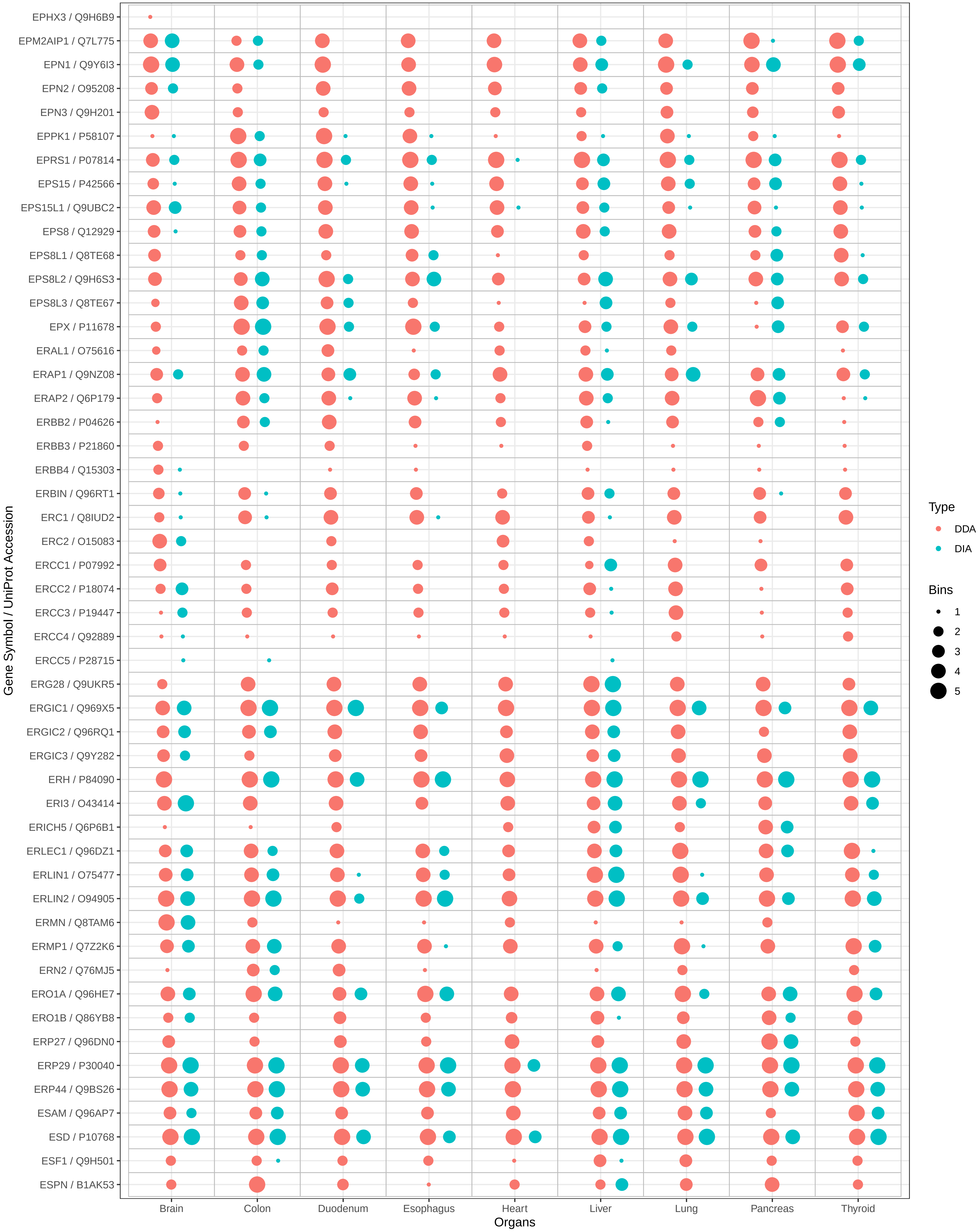
EPHX3 / Q9H6B9
EPM2AIP1 / Q7L775
EPN1 / Q9Y6I3
EPN2 / O95208
EPN3 / Q9H201
EPPK1 / P58107
EPRS1 / P07814
EPS15 / P42566
EPS15L1 / Q9UBC2
EPS8 / Q12929
EPS8L1 / Q8TE68
EPS8L2 / Q9H6S3
EPS8L3 / Q8TE67
EPX / P11678
ERAL1 / O75616
ERAP1 / Q9NZ08
ERAP2 / Q6P179
ERBB2 / P04626
ERBB3 / P21860
ERBB4 / Q15303
ERBIN / Q96RT1
ERC1 / Q8IUD2
ERC2 / O15083
ERCC1 / P07992
ERCC2 / P18074
ERCC3 / P19447
ERCC4 / Q92889
ERCC5 / P28715
ERG28 / Q9UKR5
ERGIC1 / Q969X5
ERGIC2 / Q96RQ1
ERGIC3 / Q9Y282
ERH / P84090
ERI3 / O43414
ERICH5 / Q6P6B1
ERLEC1 / Q96DZ1
ERLIN1 / O75477
ERLIN2 / O94905
ERMN / Q8TAM6
ERMP1 / Q7Z2K6
ERN2 / Q76MJ5
ERO1A / Q96HE7
ERO1B / Q86YB8
ERP27 / Q96DN0
ERP29 / P30040
ERP44 / Q9BS26
ESAM / Q96AP7
ESD / P10768
ESF1 / Q9H501
ESPN / B1AK53
Gene Symbol / UniProt Accession
Brain
Colon
Duodenum
Esophagus
Heart
Liver
Lung
Pancreas
Thyroid
Organs
Type
DDA
DIA
Bins
1
2
3
4
5

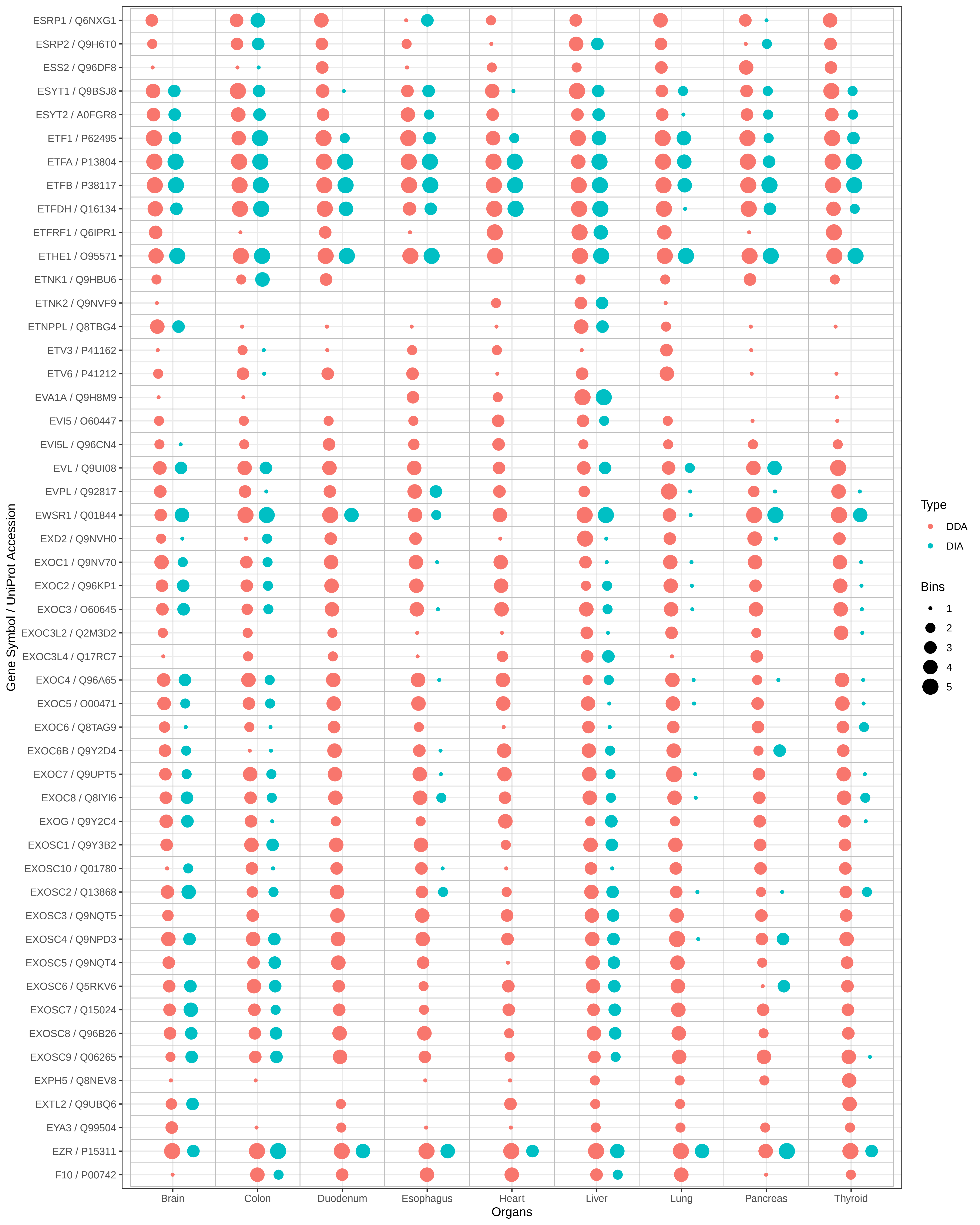

ESRP1 / Q6NXG1
ESRP2 / Q9H6T0
ESS2 / Q96DF8
ESYT1 / Q9BSJ8
ESYT2 / A0FGR8
ETF1 / P62495
ETFA / P13804
ETFB / P38117
ETFDH / Q16134
ETFRF1 / Q6IPR1
ETHE1 / O95571
ETNK1 / Q9HBU6
ETNK2 / Q9NVF9
ETNPPL / Q8TBG4
ETV3 / P41162
ETV6 / P41212
EVA1A / Q9H8M9
EVI5 / O60447
EVI5L / Q96CN4
EVL / Q9UI08
EVPL / Q92817
EWSR1 / Q01844
EXD2 / Q9NVH0
EXOC1 / Q9NV70
EXOC2 / Q96KP1
EXOC3 / O60645
EXOC3L2 / Q2M3D2
EXOC3L4 / Q17RC7
EXOC4 / Q96A65
EXOC5 / O00471
EXOC6 / Q8TAG9
EXOC6B / Q9Y2D4
EXOC7 / Q9UPT5
EXOC8 / Q8IYI6
EXOG / Q9Y2C4
EXOSC1 / Q9Y3B2
EXOSC10 / Q01780
EXOSC2 / Q13868
EXOSC3 / Q9NQT5
EXOSC4 / Q9NPD3
EXOSC5 / Q9NQT4
EXOSC6 / Q5RKV6
EXOSC7 / Q15024
EXOSC8 / Q96B26
EXOSC9 / Q06265
EXPH5 / Q8NEV8
EXTL2 / Q9UBQ6
EYA3 / Q99504
EZR / P15311
F10 / P00742
Gene Symbol / UniProt Accession
Brain
Colon
Duodenum
Esophagus
Heart
Liver
Lung
Pancreas
Thyroid
Organs
Type
DDA
DIA
Bins
1
2
3
4
5

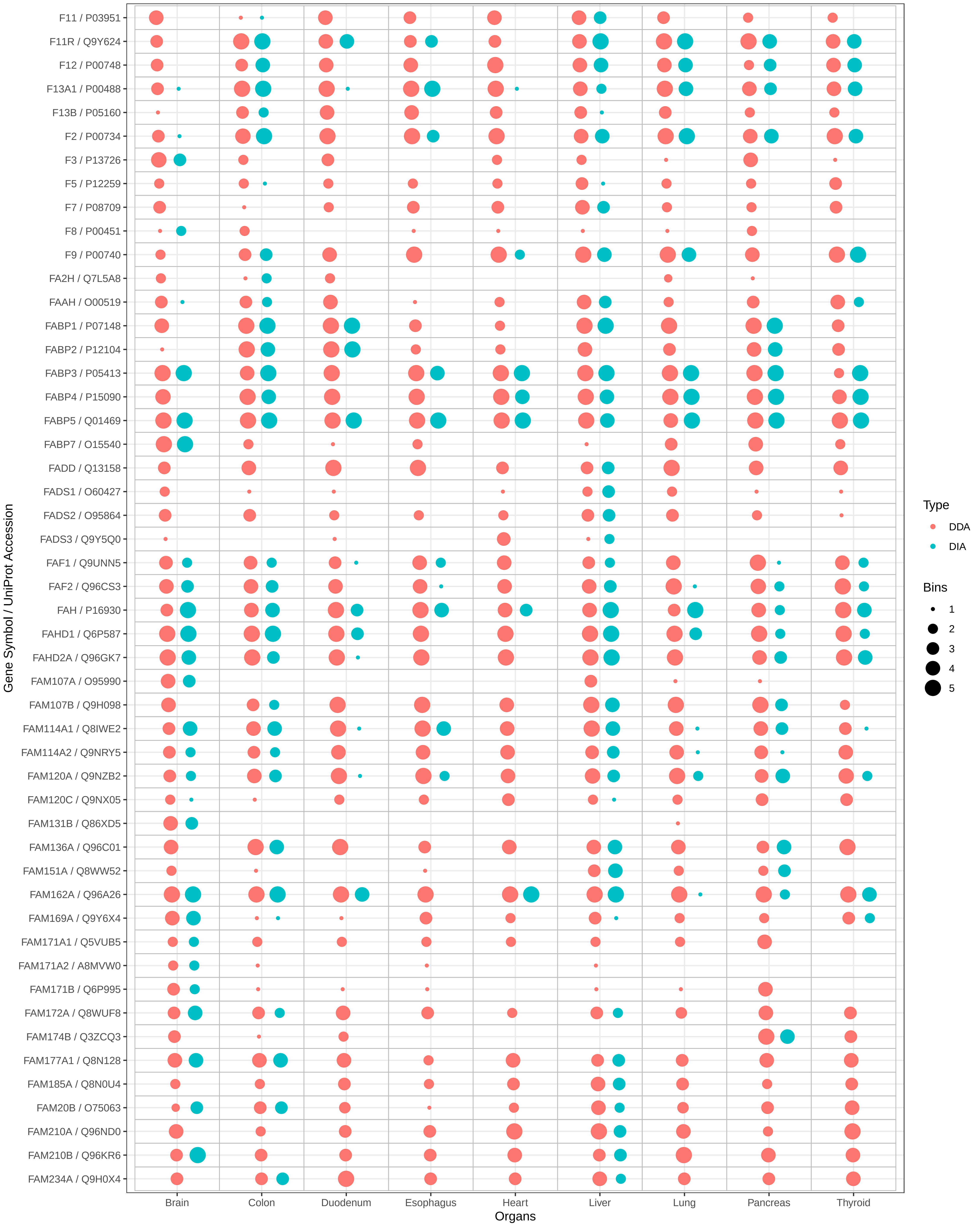

F11 / P03951
F11R / Q9Y624
F12 / P00748
F13A1 / P00488
F13B / P05160
F2 / P00734
F3 / P13726
F5 / P12259
F7 / P08709
F8 / P00451
F9 / P00740
FA2H / Q7L5A8
FAAH / O00519
FABP1 / P07148
FABP2 / P12104
FABP3 / P05413
FABP4 / P15090
FABP5 / Q01469
FABP7 / O15540
FADD / Q13158
FADS1 / O60427
FADS2 / O95864
FADS3 / Q9Y5Q0
FAF1 / Q9UNN5
FAF2 / Q96CS3
FAH / P16930
FAHD1 / Q6P587
FAHD2A / Q96GK7
FAM107A / O95990
FAM107B / Q9H098
FAM114A1 / Q8IWE2
FAM114A2 / Q9NRY5
FAM120A / Q9NZB2
FAM120C / Q9NX05
FAM131B / Q86XD5
FAM136A / Q96C01
FAM151A / Q8WW52
FAM162A / Q96A26
FAM169A / Q9Y6X4
FAM171A1 / Q5VUB5
FAM171A2 / A8MVW0
FAM171B / Q6P995
FAM172A / Q8WUF8
FAM174B / Q3ZCQ3
FAM177A1 / Q8N128
FAM185A / Q8N0U4
FAM20B / O75063
FAM210A / Q96ND0
FAM210B / Q96KR6
FAM234A / Q9H0X4
Gene Symbol / UniProt Accession
Brain
Colon
Duodenum
Esophagus
Heart
Liver
Lung
Pancreas
Thyroid
Organs
Type
DDA
DIA
Bins
1
2
3
4
5

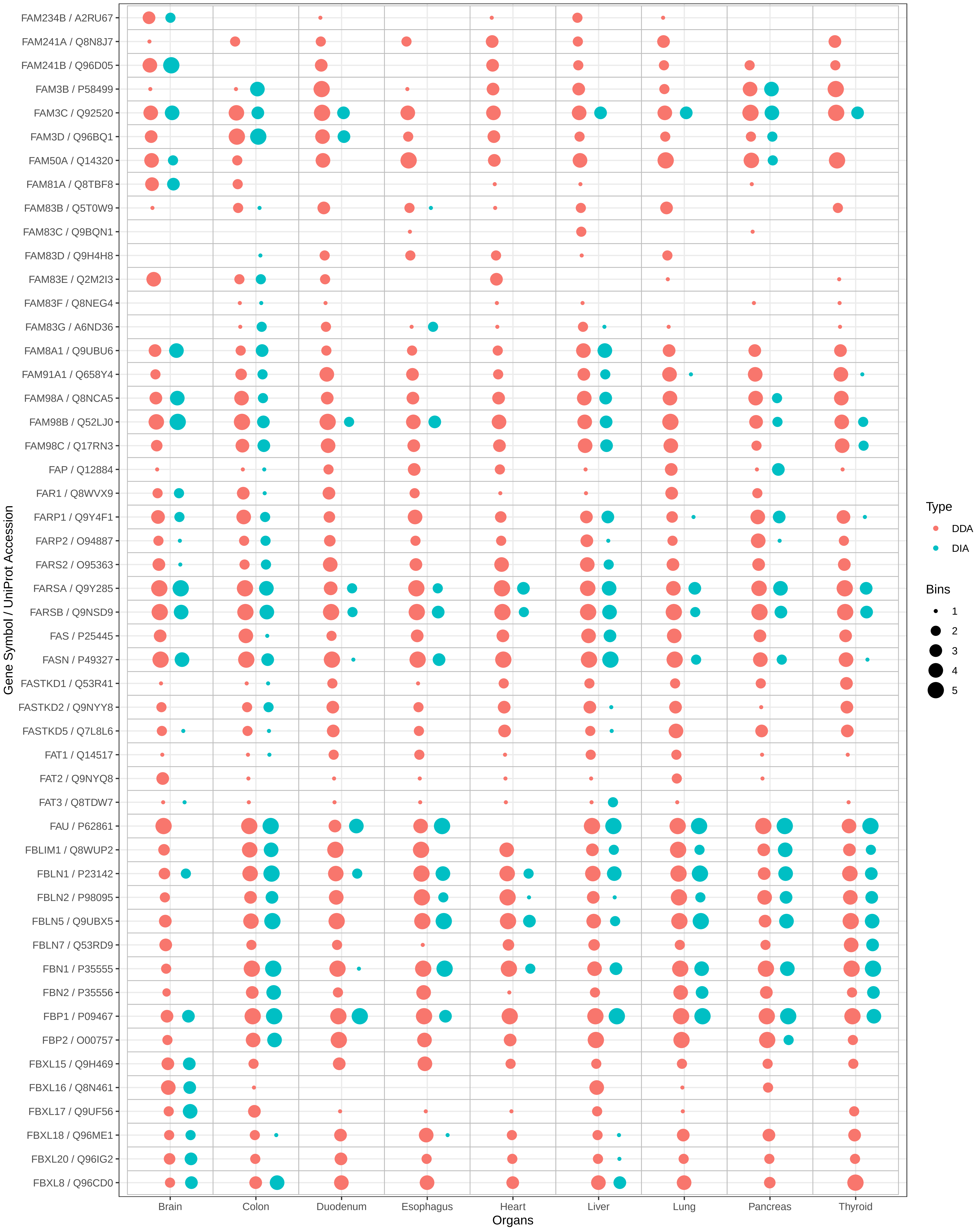

FAM234B / A2RU67
FAM241A / Q8N8J7
FAM241B / Q96D05
FAM3B / P58499
FAM3C / Q92520
FAM3D / Q96BQ1
FAM50A / Q14320
FAM81A / Q8TBF8
FAM83B / Q5T0W9
FAM83C / Q9BQN1
FAM83D / Q9H4H8
FAM83E / Q2M2I3
FAM83F / Q8NEG4
FAM83G / A6ND36
FAM8A1 / Q9UBU6
FAM91A1 / Q658Y4
FAM98A / Q8NCA5
FAM98B / Q52LJ0
FAM98C / Q17RN3
FAP / Q12884
FAR1 / Q8WVX9
FARP1 / Q9Y4F1
FARP2 / O94887
FARS2 / O95363
FARSA / Q9Y285
FARSB / Q9NSD9
FAS / P25445
FASN / P49327
FASTKD1 / Q53R41
FASTKD2 / Q9NYY8
FASTKD5 / Q7L8L6
FAT1 / Q14517
FAT2 / Q9NYQ8
FAT3 / Q8TDW7
FAU / P62861
FBLIM1 / Q8WUP2
FBLN1 / P23142
FBLN2 / P98095
FBLN5 / Q9UBX5
FBLN7 / Q53RD9
FBN1 / P35555
FBN2 / P35556
FBP1 / P09467
FBP2 / O00757
FBXL15 / Q9H469
FBXL16 / Q8N461
FBXL17 / Q9UF56
FBXL18 / Q96ME1
FBXL20 / Q96IG2
FBXL8 / Q96CD0
Gene Symbol / UniProt Accession
Brain
Colon
Duodenum
Esophagus
Heart
Liver
Lung
Pancreas
Thyroid
Organs
Type
DDA
DIA
Bins
1
2
3
4
5

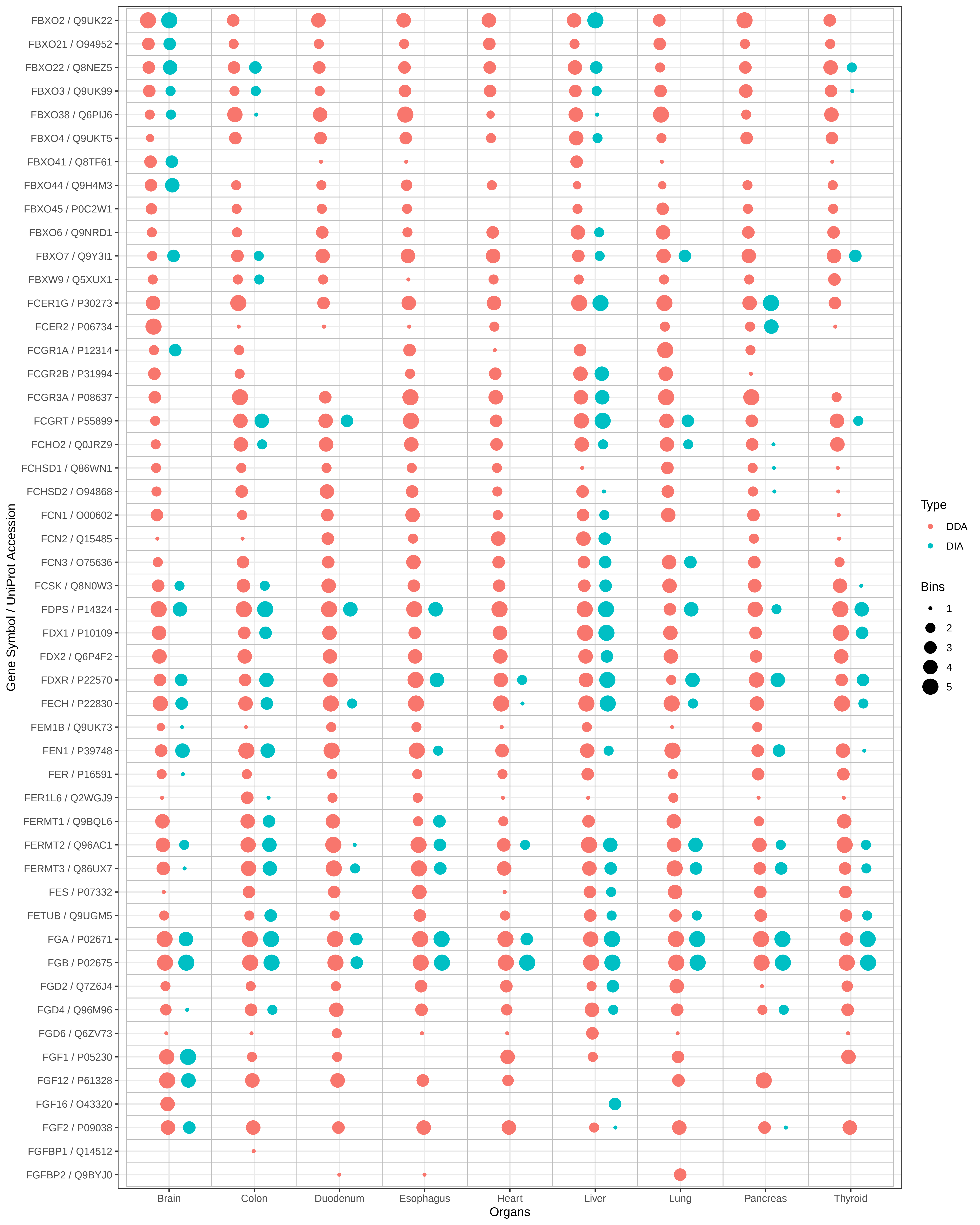

FBXO2 / Q9UK22
FBXO21 / O94952
FBXO22 / Q8NEZ5
FBXO3 / Q9UK99
FBXO38 / Q6PIJ6
FBXO4 / Q9UKT5
FBXO41 / Q8TF61
FBXO44 / Q9H4M3
FBXO45 / P0C2W1
FBXO6 / Q9NRD1
FBXO7 / Q9Y3I1
FBXW9 / Q5XUX1
FCER1G / P30273
FCER2 / P06734
FCGR1A / P12314
FCGR2B / P31994
FCGR3A / P08637
FCGRT / P55899
FCHO2 / Q0JRZ9
FCHSD1 / Q86WN1
FCHSD2 / O94868
FCN1 / O00602
FCN2 / Q15485
FCN3 / O75636
FCSK / Q8N0W3
FDPS / P14324
FDX1 / P10109
FDX2 / Q6P4F2
FDXR / P22570
FECH / P22830
FEM1B / Q9UK73
FEN1 / P39748
FER / P16591
FER1L6 / Q2WGJ9
FERMT1 / Q9BQL6
FERMT2 / Q96AC1
FERMT3 / Q86UX7
FES / P07332
FETUB / Q9UGM5
FGA / P02671
FGB / P02675
FGD2 / Q7Z6J4
FGD4 / Q96M96
FGD6 / Q6ZV73
FGF1 / P05230
FGF12 / P61328
FGF16 / O43320
FGF2 / P09038
FGFBP1 / Q14512
FGFBP2 / Q9BYJ0
Gene Symbol / UniProt Accession
Brain
Colon
Duodenum
Esophagus
Heart
Liver
Lung
Pancreas
Thyroid
Organs
Type
DDA
DIA
Bins
1
2
3
4
5

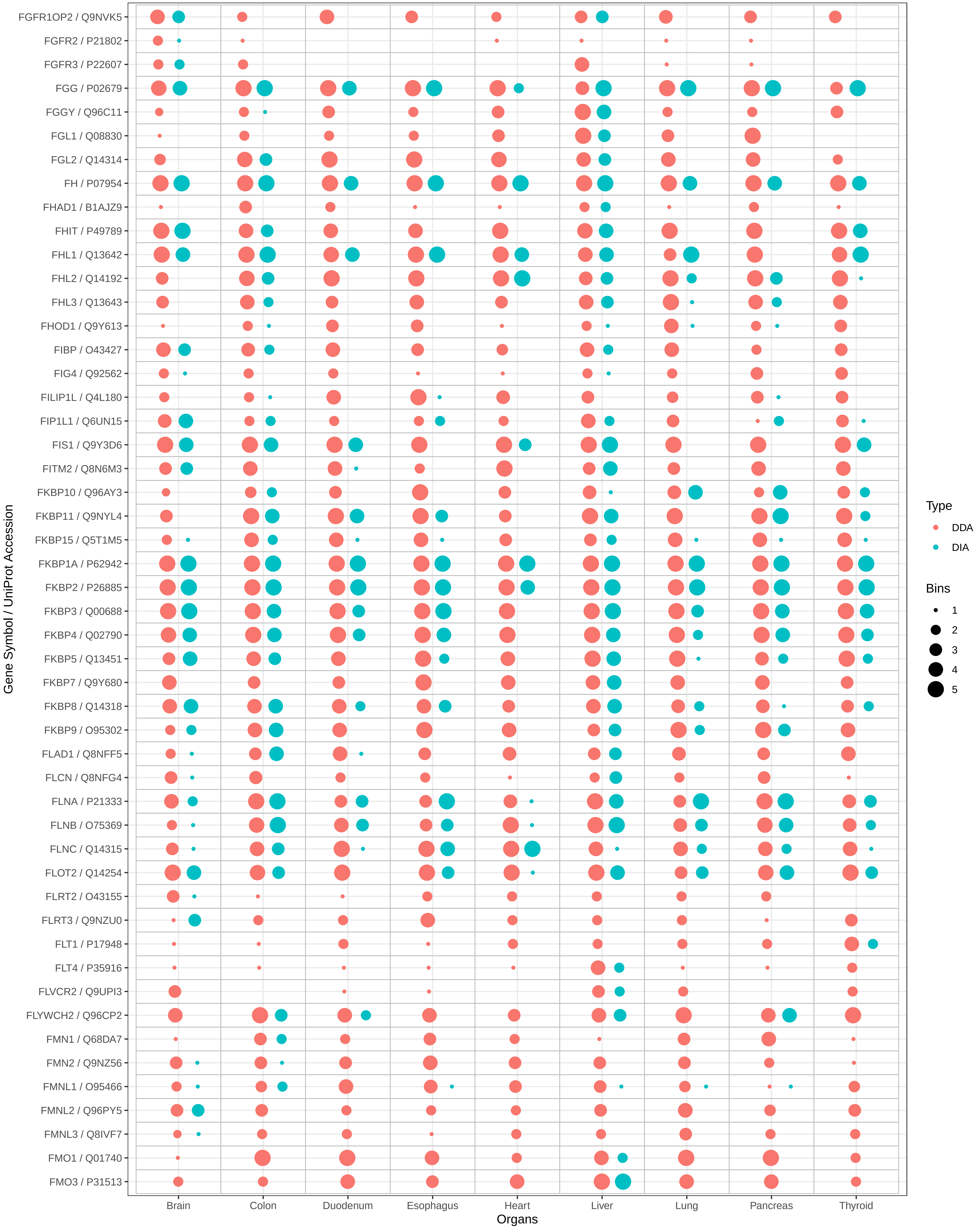

FGFR1OP2 / Q9NVK5
FGFR2 / P21802
FGFR3 / P22607
FGG / P02679
FGGY / Q96C11
FGL1 / Q08830
FGL2 / Q14314
FH / P07954
FHAD1 / B1AJZ9
FHIT / P49789
FHL1 / Q13642
FHL2 / Q14192
FHL3 / Q13643
FHOD1 / Q9Y613
FIBP / O43427
FIG4 / Q92562
FILIP1L / Q4L180
FIP1L1 / Q6UN15
FIS1 / Q9Y3D6
FITM2 / Q8N6M3
FKBP10 / Q96AY3
FKBP11 / Q9NYL4
FKBP15 / Q5T1M5
FKBP1A / P62942
FKBP2 / P26885
FKBP3 / Q00688
FKBP4 / Q02790
FKBP5 / Q13451
FKBP7 / Q9Y680
FKBP8 / Q14318
FKBP9 / O95302
FLAD1 / Q8NFF5
FLCN / Q8NFG4
FLNA / P21333
FLNB / O75369
FLNC / Q14315
FLOT2 / Q14254
FLRT2 / O43155
FLRT3 / Q9NZU0
FLT1 / P17948
FLT4 / P35916
FLVCR2 / Q9UPI3
FLYWCH2 / Q96CP2
FMN1 / Q68DA7
FMN2 / Q9NZ56
FMNL1 / O95466
FMNL2 / Q96PY5
FMNL3 / Q8IVF7
FMO1 / Q01740
FMO3 / P31513
Gene Symbol / UniProt Accession
Brain
Colon
Duodenum
Esophagus
Heart
Liver
Lung
Pancreas
Thyroid
Organs
Type
DDA
DIA
Bins
1
2
3
4
5

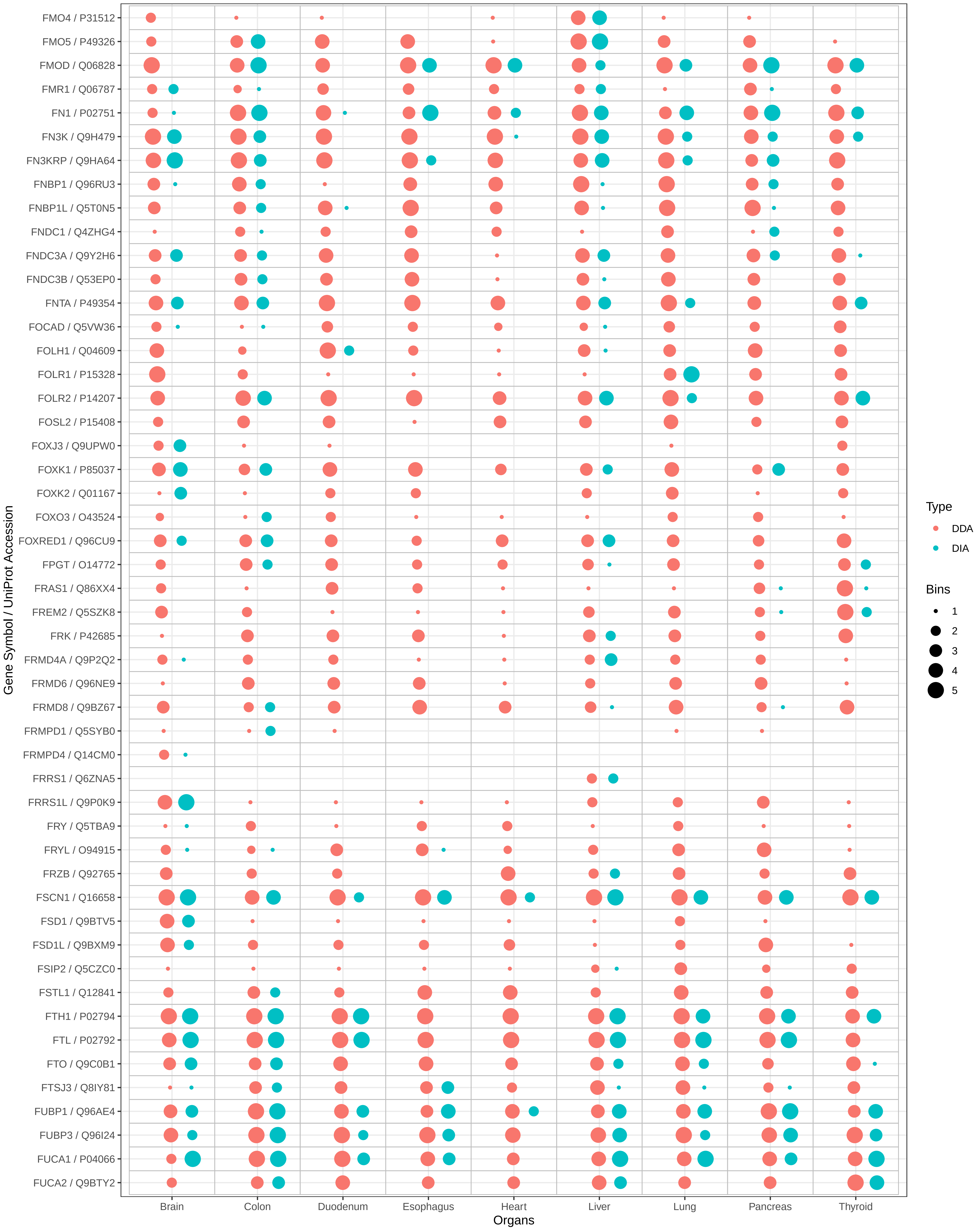

FMO4 / P31512
FMO5 / P49326
FMOD / Q06828
FMR1 / Q06787
FN1 / P02751
FN3K / Q9H479
FN3KRP / Q9HA64
FNBP1 / Q96RU3
FNBP1L / Q5T0N5
FNDC1 / Q4ZHG4
FNDC3A / Q9Y2H6
FNDC3B / Q53EP0
FNTA / P49354
FOCAD / Q5VW36
FOLH1 / Q04609
FOLR1 / P15328
FOLR2 / P14207
FOSL2 / P15408
FOXJ3 / Q9UPW0
FOXK1 / P85037
FOXK2 / Q01167
FOXO3 / O43524
FOXRED1 / Q96CU9
FPGT / O14772
FRAS1 / Q86XX4
FREM2 / Q5SZK8
FRK / P42685
FRMD4A / Q9P2Q2
FRMD6 / Q96NE9
FRMD8 / Q9BZ67
FRMPD1 / Q5SYB0
FRMPD4 / Q14CM0
FRRS1 / Q6ZNA5
FRRS1L / Q9P0K9
FRY / Q5TBA9
FRYL / O94915
FRZB / Q92765
FSCN1 / Q16658
FSD1 / Q9BTV5
FSD1L / Q9BXM9
FSIP2 / Q5CZC0
FSTL1 / Q12841
FTH1 / P02794
FTL / P02792
FTO / Q9C0B1
FTSJ3 / Q8IY81
FUBP1 / Q96AE4
FUBP3 / Q96I24
FUCA1 / P04066
FUCA2 / Q9BTY2
Gene Symbol / UniProt Accession
Brain
Colon
Duodenum
Esophagus
Heart
Liver
Lung
Pancreas
Thyroid
Organs
Type
DDA
DIA
Bins
1
2
3
4
5

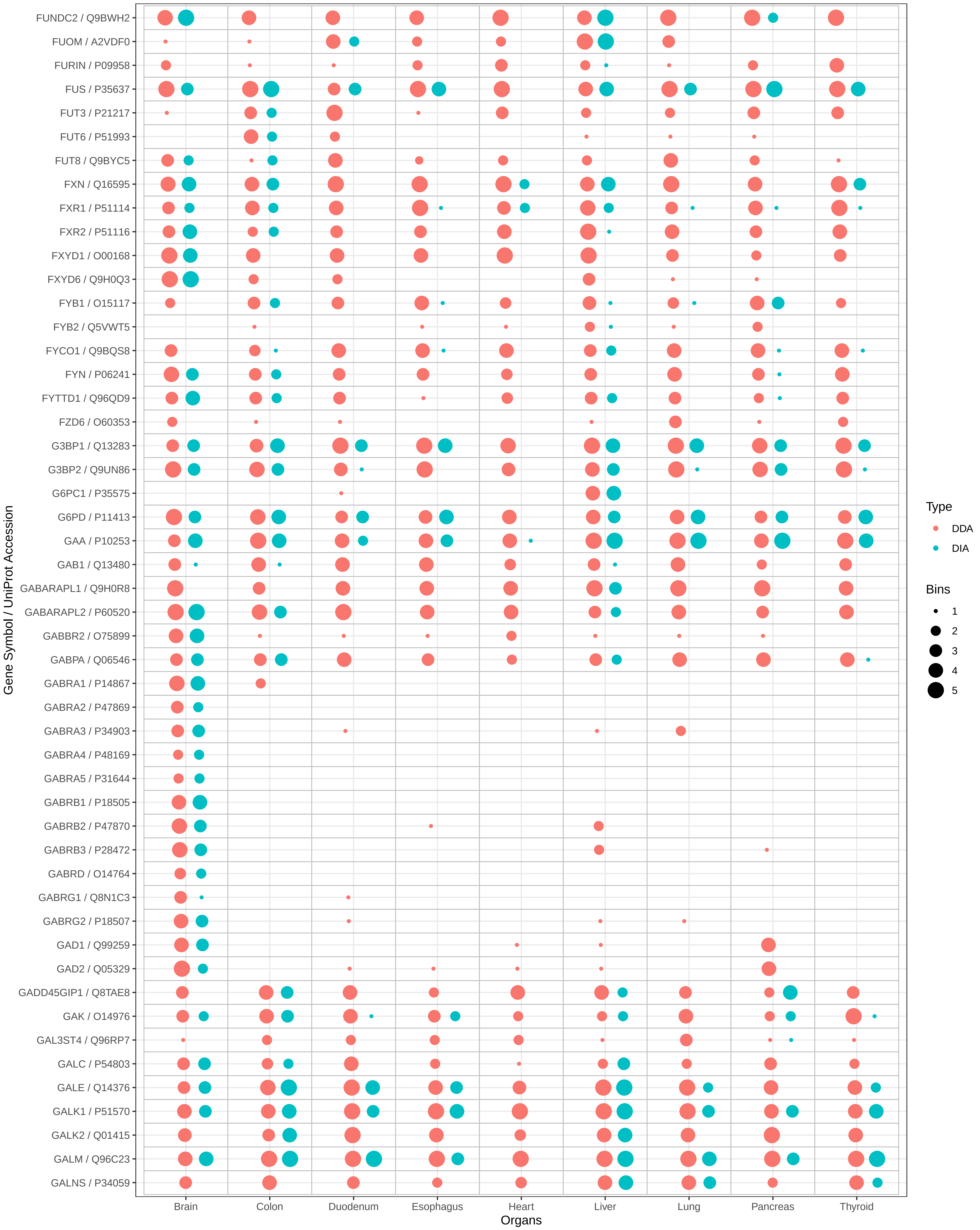
FUNDC2 / Q9BWH2
FUOM / A2VDF0
FURIN / P09958
FUS / P35637
FUT3 / P21217
FUT6 / P51993
FUT8 / Q9BYC5
FXN / Q16595
FXR1 / P51114
FXR2 / P51116
FXYD1 / O00168
FXYD6 / Q9H0Q3
FYB1 / O15117
FYB2 / Q5VWT5
FYCO1 / Q9BQS8
FYN / P06241
FYTTD1 / Q96QD9
FZD6 / O60353
G3BP1 / Q13283
G3BP2 / Q9UN86
G6PC1 / P35575
G6PD / P11413
GAA / P10253
GAB1 / Q13480
GABARAPL1 / Q9H0R8
GABARAPL2 / P60520
GABBR2 / O75899
GABPA / Q06546
GABRA1 / P14867
GABRA2 / P47869
GABRA3 / P34903
GABRA4 / P48169
GABRA5 / P31644
GABRB1 / P18505
GABRB2 / P47870
GABRB3 / P28472
GABRD / O14764
GABRG1 / Q8N1C3
GABRG2 / P18507
GAD1 / Q99259
GAD2 / Q05329
GADD45GIP1 / Q8TAE8
GAK / O14976
GAL3ST4 / Q96RP7
GALC / P54803
GALE / Q14376
GALK1 / P51570
GALK2 / Q01415
GALM / Q96C23
GALNS / P34059
Gene Symbol / UniProt Accession
Brain
Colon
Duodenum
Esophagus
Heart
Liver
Lung
Pancreas
Thyroid
Organs
Type
DDA
DIA
Bins
1
2
3
4
5

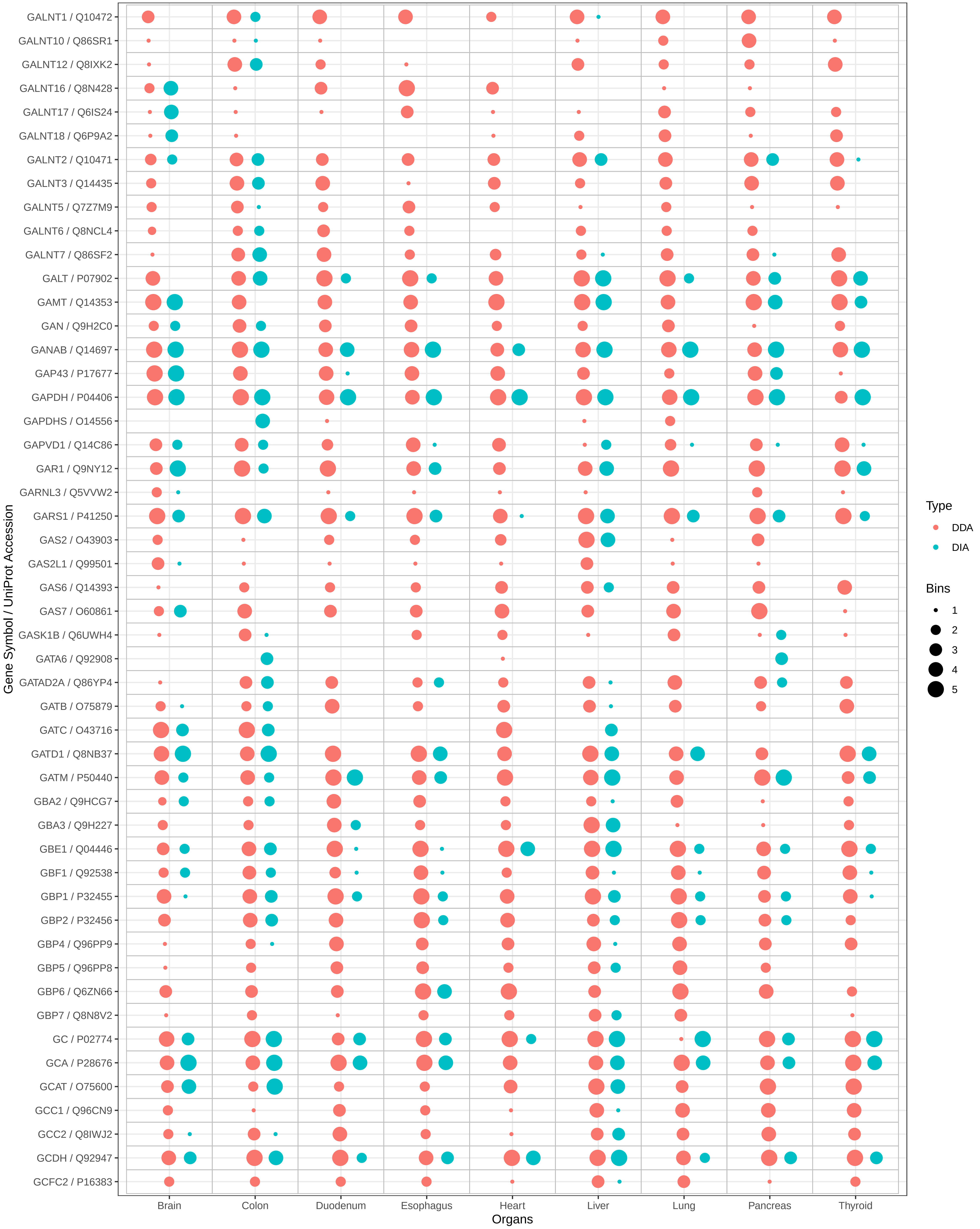

GALNT1 / Q10472
GALNT10 / Q86SR1
GALNT12 / Q8IXK2
GALNT16 / Q8N428
GALNT17 / Q6IS24
GALNT18 / Q6P9A2
GALNT2 / Q10471
GALNT3 / Q14435
GALNT5 / Q7Z7M9
GALNT6 / Q8NCL4
GALNT7 / Q86SF2
GALT / P07902
GAMT / Q14353
GAN / Q9H2C0
GANAB / Q14697
GAP43 / P17677
GAPDH / P04406
GAPDHS / O14556
GAPVD1 / Q14C86
GAR1 / Q9NY12
GARNL3 / Q5VVW2
GARS1 / P41250
GAS2 / O43903
GAS2L1 / Q99501
GAS6 / Q14393
GAS7 / O60861
GASK1B / Q6UWH4
GATA6 / Q92908
GATAD2A / Q86YP4
GATB / O75879
GATC / O43716
GATD1 / Q8NB37
GATM / P50440
GBA2 / Q9HCG7
GBA3 / Q9H227
GBE1 / Q04446
GBF1 / Q92538
GBP1 / P32455
GBP2 / P32456
GBP4 / Q96PP9
GBP5 / Q96PP8
GBP6 / Q6ZN66
GBP7 / Q8N8V2
GC / P02774
GCA / P28676
GCAT / O75600
GCC1 / Q96CN9
GCC2 / Q8IWJ2
GCDH / Q92947
GCFC2 / P16383
Gene Symbol / UniProt Accession
Brain
Colon
Duodenum
Esophagus
Heart
Liver
Lung
Pancreas
Thyroid
Organs
Type
DDA
DIA
Bins
1
2
3
4
5

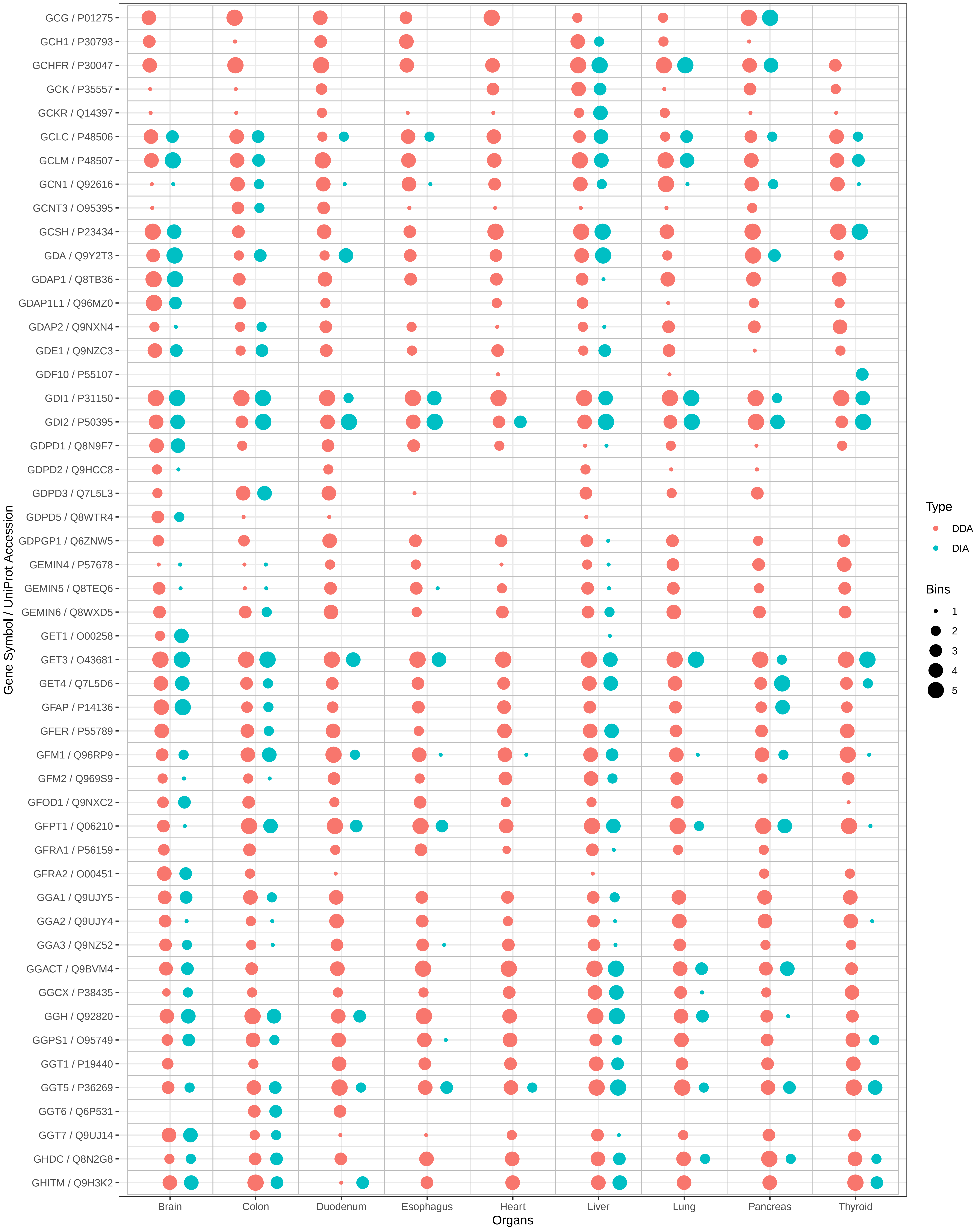
GCG / P01275
GCH1 / P30793
GCHFR / P30047
GCK / P35557
GCKR / Q14397
GCLC / P48506
GCLM / P48507
GCN1 / Q92616
GCNT3 / O95395
GCSH / P23434
GDA / Q9Y2T3
GDAP1 / Q8TB36
GDAP1L1 / Q96MZ0
GDAP2 / Q9NXN4
GDE1 / Q9NZC3
GDF10 / P55107
GDI1 / P31150
GDI2 / P50395
GDPD1 / Q8N9F7
GDPD2 / Q9HCC8
GDPD3 / Q7L5L3
GDPD5 / Q8WTR4
GDPGP1 / Q6ZNW5
GEMIN4 / P57678
GEMIN5 / Q8TEQ6
GEMIN6 / Q8WXD5
GET1 / O00258
GET3 / O43681
GET4 / Q7L5D6
GFAP / P14136
GFER / P55789
GFM1 / Q96RP9
GFM2 / Q969S9
GFOD1 / Q9NXC2
GFPT1 / Q06210
GFRA1 / P56159
GFRA2 / O00451
GGA1 / Q9UJY5
GGA2 / Q9UJY4
GGA3 / Q9NZ52
GGACT / Q9BVM4
GGCX / P38435
GGH / Q92820
GGPS1 / O95749
GGT1 / P19440
GGT5 / P36269
GGT6 / Q6P531
GGT7 / Q9UJ14
GHDC / Q8N2G8
GHITM / Q9H3K2
Gene Symbol / UniProt Accession
Brain
Colon
Duodenum
Esophagus
Heart
Liver
Lung
Pancreas
Thyroid
Organs
Type
DDA
DIA
Bins
1
2
3
4
5

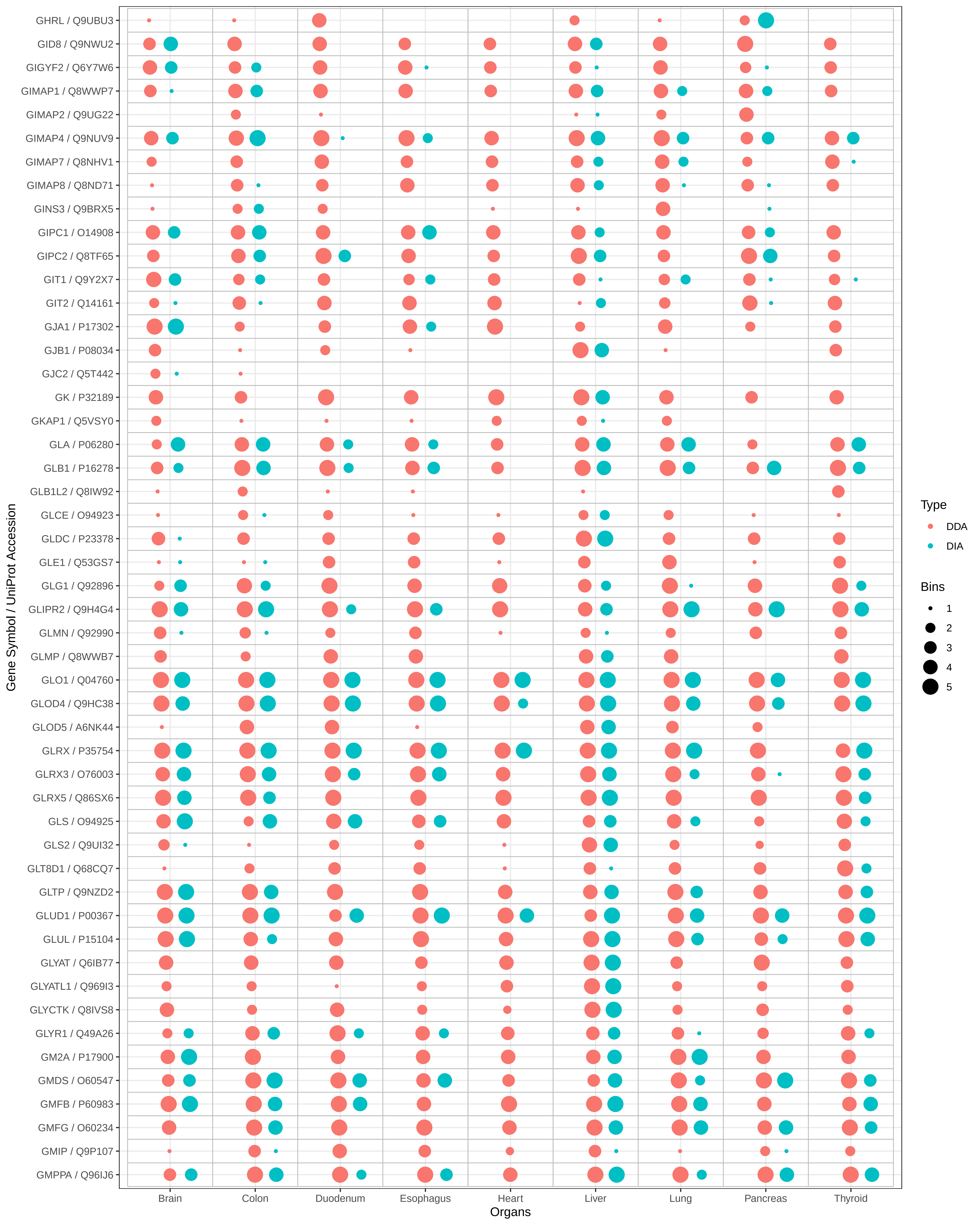

GHRL / Q9UBU3
GID8 / Q9NWU2
GIGYF2 / Q6Y7W6
GIMAP1 / Q8WWP7
GIMAP2 / Q9UG22
GIMAP4 / Q9NUV9
GIMAP7 / Q8NHV1
GIMAP8 / Q8ND71
GINS3 / Q9BRX5
GIPC1 / O14908
GIPC2 / Q8TF65
GIT1 / Q9Y2X7
GIT2 / Q14161
GJA1 / P17302
GJB1 / P08034
GJC2 / Q5T442
GK / P32189
GKAP1 / Q5VSY0
GLA / P06280
GLB1 / P16278
GLB1L2 / Q8IW92
GLCE / O94923
GLDC / P23378
GLE1 / Q53GS7
GLG1 / Q92896
GLIPR2 / Q9H4G4
GLMN / Q92990
GLMP / Q8WWB7
GLO1 / Q04760
GLOD4 / Q9HC38
GLOD5 / A6NK44
GLRX / P35754
GLRX3 / O76003
GLRX5 / Q86SX6
GLS / O94925
GLS2 / Q9UI32
GLT8D1 / Q68CQ7
GLTP / Q9NZD2
GLUD1 / P00367
GLUL / P15104
GLYAT / Q6IB77
GLYATL1 / Q969I3
GLYCTK / Q8IVS8
GLYR1 / Q49A26
GM2A / P17900
GMDS / O60547
GMFB / P60983
GMFG / O60234
GMIP / Q9P107
GMPPA / Q96IJ6
Gene Symbol / UniProt Accession
Brain
Colon
Duodenum
Esophagus
Heart
Liver
Lung
Pancreas
Thyroid
Organs
Type
DDA
DIA
Bins
1
2
3
4
5

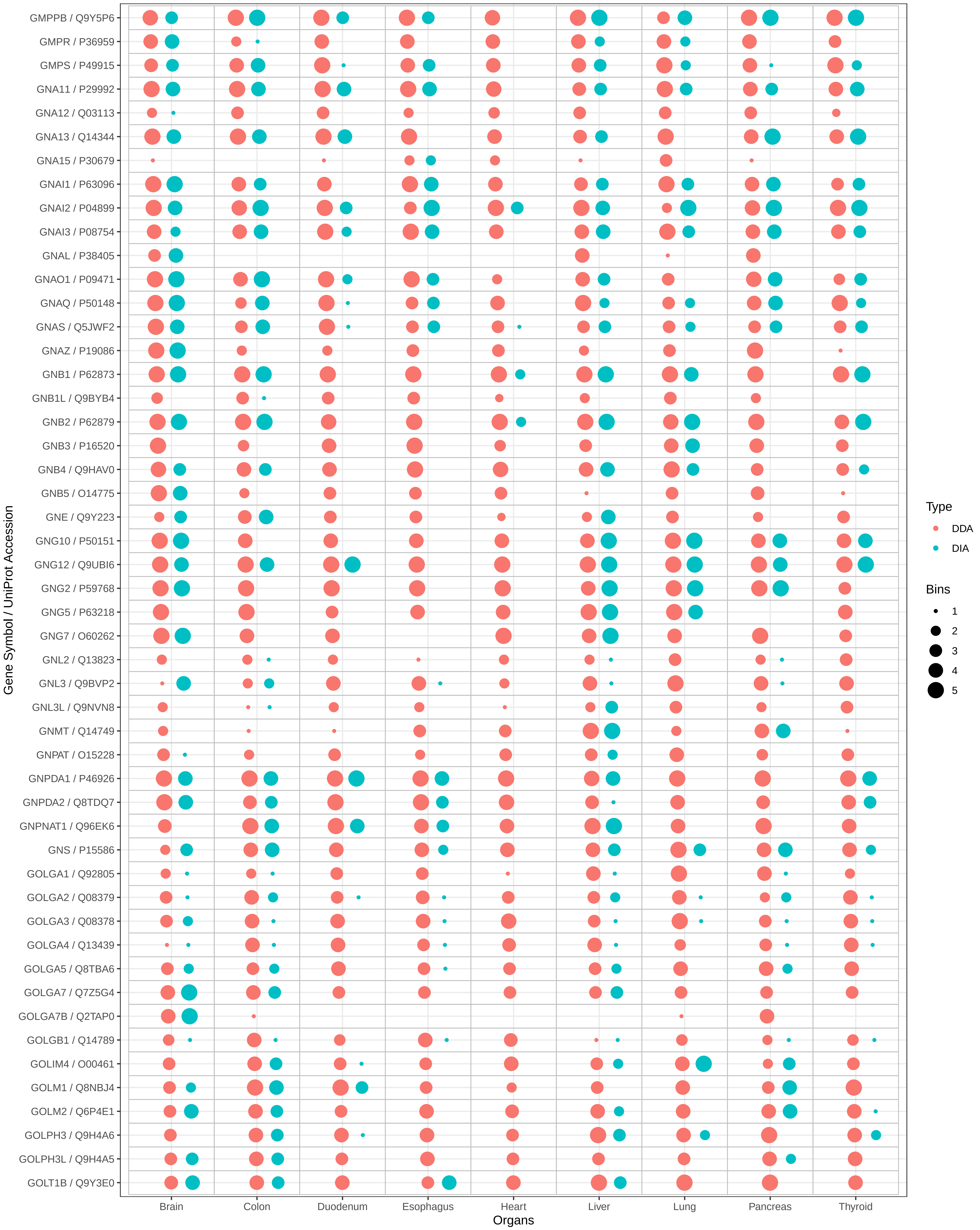

GMPPB / Q9Y5P6
GMPR / P36959
GMPS / P49915
GNA11 / P29992
GNA12 / Q03113
GNA13 / Q14344
GNA15 / P30679
GNAI1 / P63096
GNAI2 / P04899
GNAI3 / P08754
GNAL / P38405
GNAO1 / P09471
GNAQ / P50148
GNAS / Q5JWF2
GNAZ / P19086
GNB1 / P62873
GNB1L / Q9BYB4
GNB2 / P62879
GNB3 / P16520
GNB4 / Q9HAV0
GNB5 / O14775
GNE / Q9Y223
GNG10 / P50151
GNG12 / Q9UBI6
GNG2 / P59768
GNG5 / P63218
GNG7 / O60262
GNL2 / Q13823
GNL3 / Q9BVP2
GNL3L / Q9NVN8
GNMT / Q14749
GNPAT / O15228
GNPDA1 / P46926
GNPDA2 / Q8TDQ7
GNPNAT1 / Q96EK6
GNS / P15586
GOLGA1 / Q92805
GOLGA2 / Q08379
GOLGA3 / Q08378
GOLGA4 / Q13439
GOLGA5 / Q8TBA6
GOLGA7 / Q7Z5G4
GOLGA7B / Q2TAP0
GOLGB1 / Q14789
GOLIM4 / O00461
GOLM1 / Q8NBJ4
GOLM2 / Q6P4E1
GOLPH3 / Q9H4A6
GOLPH3L / Q9H4A5
GOLT1B / Q9Y3E0
Gene Symbol / UniProt Accession
Brain
Colon
Duodenum
Esophagus
Heart
Liver
Lung
Pancreas
Thyroid
Organs
Type
DDA
DIA
Bins
1
2
3
4
5

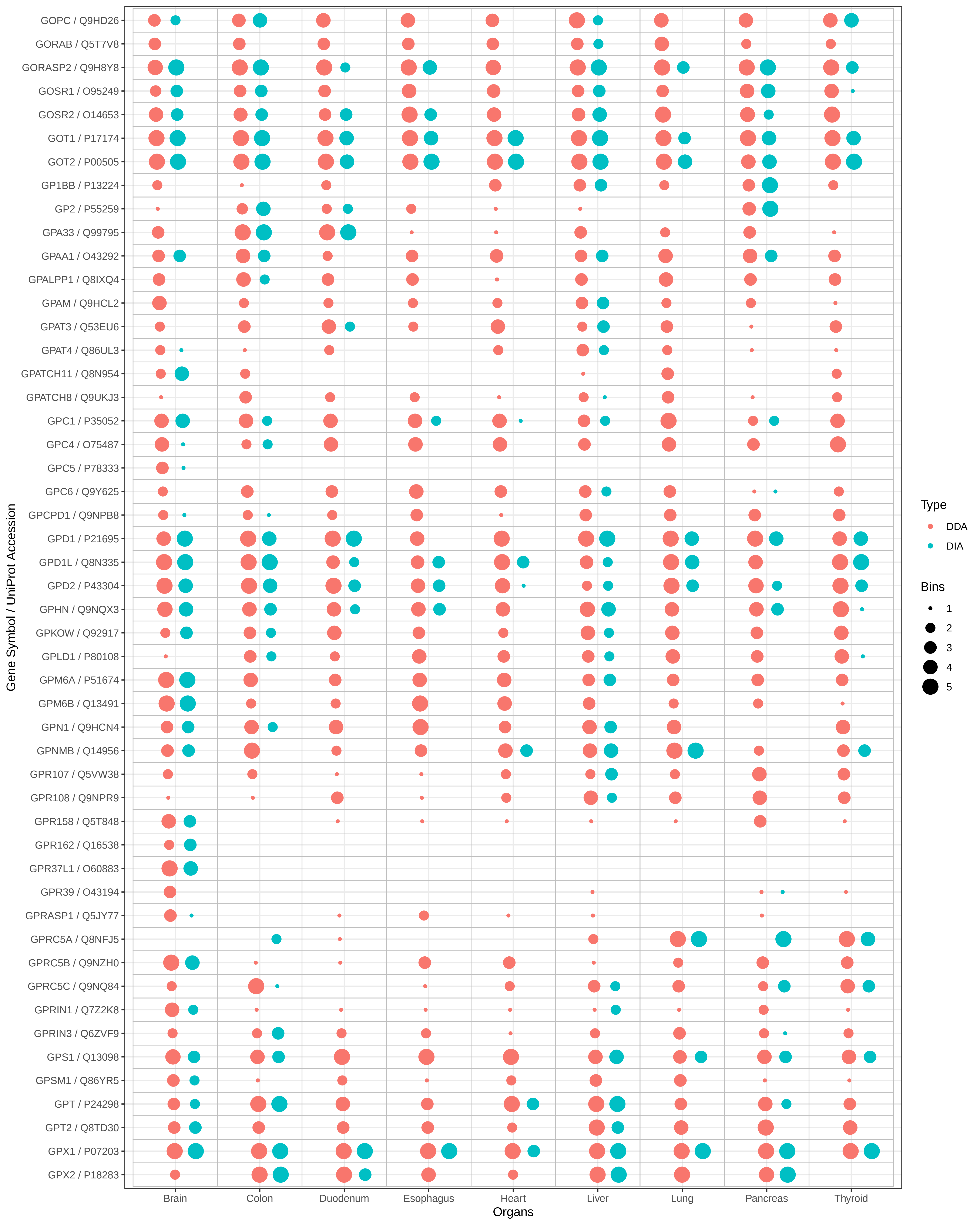

GOPC / Q9HD26
GORAB / Q5T7V8
GORASP2 / Q9H8Y8
GOSR1 / O95249
GOSR2 / O14653
GOT1 / P17174
GOT2 / P00505
GP1BB / P13224
GP2 / P55259
GPA33 / Q99795
GPAA1 / O43292
GPALPP1 / Q8IXQ4
GPAM / Q9HCL2
GPAT3 / Q53EU6
GPAT4 / Q86UL3
GPATCH11 / Q8N954
GPATCH8 / Q9UKJ3
GPC1 / P35052
GPC4 / O75487
GPC5 / P78333
GPC6 / Q9Y625
GPCPD1 / Q9NPB8
GPD1 / P21695
GPD1L / Q8N335
GPD2 / P43304
GPHN / Q9NQX3
GPKOW / Q92917
GPLD1 / P80108
GPM6A / P51674
GPM6B / Q13491
GPN1 / Q9HCN4
GPNMB / Q14956
GPR107 / Q5VW38
GPR108 / Q9NPR9
GPR158 / Q5T848
GPR162 / Q16538
GPR37L1 / O60883
GPR39 / O43194
GPRASP1 / Q5JY77
GPRC5A / Q8NFJ5
GPRC5B / Q9NZH0
GPRC5C / Q9NQ84
GPRIN1 / Q7Z2K8
GPRIN3 / Q6ZVF9
GPS1 / Q13098
GPSM1 / Q86YR5
GPT / P24298
GPT2 / Q8TD30
GPX1 / P07203
GPX2 / P18283
Gene Symbol / UniProt Accession
Brain
Colon
Duodenum
Esophagus
Heart
Liver
Lung
Pancreas
Thyroid
Organs
Type
DDA
DIA
Bins
1
2
3
4
5

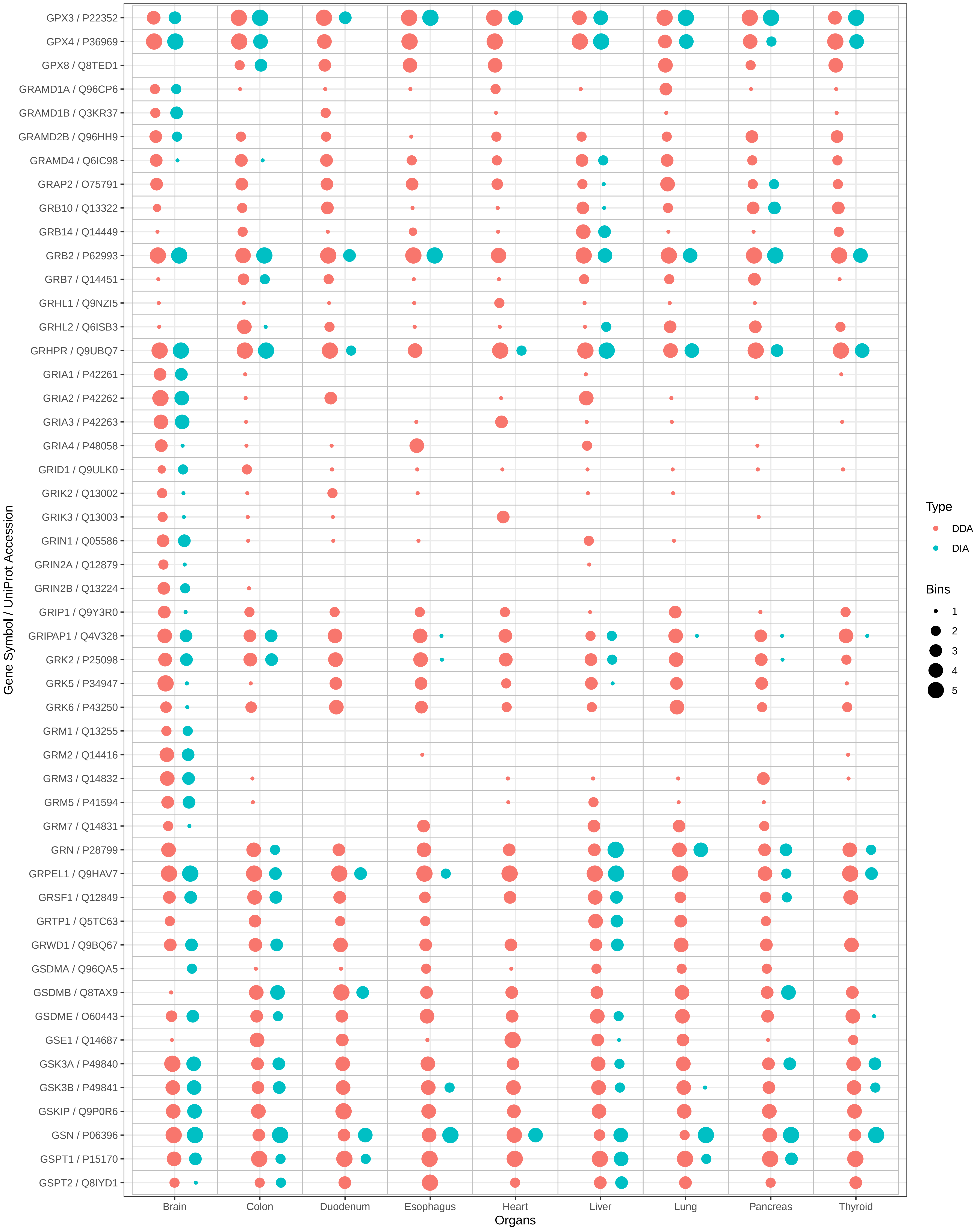

GPX3 / P22352
GPX4 / P36969
GPX8 / Q8TED1
GRAMD1A / Q96CP6
GRAMD1B / Q3KR37
GRAMD2B / Q96HH9
GRAMD4 / Q6IC98
GRAP2 / O75791
GRB10 / Q13322
GRB14 / Q14449
GRB2 / P62993
GRB7 / Q14451
GRHL1 / Q9NZI5
GRHL2 / Q6ISB3
GRHPR / Q9UBQ7
GRIA1 / P42261
GRIA2 / P42262
GRIA3 / P42263
GRIA4 / P48058
GRID1 / Q9ULK0
GRIK2 / Q13002
GRIK3 / Q13003
GRIN1 / Q05586
GRIN2A / Q12879
GRIN2B / Q13224
GRIP1 / Q9Y3R0
GRIPAP1 / Q4V328
GRK2 / P25098
GRK5 / P34947
GRK6 / P43250
GRM1 / Q13255
GRM2 / Q14416
GRM3 / Q14832
GRM5 / P41594
GRM7 / Q14831
GRN / P28799
GRPEL1 / Q9HAV7
GRSF1 / Q12849
GRTP1 / Q5TC63
GRWD1 / Q9BQ67
GSDMA / Q96QA5
GSDMB / Q8TAX9
GSDME / O60443
GSE1 / Q14687
GSK3A / P49840
GSK3B / P49841
GSKIP / Q9P0R6
GSN / P06396
GSPT1 / P15170
GSPT2 / Q8IYD1
Gene Symbol / UniProt Accession
Brain
Colon
Duodenum
Esophagus
Heart
Liver
Lung
Pancreas
Thyroid
Organs
Type
DDA
DIA
Bins
1
2
3
4
5

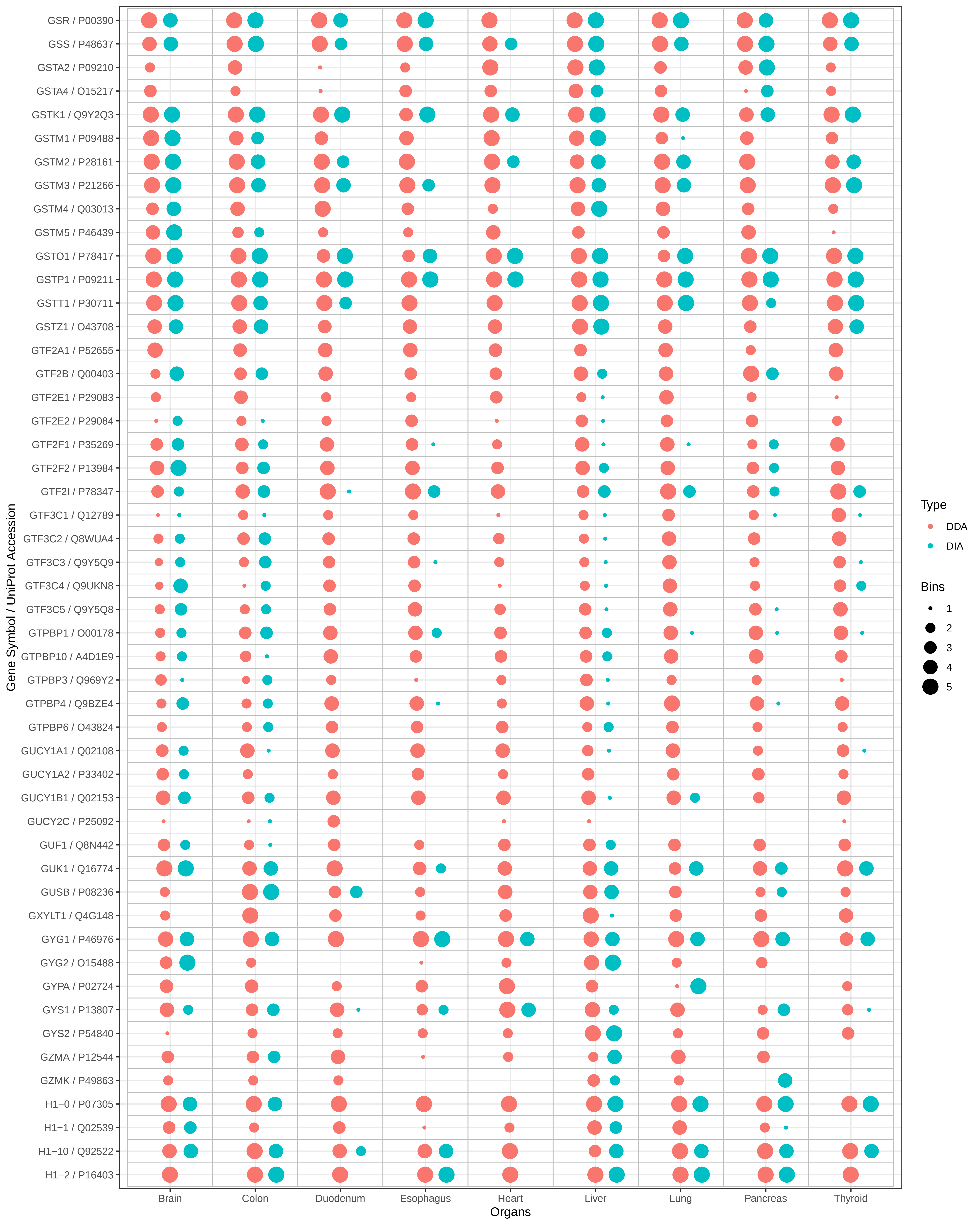

GSR / P00390
GSS / P48637
GSTA2 / P09210
GSTA4 / O15217
GSTK1 / Q9Y2Q3
GSTM1 / P09488
GSTM2 / P28161
GSTM3 / P21266
GSTM4 / Q03013
GSTM5 / P46439
GSTO1 / P78417
GSTP1 / P09211
GSTT1 / P30711
GSTZ1 / O43708
GTF2A1 / P52655
GTF2B / Q00403
GTF2E1 / P29083
GTF2E2 / P29084
GTF2F1 / P35269
GTF2F2 / P13984
GTF2I / P78347
GTF3C1 / Q12789
GTF3C2 / Q8WUA4
GTF3C3 / Q9Y5Q9
GTF3C4 / Q9UKN8
GTF3C5 / Q9Y5Q8
GTPBP1 / O00178
GTPBP10 / A4D1E9
GTPBP3 / Q969Y2
GTPBP4 / Q9BZE4
GTPBP6 / O43824
GUCY1A1 / Q02108
GUCY1A2 / P33402
GUCY1B1 / Q02153
GUCY2C / P25092
GUF1 / Q8N442
GUK1 / Q16774
GUSB / P08236
GXYLT1 / Q4G148
GYG1 / P46976
GYG2 / O15488
GYPA / P02724
GYS1 / P13807
GYS2 / P54840
GZMA / P12544
GZMK / P49863
H1−0 / P07305
H1−1 / Q02539
H1−10 / Q92522
H1−2 / P16403
Gene Symbol / UniProt Accession
Brain
Colon
Duodenum
Esophagus
Heart
Liver
Lung
Pancreas
Thyroid
Organs
Type
DDA
DIA
Bins
1
2
3
4
5

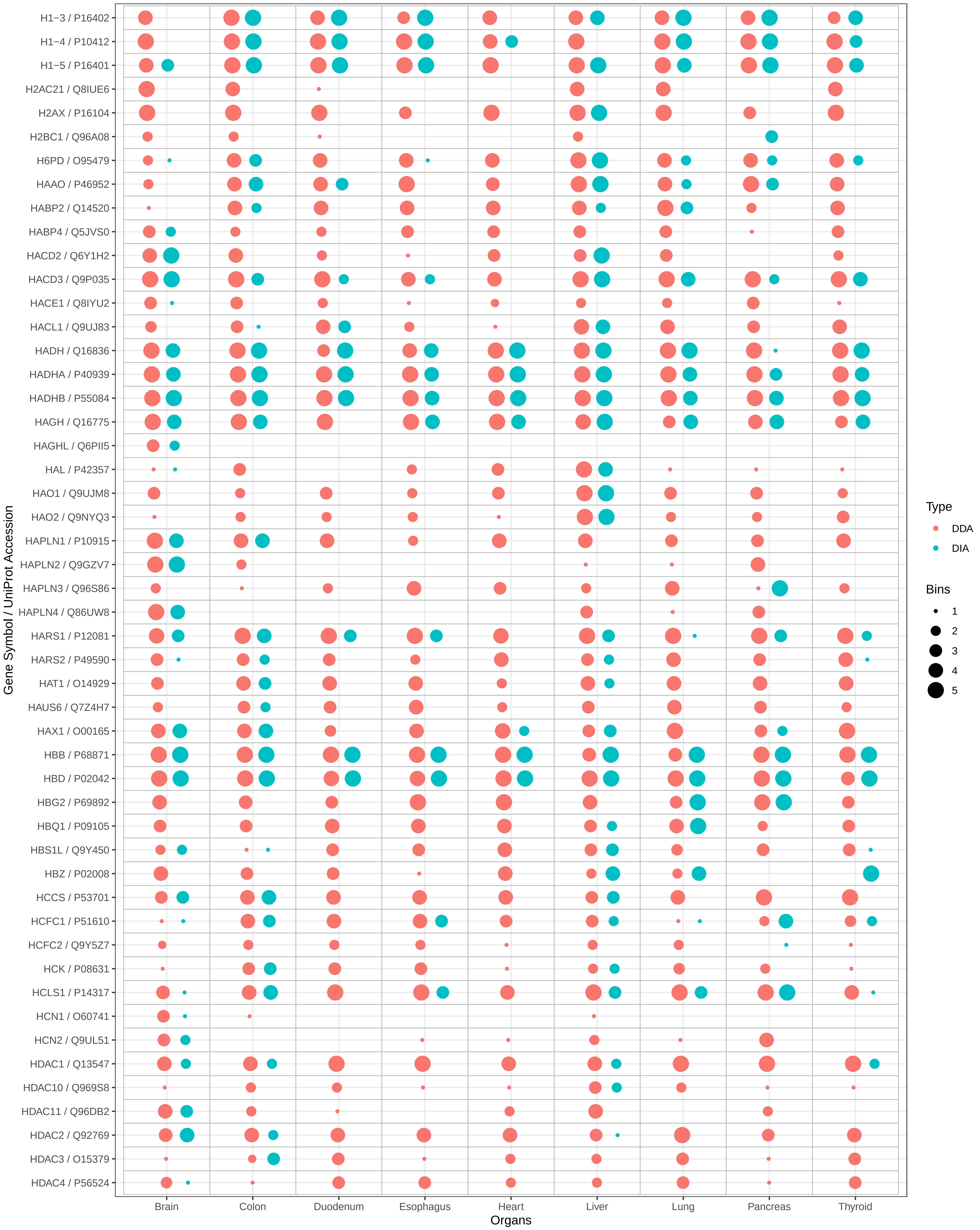

H1−3 / P16402
H1−4 / P10412
H1−5 / P16401
H2AC21 / Q8IUE6
H2AX / P16104
H2BC1 / Q96A08
H6PD / O95479
HAAO / P46952
HABP2 / Q14520
HABP4 / Q5JVS0
HACD2 / Q6Y1H2
HACD3 / Q9P035
HACE1 / Q8IYU2
HACL1 / Q9UJ83
HADH / Q16836
HADHA / P40939
HADHB / P55084
HAGH / Q16775
HAGHL / Q6PII5
HAL / P42357
HAO1 / Q9UJM8
HAO2 / Q9NYQ3
HAPLN1 / P10915
HAPLN2 / Q9GZV7
HAPLN3 / Q96S86
HAPLN4 / Q86UW8
HARS1 / P12081
HARS2 / P49590
HAT1 / O14929
HAUS6 / Q7Z4H7
HAX1 / O00165
HBB / P68871
HBD / P02042
HBG2 / P69892
HBQ1 / P09105
HBS1L / Q9Y450
HBZ / P02008
HCCS / P53701
HCFC1 / P51610
HCFC2 / Q9Y5Z7
HCK / P08631
HCLS1 / P14317
HCN1 / O60741
HCN2 / Q9UL51
HDAC1 / Q13547
HDAC10 / Q969S8
HDAC11 / Q96DB2
HDAC2 / Q92769
HDAC3 / O15379
HDAC4 / P56524
Gene Symbol / UniProt Accession
Brain
Colon
Duodenum
Esophagus
Heart
Liver
Lung
Pancreas
Thyroid
Organs
Type
DDA
DIA
Bins
1
2
3
4
5

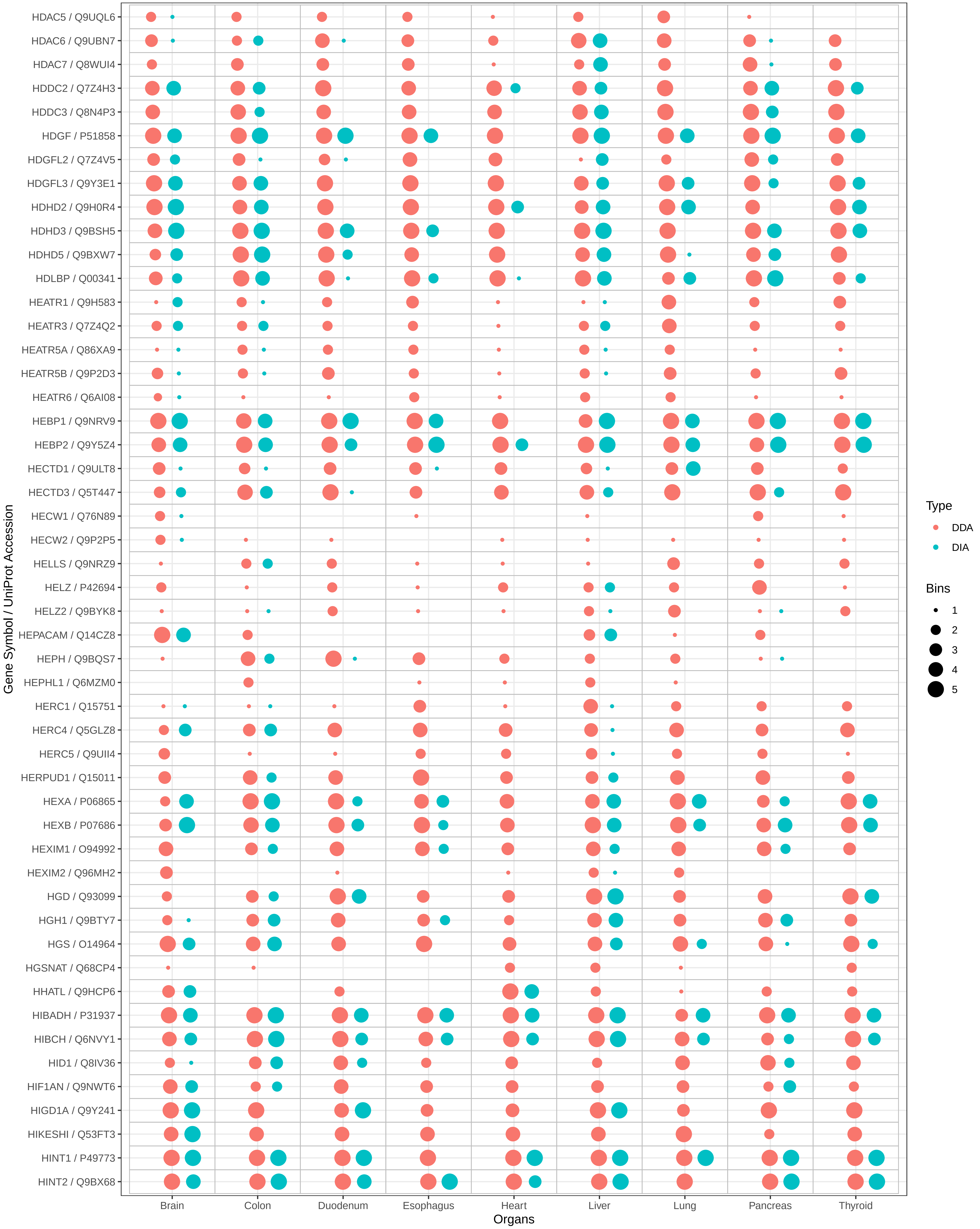

HDAC5 / Q9UQL6
HDAC6 / Q9UBN7
HDAC7 / Q8WUI4
HDDC2 / Q7Z4H3
HDDC3 / Q8N4P3
HDGF / P51858
HDGFL2 / Q7Z4V5
HDGFL3 / Q9Y3E1
HDHD2 / Q9H0R4
HDHD3 / Q9BSH5
HDHD5 / Q9BXW7
HDLBP / Q00341
HEATR1 / Q9H583
HEATR3 / Q7Z4Q2
HEATR5A / Q86XA9
HEATR5B / Q9P2D3
HEATR6 / Q6AI08
HEBP1 / Q9NRV9
HEBP2 / Q9Y5Z4
HECTD1 / Q9ULT8
HECTD3 / Q5T447
HECW1 / Q76N89
HECW2 / Q9P2P5
HELLS / Q9NRZ9
HELZ / P42694
HELZ2 / Q9BYK8
HEPACAM / Q14CZ8
HEPH / Q9BQS7
HEPHL1 / Q6MZM0
HERC1 / Q15751
HERC4 / Q5GLZ8
HERC5 / Q9UII4
HERPUD1 / Q15011
HEXA / P06865
HEXB / P07686
HEXIM1 / O94992
HEXIM2 / Q96MH2
HGD / Q93099
HGH1 / Q9BTY7
HGS / O14964
HGSNAT / Q68CP4
HHATL / Q9HCP6
HIBADH / P31937
HIBCH / Q6NVY1
HID1 / Q8IV36
HIF1AN / Q9NWT6
HIGD1A / Q9Y241
HIKESHI / Q53FT3
HINT1 / P49773
HINT2 / Q9BX68
Gene Symbol / UniProt Accession
Brain
Colon
Duodenum
Esophagus
Heart
Liver
Lung
Pancreas
Thyroid
Organs
Type
DDA
DIA
Bins
1
2
3
4
5

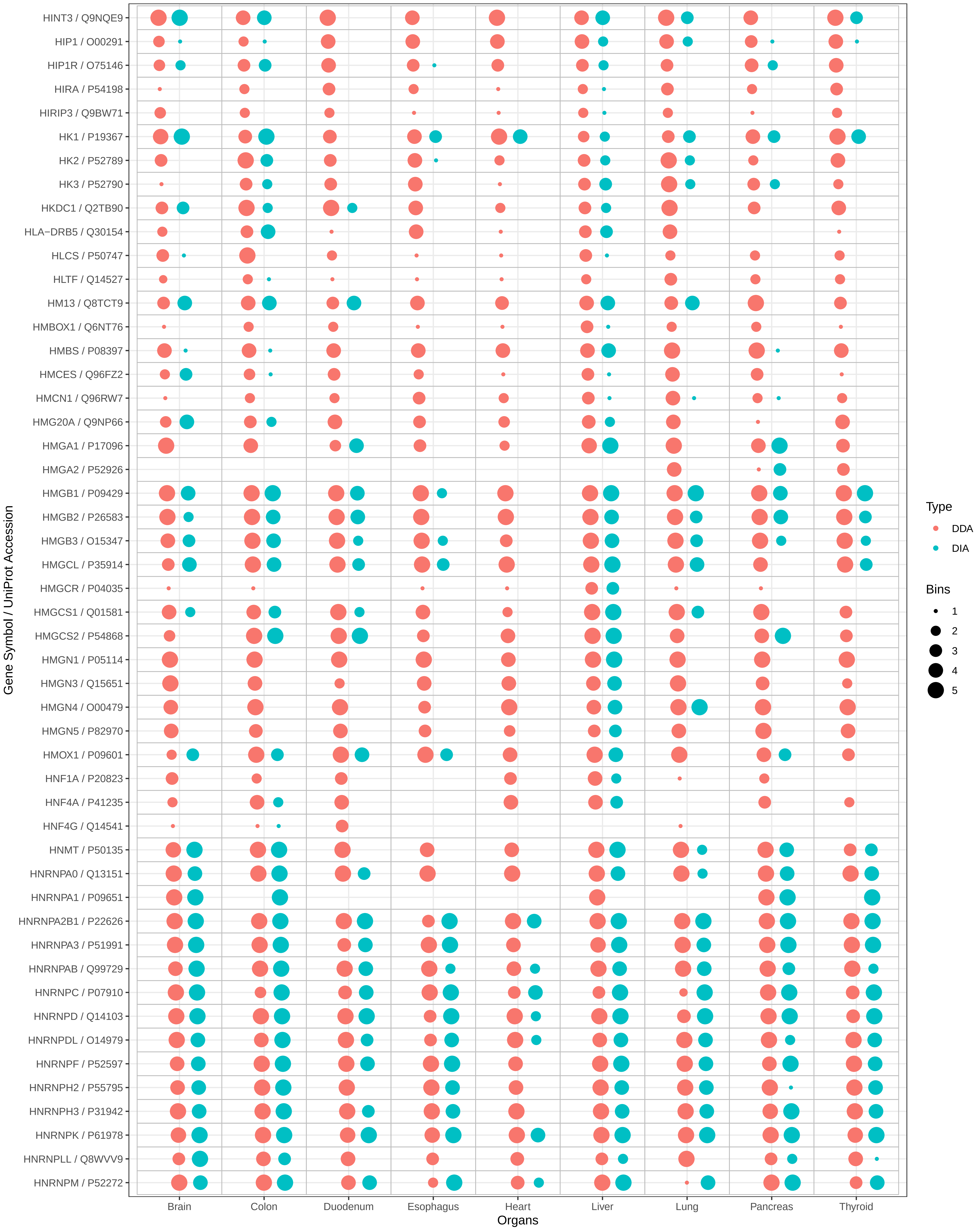
HINT3 / Q9NQE9
HIP1 / O00291
HIP1R / O75146
HIRA / P54198
HIRIP3 / Q9BW71
HK1 / P19367
HK2 / P52789
HK3 / P52790
HKDC1 / Q2TB90
HLA−DRB5 / Q30154
HLCS / P50747
HLTF / Q14527
HM13 / Q8TCT9
HMBOX1 / Q6NT76
HMBS / P08397
HMCES / Q96FZ2
HMCN1 / Q96RW7
HMG20A / Q9NP66
HMGA1 / P17096
HMGA2 / P52926
HMGB1 / P09429
HMGB2 / P26583
HMGB3 / O15347
HMGCL / P35914
HMGCR / P04035
HMGCS1 / Q01581
HMGCS2 / P54868
HMGN1 / P05114
HMGN3 / Q15651
HMGN4 / O00479
HMGN5 / P82970
HMOX1 / P09601
HNF1A / P20823
HNF4A / P41235
HNF4G / Q14541
HNMT / P50135
HNRNPA0 / Q13151
HNRNPA1 / P09651
HNRNPA2B1 / P22626
HNRNPA3 / P51991
HNRNPAB / Q99729
HNRNPC / P07910
HNRNPD / Q14103
HNRNPDL / O14979
HNRNPF / P52597
HNRNPH2 / P55795
HNRNPH3 / P31942
HNRNPK / P61978
HNRNPLL / Q8WVV9
HNRNPM / P52272
Gene Symbol / UniProt Accession
Brain
Colon
Duodenum
Esophagus
Heart
Liver
Lung
Pancreas
Thyroid
Organs
Type
DDA
DIA
Bins
1
2
3
4
5

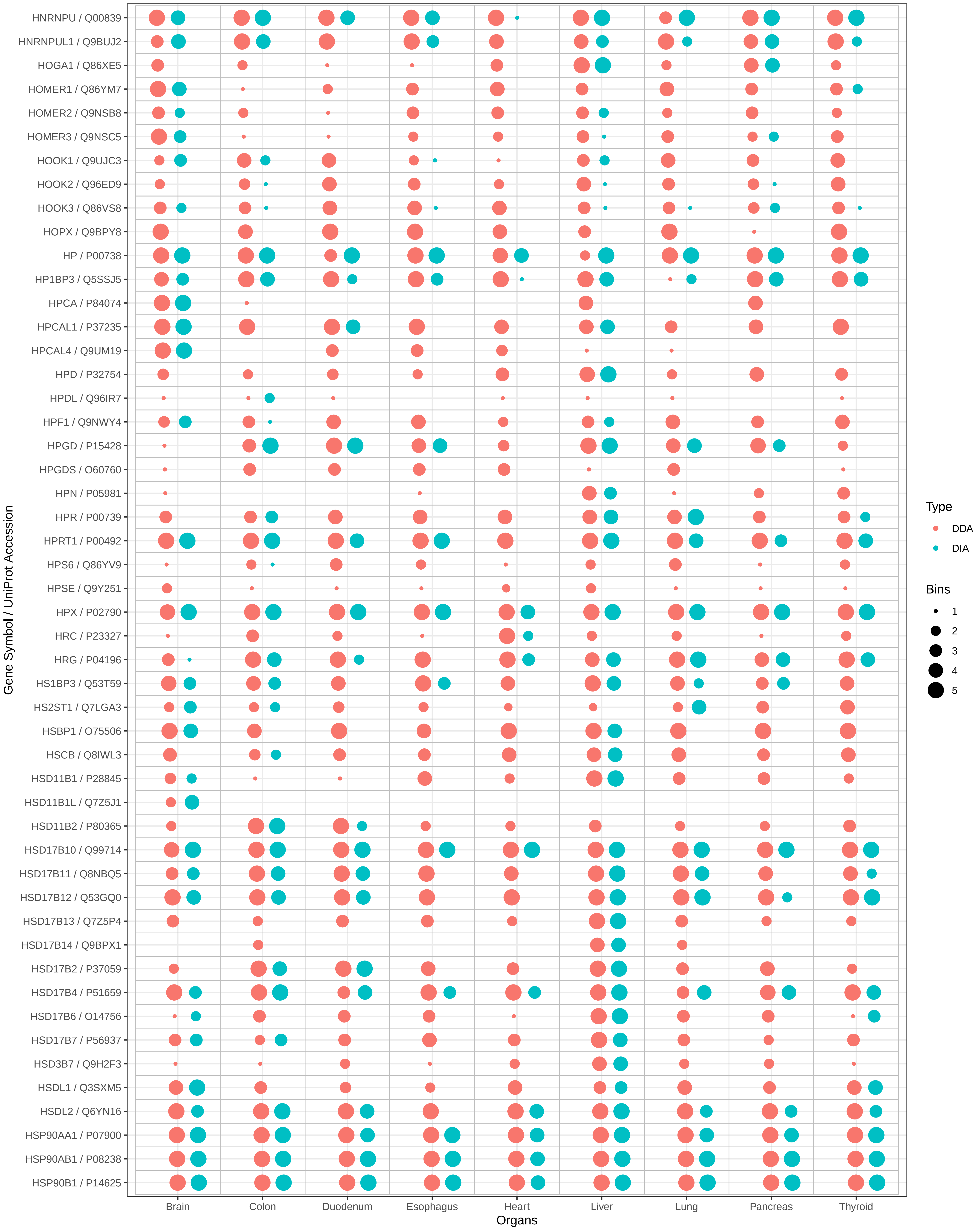

HNRNPU / Q00839
HNRNPUL1 / Q9BUJ2
HOGA1 / Q86XE5
HOMER1 / Q86YM7
HOMER2 / Q9NSB8
HOMER3 / Q9NSC5
HOOK1 / Q9UJC3
HOOK2 / Q96ED9
HOOK3 / Q86VS8
HOPX / Q9BPY8
HP / P00738
HP1BP3 / Q5SSJ5
HPCA / P84074
HPCAL1 / P37235
HPCAL4 / Q9UM19
HPD / P32754
HPDL / Q96IR7
HPF1 / Q9NWY4
HPGD / P15428
HPGDS / O60760
HPN / P05981
HPR / P00739
HPRT1 / P00492
HPS6 / Q86YV9
HPSE / Q9Y251
HPX / P02790
HRC / P23327
HRG / P04196
HS1BP3 / Q53T59
HS2ST1 / Q7LGA3
HSBP1 / O75506
HSCB / Q8IWL3
HSD11B1 / P28845
HSD11B1L / Q7Z5J1
HSD11B2 / P80365
HSD17B10 / Q99714
HSD17B11 / Q8NBQ5
HSD17B12 / Q53GQ0
HSD17B13 / Q7Z5P4
HSD17B14 / Q9BPX1
HSD17B2 / P37059
HSD17B4 / P51659
HSD17B6 / O14756
HSD17B7 / P56937
HSD3B7 / Q9H2F3
HSDL1 / Q3SXM5
HSDL2 / Q6YN16
HSP90AA1 / P07900
HSP90AB1 / P08238
HSP90B1 / P14625
Gene Symbol / UniProt Accession
Brain
Colon
Duodenum
Esophagus
Heart
Liver
Lung
Pancreas
Thyroid
Organs
Type
DDA
DIA
Bins
1
2
3
4
5

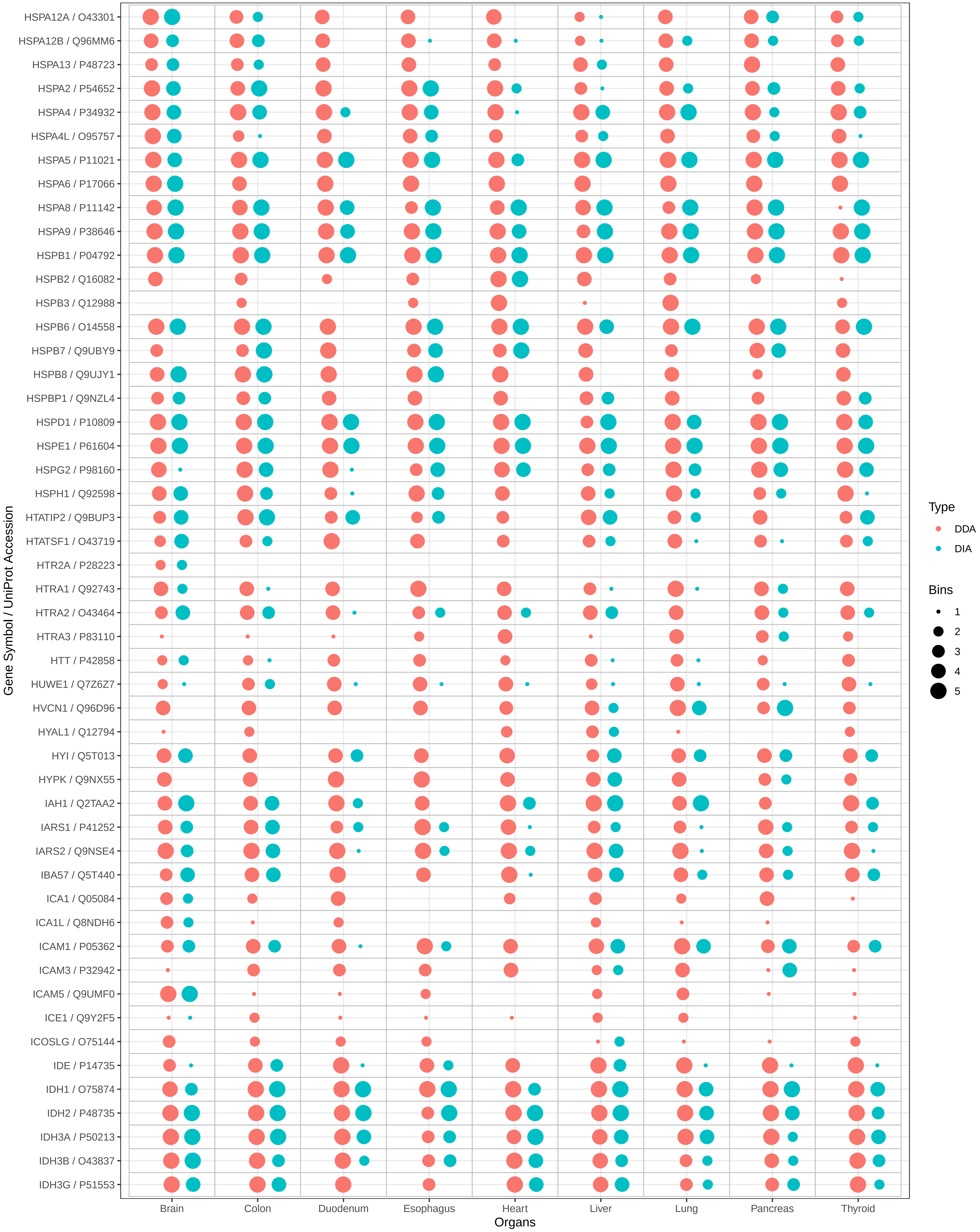

HSPA12A / O43301
HSPA12B / Q96MM6
HSPA13 / P48723
HSPA2 / P54652
HSPA4 / P34932
HSPA4L / O95757
HSPA5 / P11021
HSPA6 / P17066
HSPA8 / P11142
HSPA9 / P38646
HSPB1 / P04792
HSPB2 / Q16082
HSPB3 / Q12988
HSPB6 / O14558
HSPB7 / Q9UBY9
HSPB8 / Q9UJY1
HSPBP1 / Q9NZL4
HSPD1 / P10809
HSPE1 / P61604
HSPG2 / P98160
HSPH1 / Q92598
HTATIP2 / Q9BUP3
HTATSF1 / O43719
HTR2A / P28223
HTRA1 / Q92743
HTRA2 / O43464
HTRA3 / P83110
HTT / P42858
HUWE1 / Q7Z6Z7
HVCN1 / Q96D96
HYAL1 / Q12794
HYI / Q5T013
HYPK / Q9NX55
IAH1 / Q2TAA2
IARS1 / P41252
IARS2 / Q9NSE4
IBA57 / Q5T440
ICA1 / Q05084
ICA1L / Q8NDH6
ICAM1 / P05362
ICAM3 / P32942
ICAM5 / Q9UMF0
ICE1 / Q9Y2F5
ICOSLG / O75144
IDE / P14735
IDH1 / O75874
IDH2 / P48735
IDH3A / P50213
IDH3B / O43837
IDH3G / P51553
Gene Symbol / UniProt Accession
Brain
Colon
Duodenum
Esophagus
Heart
Liver
Lung
Pancreas
Thyroid
Organs
Type
DDA
DIA
Bins
1
2
3
4
5

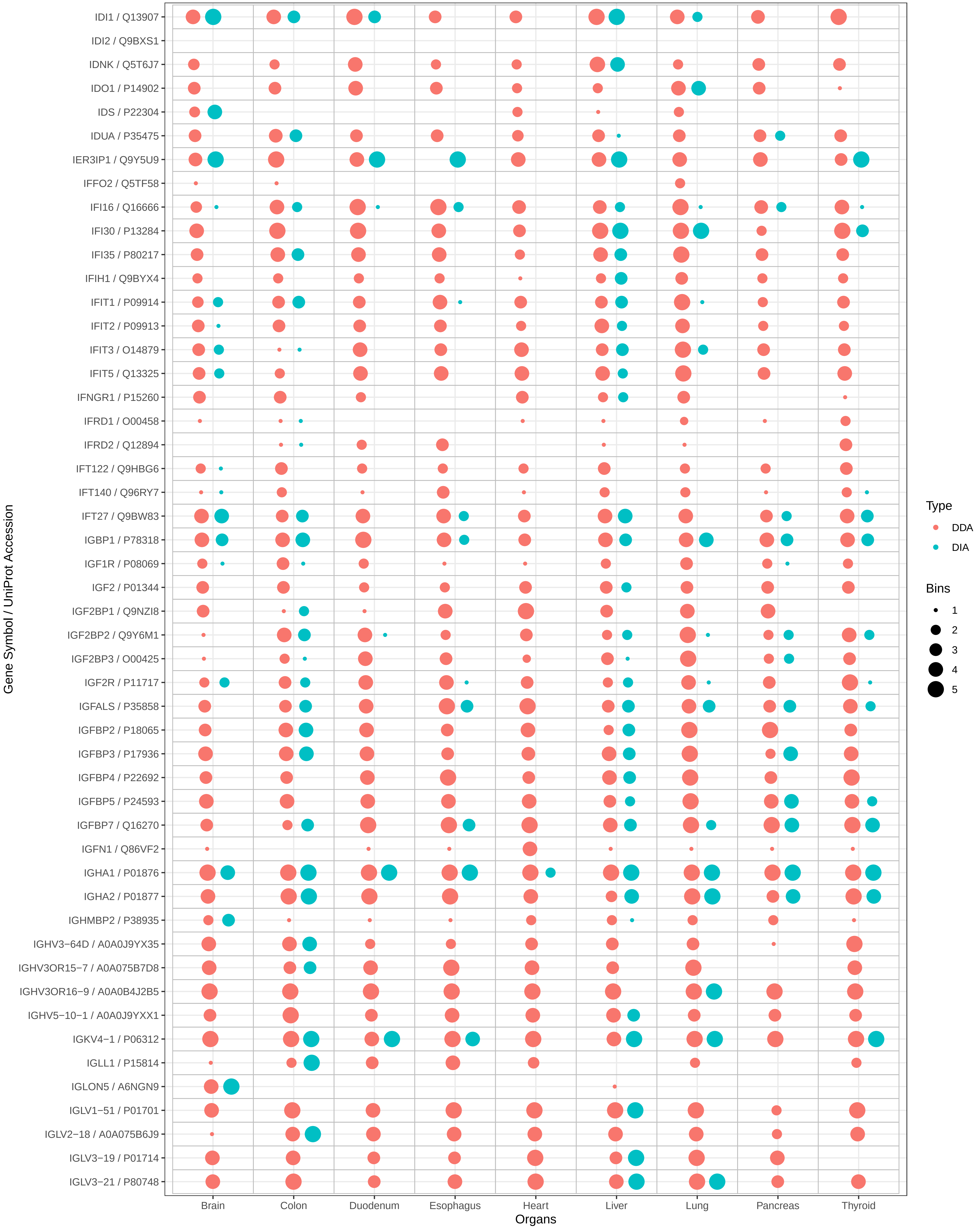
IDI1 / Q13907
IDI2 / Q9BXS1
IDNK / Q5T6J7
IDO1 / P14902
IDS / P22304
IDUA / P35475
IER3IP1 / Q9Y5U9
IFFO2 / Q5TF58
IFI16 / Q16666
IFI30 / P13284
IFI35 / P80217
IFIH1 / Q9BYX4
IFIT1 / P09914
IFIT2 / P09913
IFIT3 / O14879
IFIT5 / Q13325
IFNGR1 / P15260
IFRD1 / O00458
IFRD2 / Q12894
IFT122 / Q9HBG6
IFT140 / Q96RY7
IFT27 / Q9BW83
IGBP1 / P78318
IGF1R / P08069
IGF2 / P01344
IGF2BP1 / Q9NZI8
IGF2BP2 / Q9Y6M1
IGF2BP3 / O00425
IGF2R / P11717
IGFALS / P35858
IGFBP2 / P18065
IGFBP3 / P17936
IGFBP4 / P22692
IGFBP5 / P24593
IGFBP7 / Q16270
IGFN1 / Q86VF2
IGHA1 / P01876
IGHA2 / P01877
IGHMBP2 / P38935
IGHV3−64D / A0A0J9YX35
IGHV3OR15−7 / A0A075B7D8
IGHV3OR16−9 / A0A0B4J2B5
IGHV5−10−1 / A0A0J9YXX1
IGKV4−1 / P06312
IGLL1 / P15814
IGLON5 / A6NGN9
IGLV1−51 / P01701
IGLV2−18 / A0A075B6J9
IGLV3−19 / P01714
IGLV3−21 / P80748
Gene Symbol / UniProt Accession
Brain
Colon
Duodenum
Esophagus
Heart
Liver
Lung
Pancreas
Thyroid
Organs
Type
DDA
DIA
Bins
1
2
3
4
5

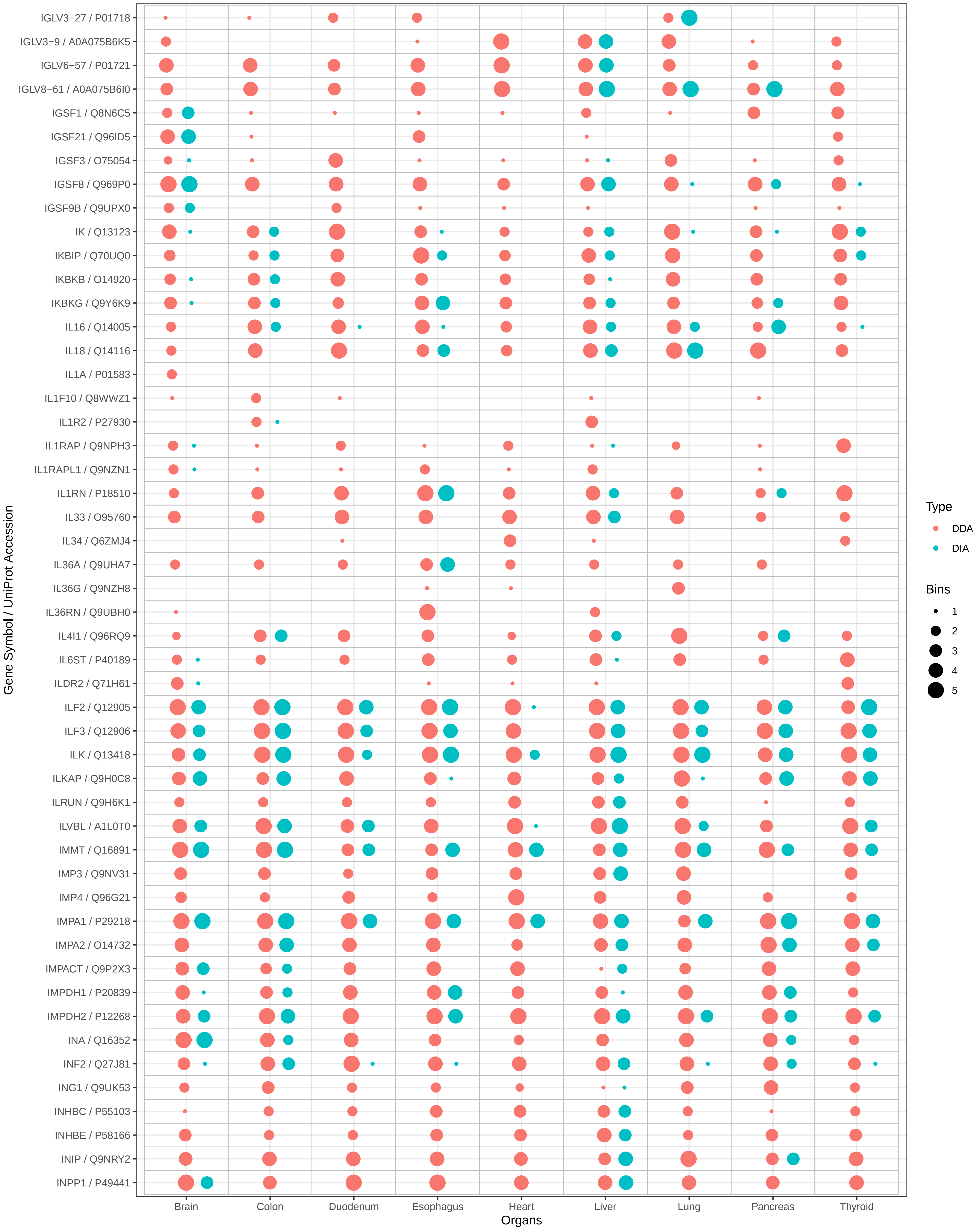
IGLV3−27 / P01718
IGLV3−9 / A0A075B6K5
IGLV6−57 / P01721
IGLV8−61 / A0A075B6I0
IGSF1 / Q8N6C5
IGSF21 / Q96ID5
IGSF3 / O75054
IGSF8 / Q969P0
IGSF9B / Q9UPX0
IK / Q13123
IKBIP / Q70UQ0
IKBKB / O14920
IKBKG / Q9Y6K9
IL16 / Q14005
IL18 / Q14116
IL1A / P01583
IL1F10 / Q8WWZ1
IL1R2 / P27930
IL1RAP / Q9NPH3
IL1RAPL1 / Q9NZN1
IL1RN / P18510
IL33 / O95760
IL34 / Q6ZMJ4
IL36A / Q9UHA7
IL36G / Q9NZH8
IL36RN / Q9UBH0
IL4I1 / Q96RQ9
IL6ST / P40189
ILDR2 / Q71H61
ILF2 / Q12905
ILF3 / Q12906
ILK / Q13418
ILKAP / Q9H0C8
ILRUN / Q9H6K1
ILVBL / A1L0T0
IMMT / Q16891
IMP3 / Q9NV31
IMP4 / Q96G21
IMPA1 / P29218
IMPA2 / O14732
IMPACT / Q9P2X3
IMPDH1 / P20839
IMPDH2 / P12268
INA / Q16352
INF2 / Q27J81
ING1 / Q9UK53
INHBC / P55103
INHBE / P58166
INIP / Q9NRY2
INPP1 / P49441
Gene Symbol / UniProt Accession
Brain
Colon
Duodenum
Esophagus
Heart
Liver
Lung
Pancreas
Thyroid
Organs
Type
DDA
DIA
Bins
1
2
3
4
5

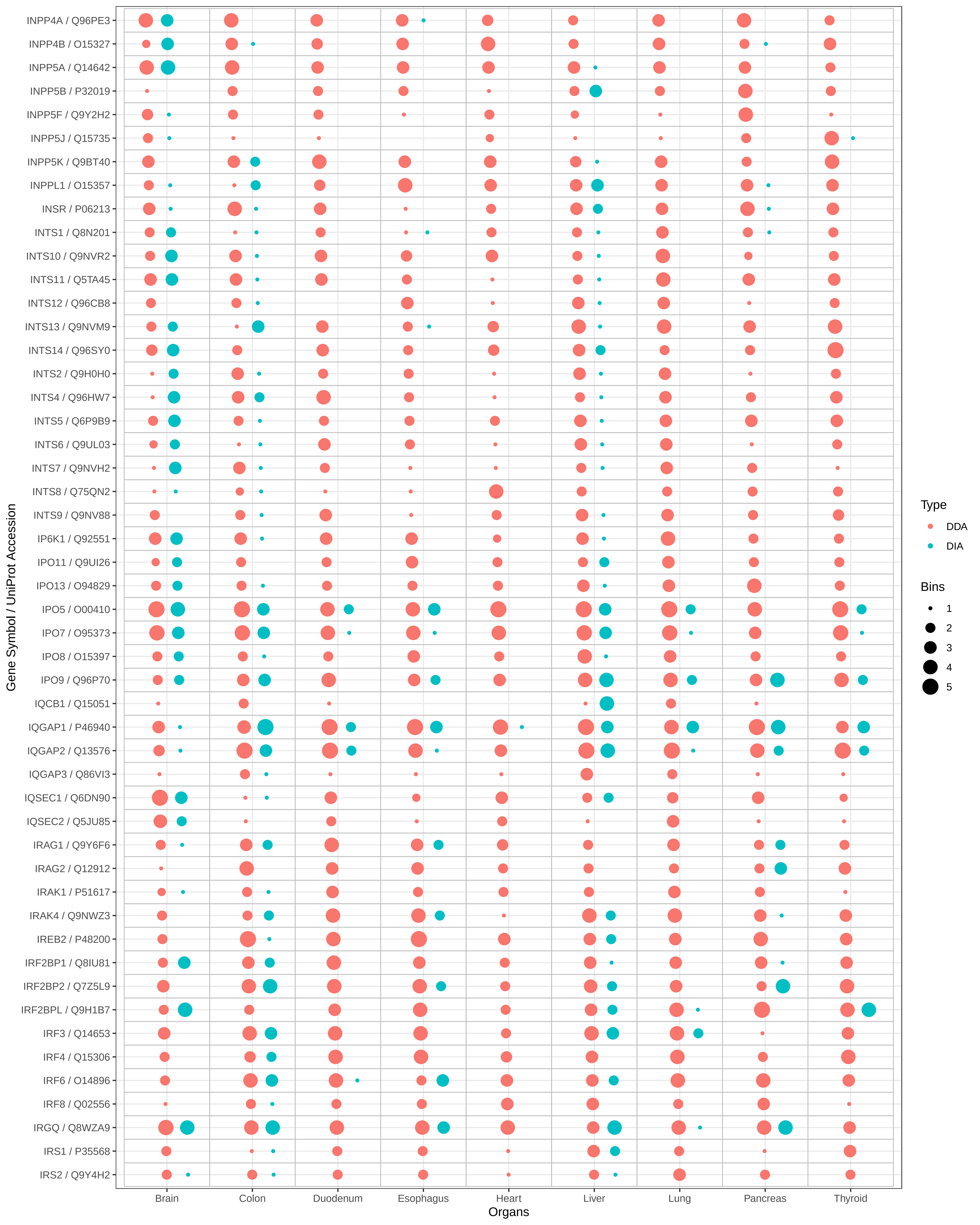
INPP4A / Q96PE3
INPP4B / O15327
INPP5A / Q14642
INPP5B / P32019
INPP5F / Q9Y2H2
INPP5J / Q15735
INPP5K / Q9BT40
INPPL1 / O15357
INSR / P06213
INTS1 / Q8N201
INTS10 / Q9NVR2
INTS11 / Q5TA45
INTS12 / Q96CB8
INTS13 / Q9NVM9
INTS14 / Q96SY0
INTS2 / Q9H0H0
INTS4 / Q96HW7
INTS5 / Q6P9B9
INTS6 / Q9UL03
INTS7 / Q9NVH2
INTS8 / Q75QN2
INTS9 / Q9NV88
IP6K1 / Q92551
IPO11 / Q9UI26
IPO13 / O94829
IPO5 / O00410
IPO7 / O95373
IPO8 / O15397
IPO9 / Q96P70
IQCB1 / Q15051
IQGAP1 / P46940
IQGAP2 / Q13576
IQGAP3 / Q86VI3
IQSEC1 / Q6DN90
IQSEC2 / Q5JU85
IRAG1 / Q9Y6F6
IRAG2 / Q12912
IRAK1 / P51617
IRAK4 / Q9NWZ3
IREB2 / P48200
IRF2BP1 / Q8IU81
IRF2BP2 / Q7Z5L9
IRF2BPL / Q9H1B7
IRF3 / Q14653
IRF4 / Q15306
IRF6 / O14896
IRF8 / Q02556
IRGQ / Q8WZA9
IRS1 / P35568
IRS2 / Q9Y4H2
Gene Symbol / UniProt Accession
Brain
Colon
Duodenum
Esophagus
Heart
Liver
Lung
Pancreas
Thyroid
Organs
Type
DDA
DIA
Bins
1
2
3
4
5

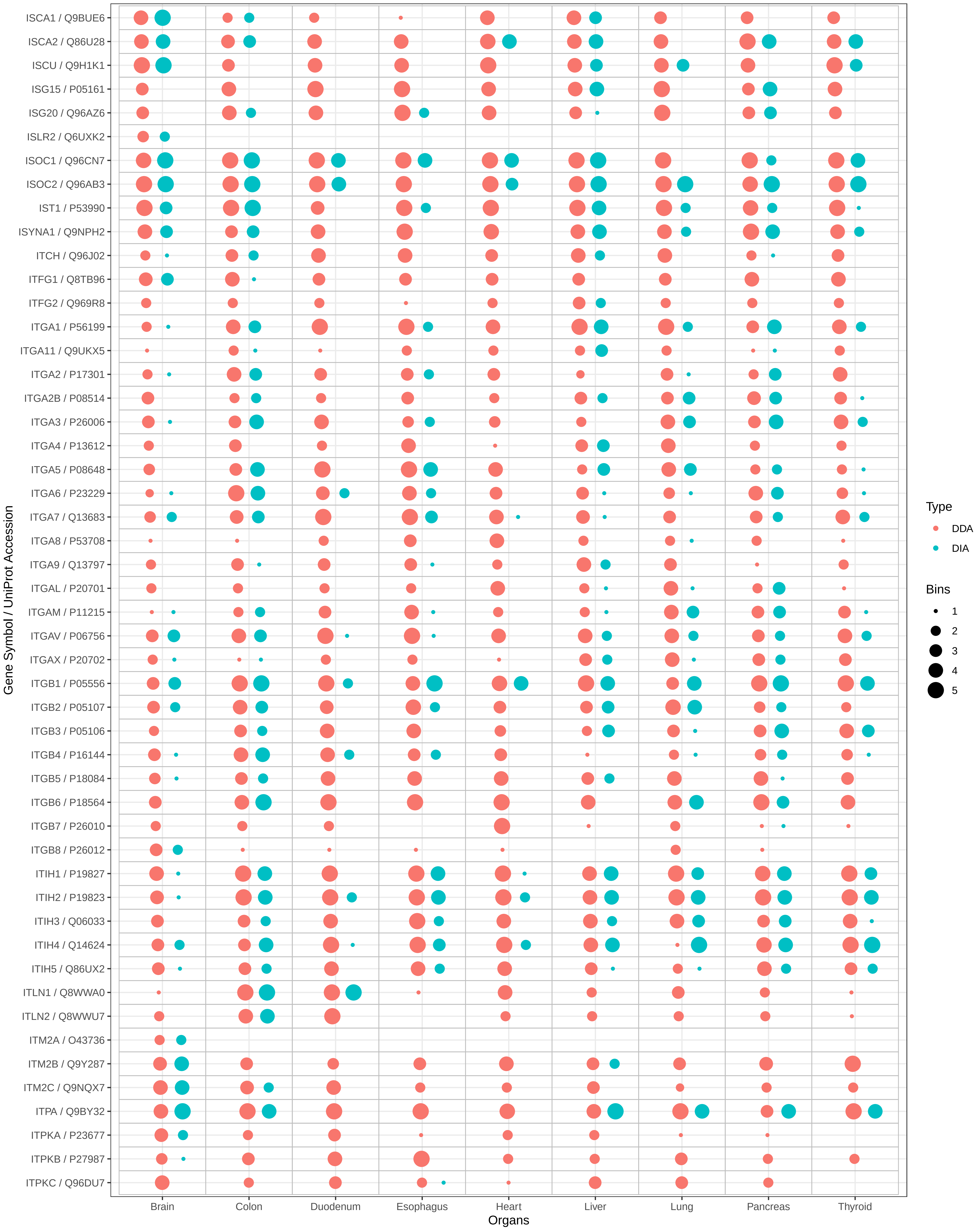

ISCA1 / Q9BUE6
ISCA2 / Q86U28
ISCU / Q9H1K1
ISG15 / P05161
ISG20 / Q96AZ6
ISLR2 / Q6UXK2
ISOC1 / Q96CN7
ISOC2 / Q96AB3
IST1 / P53990
ISYNA1 / Q9NPH2
ITCH / Q96J02
ITFG1 / Q8TB96
ITFG2 / Q969R8
ITGA1 / P56199
ITGA11 / Q9UKX5
ITGA2 / P17301
ITGA2B / P08514
ITGA3 / P26006
ITGA4 / P13612
ITGA5 / P08648
ITGA6 / P23229
ITGA7 / Q13683
ITGA8 / P53708
ITGA9 / Q13797
ITGAL / P20701
ITGAM / P11215
ITGAV / P06756
ITGAX / P20702
ITGB1 / P05556
ITGB2 / P05107
ITGB3 / P05106
ITGB4 / P16144
ITGB5 / P18084
ITGB6 / P18564
ITGB7 / P26010
ITGB8 / P26012
ITIH1 / P19827
ITIH2 / P19823
ITIH3 / Q06033
ITIH4 / Q14624
ITIH5 / Q86UX2
ITLN1 / Q8WWA0
ITLN2 / Q8WWU7
ITM2A / O43736
ITM2B / Q9Y287
ITM2C / Q9NQX7
ITPA / Q9BY32
ITPKA / P23677
ITPKB / P27987
ITPKC / Q96DU7
Gene Symbol / UniProt Accession
Brain
Colon
Duodenum
Esophagus
Heart
Liver
Lung
Pancreas
Thyroid
Organs
Type
DDA
DIA
Bins
1
2
3
4
5

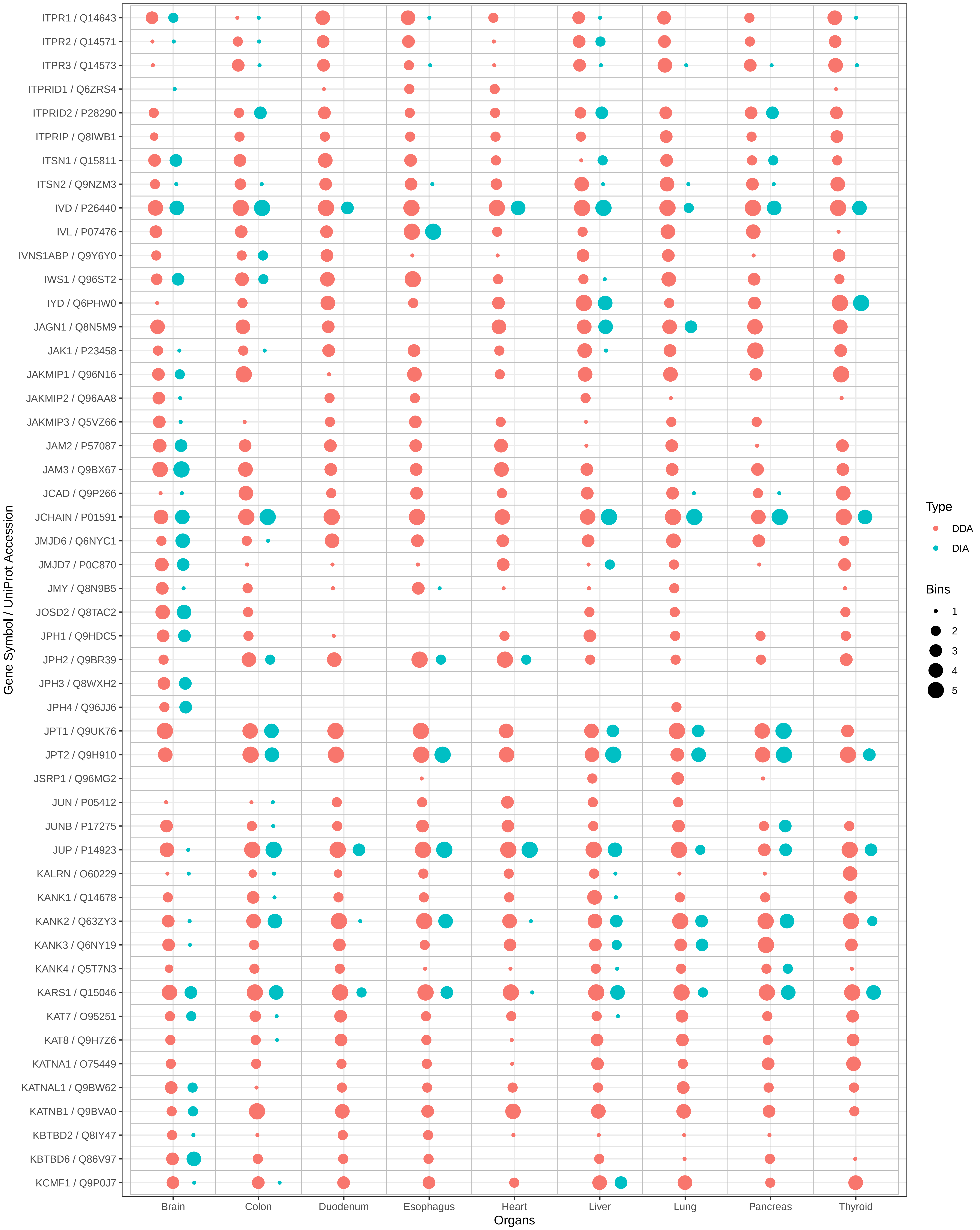

ITPR1 / Q14643
ITPR2 / Q14571
ITPR3 / Q14573
ITPRID1 / Q6ZRS4
ITPRID2 / P28290
ITPRIP / Q8IWB1
ITSN1 / Q15811
ITSN2 / Q9NZM3
IVD / P26440
IVL / P07476
IVNS1ABP / Q9Y6Y0
IWS1 / Q96ST2
IYD / Q6PHW0
JAGN1 / Q8N5M9
JAK1 / P23458
JAKMIP1 / Q96N16
JAKMIP2 / Q96AA8
JAKMIP3 / Q5VZ66
JAM2 / P57087
JAM3 / Q9BX67
JCAD / Q9P266
JCHAIN / P01591
JMJD6 / Q6NYC1
JMJD7 / P0C870
JMY / Q8N9B5
JOSD2 / Q8TAC2
JPH1 / Q9HDC5
JPH2 / Q9BR39
JPH3 / Q8WXH2
JPH4 / Q96JJ6
JPT1 / Q9UK76
JPT2 / Q9H910
JSRP1 / Q96MG2
JUN / P05412
JUNB / P17275
JUP / P14923
KALRN / O60229
KANK1 / Q14678
KANK2 / Q63ZY3
KANK3 / Q6NY19
KANK4 / Q5T7N3
KARS1 / Q15046
KAT7 / O95251
KAT8 / Q9H7Z6
KATNA1 / O75449
KATNAL1 / Q9BW62
KATNB1 / Q9BVA0
KBTBD2 / Q8IY47
KBTBD6 / Q86V97
KCMF1 / Q9P0J7
Gene Symbol / UniProt Accession
Brain
Colon
Duodenum
Esophagus
Heart
Liver
Lung
Pancreas
Thyroid
Organs
Type
DDA
DIA
Bins
1
2
3
4
5

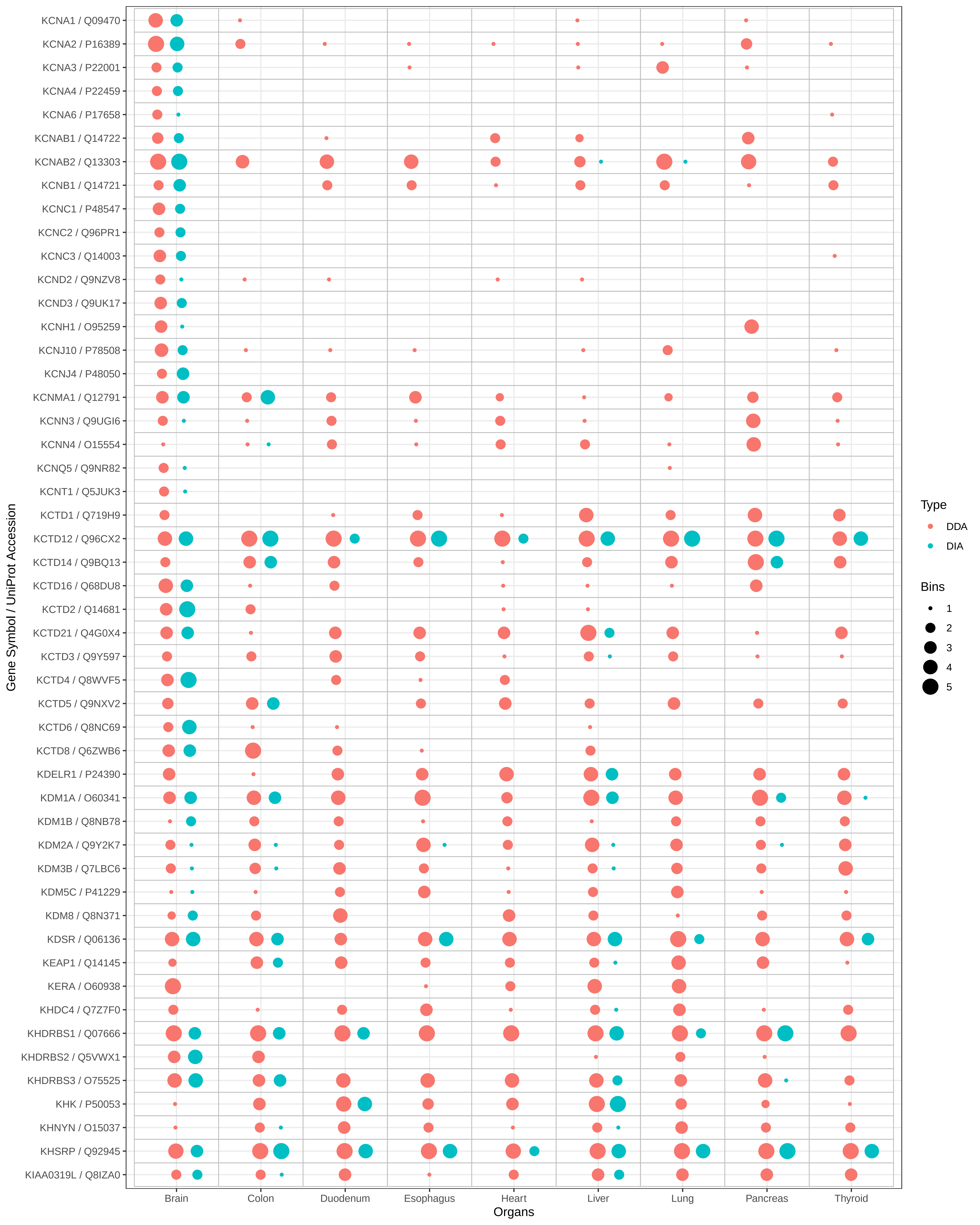

KCNA1 / Q09470
KCNA2 / P16389
KCNA3 / P22001
KCNA4 / P22459
KCNA6 / P17658
KCNAB1 / Q14722
KCNAB2 / Q13303
KCNB1 / Q14721
KCNC1 / P48547
KCNC2 / Q96PR1
KCNC3 / Q14003
KCND2 / Q9NZV8
KCND3 / Q9UK17
KCNH1 / O95259
KCNJ10 / P78508
KCNJ4 / P48050
KCNMA1 / Q12791
KCNN3 / Q9UGI6
KCNN4 / O15554
KCNQ5 / Q9NR82
KCNT1 / Q5JUK3
KCTD1 / Q719H9
KCTD12 / Q96CX2
KCTD14 / Q9BQ13
KCTD16 / Q68DU8
KCTD2 / Q14681
KCTD21 / Q4G0X4
KCTD3 / Q9Y597
KCTD4 / Q8WVF5
KCTD5 / Q9NXV2
KCTD6 / Q8NC69
KCTD8 / Q6ZWB6
KDELR1 / P24390
KDM1A / O60341
KDM1B / Q8NB78
KDM2A / Q9Y2K7
KDM3B / Q7LBC6
KDM5C / P41229
KDM8 / Q8N371
KDSR / Q06136
KEAP1 / Q14145
KERA / O60938
KHDC4 / Q7Z7F0
KHDRBS1 / Q07666
KHDRBS2 / Q5VWX1
KHDRBS3 / O75525
KHK / P50053
KHNYN / O15037
KHSRP / Q92945
KIAA0319L / Q8IZA0
Gene Symbol / UniProt Accession
Brain
Colon
Duodenum
Esophagus
Heart
Liver
Lung
Pancreas
Thyroid
Organs
Type
DDA
DIA
Bins
1
2
3
4
5

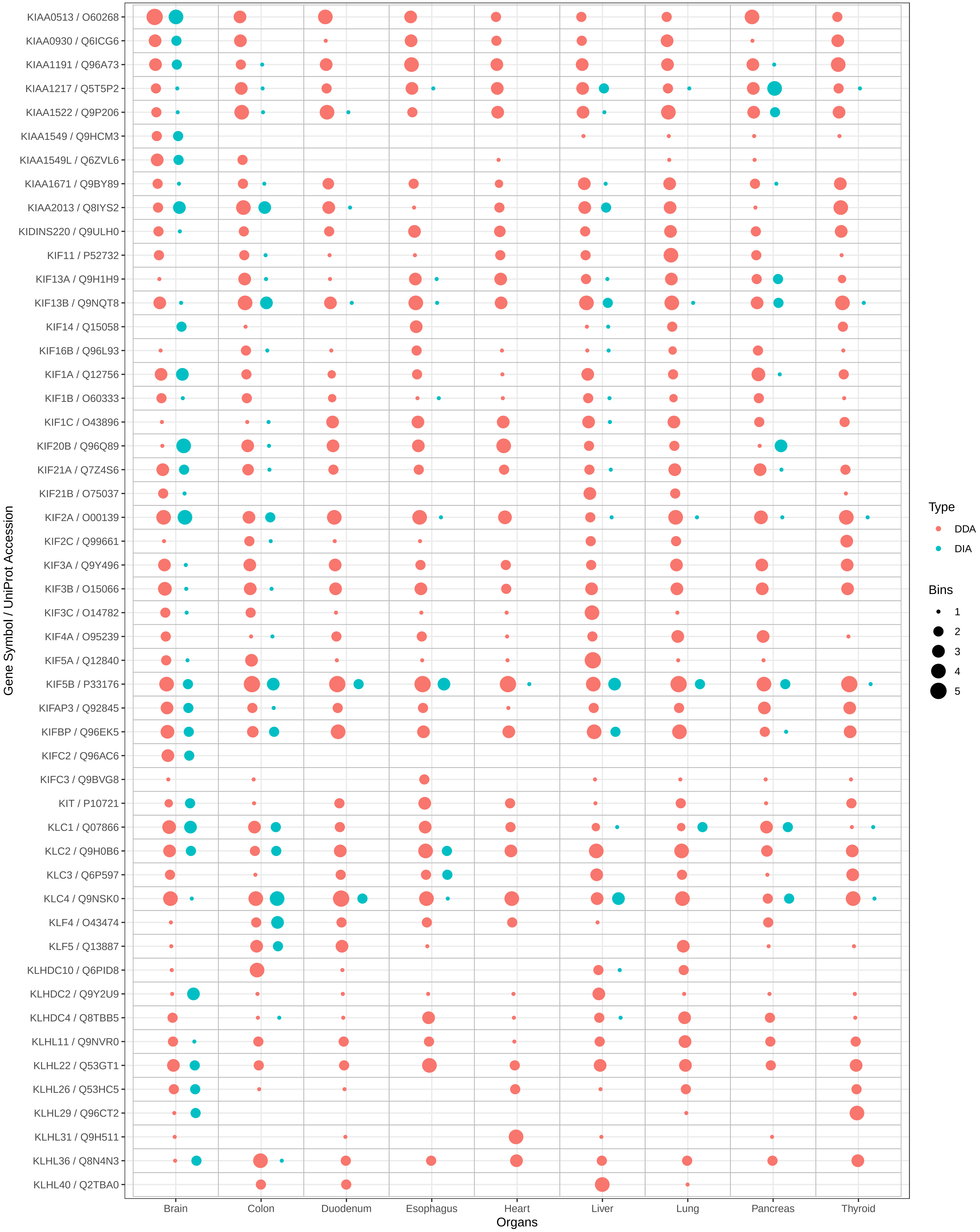
Gene Symbol / UniProt Accession
KIAA0513 / O60268
KIAA0930 / Q6ICG6
KIAA1191 / Q96A73
KIAA1217 / Q5T5P2
KIAA1522 / Q9P206
KIAA1549 / Q9HCM3
KIAA1549L / Q6ZVL6
KIAA1671 / Q9BY89
KIAA2013 / Q8IYS2
KIDINS220 / Q9ULH0
KIF11 / P52732
KIF13A / Q9H1H9
KIF13B / Q9NQT8
KIF14 / Q15058
KIF16B / Q96L93
KIF1A / Q12756
KIF1B / O60333
KIF1C / O43896
KIF20B / Q96Q89
KIF21A / Q7Z4S6
KIF21B / O75037
KIF2A / O00139
KIF2C / Q99661
KIF3A / Q9Y496
KIF3B / O15066
KIF3C / O14782
KIF4A / O95239
KIF5A / Q12840
KIF5B / P33176
KIFAP3 / Q92845
KIFBP / Q96EK5
KIFC2 / Q96AC6
KIFC3 / Q9BVG8
KIT / P10721
KLC1 / Q07866
KLC2 / Q9H0B6
KLC3 / Q6P597
KLC4 / Q9NSK0
KLF4 / O43474
KLF5 / Q13887
KLHDC10 / Q6PID8
KLHDC2 / Q9Y2U9
KLHDC4 / Q8TBB5
KLHL11 / Q9NVR0
KLHL22 / Q53GT1
KLHL26 / Q53HC5
KLHL29 / Q96CT2
KLHL31 / Q9H511
KLHL36 / Q8N4N3
KLHL40 / Q2TBA0
Brain
Colon
Duodenum
Esophagus
Heart
Liver
Lung
Pancreas
Thyroid
Organs
Type
DDA
DIA
Bins
1
2
3
4
5

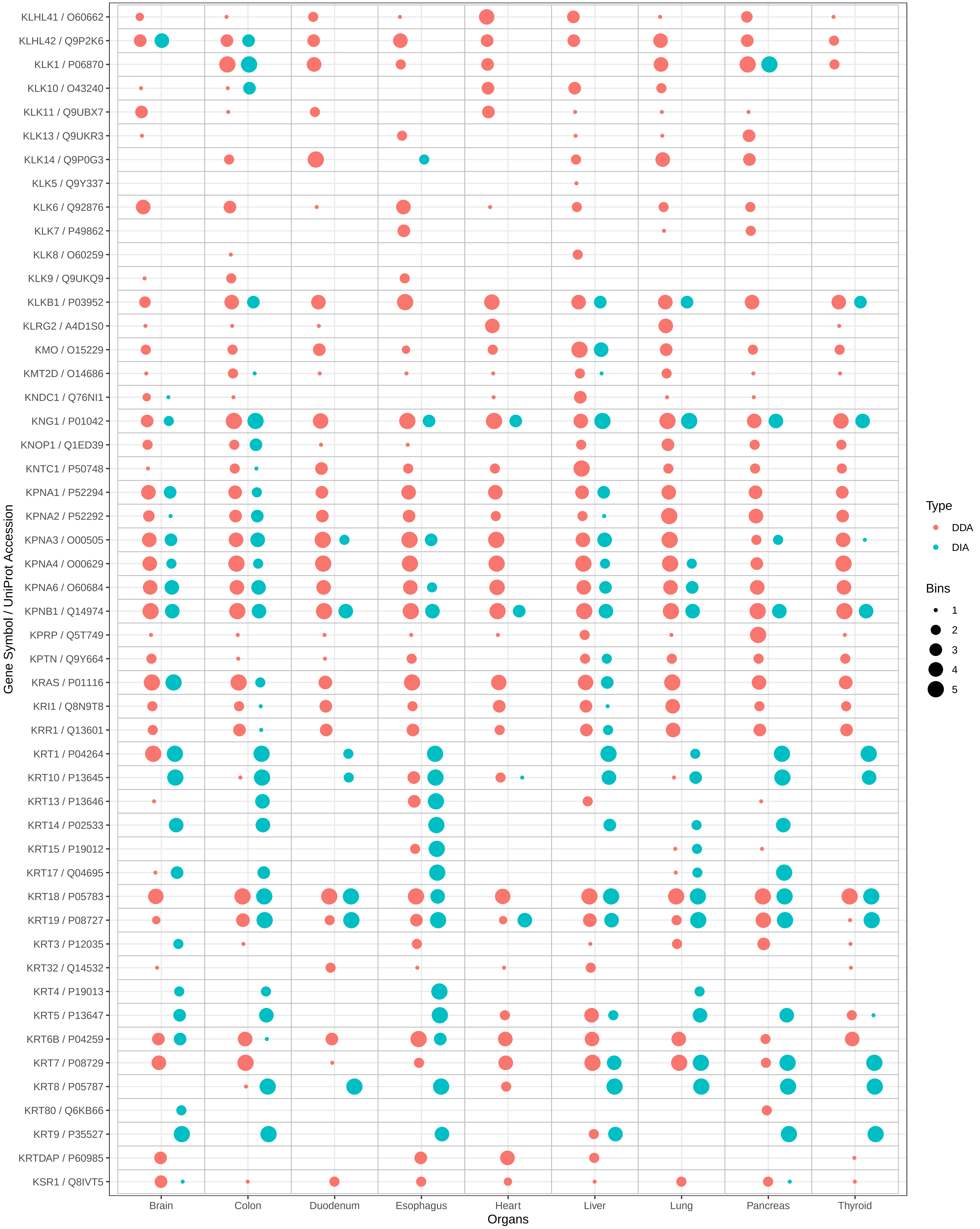
KLHL41 / O60662
KLHL42 / Q9P2K6
KLK1 / P06870
KLK10 / O43240
KLK11 / Q9UBX7
KLK13 / Q9UKR3
KLK14 / Q9P0G3
KLK5 / Q9Y337
KLK6 / Q92876
KLK7 / P49862
KLK8 / O60259
KLK9 / Q9UKQ9
KLKB1 / P03952
KLRG2 / A4D1S0
KMO / O15229
KMT2D / O14686
KNDC1 / Q76NI1
KNG1 / P01042
KNOP1 / Q1ED39
KNTC1 / P50748
KPNA1 / P52294
KPNA2 / P52292
KPNA3 / O00505
KPNA4 / O00629
KPNA6 / O60684
KPNB1 / Q14974
KPRP / Q5T749
KPTN / Q9Y664
KRAS / P01116
KRI1 / Q8N9T8
KRR1 / Q13601
KRT1 / P04264
KRT10 / P13645
KRT13 / P13646
KRT14 / P02533
KRT15 / P19012
KRT17 / Q04695
KRT18 / P05783
KRT19 / P08727
KRT3 / P12035
KRT32 / Q14532
KRT4 / P19013
KRT5 / P13647
KRT6B / P04259
KRT7 / P08729
KRT8 / P05787
KRT80 / Q6KB66
KRT9 / P35527
KRTDAP / P60985
KSR1 / Q8IVT5
Gene Symbol / UniProt Accession
Brain
Colon
Duodenum
Esophagus
Heart
Liver
Lung
Pancreas
Thyroid
Organs
Type
DDA
DIA
Bins
1
2
3
4
5

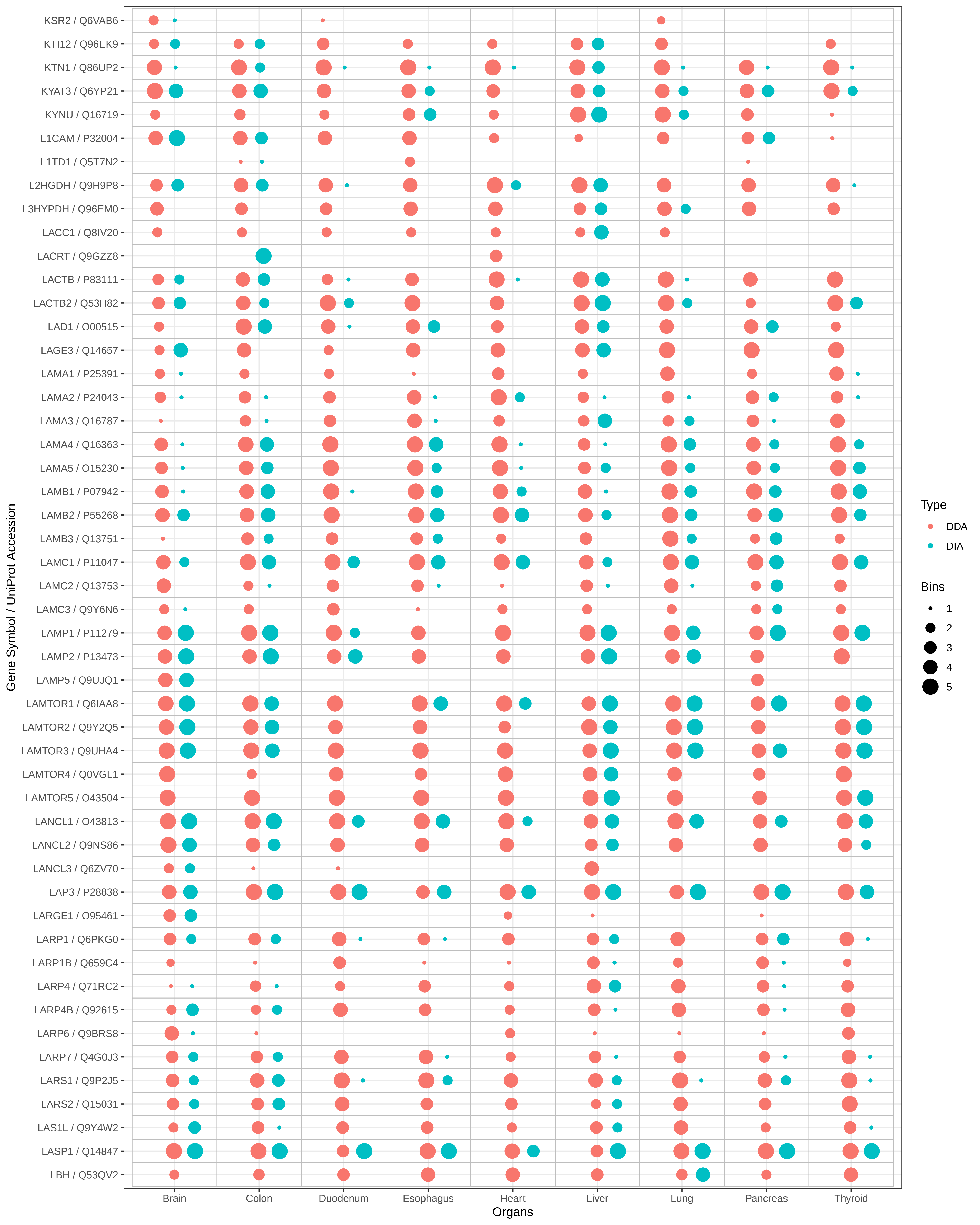

KSR2 / Q6VAB6
KTI12 / Q96EK9
KTN1 / Q86UP2
KYAT3 / Q6YP21
KYNU / Q16719
L1CAM / P32004
L1TD1 / Q5T7N2
L2HGDH / Q9H9P8
L3HYPDH / Q96EM0
LACC1 / Q8IV20
LACRT / Q9GZZ8
LACTB / P83111
LACTB2 / Q53H82
LAD1 / O00515
LAGE3 / Q14657
LAMA1 / P25391
LAMA2 / P24043
LAMA3 / Q16787
LAMA4 / Q16363
LAMA5 / O15230
LAMB1 / P07942
LAMB2 / P55268
LAMB3 / Q13751
LAMC1 / P11047
LAMC2 / Q13753
LAMC3 / Q9Y6N6
LAMP1 / P11279
LAMP2 / P13473
LAMP5 / Q9UJQ1
LAMTOR1 / Q6IAA8
LAMTOR2 / Q9Y2Q5
LAMTOR3 / Q9UHA4
LAMTOR4 / Q0VGL1
LAMTOR5 / O43504
LANCL1 / O43813
LANCL2 / Q9NS86
LANCL3 / Q6ZV70
LAP3 / P28838
LARGE1 / O95461
LARP1 / Q6PKG0
LARP1B / Q659C4
LARP4 / Q71RC2
LARP4B / Q92615
LARP6 / Q9BRS8
LARP7 / Q4G0J3
LARS1 / Q9P2J5
LARS2 / Q15031
LAS1L / Q9Y4W2
LASP1 / Q14847
LBH / Q53QV2
Gene Symbol / UniProt Accession
Brain
Colon
Duodenum
Esophagus
Heart
Liver
Lung
Pancreas
Thyroid
Organs
Type
DDA
DIA
Bins
1
2
3
4
5

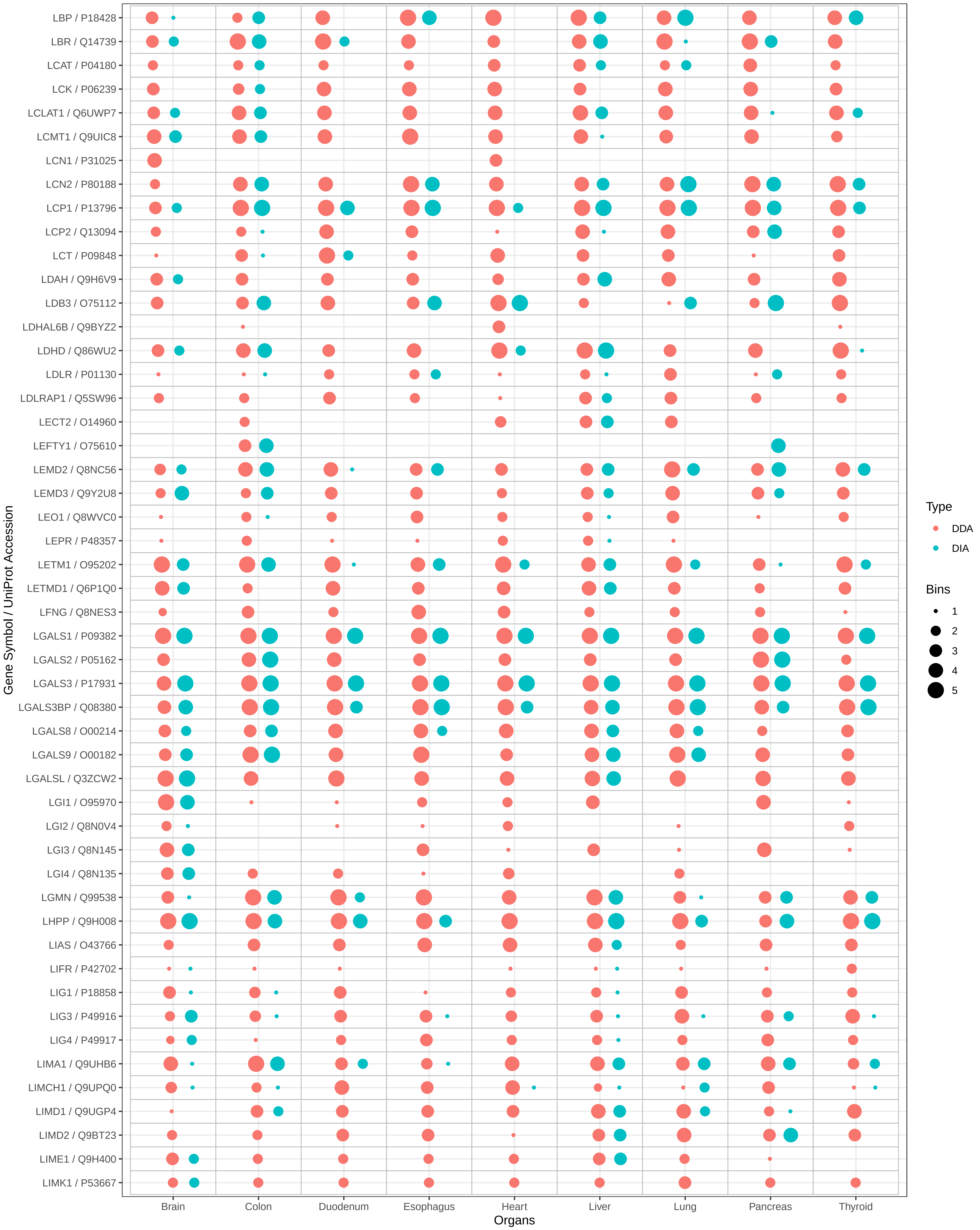

LBP / P18428
LBR / Q14739
LCAT / P04180
LCK / P06239
LCLAT1 / Q6UWP7
LCMT1 / Q9UIC8
LCN1 / P31025
LCN2 / P80188
LCP1 / P13796
LCP2 / Q13094
LCT / P09848
LDAH / Q9H6V9
LDB3 / O75112
LDHAL6B / Q9BYZ2
LDHD / Q86WU2
LDLR / P01130
LDLRAP1 / Q5SW96
LECT2 / O14960
LEFTY1 / O75610
LEMD2 / Q8NC56
LEMD3 / Q9Y2U8
LEO1 / Q8WVC0
LEPR / P48357
LETM1 / O95202
LETMD1 / Q6P1Q0
LFNG / Q8NES3
LGALS1 / P09382
LGALS2 / P05162
LGALS3 / P17931
LGALS3BP / Q08380
LGALS8 / O00214
LGALS9 / O00182
LGALSL / Q3ZCW2
LGI1 / O95970
LGI2 / Q8N0V4
LGI3 / Q8N145
LGI4 / Q8N135
LGMN / Q99538
LHPP / Q9H008
LIAS / O43766
LIFR / P42702
LIG1 / P18858
LIG3 / P49916
LIG4 / P49917
LIMA1 / Q9UHB6
LIMCH1 / Q9UPQ0
LIMD1 / Q9UGP4
LIMD2 / Q9BT23
LIME1 / Q9H400
LIMK1 / P53667
Gene Symbol / UniProt Accession
Brain
Colon
Duodenum
Esophagus
Heart
Liver
Lung
Pancreas
Thyroid
Organs
Type
DDA
DIA
Bins
1
2
3
4
5

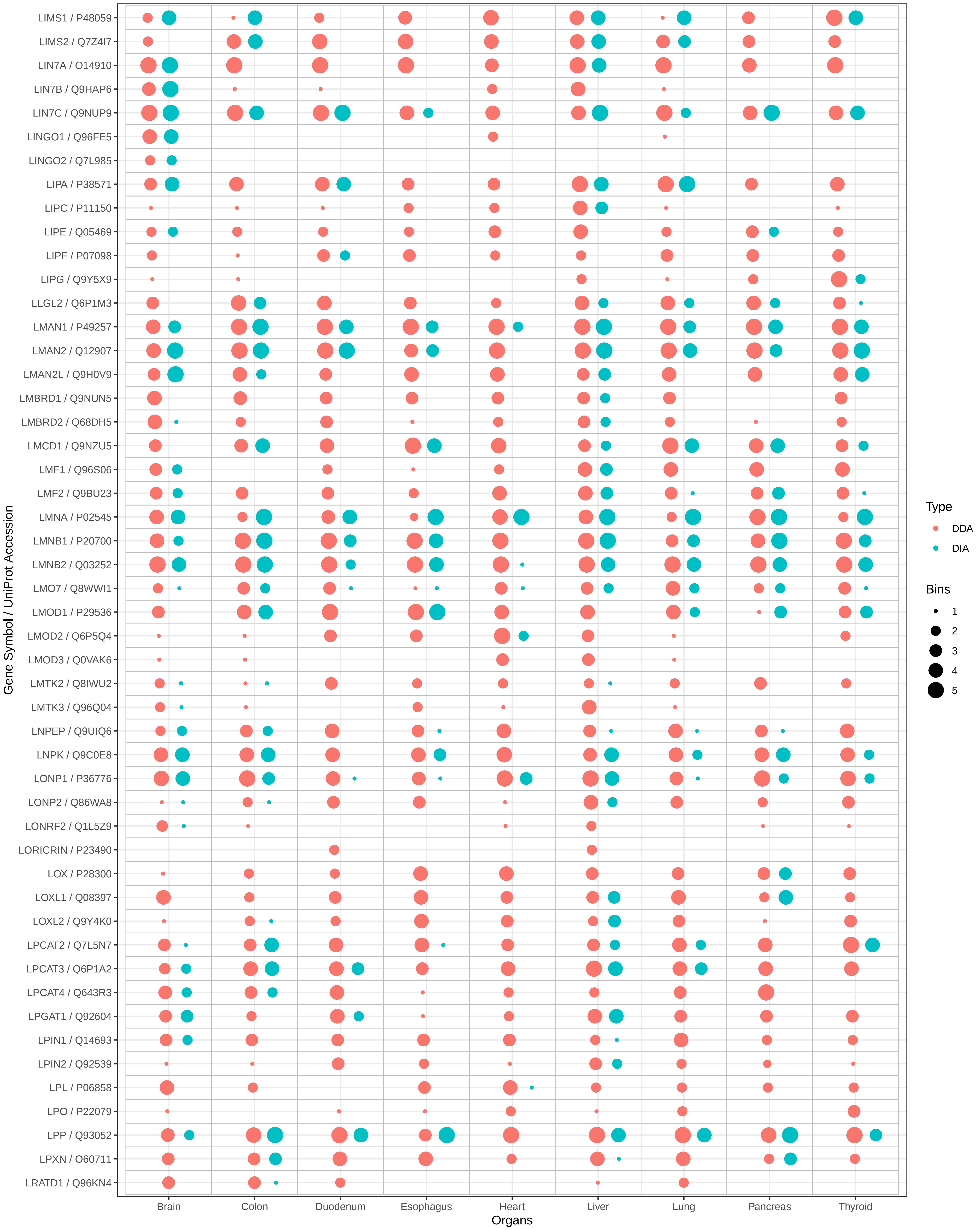

LIMS1 / P48059
LIMS2 / Q7Z4I7
LIN7A / O14910
LIN7B / Q9HAP6
LIN7C / Q9NUP9
LINGO1 / Q96FE5
LINGO2 / Q7L985
LIPA / P38571
LIPC / P11150
LIPE / Q05469
LIPF / P07098
LIPG / Q9Y5X9
LLGL2 / Q6P1M3
LMAN1 / P49257
LMAN2 / Q12907
LMAN2L / Q9H0V9
LMBRD1 / Q9NUN5
LMBRD2 / Q68DH5
LMCD1 / Q9NZU5
LMF1 / Q96S06
LMF2 / Q9BU23
LMNA / P02545
LMNB1 / P20700
LMNB2 / Q03252
LMO7 / Q8WWI1
LMOD1 / P29536
LMOD2 / Q6P5Q4
LMOD3 / Q0VAK6
LMTK2 / Q8IWU2
LMTK3 / Q96Q04
LNPEP / Q9UIQ6
LNPK / Q9C0E8
LONP1 / P36776
LONP2 / Q86WA8
LONRF2 / Q1L5Z9
LORICRIN / P23490
LOX / P28300
LOXL1 / Q08397
LOXL2 / Q9Y4K0
LPCAT2 / Q7L5N7
LPCAT3 / Q6P1A2
LPCAT4 / Q643R3
LPGAT1 / Q92604
LPIN1 / Q14693
LPIN2 / Q92539
LPL / P06858
LPO / P22079
LPP / Q93052
LPXN / O60711
LRATD1 / Q96KN4
Gene Symbol / UniProt Accession
Brain
Colon
Duodenum
Esophagus
Heart
Liver
Lung
Pancreas
Thyroid
Organs
Type
DDA
DIA
Bins
1
2
3
4
5

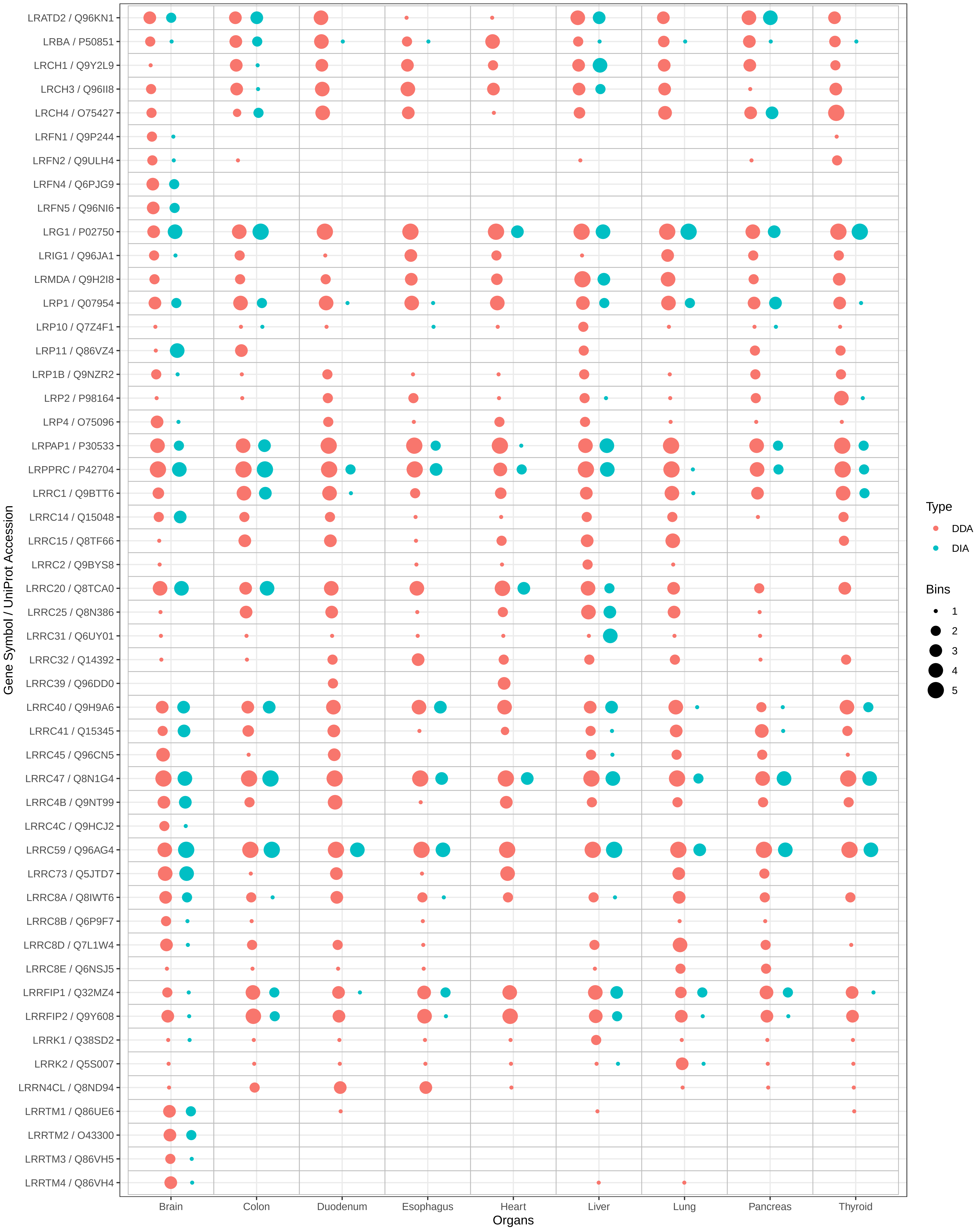

LRATD2 / Q96KN1
LRBA / P50851
LRCH1 / Q9Y2L9
LRCH3 / Q96II8
LRCH4 / O75427
LRFN1 / Q9P244
LRFN2 / Q9ULH4
LRFN4 / Q6PJG9
LRFN5 / Q96NI6
LRG1 / P02750
LRIG1 / Q96JA1
LRMDA / Q9H2I8
LRP1 / Q07954
LRP10 / Q7Z4F1
LRP11 / Q86VZ4
LRP1B / Q9NZR2
LRP2 / P98164
LRP4 / O75096
LRPAP1 / P30533
LRPPRC / P42704
LRRC1 / Q9BTT6
LRRC14 / Q15048
LRRC15 / Q8TF66
LRRC2 / Q9BYS8
LRRC20 / Q8TCA0
LRRC25 / Q8N386
LRRC31 / Q6UY01
LRRC32 / Q14392
LRRC39 / Q96DD0
LRRC40 / Q9H9A6
LRRC41 / Q15345
LRRC45 / Q96CN5
LRRC47 / Q8N1G4
LRRC4B / Q9NT99
LRRC4C / Q9HCJ2
LRRC59 / Q96AG4
LRRC73 / Q5JTD7
LRRC8A / Q8IWT6
LRRC8B / Q6P9F7
LRRC8D / Q7L1W4
LRRC8E / Q6NSJ5
LRRFIP1 / Q32MZ4
LRRFIP2 / Q9Y608
LRRK1 / Q38SD2
LRRK2 / Q5S007
LRRN4CL / Q8ND94
LRRTM1 / Q86UE6
LRRTM2 / O43300
LRRTM3 / Q86VH5
LRRTM4 / Q86VH4
Gene Symbol / UniProt Accession
Brain
Colon
Duodenum
Esophagus
Heart
Liver
Lung
Pancreas
Thyroid
Organs
Type
DDA
DIA
Bins
1
2
3
4
5

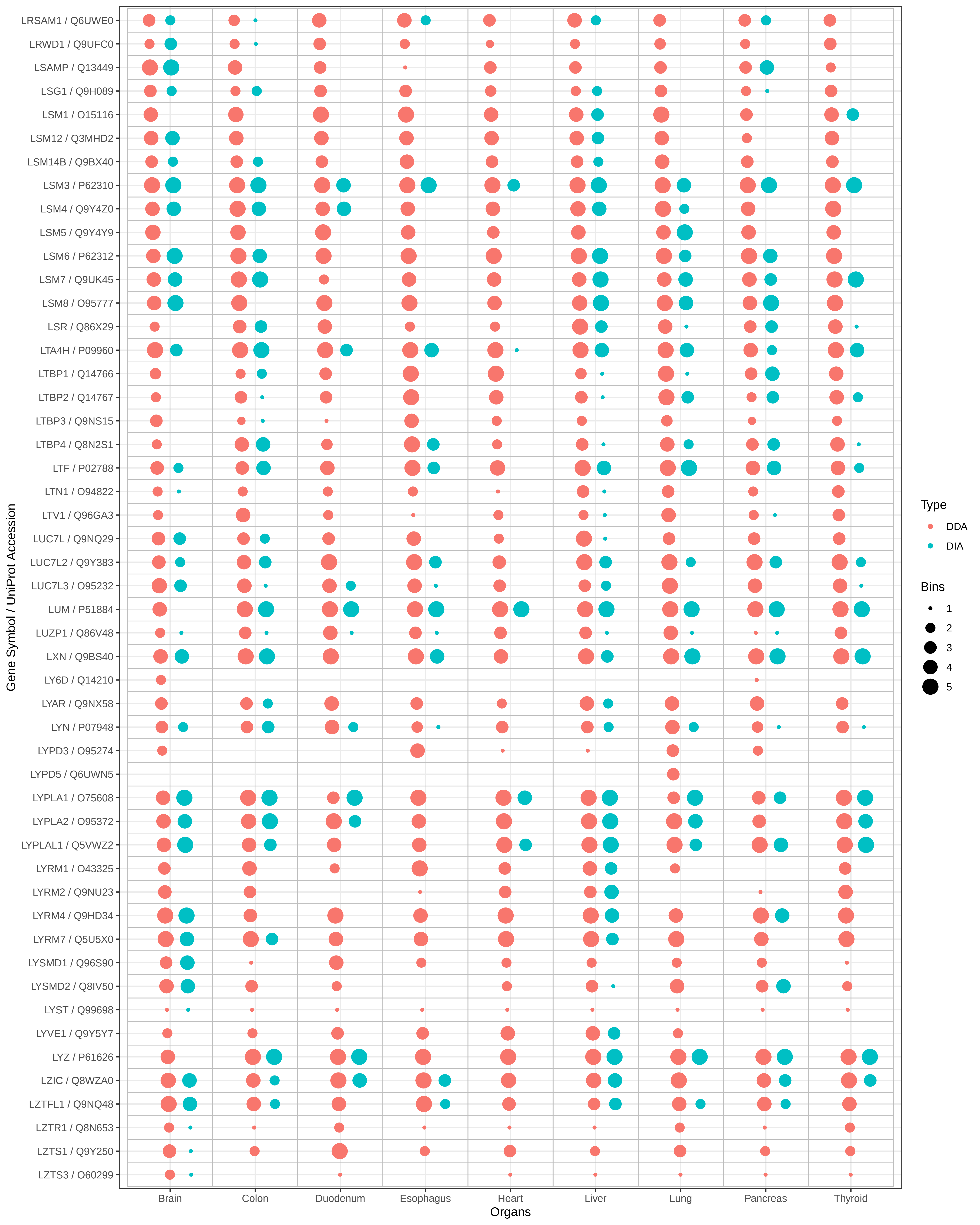
LRSAM1 / Q6UWE0
LRWD1 / Q9UFC0
LSAMP / Q13449
LSG1 / Q9H089
LSM1 / O15116
LSM12 / Q3MHD2
LSM14B / Q9BX40
LSM3 / P62310
LSM4 / Q9Y4Z0
LSM5 / Q9Y4Y9
LSM6 / P62312
LSM7 / Q9UK45
LSM8 / O95777
LSR / Q86X29
LTA4H / P09960
LTBP1 / Q14766
LTBP2 / Q14767
LTBP3 / Q9NS15
LTBP4 / Q8N2S1
LTF / P02788
LTN1 / O94822
LTV1 / Q96GA3
LUC7L / Q9NQ29
LUC7L2 / Q9Y383
LUC7L3 / O95232
LUM / P51884
LUZP1 / Q86V48
LXN / Q9BS40
LY6D / Q14210
LYAR / Q9NX58
LYN / P07948
LYPD3 / O95274
LYPD5 / Q6UWN5
LYPLA1 / O75608
LYPLA2 / O95372
LYPLAL1 / Q5VWZ2
LYRM1 / O43325
LYRM2 / Q9NU23
LYRM4 / Q9HD34
LYRM7 / Q5U5X0
LYSMD1 / Q96S90
LYSMD2 / Q8IV50
LYST / Q99698
LYVE1 / Q9Y5Y7
LYZ / P61626
LZIC / Q8WZA0
LZTFL1 / Q9NQ48
LZTR1 / Q8N653
LZTS1 / Q9Y250
LZTS3 / O60299
Gene Symbol / UniProt Accession
Brain
Colon
Duodenum
Esophagus
Heart
Liver
Lung
Pancreas
Thyroid
Organs
Type
DDA
DIA
Bins
1
2
3
4
5

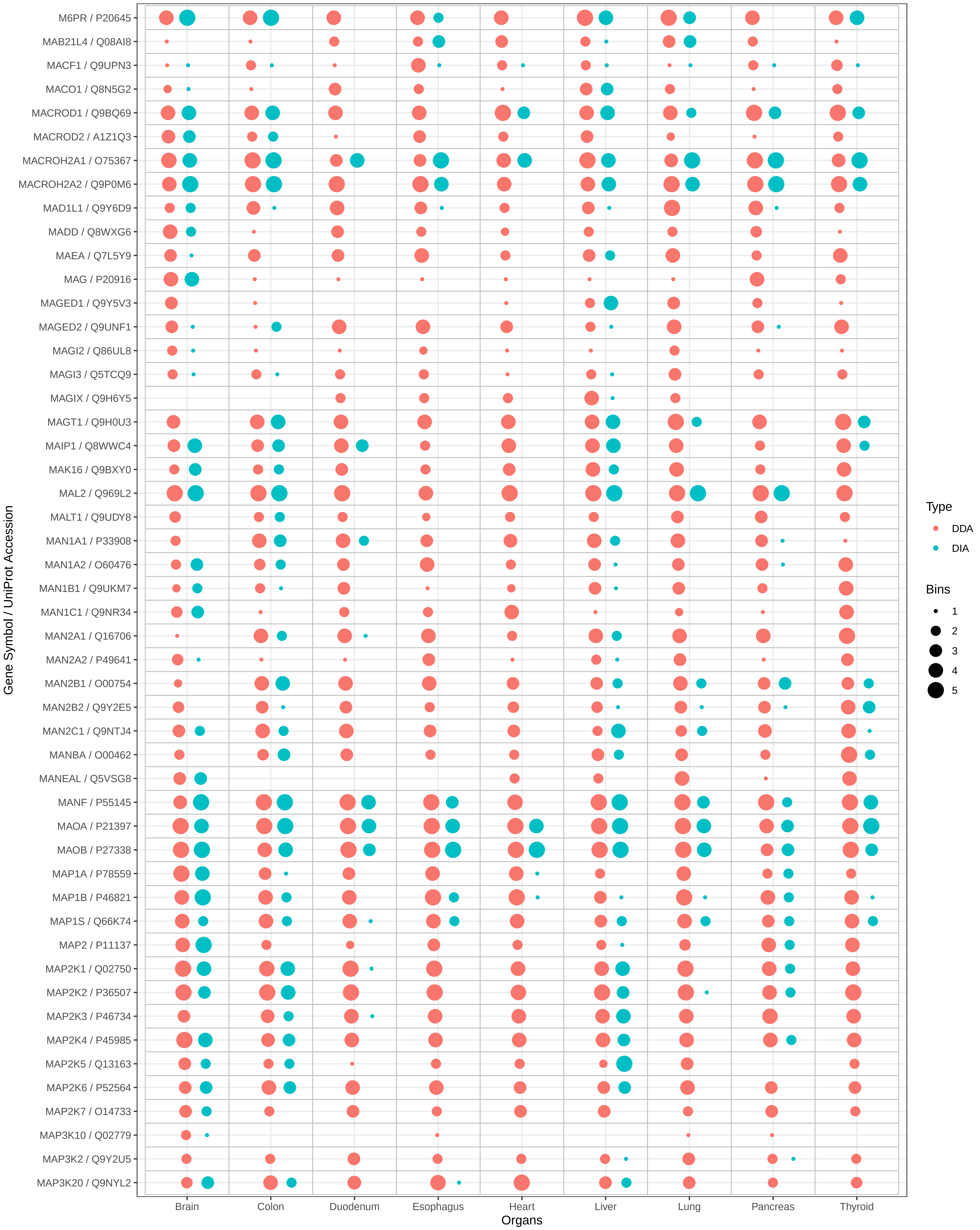

M6PR / P20645
MAB21L4 / Q08AI8
MACF1 / Q9UPN3
MACO1 / Q8N5G2
MACROD1 / Q9BQ69
MACROD2 / A1Z1Q3
MACROH2A1 / O75367
MACROH2A2 / Q9P0M6
MAD1L1 / Q9Y6D9
MADD / Q8WXG6
MAEA / Q7L5Y9
MAG / P20916
MAGED1 / Q9Y5V3
MAGED2 / Q9UNF1
MAGI2 / Q86UL8
MAGI3 / Q5TCQ9
MAGIX / Q9H6Y5
MAGT1 / Q9H0U3
MAIP1 / Q8WWC4
MAK16 / Q9BXY0
MAL2 / Q969L2
MALT1 / Q9UDY8
MAN1A1 / P33908
MAN1A2 / O60476
MAN1B1 / Q9UKM7
MAN1C1 / Q9NR34
MAN2A1 / Q16706
MAN2A2 / P49641
MAN2B1 / O00754
MAN2B2 / Q9Y2E5
MAN2C1 / Q9NTJ4
MANBA / O00462
MANEAL / Q5VSG8
MANF / P55145
MAOA / P21397
MAOB / P27338
MAP1A / P78559
MAP1B / P46821
MAP1S / Q66K74
MAP2 / P11137
MAP2K1 / Q02750
MAP2K2 / P36507
MAP2K3 / P46734
MAP2K4 / P45985
MAP2K5 / Q13163
MAP2K6 / P52564
MAP2K7 / O14733
MAP3K10 / Q02779
MAP3K2 / Q9Y2U5
MAP3K20 / Q9NYL2
Gene Symbol / UniProt Accession
Brain
Colon
Duodenum
Esophagus
Heart
Liver
Lung
Pancreas
Thyroid
Organs
Type
DDA
DIA
Bins
1
2
3
4
5

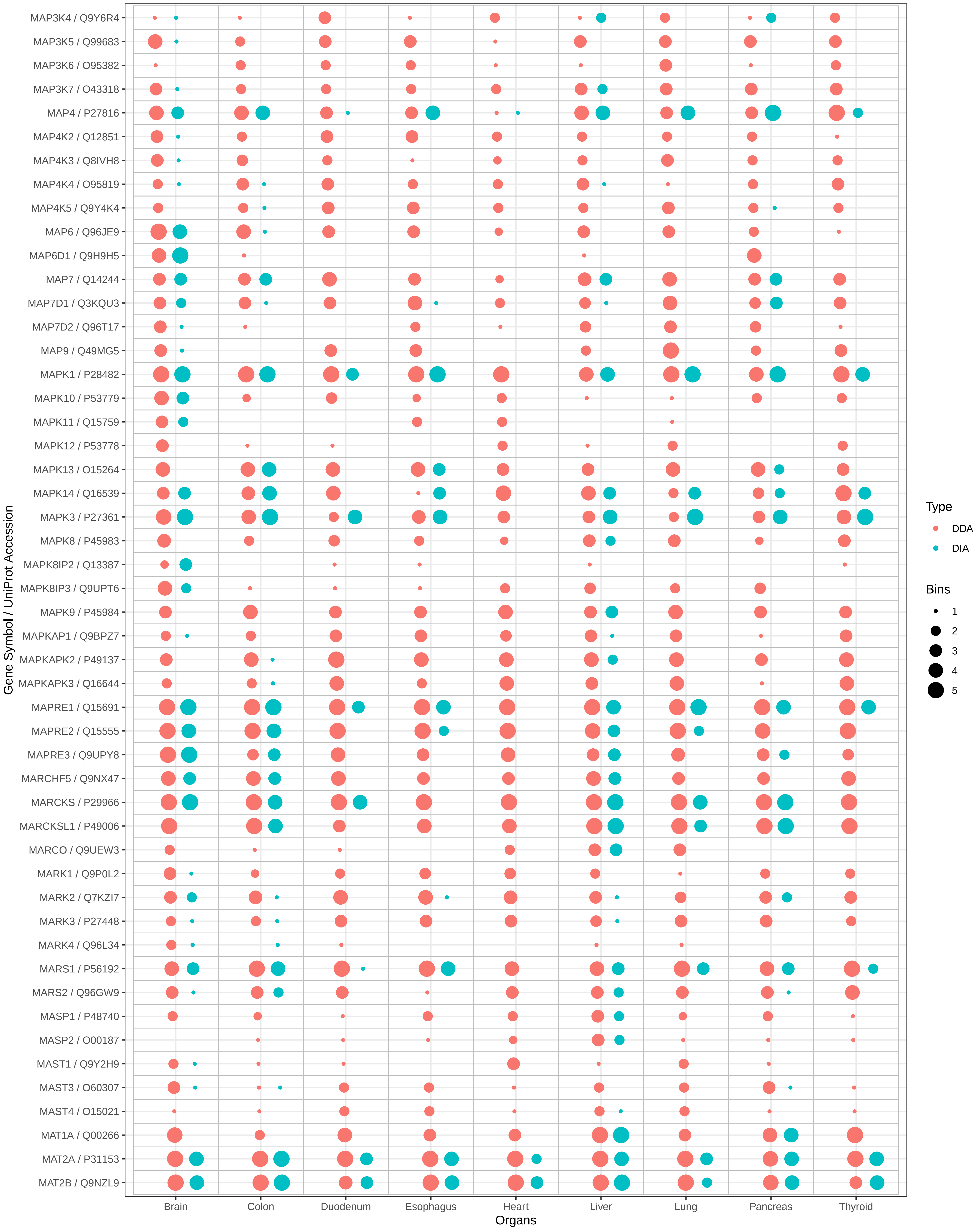

MAP3K4 / Q9Y6R4
MAP3K5 / Q99683
MAP3K6 / O95382
MAP3K7 / O43318
MAP4 / P27816
MAP4K2 / Q12851
MAP4K3 / Q8IVH8
MAP4K4 / O95819
MAP4K5 / Q9Y4K4
MAP6 / Q96JE9
MAP6D1 / Q9H9H5
MAP7 / Q14244
MAP7D1 / Q3KQU3
MAP7D2 / Q96T17
MAP9 / Q49MG5
MAPK1 / P28482
MAPK10 / P53779
MAPK11 / Q15759
MAPK12 / P53778
MAPK13 / O15264
MAPK14 / Q16539
MAPK3 / P27361
MAPK8 / P45983
MAPK8IP2 / Q13387
MAPK8IP3 / Q9UPT6
MAPK9 / P45984
MAPKAP1 / Q9BPZ7
MAPKAPK2 / P49137
MAPKAPK3 / Q16644
MAPRE1 / Q15691
MAPRE2 / Q15555
MAPRE3 / Q9UPY8
MARCHF5 / Q9NX47
MARCKS / P29966
MARCKSL1 / P49006
MARCO / Q9UEW3
MARK1 / Q9P0L2
MARK2 / Q7KZI7
MARK3 / P27448
MARK4 / Q96L34
MARS1 / P56192
MARS2 / Q96GW9
MASP1 / P48740
MASP2 / O00187
MAST1 / Q9Y2H9
MAST3 / O60307
MAST4 / O15021
MAT1A / Q00266
MAT2A / P31153
MAT2B / Q9NZL9
Gene Symbol / UniProt Accession
Brain
Colon
Duodenum
Esophagus
Heart
Liver
Lung
Pancreas
Thyroid
Organs
Type
DDA
DIA
Bins
1
2
3
4
5

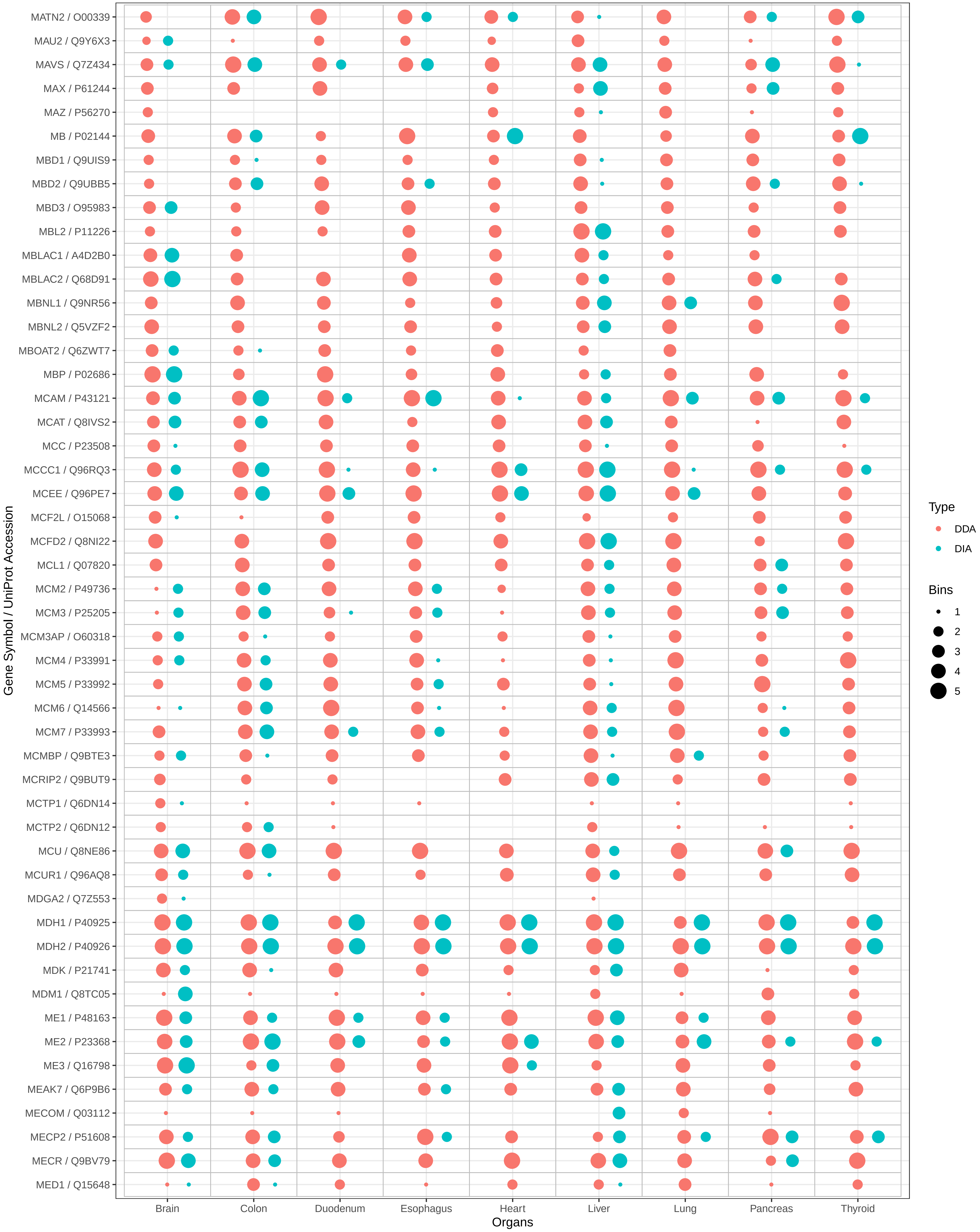

MATN2 / O00339
MAU2 / Q9Y6X3
MAVS / Q7Z434
MAX / P61244
MAZ / P56270
MB / P02144
MBD1 / Q9UIS9
MBD2 / Q9UBB5
MBD3 / O95983
MBL2 / P11226
MBLAC1 / A4D2B0
MBLAC2 / Q68D91
MBNL1 / Q9NR56
MBNL2 / Q5VZF2
MBOAT2 / Q6ZWT7
MBP / P02686
MCAM / P43121
MCAT / Q8IVS2
MCC / P23508
MCCC1 / Q96RQ3
MCEE / Q96PE7
MCF2L / O15068
MCFD2 / Q8NI22
MCL1 / Q07820
MCM2 / P49736
MCM3 / P25205
MCM3AP / O60318
MCM4 / P33991
MCM5 / P33992
MCM6 / Q14566
MCM7 / P33993
MCMBP / Q9BTE3
MCRIP2 / Q9BUT9
MCTP1 / Q6DN14
MCTP2 / Q6DN12
MCU / Q8NE86
MCUR1 / Q96AQ8
MDGA2 / Q7Z553
MDH1 / P40925
MDH2 / P40926
MDK / P21741
MDM1 / Q8TC05
ME1 / P48163
ME2 / P23368
ME3 / Q16798
MEAK7 / Q6P9B6
MECOM / Q03112
MECP2 / P51608
MECR / Q9BV79
MED1 / Q15648
Gene Symbol / UniProt Accession
Brain
Colon
Duodenum
Esophagus
Heart
Liver
Lung
Pancreas
Thyroid
Organs
Type
DDA
DIA
Bins
1
2
3
4
5

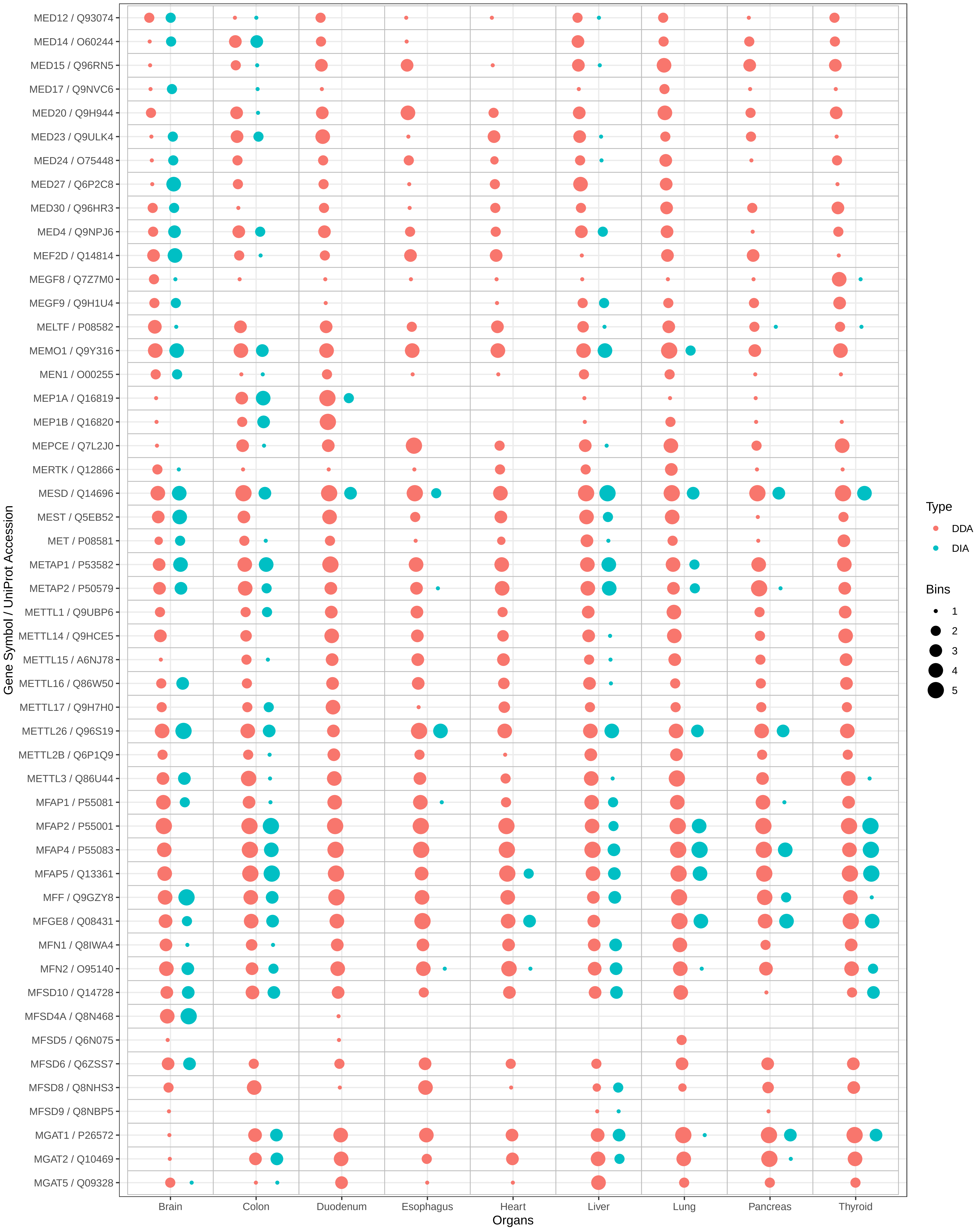
MED12 / Q93074
MED14 / O60244
MED15 / Q96RN5
MED17 / Q9NVC6
MED20 / Q9H944
MED23 / Q9ULK4
MED24 / O75448
MED27 / Q6P2C8
MED30 / Q96HR3
MED4 / Q9NPJ6
MEF2D / Q14814
MEGF8 / Q7Z7M0
MEGF9 / Q9H1U4
MELTF / P08582
MEMO1 / Q9Y316
MEN1 / O00255
MEP1A / Q16819
MEP1B / Q16820
MEPCE / Q7L2J0
MERTK / Q12866
MESD / Q14696
MEST / Q5EB52
MET / P08581
METAP1 / P53582
METAP2 / P50579
METTL1 / Q9UBP6
METTL14 / Q9HCE5
METTL15 / A6NJ78
METTL16 / Q86W50
METTL17 / Q9H7H0
METTL26 / Q96S19
METTL2B / Q6P1Q9
METTL3 / Q86U44
MFAP1 / P55081
MFAP2 / P55001
MFAP4 / P55083
MFAP5 / Q13361
MFF / Q9GZY8
MFGE8 / Q08431
MFN1 / Q8IWA4
MFN2 / O95140
MFSD10 / Q14728
MFSD4A / Q8N468
MFSD5 / Q6N075
MFSD6 / Q6ZSS7
MFSD8 / Q8NHS3
MFSD9 / Q8NBP5
MGAT1 / P26572
MGAT2 / Q10469
MGAT5 / Q09328
Gene Symbol / UniProt Accession
Brain
Colon
Duodenum
Esophagus
Heart
Liver
Lung
Pancreas
Thyroid
Organs
Type
DDA
DIA
Bins
1
2
3
4
5

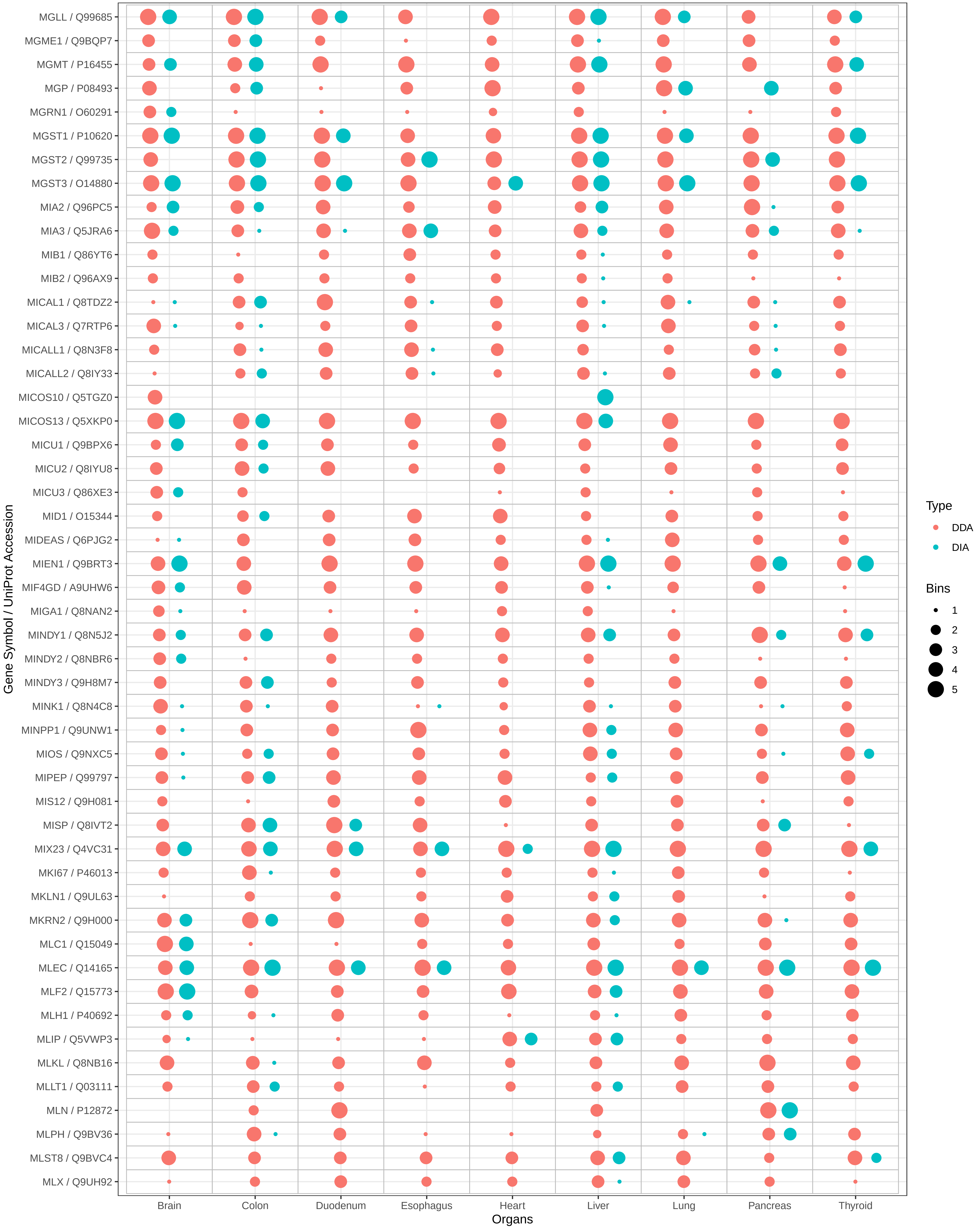

MGLL / Q99685
MGME1 / Q9BQP7
MGMT / P16455
MGP / P08493
MGRN1 / O60291
MGST1 / P10620
MGST2 / Q99735
MGST3 / O14880
MIA2 / Q96PC5
MIA3 / Q5JRA6
MIB1 / Q86YT6
MIB2 / Q96AX9
MICAL1 / Q8TDZ2
MICAL3 / Q7RTP6
MICALL1 / Q8N3F8
MICALL2 / Q8IY33
MICOS10 / Q5TGZ0
MICOS13 / Q5XKP0
MICU1 / Q9BPX6
MICU2 / Q8IYU8
MICU3 / Q86XE3
MID1 / O15344
MIDEAS / Q6PJG2
MIEN1 / Q9BRT3
MIF4GD / A9UHW6
MIGA1 / Q8NAN2
MINDY1 / Q8N5J2
MINDY2 / Q8NBR6
MINDY3 / Q9H8M7
MINK1 / Q8N4C8
MINPP1 / Q9UNW1
MIOS / Q9NXC5
MIPEP / Q99797
MIS12 / Q9H081
MISP / Q8IVT2
MIX23 / Q4VC31
MKI67 / P46013
MKLN1 / Q9UL63
MKRN2 / Q9H000
MLC1 / Q15049
MLEC / Q14165
MLF2 / Q15773
MLH1 / P40692
MLIP / Q5VWP3
MLKL / Q8NB16
MLLT1 / Q03111
MLN / P12872
MLPH / Q9BV36
MLST8 / Q9BVC4
MLX / Q9UH92
Gene Symbol / UniProt Accession
Brain
Colon
Duodenum
Esophagus
Heart
Liver
Lung
Pancreas
Thyroid
Organs
Type
DDA
DIA
Bins
1
2
3
4
5

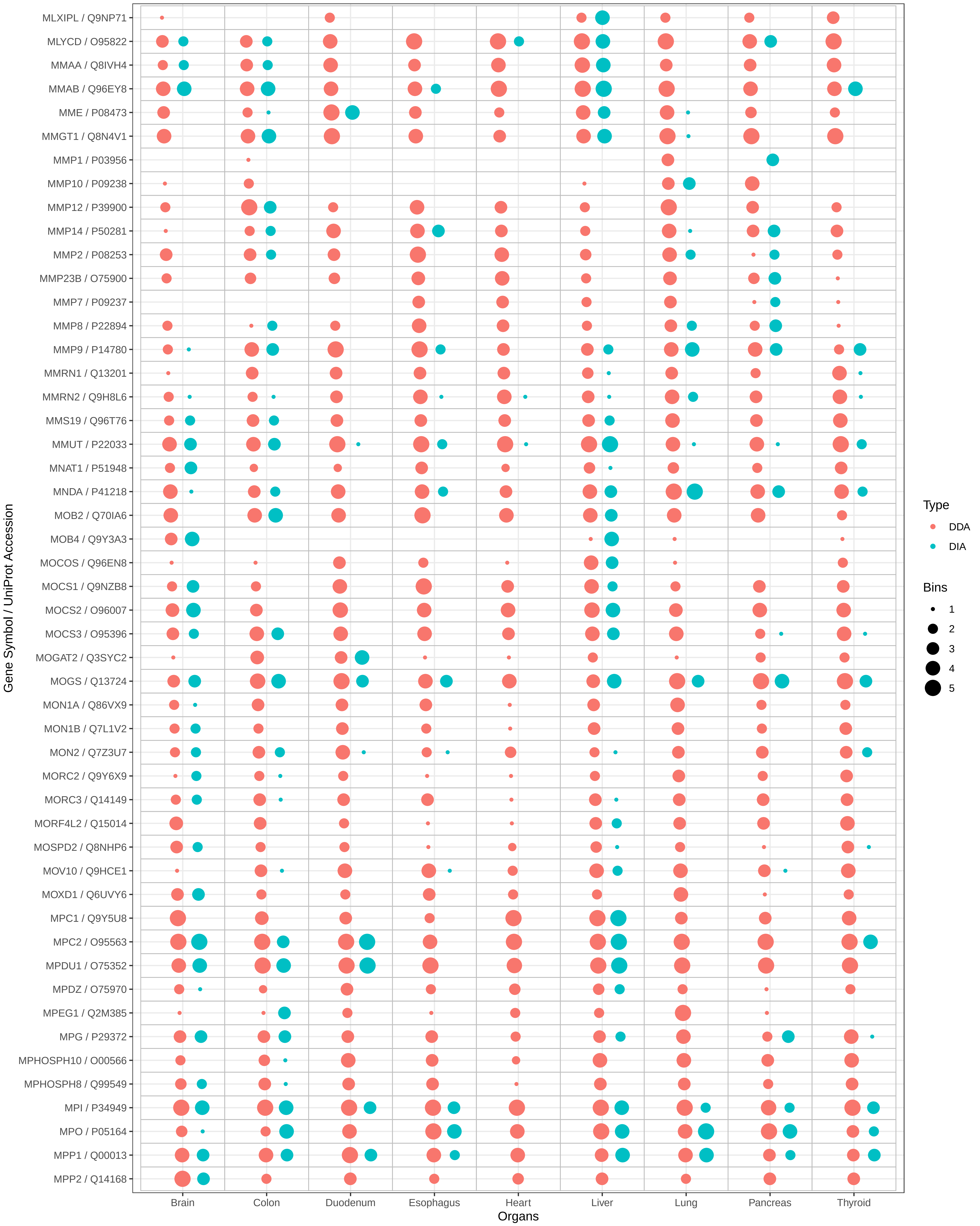

MLXIPL / Q9NP71
MLYCD / O95822
MMAA / Q8IVH4
MMAB / Q96EY8
MME / P08473
MMGT1 / Q8N4V1
MMP1 / P03956
MMP10 / P09238
MMP12 / P39900
MMP14 / P50281
MMP2 / P08253
MMP23B / O75900
MMP7 / P09237
MMP8 / P22894
MMP9 / P14780
MMRN1 / Q13201
MMRN2 / Q9H8L6
MMS19 / Q96T76
MMUT / P22033
MNAT1 / P51948
MNDA / P41218
MOB2 / Q70IA6
MOB4 / Q9Y3A3
MOCOS / Q96EN8
MOCS1 / Q9NZB8
MOCS2 / O96007
MOCS3 / O95396
MOGAT2 / Q3SYC2
MOGS / Q13724
MON1A / Q86VX9
MON1B / Q7L1V2
MON2 / Q7Z3U7
MORC2 / Q9Y6X9
MORC3 / Q14149
MORF4L2 / Q15014
MOSPD2 / Q8NHP6
MOV10 / Q9HCE1
MOXD1 / Q6UVY6
MPC1 / Q9Y5U8
MPC2 / O95563
MPDU1 / O75352
MPDZ / O75970
MPEG1 / Q2M385
MPG / P29372
MPHOSPH10 / O00566
MPHOSPH8 / Q99549
MPI / P34949
MPO / P05164
MPP1 / Q00013
MPP2 / Q14168
Gene Symbol / UniProt Accession
Brain
Colon
Duodenum
Esophagus
Heart
Liver
Lung
Pancreas
Thyroid
Organs
Type
DDA
DIA
Bins
1
2
3
4
5

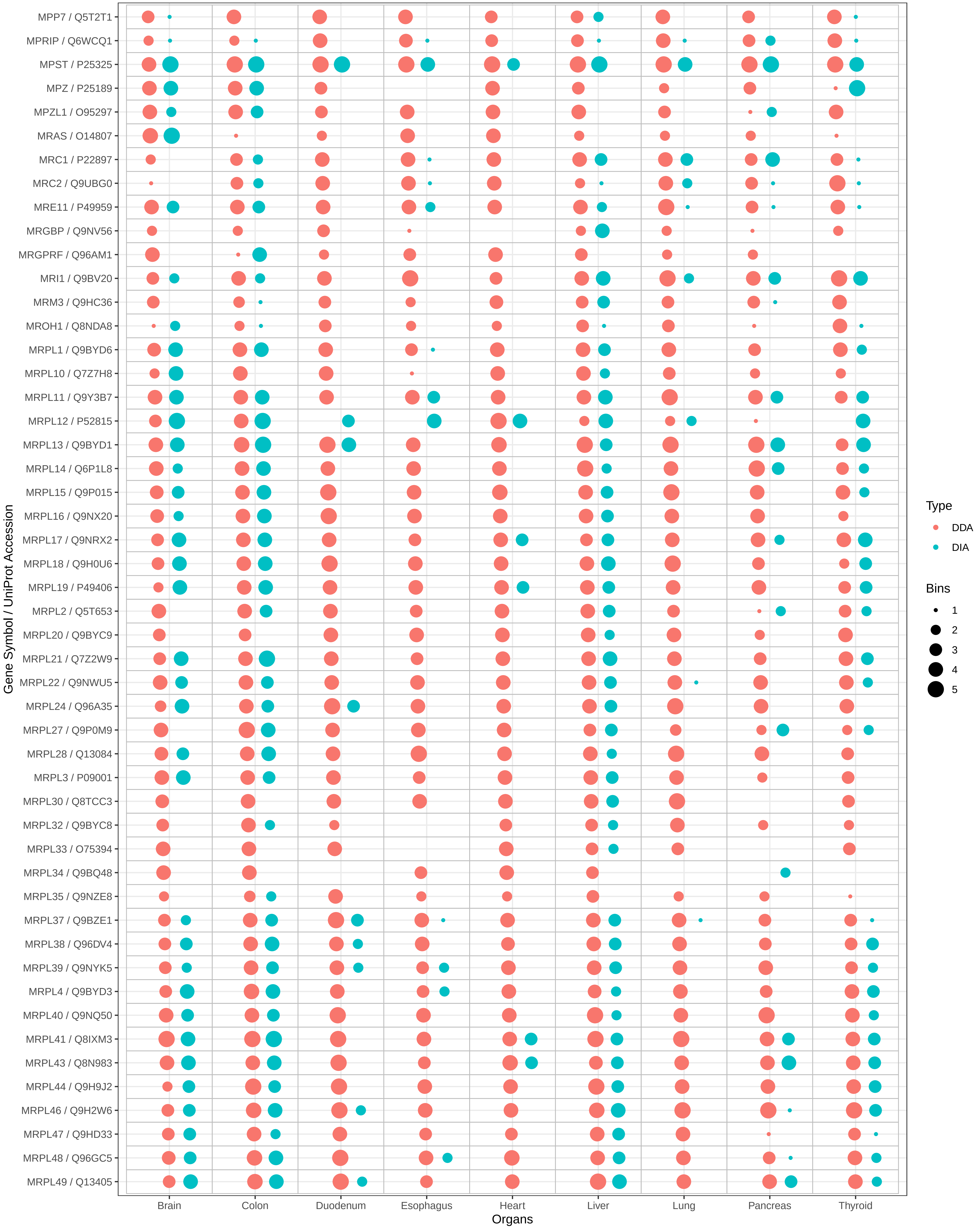

MPP7 / Q5T2T1
MPRIP / Q6WCQ1
MPST / P25325
MPZ / P25189
MPZL1 / O95297
MRAS / O14807
MRC1 / P22897
MRC2 / Q9UBG0
MRE11 / P49959
MRGBP / Q9NV56
MRGPRF / Q96AM1
MRI1 / Q9BV20
MRM3 / Q9HC36
MROH1 / Q8NDA8
MRPL1 / Q9BYD6
MRPL10 / Q7Z7H8
MRPL11 / Q9Y3B7
MRPL12 / P52815
MRPL13 / Q9BYD1
MRPL14 / Q6P1L8
MRPL15 / Q9P015
MRPL16 / Q9NX20
MRPL17 / Q9NRX2
MRPL18 / Q9H0U6
MRPL19 / P49406
MRPL2 / Q5T653
MRPL20 / Q9BYC9
MRPL21 / Q7Z2W9
MRPL22 / Q9NWU5
MRPL24 / Q96A35
MRPL27 / Q9P0M9
MRPL28 / Q13084
MRPL3 / P09001
MRPL30 / Q8TCC3
MRPL32 / Q9BYC8
MRPL33 / O75394
MRPL34 / Q9BQ48
MRPL35 / Q9NZE8
MRPL37 / Q9BZE1
MRPL38 / Q96DV4
MRPL39 / Q9NYK5
MRPL4 / Q9BYD3
MRPL40 / Q9NQ50
MRPL41 / Q8IXM3
MRPL43 / Q8N983
MRPL44 / Q9H9J2
MRPL46 / Q9H2W6
MRPL47 / Q9HD33
MRPL48 / Q96GC5
MRPL49 / Q13405
Gene Symbol / UniProt Accession
Brain
Colon
Duodenum
Esophagus
Heart
Liver
Lung
Pancreas
Thyroid
Organs
Type
DDA
DIA
Bins
1
2
3
4
5

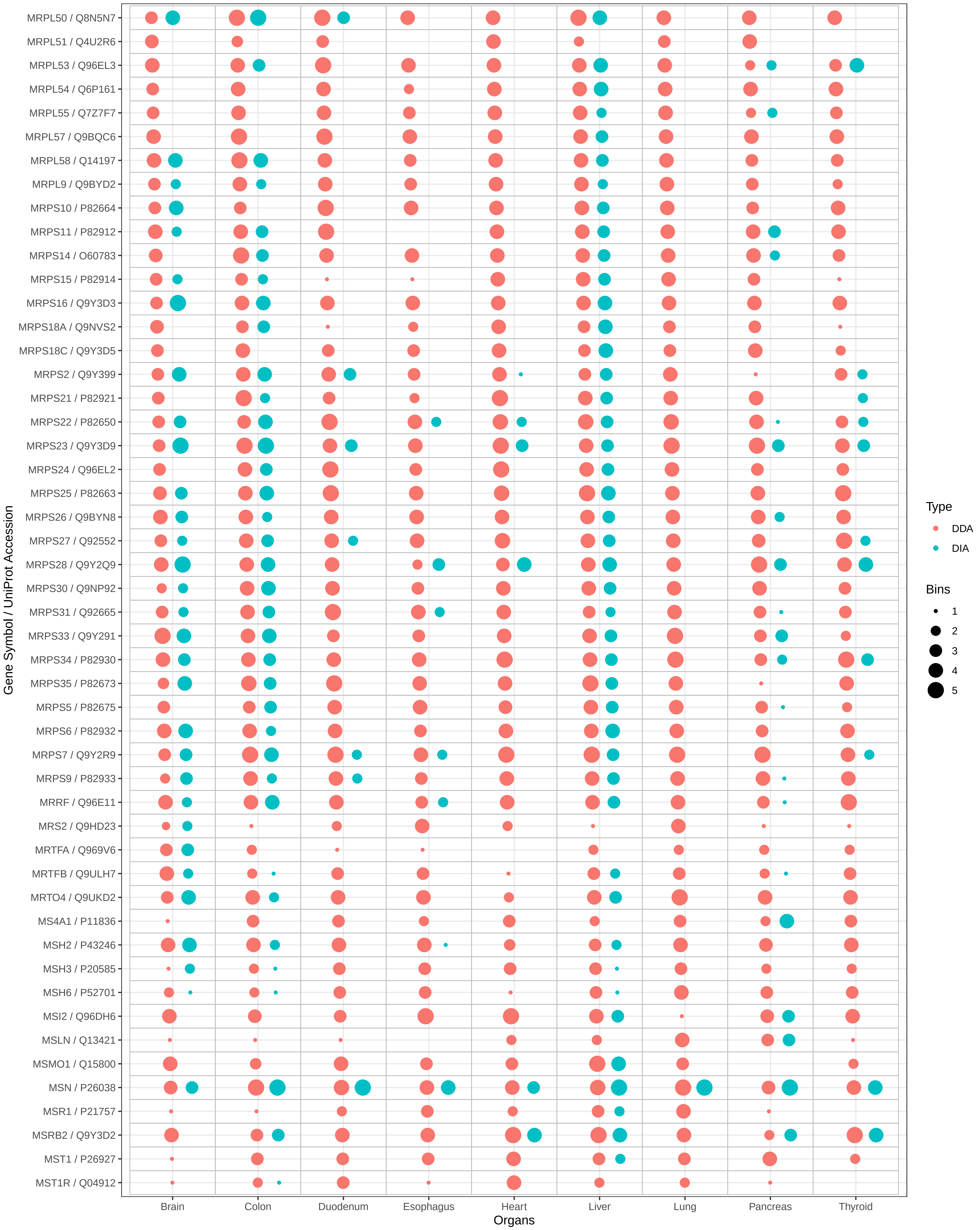

MRPL50 / Q8N5N7
MRPL51 / Q4U2R6
MRPL53 / Q96EL3
MRPL54 / Q6P161
MRPL55 / Q7Z7F7
MRPL57 / Q9BQC6
MRPL58 / Q14197
MRPL9 / Q9BYD2
MRPS10 / P82664
MRPS11 / P82912
MRPS14 / O60783
MRPS15 / P82914
MRPS16 / Q9Y3D3
MRPS18A / Q9NVS2
MRPS18C / Q9Y3D5
MRPS2 / Q9Y399
MRPS21 / P82921
MRPS22 / P82650
MRPS23 / Q9Y3D9
MRPS24 / Q96EL2
MRPS25 / P82663
MRPS26 / Q9BYN8
MRPS27 / Q92552
MRPS28 / Q9Y2Q9
MRPS30 / Q9NP92
MRPS31 / Q92665
MRPS33 / Q9Y291
MRPS34 / P82930
MRPS35 / P82673
MRPS5 / P82675
MRPS6 / P82932
MRPS7 / Q9Y2R9
MRPS9 / P82933
MRRF / Q96E11
MRS2 / Q9HD23
MRTFA / Q969V6
MRTFB / Q9ULH7
MRTO4 / Q9UKD2
MS4A1 / P11836
MSH2 / P43246
MSH3 / P20585
MSH6 / P52701
MSI2 / Q96DH6
MSLN / Q13421
MSMO1 / Q15800
MSN / P26038
MSR1 / P21757
MSRB2 / Q9Y3D2
MST1 / P26927
MST1R / Q04912
Gene Symbol / UniProt Accession
Brain
Colon
Duodenum
Esophagus
Heart
Liver
Lung
Pancreas
Thyroid
Organs
Type
DDA
DIA
Bins
1
2
3
4
5

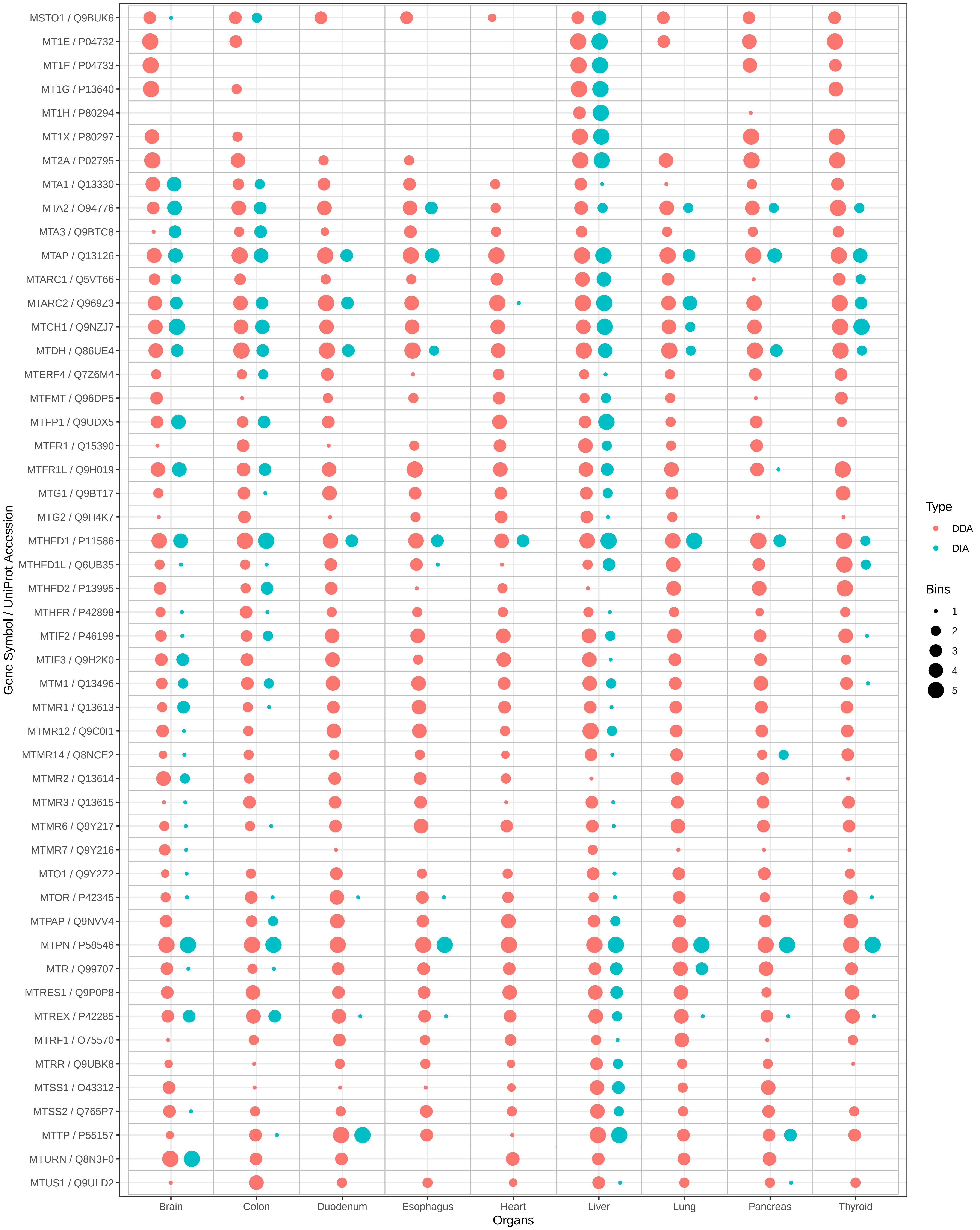
MSTO1 / Q9BUK6
MT1E / P04732
MT1F / P04733
MT1G / P13640
MT1H / P80294
MT1X / P80297
MT2A / P02795
MTA1 / Q13330
MTA2 / O94776
MTA3 / Q9BTC8
MTAP / Q13126
MTARC1 / Q5VT66
MTARC2 / Q969Z3
MTCH1 / Q9NZJ7
MTDH / Q86UE4
MTERF4 / Q7Z6M4
MTFMT / Q96DP5
MTFP1 / Q9UDX5
MTFR1 / Q15390
MTFR1L / Q9H019
MTG1 / Q9BT17
MTG2 / Q9H4K7
MTHFD1 / P11586
MTHFD1L / Q6UB35
MTHFD2 / P13995
MTHFR / P42898
MTIF2 / P46199
MTIF3 / Q9H2K0
MTM1 / Q13496
MTMR1 / Q13613
MTMR12 / Q9C0I1
MTMR14 / Q8NCE2
MTMR2 / Q13614
MTMR3 / Q13615
MTMR6 / Q9Y217
MTMR7 / Q9Y216
MTO1 / Q9Y2Z2
MTOR / P42345
MTPAP / Q9NVV4
MTPN / P58546
MTR / Q99707
MTRES1 / Q9P0P8
MTREX / P42285
MTRF1 / O75570
MTRR / Q9UBK8
MTSS1 / O43312
MTSS2 / Q765P7
MTTP / P55157
MTURN / Q8N3F0
MTUS1 / Q9ULD2
Gene Symbol / UniProt Accession
Brain
Colon
Duodenum
Esophagus
Heart
Liver
Lung
Pancreas
Thyroid
Organs
Type
DDA
DIA
Bins
1
2
3
4
5

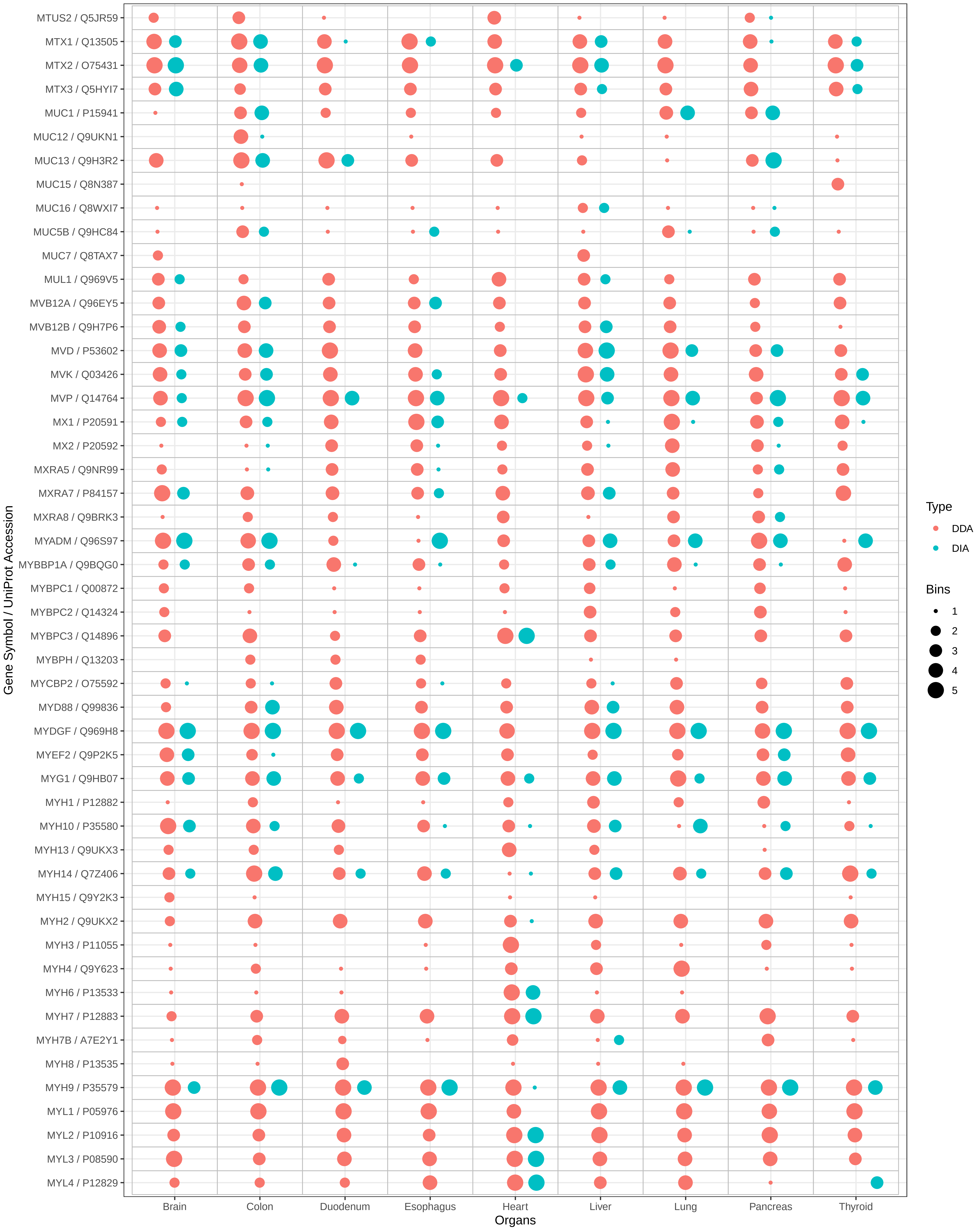

MTUS2 / Q5JR59
MTX1 / Q13505
MTX2 / O75431
MTX3 / Q5HYI7
MUC1 / P15941
MUC12 / Q9UKN1
MUC13 / Q9H3R2
MUC15 / Q8N387
MUC16 / Q8WXI7
MUC5B / Q9HC84
MUC7 / Q8TAX7
MUL1 / Q969V5
MVB12A / Q96EY5
MVB12B / Q9H7P6
MVD / P53602
MVK / Q03426
MVP / Q14764
MX1 / P20591
MX2 / P20592
MXRA5 / Q9NR99
MXRA7 / P84157
MXRA8 / Q9BRK3
MYADM / Q96S97
MYBBP1A / Q9BQG0
MYBPC1 / Q00872
MYBPC2 / Q14324
MYBPC3 / Q14896
MYBPH / Q13203
MYCBP2 / O75592
MYD88 / Q99836
MYDGF / Q969H8
MYEF2 / Q9P2K5
MYG1 / Q9HB07
MYH1 / P12882
MYH10 / P35580
MYH13 / Q9UKX3
MYH14 / Q7Z406
MYH15 / Q9Y2K3
MYH2 / Q9UKX2
MYH3 / P11055
MYH4 / Q9Y623
MYH6 / P13533
MYH7 / P12883
MYH7B / A7E2Y1
MYH8 / P13535
MYH9 / P35579
MYL1 / P05976
MYL2 / P10916
MYL3 / P08590
MYL4 / P12829
Gene Symbol / UniProt Accession
Brain
Colon
Duodenum
Esophagus
Heart
Liver
Lung
Pancreas
Thyroid
Organs
Type
DDA
DIA
Bins
1
2
3
4
5

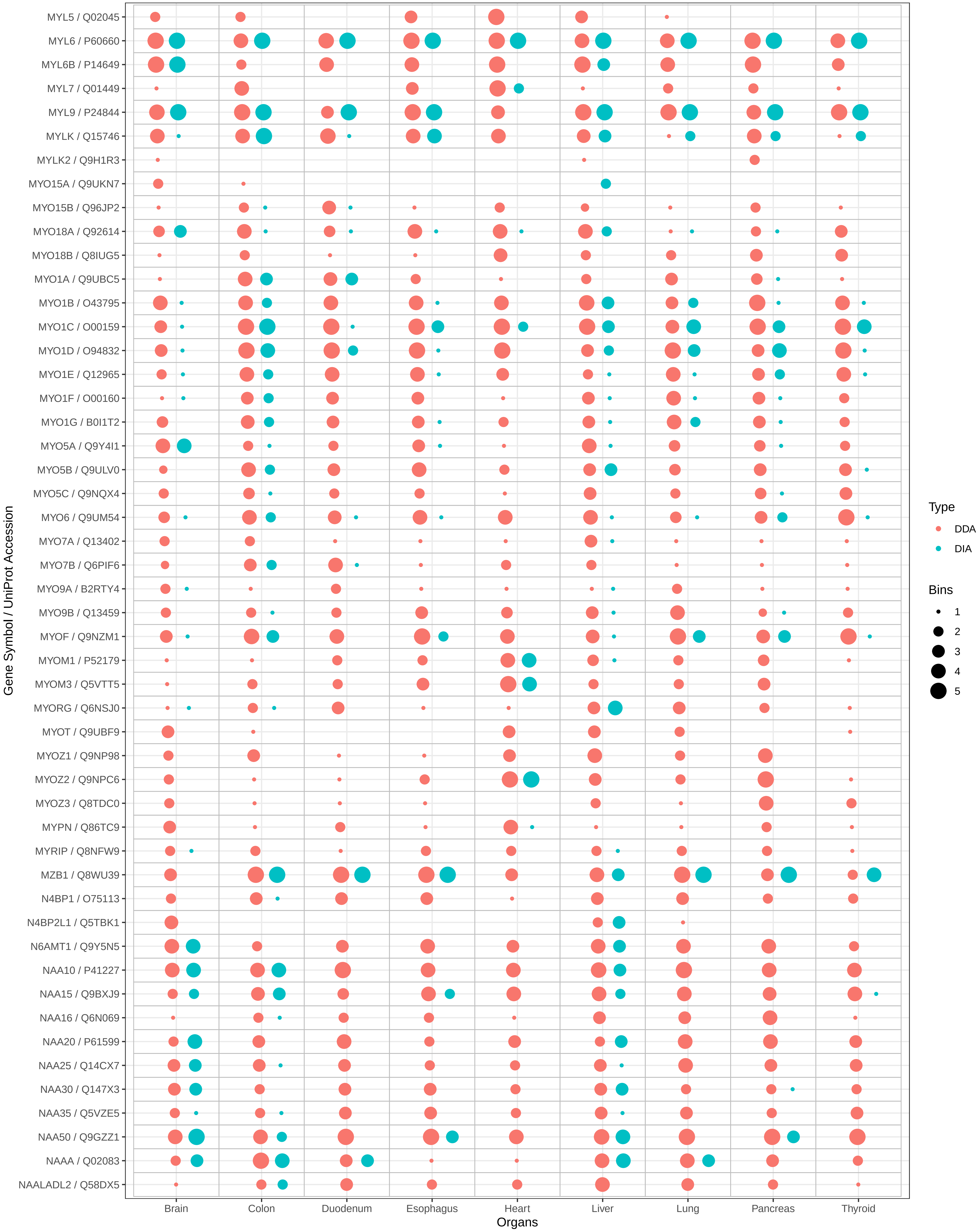

MYL5 / Q02045
MYL6 / P60660
MYL6B / P14649
MYL7 / Q01449
MYL9 / P24844
MYLK / Q15746
MYLK2 / Q9H1R3
MYO15A / Q9UKN7
MYO15B / Q96JP2
MYO18A / Q92614
MYO18B / Q8IUG5
MYO1A / Q9UBC5
MYO1B / O43795
MYO1C / O00159
MYO1D / O94832
MYO1E / Q12965
MYO1F / O00160
MYO1G / B0I1T2
MYO5A / Q9Y4I1
MYO5B / Q9ULV0
MYO5C / Q9NQX4
MYO6 / Q9UM54
MYO7A / Q13402
MYO7B / Q6PIF6
MYO9A / B2RTY4
MYO9B / Q13459
MYOF / Q9NZM1
MYOM1 / P52179
MYOM3 / Q5VTT5
MYORG / Q6NSJ0
MYOT / Q9UBF9
MYOZ1 / Q9NP98
MYOZ2 / Q9NPC6
MYOZ3 / Q8TDC0
MYPN / Q86TC9
MYRIP / Q8NFW9
MZB1 / Q8WU39
N4BP1 / O75113
N4BP2L1 / Q5TBK1
N6AMT1 / Q9Y5N5
NAA10 / P41227
NAA15 / Q9BXJ9
NAA16 / Q6N069
NAA20 / P61599
NAA25 / Q14CX7
NAA30 / Q147X3
NAA35 / Q5VZE5
NAA50 / Q9GZZ1
NAAA / Q02083
NAALADL2 / Q58DX5
Gene Symbol / UniProt Accession
Brain
Colon
Duodenum
Esophagus
Heart
Liver
Lung
Pancreas
Thyroid
Organs
Type
DDA
DIA
Bins
1
2
3
4
5

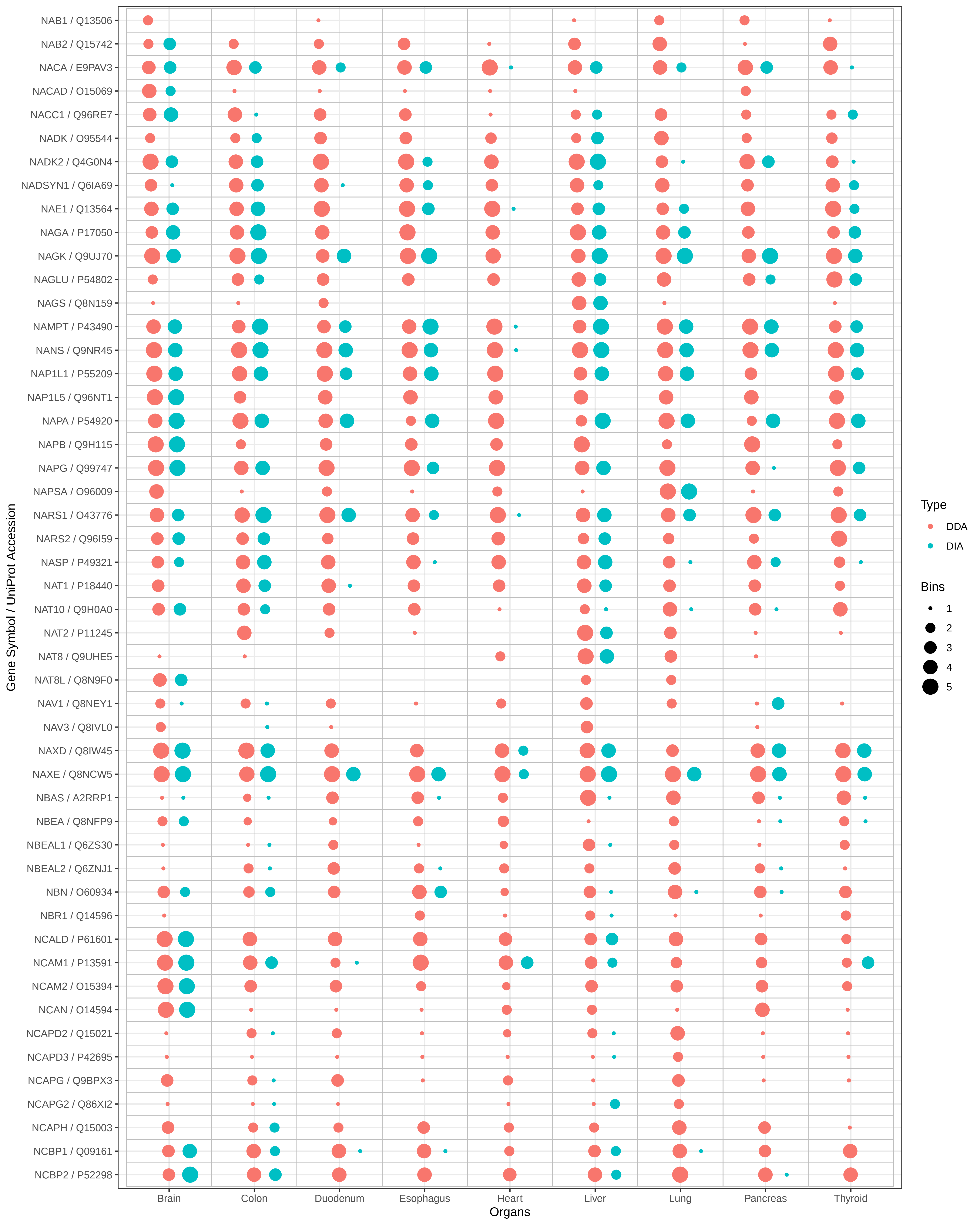
NAB1 / Q13506
NAB2 / Q15742
NACA / E9PAV3
NACAD / O15069
NACC1 / Q96RE7
NADK / O95544
NADK2 / Q4G0N4
NADSYN1 / Q6IA69
NAE1 / Q13564
NAGA / P17050
NAGK / Q9UJ70
NAGLU / P54802
NAGS / Q8N159
NAMPT / P43490
NANS / Q9NR45
NAP1L1 / P55209
NAP1L5 / Q96NT1
NAPA / P54920
NAPB / Q9H115
NAPG / Q99747
NAPSA / O96009
NARS1 / O43776
NARS2 / Q96I59
NASP / P49321
NAT1 / P18440
NAT10 / Q9H0A0
NAT2 / P11245
NAT8 / Q9UHE5
NAT8L / Q8N9F0
NAV1 / Q8NEY1
NAV3 / Q8IVL0
NAXD / Q8IW45
NAXE / Q8NCW5
NBAS / A2RRP1
NBEA / Q8NFP9
NBEAL1 / Q6ZS30
NBEAL2 / Q6ZNJ1
NBN / O60934
NBR1 / Q14596
NCALD / P61601
NCAM1 / P13591
NCAM2 / O15394
NCAN / O14594
NCAPD2 / Q15021
NCAPD3 / P42695
NCAPG / Q9BPX3
NCAPG2 / Q86XI2
NCAPH / Q15003
NCBP1 / Q09161
NCBP2 / P52298
Gene Symbol / UniProt Accession
Brain
Colon
Duodenum
Esophagus
Heart
Liver
Lung
Pancreas
Thyroid
Organs
Type
DDA
DIA
Bins
1
2
3
4
5

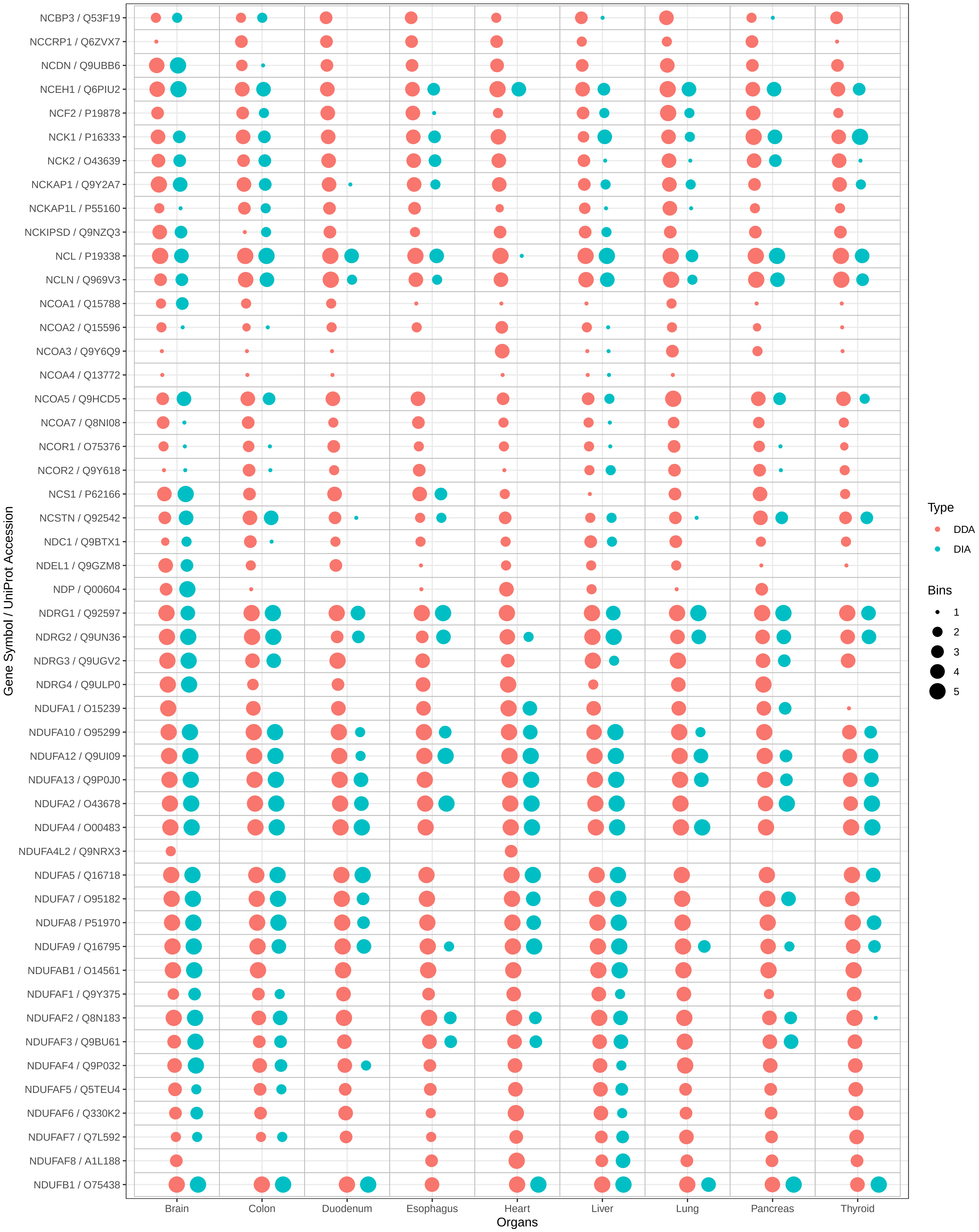
NCBP3 / Q53F19
NCCRP1 / Q6ZVX7
NCDN / Q9UBB6
NCEH1 / Q6PIU2
NCF2 / P19878
NCK1 / P16333
NCK2 / O43639
NCKAP1 / Q9Y2A7
NCKAP1L / P55160
NCKIPSD / Q9NZQ3
NCL / P19338
NCLN / Q969V3
NCOA1 / Q15788
NCOA2 / Q15596
NCOA3 / Q9Y6Q9
NCOA4 / Q13772
NCOA5 / Q9HCD5
NCOA7 / Q8NI08
NCOR1 / O75376
NCOR2 / Q9Y618
NCS1 / P62166
NCSTN / Q92542
NDC1 / Q9BTX1
NDEL1 / Q9GZM8
NDP / Q00604
NDRG1 / Q92597
NDRG2 / Q9UN36
NDRG3 / Q9UGV2
NDRG4 / Q9ULP0
NDUFA1 / O15239
NDUFA10 / O95299
NDUFA12 / Q9UI09
NDUFA13 / Q9P0J0
NDUFA2 / O43678
NDUFA4 / O00483
NDUFA4L2 / Q9NRX3
NDUFA5 / Q16718
NDUFA7 / O95182
NDUFA8 / P51970
NDUFA9 / Q16795
NDUFAB1 / O14561
NDUFAF1 / Q9Y375
NDUFAF2 / Q8N183
NDUFAF3 / Q9BU61
NDUFAF4 / Q9P032
NDUFAF5 / Q5TEU4
NDUFAF6 / Q330K2
NDUFAF7 / Q7L592
NDUFAF8 / A1L188
NDUFB1 / O75438
Gene Symbol / UniProt Accession
Brain
Colon
Duodenum
Esophagus
Heart
Liver
Lung
Pancreas
Thyroid
Organs
Type
DDA
DIA
Bins
1
2
3
4
5

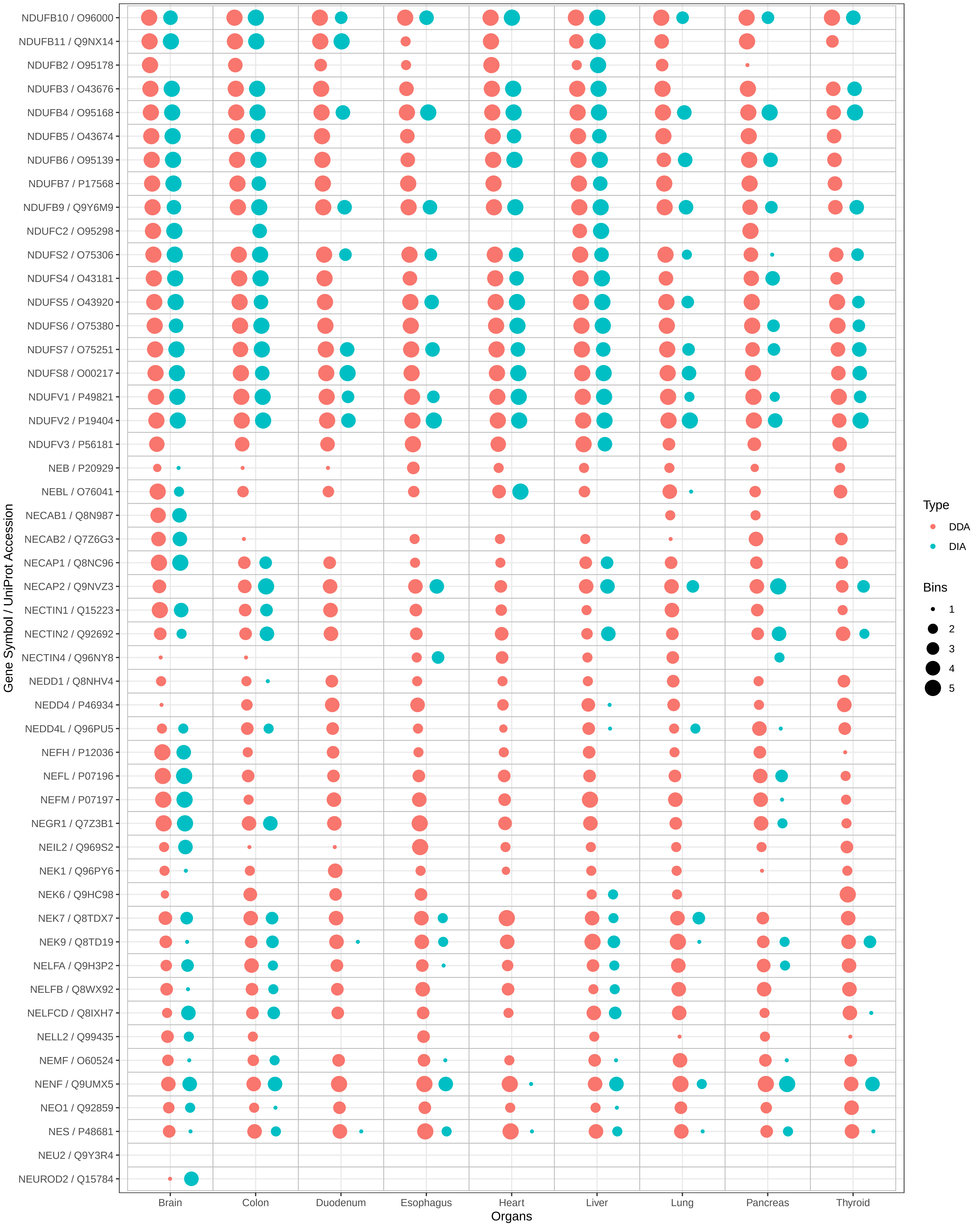

NDUFB10 / O96000
NDUFB11 / Q9NX14
NDUFB2 / O95178
NDUFB3 / O43676
NDUFB4 / O95168
NDUFB5 / O43674
NDUFB6 / O95139
NDUFB7 / P17568
NDUFB9 / Q9Y6M9
NDUFC2 / O95298
NDUFS2 / O75306
NDUFS4 / O43181
NDUFS5 / O43920
NDUFS6 / O75380
NDUFS7 / O75251
NDUFS8 / O00217
NDUFV1 / P49821
NDUFV2 / P19404
NDUFV3 / P56181
NEB / P20929
NEBL / O76041
NECAB1 / Q8N987
NECAB2 / Q7Z6G3
NECAP1 / Q8NC96
NECAP2 / Q9NVZ3
NECTIN1 / Q15223
NECTIN2 / Q92692
NECTIN4 / Q96NY8
NEDD1 / Q8NHV4
NEDD4 / P46934
NEDD4L / Q96PU5
NEFH / P12036
NEFL / P07196
NEFM / P07197
NEGR1 / Q7Z3B1
NEIL2 / Q969S2
NEK1 / Q96PY6
NEK6 / Q9HC98
NEK7 / Q8TDX7
NEK9 / Q8TD19
NELFA / Q9H3P2
NELFB / Q8WX92
NELFCD / Q8IXH7
NELL2 / Q99435
NEMF / O60524
NENF / Q9UMX5
NEO1 / Q92859
NES / P48681
NEU2 / Q9Y3R4
NEUROD2 / Q15784
Gene Symbol / UniProt Accession
Brain
Colon
Duodenum
Esophagus
Heart
Liver
Lung
Pancreas
Thyroid
Organs
Type
DDA
DIA
Bins
1
2
3
4
5

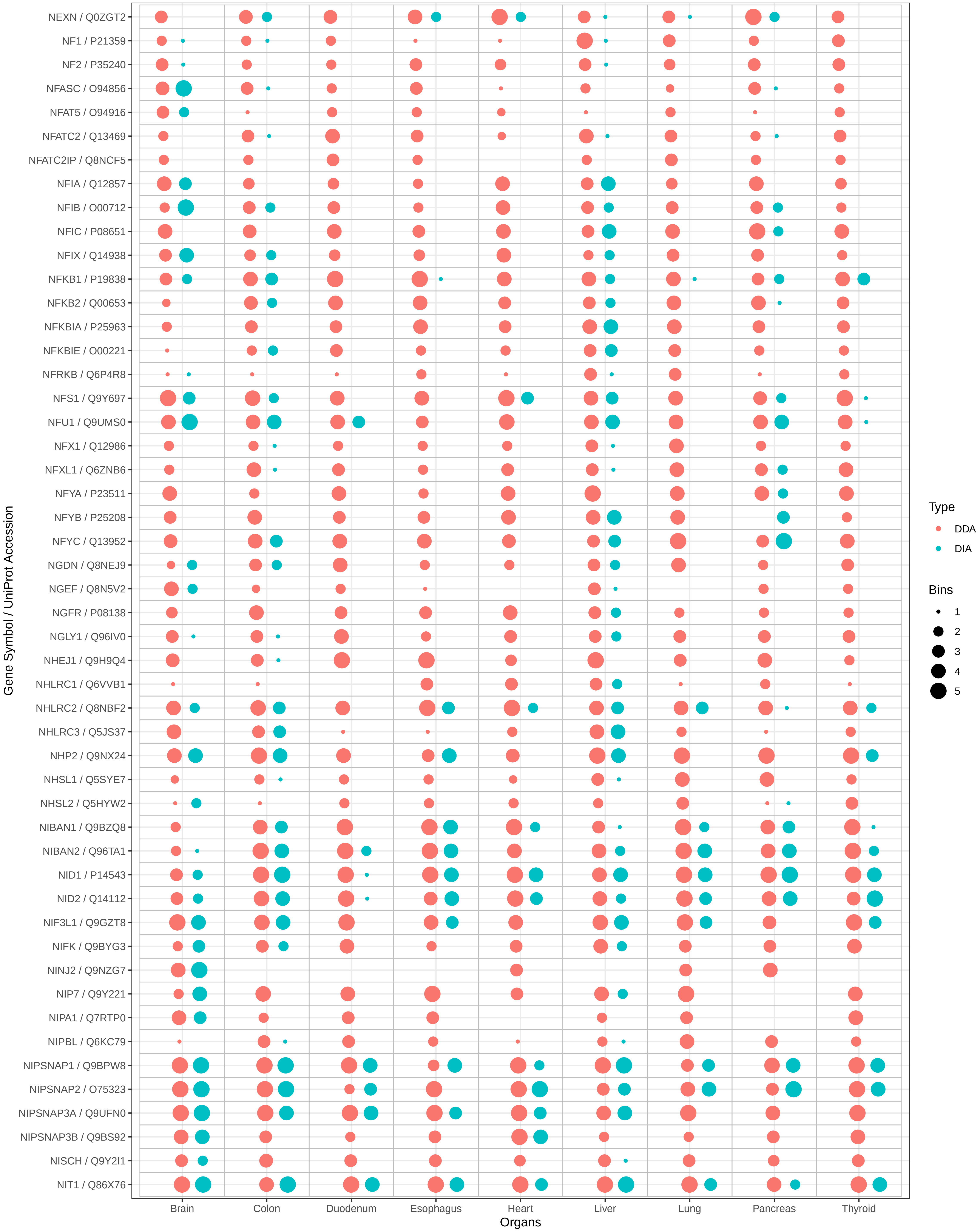
NEXN / Q0ZGT2
NF1 / P21359
NF2 / P35240
NFASC / O94856
NFAT5 / O94916
NFATC2 / Q13469
NFATC2IP / Q8NCF5
NFIA / Q12857
NFIB / O00712
NFIC / P08651
NFIX / Q14938
NFKB1 / P19838
NFKB2 / Q00653
NFKBIA / P25963
NFKBIE / O00221
NFRKB / Q6P4R8
NFS1 / Q9Y697
NFU1 / Q9UMS0
NFX1 / Q12986
NFXL1 / Q6ZNB6
NFYA / P23511
NFYB / P25208
NFYC / Q13952
NGDN / Q8NEJ9
NGEF / Q8N5V2
NGFR / P08138
NGLY1 / Q96IV0
NHEJ1 / Q9H9Q4
NHLRC1 / Q6VVB1
NHLRC2 / Q8NBF2
NHLRC3 / Q5JS37
NHP2 / Q9NX24
NHSL1 / Q5SYE7
NHSL2 / Q5HYW2
NIBAN1 / Q9BZQ8
NIBAN2 / Q96TA1
NID1 / P14543
NID2 / Q14112
NIF3L1 / Q9GZT8
NIFK / Q9BYG3
NINJ2 / Q9NZG7
NIP7 / Q9Y221
NIPA1 / Q7RTP0
NIPBL / Q6KC79
NIPSNAP1 / Q9BPW8
NIPSNAP2 / O75323
NIPSNAP3A / Q9UFN0
NIPSNAP3B / Q9BS92
NISCH / Q9Y2I1
NIT1 / Q86X76
Gene Symbol / UniProt Accession
Brain
Colon
Duodenum
Esophagus
Heart
Liver
Lung
Pancreas
Thyroid
Organs
Type
DDA
DIA
Bins
1
2
3
4
5

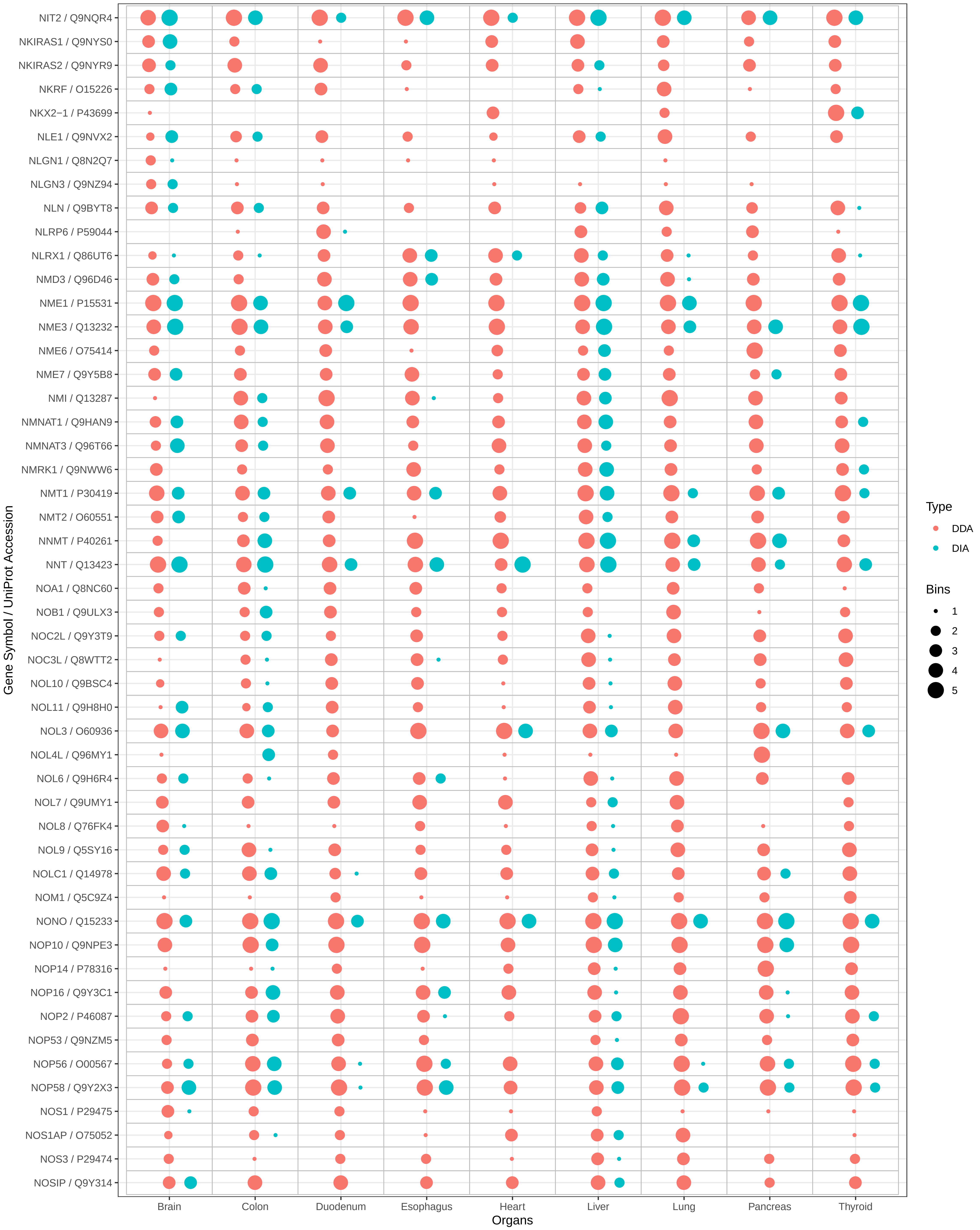
NIT2 / Q9NQR4
NKIRAS1 / Q9NYS0
NKIRAS2 / Q9NYR9
NKRF / O15226
NKX2−1 / P43699
NLE1 / Q9NVX2
NLGN1 / Q8N2Q7
NLGN3 / Q9NZ94
NLN / Q9BYT8
NLRP6 / P59044
NLRX1 / Q86UT6
NMD3 / Q96D46
NME1 / P15531
NME3 / Q13232
NME6 / O75414
NME7 / Q9Y5B8
NMI / Q13287
NMNAT1 / Q9HAN9
NMNAT3 / Q96T66
NMRK1 / Q9NWW6
NMT1 / P30419
NMT2 / O60551
NNMT / P40261
NNT / Q13423
NOA1 / Q8NC60
NOB1 / Q9ULX3
NOC2L / Q9Y3T9
NOC3L / Q8WTT2
NOL10 / Q9BSC4
NOL11 / Q9H8H0
NOL3 / O60936
NOL4L / Q96MY1
NOL6 / Q9H6R4
NOL7 / Q9UMY1
NOL8 / Q76FK4
NOL9 / Q5SY16
NOLC1 / Q14978
NOM1 / Q5C9Z4
NONO / Q15233
NOP10 / Q9NPE3
NOP14 / P78316
NOP16 / Q9Y3C1
NOP2 / P46087
NOP53 / Q9NZM5
NOP56 / O00567
NOP58 / Q9Y2X3
NOS1 / P29475
NOS1AP / O75052
NOS3 / P29474
NOSIP / Q9Y314
Gene Symbol / UniProt Accession
Brain
Colon
Duodenum
Esophagus
Heart
Liver
Lung
Pancreas
Thyroid
Organs
Type
DDA
DIA
Bins
1
2
3
4
5

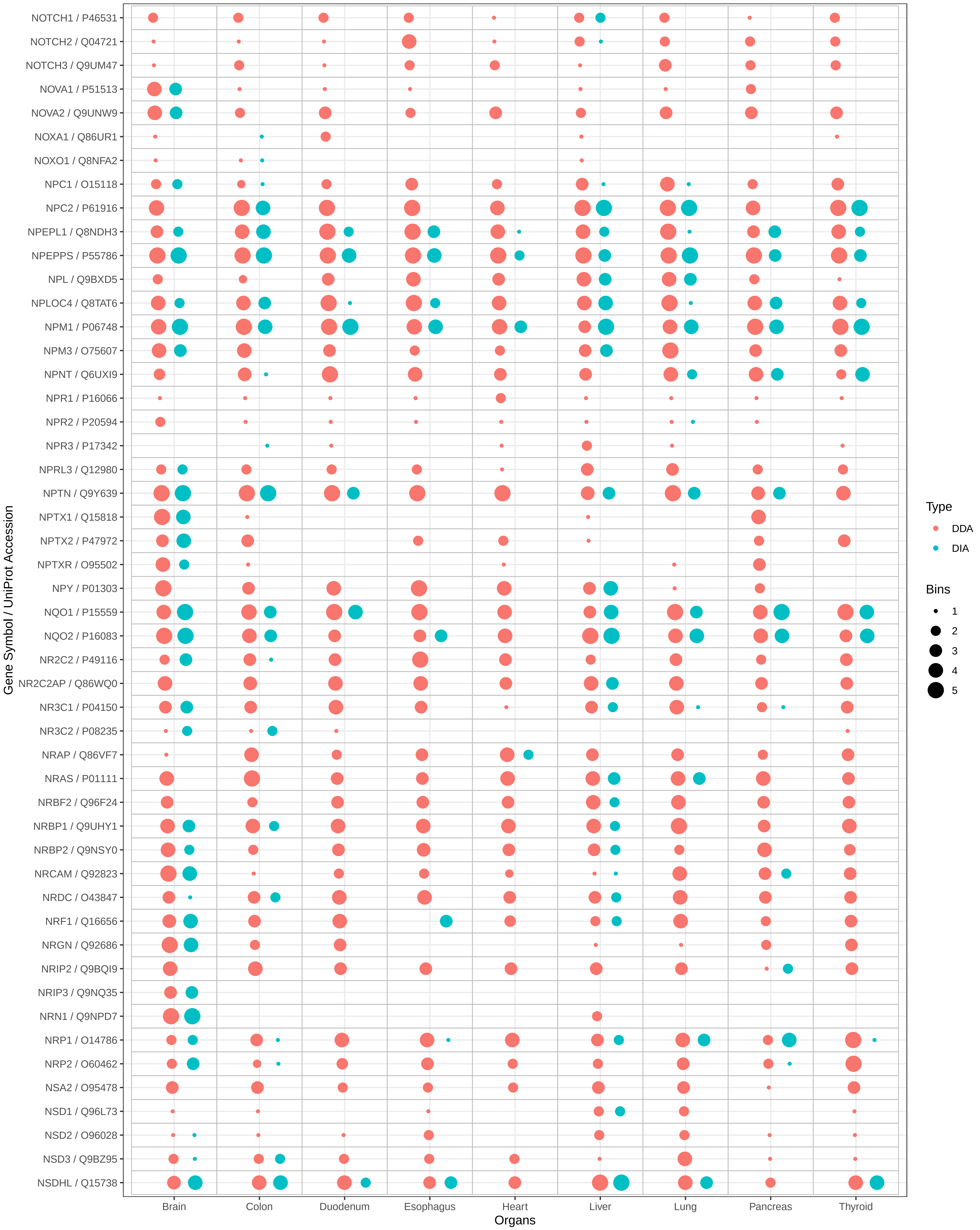

NOTCH1 / P46531
NOTCH2 / Q04721
NOTCH3 / Q9UM47
NOVA1 / P51513
NOVA2 / Q9UNW9
NOXA1 / Q86UR1
NOXO1 / Q8NFA2
NPC1 / O15118
NPC2 / P61916
NPEPL1 / Q8NDH3
NPEPPS / P55786
NPL / Q9BXD5
NPLOC4 / Q8TAT6
NPM1 / P06748
NPM3 / O75607
NPNT / Q6UXI9
NPR1 / P16066
NPR2 / P20594
NPR3 / P17342
NPRL3 / Q12980
NPTN / Q9Y639
NPTX1 / Q15818
NPTX2 / P47972
NPTXR / O95502
NPY / P01303
NQO1 / P15559
NQO2 / P16083
NR2C2 / P49116
NR2C2AP / Q86WQ0
NR3C1 / P04150
NR3C2 / P08235
NRAP / Q86VF7
NRAS / P01111
NRBF2 / Q96F24
NRBP1 / Q9UHY1
NRBP2 / Q9NSY0
NRCAM / Q92823
NRDC / O43847
NRF1 / Q16656
NRGN / Q92686
NRIP2 / Q9BQI9
NRIP3 / Q9NQ35
NRN1 / Q9NPD7
NRP1 / O14786
NRP2 / O60462
NSA2 / O95478
NSD1 / Q96L73
NSD2 / O96028
NSD3 / Q9BZ95
NSDHL / Q15738
Gene Symbol / UniProt Accession
Brain
Colon
Duodenum
Esophagus
Heart
Liver
Lung
Pancreas
Thyroid
Organs
Type
DDA
DIA
Bins
1
2
3
4
5

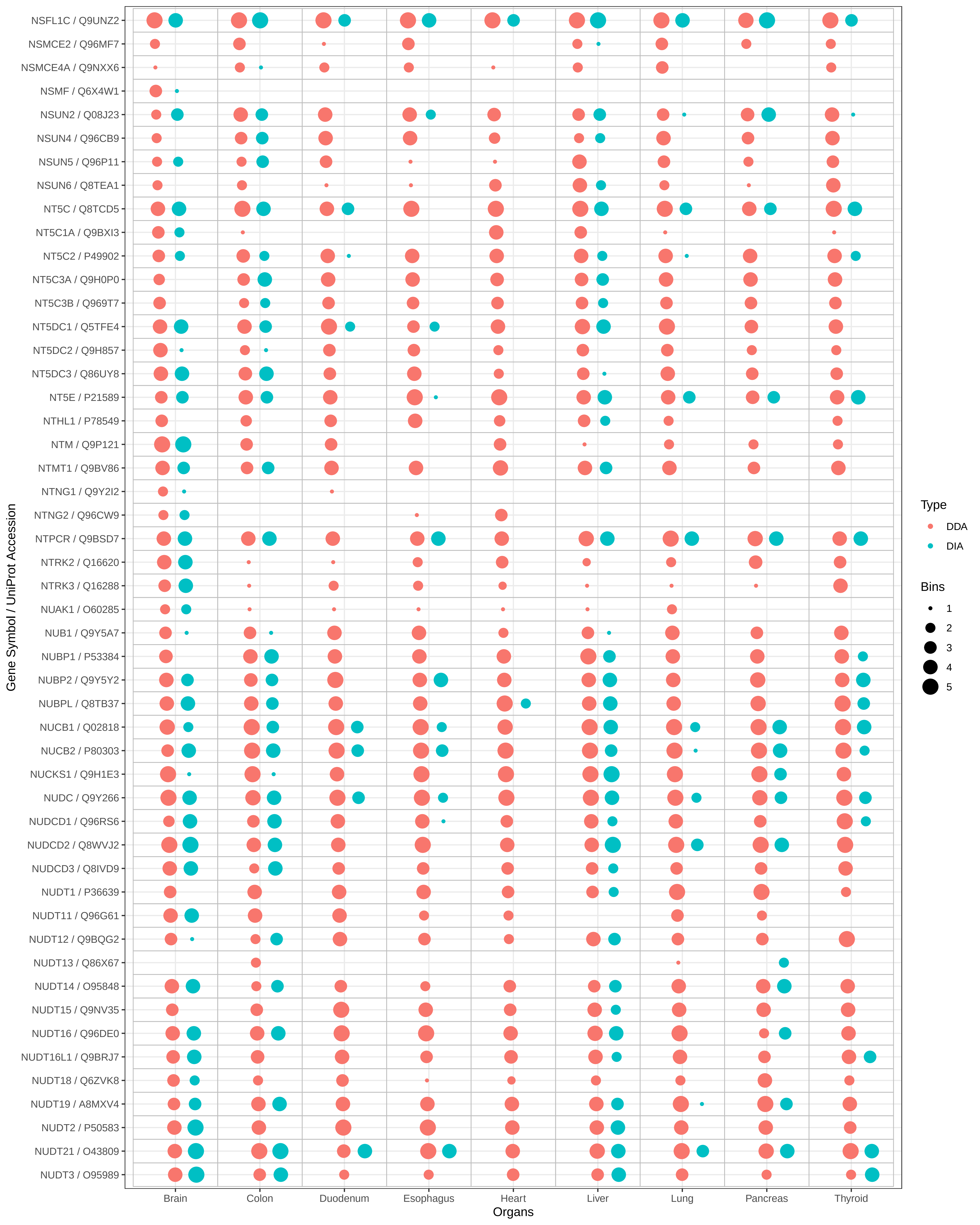

NSFL1C / Q9UNZ2
NSMCE2 / Q96MF7
NSMCE4A / Q9NXX6
NSMF / Q6X4W1
NSUN2 / Q08J23
NSUN4 / Q96CB9
NSUN5 / Q96P11
NSUN6 / Q8TEA1
NT5C / Q8TCD5
NT5C1A / Q9BXI3
NT5C2 / P49902
NT5C3A / Q9H0P0
NT5C3B / Q969T7
NT5DC1 / Q5TFE4
NT5DC2 / Q9H857
NT5DC3 / Q86UY8
NT5E / P21589
NTHL1 / P78549
NTM / Q9P121
NTMT1 / Q9BV86
NTNG1 / Q9Y2I2
NTNG2 / Q96CW9
NTPCR / Q9BSD7
NTRK2 / Q16620
NTRK3 / Q16288
NUAK1 / O60285
NUB1 / Q9Y5A7
NUBP1 / P53384
NUBP2 / Q9Y5Y2
NUBPL / Q8TB37
NUCB1 / Q02818
NUCB2 / P80303
NUCKS1 / Q9H1E3
NUDC / Q9Y266
NUDCD1 / Q96RS6
NUDCD2 / Q8WVJ2
NUDCD3 / Q8IVD9
NUDT1 / P36639
NUDT11 / Q96G61
NUDT12 / Q9BQG2
NUDT13 / Q86X67
NUDT14 / O95848
NUDT15 / Q9NV35
NUDT16 / Q96DE0
NUDT16L1 / Q9BRJ7
NUDT18 / Q6ZVK8
NUDT19 / A8MXV4
NUDT2 / P50583
NUDT21 / O43809
NUDT3 / O95989
Gene Symbol / UniProt Accession
Brain
Colon
Duodenum
Esophagus
Heart
Liver
Lung
Pancreas
Thyroid
Organs
Type
DDA
DIA
Bins
1
2
3
4
5

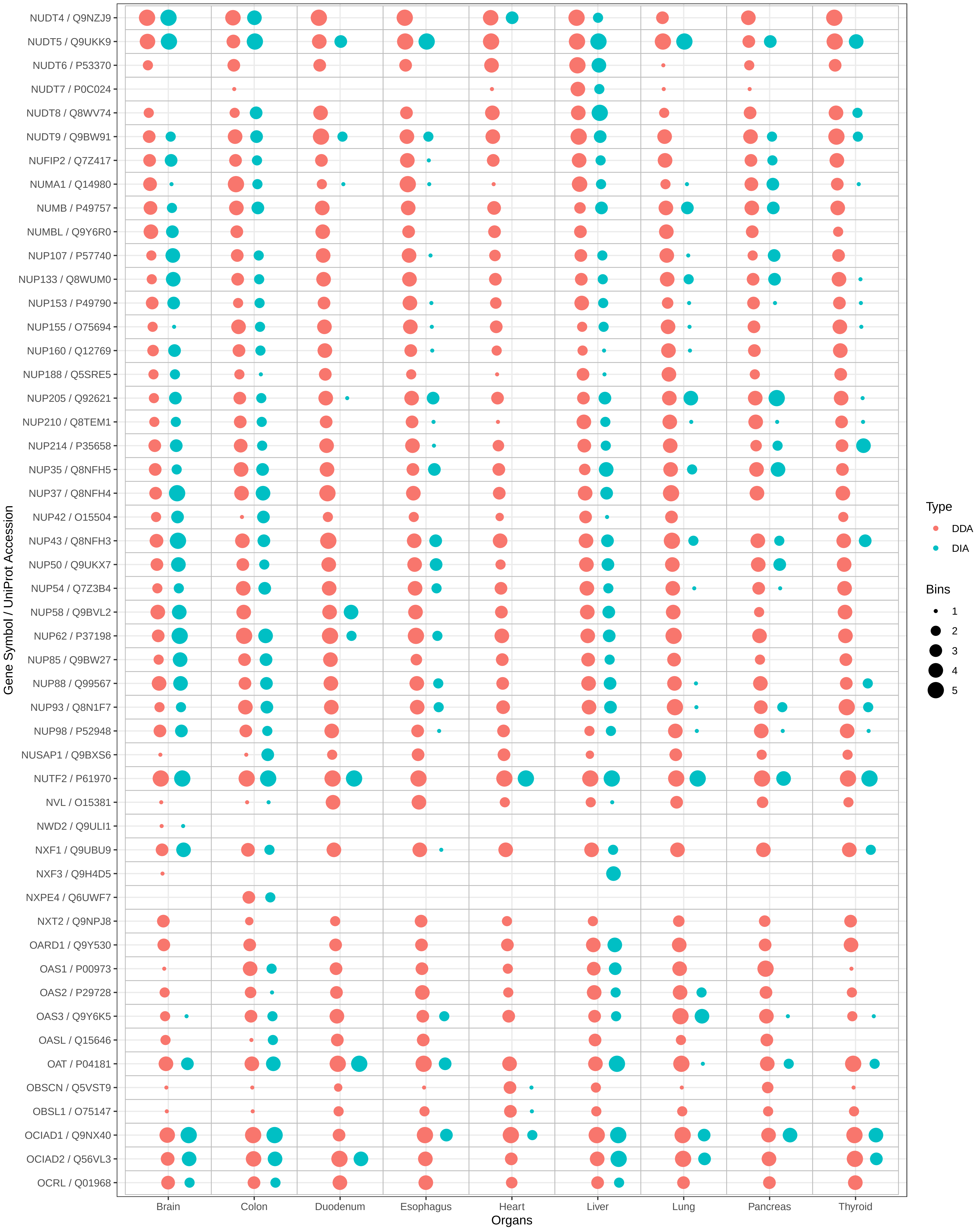

NUDT4 / Q9NZJ9
NUDT5 / Q9UKK9
NUDT6 / P53370
NUDT7 / P0C024
NUDT8 / Q8WV74
NUDT9 / Q9BW91
NUFIP2 / Q7Z417
NUMA1 / Q14980
NUMB / P49757
NUMBL / Q9Y6R0
NUP107 / P57740
NUP133 / Q8WUM0
NUP153 / P49790
NUP155 / O75694
NUP160 / Q12769
NUP188 / Q5SRE5
NUP205 / Q92621
NUP210 / Q8TEM1
NUP214 / P35658
NUP35 / Q8NFH5
NUP37 / Q8NFH4
NUP42 / O15504
NUP43 / Q8NFH3
NUP50 / Q9UKX7
NUP54 / Q7Z3B4
NUP58 / Q9BVL2
NUP62 / P37198
NUP85 / Q9BW27
NUP88 / Q99567
NUP93 / Q8N1F7
NUP98 / P52948
NUSAP1 / Q9BXS6
NUTF2 / P61970
NVL / O15381
NWD2 / Q9ULI1
NXF1 / Q9UBU9
NXF3 / Q9H4D5
NXPE4 / Q6UWF7
NXT2 / Q9NPJ8
OARD1 / Q9Y530
OAS1 / P00973
OAS2 / P29728
OAS3 / Q9Y6K5
OASL / Q15646
OAT / P04181
OBSCN / Q5VST9
OBSL1 / O75147
OCIAD1 / Q9NX40
OCIAD2 / Q56VL3
OCRL / Q01968
Gene Symbol / UniProt Accession
Brain
Colon
Duodenum
Esophagus
Heart
Liver
Lung
Pancreas
Thyroid
Organs
Type
DDA
DIA
Bins
1
2
3
4
5

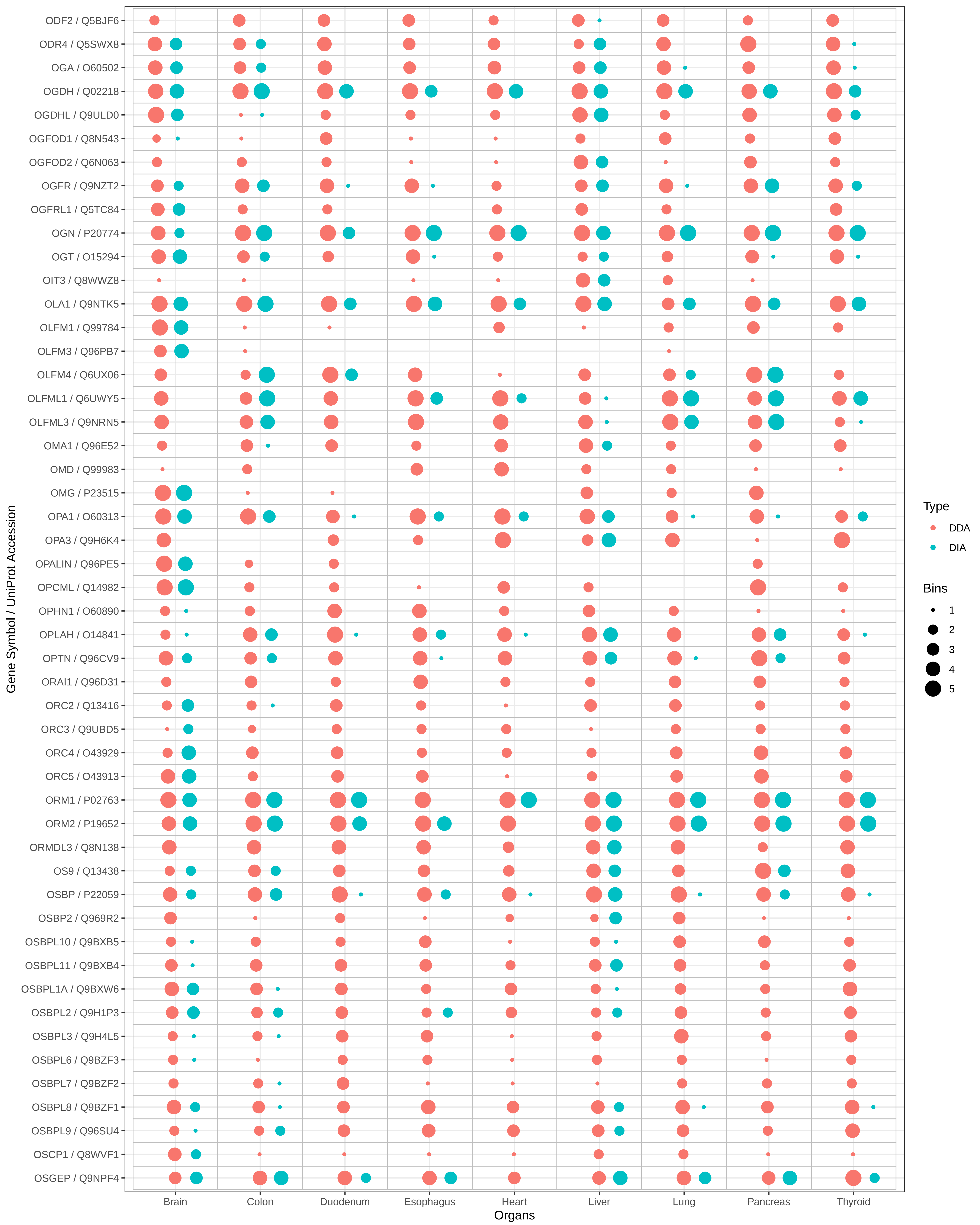

ODF2 / Q5BJF6
ODR4 / Q5SWX8
OGA / O60502
OGDH / Q02218
OGDHL / Q9ULD0
OGFOD1 / Q8N543
OGFOD2 / Q6N063
OGFR / Q9NZT2
OGFRL1 / Q5TC84
OGN / P20774
OGT / O15294
OIT3 / Q8WWZ8
OLA1 / Q9NTK5
OLFM1 / Q99784
OLFM3 / Q96PB7
OLFM4 / Q6UX06
OLFML1 / Q6UWY5
OLFML3 / Q9NRN5
OMA1 / Q96E52
OMD / Q99983
OMG / P23515
OPA1 / O60313
OPA3 / Q9H6K4
OPALIN / Q96PE5
OPCML / Q14982
OPHN1 / O60890
OPLAH / O14841
OPTN / Q96CV9
ORAI1 / Q96D31
ORC2 / Q13416
ORC3 / Q9UBD5
ORC4 / O43929
ORC5 / O43913
ORM1 / P02763
ORM2 / P19652
ORMDL3 / Q8N138
OS9 / Q13438
OSBP / P22059
OSBP2 / Q969R2
OSBPL10 / Q9BXB5
OSBPL11 / Q9BXB4
OSBPL1A / Q9BXW6
OSBPL2 / Q9H1P3
OSBPL3 / Q9H4L5
OSBPL6 / Q9BZF3
OSBPL7 / Q9BZF2
OSBPL8 / Q9BZF1
OSBPL9 / Q96SU4
OSCP1 / Q8WVF1
OSGEP / Q9NPF4
Gene Symbol / UniProt Accession
Brain
Colon
Duodenum
Esophagus
Heart
Liver
Lung
Pancreas
Thyroid
Organs
Type
DDA
DIA
Bins
1
2
3
4
5

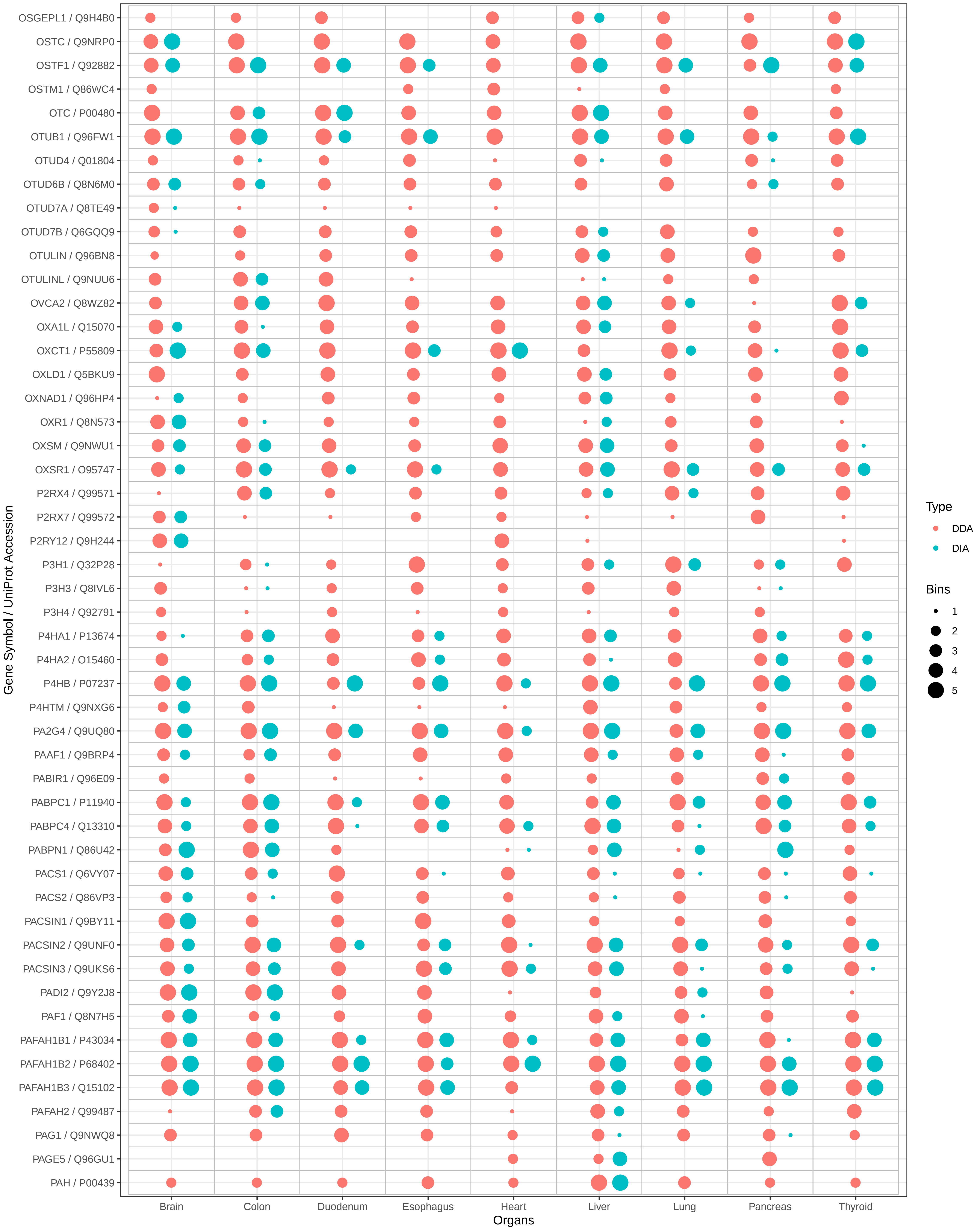

OSGEPL1 / Q9H4B0
OSTC / Q9NRP0
OSTF1 / Q92882
OSTM1 / Q86WC4
OTC / P00480
OTUB1 / Q96FW1
OTUD4 / Q01804
OTUD6B / Q8N6M0
OTUD7A / Q8TE49
OTUD7B / Q6GQQ9
OTULIN / Q96BN8
OTULINL / Q9NUU6
OVCA2 / Q8WZ82
OXA1L / Q15070
OXCT1 / P55809
OXLD1 / Q5BKU9
OXNAD1 / Q96HP4
OXR1 / Q8N573
OXSM / Q9NWU1
OXSR1 / O95747
P2RX4 / Q99571
P2RX7 / Q99572
P2RY12 / Q9H244
P3H1 / Q32P28
P3H3 / Q8IVL6
P3H4 / Q92791
P4HA1 / P13674
P4HA2 / O15460
P4HB / P07237
P4HTM / Q9NXG6
PA2G4 / Q9UQ80
PAAF1 / Q9BRP4
PABIR1 / Q96E09
PABPC1 / P11940
PABPC4 / Q13310
PABPN1 / Q86U42
PACS1 / Q6VY07
PACS2 / Q86VP3
PACSIN1 / Q9BY11
PACSIN2 / Q9UNF0
PACSIN3 / Q9UKS6
PADI2 / Q9Y2J8
PAF1 / Q8N7H5
PAFAH1B1 / P43034
PAFAH1B2 / P68402
PAFAH1B3 / Q15102
PAFAH2 / Q99487
PAG1 / Q9NWQ8
PAGE5 / Q96GU1
PAH / P00439
Gene Symbol / UniProt Accession
Brain
Colon
Duodenum
Esophagus
Heart
Liver
Lung
Pancreas
Thyroid
Organs
Type
DDA
DIA
Bins
1
2
3
4
5

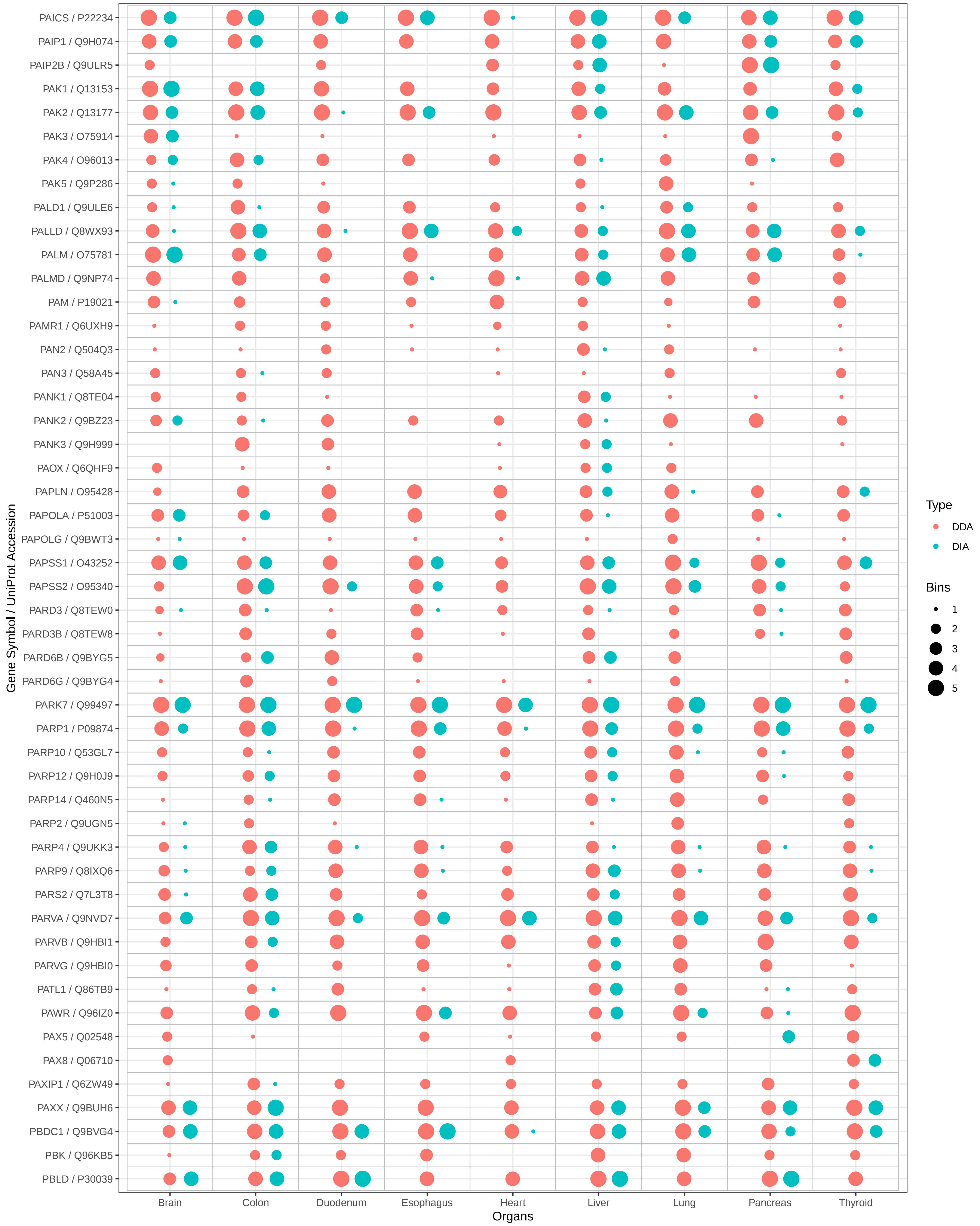

PAICS / P22234
PAIP1 / Q9H074
PAIP2B / Q9ULR5
PAK1 / Q13153
PAK2 / Q13177
PAK3 / O75914
PAK4 / O96013
PAK5 / Q9P286
PALD1 / Q9ULE6
PALLD / Q8WX93
PALM / O75781
PALMD / Q9NP74
PAM / P19021
PAMR1 / Q6UXH9
PAN2 / Q504Q3
PAN3 / Q58A45
PANK1 / Q8TE04
PANK2 / Q9BZ23
PANK3 / Q9H999
PAOX / Q6QHF9
PAPLN / O95428
PAPOLA / P51003
PAPOLG / Q9BWT3
PAPSS1 / O43252
PAPSS2 / O95340
PARD3 / Q8TEW0
PARD3B / Q8TEW8
PARD6B / Q9BYG5
PARD6G / Q9BYG4
PARK7 / Q99497
PARP1 / P09874
PARP10 / Q53GL7
PARP12 / Q9H0J9
PARP14 / Q460N5
PARP2 / Q9UGN5
PARP4 / Q9UKK3
PARP9 / Q8IXQ6
PARS2 / Q7L3T8
PARVA / Q9NVD7
PARVB / Q9HBI1
PARVG / Q9HBI0
PATL1 / Q86TB9
PAWR / Q96IZ0
PAX5 / Q02548
PAX8 / Q06710
PAXIP1 / Q6ZW49
PAXX / Q9BUH6
PBDC1 / Q9BVG4
PBK / Q96KB5
PBLD / P30039
Gene Symbol / UniProt Accession
Brain
Colon
Duodenum
Esophagus
Heart
Liver
Lung
Pancreas
Thyroid
Organs
Type
DDA
DIA
Bins
1
2
3
4
5

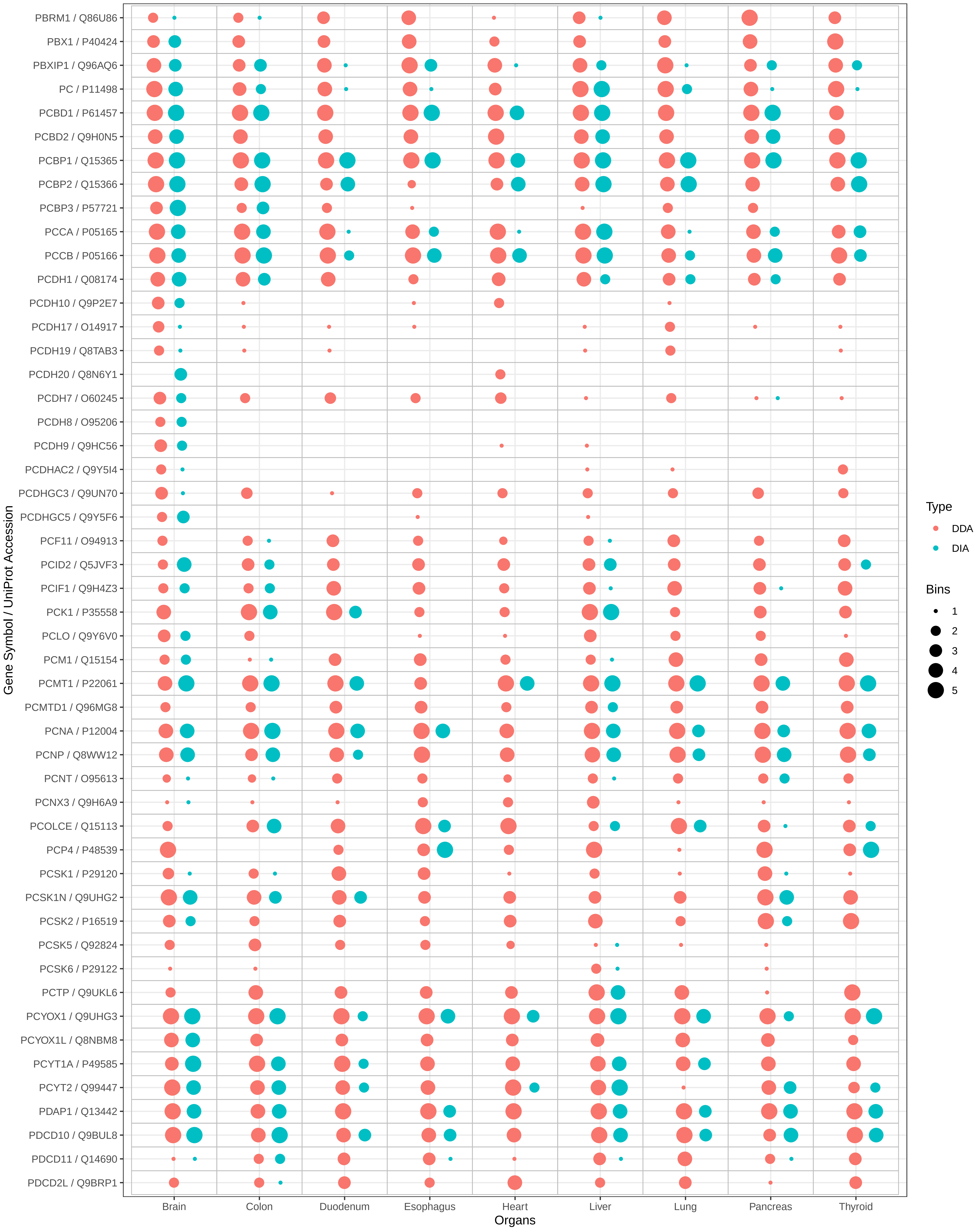
PBRM1 / Q86U86
PBX1 / P40424
PBXIP1 / Q96AQ6
PC / P11498
PCBD1 / P61457
PCBD2 / Q9H0N5
PCBP1 / Q15365
PCBP2 / Q15366
PCBP3 / P57721
PCCA / P05165
PCCB / P05166
PCDH1 / Q08174
PCDH10 / Q9P2E7
PCDH17 / O14917
PCDH19 / Q8TAB3
PCDH20 / Q8N6Y1
PCDH7 / O60245
PCDH8 / O95206
PCDH9 / Q9HC56
PCDHAC2 / Q9Y5I4
PCDHGC3 / Q9UN70
PCDHGC5 / Q9Y5F6
PCF11 / O94913
PCID2 / Q5JVF3
PCIF1 / Q9H4Z3
PCK1 / P35558
PCLO / Q9Y6V0
PCM1 / Q15154
PCMT1 / P22061
PCMTD1 / Q96MG8
PCNA / P12004
PCNP / Q8WW12
PCNT / O95613
PCNX3 / Q9H6A9
PCOLCE / Q15113
PCP4 / P48539
PCSK1 / P29120
PCSK1N / Q9UHG2
PCSK2 / P16519
PCSK5 / Q92824
PCSK6 / P29122
PCTP / Q9UKL6
PCYOX1 / Q9UHG3
PCYOX1L / Q8NBM8
PCYT1A / P49585
PCYT2 / Q99447
PDAP1 / Q13442
PDCD10 / Q9BUL8
PDCD11 / Q14690
PDCD2L / Q9BRP1
Gene Symbol / UniProt Accession
Brain
Colon
Duodenum
Esophagus
Heart
Liver
Lung
Pancreas
Thyroid
Organs
Type
DDA
DIA
Bins
1
2
3
4
5

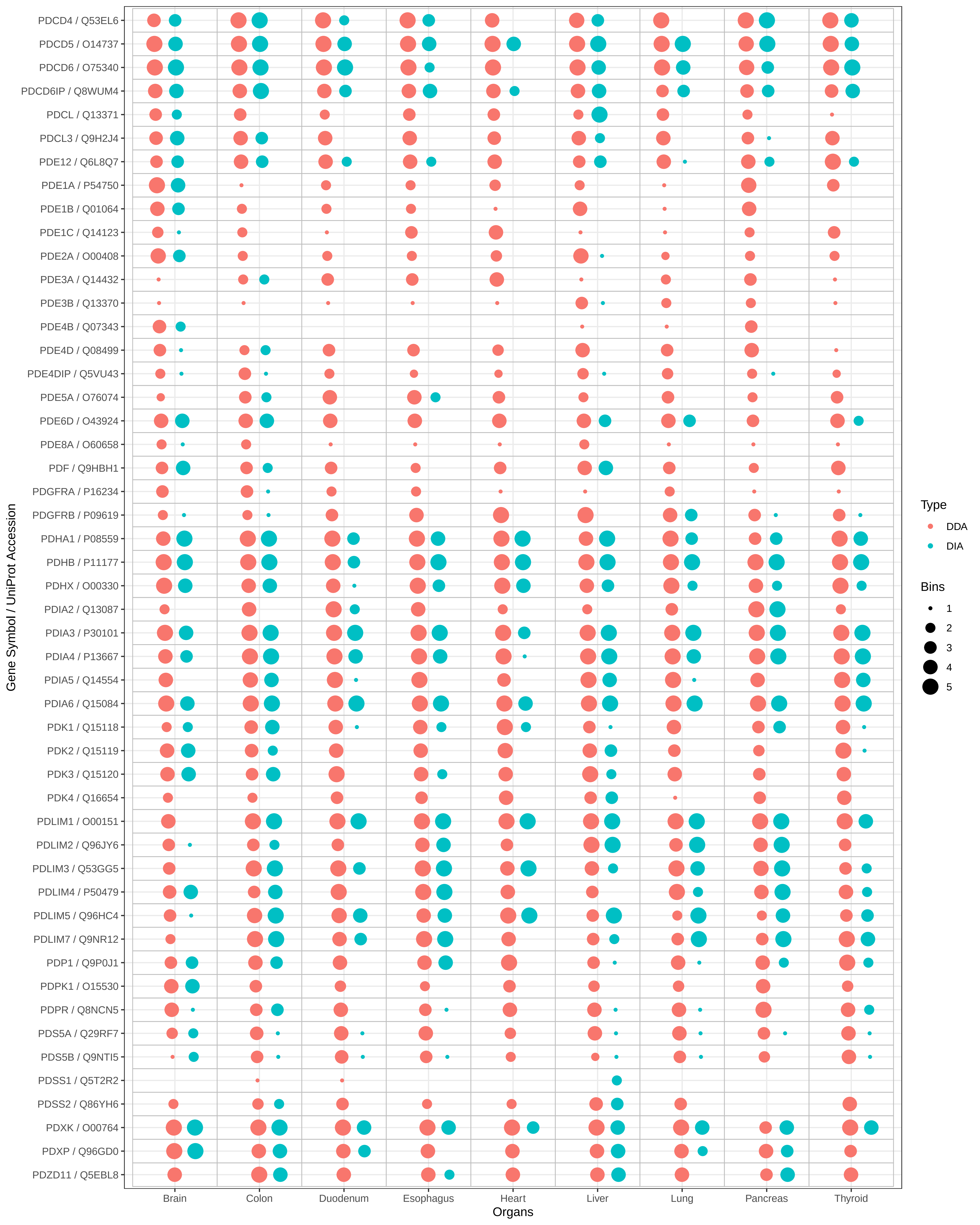
PDCD4 / Q53EL6
PDCD5 / O14737
PDCD6 / O75340
PDCD6IP / Q8WUM4
PDCL / Q13371
PDCL3 / Q9H2J4
PDE12 / Q6L8Q7
PDE1A / P54750
PDE1B / Q01064
PDE1C / Q14123
PDE2A / O00408
PDE3A / Q14432
PDE3B / Q13370
PDE4B / Q07343
PDE4D / Q08499
PDE4DIP / Q5VU43
PDE5A / O76074
PDE6D / O43924
PDE8A / O60658
PDF / Q9HBH1
PDGFRA / P16234
PDGFRB / P09619
PDHA1 / P08559
PDHB / P11177
PDHX / O00330
PDIA2 / Q13087
PDIA3 / P30101
PDIA4 / P13667
PDIA5 / Q14554
PDIA6 / Q15084
PDK1 / Q15118
PDK2 / Q15119
PDK3 / Q15120
PDK4 / Q16654
PDLIM1 / O00151
PDLIM2 / Q96JY6
PDLIM3 / Q53GG5
PDLIM4 / P50479
PDLIM5 / Q96HC4
PDLIM7 / Q9NR12
PDP1 / Q9P0J1
PDPK1 / O15530
PDPR / Q8NCN5
PDS5A / Q29RF7
PDS5B / Q9NTI5
PDSS1 / Q5T2R2
PDSS2 / Q86YH6
PDXK / O00764
PDXP / Q96GD0
PDZD11 / Q5EBL8
Gene Symbol / UniProt Accession
Brain
Colon
Duodenum
Esophagus
Heart
Liver
Lung
Pancreas
Thyroid
Organs
Type
DDA
DIA
Bins
1
2
3
4
5

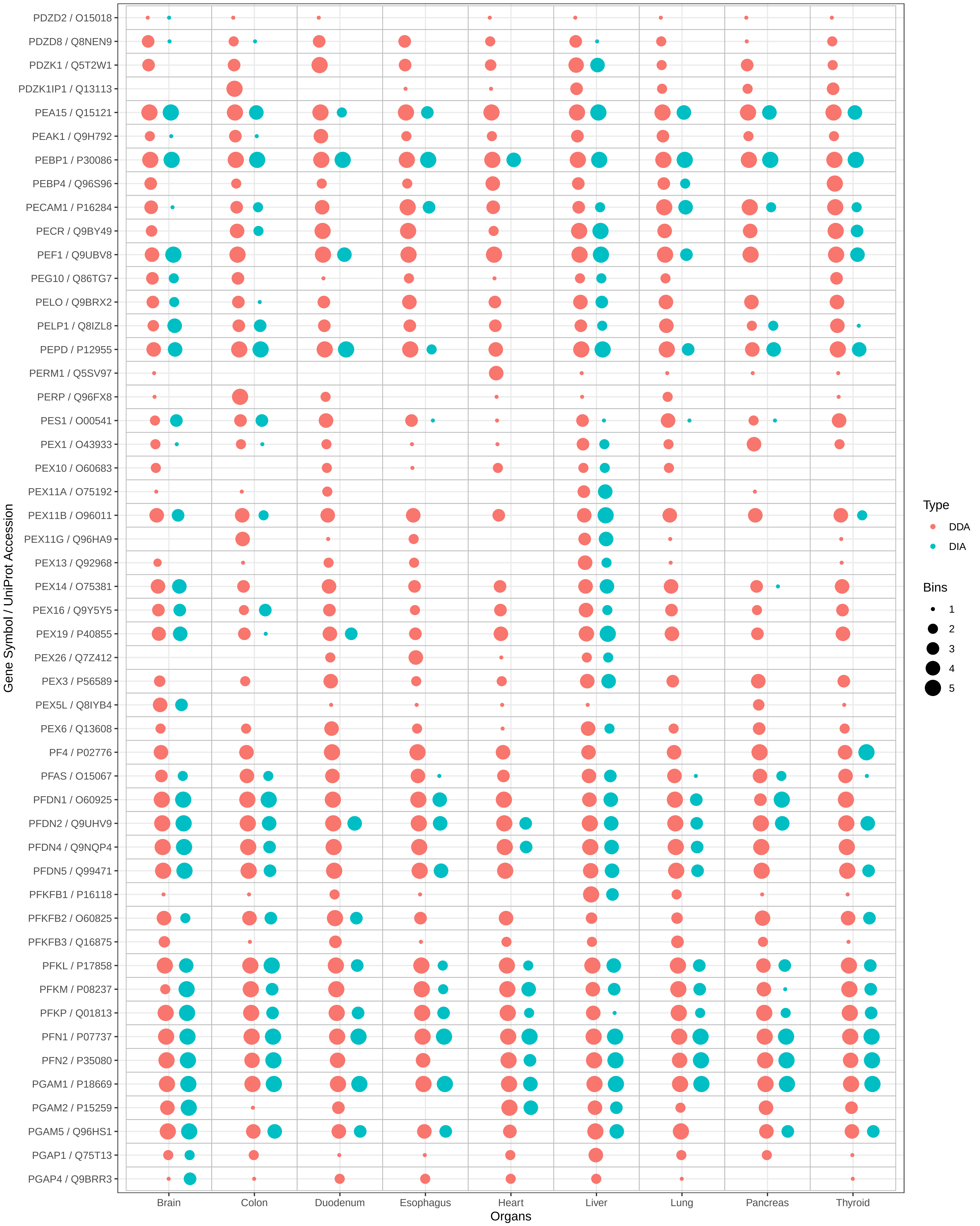

PDZD2 / O15018
PDZD8 / Q8NEN9
PDZK1 / Q5T2W1
PDZK1IP1 / Q13113
PEA15 / Q15121
PEAK1 / Q9H792
PEBP1 / P30086
PEBP4 / Q96S96
PECAM1 / P16284
PECR / Q9BY49
PEF1 / Q9UBV8
PEG10 / Q86TG7
PELO / Q9BRX2
PELP1 / Q8IZL8
PEPD / P12955
PERM1 / Q5SV97
PERP / Q96FX8
PES1 / O00541
PEX1 / O43933
PEX10 / O60683
PEX11A / O75192
PEX11B / O96011
PEX11G / Q96HA9
PEX13 / Q92968
PEX14 / O75381
PEX16 / Q9Y5Y5
PEX19 / P40855
PEX26 / Q7Z412
PEX3 / P56589
PEX5L / Q8IYB4
PEX6 / Q13608
PF4 / P02776
PFAS / O15067
PFDN1 / O60925
PFDN2 / Q9UHV9
PFDN4 / Q9NQP4
PFDN5 / Q99471
PFKFB1 / P16118
PFKFB2 / O60825
PFKFB3 / Q16875
PFKL / P17858
PFKM / P08237
PFKP / Q01813
PFN1 / P07737
PFN2 / P35080
PGAM1 / P18669
PGAM2 / P15259
PGAM5 / Q96HS1
PGAP1 / Q75T13
PGAP4 / Q9BRR3
Gene Symbol / UniProt Accession
Brain
Colon
Duodenum
Esophagus
Heart
Liver
Lung
Pancreas
Thyroid
Organs
Type
DDA
DIA
Bins
1
2
3
4
5

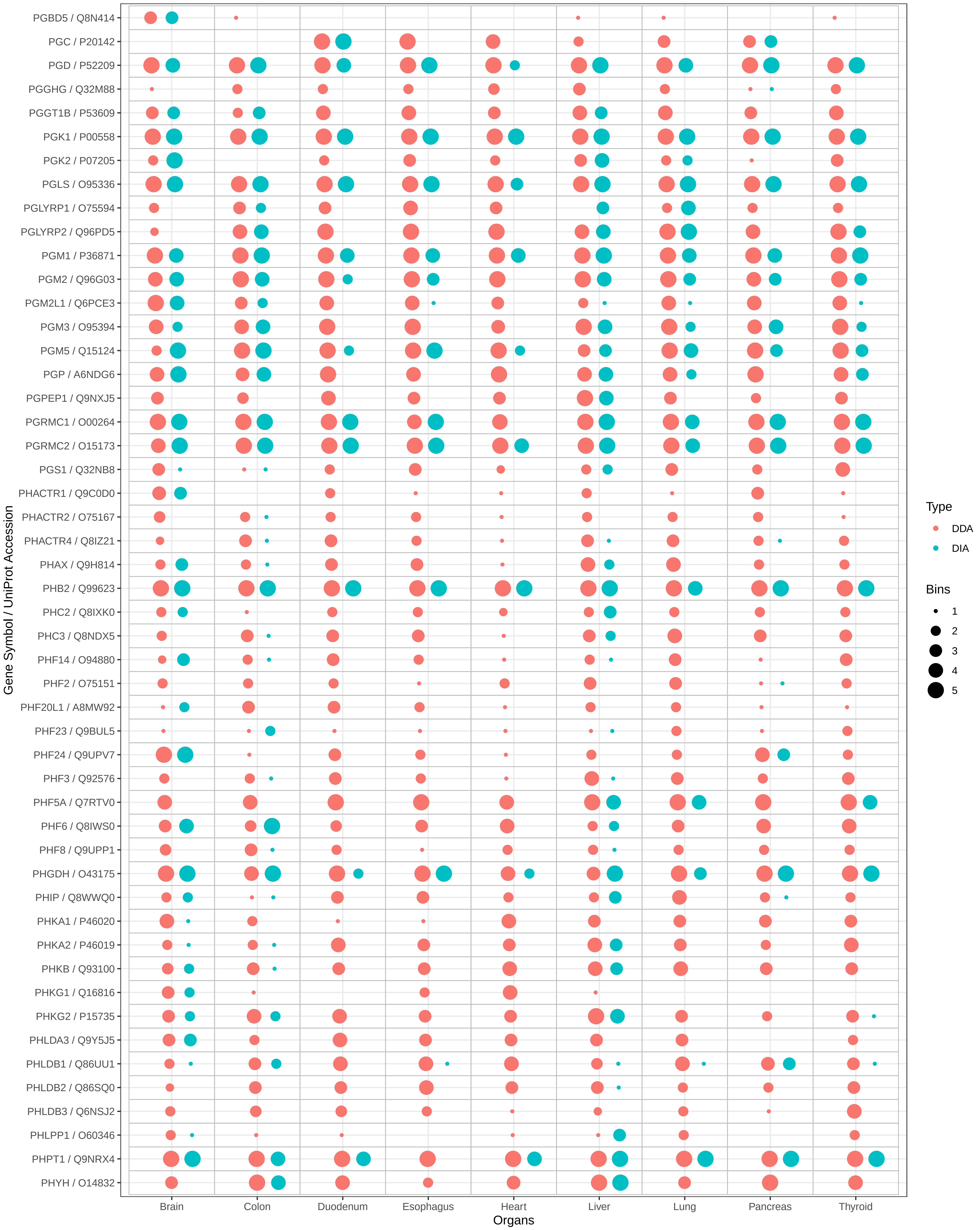
PGBD5 / Q8N414
PGC / P20142
PGD / P52209
PGGHG / Q32M88
PGGT1B / P53609
PGK1 / P00558
PGK2 / P07205
PGLS / O95336
PGLYRP1 / O75594
PGLYRP2 / Q96PD5
PGM1 / P36871
PGM2 / Q96G03
PGM2L1 / Q6PCE3
PGM3 / O95394
PGM5 / Q15124
PGP / A6NDG6
PGPEP1 / Q9NXJ5
PGRMC1 / O00264
PGRMC2 / O15173
PGS1 / Q32NB8
PHACTR1 / Q9C0D0
PHACTR2 / O75167
PHACTR4 / Q8IZ21
PHAX / Q9H814
PHB2 / Q99623
PHC2 / Q8IXK0
PHC3 / Q8NDX5
PHF14 / O94880
PHF2 / O75151
PHF20L1 / A8MW92
PHF23 / Q9BUL5
PHF24 / Q9UPV7
PHF3 / Q92576
PHF5A / Q7RTV0
PHF6 / Q8IWS0
PHF8 / Q9UPP1
PHGDH / O43175
PHIP / Q8WWQ0
PHKA1 / P46020
PHKA2 / P46019
PHKB / Q93100
PHKG1 / Q16816
PHKG2 / P15735
PHLDA3 / Q9Y5J5
PHLDB1 / Q86UU1
PHLDB2 / Q86SQ0
PHLDB3 / Q6NSJ2
PHLPP1 / O60346
PHPT1 / Q9NRX4
PHYH / O14832
Gene Symbol / UniProt Accession
Brain
Colon
Duodenum
Esophagus
Heart
Liver
Lung
Pancreas
Thyroid
Organs
Type
DDA
DIA
Bins
1
2
3
4
5

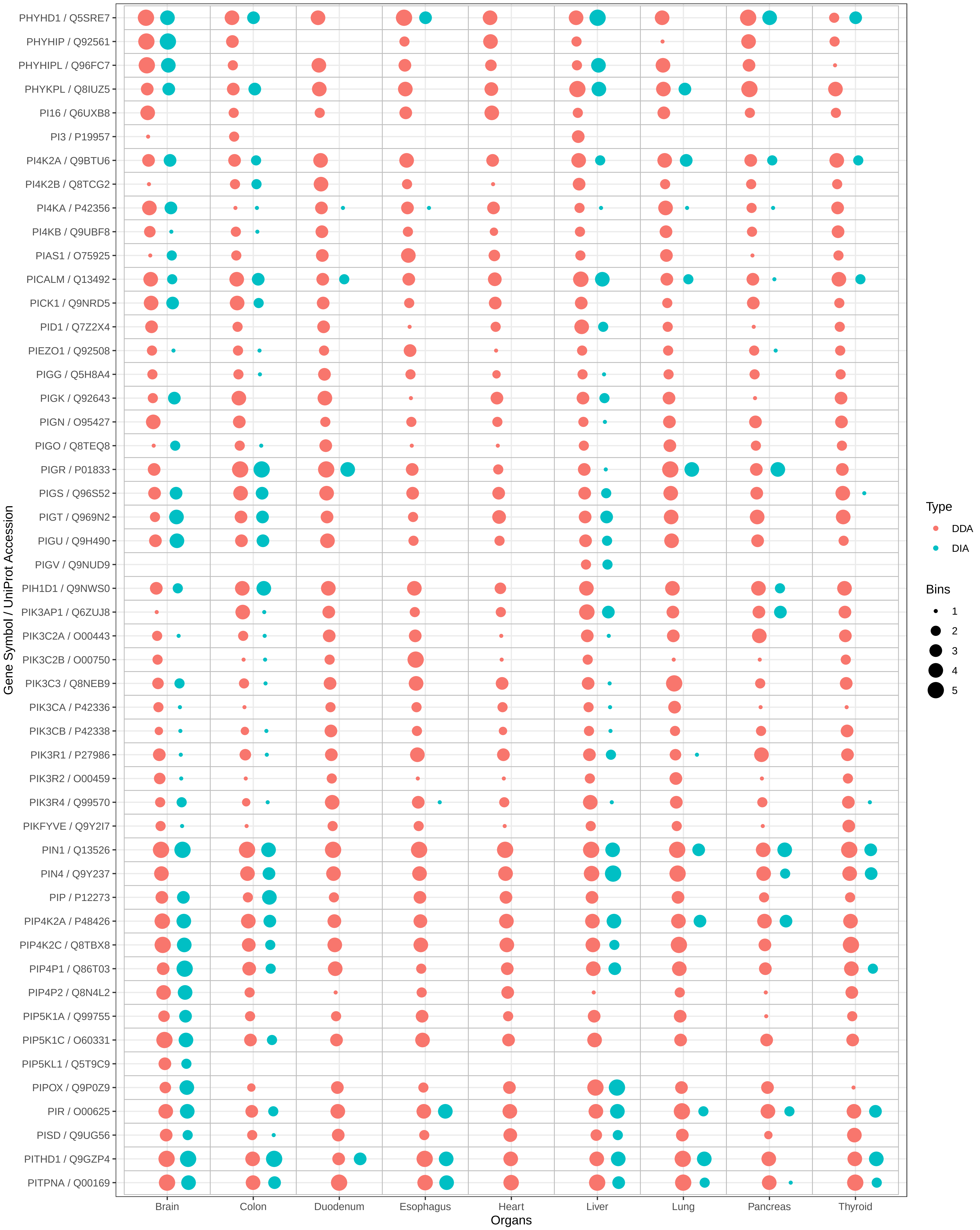
PHYHD1 / Q5SRE7
PHYHIP / Q92561
PHYHIPL / Q96FC7
PHYKPL / Q8IUZ5
PI16 / Q6UXB8
PI3 / P19957
PI4K2A / Q9BTU6
PI4K2B / Q8TCG2
PI4KA / P42356
PI4KB / Q9UBF8
PIAS1 / O75925
PICALM / Q13492
PICK1 / Q9NRD5
PID1 / Q7Z2X4
PIEZO1 / Q92508
PIGG / Q5H8A4
PIGK / Q92643
PIGN / O95427
PIGO / Q8TEQ8
PIGR / P01833
PIGS / Q96S52
PIGT / Q969N2
PIGU / Q9H490
PIGV / Q9NUD9
PIH1D1 / Q9NWS0
PIK3AP1 / Q6ZUJ8
PIK3C2A / O00443
PIK3C2B / O00750
PIK3C3 / Q8NEB9
PIK3CA / P42336
PIK3CB / P42338
PIK3R1 / P27986
PIK3R2 / O00459
PIK3R4 / Q99570
PIKFYVE / Q9Y2I7
PIN1 / Q13526
PIN4 / Q9Y237
PIP / P12273
PIP4K2A / P48426
PIP4K2C / Q8TBX8
PIP4P1 / Q86T03
PIP4P2 / Q8N4L2
PIP5K1A / Q99755
PIP5K1C / O60331
PIP5KL1 / Q5T9C9
PIPOX / Q9P0Z9
PIR / O00625
PISD / Q9UG56
PITHD1 / Q9GZP4
PITPNA / Q00169
Gene Symbol / UniProt Accession
Brain
Colon
Duodenum
Esophagus
Heart
Liver
Lung
Pancreas
Thyroid
Organs
Type
DDA
DIA
Bins
1
2
3
4
5

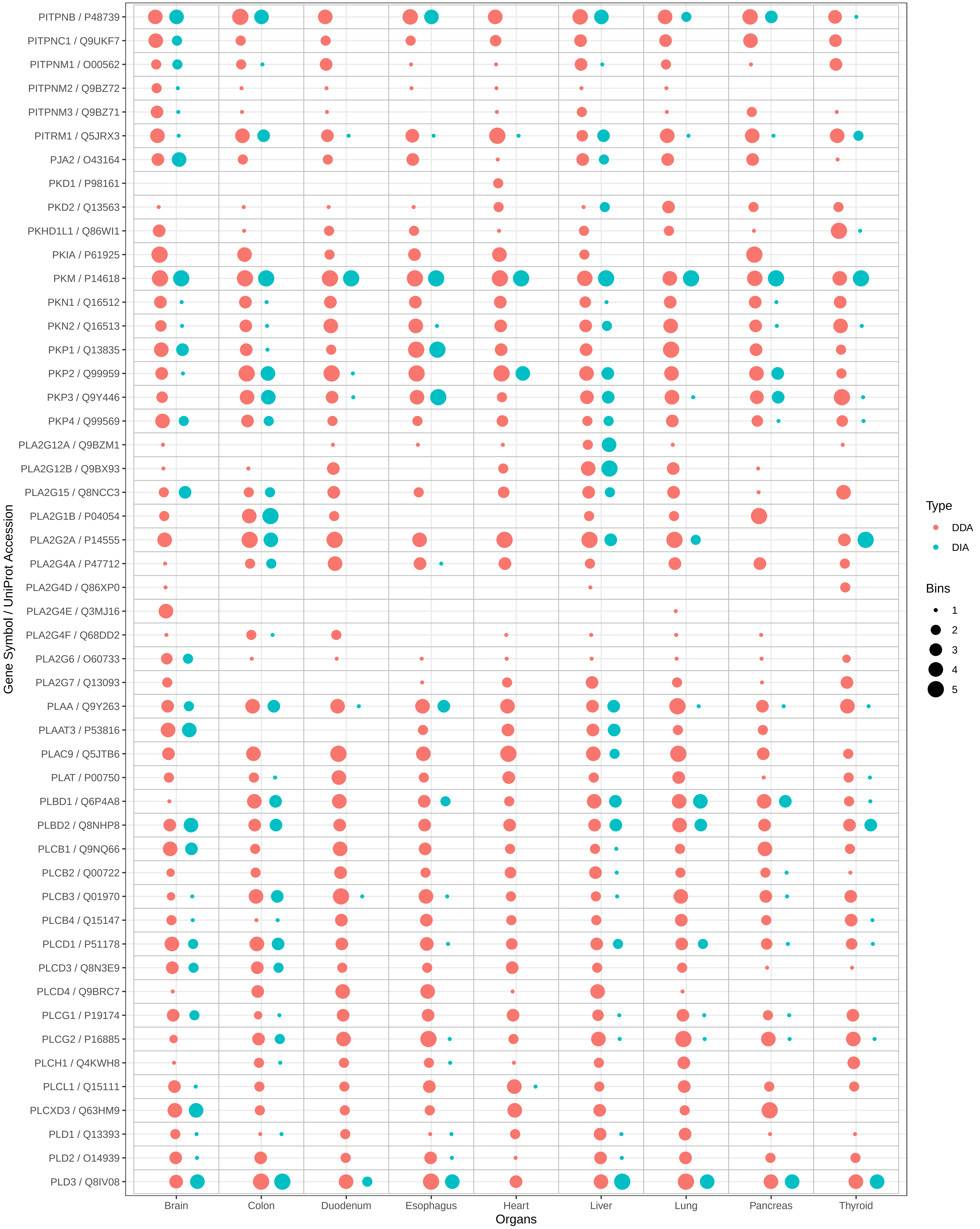

PITPNB / P48739
PITPNC1 / Q9UKF7
PITPNM1 / O00562
PITPNM2 / Q9BZ72
PITPNM3 / Q9BZ71
PITRM1 / Q5JRX3
PJA2 / O43164
PKD1 / P98161
PKD2 / Q13563
PKHD1L1 / Q86WI1
PKIA / P61925
PKM / P14618
PKN1 / Q16512
PKN2 / Q16513
PKP1 / Q13835
PKP2 / Q99959
PKP3 / Q9Y446
PKP4 / Q99569
PLA2G12A / Q9BZM1
PLA2G12B / Q9BX93
PLA2G15 / Q8NCC3
PLA2G1B / P04054
PLA2G2A / P14555
PLA2G4A / P47712
PLA2G4D / Q86XP0
PLA2G4E / Q3MJ16
PLA2G4F / Q68DD2
PLA2G6 / O60733
PLA2G7 / Q13093
PLAA / Q9Y263
PLAAT3 / P53816
PLAC9 / Q5JTB6
PLAT / P00750
PLBD1 / Q6P4A8
PLBD2 / Q8NHP8
PLCB1 / Q9NQ66
PLCB2 / Q00722
PLCB3 / Q01970
PLCB4 / Q15147
PLCD1 / P51178
PLCD3 / Q8N3E9
PLCD4 / Q9BRC7
PLCG1 / P19174
PLCG2 / P16885
PLCH1 / Q4KWH8
PLCL1 / Q15111
PLCXD3 / Q63HM9
PLD1 / Q13393
PLD2 / O14939
PLD3 / Q8IV08
Gene Symbol / UniProt Accession
Brain
Colon
Duodenum
Esophagus
Heart
Liver
Lung
Pancreas
Thyroid
Organs
Type
DDA
DIA
Bins
1
2
3
4
5

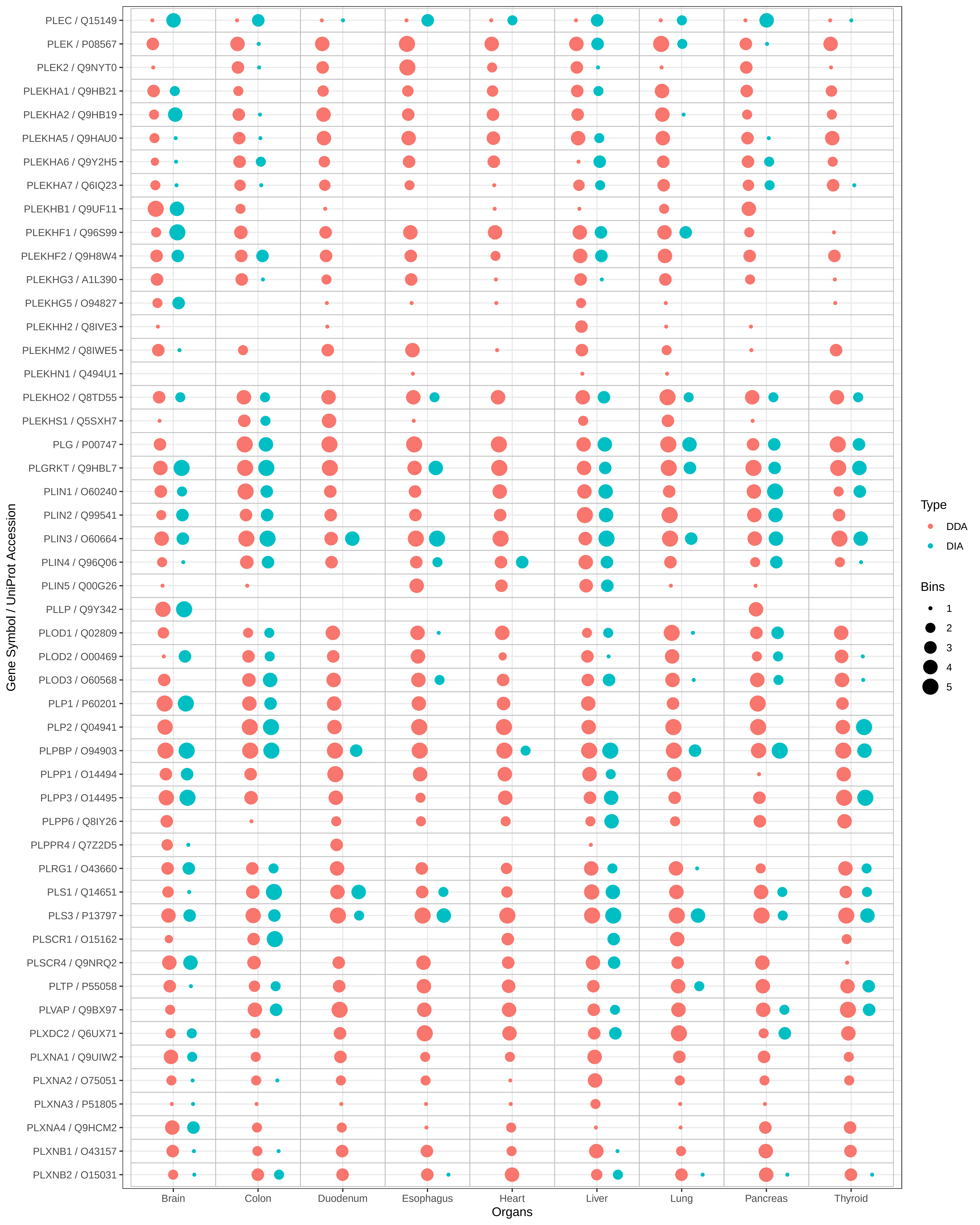

PLEC / Q15149
PLEK / P08567
PLEK2 / Q9NYT0
PLEKHA1 / Q9HB21
PLEKHA2 / Q9HB19
PLEKHA5 / Q9HAU0
PLEKHA6 / Q9Y2H5
PLEKHA7 / Q6IQ23
PLEKHB1 / Q9UF11
PLEKHF1 / Q96S99
PLEKHF2 / Q9H8W4
PLEKHG3 / A1L390
PLEKHG5 / O94827
PLEKHH2 / Q8IVE3
PLEKHM2 / Q8IWE5
PLEKHN1 / Q494U1
PLEKHO2 / Q8TD55
PLEKHS1 / Q5SXH7
PLG / P00747
PLGRKT / Q9HBL7
PLIN1 / O60240
PLIN2 / Q99541
PLIN3 / O60664
PLIN4 / Q96Q06
PLIN5 / Q00G26
PLLP / Q9Y342
PLOD1 / Q02809
PLOD2 / O00469
PLOD3 / O60568
PLP1 / P60201
PLP2 / Q04941
PLPBP / O94903
PLPP1 / O14494
PLPP3 / O14495
PLPP6 / Q8IY26
PLPPR4 / Q7Z2D5
PLRG1 / O43660
PLS1 / Q14651
PLS3 / P13797
PLSCR1 / O15162
PLSCR4 / Q9NRQ2
PLTP / P55058
PLVAP / Q9BX97
PLXDC2 / Q6UX71
PLXNA1 / Q9UIW2
PLXNA2 / O75051
PLXNA3 / P51805
PLXNA4 / Q9HCM2
PLXNB1 / O43157
PLXNB2 / O15031
Gene Symbol / UniProt Accession
Brain
Colon
Duodenum
Esophagus
Heart
Liver
Lung
Pancreas
Thyroid
Organs
Type
DDA
DIA
Bins
1
2
3
4
5

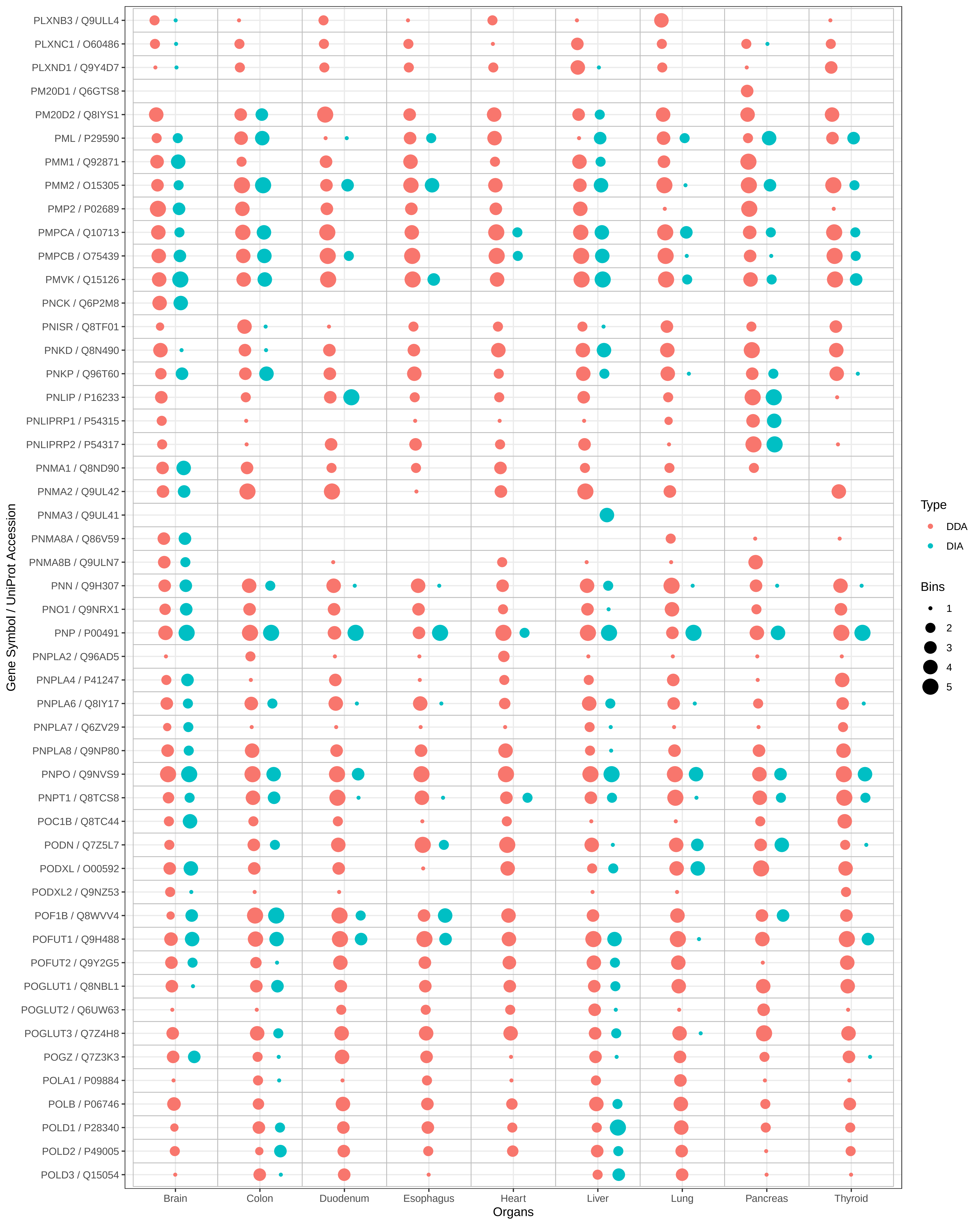

PLXNB3 / Q9ULL4
PLXNC1 / O60486
PLXND1 / Q9Y4D7
PM20D1 / Q6GTS8
PM20D2 / Q8IYS1
PML / P29590
PMM1 / Q92871
PMM2 / O15305
PMP2 / P02689
PMPCA / Q10713
PMPCB / O75439
PMVK / Q15126
PNCK / Q6P2M8
PNISR / Q8TF01
PNKD / Q8N490
PNKP / Q96T60
PNLIP / P16233
PNLIPRP1 / P54315
PNLIPRP2 / P54317
PNMA1 / Q8ND90
PNMA2 / Q9UL42
PNMA3 / Q9UL41
PNMA8A / Q86V59
PNMA8B / Q9ULN7
PNN / Q9H307
PNO1 / Q9NRX1
PNP / P00491
PNPLA2 / Q96AD5
PNPLA4 / P41247
PNPLA6 / Q8IY17
PNPLA7 / Q6ZV29
PNPLA8 / Q9NP80
PNPO / Q9NVS9
PNPT1 / Q8TCS8
POC1B / Q8TC44
PODN / Q7Z5L7
PODXL / O00592
PODXL2 / Q9NZ53
POF1B / Q8WVV4
POFUT1 / Q9H488
POFUT2 / Q9Y2G5
POGLUT1 / Q8NBL1
POGLUT2 / Q6UW63
POGLUT3 / Q7Z4H8
POGZ / Q7Z3K3
POLA1 / P09884
POLB / P06746
POLD1 / P28340
POLD2 / P49005
POLD3 / Q15054
Gene Symbol / UniProt Accession
Brain
Colon
Duodenum
Esophagus
Heart
Liver
Lung
Pancreas
Thyroid
Organs
Type
DDA
DIA
Bins
1
2
3
4
5

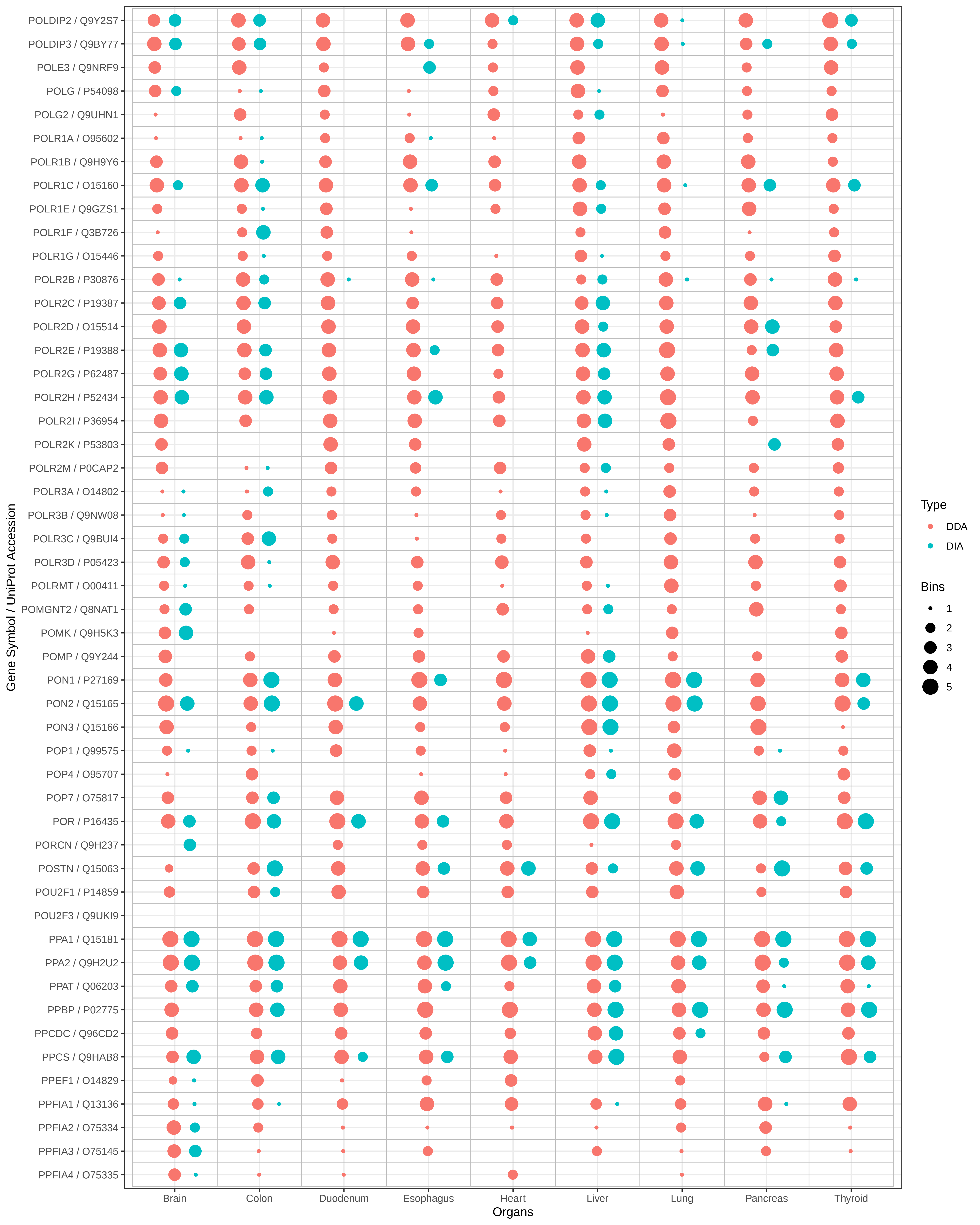

POLDIP2 / Q9Y2S7
POLDIP3 / Q9BY77
POLE3 / Q9NRF9
POLG / P54098
POLG2 / Q9UHN1
POLR1A / O95602
POLR1B / Q9H9Y6
POLR1C / O15160
POLR1E / Q9GZS1
POLR1F / Q3B726
POLR1G / O15446
POLR2B / P30876
POLR2C / P19387
POLR2D / O15514
POLR2E / P19388
POLR2G / P62487
POLR2H / P52434
POLR2I / P36954
POLR2K / P53803
POLR2M / P0CAP2
POLR3A / O14802
POLR3B / Q9NW08
POLR3C / Q9BUI4
POLR3D / P05423
POLRMT / O00411
POMGNT2 / Q8NAT1
POMK / Q9H5K3
POMP / Q9Y244
PON1 / P27169
PON2 / Q15165
PON3 / Q15166
POP1 / Q99575
POP4 / O95707
POP7 / O75817
POR / P16435
PORCN / Q9H237
POSTN / Q15063
POU2F1 / P14859
POU2F3 / Q9UKI9
PPA1 / Q15181
PPA2 / Q9H2U2
PPAT / Q06203
PPBP / P02775
PPCDC / Q96CD2
PPCS / Q9HAB8
PPEF1 / O14829
PPFIA1 / Q13136
PPFIA2 / O75334
PPFIA3 / O75145
PPFIA4 / O75335
Gene Symbol / UniProt Accession
Brain
Colon
Duodenum
Esophagus
Heart
Liver
Lung
Pancreas
Thyroid
Organs
Type
DDA
DIA
Bins
1
2
3
4
5

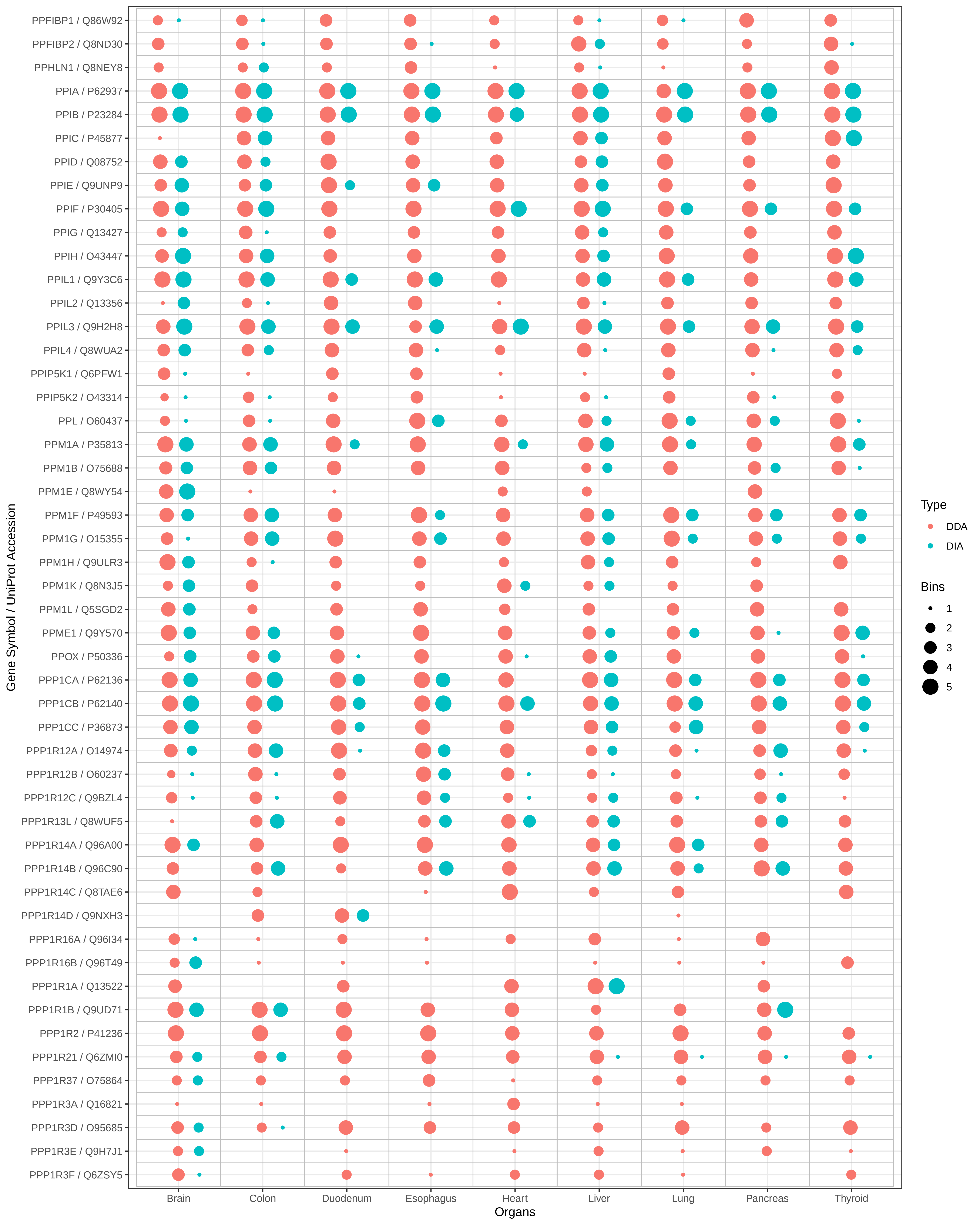

PPFIBP1 / Q86W92
PPFIBP2 / Q8ND30
PPHLN1 / Q8NEY8
PPIA / P62937
PPIB / P23284
PPIC / P45877
PPID / Q08752
PPIE / Q9UNP9
PPIF / P30405
PPIG / Q13427
PPIH / O43447
PPIL1 / Q9Y3C6
PPIL2 / Q13356
PPIL3 / Q9H2H8
PPIL4 / Q8WUA2
PPIP5K1 / Q6PFW1
PPIP5K2 / O43314
PPL / O60437
PPM1A / P35813
PPM1B / O75688
PPM1E / Q8WY54
PPM1F / P49593
PPM1G / O15355
PPM1H / Q9ULR3
PPM1K / Q8N3J5
PPM1L / Q5SGD2
PPME1 / Q9Y570
PPOX / P50336
PPP1CA / P62136
PPP1CB / P62140
PPP1CC / P36873
PPP1R12A / O14974
PPP1R12B / O60237
PPP1R12C / Q9BZL4
PPP1R13L / Q8WUF5
PPP1R14A / Q96A00
PPP1R14B / Q96C90
PPP1R14C / Q8TAE6
PPP1R14D / Q9NXH3
PPP1R16A / Q96I34
PPP1R16B / Q96T49
PPP1R1A / Q13522
PPP1R1B / Q9UD71
PPP1R2 / P41236
PPP1R21 / Q6ZMI0
PPP1R37 / O75864
PPP1R3A / Q16821
PPP1R3D / O95685
PPP1R3E / Q9H7J1
PPP1R3F / Q6ZSY5
Gene Symbol / UniProt Accession
Brain
Colon
Duodenum
Esophagus
Heart
Liver
Lung
Pancreas
Thyroid
Organs
Type
DDA
DIA
Bins
1
2
3
4
5

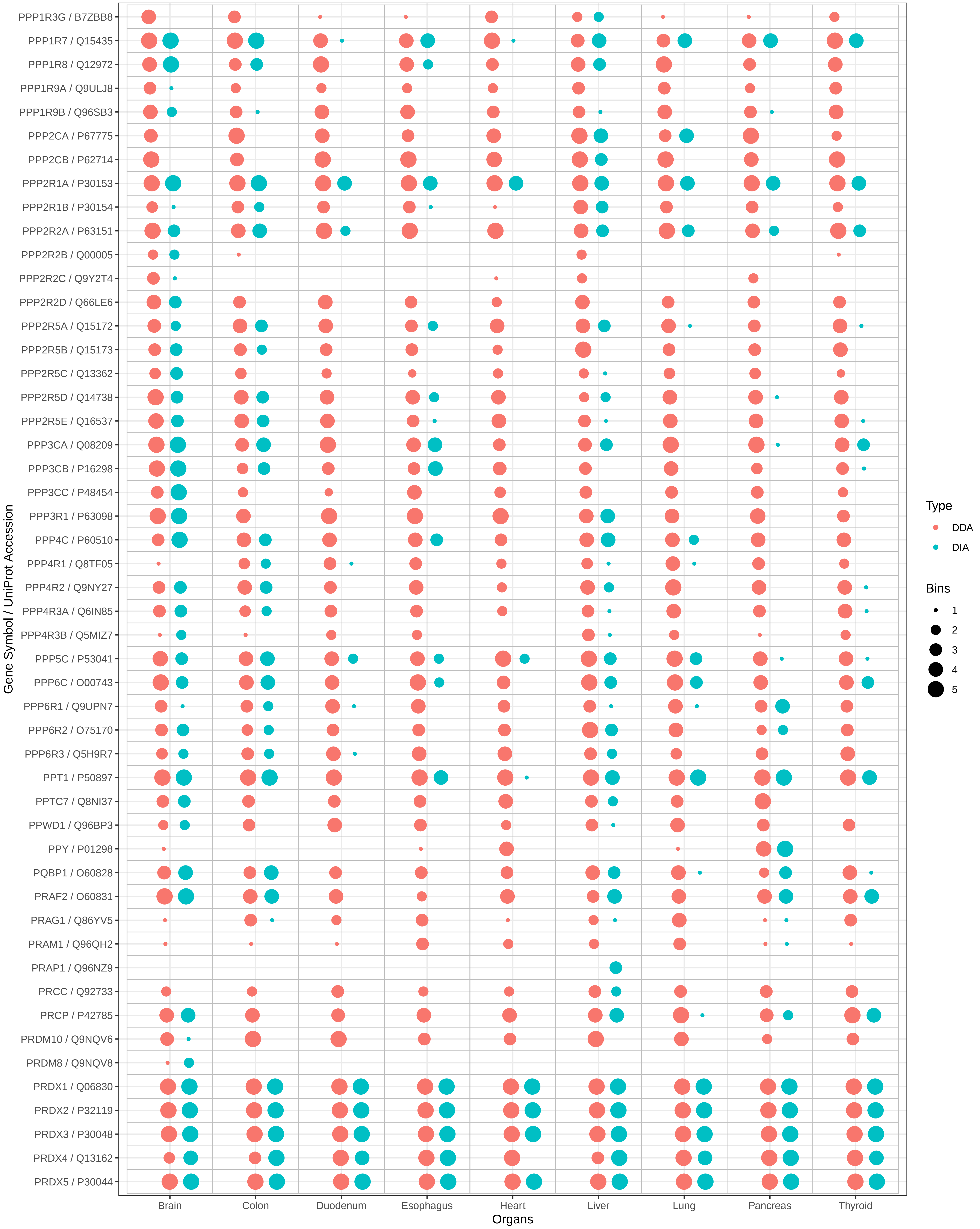

PPP1R3G / B7ZBB8
PPP1R7 / Q15435
PPP1R8 / Q12972
PPP1R9A / Q9ULJ8
PPP1R9B / Q96SB3
PPP2CA / P67775
PPP2CB / P62714
PPP2R1A / P30153
PPP2R1B / P30154
PPP2R2A / P63151
PPP2R2B / Q00005
PPP2R2C / Q9Y2T4
PPP2R2D / Q66LE6
PPP2R5A / Q15172
PPP2R5B / Q15173
PPP2R5C / Q13362
PPP2R5D / Q14738
PPP2R5E / Q16537
PPP3CA / Q08209
PPP3CB / P16298
PPP3CC / P48454
PPP3R1 / P63098
PPP4C / P60510
PPP4R1 / Q8TF05
PPP4R2 / Q9NY27
PPP4R3A / Q6IN85
PPP4R3B / Q5MIZ7
PPP5C / P53041
PPP6C / O00743
PPP6R1 / Q9UPN7
PPP6R2 / O75170
PPP6R3 / Q5H9R7
PPT1 / P50897
PPTC7 / Q8NI37
PPWD1 / Q96BP3
PPY / P01298
PQBP1 / O60828
PRAF2 / O60831
PRAG1 / Q86YV5
PRAM1 / Q96QH2
PRAP1 / Q96NZ9
PRCC / Q92733
PRCP / P42785
PRDM10 / Q9NQV6
PRDM8 / Q9NQV8
PRDX1 / Q06830
PRDX2 / P32119
PRDX3 / P30048
PRDX4 / Q13162
PRDX5 / P30044
Gene Symbol / UniProt Accession
Brain
Colon
Duodenum
Esophagus
Heart
Liver
Lung
Pancreas
Thyroid
Organs
Type
DDA
DIA
Bins
1
2
3
4
5

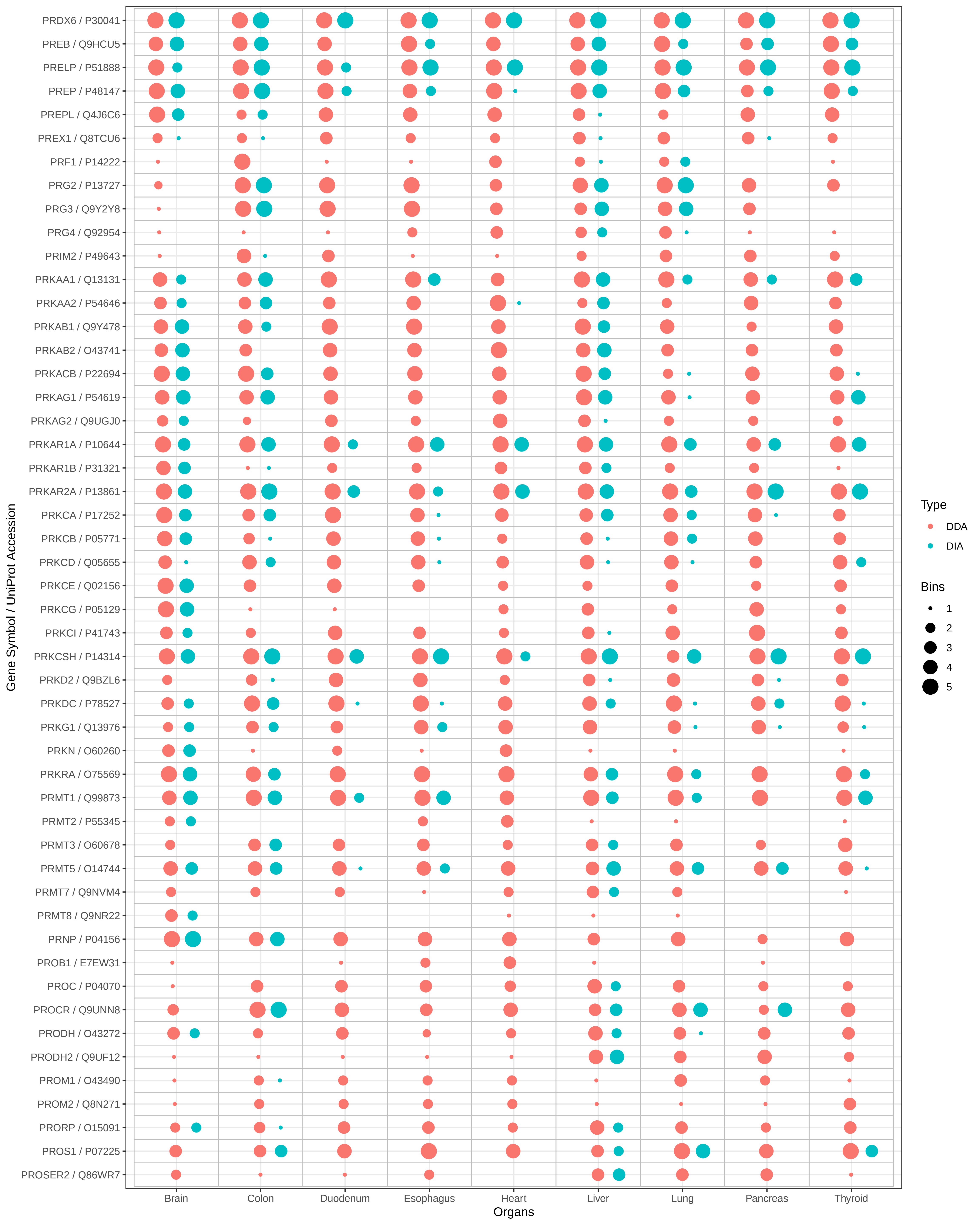

PRDX6 / P30041
PREB / Q9HCU5
PRELP / P51888
PREP / P48147
PREPL / Q4J6C6
PREX1 / Q8TCU6
PRF1 / P14222
PRG2 / P13727
PRG3 / Q9Y2Y8
PRG4 / Q92954
PRIM2 / P49643
PRKAA1 / Q13131
PRKAA2 / P54646
PRKAB1 / Q9Y478
PRKAB2 / O43741
PRKACB / P22694
PRKAG1 / P54619
PRKAG2 / Q9UGJ0
PRKAR1A / P10644
PRKAR1B / P31321
PRKAR2A / P13861
PRKCA / P17252
PRKCB / P05771
PRKCD / Q05655
PRKCE / Q02156
PRKCG / P05129
PRKCI / P41743
PRKCSH / P14314
PRKD2 / Q9BZL6
PRKDC / P78527
PRKG1 / Q13976
PRKN / O60260
PRKRA / O75569
PRMT1 / Q99873
PRMT2 / P55345
PRMT3 / O60678
PRMT5 / O14744
PRMT7 / Q9NVM4
PRMT8 / Q9NR22
PRNP / P04156
PROB1 / E7EW31
PROC / P04070
PROCR / Q9UNN8
PRODH / O43272
PRODH2 / Q9UF12
PROM1 / O43490
PROM2 / Q8N271
PRORP / O15091
PROS1 / P07225
PROSER2 / Q86WR7
Gene Symbol / UniProt Accession
Brain
Colon
Duodenum
Esophagus
Heart
Liver
Lung
Pancreas
Thyroid
Organs
Type
DDA
DIA
Bins
1
2
3
4
5

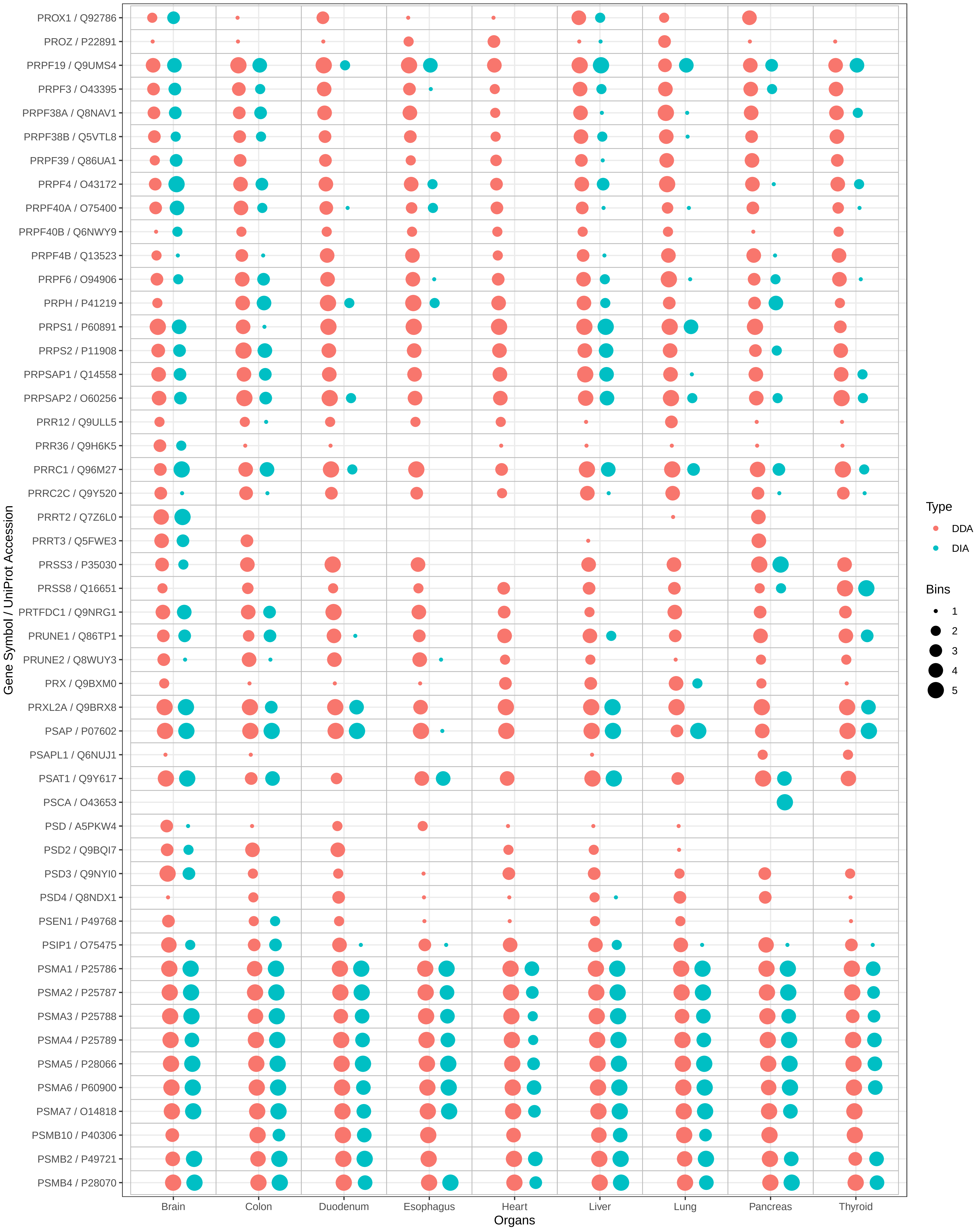
PROX1 / Q92786
PROZ / P22891
PRPF19 / Q9UMS4
PRPF3 / O43395
PRPF38A / Q8NAV1
PRPF38B / Q5VTL8
PRPF39 / Q86UA1
PRPF4 / O43172
PRPF40A / O75400
PRPF40B / Q6NWY9
PRPF4B / Q13523
PRPF6 / O94906
PRPH / P41219
PRPS1 / P60891
PRPS2 / P11908
PRPSAP1 / Q14558
PRPSAP2 / O60256
PRR12 / Q9ULL5
PRR36 / Q9H6K5
PRRC1 / Q96M27
PRRC2C / Q9Y520
PRRT2 / Q7Z6L0
PRRT3 / Q5FWE3
PRSS3 / P35030
PRSS8 / Q16651
PRTFDC1 / Q9NRG1
PRUNE1 / Q86TP1
PRUNE2 / Q8WUY3
PRX / Q9BXM0
PRXL2A / Q9BRX8
PSAP / P07602
PSAPL1 / Q6NUJ1
PSAT1 / Q9Y617
PSCA / O43653
PSD / A5PKW4
PSD2 / Q9BQI7
PSD3 / Q9NYI0
PSD4 / Q8NDX1
PSEN1 / P49768
PSIP1 / O75475
PSMA1 / P25786
PSMA2 / P25787
PSMA3 / P25788
PSMA4 / P25789
PSMA5 / P28066
PSMA6 / P60900
PSMA7 / O14818
PSMB10 / P40306
PSMB2 / P49721
PSMB4 / P28070
Gene Symbol / UniProt Accession
Brain
Colon
Duodenum
Esophagus
Heart
Liver
Lung
Pancreas
Thyroid
Organs
Type
DDA
DIA
Bins
1
2
3
4
5

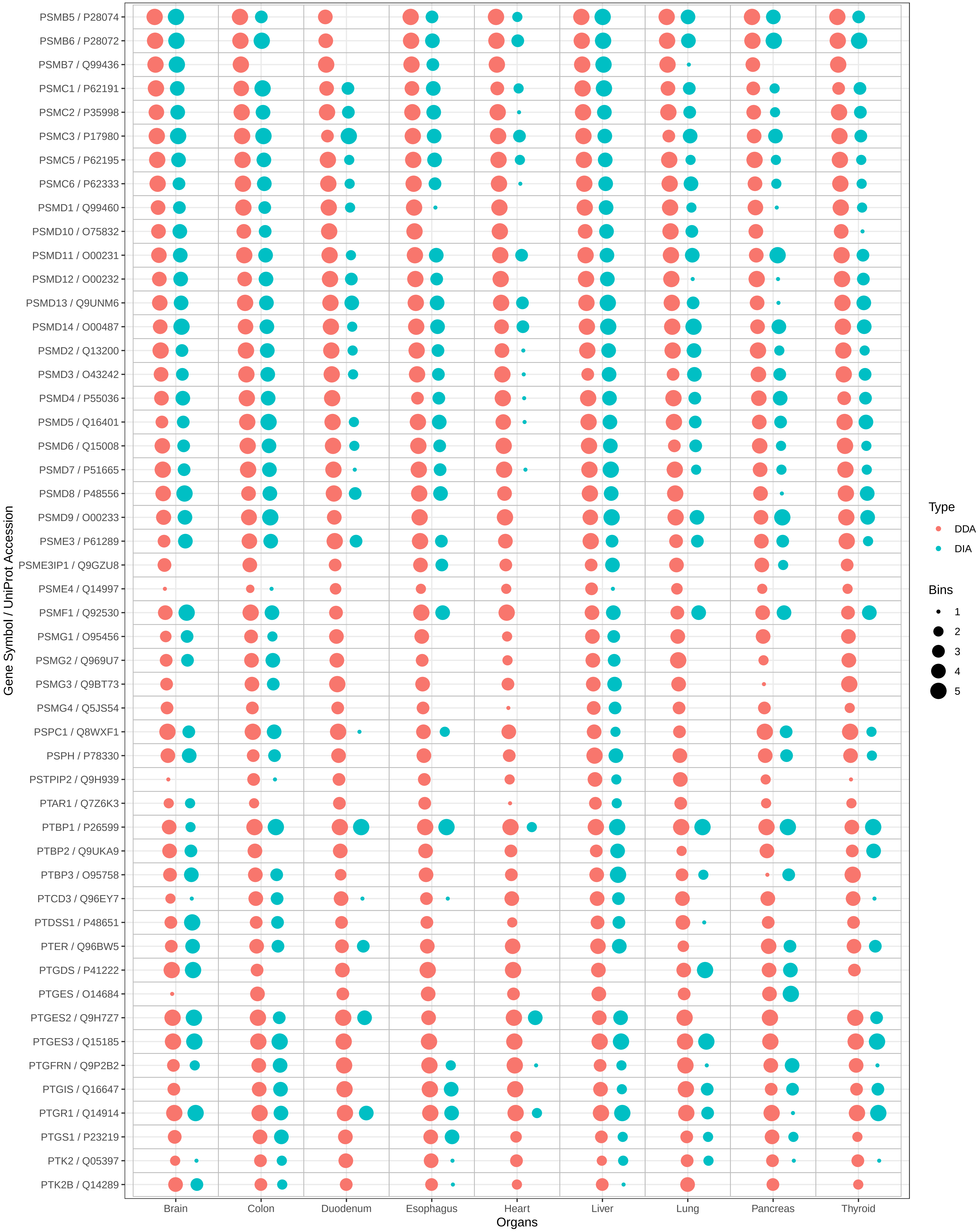
PSMB5 / P28074
PSMB6 / P28072
PSMB7 / Q99436
PSMC1 / P62191
PSMC2 / P35998
PSMC3 / P17980
PSMC5 / P62195
PSMC6 / P62333
PSMD1 / Q99460
PSMD10 / O75832
PSMD11 / O00231
PSMD12 / O00232
PSMD13 / Q9UNM6
PSMD14 / O00487
PSMD2 / Q13200
PSMD3 / O43242
PSMD4 / P55036
PSMD5 / Q16401
PSMD6 / Q15008
PSMD7 / P51665
PSMD8 / P48556
PSMD9 / O00233
PSME3 / P61289
PSME3IP1 / Q9GZU8
PSME4 / Q14997
PSMF1 / Q92530
PSMG1 / O95456
PSMG2 / Q969U7
PSMG3 / Q9BT73
PSMG4 / Q5JS54
PSPC1 / Q8WXF1
PSPH / P78330
PSTPIP2 / Q9H939
PTAR1 / Q7Z6K3
PTBP1 / P26599
PTBP2 / Q9UKA9
PTBP3 / O95758
PTCD3 / Q96EY7
PTDSS1 / P48651
PTER / Q96BW5
PTGDS / P41222
PTGES / O14684
PTGES2 / Q9H7Z7
PTGES3 / Q15185
PTGFRN / Q9P2B2
PTGIS / Q16647
PTGR1 / Q14914
PTGS1 / P23219
PTK2 / Q05397
PTK2B / Q14289
Gene Symbol / UniProt Accession
Brain
Colon
Duodenum
Esophagus
Heart
Liver
Lung
Pancreas
Thyroid
Organs
Type
DDA
DIA
Bins
1
2
3
4
5

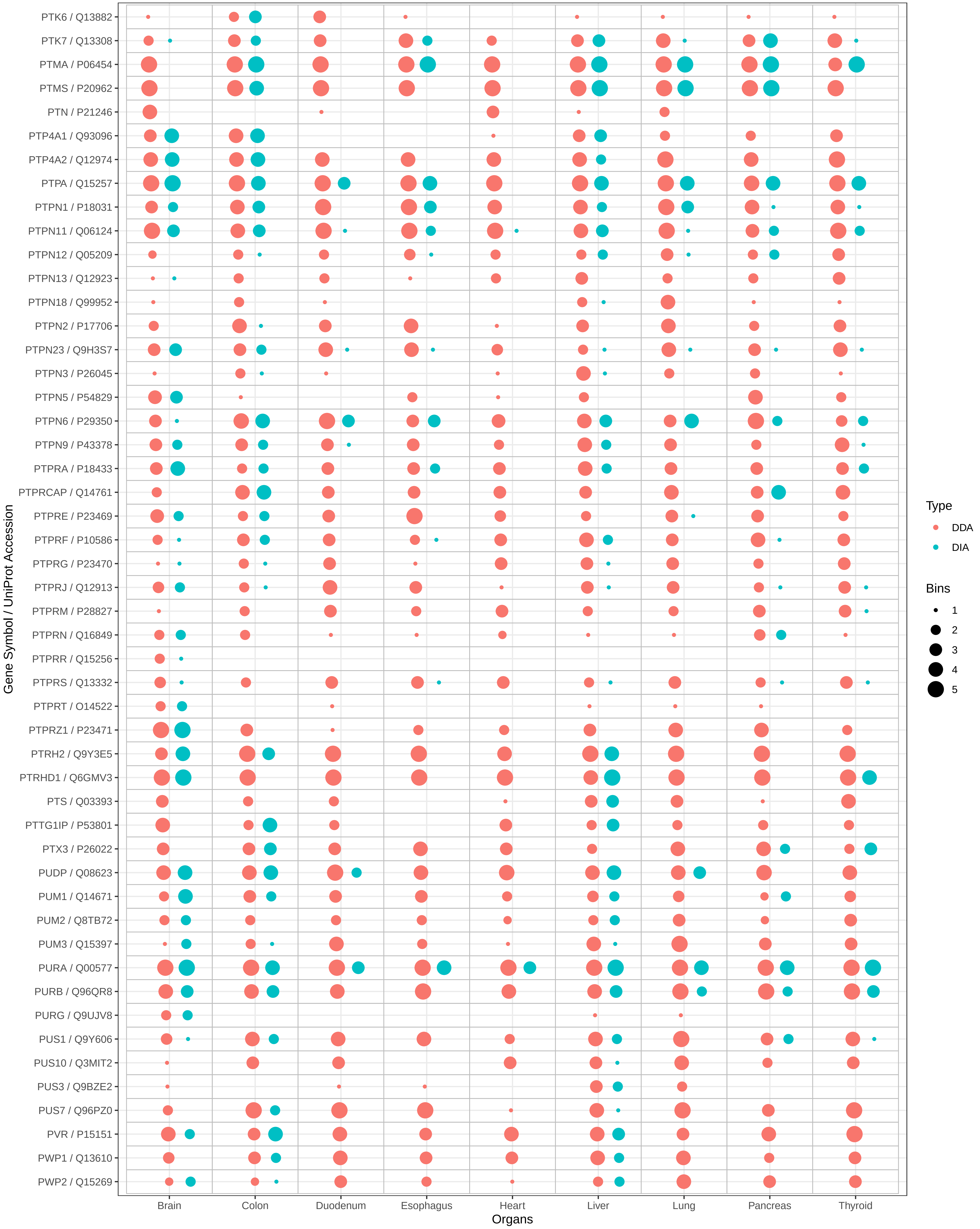

PTK6 / Q13882
PTK7 / Q13308
PTMA / P06454
PTMS / P20962
PTN / P21246
PTP4A1 / Q93096
PTP4A2 / Q12974
PTPA / Q15257
PTPN1 / P18031
PTPN11 / Q06124
PTPN12 / Q05209
PTPN13 / Q12923
PTPN18 / Q99952
PTPN2 / P17706
PTPN23 / Q9H3S7
PTPN3 / P26045
PTPN5 / P54829
PTPN6 / P29350
PTPN9 / P43378
PTPRA / P18433
PTPRCAP / Q14761
PTPRE / P23469
PTPRF / P10586
PTPRG / P23470
PTPRJ / Q12913
PTPRM / P28827
PTPRN / Q16849
PTPRR / Q15256
PTPRS / Q13332
PTPRT / O14522
PTPRZ1 / P23471
PTRH2 / Q9Y3E5
PTRHD1 / Q6GMV3
PTS / Q03393
PTTG1IP / P53801
PTX3 / P26022
PUDP / Q08623
PUM1 / Q14671
PUM2 / Q8TB72
PUM3 / Q15397
PURA / Q00577
PURB / Q96QR8
PURG / Q9UJV8
PUS1 / Q9Y606
PUS10 / Q3MIT2
PUS3 / Q9BZE2
PUS7 / Q96PZ0
PVR / P15151
PWP1 / Q13610
PWP2 / Q15269
Gene Symbol / UniProt Accession
Brain
Colon
Duodenum
Esophagus
Heart
Liver
Lung
Pancreas
Thyroid
Organs
Type
DDA
DIA
Bins
1
2
3
4
5

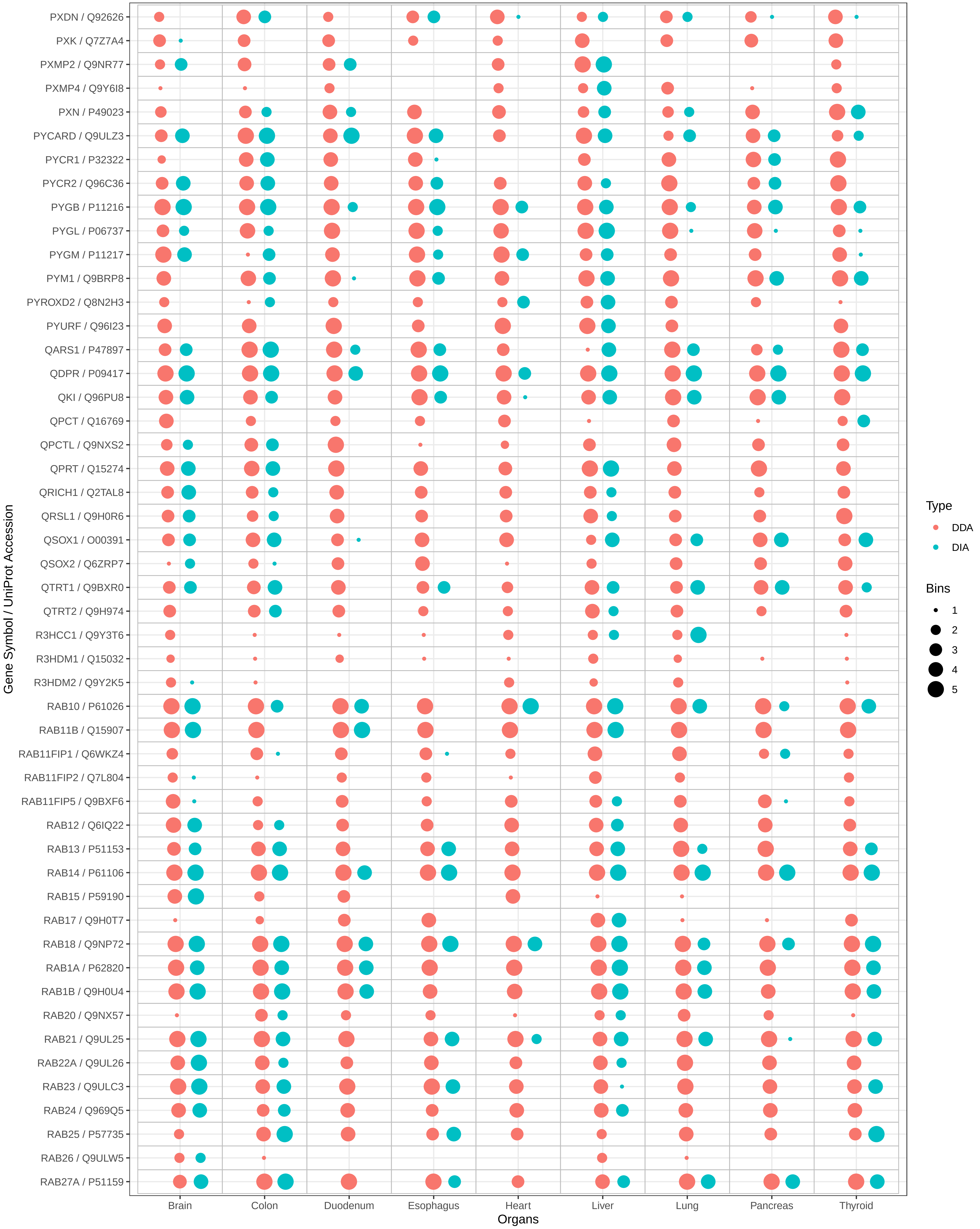

PXDN / Q92626
PXK / Q7Z7A4
PXMP2 / Q9NR77
PXMP4 / Q9Y6I8
PXN / P49023
PYCARD / Q9ULZ3
PYCR1 / P32322
PYCR2 / Q96C36
PYGB / P11216
PYGL / P06737
PYGM / P11217
PYM1 / Q9BRP8
PYROXD2 / Q8N2H3
PYURF / Q96I23
QARS1 / P47897
QDPR / P09417
QKI / Q96PU8
QPCT / Q16769
QPCTL / Q9NXS2
QPRT / Q15274
QRICH1 / Q2TAL8
QRSL1 / Q9H0R6
QSOX1 / O00391
QSOX2 / Q6ZRP7
QTRT1 / Q9BXR0
QTRT2 / Q9H974
R3HCC1 / Q9Y3T6
R3HDM1 / Q15032
R3HDM2 / Q9Y2K5
RAB10 / P61026
RAB11B / Q15907
RAB11FIP1 / Q6WKZ4
RAB11FIP2 / Q7L804
RAB11FIP5 / Q9BXF6
RAB12 / Q6IQ22
RAB13 / P51153
RAB14 / P61106
RAB15 / P59190
RAB17 / Q9H0T7
RAB18 / Q9NP72
RAB1A / P62820
RAB1B / Q9H0U4
RAB20 / Q9NX57
RAB21 / Q9UL25
RAB22A / Q9UL26
RAB23 / Q9ULC3
RAB24 / Q969Q5
RAB25 / P57735
RAB26 / Q9ULW5
RAB27A / P51159
Gene Symbol / UniProt Accession
Brain
Colon
Duodenum
Esophagus
Heart
Liver
Lung
Pancreas
Thyroid
Organs
Type
DDA
DIA
Bins
1
2
3
4
5

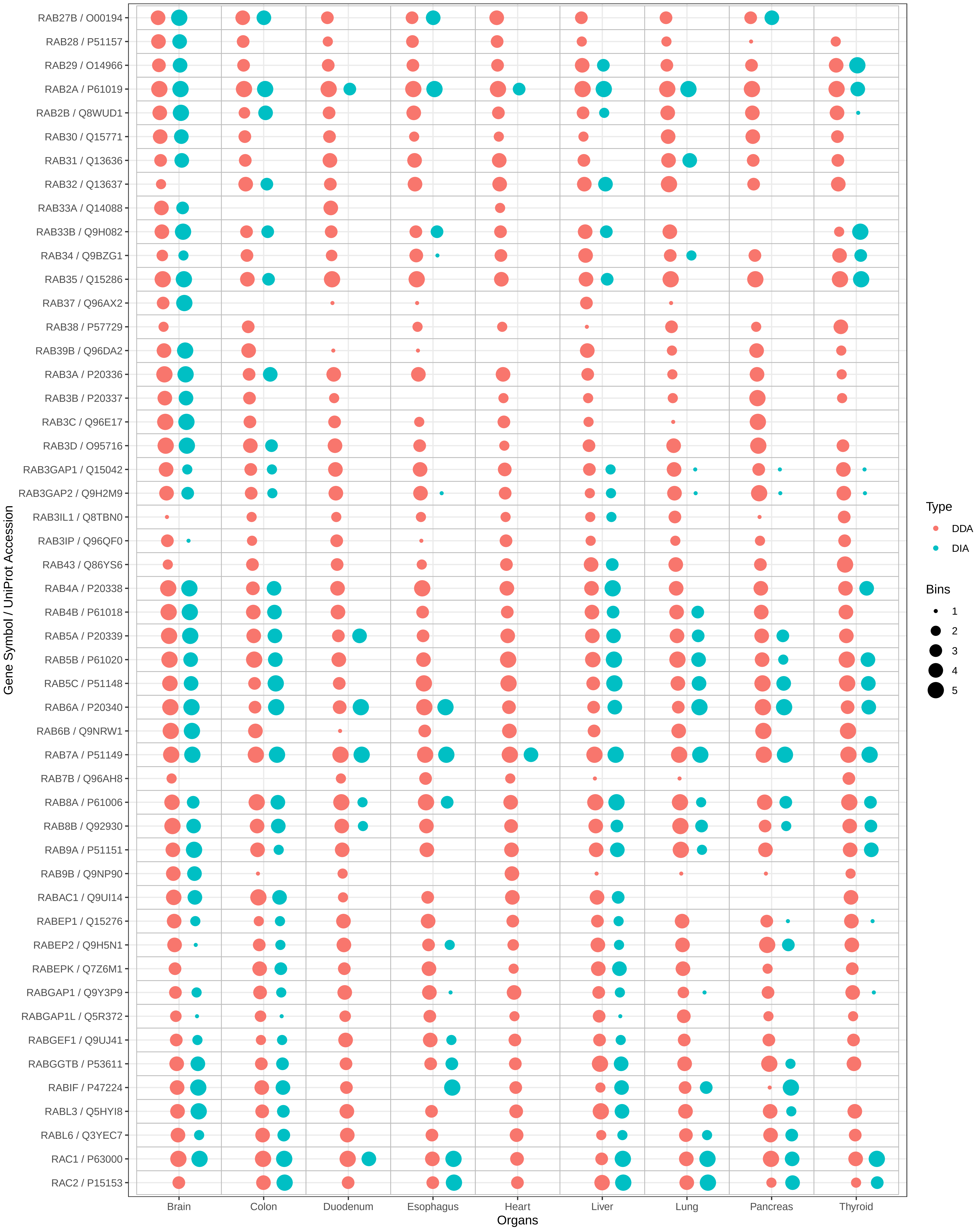
RAB27B / O00194
RAB28 / P51157
RAB29 / O14966
RAB2A / P61019
RAB2B / Q8WUD1
RAB30 / Q15771
RAB31 / Q13636
RAB32 / Q13637
RAB33A / Q14088
RAB33B / Q9H082
RAB34 / Q9BZG1
RAB35 / Q15286
RAB37 / Q96AX2
RAB38 / P57729
RAB39B / Q96DA2
RAB3A / P20336
RAB3B / P20337
RAB3C / Q96E17
RAB3D / O95716
RAB3GAP1 / Q15042
RAB3GAP2 / Q9H2M9
RAB3IL1 / Q8TBN0
RAB3IP / Q96QF0
RAB43 / Q86YS6
RAB4A / P20338
RAB4B / P61018
RAB5A / P20339
RAB5B / P61020
RAB5C / P51148
RAB6A / P20340
RAB6B / Q9NRW1
RAB7A / P51149
RAB7B / Q96AH8
RAB8A / P61006
RAB8B / Q92930
RAB9A / P51151
RAB9B / Q9NP90
RABAC1 / Q9UI14
RABEP1 / Q15276
RABEP2 / Q9H5N1
RABEPK / Q7Z6M1
RABGAP1 / Q9Y3P9
RABGAP1L / Q5R372
RABGEF1 / Q9UJ41
RABGGTB / P53611
RABIF / P47224
RABL3 / Q5HYI8
RABL6 / Q3YEC7
RAC1 / P63000
RAC2 / P15153
Gene Symbol / UniProt Accession
Brain
Colon
Duodenum
Esophagus
Heart
Liver
Lung
Pancreas
Thyroid
Organs
Type
DDA
DIA
Bins
1
2
3
4
5

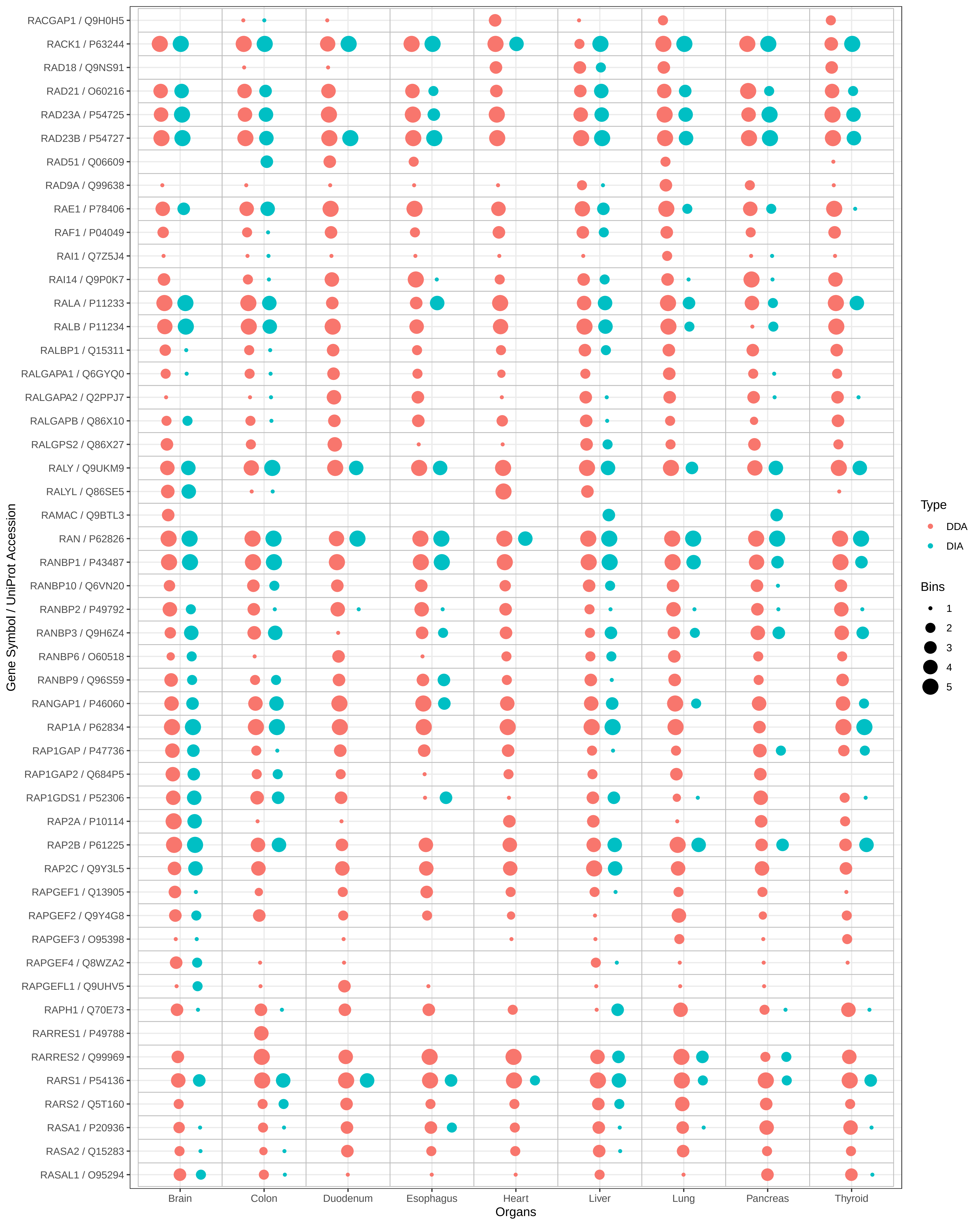

RACGAP1 / Q9H0H5
RACK1 / P63244
RAD18 / Q9NS91
RAD21 / O60216
RAD23A / P54725
RAD23B / P54727
RAD51 / Q06609
RAD9A / Q99638
RAE1 / P78406
RAF1 / P04049
RAI1 / Q7Z5J4
RAI14 / Q9P0K7
RALA / P11233
RALB / P11234
RALBP1 / Q15311
RALGAPA1 / Q6GYQ0
RALGAPA2 / Q2PPJ7
RALGAPB / Q86X10
RALGPS2 / Q86X27
RALY / Q9UKM9
RALYL / Q86SE5
RAMAC / Q9BTL3
RAN / P62826
RANBP1 / P43487
RANBP10 / Q6VN20
RANBP2 / P49792
RANBP3 / Q9H6Z4
RANBP6 / O60518
RANBP9 / Q96S59
RANGAP1 / P46060
RAP1A / P62834
RAP1GAP / P47736
RAP1GAP2 / Q684P5
RAP1GDS1 / P52306
RAP2A / P10114
RAP2B / P61225
RAP2C / Q9Y3L5
RAPGEF1 / Q13905
RAPGEF2 / Q9Y4G8
RAPGEF3 / O95398
RAPGEF4 / Q8WZA2
RAPGEFL1 / Q9UHV5
RAPH1 / Q70E73
RARRES1 / P49788
RARRES2 / Q99969
RARS1 / P54136
RARS2 / Q5T160
RASA1 / P20936
RASA2 / Q15283
RASAL1 / O95294
Gene Symbol / UniProt Accession
Brain
Colon
Duodenum
Esophagus
Heart
Liver
Lung
Pancreas
Thyroid
Organs
Type
DDA
DIA
Bins
1
2
3
4
5

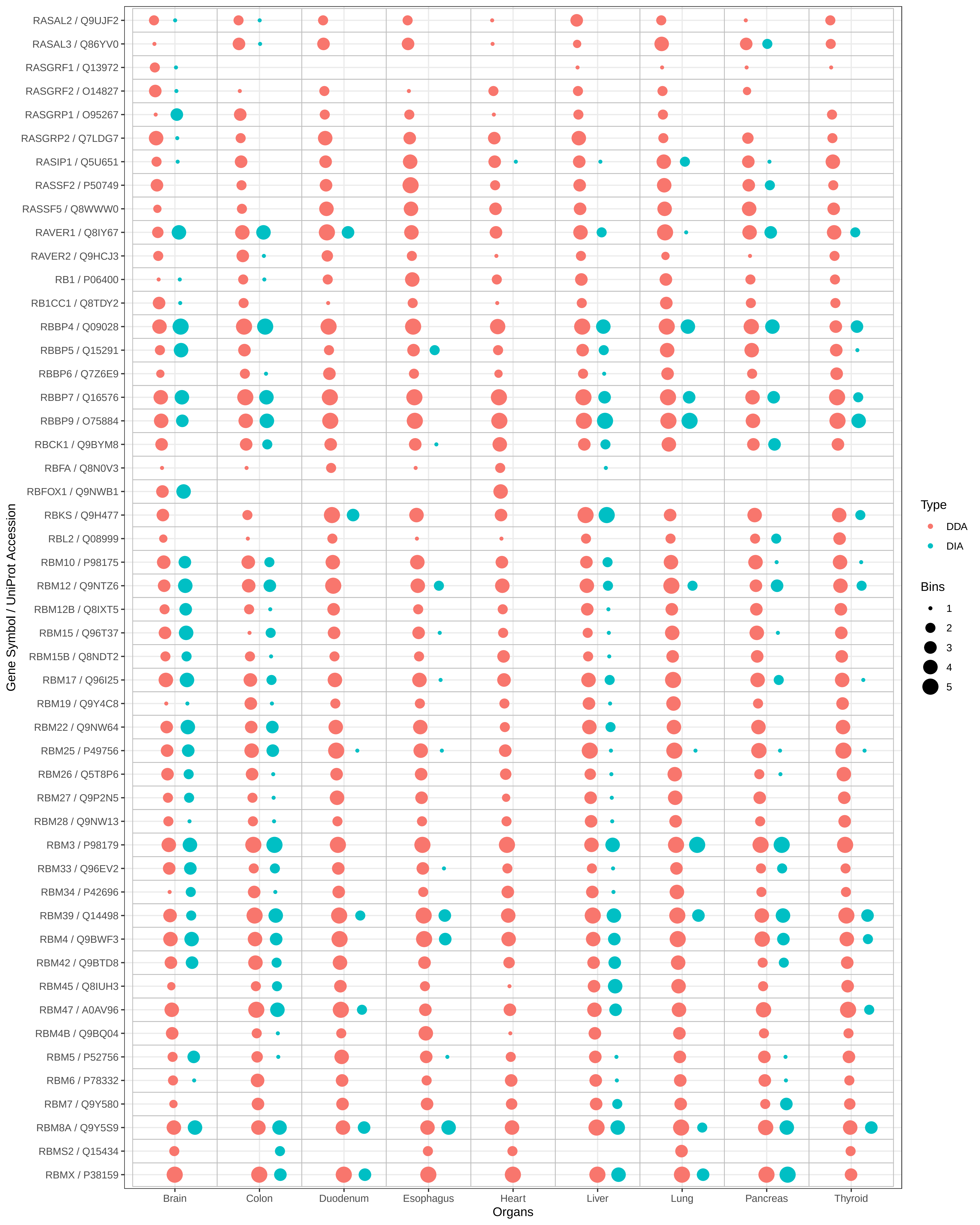
RASAL2 / Q9UJF2
RASAL3 / Q86YV0
RASGRF1 / Q13972
RASGRF2 / O14827
RASGRP1 / O95267
RASGRP2 / Q7LDG7
RASIP1 / Q5U651
RASSF2 / P50749
RASSF5 / Q8WWW0
RAVER1 / Q8IY67
RAVER2 / Q9HCJ3
RB1 / P06400
RB1CC1 / Q8TDY2
RBBP4 / Q09028
RBBP5 / Q15291
RBBP6 / Q7Z6E9
RBBP7 / Q16576
RBBP9 / O75884
RBCK1 / Q9BYM8
RBFA / Q8N0V3
RBFOX1 / Q9NWB1
RBKS / Q9H477
RBL2 / Q08999
RBM10 / P98175
RBM12 / Q9NTZ6
RBM12B / Q8IXT5
RBM15 / Q96T37
RBM15B / Q8NDT2
RBM17 / Q96I25
RBM19 / Q9Y4C8
RBM22 / Q9NW64
RBM25 / P49756
RBM26 / Q5T8P6
RBM27 / Q9P2N5
RBM28 / Q9NW13
RBM3 / P98179
RBM33 / Q96EV2
RBM34 / P42696
RBM39 / Q14498
RBM4 / Q9BWF3
RBM42 / Q9BTD8
RBM45 / Q8IUH3
RBM47 / A0AV96
RBM4B / Q9BQ04
RBM5 / P52756
RBM6 / P78332
RBM7 / Q9Y580
RBM8A / Q9Y5S9
RBMS2 / Q15434
RBMX / P38159
Gene Symbol / UniProt Accession
Brain
Colon
Duodenum
Esophagus
Heart
Liver
Lung
Pancreas
Thyroid
Organs
Type
DDA
DIA
Bins
1
2
3
4
5

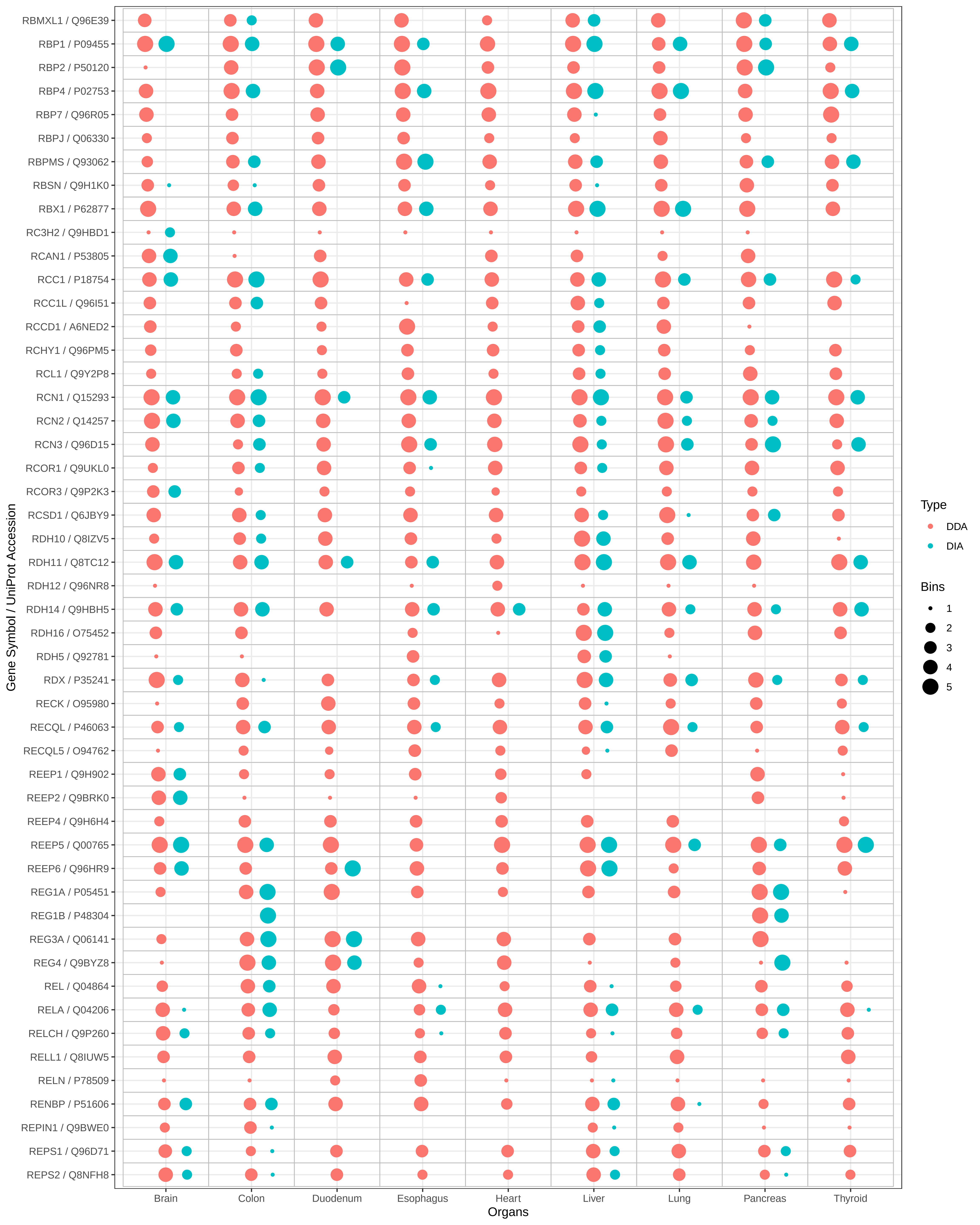
RBMXL1 / Q96E39
RBP1 / P09455
RBP2 / P50120
RBP4 / P02753
RBP7 / Q96R05
RBPJ / Q06330
RBPMS / Q93062
RBSN / Q9H1K0
RBX1 / P62877
RC3H2 / Q9HBD1
RCAN1 / P53805
RCC1 / P18754
RCC1L / Q96I51
RCCD1 / A6NED2
RCHY1 / Q96PM5
RCL1 / Q9Y2P8
RCN1 / Q15293
RCN2 / Q14257
RCN3 / Q96D15
RCOR1 / Q9UKL0
RCOR3 / Q9P2K3
RCSD1 / Q6JBY9
RDH10 / Q8IZV5
RDH11 / Q8TC12
RDH12 / Q96NR8
RDH14 / Q9HBH5
RDH16 / O75452
RDH5 / Q92781
RDX / P35241
RECK / O95980
RECQL / P46063
RECQL5 / O94762
REEP1 / Q9H902
REEP2 / Q9BRK0
REEP4 / Q9H6H4
REEP5 / Q00765
REEP6 / Q96HR9
REG1A / P05451
REG1B / P48304
REG3A / Q06141
REG4 / Q9BYZ8
REL / Q04864
RELA / Q04206
RELCH / Q9P260
RELL1 / Q8IUW5
RELN / P78509
RENBP / P51606
REPIN1 / Q9BWE0
REPS1 / Q96D71
REPS2 / Q8NFH8
Gene Symbol / UniProt Accession
Brain
Colon
Duodenum
Esophagus
Heart
Liver
Lung
Pancreas
Thyroid
Organs
Type
DDA
DIA
Bins
1
2
3
4
5

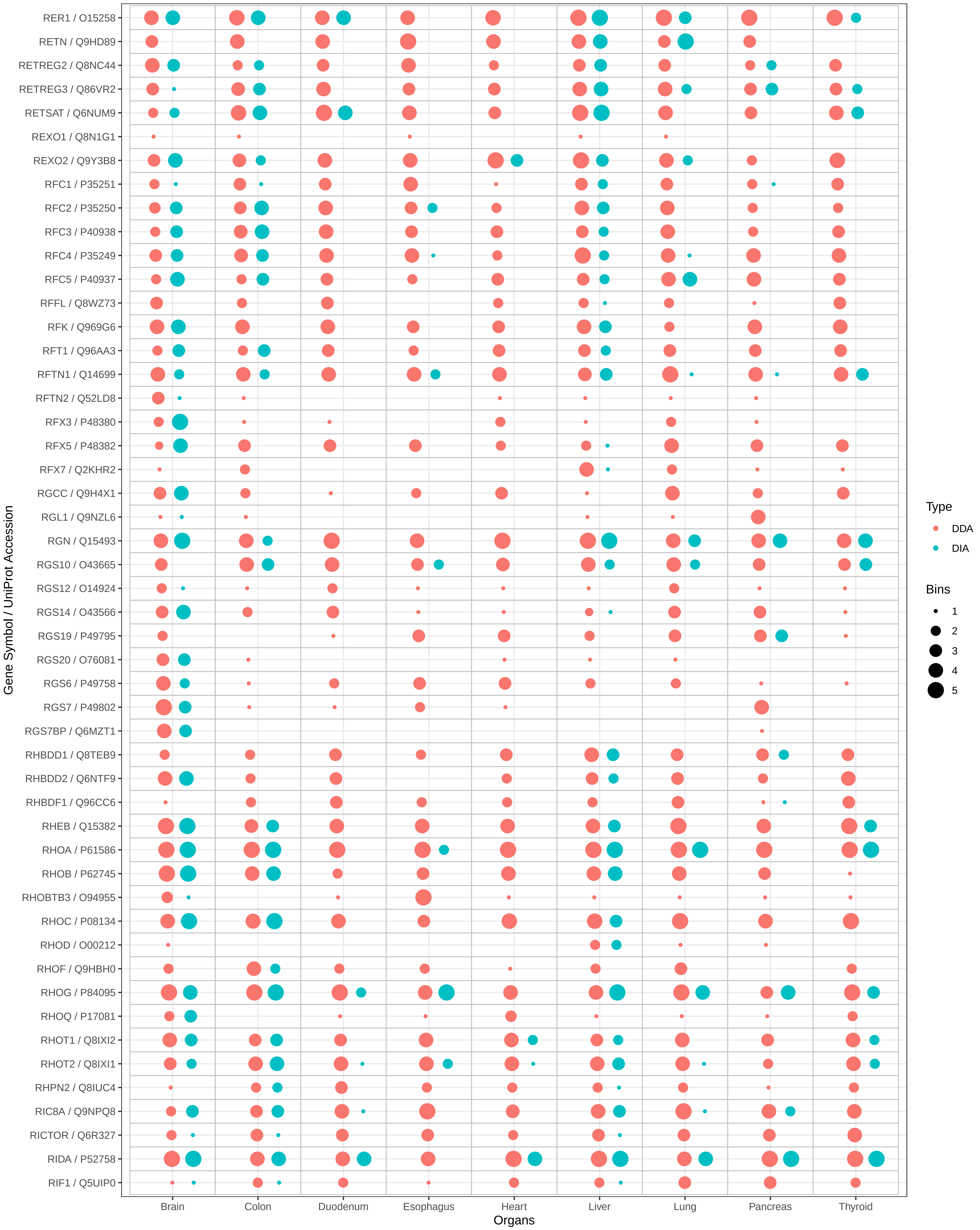

RER1 / O15258
RETN / Q9HD89
RETREG2 / Q8NC44
RETREG3 / Q86VR2
RETSAT / Q6NUM9
REXO1 / Q8N1G1
REXO2 / Q9Y3B8
RFC1 / P35251
RFC2 / P35250
RFC3 / P40938
RFC4 / P35249
RFC5 / P40937
RFFL / Q8WZ73
RFK / Q969G6
RFT1 / Q96AA3
RFTN1 / Q14699
RFTN2 / Q52LD8
RFX3 / P48380
RFX5 / P48382
RFX7 / Q2KHR2
RGCC / Q9H4X1
RGL1 / Q9NZL6
RGN / Q15493
RGS10 / O43665
RGS12 / O14924
RGS14 / O43566
RGS19 / P49795
RGS20 / O76081
RGS6 / P49758
RGS7 / P49802
RGS7BP / Q6MZT1
RHBDD1 / Q8TEB9
RHBDD2 / Q6NTF9
RHBDF1 / Q96CC6
RHEB / Q15382
RHOA / P61586
RHOB / P62745
RHOBTB3 / O94955
RHOC / P08134
RHOD / O00212
RHOF / Q9HBH0
RHOG / P84095
RHOQ / P17081
RHOT1 / Q8IXI2
RHOT2 / Q8IXI1
RHPN2 / Q8IUC4
RIC8A / Q9NPQ8
RICTOR / Q6R327
RIDA / P52758
RIF1 / Q5UIP0
Gene Symbol / UniProt Accession
Brain
Colon
Duodenum
Esophagus
Heart
Liver
Lung
Pancreas
Thyroid
Organs
Type
DDA
DIA
Bins
1
2
3
4
5

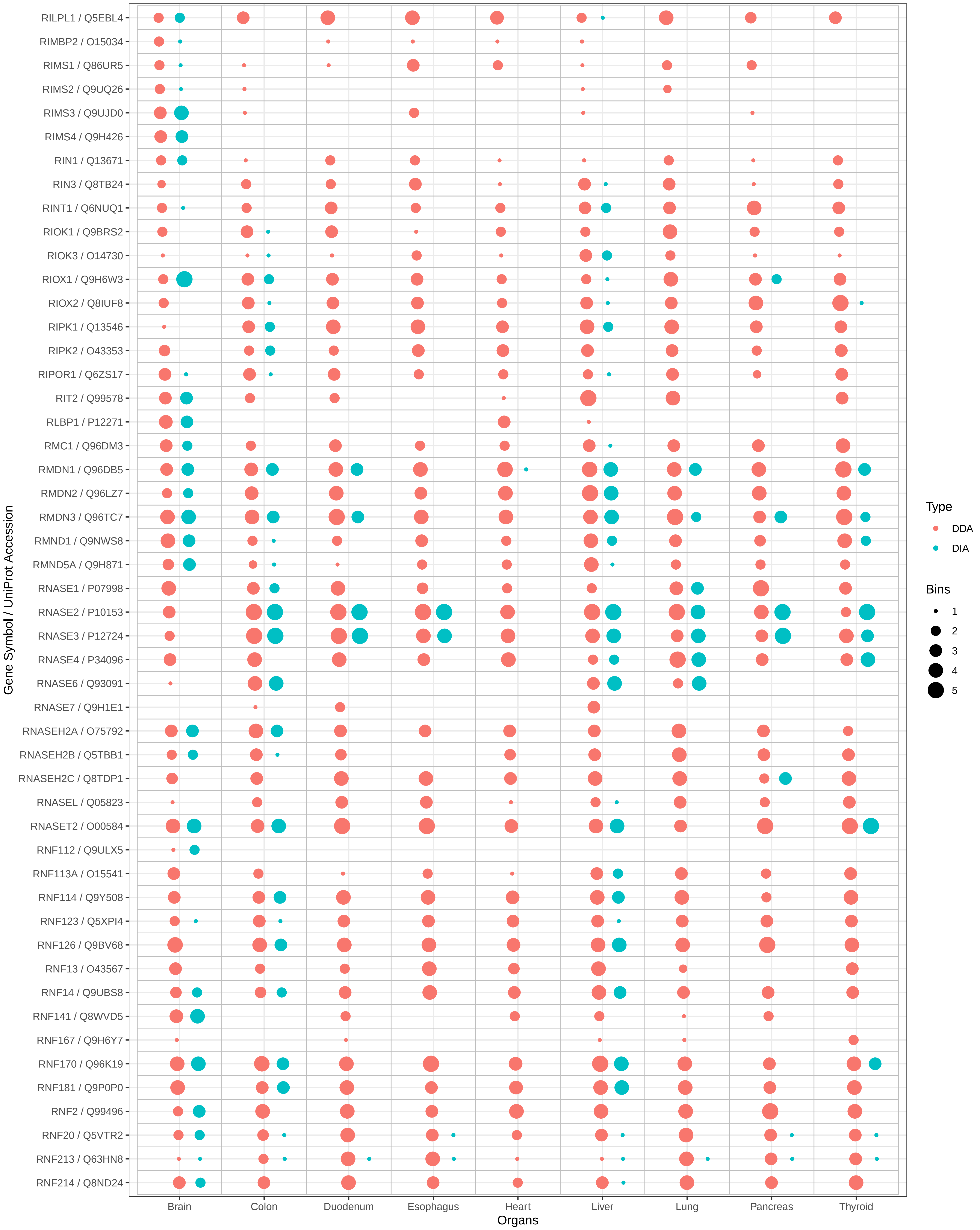

RILPL1 / Q5EBL4
RIMBP2 / O15034
RIMS1 / Q86UR5
RIMS2 / Q9UQ26
RIMS3 / Q9UJD0
RIMS4 / Q9H426
RIN1 / Q13671
RIN3 / Q8TB24
RINT1 / Q6NUQ1
RIOK1 / Q9BRS2
RIOK3 / O14730
RIOX1 / Q9H6W3
RIOX2 / Q8IUF8
RIPK1 / Q13546
RIPK2 / O43353
RIPOR1 / Q6ZS17
RIT2 / Q99578
RLBP1 / P12271
RMC1 / Q96DM3
RMDN1 / Q96DB5
RMDN2 / Q96LZ7
RMDN3 / Q96TC7
RMND1 / Q9NWS8
RMND5A / Q9H871
RNASE1 / P07998
RNASE2 / P10153
RNASE3 / P12724
RNASE4 / P34096
RNASE6 / Q93091
RNASE7 / Q9H1E1
RNASEH2A / O75792
RNASEH2B / Q5TBB1
RNASEH2C / Q8TDP1
RNASEL / Q05823
RNASET2 / O00584
RNF112 / Q9ULX5
RNF113A / O15541
RNF114 / Q9Y508
RNF123 / Q5XPI4
RNF126 / Q9BV68
RNF13 / O43567
RNF14 / Q9UBS8
RNF141 / Q8WVD5
RNF167 / Q9H6Y7
RNF170 / Q96K19
RNF181 / Q9P0P0
RNF2 / Q99496
RNF20 / Q5VTR2
RNF213 / Q63HN8
RNF214 / Q8ND24
Gene Symbol / UniProt Accession
Brain
Colon
Duodenum
Esophagus
Heart
Liver
Lung
Pancreas
Thyroid
Organs
Type
DDA
DIA
Bins
1
2
3
4
5

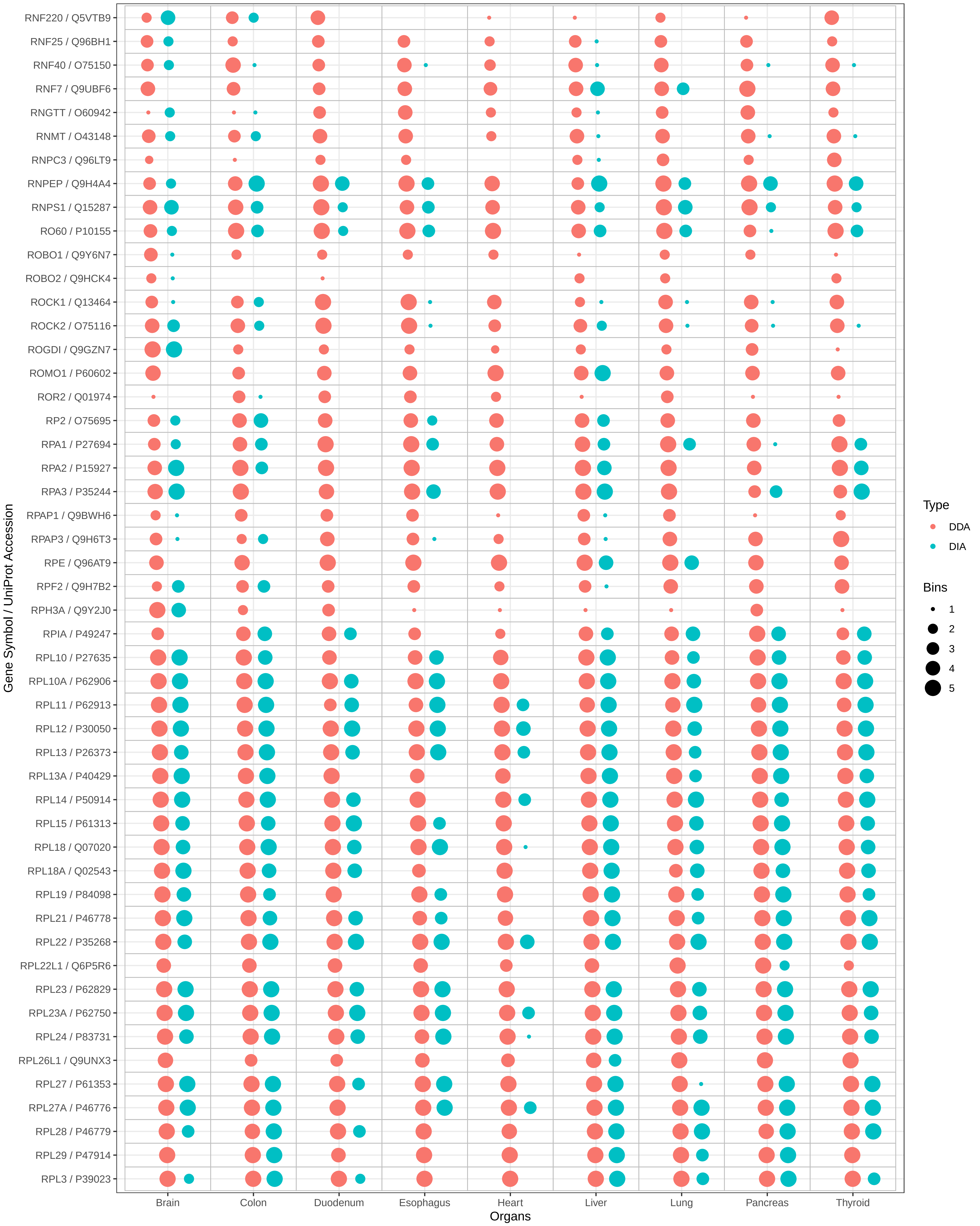
RNF220 / Q5VTB9
RNF25 / Q96BH1
RNF40 / O75150
RNF7 / Q9UBF6
RNGTT / O60942
RNMT / O43148
RNPC3 / Q96LT9
RNPEP / Q9H4A4
RNPS1 / Q15287
RO60 / P10155
ROBO1 / Q9Y6N7
ROBO2 / Q9HCK4
ROCK1 / Q13464
ROCK2 / O75116
ROGDI / Q9GZN7
ROMO1 / P60602
ROR2 / Q01974
RP2 / O75695
RPA1 / P27694
RPA2 / P15927
RPA3 / P35244
RPAP1 / Q9BWH6
RPAP3 / Q9H6T3
RPE / Q96AT9
RPF2 / Q9H7B2
RPH3A / Q9Y2J0
RPIA / P49247
RPL10 / P27635
RPL10A / P62906
RPL11 / P62913
RPL12 / P30050
RPL13 / P26373
RPL13A / P40429
RPL14 / P50914
RPL15 / P61313
RPL18 / Q07020
RPL18A / Q02543
RPL19 / P84098
RPL21 / P46778
RPL22 / P35268
RPL22L1 / Q6P5R6
RPL23 / P62829
RPL23A / P62750
RPL24 / P83731
RPL26L1 / Q9UNX3
RPL27 / P61353
RPL27A / P46776
RPL28 / P46779
RPL29 / P47914
RPL3 / P39023
Gene Symbol / UniProt Accession
Brain
Colon
Duodenum
Esophagus
Heart
Liver
Lung
Pancreas
Thyroid
Organs
Type
DDA
DIA
Bins
1
2
3
4
5

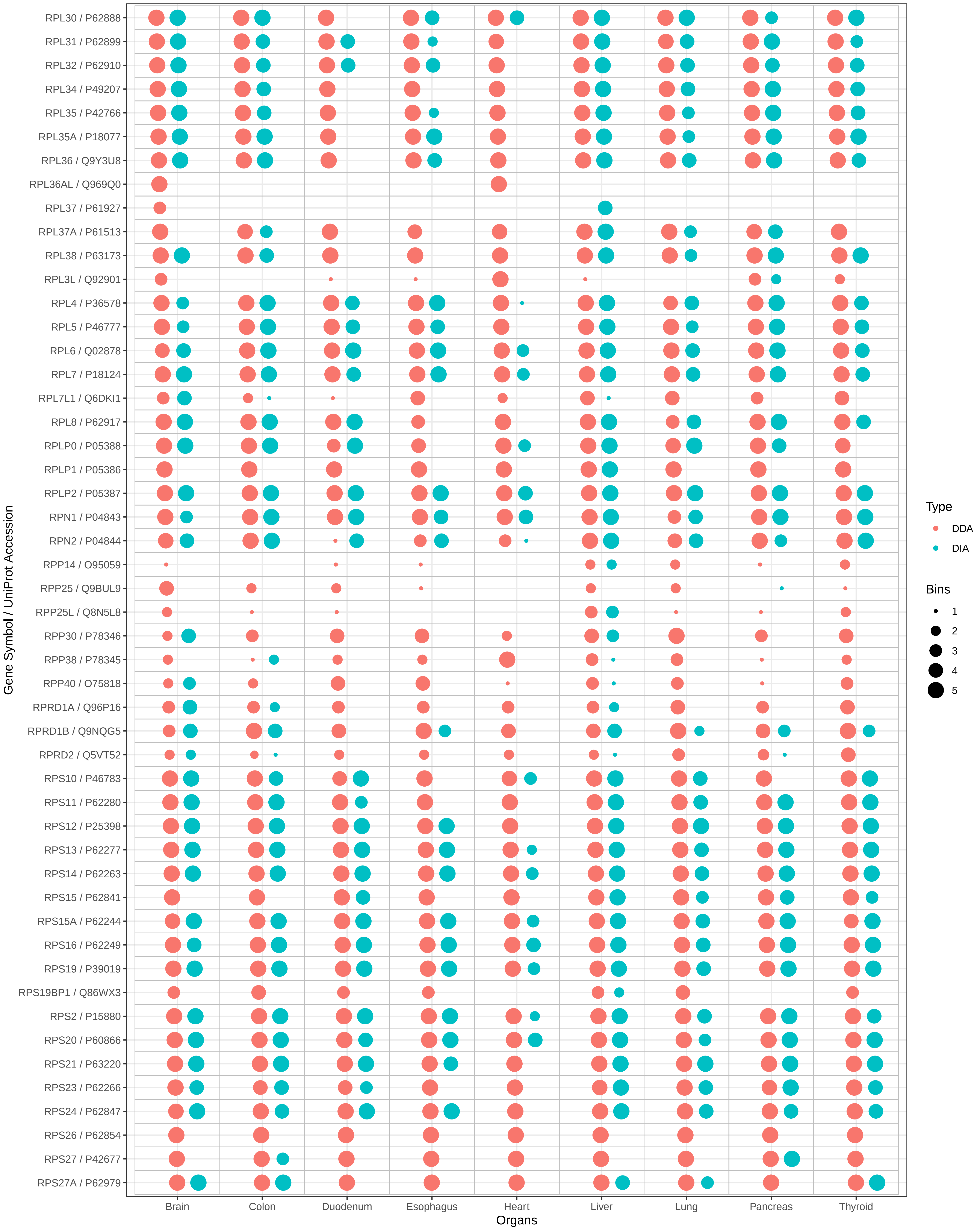

RPL30 / P62888
RPL31 / P62899
RPL32 / P62910
RPL34 / P49207
RPL35 / P42766
RPL35A / P18077
RPL36 / Q9Y3U8
RPL36AL / Q969Q0
RPL37 / P61927
RPL37A / P61513
RPL38 / P63173
RPL3L / Q92901
RPL4 / P36578
RPL5 / P46777
RPL6 / Q02878
RPL7 / P18124
RPL7L1 / Q6DKI1
RPL8 / P62917
RPLP0 / P05388
RPLP1 / P05386
RPLP2 / P05387
RPN1 / P04843
RPN2 / P04844
RPP14 / O95059
RPP25 / Q9BUL9
RPP25L / Q8N5L8
RPP30 / P78346
RPP38 / P78345
RPP40 / O75818
RPRD1A / Q96P16
RPRD1B / Q9NQG5
RPRD2 / Q5VT52
RPS10 / P46783
RPS11 / P62280
RPS12 / P25398
RPS13 / P62277
RPS14 / P62263
RPS15 / P62841
RPS15A / P62244
RPS16 / P62249
RPS19 / P39019
RPS19BP1 / Q86WX3
RPS2 / P15880
RPS20 / P60866
RPS21 / P63220
RPS23 / P62266
RPS24 / P62847
RPS26 / P62854
RPS27 / P42677
RPS27A / P62979
Gene Symbol / UniProt Accession
Brain
Colon
Duodenum
Esophagus
Heart
Liver
Lung
Pancreas
Thyroid
Organs
Type
DDA
DIA
Bins
1
2
3
4
5

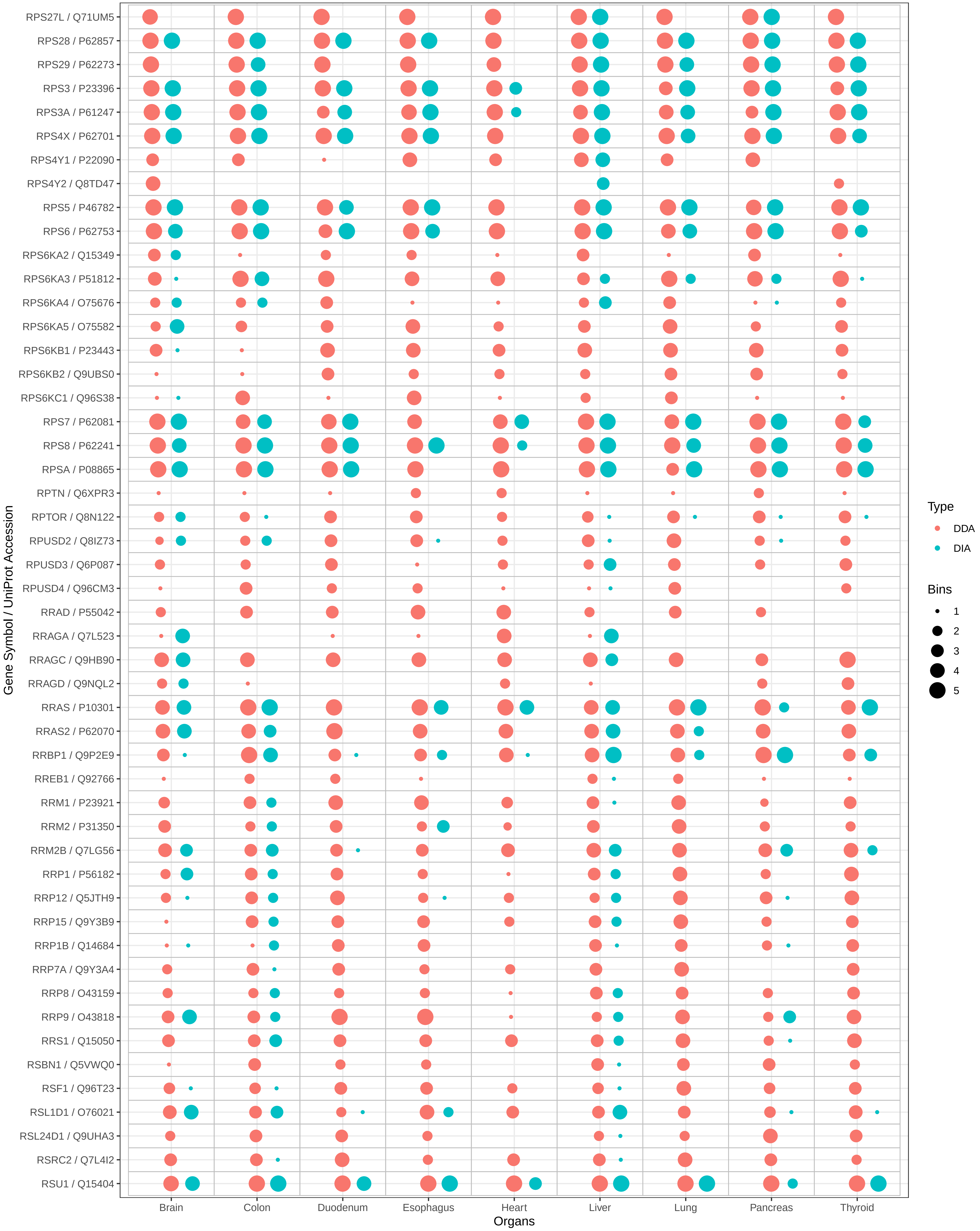

RPS27L / Q71UM5
RPS28 / P62857
RPS29 / P62273
RPS3 / P23396
RPS3A / P61247
RPS4X / P62701
RPS4Y1 / P22090
RPS4Y2 / Q8TD47
RPS5 / P46782
RPS6 / P62753
RPS6KA2 / Q15349
RPS6KA3 / P51812
RPS6KA4 / O75676
RPS6KA5 / O75582
RPS6KB1 / P23443
RPS6KB2 / Q9UBS0
RPS6KC1 / Q96S38
RPS7 / P62081
RPS8 / P62241
RPSA / P08865
RPTN / Q6XPR3
RPTOR / Q8N122
RPUSD2 / Q8IZ73
RPUSD3 / Q6P087
RPUSD4 / Q96CM3
RRAD / P55042
RRAGA / Q7L523
RRAGC / Q9HB90
RRAGD / Q9NQL2
RRAS / P10301
RRAS2 / P62070
RRBP1 / Q9P2E9
RREB1 / Q92766
RRM1 / P23921
RRM2 / P31350
RRM2B / Q7LG56
RRP1 / P56182
RRP12 / Q5JTH9
RRP15 / Q9Y3B9
RRP1B / Q14684
RRP7A / Q9Y3A4
RRP8 / O43159
RRP9 / O43818
RRS1 / Q15050
RSBN1 / Q5VWQ0
RSF1 / Q96T23
RSL1D1 / O76021
RSL24D1 / Q9UHA3
RSRC2 / Q7L4I2
RSU1 / Q15404
Gene Symbol / UniProt Accession
Brain
Colon
Duodenum
Esophagus
Heart
Liver
Lung
Pancreas
Thyroid
Organs
Type
DDA
DIA
Bins
1
2
3
4
5

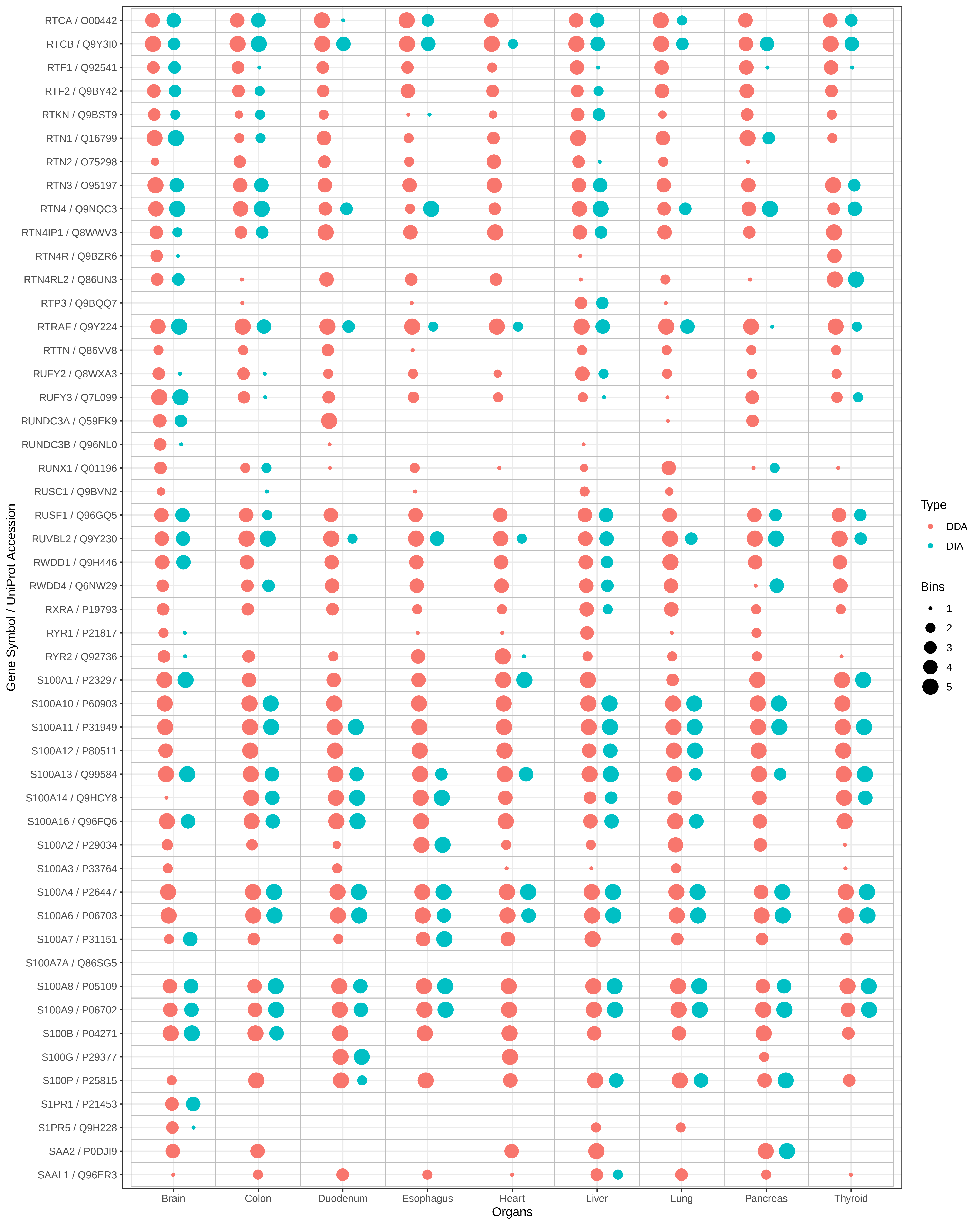
RTCA / O00442
RTCB / Q9Y3I0
RTF1 / Q92541
RTF2 / Q9BY42
RTKN / Q9BST9
RTN1 / Q16799
RTN2 / O75298
RTN3 / O95197
RTN4 / Q9NQC3
RTN4IP1 / Q8WWV3
RTN4R / Q9BZR6
RTN4RL2 / Q86UN3
RTP3 / Q9BQQ7
RTRAF / Q9Y224
RTTN / Q86VV8
RUFY2 / Q8WXA3
RUFY3 / Q7L099
RUNDC3A / Q59EK9
RUNDC3B / Q96NL0
RUNX1 / Q01196
RUSC1 / Q9BVN2
RUSF1 / Q96GQ5
RUVBL2 / Q9Y230
RWDD1 / Q9H446
RWDD4 / Q6NW29
RXRA / P19793
RYR1 / P21817
RYR2 / Q92736
S100A1 / P23297
S100A10 / P60903
S100A11 / P31949
S100A12 / P80511
S100A13 / Q99584
S100A14 / Q9HCY8
S100A16 / Q96FQ6
S100A2 / P29034
S100A3 / P33764
S100A4 / P26447
S100A6 / P06703
S100A7 / P31151
S100A7A / Q86SG5
S100A8 / P05109
S100A9 / P06702
S100B / P04271
S100G / P29377
S100P / P25815
S1PR1 / P21453
S1PR5 / Q9H228
SAA2 / P0DJI9
SAAL1 / Q96ER3
Gene Symbol / UniProt Accession
Brain
Colon
Duodenum
Esophagus
Heart
Liver
Lung
Pancreas
Thyroid
Organs
Type
DDA
DIA
Bins
1
2
3
4
5

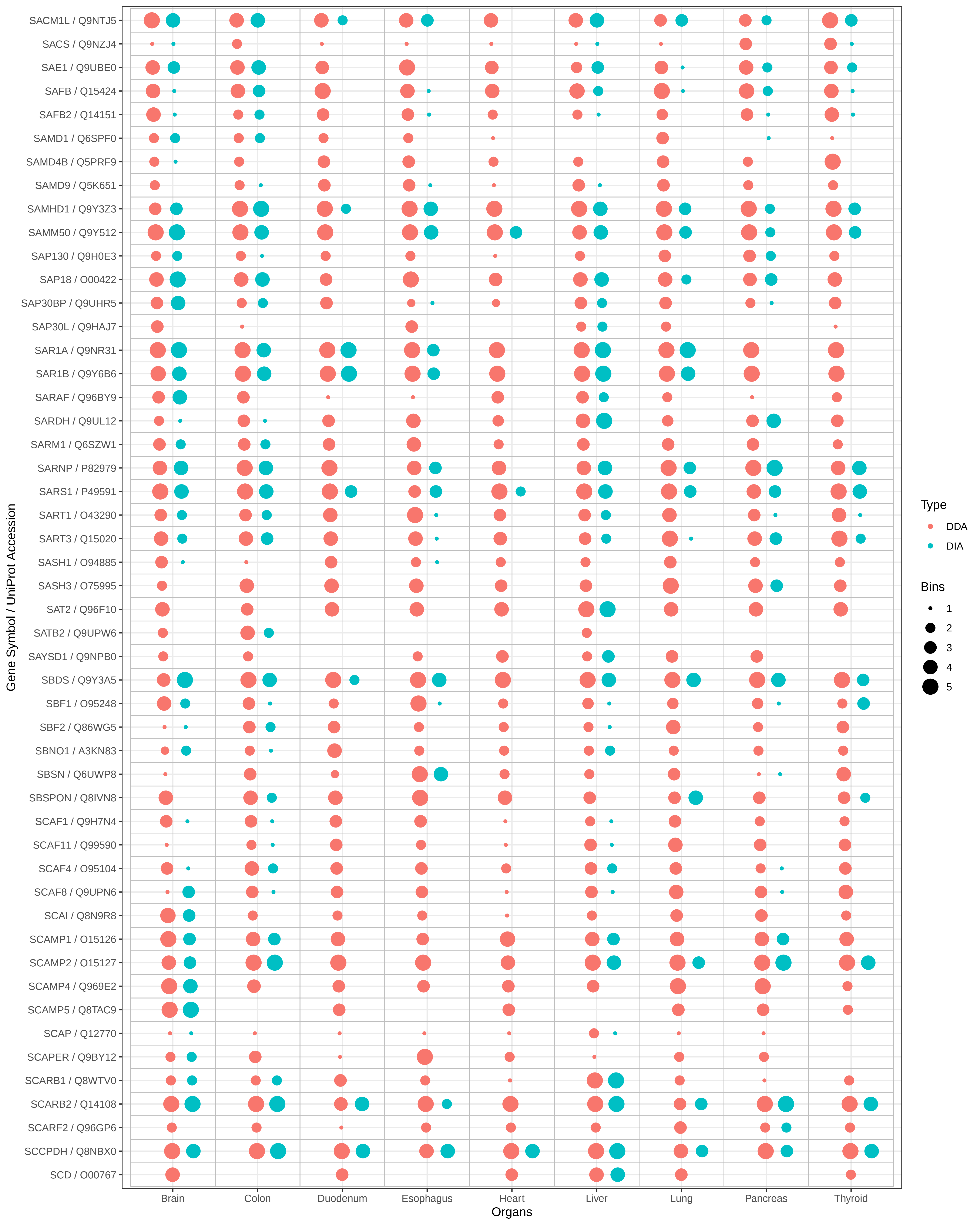
SACM1L / Q9NTJ5
SACS / Q9NZJ4
SAE1 / Q9UBE0
SAFB / Q15424
SAFB2 / Q14151
SAMD1 / Q6SPF0
SAMD4B / Q5PRF9
SAMD9 / Q5K651
SAMHD1 / Q9Y3Z3
SAMM50 / Q9Y512
SAP130 / Q9H0E3
SAP18 / O00422
SAP30BP / Q9UHR5
SAP30L / Q9HAJ7
SAR1A / Q9NR31
SAR1B / Q9Y6B6
SARAF / Q96BY9
SARDH / Q9UL12
SARM1 / Q6SZW1
SARNP / P82979
SARS1 / P49591
SART1 / O43290
SART3 / Q15020
SASH1 / O94885
SASH3 / O75995
SAT2 / Q96F10
SATB2 / Q9UPW6
SAYSD1 / Q9NPB0
SBDS / Q9Y3A5
SBF1 / O95248
SBF2 / Q86WG5
SBNO1 / A3KN83
SBSN / Q6UWP8
SBSPON / Q8IVN8
SCAF1 / Q9H7N4
SCAF11 / Q99590
SCAF4 / O95104
SCAF8 / Q9UPN6
SCAI / Q8N9R8
SCAMP1 / O15126
SCAMP2 / O15127
SCAMP4 / Q969E2
SCAMP5 / Q8TAC9
SCAP / Q12770
SCAPER / Q9BY12
SCARB1 / Q8WTV0
SCARB2 / Q14108
SCARF2 / Q96GP6
SCCPDH / Q8NBX0
SCD / O00767
Gene Symbol / UniProt Accession
Brain
Colon
Duodenum
Esophagus
Heart
Liver
Lung
Pancreas
Thyroid
Organs
Type
DDA
DIA
Bins
1
2
3
4
5

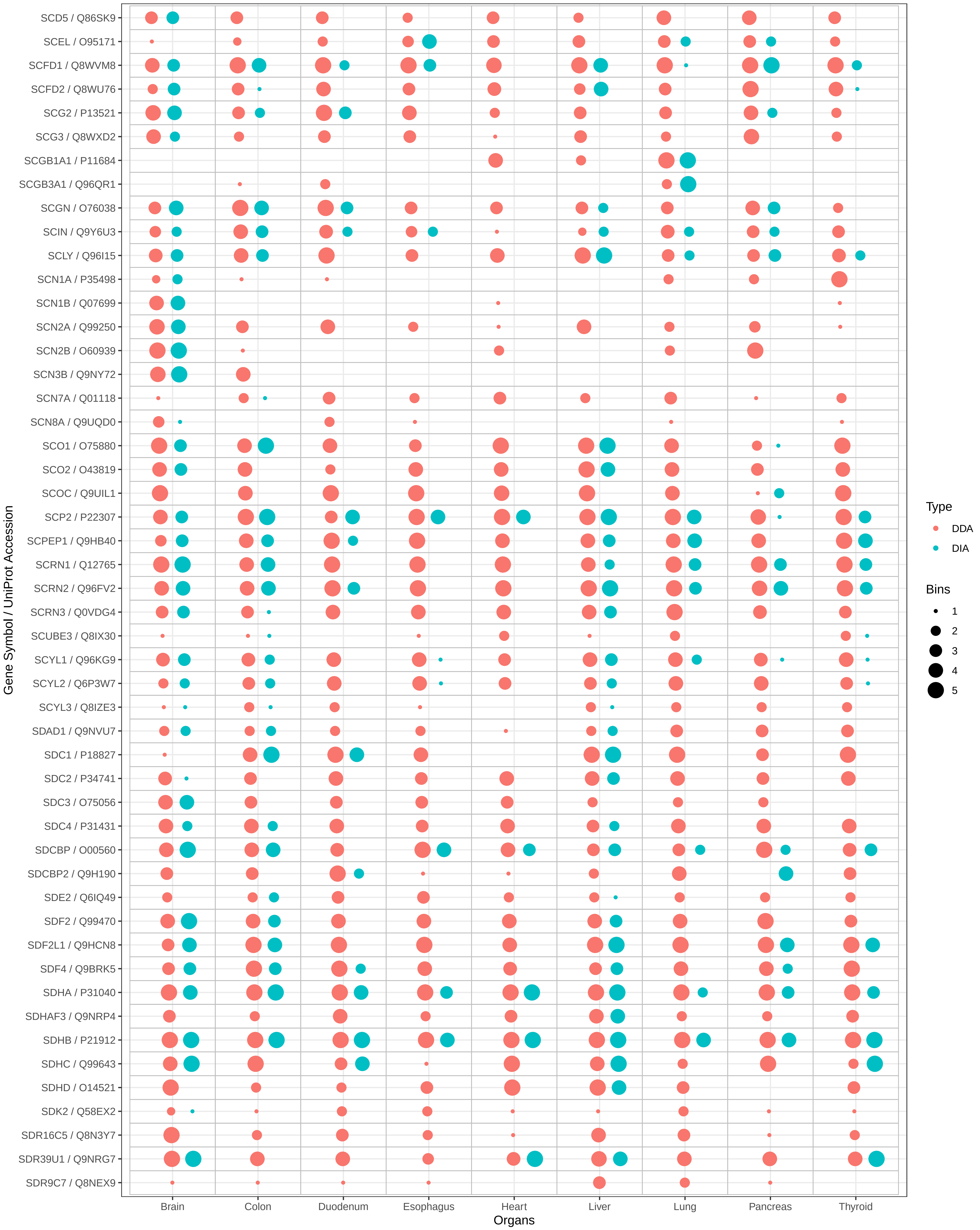

SCD5 / Q86SK9
SCEL / O95171
SCFD1 / Q8WVM8
SCFD2 / Q8WU76
SCG2 / P13521
SCG3 / Q8WXD2
SCGB1A1 / P11684
SCGB3A1 / Q96QR1
SCGN / O76038
SCIN / Q9Y6U3
SCLY / Q96I15
SCN1A / P35498
SCN1B / Q07699
SCN2A / Q99250
SCN2B / O60939
SCN3B / Q9NY72
SCN7A / Q01118
SCN8A / Q9UQD0
SCO1 / O75880
SCO2 / O43819
SCOC / Q9UIL1
SCP2 / P22307
SCPEP1 / Q9HB40
SCRN1 / Q12765
SCRN2 / Q96FV2
SCRN3 / Q0VDG4
SCUBE3 / Q8IX30
SCYL1 / Q96KG9
SCYL2 / Q6P3W7
SCYL3 / Q8IZE3
SDAD1 / Q9NVU7
SDC1 / P18827
SDC2 / P34741
SDC3 / O75056
SDC4 / P31431
SDCBP / O00560
SDCBP2 / Q9H190
SDE2 / Q6IQ49
SDF2 / Q99470
SDF2L1 / Q9HCN8
SDF4 / Q9BRK5
SDHA / P31040
SDHAF3 / Q9NRP4
SDHB / P21912
SDHC / Q99643
SDHD / O14521
SDK2 / Q58EX2
SDR16C5 / Q8N3Y7
SDR39U1 / Q9NRG7
SDR9C7 / Q8NEX9
Gene Symbol / UniProt Accession
Brain
Colon
Duodenum
Esophagus
Heart
Liver
Lung
Pancreas
Thyroid
Organs
Type
DDA
DIA
Bins
1
2
3
4
5

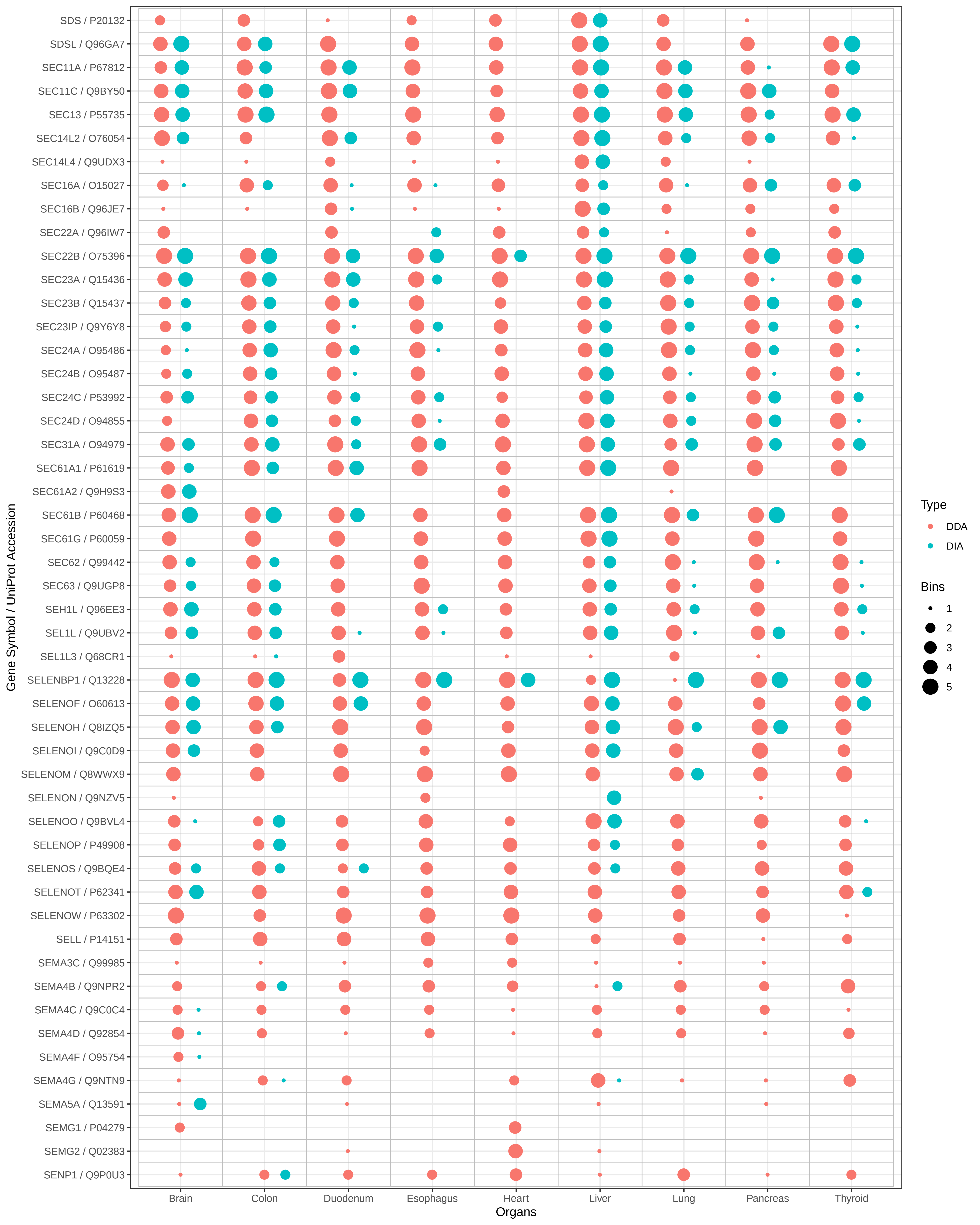
SDS / P20132
SDSL / Q96GA7
SEC11A / P67812
SEC11C / Q9BY50
SEC13 / P55735
SEC14L2 / O76054
SEC14L4 / Q9UDX3
SEC16A / O15027
SEC16B / Q96JE7
SEC22A / Q96IW7
SEC22B / O75396
SEC23A / Q15436
SEC23B / Q15437
SEC23IP / Q9Y6Y8
SEC24A / O95486
SEC24B / O95487
SEC24C / P53992
SEC24D / O94855
SEC31A / O94979
SEC61A1 / P61619
SEC61A2 / Q9H9S3
SEC61B / P60468
SEC61G / P60059
SEC62 / Q99442
SEC63 / Q9UGP8
SEH1L / Q96EE3
SEL1L / Q9UBV2
SEL1L3 / Q68CR1
SELENBP1 / Q13228
SELENOF / O60613
SELENOH / Q8IZQ5
SELENOI / Q9C0D9
SELENOM / Q8WWX9
SELENON / Q9NZV5
SELENOO / Q9BVL4
SELENOP / P49908
SELENOS / Q9BQE4
SELENOT / P62341
SELENOW / P63302
SELL / P14151
SEMA3C / Q99985
SEMA4B / Q9NPR2
SEMA4C / Q9C0C4
SEMA4D / Q92854
SEMA4F / O95754
SEMA4G / Q9NTN9
SEMA5A / Q13591
SEMG1 / P04279
SEMG2 / Q02383
SENP1 / Q9P0U3
Gene Symbol / UniProt Accession
Brain
Colon
Duodenum
Esophagus
Heart
Liver
Lung
Pancreas
Thyroid
Organs
Type
DDA
DIA
Bins
1
2
3
4
5

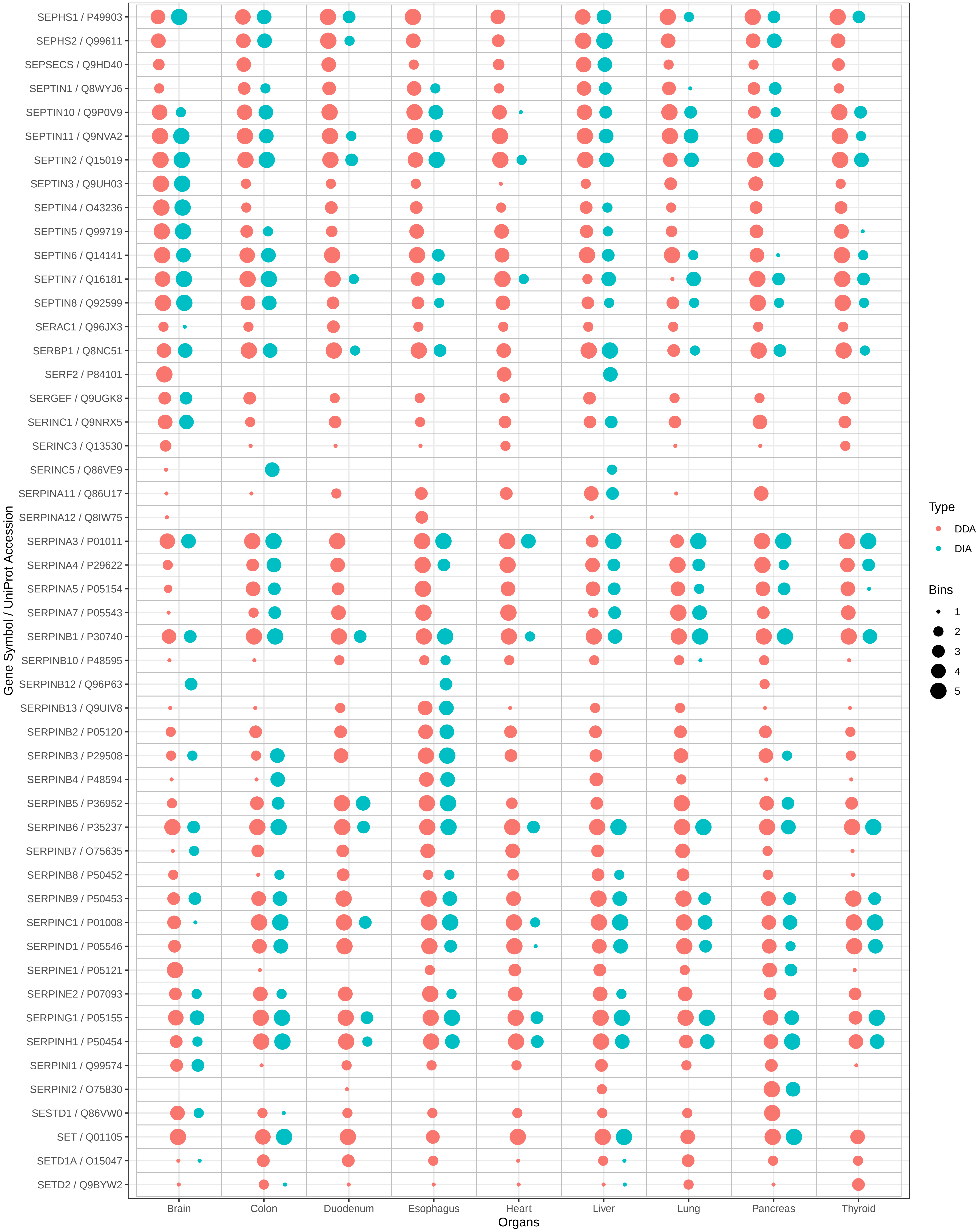

SEPHS1 / P49903
SEPHS2 / Q99611
SEPSECS / Q9HD40
SEPTIN1 / Q8WYJ6
SEPTIN10 / Q9P0V9
SEPTIN11 / Q9NVA2
SEPTIN2 / Q15019
SEPTIN3 / Q9UH03
SEPTIN4 / O43236
SEPTIN5 / Q99719
SEPTIN6 / Q14141
SEPTIN7 / Q16181
SEPTIN8 / Q92599
SERAC1 / Q96JX3
SERBP1 / Q8NC51
SERF2 / P84101
SERGEF / Q9UGK8
SERINC1 / Q9NRX5
SERINC3 / Q13530
SERINC5 / Q86VE9
SERPINA11 / Q86U17
SERPINA12 / Q8IW75
SERPINA3 / P01011
SERPINA4 / P29622
SERPINA5 / P05154
SERPINA7 / P05543
SERPINB1 / P30740
SERPINB10 / P48595
SERPINB12 / Q96P63
SERPINB13 / Q9UIV8
SERPINB2 / P05120
SERPINB3 / P29508
SERPINB4 / P48594
SERPINB5 / P36952
SERPINB6 / P35237
SERPINB7 / O75635
SERPINB8 / P50452
SERPINB9 / P50453
SERPINC1 / P01008
SERPIND1 / P05546
SERPINE1 / P05121
SERPINE2 / P07093
SERPING1 / P05155
SERPINH1 / P50454
SERPINI1 / Q99574
SERPINI2 / O75830
SESTD1 / Q86VW0
SET / Q01105
SETD1A / O15047
SETD2 / Q9BYW2
Gene Symbol / UniProt Accession
Brain
Colon
Duodenum
Esophagus
Heart
Liver
Lung
Pancreas
Thyroid
Organs
Type
DDA
DIA
Bins
1
2
3
4
5

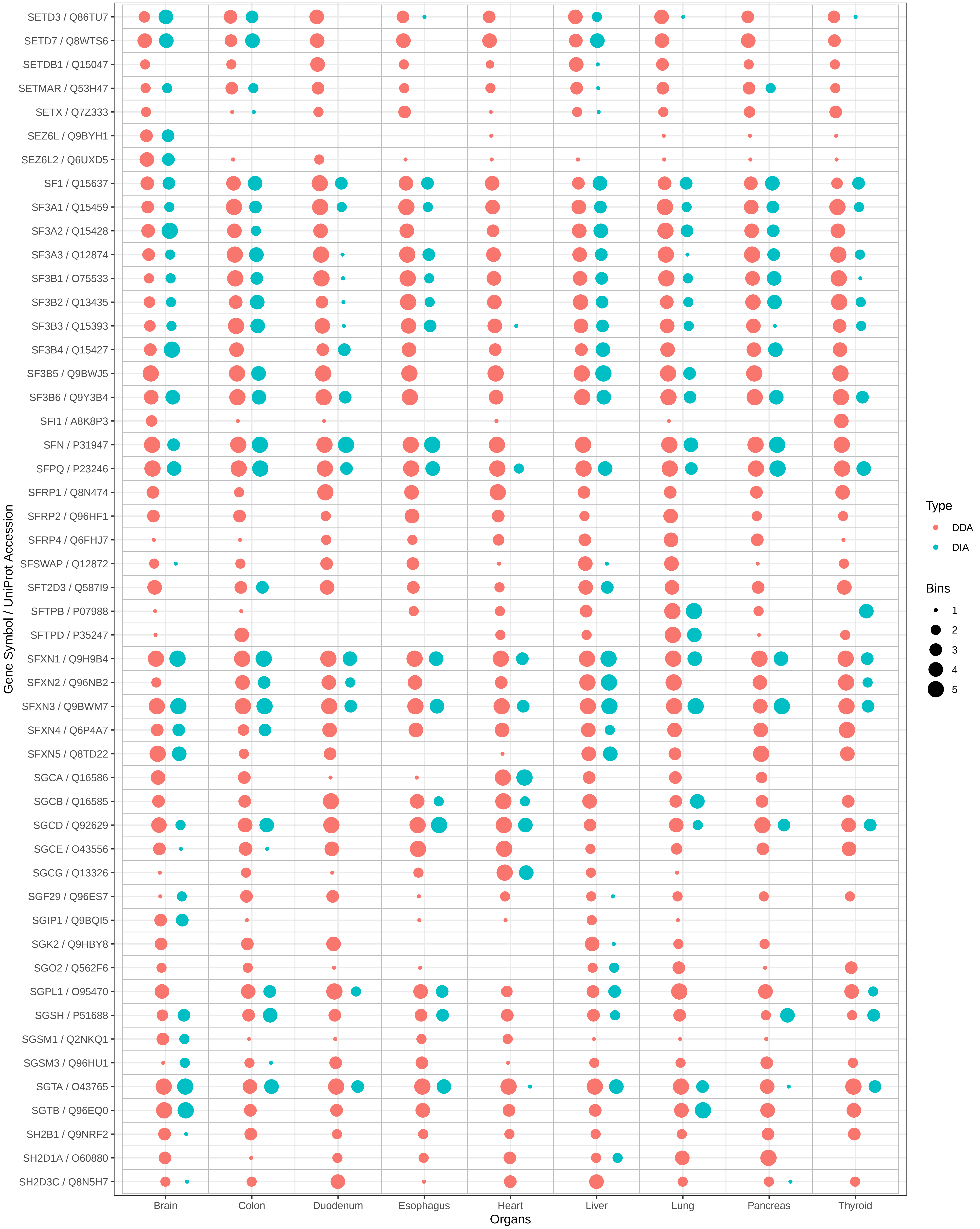
SETD3 / Q86TU7
SETD7 / Q8WTS6
SETDB1 / Q15047
SETMAR / Q53H47
SETX / Q7Z333
SEZ6L / Q9BYH1
SEZ6L2 / Q6UXD5
SF1 / Q15637
SF3A1 / Q15459
SF3A2 / Q15428
SF3A3 / Q12874
SF3B1 / O75533
SF3B2 / Q13435
SF3B3 / Q15393
SF3B4 / Q15427
SF3B5 / Q9BWJ5
SF3B6 / Q9Y3B4
SFI1 / A8K8P3
SFN / P31947
SFPQ / P23246
SFRP1 / Q8N474
SFRP2 / Q96HF1
SFRP4 / Q6FHJ7
SFSWAP / Q12872
SFT2D3 / Q587I9
SFTPB / P07988
SFTPD / P35247
SFXN1 / Q9H9B4
SFXN2 / Q96NB2
SFXN3 / Q9BWM7
SFXN4 / Q6P4A7
SFXN5 / Q8TD22
SGCA / Q16586
SGCB / Q16585
SGCD / Q92629
SGCE / O43556
SGCG / Q13326
SGF29 / Q96ES7
SGIP1 / Q9BQI5
SGK2 / Q9HBY8
SGO2 / Q562F6
SGPL1 / O95470
SGSH / P51688
SGSM1 / Q2NKQ1
SGSM3 / Q96HU1
SGTA / O43765
SGTB / Q96EQ0
SH2B1 / Q9NRF2
SH2D1A / O60880
SH2D3C / Q8N5H7
Gene Symbol / UniProt Accession
Brain
Colon
Duodenum
Esophagus
Heart
Liver
Lung
Pancreas
Thyroid
Organs
Type
DDA
DIA
Bins
1
2
3
4
5

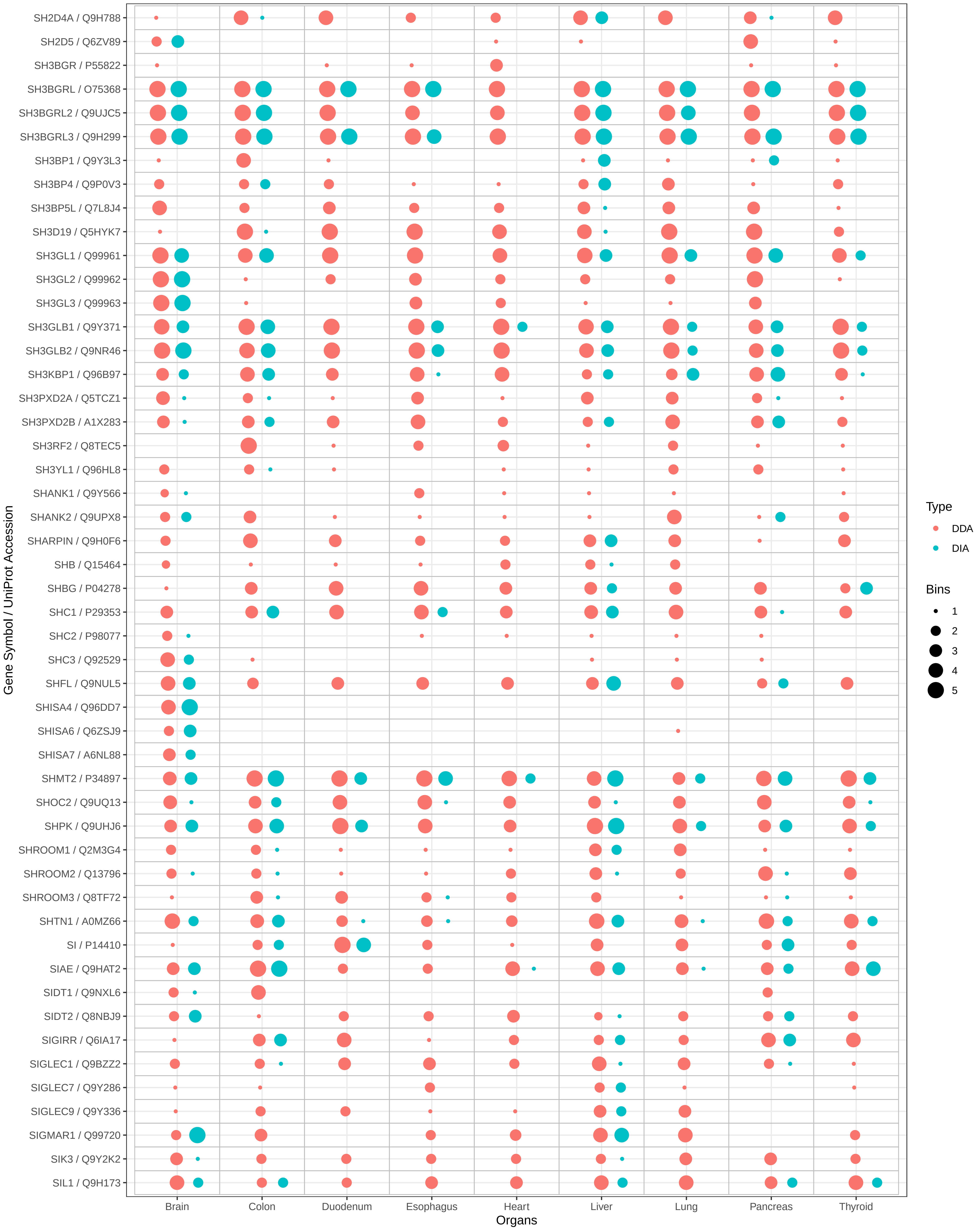

SH2D4A / Q9H788
SH2D5 / Q6ZV89
SH3BGR / P55822
SH3BGRL / O75368
SH3BGRL2 / Q9UJC5
SH3BGRL3 / Q9H299
SH3BP1 / Q9Y3L3
SH3BP4 / Q9P0V3
SH3BP5L / Q7L8J4
SH3D19 / Q5HYK7
SH3GL1 / Q99961
SH3GL2 / Q99962
SH3GL3 / Q99963
SH3GLB1 / Q9Y371
SH3GLB2 / Q9NR46
SH3KBP1 / Q96B97
SH3PXD2A / Q5TCZ1
SH3PXD2B / A1X283
SH3RF2 / Q8TEC5
SH3YL1 / Q96HL8
SHANK1 / Q9Y566
SHANK2 / Q9UPX8
SHARPIN / Q9H0F6
SHB / Q15464
SHBG / P04278
SHC1 / P29353
SHC2 / P98077
SHC3 / Q92529
SHFL / Q9NUL5
SHISA4 / Q96DD7
SHISA6 / Q6ZSJ9
SHISA7 / A6NL88
SHMT2 / P34897
SHOC2 / Q9UQ13
SHPK / Q9UHJ6
SHROOM1 / Q2M3G4
SHROOM2 / Q13796
SHROOM3 / Q8TF72
SHTN1 / A0MZ66
SI / P14410
SIAE / Q9HAT2
SIDT1 / Q9NXL6
SIDT2 / Q8NBJ9
SIGIRR / Q6IA17
SIGLEC1 / Q9BZZ2
SIGLEC7 / Q9Y286
SIGLEC9 / Q9Y336
SIGMAR1 / Q99720
SIK3 / Q9Y2K2
SIL1 / Q9H173
Gene Symbol / UniProt Accession
Brain
Colon
Duodenum
Esophagus
Heart
Liver
Lung
Pancreas
Thyroid
Organs
Type
DDA
DIA
Bins
1
2
3
4
5

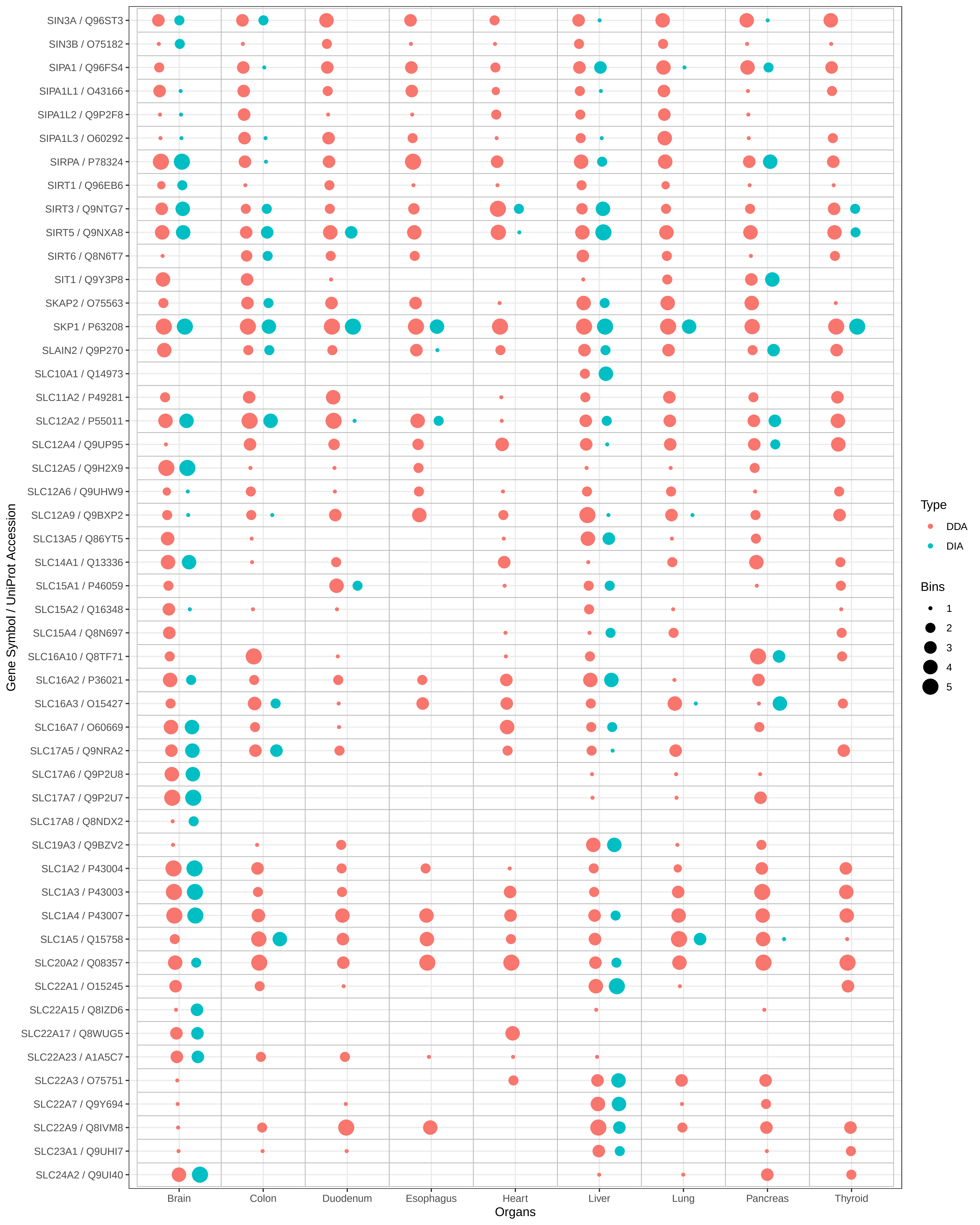

SIN3A / Q96ST3
SIN3B / O75182
SIPA1 / Q96FS4
SIPA1L1 / O43166
SIPA1L2 / Q9P2F8
SIPA1L3 / O60292
SIRPA / P78324
SIRT1 / Q96EB6
SIRT3 / Q9NTG7
SIRT5 / Q9NXA8
SIRT6 / Q8N6T7
SIT1 / Q9Y3P8
SKAP2 / O75563
SKP1 / P63208
SLAIN2 / Q9P270
SLC10A1 / Q14973
SLC11A2 / P49281
SLC12A2 / P55011
SLC12A4 / Q9UP95
SLC12A5 / Q9H2X9
SLC12A6 / Q9UHW9
SLC12A9 / Q9BXP2
SLC13A5 / Q86YT5
SLC14A1 / Q13336
SLC15A1 / P46059
SLC15A2 / Q16348
SLC15A4 / Q8N697
SLC16A10 / Q8TF71
SLC16A2 / P36021
SLC16A3 / O15427
SLC16A7 / O60669
SLC17A5 / Q9NRA2
SLC17A6 / Q9P2U8
SLC17A7 / Q9P2U7
SLC17A8 / Q8NDX2
SLC19A3 / Q9BZV2
SLC1A2 / P43004
SLC1A3 / P43003
SLC1A4 / P43007
SLC1A5 / Q15758
SLC20A2 / Q08357
SLC22A1 / O15245
SLC22A15 / Q8IZD6
SLC22A17 / Q8WUG5
SLC22A23 / A1A5C7
SLC22A3 / O75751
SLC22A7 / Q9Y694
SLC22A9 / Q8IVM8
SLC23A1 / Q9UHI7
SLC24A2 / Q9UI40
Gene Symbol / UniProt Accession
Brain
Colon
Duodenum
Esophagus
Heart
Liver
Lung
Pancreas
Thyroid
Organs
Type
DDA
DIA
Bins
1
2
3
4
5

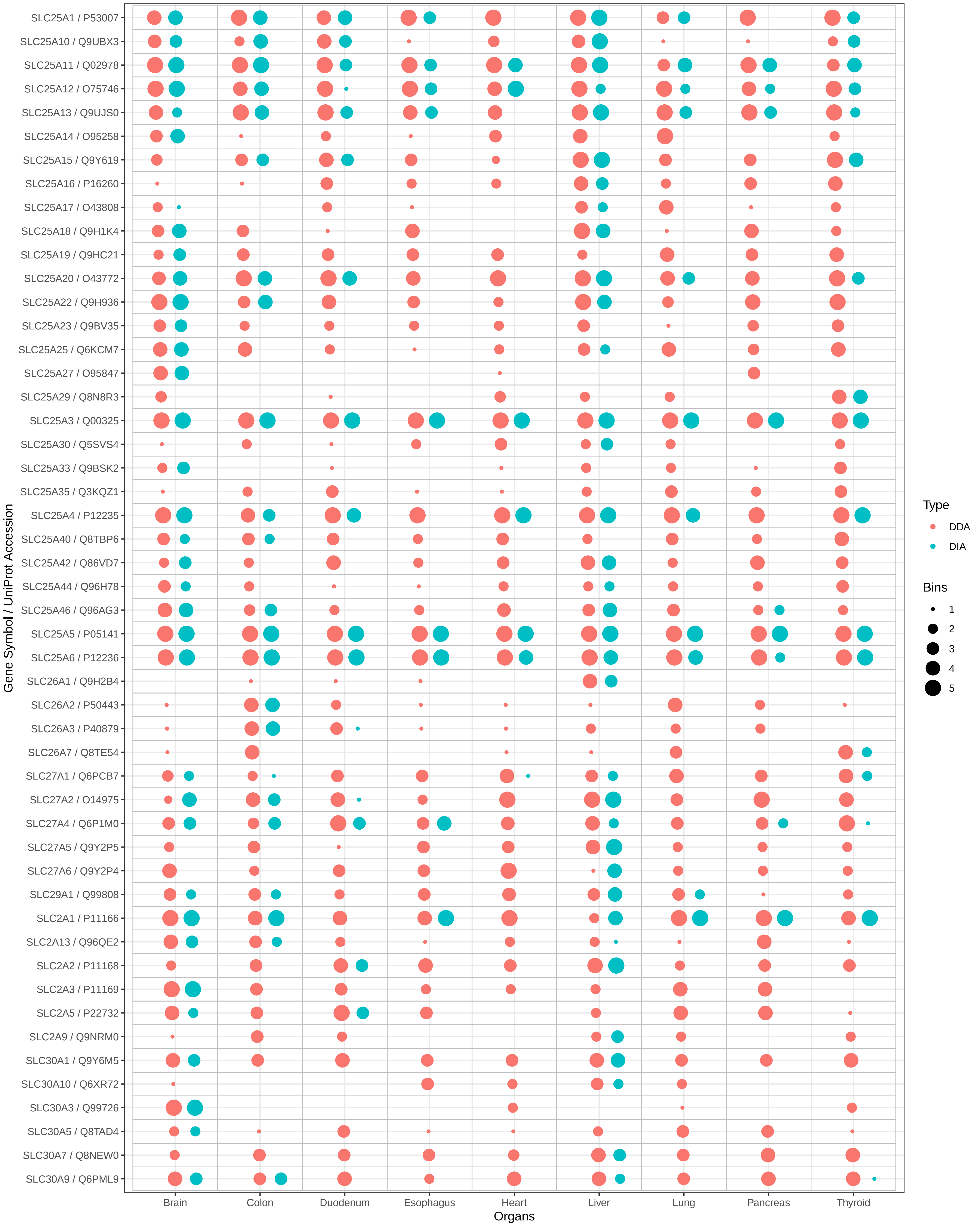
SLC25A1 / P53007
SLC25A10 / Q9UBX3
SLC25A11 / Q02978
SLC25A12 / O75746
SLC25A13 / Q9UJS0
SLC25A14 / O95258
SLC25A15 / Q9Y619
SLC25A16 / P16260
SLC25A17 / O43808
SLC25A18 / Q9H1K4
SLC25A19 / Q9HC21
SLC25A20 / O43772
SLC25A22 / Q9H936
SLC25A23 / Q9BV35
SLC25A25 / Q6KCM7
SLC25A27 / O95847
SLC25A29 / Q8N8R3
SLC25A3 / Q00325
SLC25A30 / Q5SVS4
SLC25A33 / Q9BSK2
SLC25A35 / Q3KQZ1
SLC25A4 / P12235
SLC25A40 / Q8TBP6
SLC25A42 / Q86VD7
SLC25A44 / Q96H78
SLC25A46 / Q96AG3
SLC25A5 / P05141
SLC25A6 / P12236
SLC26A1 / Q9H2B4
SLC26A2 / P50443
SLC26A3 / P40879
SLC26A7 / Q8TE54
SLC27A1 / Q6PCB7
SLC27A2 / O14975
SLC27A4 / Q6P1M0
SLC27A5 / Q9Y2P5
SLC27A6 / Q9Y2P4
SLC29A1 / Q99808
SLC2A1 / P11166
SLC2A13 / Q96QE2
SLC2A2 / P11168
SLC2A3 / P11169
SLC2A5 / P22732
SLC2A9 / Q9NRM0
SLC30A1 / Q9Y6M5
SLC30A10 / Q6XR72
SLC30A3 / Q99726
SLC30A5 / Q8TAD4
SLC30A7 / Q8NEW0
SLC30A9 / Q6PML9
Gene Symbol / UniProt Accession
Brain
Colon
Duodenum
Esophagus
Heart
Liver
Lung
Pancreas
Thyroid
Organs
Type
DDA
DIA
Bins
1
2
3
4
5

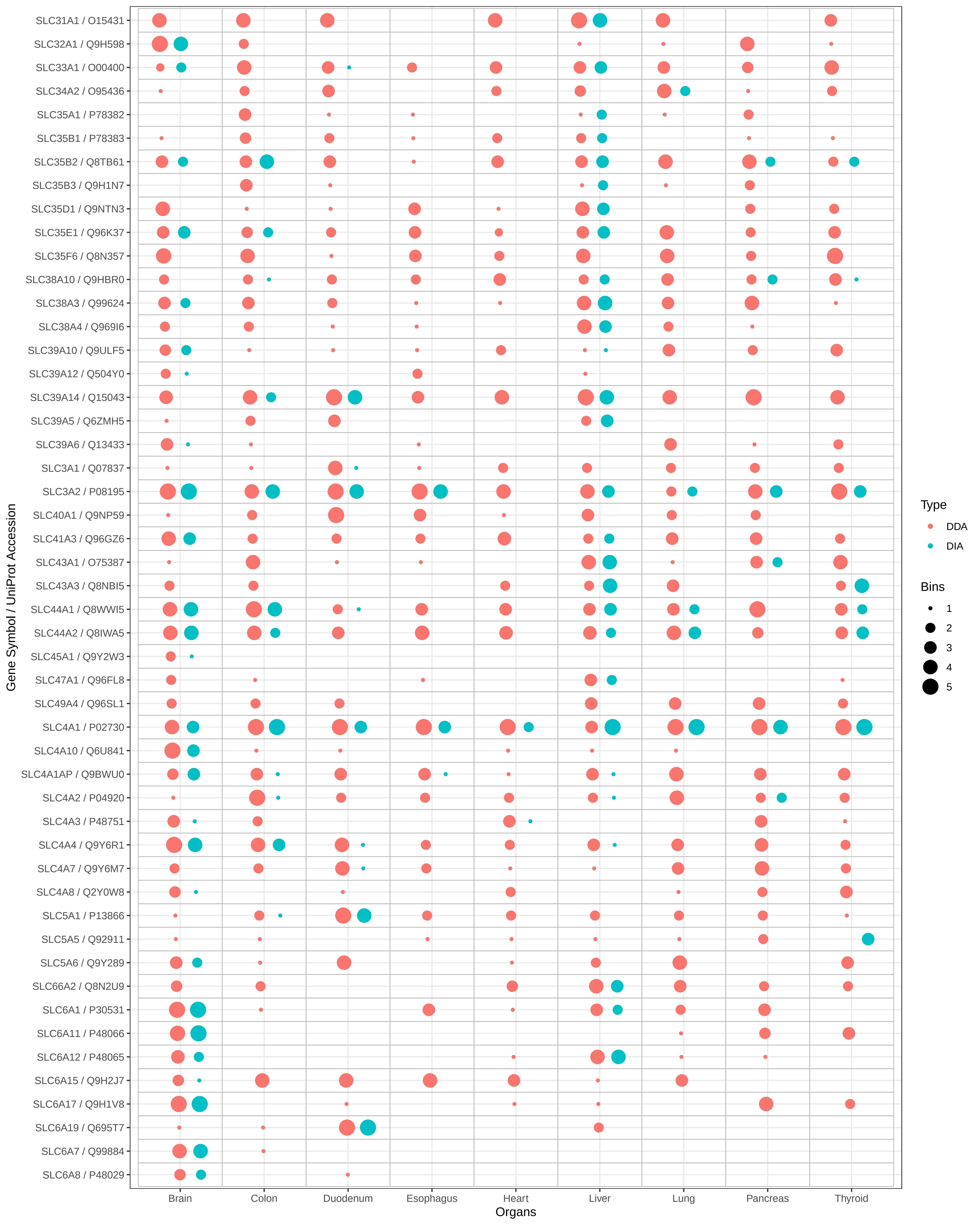

SLC31A1 / O15431
SLC32A1 / Q9H598
SLC33A1 / O00400
SLC34A2 / O95436
SLC35A1 / P78382
SLC35B1 / P78383
SLC35B2 / Q8TB61
SLC35B3 / Q9H1N7
SLC35D1 / Q9NTN3
SLC35E1 / Q96K37
SLC35F6 / Q8N357
SLC38A10 / Q9HBR0
SLC38A3 / Q99624
SLC38A4 / Q969I6
SLC39A10 / Q9ULF5
SLC39A12 / Q504Y0
SLC39A14 / Q15043
SLC39A5 / Q6ZMH5
SLC39A6 / Q13433
SLC3A1 / Q07837
SLC3A2 / P08195
SLC40A1 / Q9NP59
SLC41A3 / Q96GZ6
SLC43A1 / O75387
SLC43A3 / Q8NBI5
SLC44A1 / Q8WWI5
SLC44A2 / Q8IWA5
SLC45A1 / Q9Y2W3
SLC47A1 / Q96FL8
SLC49A4 / Q96SL1
SLC4A1 / P02730
SLC4A10 / Q6U841
SLC4A1AP / Q9BWU0
SLC4A2 / P04920
SLC4A3 / P48751
SLC4A4 / Q9Y6R1
SLC4A7 / Q9Y6M7
SLC4A8 / Q2Y0W8
SLC5A1 / P13866
SLC5A5 / Q92911
SLC5A6 / Q9Y289
SLC66A2 / Q8N2U9
SLC6A1 / P30531
SLC6A11 / P48066
SLC6A12 / P48065
SLC6A15 / Q9H2J7
SLC6A17 / Q9H1V8
SLC6A19 / Q695T7
SLC6A7 / Q99884
SLC6A8 / P48029
Gene Symbol / UniProt Accession
Brain
Colon
Duodenum
Esophagus
Heart
Liver
Lung
Pancreas
Thyroid
Organs
Type
DDA
DIA
Bins
1
2
3
4
5

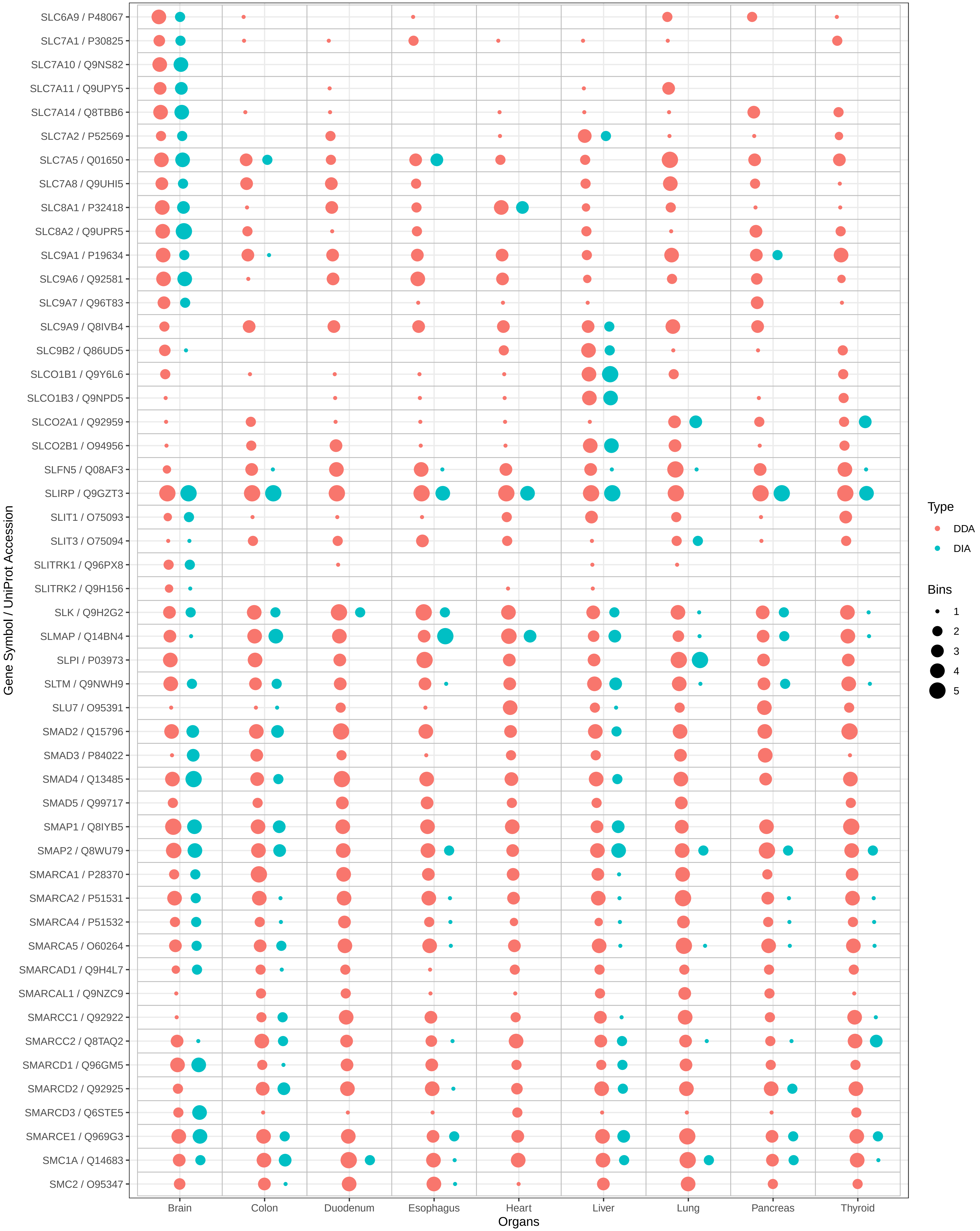
SLC6A9 / P48067
SLC7A1 / P30825
SLC7A10 / Q9NS82
SLC7A11 / Q9UPY5
SLC7A14 / Q8TBB6
SLC7A2 / P52569
SLC7A5 / Q01650
SLC7A8 / Q9UHI5
SLC8A1 / P32418
SLC8A2 / Q9UPR5
SLC9A1 / P19634
SLC9A6 / Q92581
SLC9A7 / Q96T83
SLC9A9 / Q8IVB4
SLC9B2 / Q86UD5
SLCO1B1 / Q9Y6L6
SLCO1B3 / Q9NPD5
SLCO2A1 / Q92959
SLCO2B1 / O94956
SLFN5 / Q08AF3
SLIRP / Q9GZT3
SLIT1 / O75093
SLIT3 / O75094
SLITRK1 / Q96PX8
SLITRK2 / Q9H156
SLK / Q9H2G2
SLMAP / Q14BN4
SLPI / P03973
SLTM / Q9NWH9
SLU7 / O95391
SMAD2 / Q15796
SMAD3 / P84022
SMAD4 / Q13485
SMAD5 / Q99717
SMAP1 / Q8IYB5
SMAP2 / Q8WU79
SMARCA1 / P28370
SMARCA2 / P51531
SMARCA4 / P51532
SMARCA5 / O60264
SMARCAD1 / Q9H4L7
SMARCAL1 / Q9NZC9
SMARCC1 / Q92922
SMARCC2 / Q8TAQ2
SMARCD1 / Q96GM5
SMARCD2 / Q92925
SMARCD3 / Q6STE5
SMARCE1 / Q969G3
SMC1A / Q14683
SMC2 / O95347
Gene Symbol / UniProt Accession
Brain
Colon
Duodenum
Esophagus
Heart
Liver
Lung
Pancreas
Thyroid
Organs
Type
DDA
DIA
Bins
1
2
3
4
5

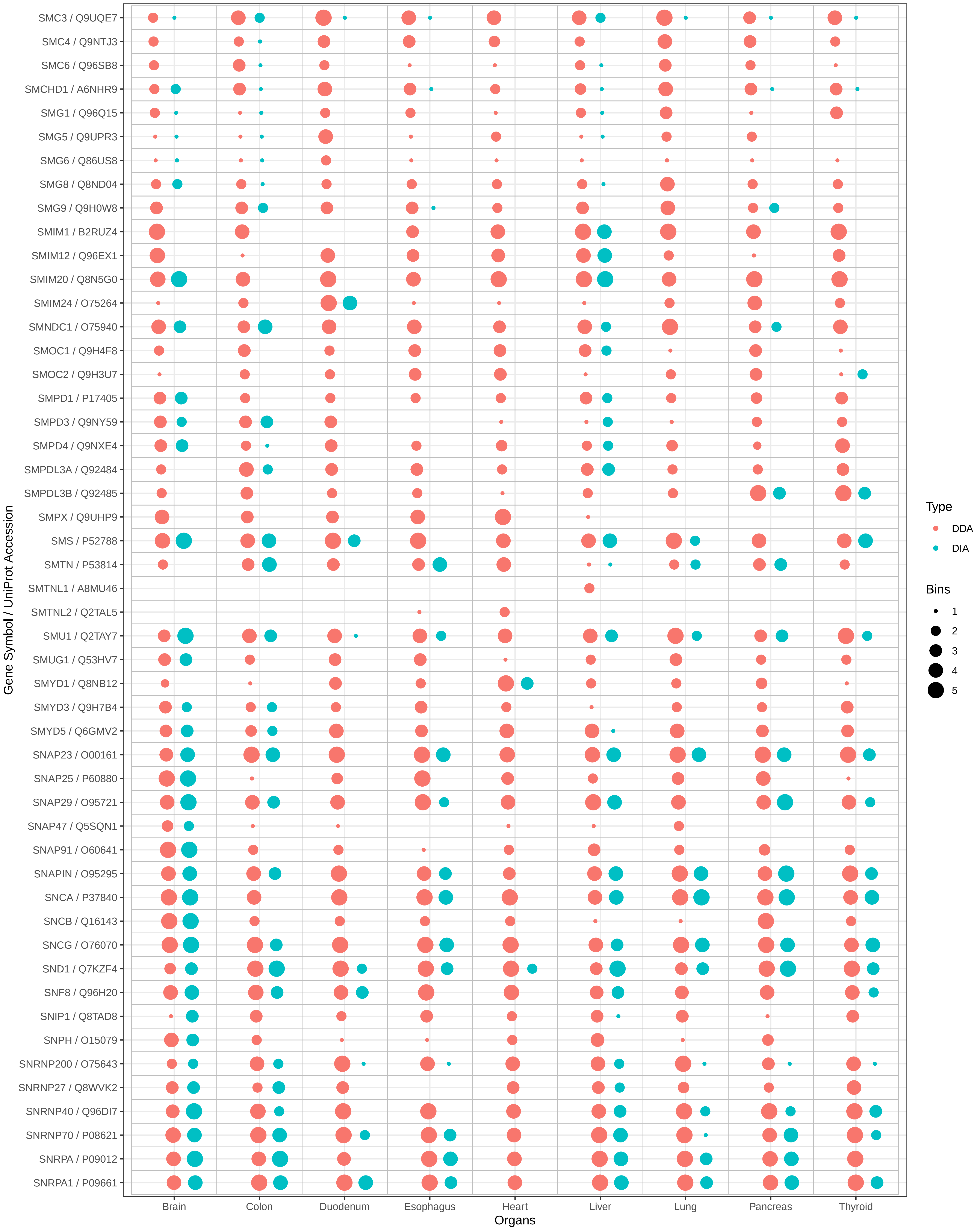

SMC3 / Q9UQE7
SMC4 / Q9NTJ3
SMC6 / Q96SB8
SMCHD1 / A6NHR9
SMG1 / Q96Q15
SMG5 / Q9UPR3
SMG6 / Q86US8
SMG8 / Q8ND04
SMG9 / Q9H0W8
SMIM1 / B2RUZ4
SMIM12 / Q96EX1
SMIM20 / Q8N5G0
SMIM24 / O75264
SMNDC1 / O75940
SMOC1 / Q9H4F8
SMOC2 / Q9H3U7
SMPD1 / P17405
SMPD3 / Q9NY59
SMPD4 / Q9NXE4
SMPDL3A / Q92484
SMPDL3B / Q92485
SMPX / Q9UHP9
SMS / P52788
SMTN / P53814
SMTNL1 / A8MU46
SMTNL2 / Q2TAL5
SMU1 / Q2TAY7
SMUG1 / Q53HV7
SMYD1 / Q8NB12
SMYD3 / Q9H7B4
SMYD5 / Q6GMV2
SNAP23 / O00161
SNAP25 / P60880
SNAP29 / O95721
SNAP47 / Q5SQN1
SNAP91 / O60641
SNAPIN / O95295
SNCA / P37840
SNCB / Q16143
SNCG / O76070
SND1 / Q7KZF4
SNF8 / Q96H20
SNIP1 / Q8TAD8
SNPH / O15079
SNRNP200 / O75643
SNRNP27 / Q8WVK2
SNRNP40 / Q96DI7
SNRNP70 / P08621
SNRPA / P09012
SNRPA1 / P09661
Gene Symbol / UniProt Accession
Brain
Colon
Duodenum
Esophagus
Heart
Liver
Lung
Pancreas
Thyroid
Organs
Type
DDA
DIA
Bins
1
2
3
4
5

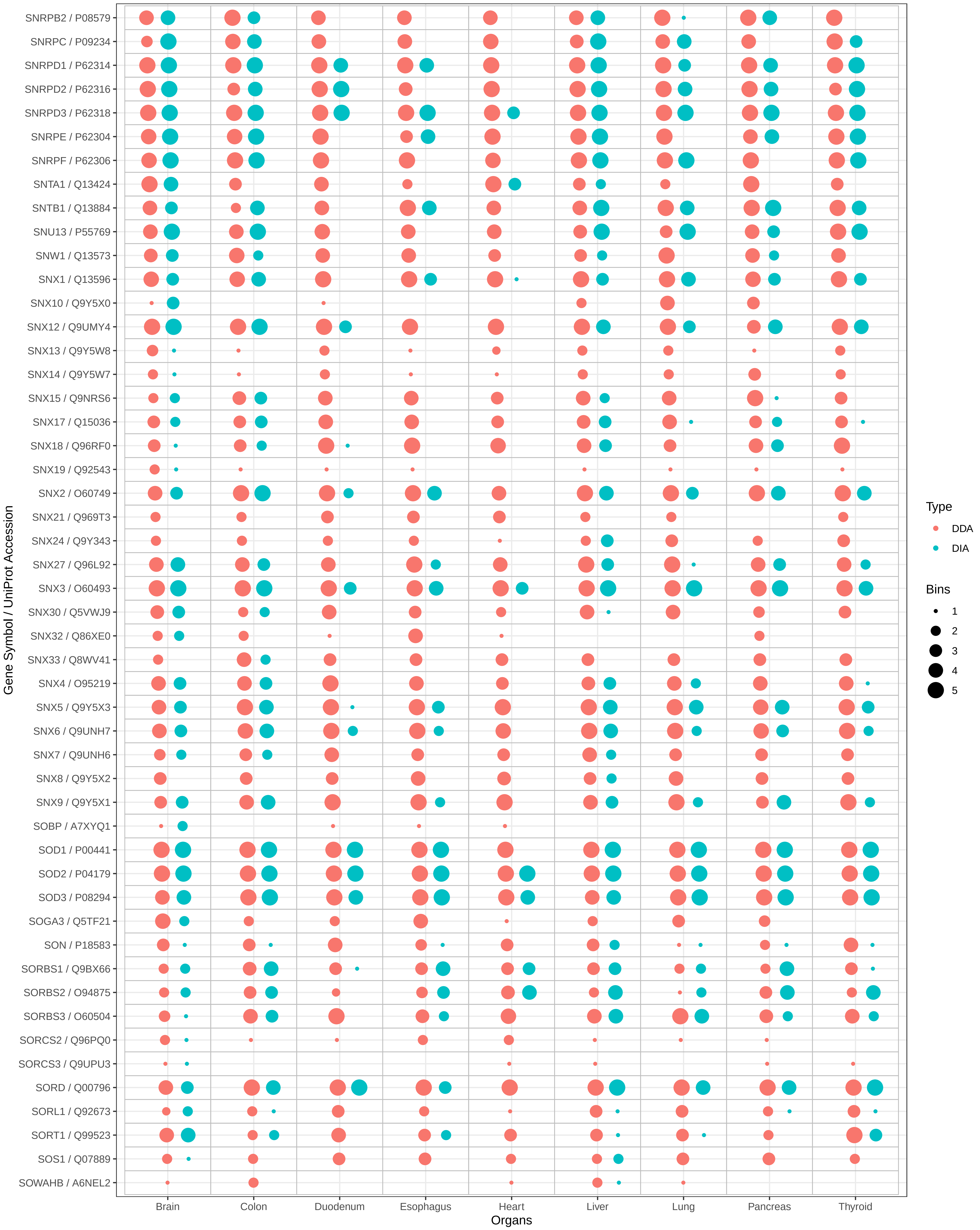
SNRPB2 / P08579
SNRPC / P09234
SNRPD1 / P62314
SNRPD2 / P62316
SNRPD3 / P62318
SNRPE / P62304
SNRPF / P62306
SNTA1 / Q13424
SNTB1 / Q13884
SNU13 / P55769
SNW1 / Q13573
SNX1 / Q13596
SNX10 / Q9Y5X0
SNX12 / Q9UMY4
SNX13 / Q9Y5W8
SNX14 / Q9Y5W7
SNX15 / Q9NRS6
SNX17 / Q15036
SNX18 / Q96RF0
SNX19 / Q92543
SNX2 / O60749
SNX21 / Q969T3
SNX24 / Q9Y343
SNX27 / Q96L92
SNX3 / O60493
SNX30 / Q5VWJ9
SNX32 / Q86XE0
SNX33 / Q8WV41
SNX4 / O95219
SNX5 / Q9Y5X3
SNX6 / Q9UNH7
SNX7 / Q9UNH6
SNX8 / Q9Y5X2
SNX9 / Q9Y5X1
SOBP / A7XYQ1
SOD1 / P00441
SOD2 / P04179
SOD3 / P08294
SOGA3 / Q5TF21
SON / P18583
SORBS1 / Q9BX66
SORBS2 / O94875
SORBS3 / O60504
SORCS2 / Q96PQ0
SORCS3 / Q9UPU3
SORD / Q00796
SORL1 / Q92673
SORT1 / Q99523
SOS1 / Q07889
SOWAHB / A6NEL2
Gene Symbol / UniProt Accession
Brain
Colon
Duodenum
Esophagus
Heart
Liver
Lung
Pancreas
Thyroid
Organs
Type
DDA
DIA
Bins
1
2
3
4
5

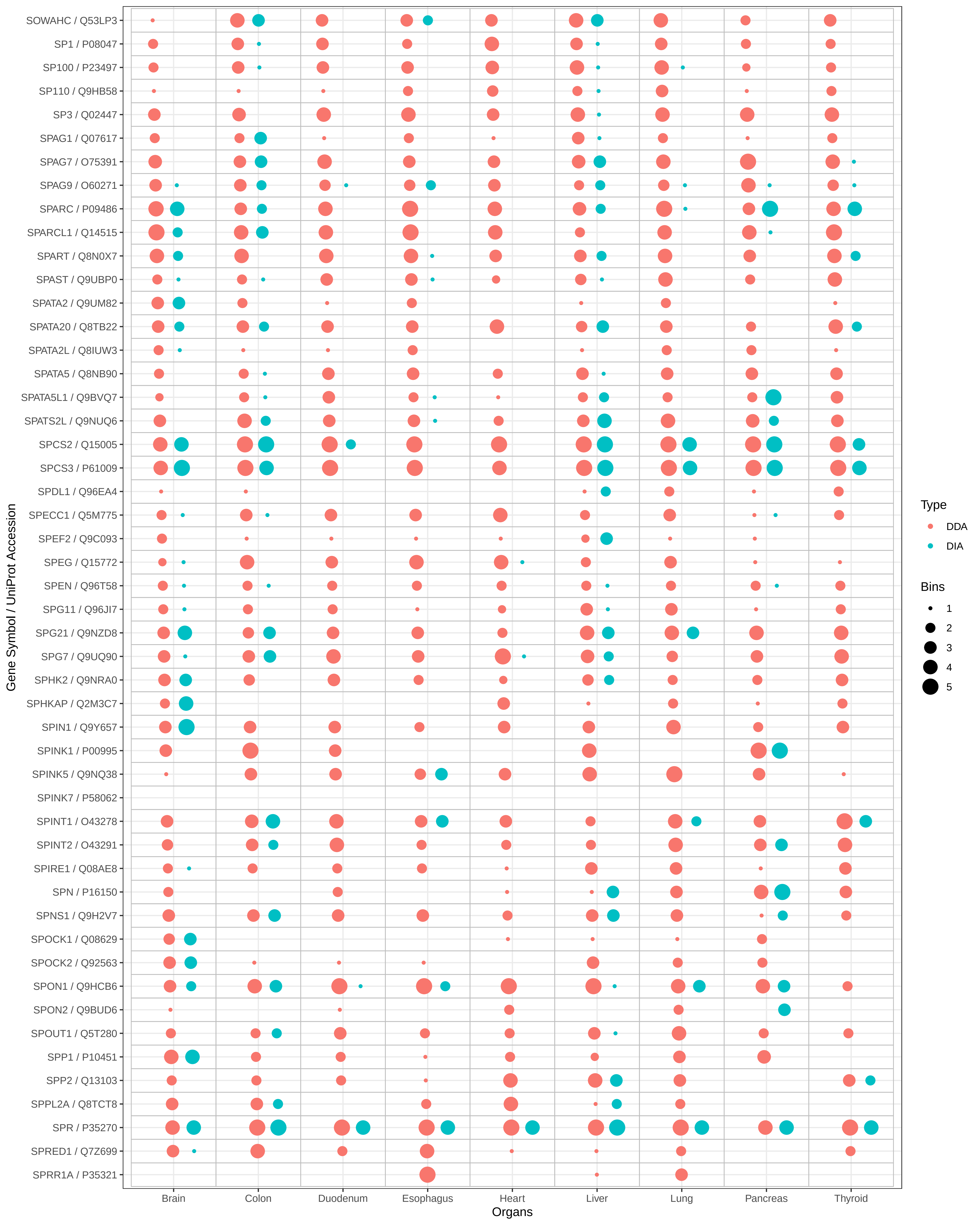

SOWAHC / Q53LP3
SP1 / P08047
SP100 / P23497
SP110 / Q9HB58
SP3 / Q02447
SPAG1 / Q07617
SPAG7 / O75391
SPAG9 / O60271
SPARC / P09486
SPARCL1 / Q14515
SPART / Q8N0X7
SPAST / Q9UBP0
SPATA2 / Q9UM82
SPATA20 / Q8TB22
SPATA2L / Q8IUW3
SPATA5 / Q8NB90
SPATA5L1 / Q9BVQ7
SPATS2L / Q9NUQ6
SPCS2 / Q15005
SPCS3 / P61009
SPDL1 / Q96EA4
SPECC1 / Q5M775
SPEF2 / Q9C093
SPEG / Q15772
SPEN / Q96T58
SPG11 / Q96JI7
SPG21 / Q9NZD8
SPG7 / Q9UQ90
SPHK2 / Q9NRA0
SPHKAP / Q2M3C7
SPIN1 / Q9Y657
SPINK1 / P00995
SPINK5 / Q9NQ38
SPINK7 / P58062
SPINT1 / O43278
SPINT2 / O43291
SPIRE1 / Q08AE8
SPN / P16150
SPNS1 / Q9H2V7
SPOCK1 / Q08629
SPOCK2 / Q92563
SPON1 / Q9HCB6
SPON2 / Q9BUD6
SPOUT1 / Q5T280
SPP1 / P10451
SPP2 / Q13103
SPPL2A / Q8TCT8
SPR / P35270
SPRED1 / Q7Z699
SPRR1A / P35321
Gene Symbol / UniProt Accession
Brain
Colon
Duodenum
Esophagus
Heart
Liver
Lung
Pancreas
Thyroid
Organs
Type
DDA
DIA
Bins
1
2
3
4
5

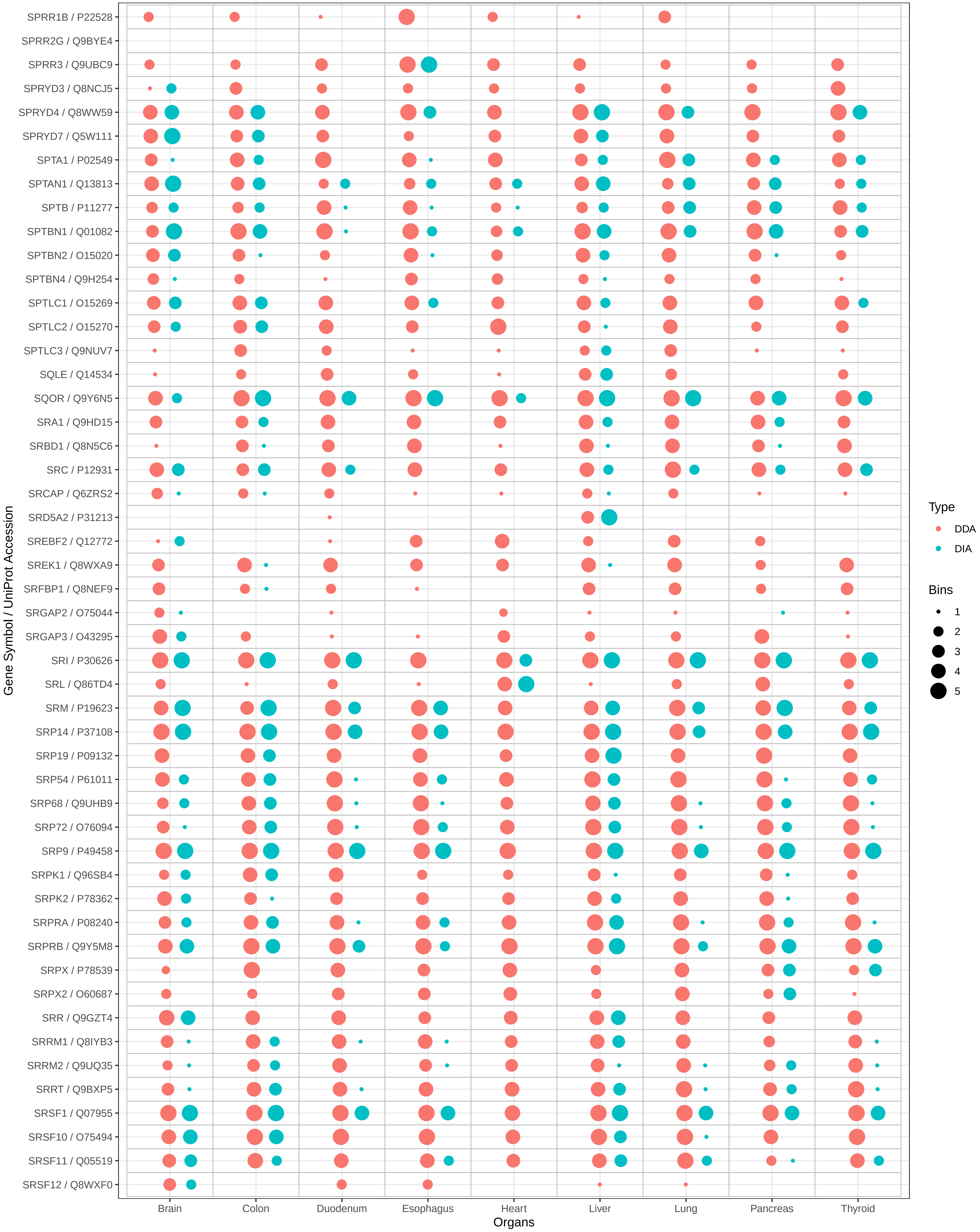

SPRR1B / P22528
SPRR2G / Q9BYE4
SPRR3 / Q9UBC9
SPRYD3 / Q8NCJ5
SPRYD4 / Q8WW59
SPRYD7 / Q5W111
SPTA1 / P02549
SPTAN1 / Q13813
SPTB / P11277
SPTBN1 / Q01082
SPTBN2 / O15020
SPTBN4 / Q9H254
SPTLC1 / O15269
SPTLC2 / O15270
SPTLC3 / Q9NUV7
SQLE / Q14534
SQOR / Q9Y6N5
SRA1 / Q9HD15
SRBD1 / Q8N5C6
SRC / P12931
SRCAP / Q6ZRS2
SRD5A2 / P31213
SREBF2 / Q12772
SREK1 / Q8WXA9
SRFBP1 / Q8NEF9
SRGAP2 / O75044
SRGAP3 / O43295
SRI / P30626
SRL / Q86TD4
SRM / P19623
SRP14 / P37108
SRP19 / P09132
SRP54 / P61011
SRP68 / Q9UHB9
SRP72 / O76094
SRP9 / P49458
SRPK1 / Q96SB4
SRPK2 / P78362
SRPRA / P08240
SRPRB / Q9Y5M8
SRPX / P78539
SRPX2 / O60687
SRR / Q9GZT4
SRRM1 / Q8IYB3
SRRM2 / Q9UQ35
SRRT / Q9BXP5
SRSF1 / Q07955
SRSF10 / O75494
SRSF11 / Q05519
SRSF12 / Q8WXF0
Gene Symbol / UniProt Accession
Brain
Colon
Duodenum
Esophagus
Heart
Liver
Lung
Pancreas
Thyroid
Organs
Type
DDA
DIA
Bins
1
2
3
4
5

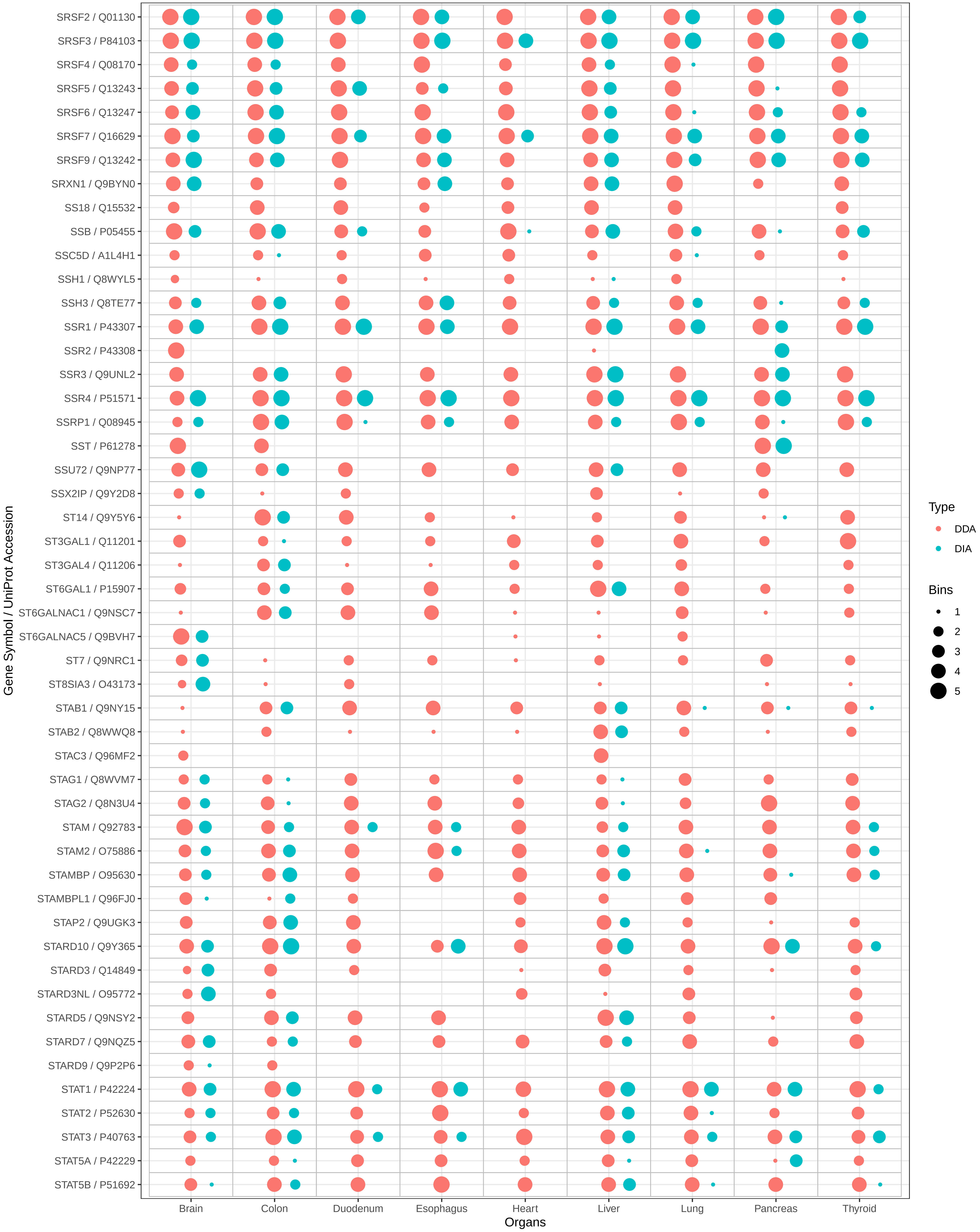
SRSF2 / Q01130
SRSF3 / P84103
SRSF4 / Q08170
SRSF5 / Q13243
SRSF6 / Q13247
SRSF7 / Q16629
SRSF9 / Q13242
SRXN1 / Q9BYN0
SS18 / Q15532
SSB / P05455
SSC5D / A1L4H1
SSH1 / Q8WYL5
SSH3 / Q8TE77
SSR1 / P43307
SSR2 / P43308
SSR3 / Q9UNL2
SSR4 / P51571
SSRP1 / Q08945
SST / P61278
SSU72 / Q9NP77
SSX2IP / Q9Y2D8
ST14 / Q9Y5Y6
ST3GAL1 / Q11201
ST3GAL4 / Q11206
ST6GAL1 / P15907
ST6GALNAC1 / Q9NSC7
ST6GALNAC5 / Q9BVH7
ST7 / Q9NRC1
ST8SIA3 / O43173
STAB1 / Q9NY15
STAB2 / Q8WWQ8
STAC3 / Q96MF2
STAG1 / Q8WVM7
STAG2 / Q8N3U4
STAM / Q92783
STAM2 / O75886
STAMBP / O95630
STAMBPL1 / Q96FJ0
STAP2 / Q9UGK3
STARD10 / Q9Y365
STARD3 / Q14849
STARD3NL / O95772
STARD5 / Q9NSY2
STARD7 / Q9NQZ5
STARD9 / Q9P2P6
STAT1 / P42224
STAT2 / P52630
STAT3 / P40763
STAT5A / P42229
STAT5B / P51692
Gene Symbol / UniProt Accession
Brain
Colon
Duodenum
Esophagus
Heart
Liver
Lung
Pancreas
Thyroid
Organs
Type
DDA
DIA
Bins
1
2
3
4
5

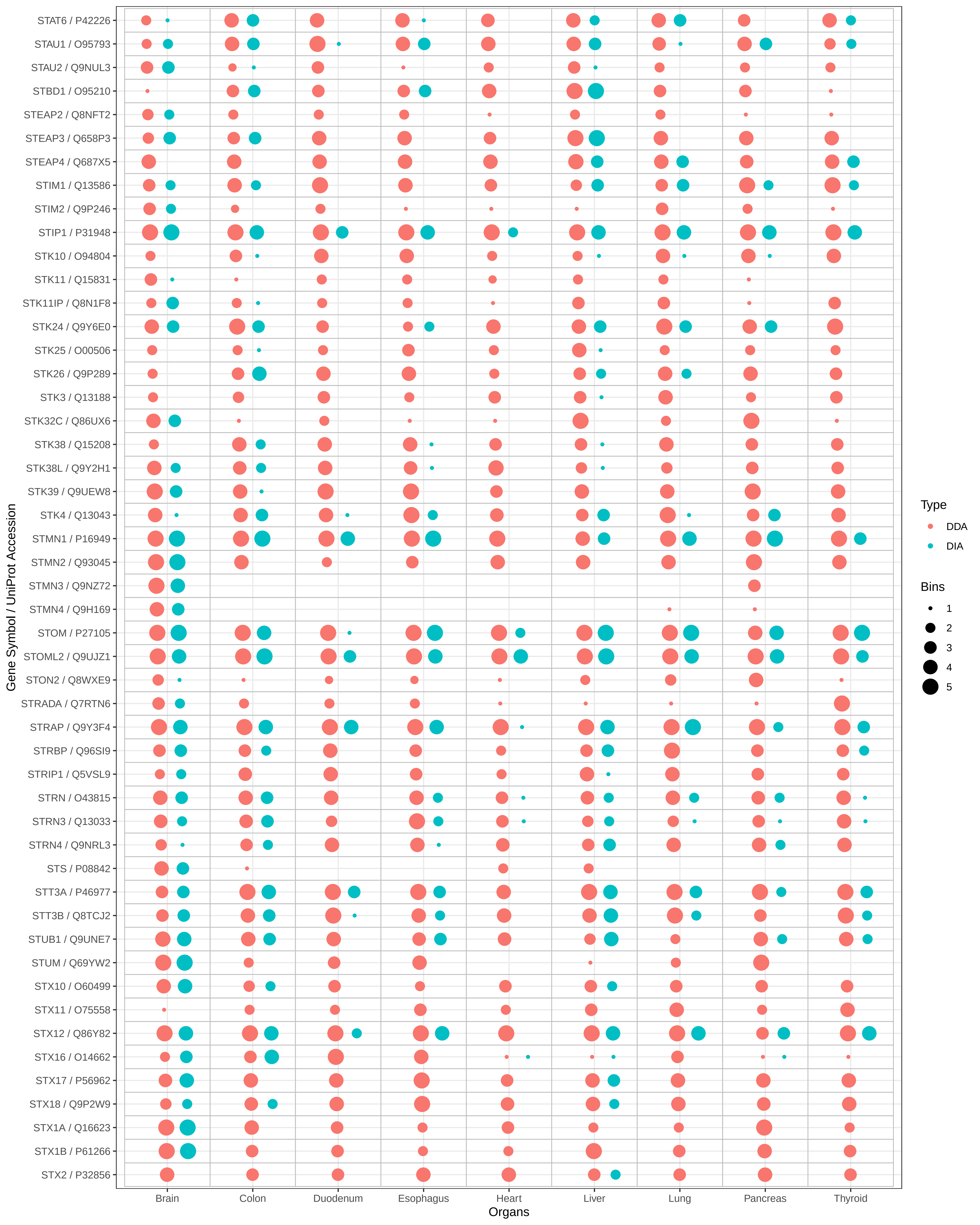

STAT6 / P42226
STAU1 / O95793
STAU2 / Q9NUL3
STBD1 / O95210
STEAP2 / Q8NFT2
STEAP3 / Q658P3
STEAP4 / Q687X5
STIM1 / Q13586
STIM2 / Q9P246
STIP1 / P31948
STK10 / O94804
STK11 / Q15831
STK11IP / Q8N1F8
STK24 / Q9Y6E0
STK25 / O00506
STK26 / Q9P289
STK3 / Q13188
STK32C / Q86UX6
STK38 / Q15208
STK38L / Q9Y2H1
STK39 / Q9UEW8
STK4 / Q13043
STMN1 / P16949
STMN2 / Q93045
STMN3 / Q9NZ72
STMN4 / Q9H169
STOM / P27105
STOML2 / Q9UJZ1
STON2 / Q8WXE9
STRADA / Q7RTN6
STRAP / Q9Y3F4
STRBP / Q96SI9
STRIP1 / Q5VSL9
STRN / O43815
STRN3 / Q13033
STRN4 / Q9NRL3
STS / P08842
STT3A / P46977
STT3B / Q8TCJ2
STUB1 / Q9UNE7
STUM / Q69YW2
STX10 / O60499
STX11 / O75558
STX12 / Q86Y82
STX16 / O14662
STX17 / P56962
STX18 / Q9P2W9
STX1A / Q16623
STX1B / P61266
STX2 / P32856
Gene Symbol / UniProt Accession
Brain
Colon
Duodenum
Esophagus
Heart
Liver
Lung
Pancreas
Thyroid
Organs
Type
DDA
DIA
Bins
1
2
3
4
5

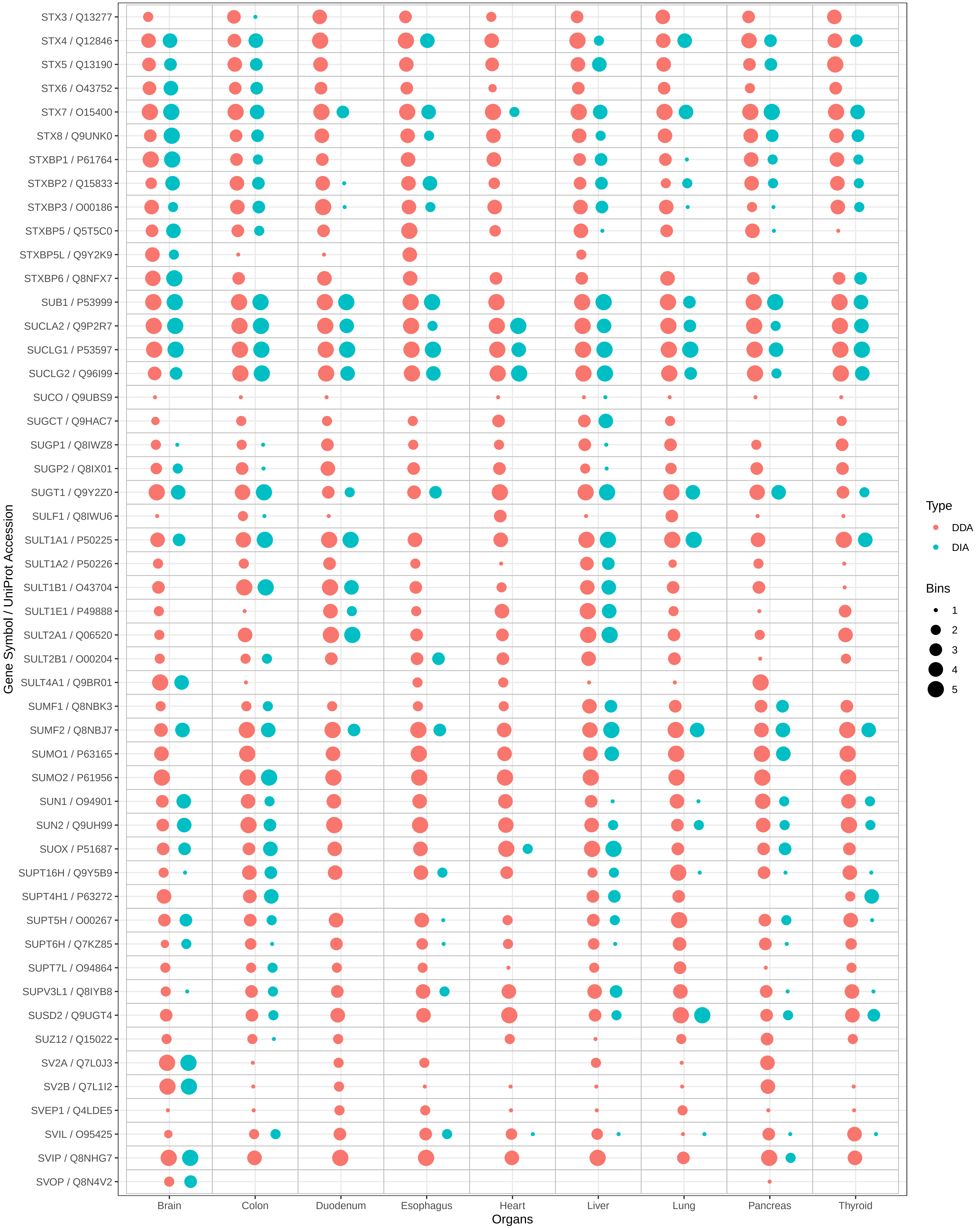

STX3 / Q13277
STX4 / Q12846
STX5 / Q13190
STX6 / O43752
STX7 / O15400
STX8 / Q9UNK0
STXBP1 / P61764
STXBP2 / Q15833
STXBP3 / O00186
STXBP5 / Q5T5C0
STXBP5L / Q9Y2K9
STXBP6 / Q8NFX7
SUB1 / P53999
SUCLA2 / Q9P2R7
SUCLG1 / P53597
SUCLG2 / Q96I99
SUCO / Q9UBS9
SUGCT / Q9HAC7
SUGP1 / Q8IWZ8
SUGP2 / Q8IX01
SUGT1 / Q9Y2Z0
SULF1 / Q8IWU6
SULT1A1 / P50225
SULT1A2 / P50226
SULT1B1 / O43704
SULT1E1 / P49888
SULT2A1 / Q06520
SULT2B1 / O00204
SULT4A1 / Q9BR01
SUMF1 / Q8NBK3
SUMF2 / Q8NBJ7
SUMO1 / P63165
SUMO2 / P61956
SUN1 / O94901
SUN2 / Q9UH99
SUOX / P51687
SUPT16H / Q9Y5B9
SUPT4H1 / P63272
SUPT5H / O00267
SUPT6H / Q7KZ85
SUPT7L / O94864
SUPV3L1 / Q8IYB8
SUSD2 / Q9UGT4
SUZ12 / Q15022
SV2A / Q7L0J3
SV2B / Q7L1I2
SVEP1 / Q4LDE5
SVIL / O95425
SVIP / Q8NHG7
SVOP / Q8N4V2
Gene Symbol / UniProt Accession
Brain
Colon
Duodenum
Esophagus
Heart
Liver
Lung
Pancreas
Thyroid
Organs
Type
DDA
DIA
Bins
1
2
3
4
5

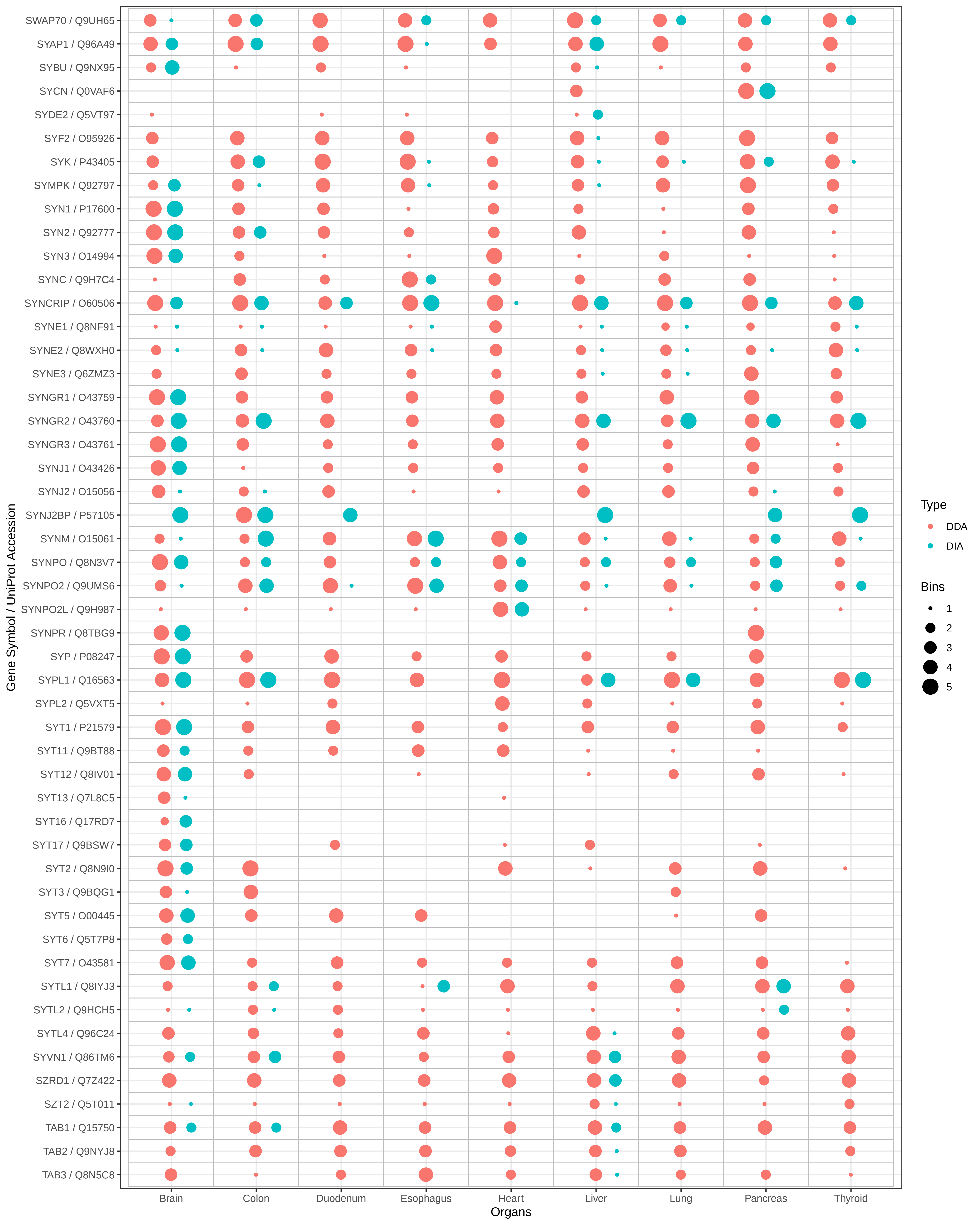
SWAP70 / Q9UH65
SYAP1 / Q96A49
SYBU / Q9NX95
SYCN / Q0VAF6
SYDE2 / Q5VT97
SYF2 / O95926
SYK / P43405
SYMPK / Q92797
SYN1 / P17600
SYN2 / Q92777
SYN3 / O14994
SYNC / Q9H7C4
SYNCRIP / O60506
SYNE1 / Q8NF91
SYNE2 / Q8WXH0
SYNE3 / Q6ZMZ3
SYNGR1 / O43759
SYNGR2 / O43760
SYNGR3 / O43761
SYNJ1 / O43426
SYNJ2 / O15056
SYNJ2BP / P57105
SYNM / O15061
SYNPO / Q8N3V7
SYNPO2 / Q9UMS6
SYNPO2L / Q9H987
SYNPR / Q8TBG9
SYP / P08247
SYPL1 / Q16563
SYPL2 / Q5VXT5
SYT1 / P21579
SYT11 / Q9BT88
SYT12 / Q8IV01
SYT13 / Q7L8C5
SYT16 / Q17RD7
SYT17 / Q9BSW7
SYT2 / Q8N9I0
SYT3 / Q9BQG1
SYT5 / O00445
SYT6 / Q5T7P8
SYT7 / O43581
SYTL1 / Q8IYJ3
SYTL2 / Q9HCH5
SYTL4 / Q96C24
SYVN1 / Q86TM6
SZRD1 / Q7Z422
SZT2 / Q5T011
TAB1 / Q15750
TAB2 / Q9NYJ8
TAB3 / Q8N5C8
Gene Symbol / UniProt Accession
Brain
Colon
Duodenum
Esophagus
Heart
Liver
Lung
Pancreas
Thyroid
Organs
Type
DDA
DIA
Bins
1
2
3
4
5

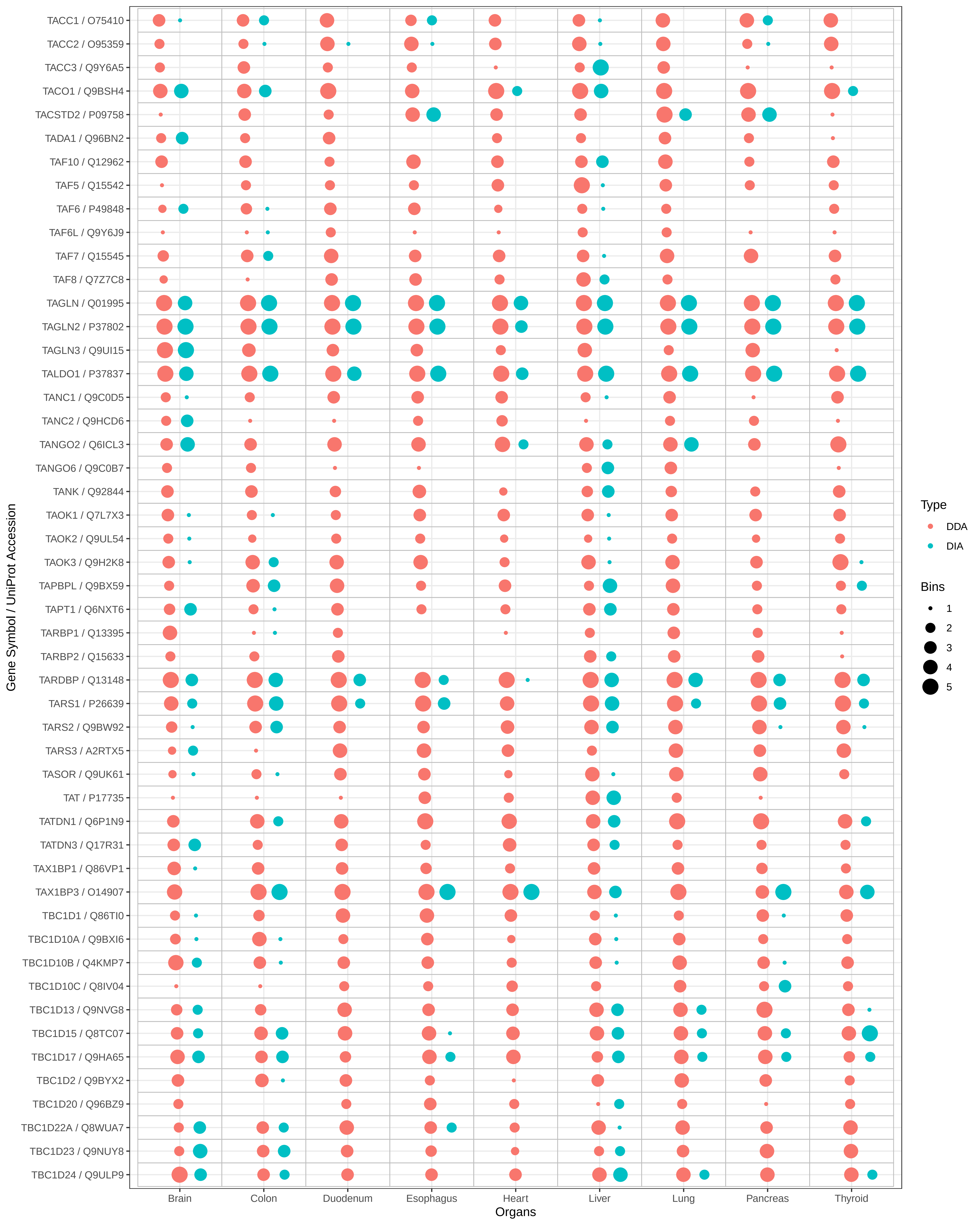

TACC1 / O75410
TACC2 / O95359
TACC3 / Q9Y6A5
TACO1 / Q9BSH4
TACSTD2 / P09758
TADA1 / Q96BN2
TAF10 / Q12962
TAF5 / Q15542
TAF6 / P49848
TAF6L / Q9Y6J9
TAF7 / Q15545
TAF8 / Q7Z7C8
TAGLN / Q01995
TAGLN2 / P37802
TAGLN3 / Q9UI15
TALDO1 / P37837
TANC1 / Q9C0D5
TANC2 / Q9HCD6
TANGO2 / Q6ICL3
TANGO6 / Q9C0B7
TANK / Q92844
TAOK1 / Q7L7X3
TAOK2 / Q9UL54
TAOK3 / Q9H2K8
TAPBPL / Q9BX59
TAPT1 / Q6NXT6
TARBP1 / Q13395
TARBP2 / Q15633
TARDBP / Q13148
TARS1 / P26639
TARS2 / Q9BW92
TARS3 / A2RTX5
TASOR / Q9UK61
TAT / P17735
TATDN1 / Q6P1N9
TATDN3 / Q17R31
TAX1BP1 / Q86VP1
TAX1BP3 / O14907
TBC1D1 / Q86TI0
TBC1D10A / Q9BXI6
TBC1D10B / Q4KMP7
TBC1D10C / Q8IV04
TBC1D13 / Q9NVG8
TBC1D15 / Q8TC07
TBC1D17 / Q9HA65
TBC1D2 / Q9BYX2
TBC1D20 / Q96BZ9
TBC1D22A / Q8WUA7
TBC1D23 / Q9NUY8
TBC1D24 / Q9ULP9
Gene Symbol / UniProt Accession
Brain
Colon
Duodenum
Esophagus
Heart
Liver
Lung
Pancreas
Thyroid
Organs
Type
DDA
DIA
Bins
1
2
3
4
5

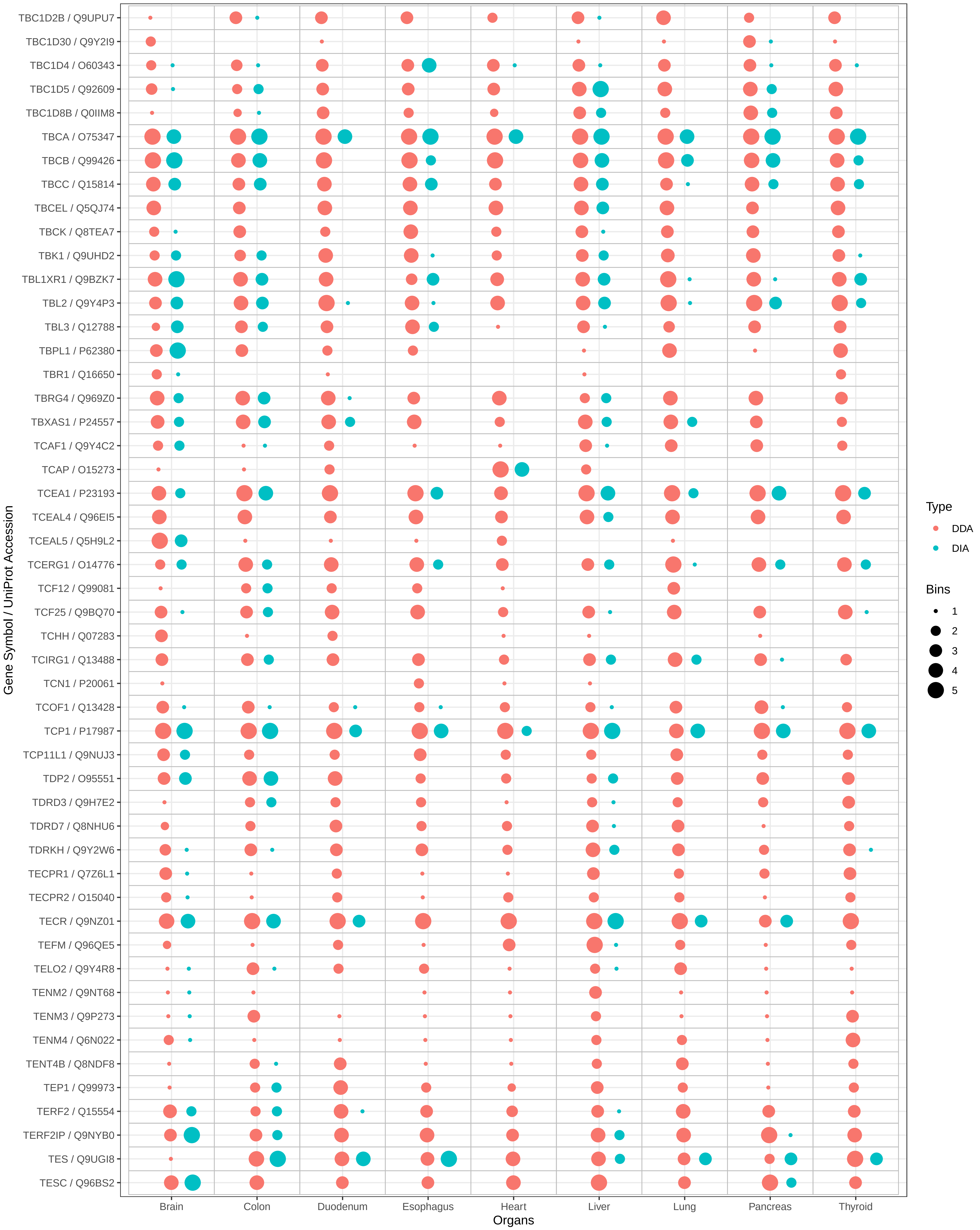
TBC1D2B / Q9UPU7
TBC1D30 / Q9Y2I9
TBC1D4 / O60343
TBC1D5 / Q92609
TBC1D8B / Q0IIM8
TBCA / O75347
TBCB / Q99426
TBCC / Q15814
TBCEL / Q5QJ74
TBCK / Q8TEA7
TBK1 / Q9UHD2
TBL1XR1 / Q9BZK7
TBL2 / Q9Y4P3
TBL3 / Q12788
TBPL1 / P62380
TBR1 / Q16650
TBRG4 / Q969Z0
TBXAS1 / P24557
TCAF1 / Q9Y4C2
TCAP / O15273
TCEA1 / P23193
TCEAL4 / Q96EI5
TCEAL5 / Q5H9L2
TCERG1 / O14776
TCF12 / Q99081
TCF25 / Q9BQ70
TCHH / Q07283
TCIRG1 / Q13488
TCN1 / P20061
TCOF1 / Q13428
TCP1 / P17987
TCP11L1 / Q9NUJ3
TDP2 / O95551
TDRD3 / Q9H7E2
TDRD7 / Q8NHU6
TDRKH / Q9Y2W6
TECPR1 / Q7Z6L1
TECPR2 / O15040
TECR / Q9NZ01
TEFM / Q96QE5
TELO2 / Q9Y4R8
TENM2 / Q9NT68
TENM3 / Q9P273
TENM4 / Q6N022
TENT4B / Q8NDF8
TEP1 / Q99973
TERF2 / Q15554
TERF2IP / Q9NYB0
TES / Q9UGI8
TESC / Q96BS2
Gene Symbol / UniProt Accession
Brain
Colon
Duodenum
Esophagus
Heart
Liver
Lung
Pancreas
Thyroid
Organs
Type
DDA
DIA
Bins
1
2
3
4
5

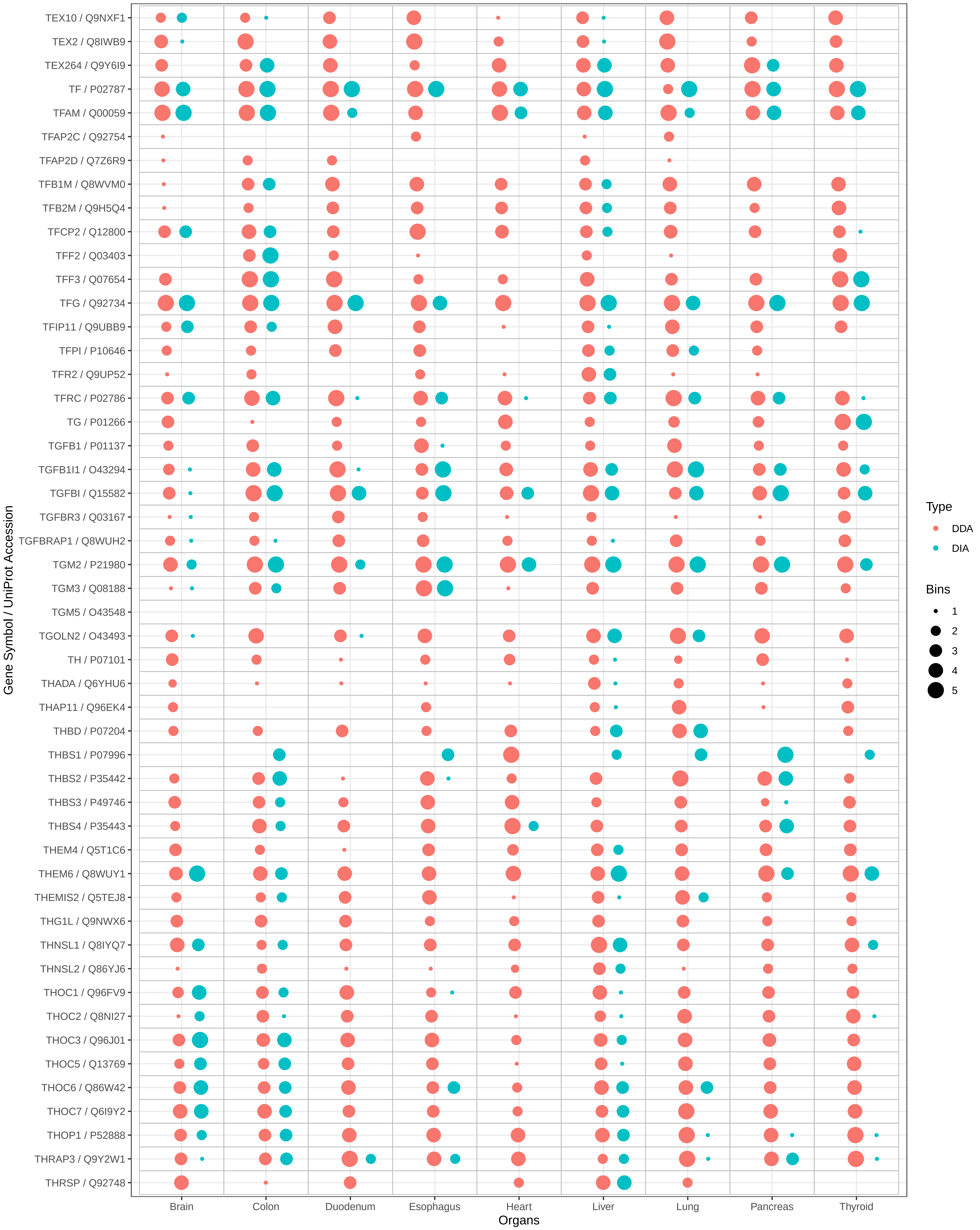
TEX10 / Q9NXF1
TEX2 / Q8IWB9
TEX264 / Q9Y6I9
TF / P02787
TFAM / Q00059
TFAP2C / Q92754
TFAP2D / Q7Z6R9
TFB1M / Q8WVM0
TFB2M / Q9H5Q4
TFCP2 / Q12800
TFF2 / Q03403
TFF3 / Q07654
TFG / Q92734
TFIP11 / Q9UBB9
TFPI / P10646
TFR2 / Q9UP52
TFRC / P02786
TG / P01266
TGFB1 / P01137
TGFB1I1 / O43294
TGFBI / Q15582
TGFBR3 / Q03167
TGFBRAP1 / Q8WUH2
TGM2 / P21980
TGM3 / Q08188
TGM5 / O43548
TGOLN2 / O43493
TH / P07101
THADA / Q6YHU6
THAP11 / Q96EK4
THBD / P07204
THBS1 / P07996
THBS2 / P35442
THBS3 / P49746
THBS4 / P35443
THEM4 / Q5T1C6
THEM6 / Q8WUY1
THEMIS2 / Q5TEJ8
THG1L / Q9NWX6
THNSL1 / Q8IYQ7
THNSL2 / Q86YJ6
THOC1 / Q96FV9
THOC2 / Q8NI27
THOC3 / Q96J01
THOC5 / Q13769
THOC6 / Q86W42
THOC7 / Q6I9Y2
THOP1 / P52888
THRAP3 / Q9Y2W1
THRSP / Q92748
Gene Symbol / UniProt Accession
Brain
Colon
Duodenum
Esophagus
Heart
Liver
Lung
Pancreas
Thyroid
Organs
Type
DDA
DIA
Bins
1
2
3
4
5

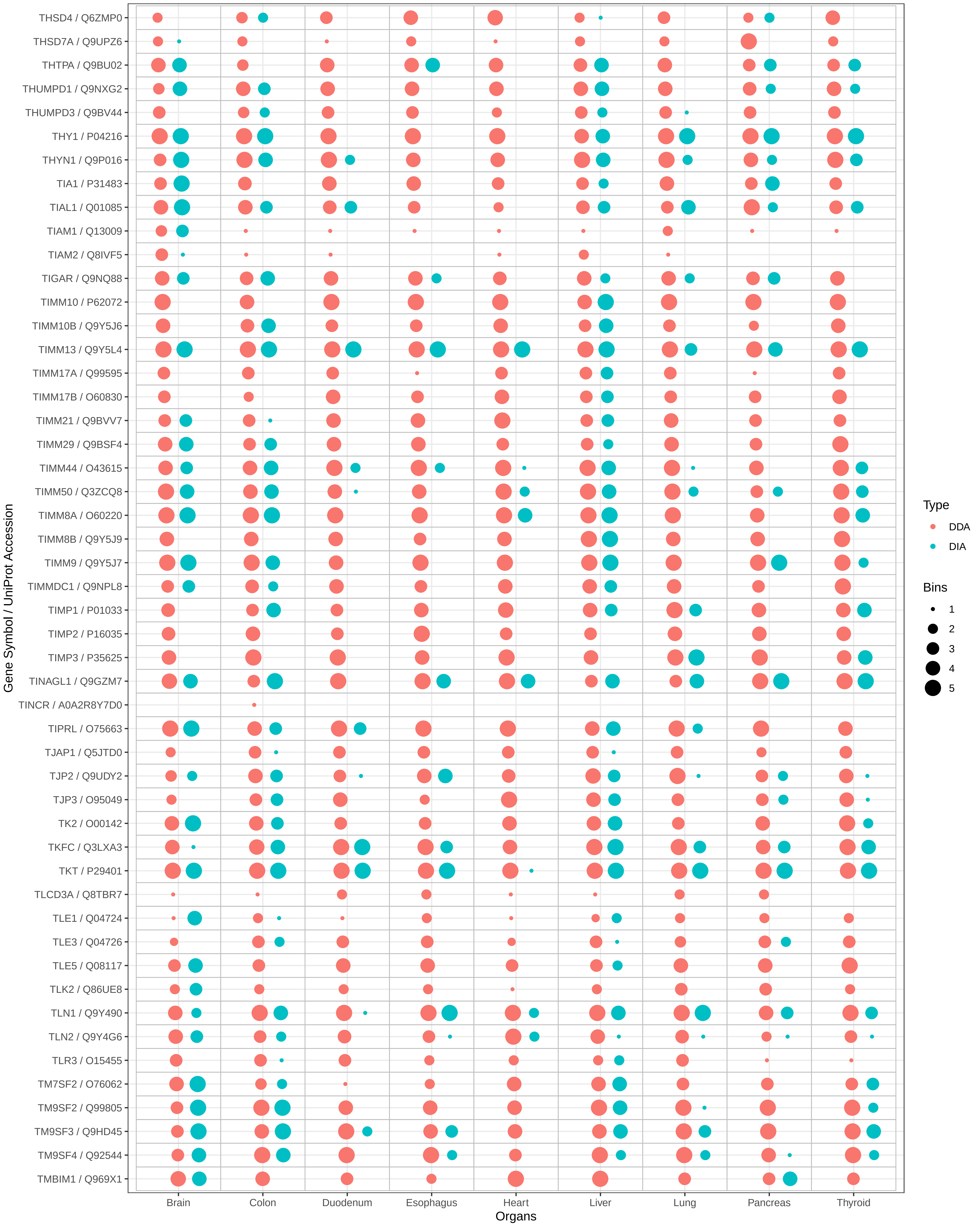

THSD4 / Q6ZMP0
THSD7A / Q9UPZ6
THTPA / Q9BU02
THUMPD1 / Q9NXG2
THUMPD3 / Q9BV44
THY1 / P04216
THYN1 / Q9P016
TIA1 / P31483
TIAL1 / Q01085
TIAM1 / Q13009
TIAM2 / Q8IVF5
TIGAR / Q9NQ88
TIMM10 / P62072
TIMM10B / Q9Y5J6
TIMM13 / Q9Y5L4
TIMM17A / Q99595
TIMM17B / O60830
TIMM21 / Q9BVV7
TIMM29 / Q9BSF4
TIMM44 / O43615
TIMM50 / Q3ZCQ8
TIMM8A / O60220
TIMM8B / Q9Y5J9
TIMM9 / Q9Y5J7
TIMMDC1 / Q9NPL8
TIMP1 / P01033
TIMP2 / P16035
TIMP3 / P35625
TINAGL1 / Q9GZM7
TINCR / A0A2R8Y7D0
TIPRL / O75663
TJAP1 / Q5JTD0
TJP2 / Q9UDY2
TJP3 / O95049
TK2 / O00142
TKFC / Q3LXA3
TKT / P29401
TLCD3A / Q8TBR7
TLE1 / Q04724
TLE3 / Q04726
TLE5 / Q08117
TLK2 / Q86UE8
TLN1 / Q9Y490
TLN2 / Q9Y4G6
TLR3 / O15455
TM7SF2 / O76062
TM9SF2 / Q99805
TM9SF3 / Q9HD45
TM9SF4 / Q92544
TMBIM1 / Q969X1
Gene Symbol / UniProt Accession
Brain
Colon
Duodenum
Esophagus
Heart
Liver
Lung
Pancreas
Thyroid
Organs
Type
DDA
DIA
Bins
1
2
3
4
5

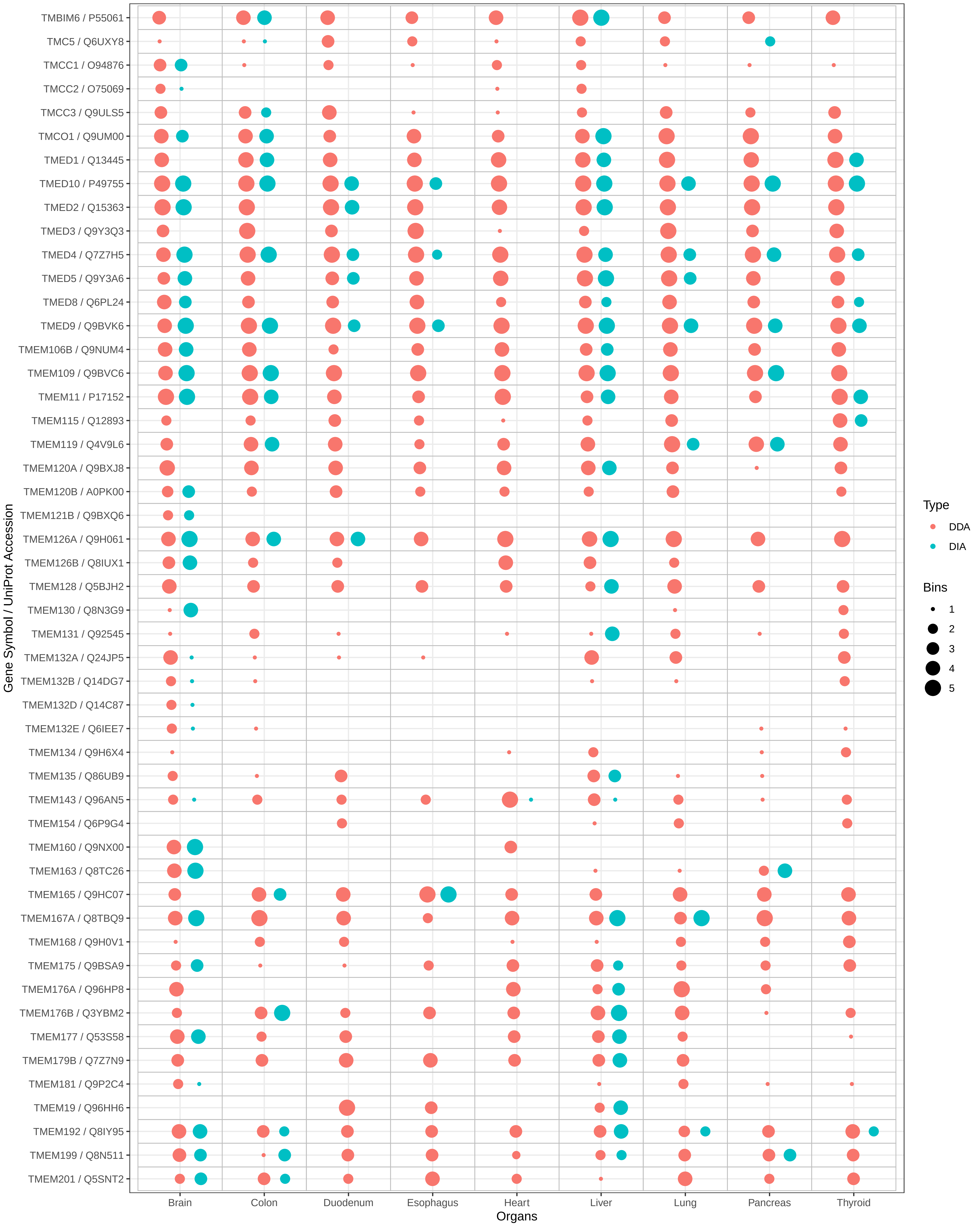

TMBIM6 / P55061
TMC5 / Q6UXY8
TMCC1 / O94876
TMCC2 / O75069
TMCC3 / Q9ULS5
TMCO1 / Q9UM00
TMED1 / Q13445
TMED10 / P49755
TMED2 / Q15363
TMED3 / Q9Y3Q3
TMED4 / Q7Z7H5
TMED5 / Q9Y3A6
TMED8 / Q6PL24
TMED9 / Q9BVK6
TMEM106B / Q9NUM4
TMEM109 / Q9BVC6
TMEM11 / P17152
TMEM115 / Q12893
TMEM119 / Q4V9L6
TMEM120A / Q9BXJ8
TMEM120B / A0PK00
TMEM121B / Q9BXQ6
TMEM126A / Q9H061
TMEM126B / Q8IUX1
TMEM128 / Q5BJH2
TMEM130 / Q8N3G9
TMEM131 / Q92545
TMEM132A / Q24JP5
TMEM132B / Q14DG7
TMEM132D / Q14C87
TMEM132E / Q6IEE7
TMEM134 / Q9H6X4
TMEM135 / Q86UB9
TMEM143 / Q96AN5
TMEM154 / Q6P9G4
TMEM160 / Q9NX00
TMEM163 / Q8TC26
TMEM165 / Q9HC07
TMEM167A / Q8TBQ9
TMEM168 / Q9H0V1
TMEM175 / Q9BSA9
TMEM176A / Q96HP8
TMEM176B / Q3YBM2
TMEM177 / Q53S58
TMEM179B / Q7Z7N9
TMEM181 / Q9P2C4
TMEM19 / Q96HH6
TMEM192 / Q8IY95
TMEM199 / Q8N511
TMEM201 / Q5SNT2
Gene Symbol / UniProt Accession
Brain
Colon
Duodenum
Esophagus
Heart
Liver
Lung
Pancreas
Thyroid
Organs
Type
DDA
DIA
Bins
1
2
3
4
5

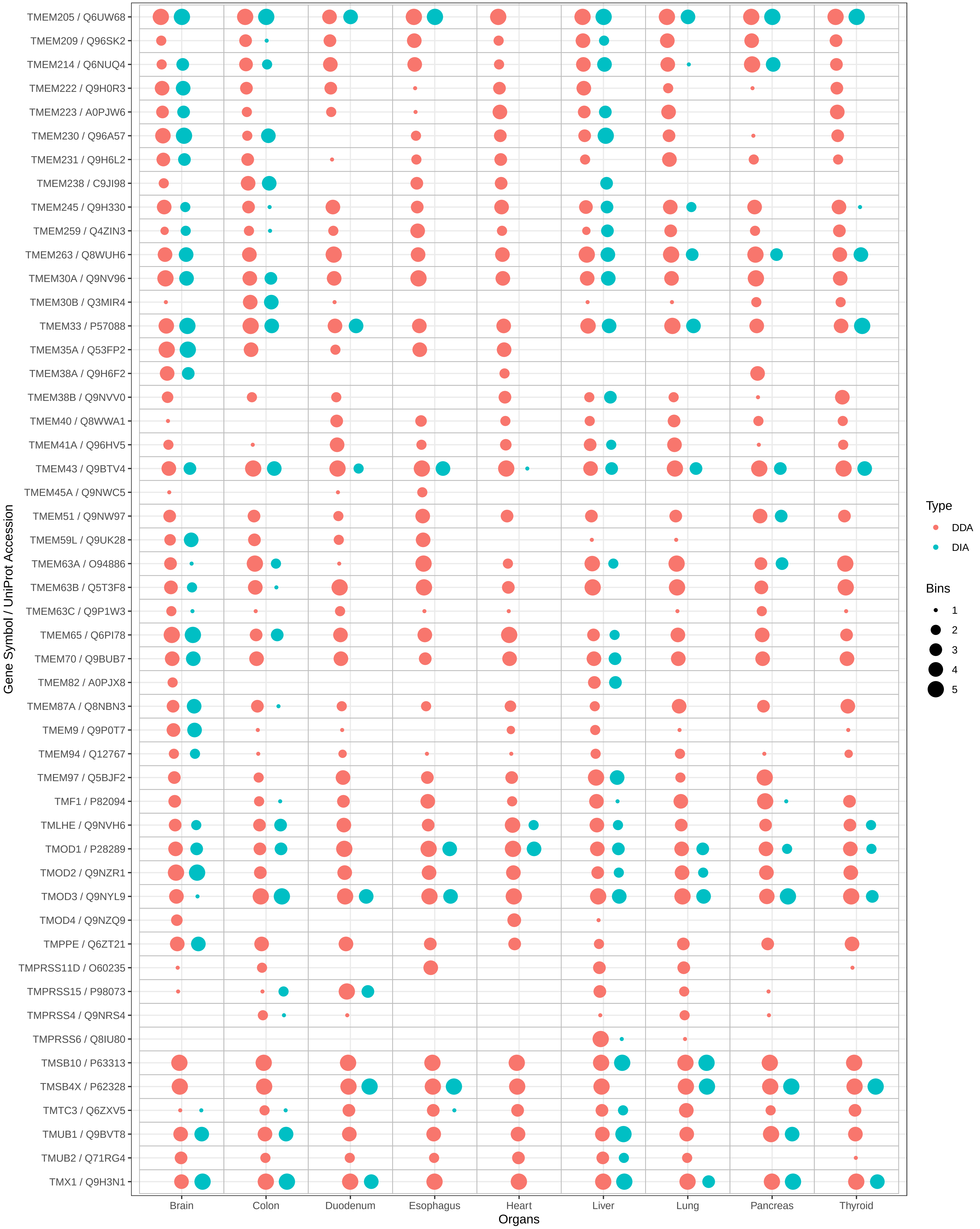
TMEM205 / Q6UW68
TMEM209 / Q96SK2
TMEM214 / Q6NUQ4
TMEM222 / Q9H0R3
TMEM223 / A0PJW6
TMEM230 / Q96A57
TMEM231 / Q9H6L2
TMEM238 / C9JI98
TMEM245 / Q9H330
TMEM259 / Q4ZIN3
TMEM263 / Q8WUH6
TMEM30A / Q9NV96
TMEM30B / Q3MIR4
TMEM33 / P57088
TMEM35A / Q53FP2
TMEM38A / Q9H6F2
TMEM38B / Q9NVV0
TMEM40 / Q8WWA1
TMEM41A / Q96HV5
TMEM43 / Q9BTV4
TMEM45A / Q9NWC5
TMEM51 / Q9NW97
TMEM59L / Q9UK28
TMEM63A / O94886
TMEM63B / Q5T3F8
TMEM63C / Q9P1W3
TMEM65 / Q6PI78
TMEM70 / Q9BUB7
TMEM82 / A0PJX8
TMEM87A / Q8NBN3
TMEM9 / Q9P0T7
TMEM94 / Q12767
TMEM97 / Q5BJF2
TMF1 / P82094
TMLHE / Q9NVH6
TMOD1 / P28289
TMOD2 / Q9NZR1
TMOD3 / Q9NYL9
TMOD4 / Q9NZQ9
TMPPE / Q6ZT21
TMPRSS11D / O60235
TMPRSS15 / P98073
TMPRSS4 / Q9NRS4
TMPRSS6 / Q8IU80
TMSB10 / P63313
TMSB4X / P62328
TMTC3 / Q6ZXV5
TMUB1 / Q9BVT8
TMUB2 / Q71RG4
TMX1 / Q9H3N1
Gene Symbol / UniProt Accession
Brain
Colon
Duodenum
Esophagus
Heart
Liver
Lung
Pancreas
Thyroid
Organs
Type
DDA
DIA
Bins
1
2
3
4
5

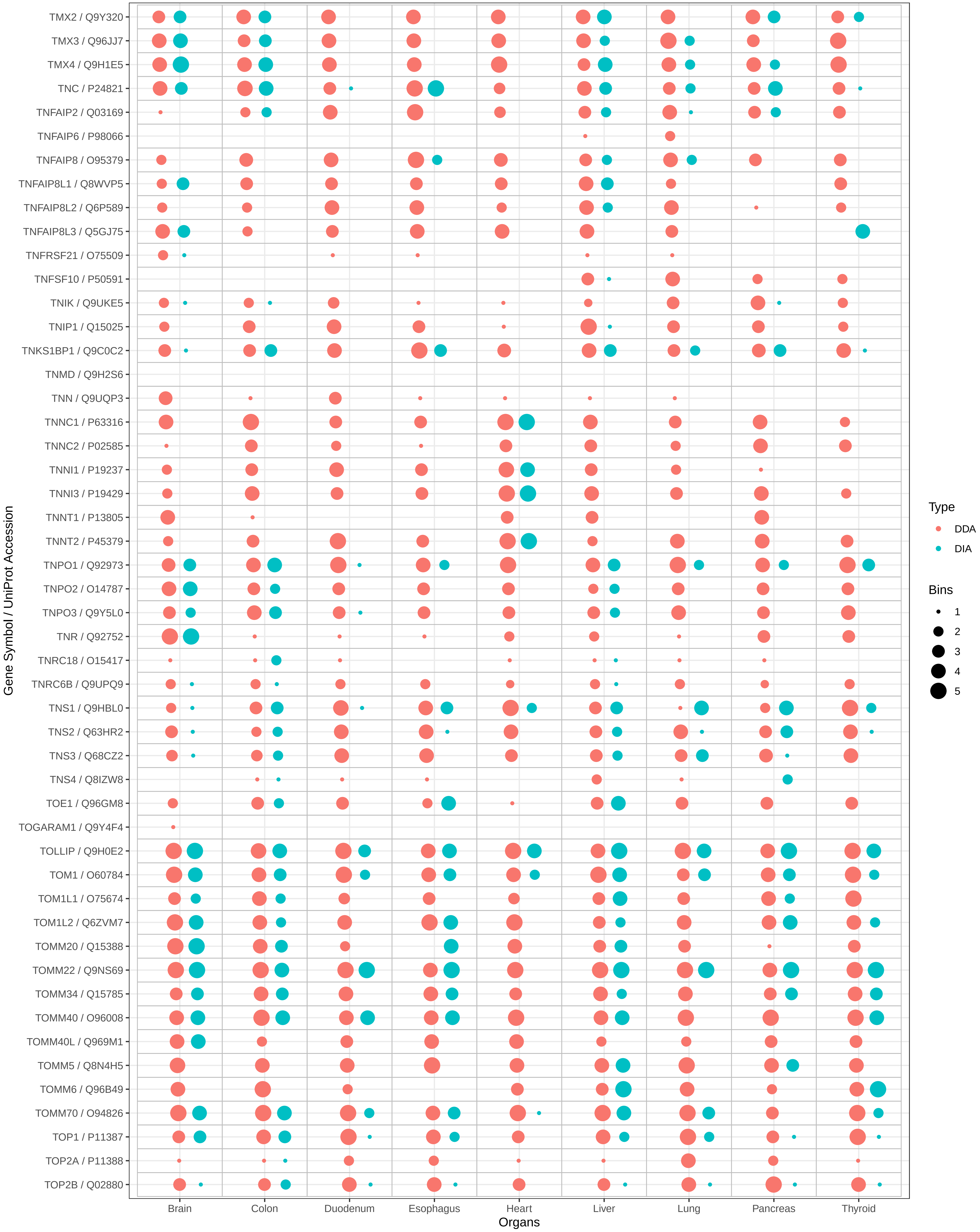

TMX2 / Q9Y320
TMX3 / Q96JJ7
TMX4 / Q9H1E5
TNC / P24821
TNFAIP2 / Q03169
TNFAIP6 / P98066
TNFAIP8 / O95379
TNFAIP8L1 / Q8WVP5
TNFAIP8L2 / Q6P589
TNFAIP8L3 / Q5GJ75
TNFRSF21 / O75509
TNFSF10 / P50591
TNIK / Q9UKE5
TNIP1 / Q15025
TNKS1BP1 / Q9C0C2
TNMD / Q9H2S6
TNN / Q9UQP3
TNNC1 / P63316
TNNC2 / P02585
TNNI1 / P19237
TNNI3 / P19429
TNNT1 / P13805
TNNT2 / P45379
TNPO1 / Q92973
TNPO2 / O14787
TNPO3 / Q9Y5L0
TNR / Q92752
TNRC18 / O15417
TNRC6B / Q9UPQ9
TNS1 / Q9HBL0
TNS2 / Q63HR2
TNS3 / Q68CZ2
TNS4 / Q8IZW8
TOE1 / Q96GM8
TOGARAM1 / Q9Y4F4
TOLLIP / Q9H0E2
TOM1 / O60784
TOM1L1 / O75674
TOM1L2 / Q6ZVM7
TOMM20 / Q15388
TOMM22 / Q9NS69
TOMM34 / Q15785
TOMM40 / O96008
TOMM40L / Q969M1
TOMM5 / Q8N4H5
TOMM6 / Q96B49
TOMM70 / O94826
TOP1 / P11387
TOP2A / P11388
TOP2B / Q02880
Gene Symbol / UniProt Accession
Brain
Colon
Duodenum
Esophagus
Heart
Liver
Lung
Pancreas
Thyroid
Organs
Type
DDA
DIA
Bins
1
2
3
4
5

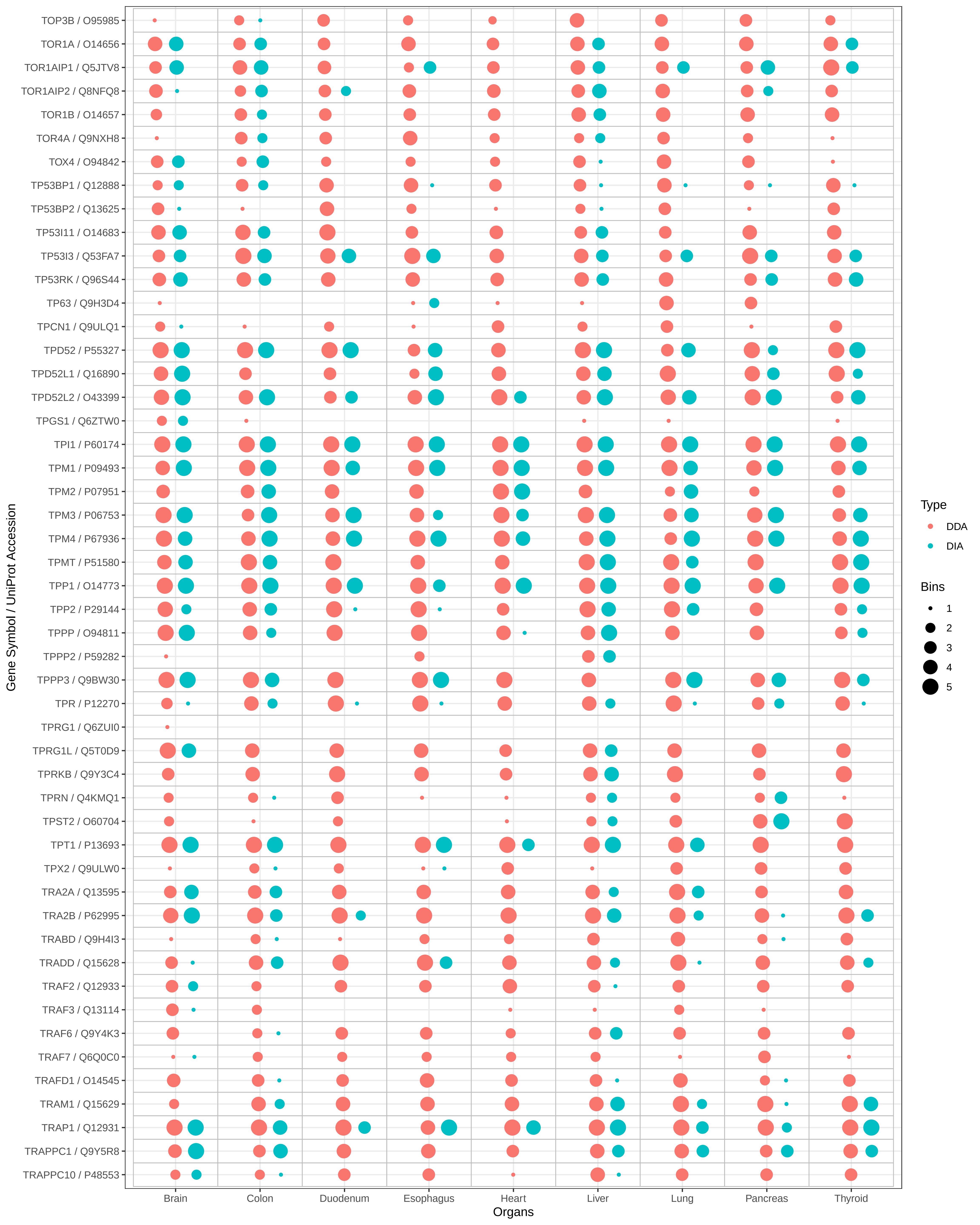

TOP3B / O95985
TOR1A / O14656
TOR1AIP1 / Q5JTV8
TOR1AIP2 / Q8NFQ8
TOR1B / O14657
TOR4A / Q9NXH8
TOX4 / O94842
TP53BP1 / Q12888
TP53BP2 / Q13625
TP53I11 / O14683
TP53I3 / Q53FA7
TP53RK / Q96S44
TP63 / Q9H3D4
TPCN1 / Q9ULQ1
TPD52 / P55327
TPD52L1 / Q16890
TPD52L2 / O43399
TPGS1 / Q6ZTW0
TPI1 / P60174
TPM1 / P09493
TPM2 / P07951
TPM3 / P06753
TPM4 / P67936
TPMT / P51580
TPP1 / O14773
TPP2 / P29144
TPPP / O94811
TPPP2 / P59282
TPPP3 / Q9BW30
TPR / P12270
TPRG1 / Q6ZUI0
TPRG1L / Q5T0D9
TPRKB / Q9Y3C4
TPRN / Q4KMQ1
TPST2 / O60704
TPT1 / P13693
TPX2 / Q9ULW0
TRA2A / Q13595
TRA2B / P62995
TRABD / Q9H4I3
TRADD / Q15628
TRAF2 / Q12933
TRAF3 / Q13114
TRAF6 / Q9Y4K3
TRAF7 / Q6Q0C0
TRAFD1 / O14545
TRAM1 / Q15629
TRAP1 / Q12931
TRAPPC1 / Q9Y5R8
TRAPPC10 / P48553
Gene Symbol / UniProt Accession
Brain
Colon
Duodenum
Esophagus
Heart
Liver
Lung
Pancreas
Thyroid
Organs
Type
DDA
DIA
Bins
1
2
3
4
5

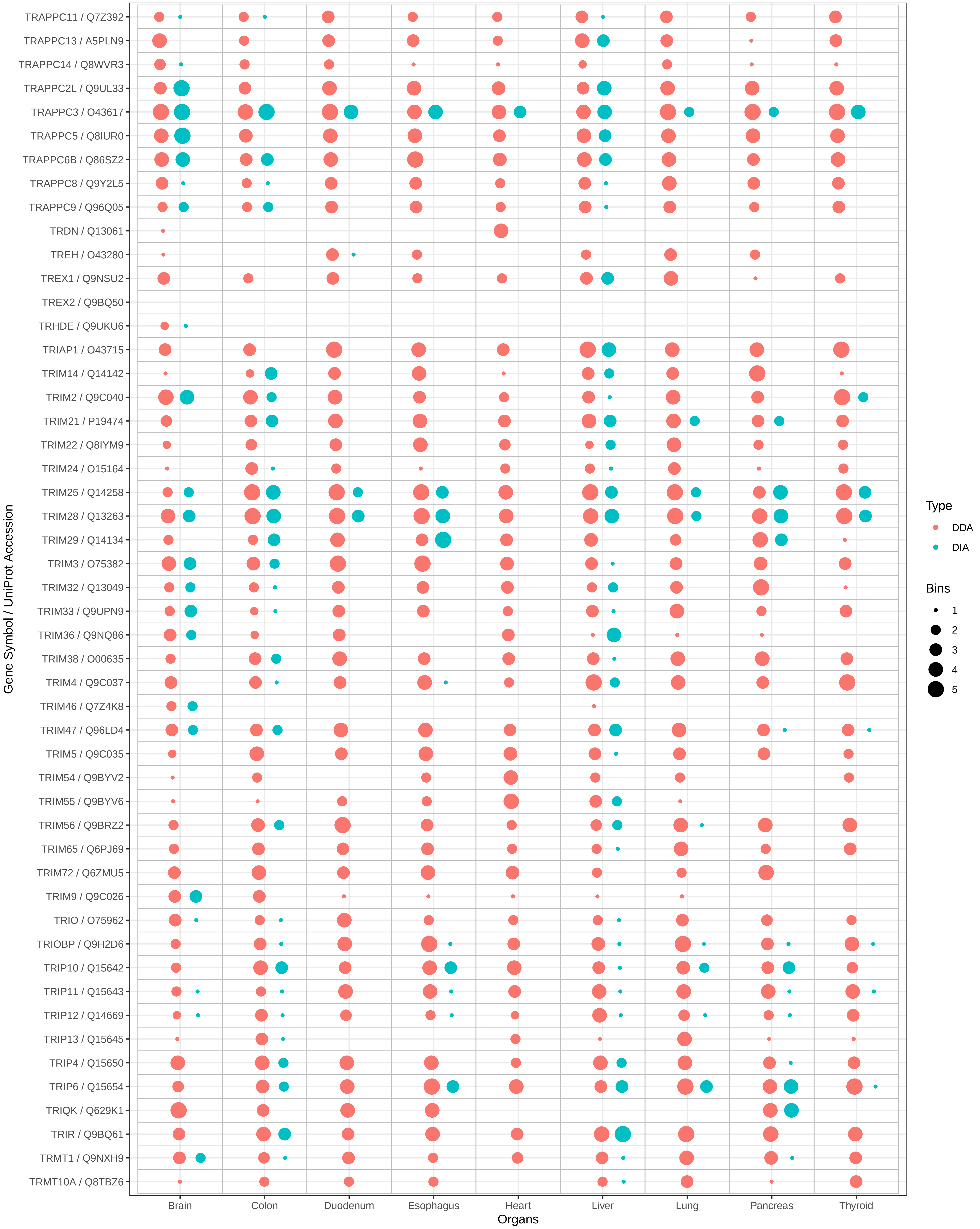
TRAPPC11 / Q7Z392
TRAPPC13 / A5PLN9
TRAPPC14 / Q8WVR3
TRAPPC2L / Q9UL33
TRAPPC3 / O43617
TRAPPC5 / Q8IUR0
TRAPPC6B / Q86SZ2
TRAPPC8 / Q9Y2L5
TRAPPC9 / Q96Q05
TRDN / Q13061
TREH / O43280
TREX1 / Q9NSU2
TREX2 / Q9BQ50
TRHDE / Q9UKU6
TRIAP1 / O43715
TRIM14 / Q14142
TRIM2 / Q9C040
TRIM21 / P19474
TRIM22 / Q8IYM9
TRIM24 / O15164
TRIM25 / Q14258
TRIM28 / Q13263
TRIM29 / Q14134
TRIM3 / O75382
TRIM32 / Q13049
TRIM33 / Q9UPN9
TRIM36 / Q9NQ86
TRIM38 / O00635
TRIM4 / Q9C037
TRIM46 / Q7Z4K8
TRIM47 / Q96LD4
TRIM5 / Q9C035
TRIM54 / Q9BYV2
TRIM55 / Q9BYV6
TRIM56 / Q9BRZ2
TRIM65 / Q6PJ69
TRIM72 / Q6ZMU5
TRIM9 / Q9C026
TRIO / O75962
TRIOBP / Q9H2D6
TRIP10 / Q15642
TRIP11 / Q15643
TRIP12 / Q14669
TRIP13 / Q15645
TRIP4 / Q15650
TRIP6 / Q15654
TRIQK / Q629K1
TRIR / Q9BQ61
TRMT1 / Q9NXH9
TRMT10A / Q8TBZ6
Gene Symbol / UniProt Accession
Brain
Colon
Duodenum
Esophagus
Heart
Liver
Lung
Pancreas
Thyroid
Organs
Type
DDA
DIA
Bins
1
2
3
4
5

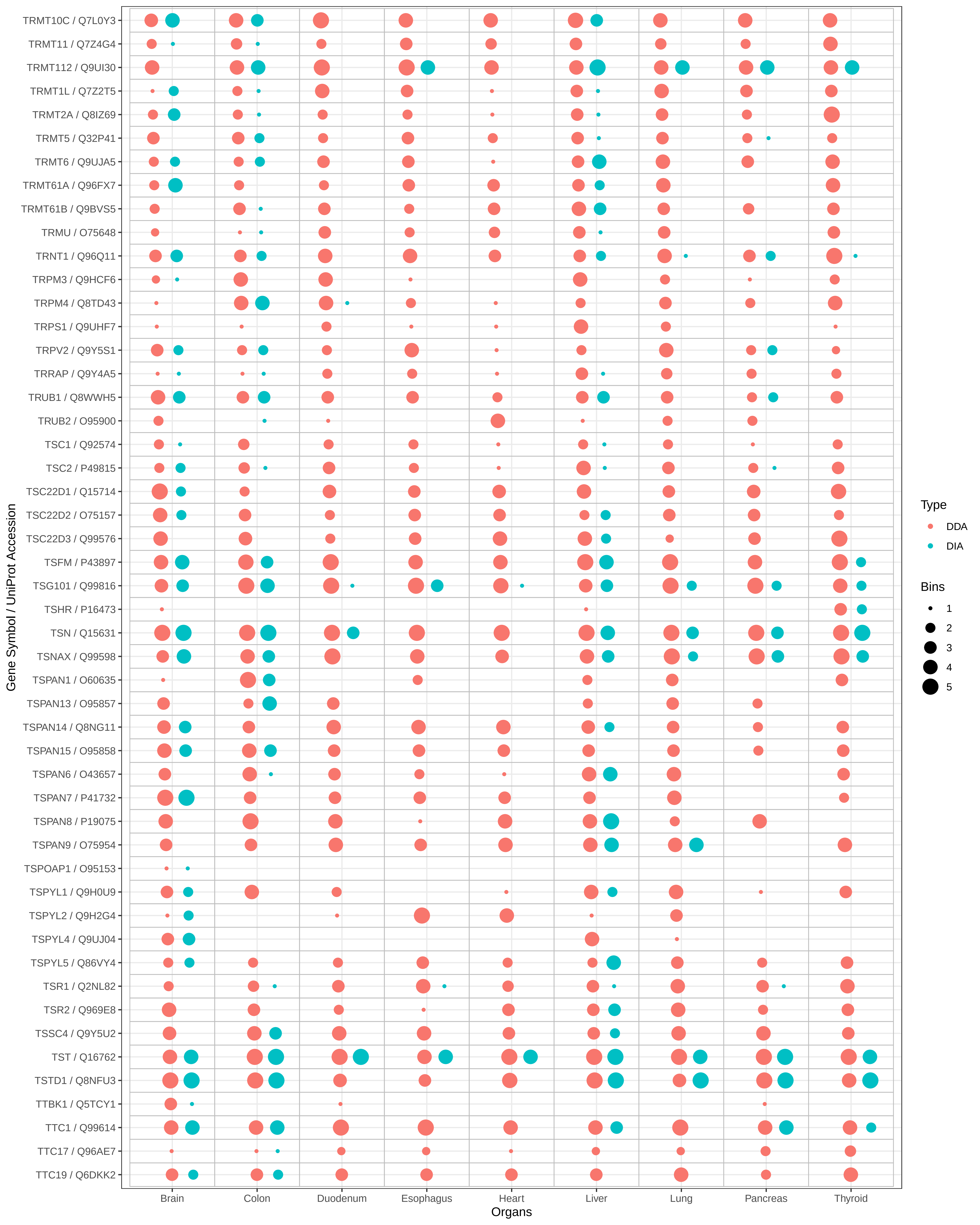
Gene Symbol / UniProt Accession
TRMT10C / Q7L0Y3
TRMT11 / Q7Z4G4
TRMT112 / Q9UI30
TRMT1L / Q7Z2T5
TRMT2A / Q8IZ69
TRMT5 / Q32P41
TRMT6 / Q9UJA5
TRMT61A / Q96FX7
TRMT61B / Q9BVS5
TRMU / O75648
TRNT1 / Q96Q11
TRPM3 / Q9HCF6
TRPM4 / Q8TD43
TRPS1 / Q9UHF7
TRPV2 / Q9Y5S1
TRRAP / Q9Y4A5
TRUB1 / Q8WWH5
TRUB2 / O95900
TSC1 / Q92574
TSC2 / P49815
TSC22D1 / Q15714
TSC22D2 / O75157
TSC22D3 / Q99576
TSFM / P43897
TSG101 / Q99816
TSHR / P16473
TSN / Q15631
TSNAX / Q99598
TSPAN1 / O60635
TSPAN13 / O95857
TSPAN14 / Q8NG11
TSPAN15 / O95858
TSPAN6 / O43657
TSPAN7 / P41732
TSPAN8 / P19075
TSPAN9 / O75954
TSPOAP1 / O95153
TSPYL1 / Q9H0U9
TSPYL2 / Q9H2G4
TSPYL4 / Q9UJ04
TSPYL5 / Q86VY4
TSR1 / Q2NL82
TSR2 / Q969E8
TSSC4 / Q9Y5U2
TST / Q16762
TSTD1 / Q8NFU3
TTBK1 / Q5TCY1
TTC1 / Q99614
TTC17 / Q96AE7
TTC19 / Q6DKK2
Brain
Colon
Duodenum
Esophagus
Heart
Liver
Lung
Pancreas
Thyroid
Organs
Type
DDA
DIA
Bins
1
2
3
4
5

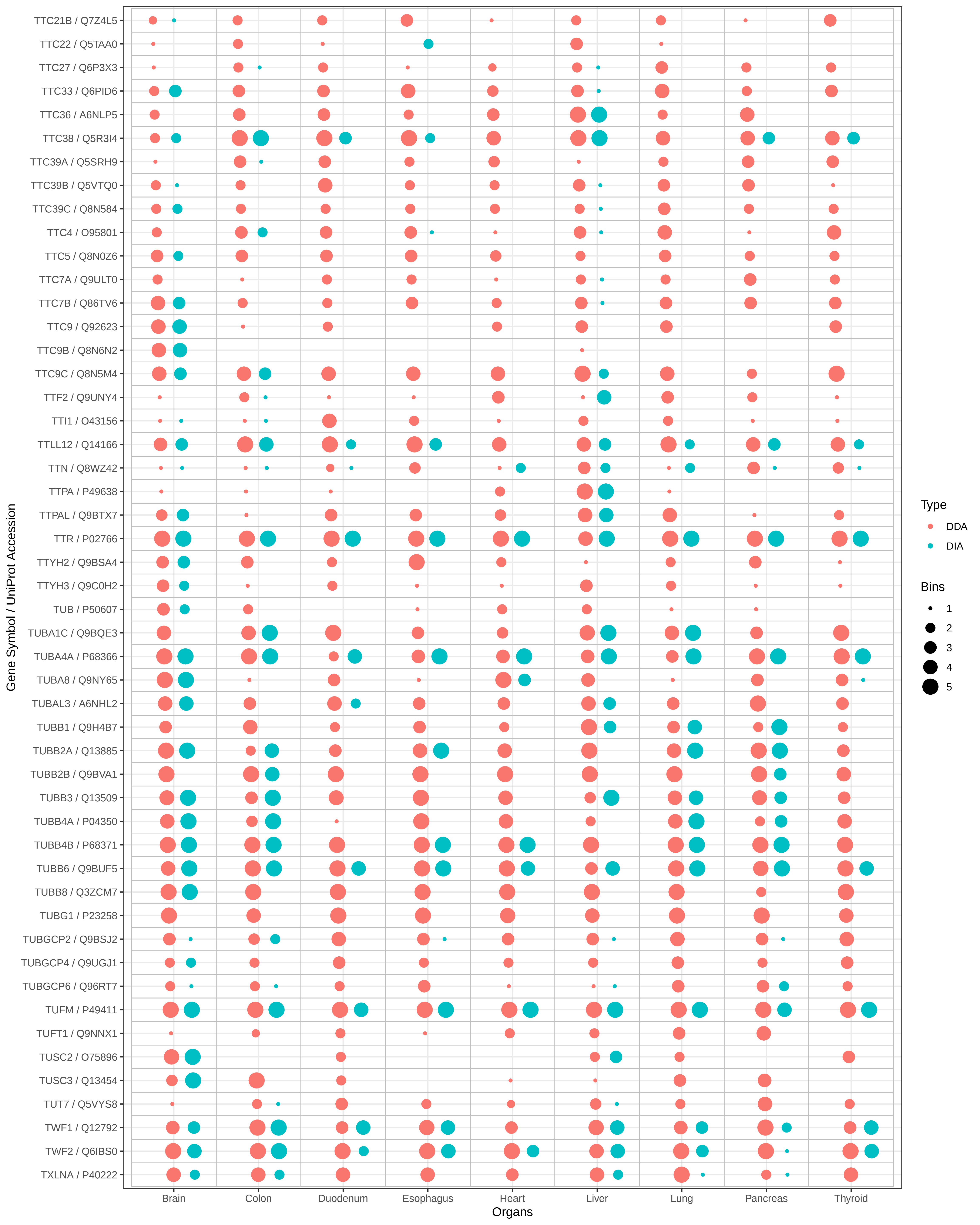

TTC21B / Q7Z4L5
TTC22 / Q5TAA0
TTC27 / Q6P3X3
TTC33 / Q6PID6
TTC36 / A6NLP5
TTC38 / Q5R3I4
TTC39A / Q5SRH9
TTC39B / Q5VTQ0
TTC39C / Q8N584
TTC4 / O95801
TTC5 / Q8N0Z6
TTC7A / Q9ULT0
TTC7B / Q86TV6
TTC9 / Q92623
TTC9B / Q8N6N2
TTC9C / Q8N5M4
TTF2 / Q9UNY4
TTI1 / O43156
TTLL12 / Q14166
TTN / Q8WZ42
TTPA / P49638
TTPAL / Q9BTX7
TTR / P02766
TTYH2 / Q9BSA4
TTYH3 / Q9C0H2
TUB / P50607
TUBA1C / Q9BQE3
TUBA4A / P68366
TUBA8 / Q9NY65
TUBAL3 / A6NHL2
TUBB1 / Q9H4B7
TUBB2A / Q13885
TUBB2B / Q9BVA1
TUBB3 / Q13509
TUBB4A / P04350
TUBB4B / P68371
TUBB6 / Q9BUF5
TUBB8 / Q3ZCM7
TUBG1 / P23258
TUBGCP2 / Q9BSJ2
TUBGCP4 / Q9UGJ1
TUBGCP6 / Q96RT7
TUFM / P49411
TUFT1 / Q9NNX1
TUSC2 / O75896
TUSC3 / Q13454
TUT7 / Q5VYS8
TWF1 / Q12792
TWF2 / Q6IBS0
TXLNA / P40222
Gene Symbol / UniProt Accession
Brain
Colon
Duodenum
Esophagus
Heart
Liver
Lung
Pancreas
Thyroid
Organs
Type
DDA
DIA
Bins
1
2
3
4
5

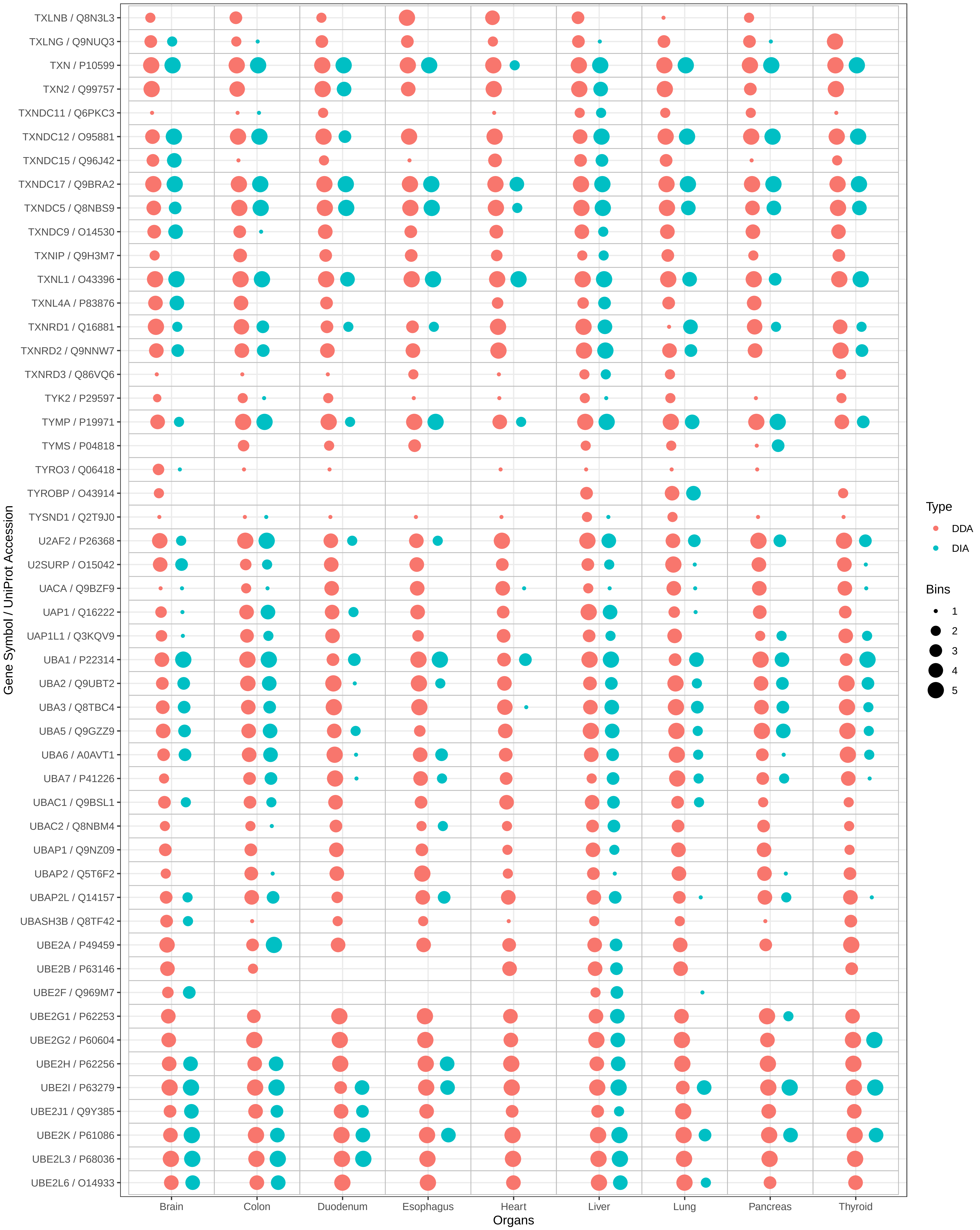

TXLNB / Q8N3L3
TXLNG / Q9NUQ3
TXN / P10599
TXN2 / Q99757
TXNDC11 / Q6PKC3
TXNDC12 / O95881
TXNDC15 / Q96J42
TXNDC17 / Q9BRA2
TXNDC5 / Q8NBS9
TXNDC9 / O14530
TXNIP / Q9H3M7
TXNL1 / O43396
TXNL4A / P83876
TXNRD1 / Q16881
TXNRD2 / Q9NNW7
TXNRD3 / Q86VQ6
TYK2 / P29597
TYMP / P19971
TYMS / P04818
TYRO3 / Q06418
TYROBP / O43914
TYSND1 / Q2T9J0
U2AF2 / P26368
U2SURP / O15042
UACA / Q9BZF9
UAP1 / Q16222
UAP1L1 / Q3KQV9
UBA1 / P22314
UBA2 / Q9UBT2
UBA3 / Q8TBC4
UBA5 / Q9GZZ9
UBA6 / A0AVT1
UBA7 / P41226
UBAC1 / Q9BSL1
UBAC2 / Q8NBM4
UBAP1 / Q9NZ09
UBAP2 / Q5T6F2
UBAP2L / Q14157
UBASH3B / Q8TF42
UBE2A / P49459
UBE2B / P63146
UBE2F / Q969M7
UBE2G1 / P62253
UBE2G2 / P60604
UBE2H / P62256
UBE2I / P63279
UBE2J1 / Q9Y385
UBE2K / P61086
UBE2L3 / P68036
UBE2L6 / O14933
Gene Symbol / UniProt Accession
Brain
Colon
Duodenum
Esophagus
Heart
Liver
Lung
Pancreas
Thyroid
Organs
Type
DDA
DIA
Bins
1
2
3
4
5

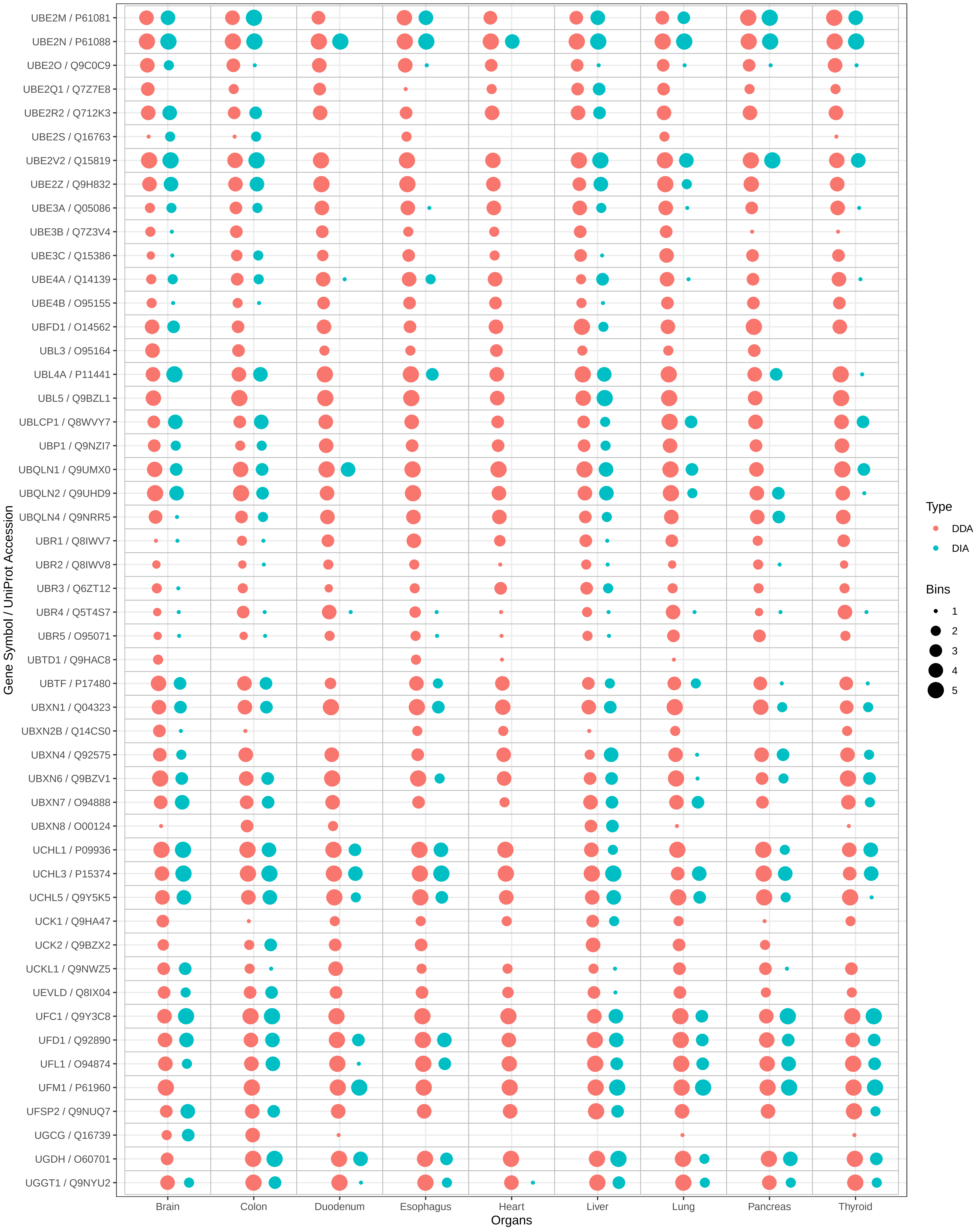

UBE2M / P61081
UBE2N / P61088
UBE2O / Q9C0C9
UBE2Q1 / Q7Z7E8
UBE2R2 / Q712K3
UBE2S / Q16763
UBE2V2 / Q15819
UBE2Z / Q9H832
UBE3A / Q05086
UBE3B / Q7Z3V4
UBE3C / Q15386
UBE4A / Q14139
UBE4B / O95155
UBFD1 / O14562
UBL3 / O95164
UBL4A / P11441
UBL5 / Q9BZL1
UBLCP1 / Q8WVY7
UBP1 / Q9NZI7
UBQLN1 / Q9UMX0
UBQLN2 / Q9UHD9
UBQLN4 / Q9NRR5
UBR1 / Q8IWV7
UBR2 / Q8IWV8
UBR3 / Q6ZT12
UBR4 / Q5T4S7
UBR5 / O95071
UBTD1 / Q9HAC8
UBTF / P17480
UBXN1 / Q04323
UBXN2B / Q14CS0
UBXN4 / Q92575
UBXN6 / Q9BZV1
UBXN7 / O94888
UBXN8 / O00124
UCHL1 / P09936
UCHL3 / P15374
UCHL5 / Q9Y5K5
UCK1 / Q9HA47
UCK2 / Q9BZX2
UCKL1 / Q9NWZ5
UEVLD / Q8IX04
UFC1 / Q9Y3C8
UFD1 / Q92890
UFL1 / O94874
UFM1 / P61960
UFSP2 / Q9NUQ7
UGCG / Q16739
UGDH / O60701
UGGT1 / Q9NYU2
Gene Symbol / UniProt Accession
Brain
Colon
Duodenum
Esophagus
Heart
Liver
Lung
Pancreas
Thyroid
Organs
Type
DDA
DIA
Bins
1
2
3
4
5

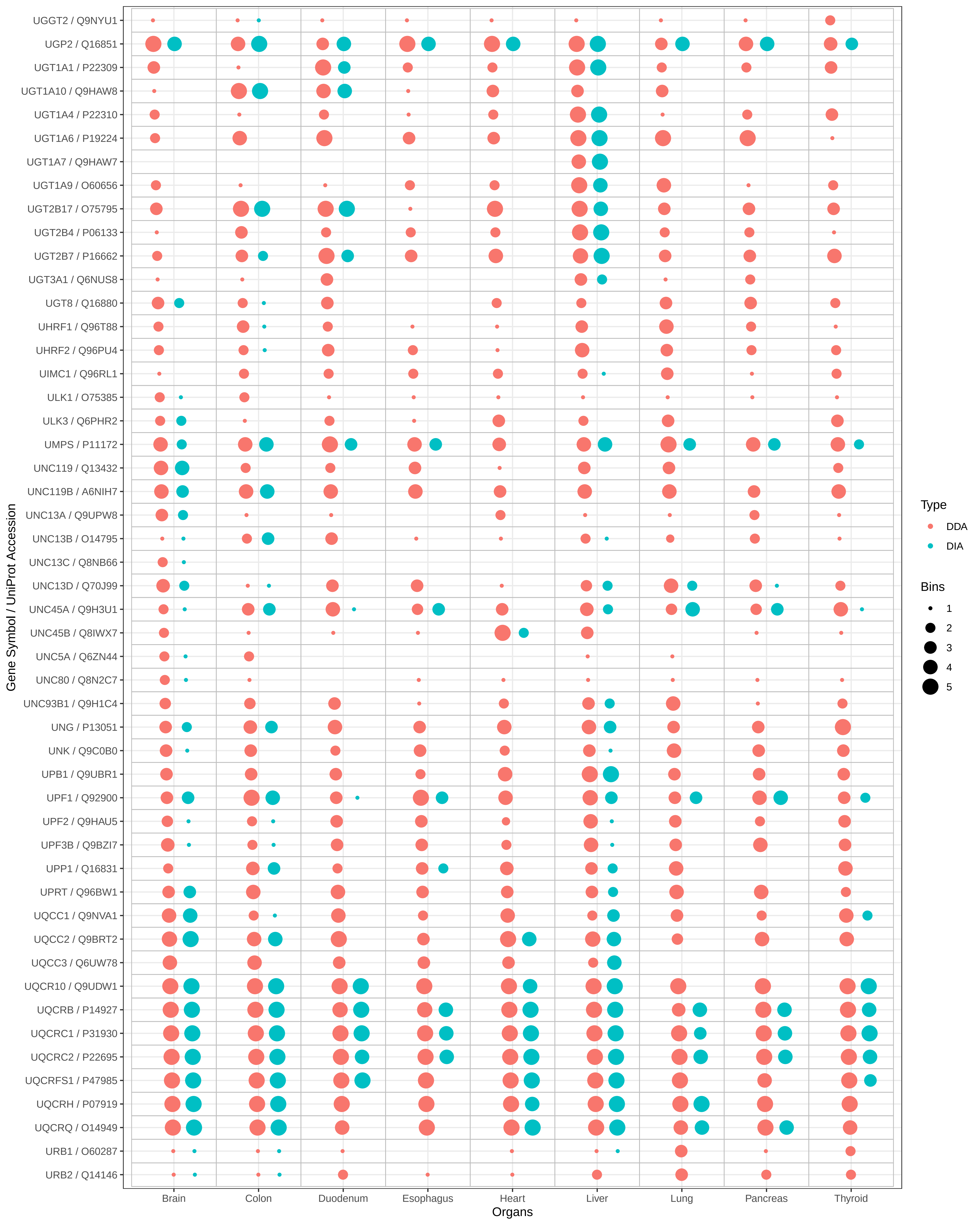

UGGT2 / Q9NYU1
UGP2 / Q16851
UGT1A1 / P22309
UGT1A10 / Q9HAW8
UGT1A4 / P22310
UGT1A6 / P19224
UGT1A7 / Q9HAW7
UGT1A9 / O60656
UGT2B17 / O75795
UGT2B4 / P06133
UGT2B7 / P16662
UGT3A1 / Q6NUS8
UGT8 / Q16880
UHRF1 / Q96T88
UHRF2 / Q96PU4
UIMC1 / Q96RL1
ULK1 / O75385
ULK3 / Q6PHR2
UMPS / P11172
UNC119 / Q13432
UNC119B / A6NIH7
UNC13A / Q9UPW8
UNC13B / O14795
UNC13C / Q8NB66
UNC13D / Q70J99
UNC45A / Q9H3U1
UNC45B / Q8IWX7
UNC5A / Q6ZN44
UNC80 / Q8N2C7
UNC93B1 / Q9H1C4
UNG / P13051
UNK / Q9C0B0
UPB1 / Q9UBR1
UPF1 / Q92900
UPF2 / Q9HAU5
UPF3B / Q9BZI7
UPP1 / Q16831
UPRT / Q96BW1
UQCC1 / Q9NVA1
UQCC2 / Q9BRT2
UQCC3 / Q6UW78
UQCR10 / Q9UDW1
UQCRB / P14927
UQCRC1 / P31930
UQCRC2 / P22695
UQCRFS1 / P47985
UQCRH / P07919
UQCRQ / O14949
URB1 / O60287
URB2 / Q14146
Gene Symbol / UniProt Accession
Brain
Colon
Duodenum
Esophagus
Heart
Liver
Lung
Pancreas
Thyroid
Organs
Type
DDA
DIA
Bins
1
2
3
4
5

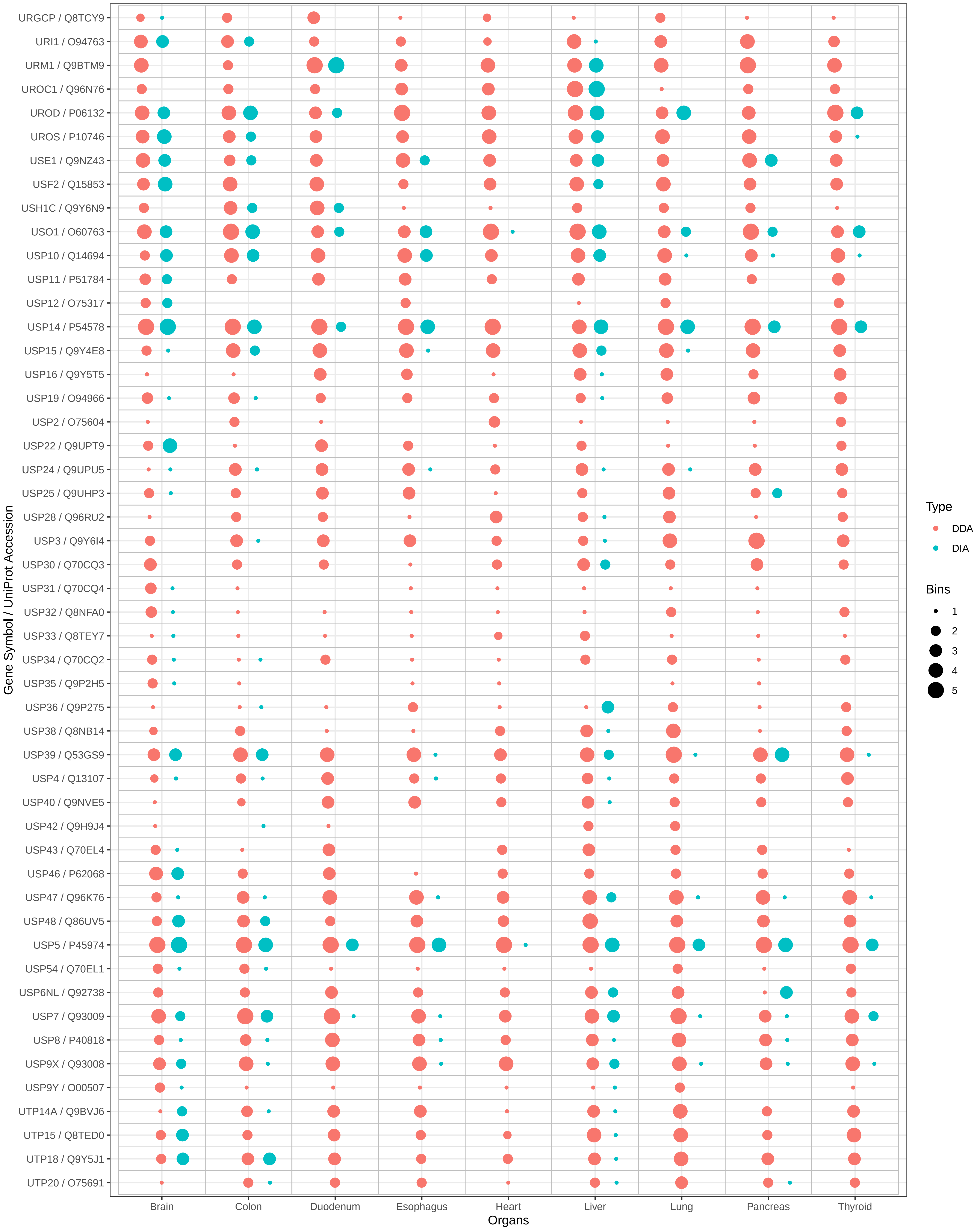
URGCP / Q8TCY9
URI1 / O94763
URM1 / Q9BTM9
UROC1 / Q96N76
UROD / P06132
UROS / P10746
USE1 / Q9NZ43
USF2 / Q15853
USH1C / Q9Y6N9
USO1 / O60763
USP10 / Q14694
USP11 / P51784
USP12 / O75317
USP14 / P54578
USP15 / Q9Y4E8
USP16 / Q9Y5T5
USP19 / O94966
USP2 / O75604
USP22 / Q9UPT9
USP24 / Q9UPU5
USP25 / Q9UHP3
USP28 / Q96RU2
USP3 / Q9Y6I4
USP30 / Q70CQ3
USP31 / Q70CQ4
USP32 / Q8NFA0
USP33 / Q8TEY7
USP34 / Q70CQ2
USP35 / Q9P2H5
USP36 / Q9P275
USP38 / Q8NB14
USP39 / Q53GS9
USP4 / Q13107
USP40 / Q9NVE5
USP42 / Q9H9J4
USP43 / Q70EL4
USP46 / P62068
USP47 / Q96K76
USP48 / Q86UV5
USP5 / P45974
USP54 / Q70EL1
USP6NL / Q92738
USP7 / Q93009
USP8 / P40818
USP9X / Q93008
USP9Y / O00507
UTP14A / Q9BVJ6
UTP15 / Q8TED0
UTP18 / Q9Y5J1
UTP20 / O75691
Gene Symbol / UniProt Accession
Brain
Colon
Duodenum
Esophagus
Heart
Liver
Lung
Pancreas
Thyroid
Organs
Type
DDA
DIA
Bins
1
2
3
4
5

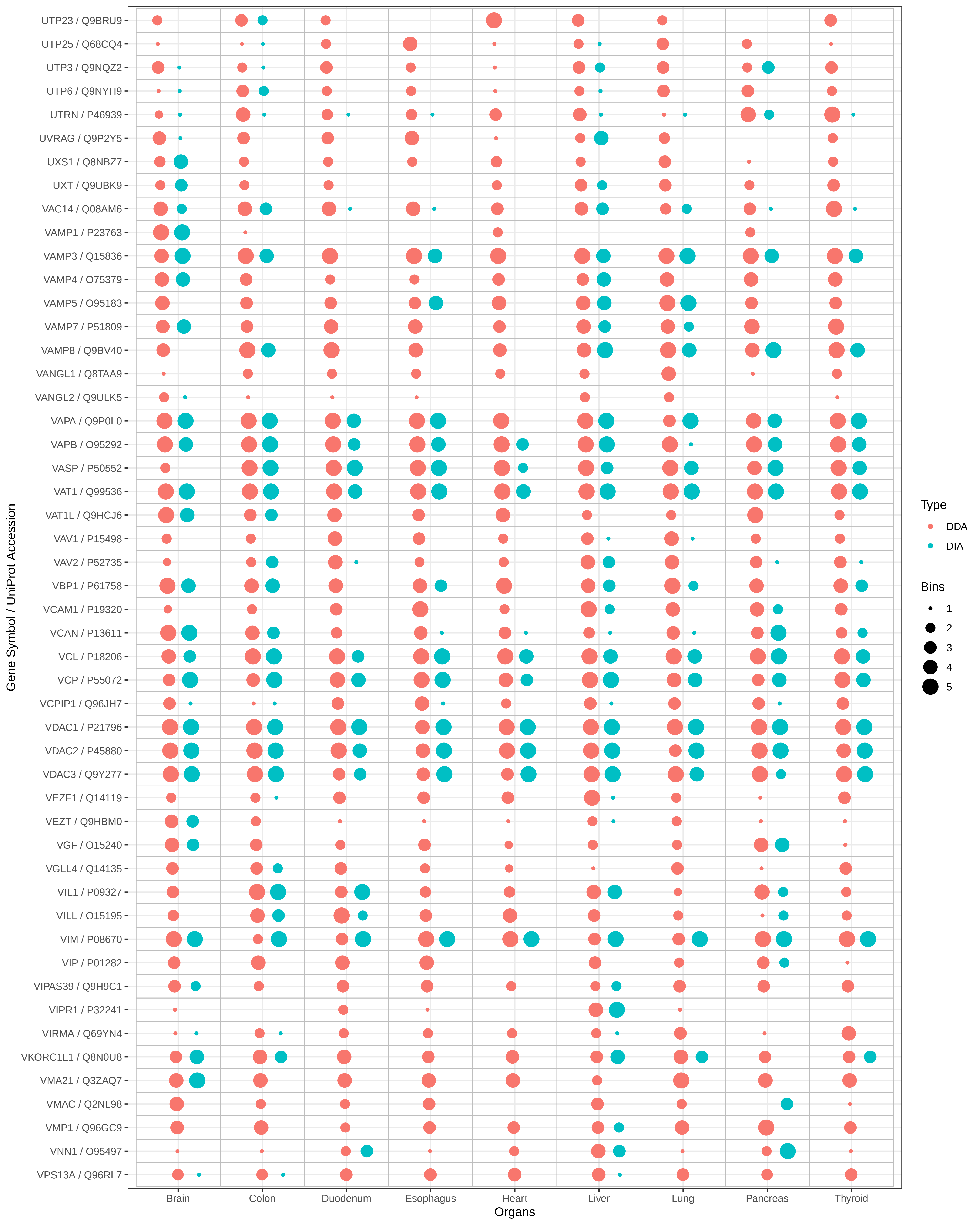
Gene Symbol / UniProt Accession
UTP23 / Q9BRU9
UTP25 / Q68CQ4
UTP3 / Q9NQZ2
UTP6 / Q9NYH9
UTRN / P46939
UVRAG / Q9P2Y5
UXS1 / Q8NBZ7
UXT / Q9UBK9
VAC14 / Q08AM6
VAMP1 / P23763
VAMP3 / Q15836
VAMP4 / O75379
VAMP5 / O95183
VAMP7 / P51809
VAMP8 / Q9BV40
VANGL1 / Q8TAA9
VANGL2 / Q9ULK5
VAPA / Q9P0L0
VAPB / O95292
VASP / P50552
VAT1 / Q99536
VAT1L / Q9HCJ6
VAV1 / P15498
VAV2 / P52735
VBP1 / P61758
VCAM1 / P19320
VCAN / P13611
VCL / P18206
VCP / P55072
VCPIP1 / Q96JH7
VDAC1 / P21796
VDAC2 / P45880
VDAC3 / Q9Y277
VEZF1 / Q14119
VEZT / Q9HBM0
VGF / O15240
VGLL4 / Q14135
VIL1 / P09327
VILL / O15195
VIM / P08670
VIP / P01282
VIPAS39 / Q9H9C1
VIPR1 / P32241
VIRMA / Q69YN4
VKORC1L1 / Q8N0U8
VMA21 / Q3ZAQ7
VMAC / Q2NL98
VMP1 / Q96GC9
VNN1 / O95497
VPS13A / Q96RL7
Brain
Colon
Duodenum
Esophagus
Heart
Liver
Lung
Pancreas
Thyroid
Organs
Type
DDA
DIA
Bins
1
2
3
4
5

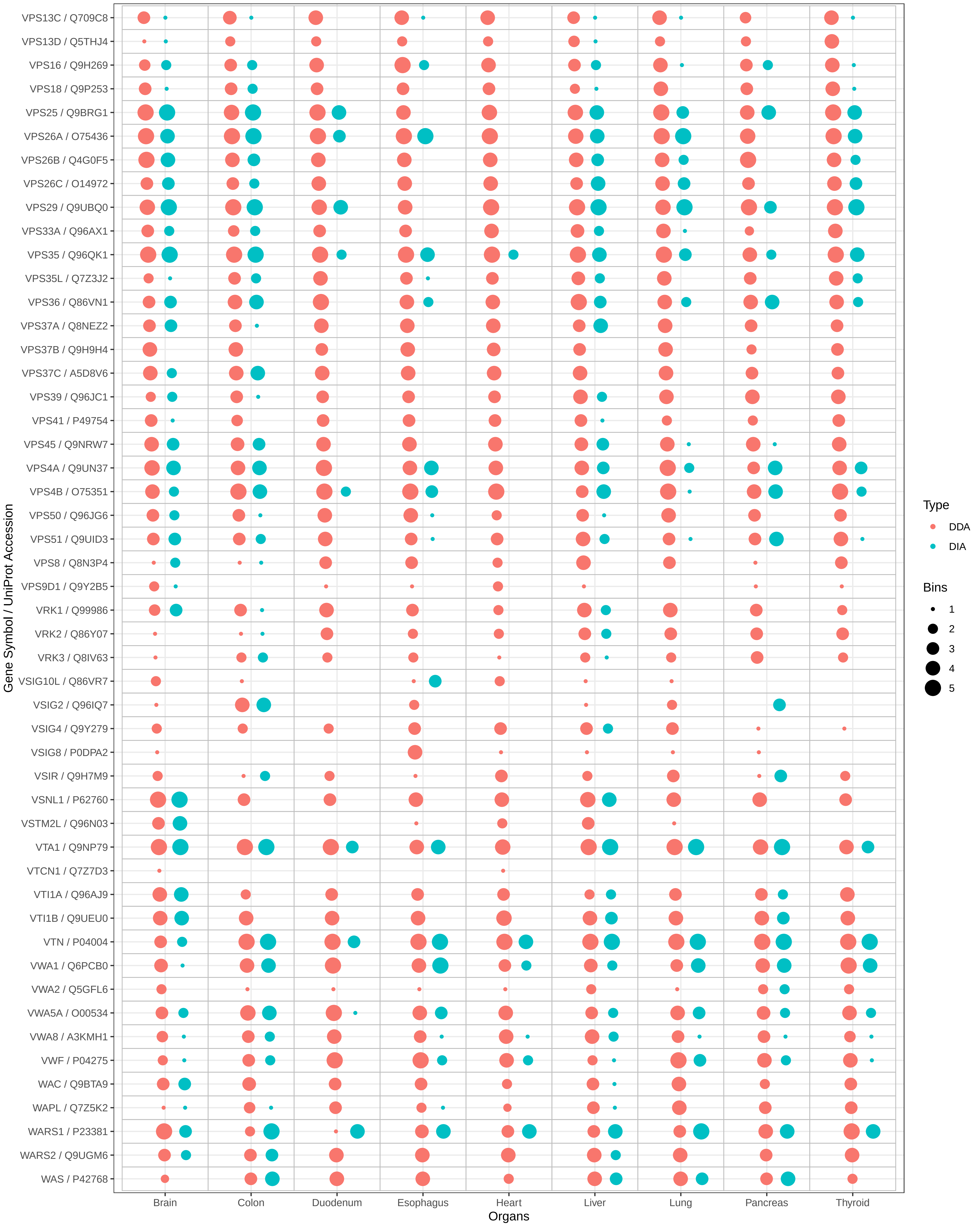

VPS13C / Q709C8
VPS13D / Q5THJ4
VPS16 / Q9H269
VPS18 / Q9P253
VPS25 / Q9BRG1
VPS26A / O75436
VPS26B / Q4G0F5
VPS26C / O14972
VPS29 / Q9UBQ0
VPS33A / Q96AX1
VPS35 / Q96QK1
VPS35L / Q7Z3J2
VPS36 / Q86VN1
VPS37A / Q8NEZ2
VPS37B / Q9H9H4
VPS37C / A5D8V6
VPS39 / Q96JC1
VPS41 / P49754
VPS45 / Q9NRW7
VPS4A / Q9UN37
VPS4B / O75351
VPS50 / Q96JG6
VPS51 / Q9UID3
VPS8 / Q8N3P4
VPS9D1 / Q9Y2B5
VRK1 / Q99986
VRK2 / Q86Y07
VRK3 / Q8IV63
VSIG10L / Q86VR7
VSIG2 / Q96IQ7
VSIG4 / Q9Y279
VSIG8 / P0DPA2
VSIR / Q9H7M9
VSNL1 / P62760
VSTM2L / Q96N03
VTA1 / Q9NP79
VTCN1 / Q7Z7D3
VTI1A / Q96AJ9
VTI1B / Q9UEU0
VTN / P04004
VWA1 / Q6PCB0
VWA2 / Q5GFL6
VWA5A / O00534
VWA8 / A3KMH1
VWF / P04275
WAC / Q9BTA9
WAPL / Q7Z5K2
WARS1 / P23381
WARS2 / Q9UGM6
WAS / P42768
Gene Symbol / UniProt Accession
Brain
Colon
Duodenum
Esophagus
Heart
Liver
Lung
Pancreas
Thyroid
Organs
Type
DDA
DIA
Bins
1
2
3
4
5

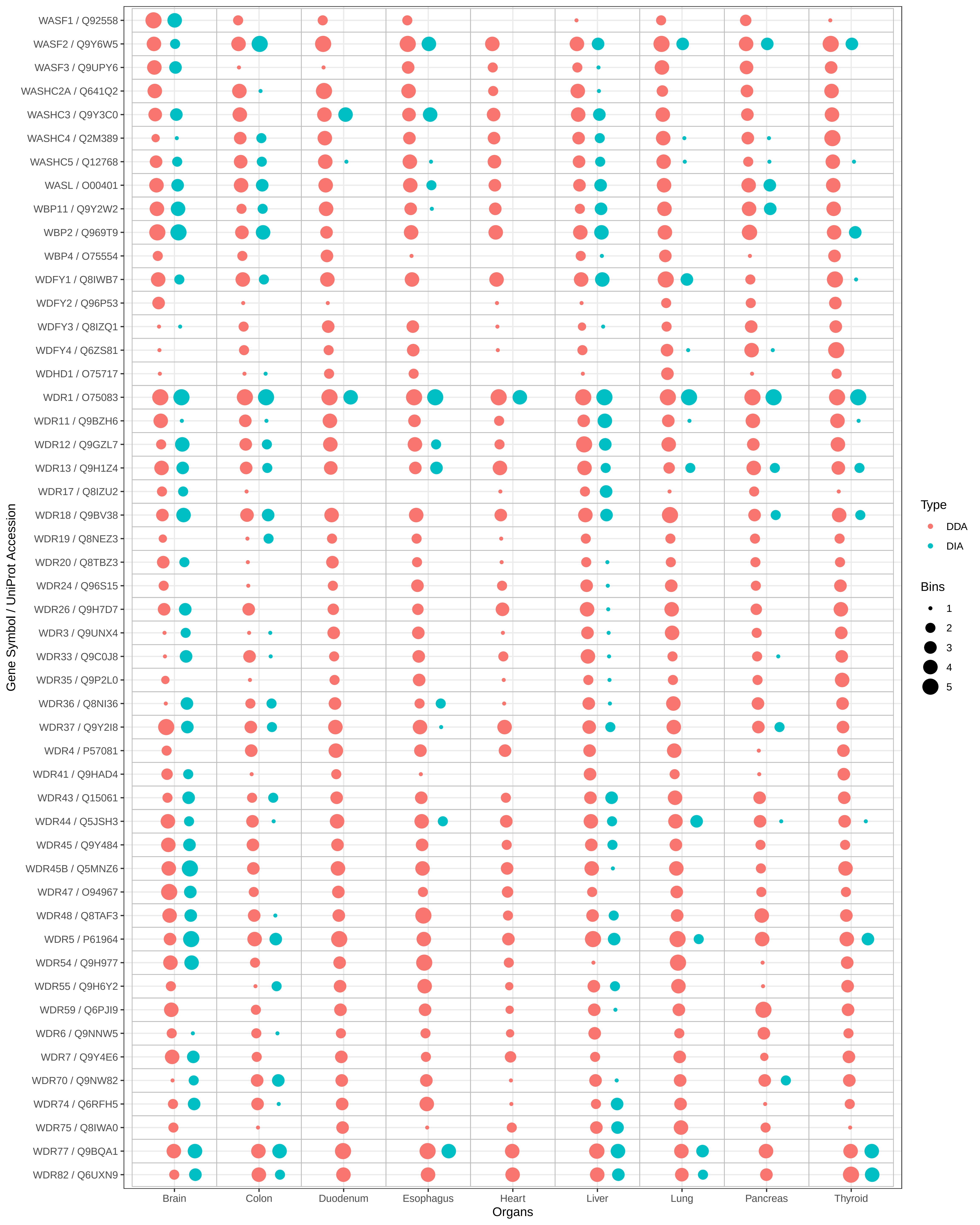

WASF1 / Q92558
WASF2 / Q9Y6W5
WASF3 / Q9UPY6
WASHC2A / Q641Q2
WASHC3 / Q9Y3C0
WASHC4 / Q2M389
WASHC5 / Q12768
WASL / O00401
WBP11 / Q9Y2W2
WBP2 / Q969T9
WBP4 / O75554
WDFY1 / Q8IWB7
WDFY2 / Q96P53
WDFY3 / Q8IZQ1
WDFY4 / Q6ZS81
WDHD1 / O75717
WDR1 / O75083
WDR11 / Q9BZH6
WDR12 / Q9GZL7
WDR13 / Q9H1Z4
WDR17 / Q8IZU2
WDR18 / Q9BV38
WDR19 / Q8NEZ3
WDR20 / Q8TBZ3
WDR24 / Q96S15
WDR26 / Q9H7D7
WDR3 / Q9UNX4
WDR33 / Q9C0J8
WDR35 / Q9P2L0
WDR36 / Q8NI36
WDR37 / Q9Y2I8
WDR4 / P57081
WDR41 / Q9HAD4
WDR43 / Q15061
WDR44 / Q5JSH3
WDR45 / Q9Y484
WDR45B / Q5MNZ6
WDR47 / O94967
WDR48 / Q8TAF3
WDR5 / P61964
WDR54 / Q9H977
WDR55 / Q9H6Y2
WDR59 / Q6PJI9
WDR6 / Q9NNW5
WDR7 / Q9Y4E6
WDR70 / Q9NW82
WDR74 / Q6RFH5
WDR75 / Q8IWA0
WDR77 / Q9BQA1
WDR82 / Q6UXN9
Gene Symbol / UniProt Accession
Brain
Colon
Duodenum
Esophagus
Heart
Liver
Lung
Pancreas
Thyroid
Organs
Type
DDA
DIA
Bins
1
2
3
4
5

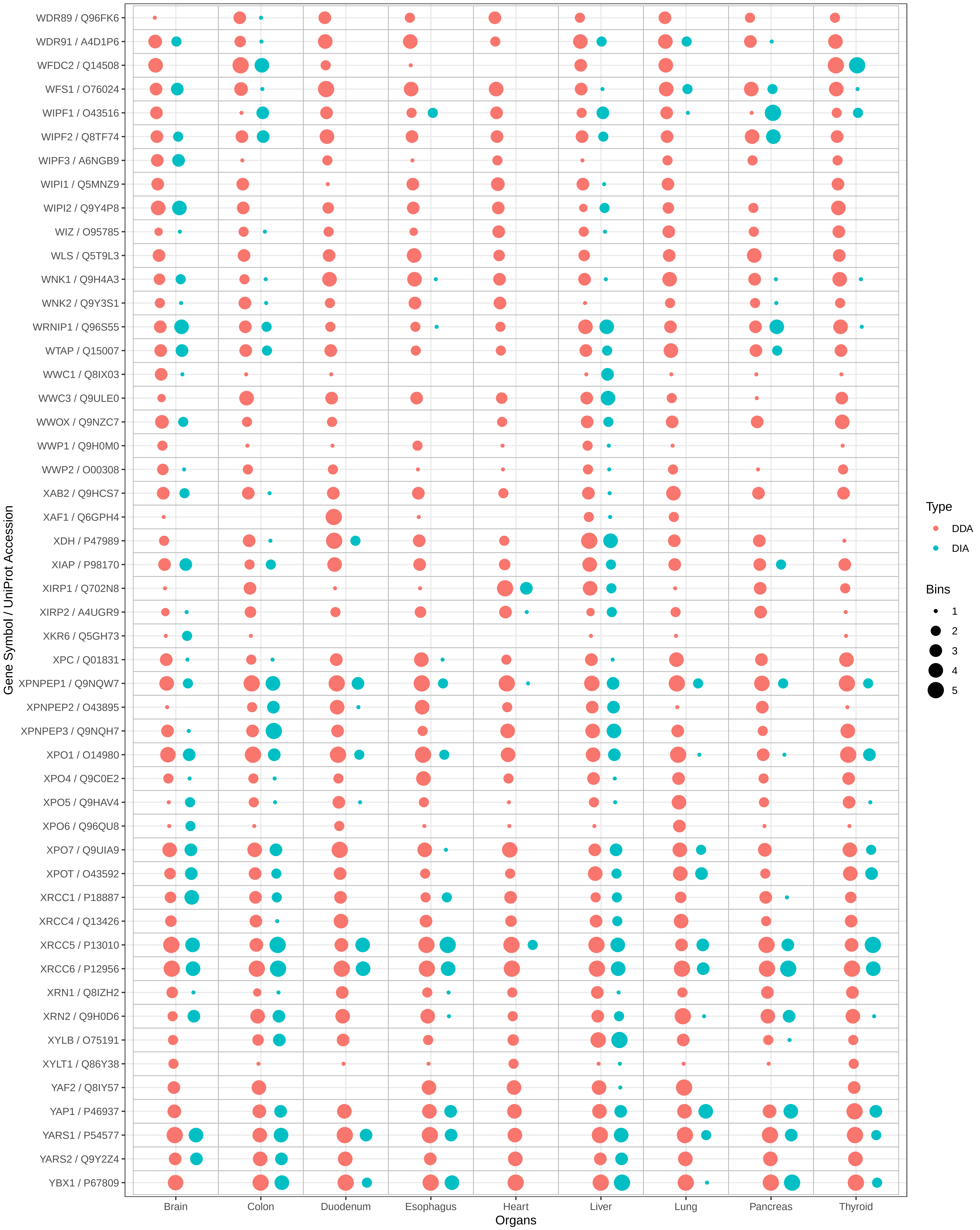

WDR89 / Q96FK6
WDR91 / A4D1P6
WFDC2 / Q14508
WFS1 / O76024
WIPF1 / O43516
WIPF2 / Q8TF74
WIPF3 / A6NGB9
WIPI1 / Q5MNZ9
WIPI2 / Q9Y4P8
WIZ / O95785
WLS / Q5T9L3
WNK1 / Q9H4A3
WNK2 / Q9Y3S1
WRNIP1 / Q96S55
WTAP / Q15007
WWC1 / Q8IX03
WWC3 / Q9ULE0
WWOX / Q9NZC7
WWP1 / Q9H0M0
WWP2 / O00308
XAB2 / Q9HCS7
XAF1 / Q6GPH4
XDH / P47989
XIAP / P98170
XIRP1 / Q702N8
XIRP2 / A4UGR9
XKR6 / Q5GH73
XPC / Q01831
XPNPEP1 / Q9NQW7
XPNPEP2 / O43895
XPNPEP3 / Q9NQH7
XPO1 / O14980
XPO4 / Q9C0E2
XPO5 / Q9HAV4
XPO6 / Q96QU8
XPO7 / Q9UIA9
XPOT / O43592
XRCC1 / P18887
XRCC4 / Q13426
XRCC5 / P13010
XRCC6 / P12956
XRN1 / Q8IZH2
XRN2 / Q9H0D6
XYLB / O75191
XYLT1 / Q86Y38
YAF2 / Q8IY57
YAP1 / P46937
YARS1 / P54577
YARS2 / Q9Y2Z4
YBX1 / P67809
Gene Symbol / UniProt Accession
Brain
Colon
Duodenum
Esophagus
Heart
Liver
Lung
Pancreas
Thyroid
Organs
Type
DDA
DIA
Bins
1
2
3
4
5

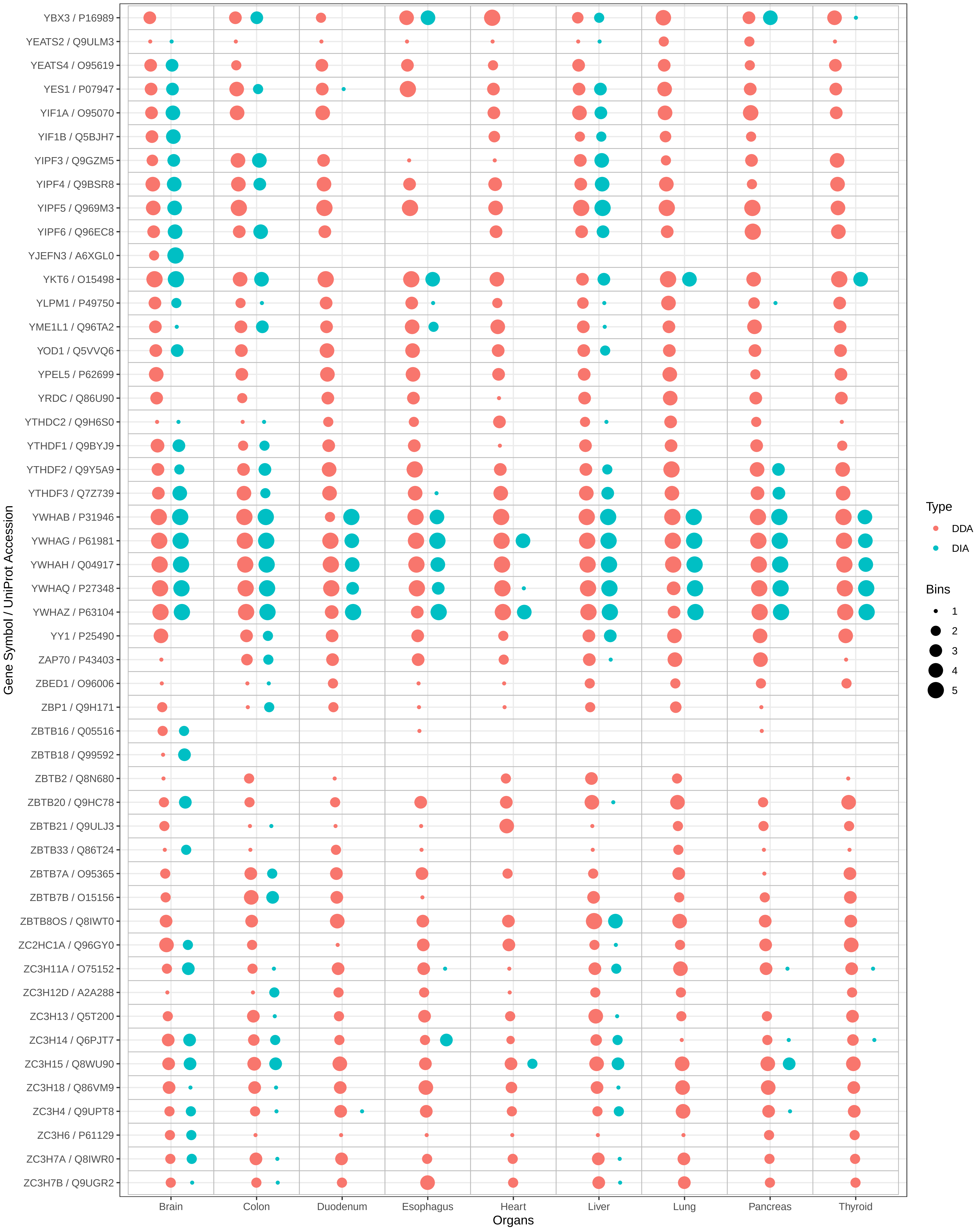

YBX3 / P16989
YEATS2 / Q9ULM3
YEATS4 / O95619
YES1 / P07947
YIF1A / O95070
YIF1B / Q5BJH7
YIPF3 / Q9GZM5
YIPF4 / Q9BSR8
YIPF5 / Q969M3
YIPF6 / Q96EC8
YJEFN3 / A6XGL0
YKT6 / O15498
YLPM1 / P49750
YME1L1 / Q96TA2
YOD1 / Q5VVQ6
YPEL5 / P62699
YRDC / Q86U90
YTHDC2 / Q9H6S0
YTHDF1 / Q9BYJ9
YTHDF2 / Q9Y5A9
YTHDF3 / Q7Z739
YWHAB / P31946
YWHAG / P61981
YWHAH / Q04917
YWHAQ / P27348
YWHAZ / P63104
YY1 / P25490
ZAP70 / P43403
ZBED1 / O96006
ZBP1 / Q9H171
ZBTB16 / Q05516
ZBTB18 / Q99592
ZBTB2 / Q8N680
ZBTB20 / Q9HC78
ZBTB21 / Q9ULJ3
ZBTB33 / Q86T24
ZBTB7A / O95365
ZBTB7B / O15156
ZBTB8OS / Q8IWT0
ZC2HC1A / Q96GY0
ZC3H11A / O75152
ZC3H12D / A2A288
ZC3H13 / Q5T200
ZC3H14 / Q6PJT7
ZC3H15 / Q8WU90
ZC3H18 / Q86VM9
ZC3H4 / Q9UPT8
ZC3H6 / P61129
ZC3H7A / Q8IWR0
ZC3H7B / Q9UGR2
Gene Symbol / UniProt Accession
Brain
Colon
Duodenum
Esophagus
Heart
Liver
Lung
Pancreas
Thyroid
Organs
Type
DDA
DIA
Bins
1
2
3
4
5

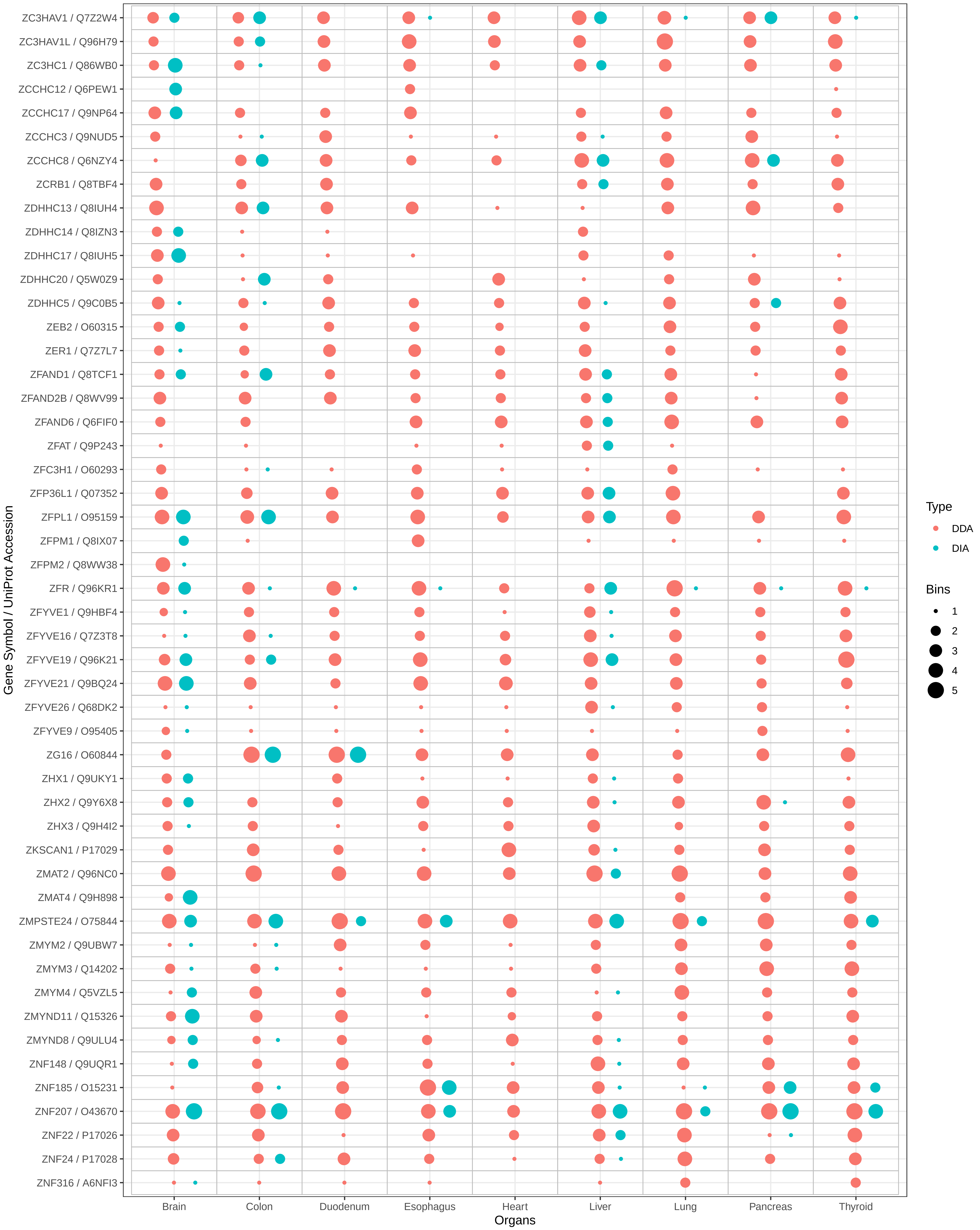

ZC3HAV1 / Q7Z2W4
ZC3HAV1L / Q96H79
ZC3HC1 / Q86WB0
ZCCHC12 / Q6PEW1
ZCCHC17 / Q9NP64
ZCCHC3 / Q9NUD5
ZCCHC8 / Q6NZY4
ZCRB1 / Q8TBF4
ZDHHC13 / Q8IUH4
ZDHHC14 / Q8IZN3
ZDHHC17 / Q8IUH5
ZDHHC20 / Q5W0Z9
ZDHHC5 / Q9C0B5
ZEB2 / O60315
ZER1 / Q7Z7L7
ZFAND1 / Q8TCF1
ZFAND2B / Q8WV99
ZFAND6 / Q6FIF0
ZFAT / Q9P243
ZFC3H1 / O60293
ZFP36L1 / Q07352
ZFPL1 / O95159
ZFPM1 / Q8IX07
ZFPM2 / Q8WW38
ZFR / Q96KR1
ZFYVE1 / Q9HBF4
ZFYVE16 / Q7Z3T8
ZFYVE19 / Q96K21
ZFYVE21 / Q9BQ24
ZFYVE26 / Q68DK2
ZFYVE9 / O95405
ZG16 / O60844
ZHX1 / Q9UKY1
ZHX2 / Q9Y6X8
ZHX3 / Q9H4I2
ZKSCAN1 / P17029
ZMAT2 / Q96NC0
ZMAT4 / Q9H898
ZMPSTE24 / O75844
ZMYM2 / Q9UBW7
ZMYM3 / Q14202
ZMYM4 / Q5VZL5
ZMYND11 / Q15326
ZMYND8 / Q9ULU4
ZNF148 / Q9UQR1
ZNF185 / O15231
ZNF207 / O43670
ZNF22 / P17026
ZNF24 / P17028
ZNF316 / A6NFI3
Gene Symbol / UniProt Accession
Brain
Colon
Duodenum
Esophagus
Heart
Liver
Lung
Pancreas
Thyroid
Organs
Type
DDA
DIA
Bins
1
2
3
4
5

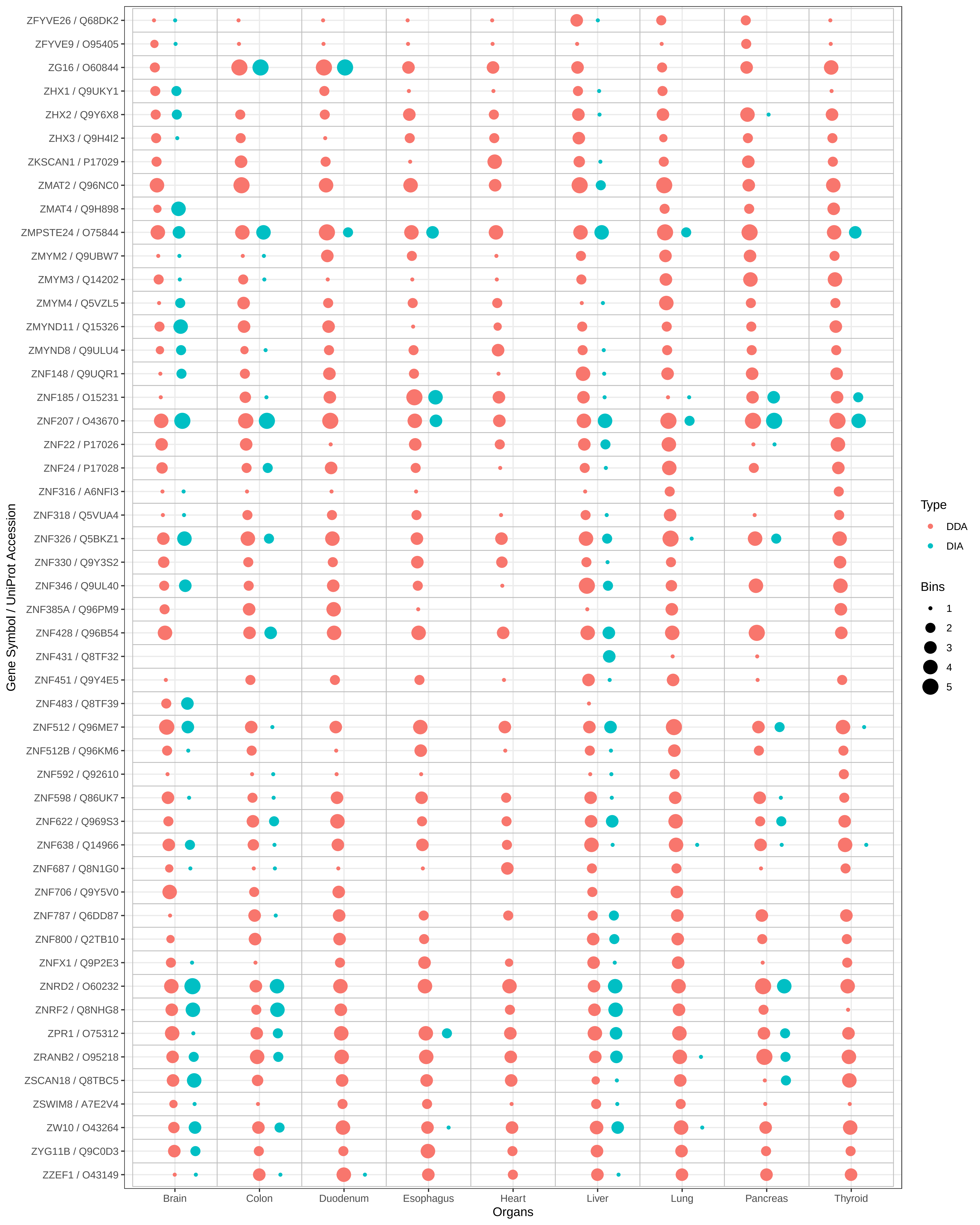

ZFYVE26 / Q68DK2
ZFYVE9 / O95405
ZG16 / O60844
ZHX1 / Q9UKY1
ZHX2 / Q9Y6X8
ZHX3 / Q9H4I2
ZKSCAN1 / P17029
ZMAT2 / Q96NC0
ZMAT4 / Q9H898
ZMPSTE24 / O75844
ZMYM2 / Q9UBW7
ZMYM3 / Q14202
ZMYM4 / Q5VZL5
ZMYND11 / Q15326
ZMYND8 / Q9ULU4
ZNF148 / Q9UQR1
ZNF185 / O15231
ZNF207 / O43670
ZNF22 / P17026
ZNF24 / P17028
ZNF316 / A6NFI3
ZNF318 / Q5VUA4
ZNF326 / Q5BKZ1
ZNF330 / Q9Y3S2
ZNF346 / Q9UL40
ZNF385A / Q96PM9
ZNF428 / Q96B54
ZNF431 / Q8TF32
ZNF451 / Q9Y4E5
ZNF483 / Q8TF39
ZNF512 / Q96ME7
ZNF512B / Q96KM6
ZNF592 / Q92610
ZNF598 / Q86UK7
ZNF622 / Q969S3
ZNF638 / Q14966
ZNF687 / Q8N1G0
ZNF706 / Q9Y5V0
ZNF787 / Q6DD87
ZNF800 / Q2TB10
ZNFX1 / Q9P2E3
ZNRD2 / O60232
ZNRF2 / Q8NHG8
ZPR1 / O75312
ZRANB2 / O95218
ZSCAN18 / Q8TBC5
ZSWIM8 / A7E2V4
ZW10 / O43264
ZYG11B / Q9C0D3
ZZEF1 / O43149
Gene Symbol / UniProt Accession
Brain
Colon
Duodenum
Esophagus
Heart
Liver
Lung
Pancreas
Thyroid
Organs
Type
DDA
DIA
Bins
1
2
3
4
5
